# Geology and discovery record of the Trinil *Pithecanthropus erectus* site, Java

**DOI:** 10.1101/2022.03.15.484451

**Authors:** O. Frank Huffman, Aart W.J. Berkhout, Paul C.H. Albers, John de Vos, Fachroel Aziz

## Abstract

The scientific utility of Eugène Dubois’ *Pithecanthropus erectus* (*P.e.*) Skullcap (Trinil 1), Femur I (Trinil 3) and associated paleontological specimens has been impaired for over a century by questions about their provenience. Firsthand accounts and contemporaneous field photographs, presented here, extensively document the site geology and discovery history.

The *P.e.* specimens and numerous-other fossils were unearthed in 1891-1893 from small excavations dug into a flat-lying bonebed exposed near the seasonal low-water level of the Solo River along its incised left embankment. Dubois’ on-site supervisors specified that the two *P.e.* fossils came from a ∼0.2-m-thick bonebed subunit traced at a single elevation for ∼12m from the 1891 Skullcap pit (∼30m^2^) to the 1892 Femur-discovery excavation and across an enlarged 1892-1893 trench (∼170m^2^). The depositional co-occurrence of the finds is supported by key documentation: the supervisors’ letters to Dubois about Femur I; his initial reporting to the Indies government; 1892-1893 accounts about expanding excavation of the Femur I stratum; Dubois’ 1891-1893 government submissions and 1894-1896 publications; confirmation by the Selenka Expedition in 1907-1908; Dubois’ annotations on unpublished site photographs; and a letter he wrote the year he died. Field studies in the 1930s to 1970s confirmed the essential aspects of the site geology.

The bonebed of 1891-1893 contained fossils referable to the extinct Trinil fauna species *Axis lydekkeri*, *Duboisia santeng* and *Stegodon trigonocephalus*. The Selenka Expedition excavations had a similar assemblage in the same stratigraphic position which they named the Hauptknochenschicht. The bonebed was thin bioclast-rich gravelly volcaniclastic sandstone with taphonomic and sedimentary features indicating an unusual origin. Bioclasts range from proboscidean craniums and logs to rat teeth, freshwater mollusc shells and leaves. The terrestrial-vertebrate skeletal elements are overwhelmingly disarticulated and frequently broken. Their surfaces are little-abraded by fluvial transport. The bone fossilization is quite uniform. More than one-hundred ungulate individuals perished. No evidence has been found of hominin- or terrestrial-carnivore involvement. The bioclasts varied in density from place-to-place and vertically, and were matrix supported in the bonebed. No substantial internal depositional hiatus was reported. In combination with Trinil’s paleogeographic context, these features implicate a catastrophic mortality of ungulates in a population aggregation along the floodplain of a perennial paleo-river, followed by lahar-flood transport and deposition of gravel-size lithic- and biotic-materials.

Trinil provides evidence favoring a broad archaic-hominin presence in southern Sundaland. The Trinil fauna is a lynch-pin in a long-lasting paleobiogeographic association between *H. erectus* and certain lineages of large bovids, cervids, proboscideans, rhinoceros, suids and tiger. The bonebed’s paleogeographic setting exemplifies the stratovolcanic drainages that *H. ere*ctus occupied for >0.8 million years in Java, including the watershed of a marine delta ∼150km east of Trinil, a volcanic island ∼100km north of Trinil, and areas to its west for 500km (where species associated with Trinil *H. erectus* occur). In the Java Sea (Sunda Shelf), seismic data image immense Pleistocene river- and coastal-terranes which archaic hominins and other large-mammals, like those at Trinil, might have inhabited.

## INTRODUCTION

September 23, 1892, was a seminal day in paleoanthropology. Eugène Dubois wrote to the Indies Government that he had unearthed a skullcap, femora and tooth at Trinil that *“brings humans in closer relation”* to *“the most advanced extant anthropoids.”* He hoped to *“get us started along the road to resolving the great mystery of human descent via paleontology,”* having concluded that *“the evolution of the femur … predated that of the skull”* (Dubois 1896b: 260, 270, translated). The Trinil Skullcap, known as *Pithecanthropus erectus* after 1894, became the first fossil to be widely accepted as representing humanities’ deep evolutionary past. But paleoanthropology continues to need reassurances about the provenience of Dubois’ finds.

The advances Dubois hope for have been realized. His search in Sumatra and Java, which was guided by geological mapping, spurred science towards the purposeful discovery and metric analysis of primate and hominin fossils (de Vos 2002, 2008, 2014, Henke 2007, Leakey and Slikkerveer 1993, Morwood et al. 2004, Shipman and Storm 2002, Theunissen 1985, 1989, Wood 2020). His discoveries started anthropology towards establishing low-profile craniums, erect-bipedal posture, limited-arboreal capabilities, modest-brain expansion, hands-freed-for-tool use and wide geographic dispersal as benchmarks in early human evolution. The limestone caves and volcaniclastic contexts that he explored are today the go-to targets for archaic-hominin discovery in Borneo, Flores, Sulawesi, Sumatra and the Philippines.

For decades, scientists accepted the Skullcap, Femur I and most other Trinil fossils as sourced from one stratum (e.g., Aziz et al. 1995, Bartstra 1982b, Bartstra et al. 1976, Boule 1923, De Terra 1943, de Vos 2008, Huffman et al. 2005, Matthew 1928, Osborn 1915, 1924, Reader 1981, Shipman 2001, Soeradi et al. 1985, Sollas 1908, van Es 1931). But the utility of the finds was impaired by uncertainty in provenience information Dubois provided and deficiencies in his underlying documentation, in part due to the submerged location of the discovery stratum in the Solo River, Java’s largest (Figures 1 and 2; Bartstra 1982, Brodrick 1948, 1964, Brongersma 1941 in de Vos 2014: 78, de Vos 1982, de Vos and Sondaar 1982, de Vos and Aziz 1989, Theunissen 1989; also, Alink et al. 2016). Here we use firsthand accounts and century-old photographs to document the geology and discovery history of *Pithecanthropus erectus* (*P.e.*).

**Figure 1.**
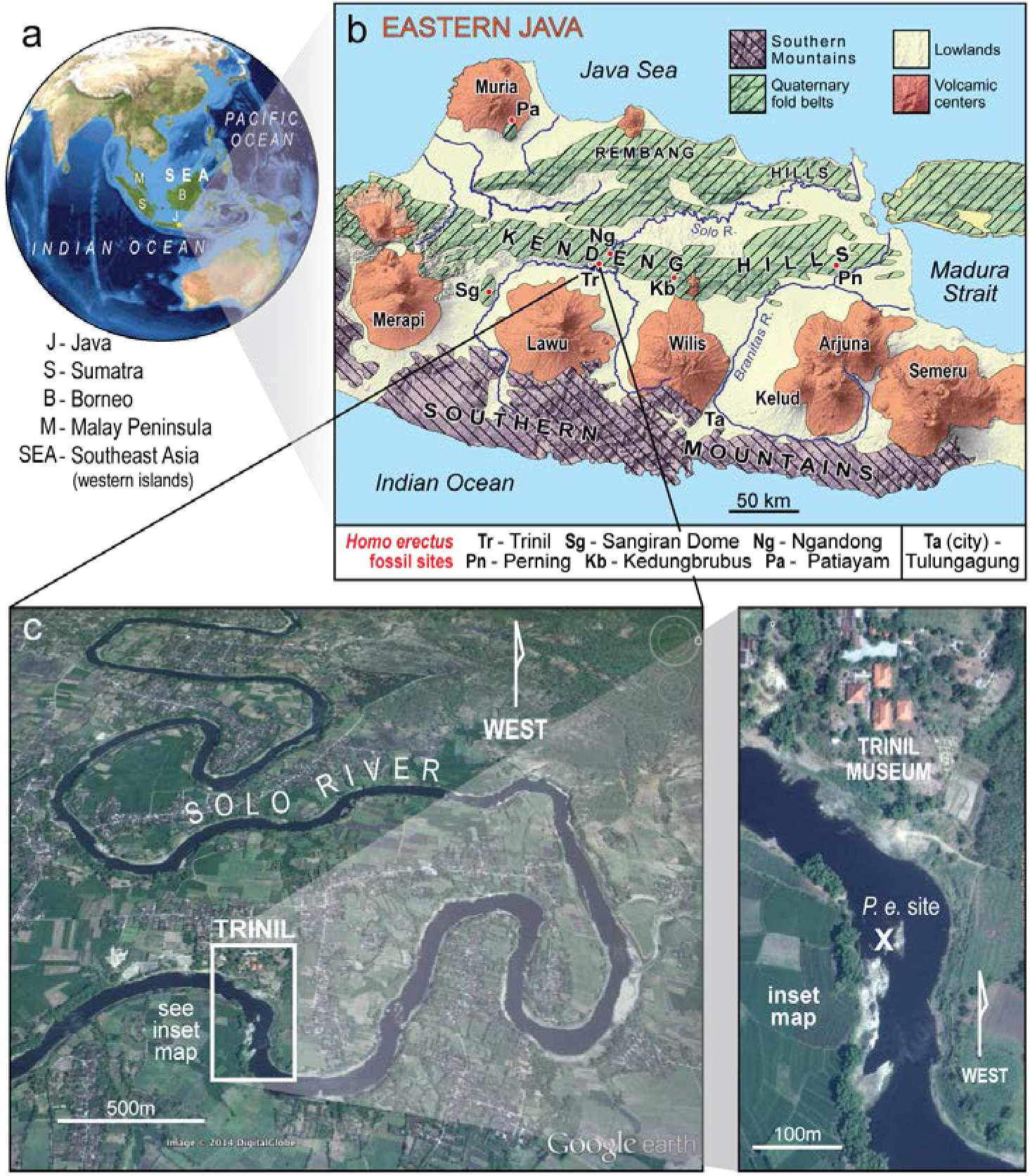
Trinil’s location. (**a**) In Southeast Asia. (**b**) In eastern Java with key *Homo erectus* fossil sites (Tables 5 and 6) on a generalized geological map (also, Huffman et al. 2010a: Figure 1). The hominin fossils from Trinil are referred to here as *Pithecanthropus erectus* (*P.e.*) to maintain an emphasis on their discovery (Figure 2; also, Supplement I Figure 1 and Supplement [Notes] II-F9; Supplement I citations have the form of S I Figure… and Supplement II-… citations are S II-… hereafter). (**c**) The spot (**X**) where the *P.e.* fossils were excavated during seasonal low-water level of the Solo River. The hominin fossils excavated amid other large vertebrate bioclasts in 1891-1892 pit and trenches (e.g., Tables 1 and 2, Figure 2a, Endnotes; de Vos and Sondaar 1982, Huffman et al. 2018). The river in the vicinity is largely bounded by incised banks (e.g., Figures 3c to 8). Since the 1920s, geologists have been able to see scars of the original excavations in the riverbed during times of particularly low water (Figure 9; S I Figures 3 and 7).

**Figure 2.**
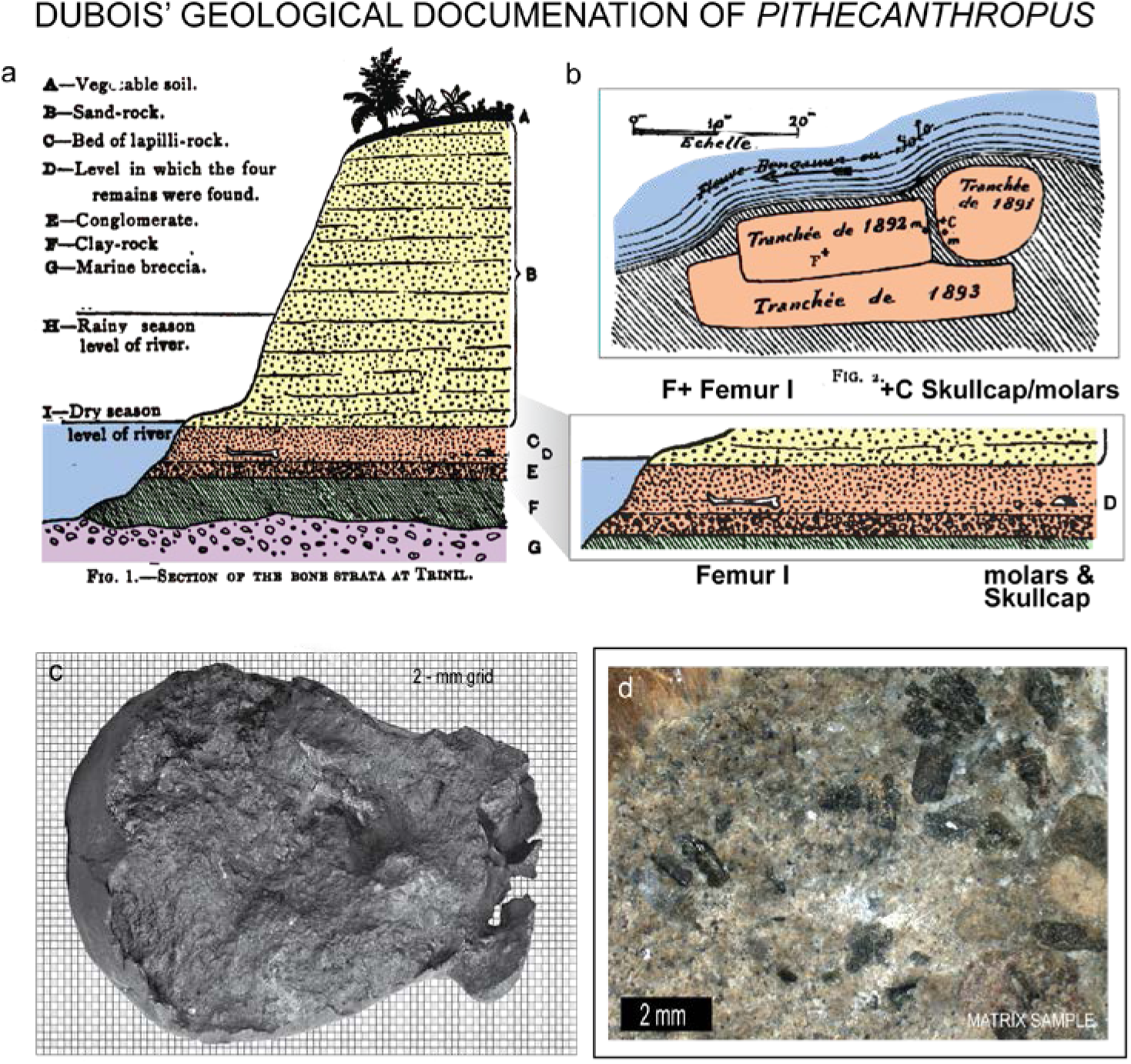
Eugène Dubois published little information on the provenience of *Pithecanthropus erectus*, so that a precise re-description of stratigraphic context and discovery history using unpublished sources is necessary. (**a**) Dubois’ principal geological display, which is a partially schematic cross section of the left bank (Figure 1c), attributed the hominin and other fossils to a *“Bed of lapilli-rock”* (the cross section was published in multiple versions; this one is from Dubois 1896c, d and is rearranged and colorized after Huffman et al., 2018; also, Dubois 1895a, b, c, 1896a, b, e in S II-F). The discovery stratum, our Lapilli Bed (**LB**), included Dubois’ *“Level in which the four* [hominin] *remains were found,”* a subunit that we term the Principal Fossil Zone (**PFZ**). He (1896b: 251; S II-F3) characterized the *“Sand-rock”* above the **LB** as *“hardened volcanic tuffs consisting of clay, sand and lapilli.”* (**b**) Dubois (1895a: 158; S II-F2) published only one map of his excavations, an 1895 sketch of the 1891-1893 pit and trenches on the left bank. We rename the excavations to conform to firsthand accounts. His *“Tranchée de 1891”* is our Skullcap Pit; it is where the 1891 Molar (‘*m*’ in the map) and 1891 *Pithecanthropus erectus* Skullcap (‘*c*’) were discovered (specimens sometimes referred to as Trinil 1 and 2, respectively). His *“Tranchée de 1892”* is the 25-m Trench, where Femur I (‘*F*’) and the 1892 Molar (“*m*”) were found (the specimens are sometimes referred to as Trinil 3 and 4, respectively). His *“Tranche de 1893”* is the 40-m Trench (Figure 3a). (**c**, **d**) Dubois had voluminous unpublished materials on Trinil, such as photographs of the Skullcap filled with coarse-grained sandstone, and sandstone specimens stored with the Skullcap (photograph of rock courtesy of F.P. Wesselingh). See also S I Figure 1. The identity of scans of photographs done at Naturalis Biodiversity Center is generally given the form DUBO#### (Albers and de Vos 2010). The ‘c’ is image DUBO1303 with a grid added to highlight granule-sized gravel clasts (2-4 mm).

**Table 1.** Prominent taxa excavated at Trinil.**^1^** (**a**) From Eugène Dubois’ 1891-1893 “Bed of Lapilli” (**LB**), Selenka Expedition’s “Hauptknochenschicht” (**HK**), and a bed near Solo River low-water levels during 1895-1900 (**LB**-**HK**). (**b**) In the Dubois Collection, Naturalis.

|  | a. Noted in Dubois field supervisors' letters and Dubois' and Selenka's records and publications <sup>1</sup> |  |  |  |  |  |  | b. <sup>2</sup> |  |
| --- | --- | --- | --- | --- | --- | --- | --- | --- | --- |
| Location of the excavation > | Left bank (south side) of the Solo River (Figure 1c) |  |  |  | Right bank of the river |  | Left or right | Percentages of Trinil specimens in the Dubois Collection |  |
| Years of excavation <sup>3</sup> > | '91 '92 '93 | '95 '96 '97 | '99 '00 | '07 | '91, '92, '95 '96 | '07 | '91 '92 '95 |  |  |
| Source bed > | LB | LB-HK | LB-HK | HK | LB-HK | HK | LB, LB-HK |  |  |
| TAXA vvvvvvvv |  |  |  |  |  |  |  |  |  |
| CERVIDAE (Cervini)<br>*Axis lydekkeri* <sup>2</sup> | '91, '92, '93 | '95, '96, '97 | '99, '00 | '07 | '91, '92, '95, '96 | '07 | '91, '92 | 28% | 68% |
| BOVIDAE (Bovini) <sup>2</sup><br>*Bibos palaesondaicus*<br>*Bubalus palaeokerabau* | '91, '92, ---<br>'91, '92, '93 | ---, --- '97<br>---, ---, '97 | '99, '00<br>'99, '00 | '07<br>'07 | '91, '95<br>'91, '92, '95 | ---<br>'07 | '91<br>'91, '92 | 40% |  |
| PROBOSCIDEA<br>*Stegodon trigonocephalus* | '91, '92, '93 | '95, ---, '97 | '99, '00 | '07 | '91, '95 | '07 | '91 | 13% | 19% |
| BOVIDAE (small)<br>*Duboisia santeng* | '91, '92, '93 | '95, ---, --- | '99, '00 | '07 | '95, '96 | '07 | '91 | 6% |  |
| CROCODYLIA<br>*Crocodylus siamensis*<br>*Gavialis bengawanicus* | ---, '92, '93<br>---, ---, --- | ---, '96, ---<br>'95, '96, --- | ---, '00<br>---, --- | '07<br>--- | '95, '96<br>---, --- | '07<br>--- | '91, '95<br>'91, --- | 2% | 7% |
| TESTUDINES<br>(Geoemydidae/Geoemydina;<br>Trionychidae) | '91, '93 | ---, '96, --- | ---, '00 | '07 | ---, '96 | '07 | '91 | 5% |  |
| HOMINID<br>*Pithecanthropus erectus* <sup>4</sup> | '91, '92 | ---, ---, --- | '00 | --- | ---, ---, --- | --- | ---, ---, --- |  |  |

<sup>1</sup> Taxonomic identifications, as originally reported, have been converted to current taxonomic nomenclature on the basis of the Trinil species that are currently recognized in the Dubois Collection (Naturalis Biodiversity Center, Leiden; Table 3). The entries for the Selenka Expedition excavations are from Selenka and Blanckenhorn (1911) and an unpublished listing of finds, the 1907 Listing ("PM\_S\_II\_Selenka\_FB\_1-78.pdf" at the Museum für Naturkunde Berlin; MNB), in part based on input from examination of the fossils at the MNB by Lawrence Todd (pers. comm., 2015).
<sup>2</sup> Based on Storm's (2012) tabulations of 3857 identified non-hominin specimens from Trinil, as recorded in the 2002 electronic catalogue of the Dubois Collection.
<sup>3</sup> Dubois' left-bank excavations are represented following way: 1891 Skullcap Pit by '91. 1892 25-m Trench by '92. 1893 40-m Trench by '93. 1895 Ledge by '95. 1896 Left-bank Pit by '96. 1897 Downstream and Upstream Pits by '97. 1899 Trench by '99. 1900 Trench by '00. There were no Trinil excavations in 1898. Selenka Expedition right-bank and left-bank Pits I and II, respectively, are indicated by '07. The ---s mean that there are no finds referable to the indicated species in that year and excavation.
<sup>4</sup> *P. erectus* is now *Homo erectus*. Dubois (1894a) attributed three fossils from Trinil to the new hominin species *Pithecanthropus erectus* without designating a single holotype specimen, so that three equally ranking syntypes form the holotype. No lectotype that conforms the standards of the International Code of Zoological Nomenclature has been assigned. The Dubois Collection at Naturalis retains the Skullcap but it does not have an accession number. Jacob (1975a) terms the 1891 Molar (Figure 2) Trinil 1, the Skullcap Trinil 2 and Femur I Trinil 3.

**Table 2.** (**a**) Vertebrate remains in the Hauptknochenschicht, **HK**, compared to superjacent stratigraphic units (‘layers’) in Selenka Expedition of Pits I and II in 1907 (preliminary evaluation). (**b**) Three depositional subdivisions were recognized in the **HK** of 1907 Pit I (right bank).

| a Number of finds listed by excavation area and layer (field designations), showing the fossil concentration in the HK <sup>1</sup> |  |  |  |  |  |  |  |  |  |  |
| --- | --- | --- | --- | --- | --- | --- | --- | --- | --- | --- |
| Excavation (location) | PIT I (right bank of the Solo River) |  |  |  |  | PIT II (left bank of the Solo River) |  |  |  |  |
| Stratigraphic position | above HK | Hauptknochenschicht (HK) |  |  | below HK | Units 2-4, Figure 4a |  | Hauptknochenschicht (HK) |  | below HK (?) |
| Layer #s in 1907 Listing | 10-14 | 15 | 16 | 17 | 18-20 | 1 <sup>2</sup> | 2 | 3 <sup>2</sup> | 4 | 5-9 |
| # of finds in the layer(s) | 50 | 545 | 86 | 429 | 26 | 17 | 67 | 299 | 106 | 17 |
| # of identifiable finds <sup>3</sup> | 42 | 449 | 68 | 305 | 17 | 12 | 38 | 191 | 52 | 11 |
| % of identified finds | 84% | 82% | 79% | 71% | 65% | 71% | 57% | 64% | 49% | 65% |
| CERVIDAE (Cervini) | 16 | 285 | 50 | 190 | 7 | 0 | 19 | 116 | 21 | 5 |
| BOVIDAE (Bovini) | 11 | 73 | 10 | 79 | 7 | 0 | 1 | 35 | 22 | 4 |
| *Stegodon trigonocephalus* | 10 | 47 | 3 | 13 | 0 | 9 | 0 | 9 | 2 | 0 |
| *Duboisia santeng* | 1 | 4 | 1 | 7 | 1 | 0 | 3 | 6 | 2 | 0 |
| CROCODYLIA | 1 | 6 | 2 | 6 | 0 | 2 | 2 | 3 | 0 | 1 |
| TESTUDINOIDEA | 3 | 18 | 0 | 4 | 1 | 0 | 0 | 0 | 1 | 1 |
| *Sus brachygnathus* | 0 | 1 | 0 | 2 | 0 | 0 | 4 | 8 | 2 | 0 |
| *Rhinoceros sondaicus* | 0 | 8 | 1 | 0 | 1 | 0 | 0 | 1 | 1 | 0 |
| *Hexaprotodon* | 0 | 0 | 0 | 0 | 0 | # | 0 | 0 | 0 | 0 |
| Fishes | 0 | 0 | 0 | 1 | 0 | 1 | 5 | 9 | 1 | 0 |
| [Antlers <sup>3</sup>] | [9] | [243] | [41] | [140] | [3] | [0] | [2] | [11] | [3] | [1] |

| b Details on Hauptknochenschicht (HK) finds, highlighting Pits I and II paleontological similarities and differences |  |  |  |  |
| --- | --- | --- | --- | --- |
| Name (location) of the excavation | PIT I (right bank) | PIT II (left bank) | PITS I AND PIT II |  |
| Stratigraphic terminology | Hauptknochenschicht (HK) |  |  |  |
| Layer #s in the 1907 Listing | 15-17 | 3-4 <sup>4</sup> | 3-4 and 15-17 |  |
| Percent MB finds that are identified | 78% | 59% | 72% |  |
| Number of finds (and % of total finds) by taxon |  |  |  |  |
| CERVIDAE (Cervini) <sup>3</sup> | 525 (64%) | 133 (56%) | 658 (62%) | (83%) |
| BOVIDAE (Bovini) | 162 (20%) | 57 (23%) | 219 (21%) |  |
| *Stegodon trigonocephalus* | 63 (8%) | 11(4%) | 74 (7%) | (9%) |
| *Duboisia santeng* | 12 (1%) | 8 (3%) | 20 (2%) |  |
| CROCODYLIA | 14 (1%) | 3 (2%) | 17 (2%) | (4%) |
| TESTUDINOIDEA | 22 (3%) | 1(-%) | 23 (2%) |  |
| *Sus brachygnathus* | 3 (-%) | 10 (4%) | 13 (1%) | (2%) |
| *Rhinoceros sondaicus* | 9 (1%) | 2 (1%) | 11 (1%) |  |
| Fishes | 1 (-%) | 10 (4%) | 11 (1%) | (2%) |
| Other | 11 (1%) | 4 (2%) | 15 (1%) |  |
| Total identified entries | 822 (100%) | 239 (100%) | 1061 (100%) | (100%) |

<sup>1</sup> From the 1907 Listing; the Museum für Naturkunde Berlin, MNB, document ‘PM\_S\_II\_Selenka\_FB\_1-78.pdf’ and in part L. Todd (pers. comm., 2015) and Selenka and Blanckenhorn (1911, translated in Berkhout and Huffman 2021). The number of finds with layer attributions is 96% for Pit I and 75% for Pit II. Oppenoorth (1908a,b, 1911) reported 2000 finds in 1908 but gave no layer details.
<sup>2</sup> Layer 1 is the “*Stegodon* bonebed” (SB; Figure 8, S I Figure 10).
<sup>3</sup> ‘Layer 3’ includes 7 “Pithe[can]thropus” bed entries; excluded here are eighteen June 5-6 entries that are ambiguous as to source Pit and bed. Endnote F(viii). Taxonomic identification based on various sources. Stremme (1911) examined 527 *Axis* antler beams (with 230 shed) and ~375 bony cervid fossils.
<sup>1</sup> From the 1907 Listing; the Museum für Naturkunde Berlin, MNB, document 'PM\_S\_II\_Selenka\_FB\_1-78.pdf' and in part L. Todd (pers. comm., 2015) and Selenka and Blanckenhorn (1911, translated in Berkhout and Huffman 2021). The number of finds with layer attributions is 96% for Pit I and 75% for Pit II. Oppenoorth (1908a,b, 1911) reported 2000 finds in 1908 but gave no layer details.
<sup>2</sup> Layer 1 is the "Stegodon bonebed" (SB; Figure 8, S I Figure 10).
<sup>3</sup> 'Layer 3' includes 7 "Pithe[can]thropus" bed entries; excluded here are eighteen June 5-6 entries that are ambiguous as to source Pit and bed. Endnote F(viii). Taxonomic identification based on various sources. Stremme (1911) examined 527 *Axis* antler beams (with 230 shed) and ~375 bony cervid fossils.
<sup>4</sup> The Selenka Expedition unearthed ~2,200m<sup>3</sup> of rock in 1907, having exposed ~260 m<sup>2</sup> of **HK** and having collected ~700 fossils. In 1908, the Expedition removed ~950 m<sup>3</sup> of material, dug ~110 m<sup>2</sup> of **HK**, and thus should have made ~300 finds (Dozy 1911a: xlii, Oppenoorth 1908b: 145, 1911: xxxviii, Berkhout and Huffman 2021: 53). The average vertebrate-fossil density in the **HK** was ~2.7m<sup>-2</sup> based on ~1000 specimens from ~370 m<sup>2</sup>. Regarding the Testudinoidea and *Stegodon*, see Janensch (1911a,b).

Among the uncertainties has been the *“contemporaneity”* of the *P.e.* Skullcap and Femur I (Rightmire 1990: 16), a controversy stemming from the *Homo sapiens*-like anatomy of the long bone. More than a hundred fossils attributed to *H. erectus* have now come from Java, seeming to make the *P.e.* finds less relevant, even though the Java assemblage might well represent several hominin species (Antón 2013, Antón et al. 2007, 2014, Baab and Zaim 2017, Indriati 2004, Kaifu 2017, Kaifu et al. 2008, 2015, Mayr 1950, Noerwidi et al. 2016, Santa Luca 1980, Schwartz 2016, Tyler 2003, Washburn 1951, Weidenreich 1946, Zanolli et al. 2019). For some, Trinil seems to be a tangential component of the hominin record in Java (e.g., Sémah et al. 2016).

In the following pages, we repair former shortcomings in the geological and paleontological record at Trinil. We define the discovery bed more precisely, beginning with Dubois’ only geological display of the *Pithecanthropus erectus* site, a partially schematic geological cross section (Figure 2a). In it, he was explicit about the provenience of the *P.e.* materials and associated fossils. They were concentrated in a thin *“Bed of lapilli-rock,”* a stratum that we term the Lapilli Bed (**LB**). It had been exposed on the left (south) bank of the Solo River by deep incision of the valley (Figure 2a). The **LB** was *“about 1 meter thick”* (Dubois 1896c: 3 in Supplementary Materials II-F3i). Both the Skullcap and the *“left femur* [originated from] *… the same level”* within the **LB** (Figure 2a; Dubois 1894a: 1, in Supplementary Materials II-B6j).

The Selenka Trinil Expedition re-excavated at Trinil in 1907-1908. One of their excavations (Pit II) was adjacent to Dubois’ 1900 Trench (Selenka and Blanckenhorn 1911). For decades thereafter, the Selenka publications e stood as confirmation of Dubois’ stratigraphic portrayal of the *P.e.* discovery context (e.g., Boule 1923, Osborn 1915, de Terra 1943). One Expedition member had coined the term Hauptknochenschicht (**HK**), the main bone-bearing layer (Branca 1908, Carthaus 1911b: 14, Dubois 1908: 1242). This designation spotlighted the high vertebrate fossil content of the bonebed relative to other levels dug in the 1907-1908.

The Selenka geologists were confident that their **HK** was the same sedimentary deposit as Dubois’ **LB** (Oppenoorth 1907, Selenka and Blanckenhorn 1911). While we ultimately agree, the term Hauptknochenschicht is not a satisfactory stratigraphic substitute for the **LB** as Dubois defined it because *P.e.* fossils came from the **LB**, not Selenka’s **HK** (Figure 2a).

Moreover, between 1895-1900, Dubois’ crews excavated the largest area ever at Trinil (Figure 3a), and discovered the great majority of the fossils from the site (Tables 1, Endnotes A through E). Dubois was in the Netherlands then, and did not later publish on the geology of the 1895-1900 excavations. The 1891-1893 discovery pit and trenches he had seen personally were buried by spoils and not visible to the Selenka geologists (Figures 6b and 7). Thus, to maintain precision in reporting on the discovery record, we refer to the fossil-rich stratum encountered in 1895-1900 as the **LB**-**HK** (Table 1). The term main bonebed is used here as a general referent.

**Figure 3.**
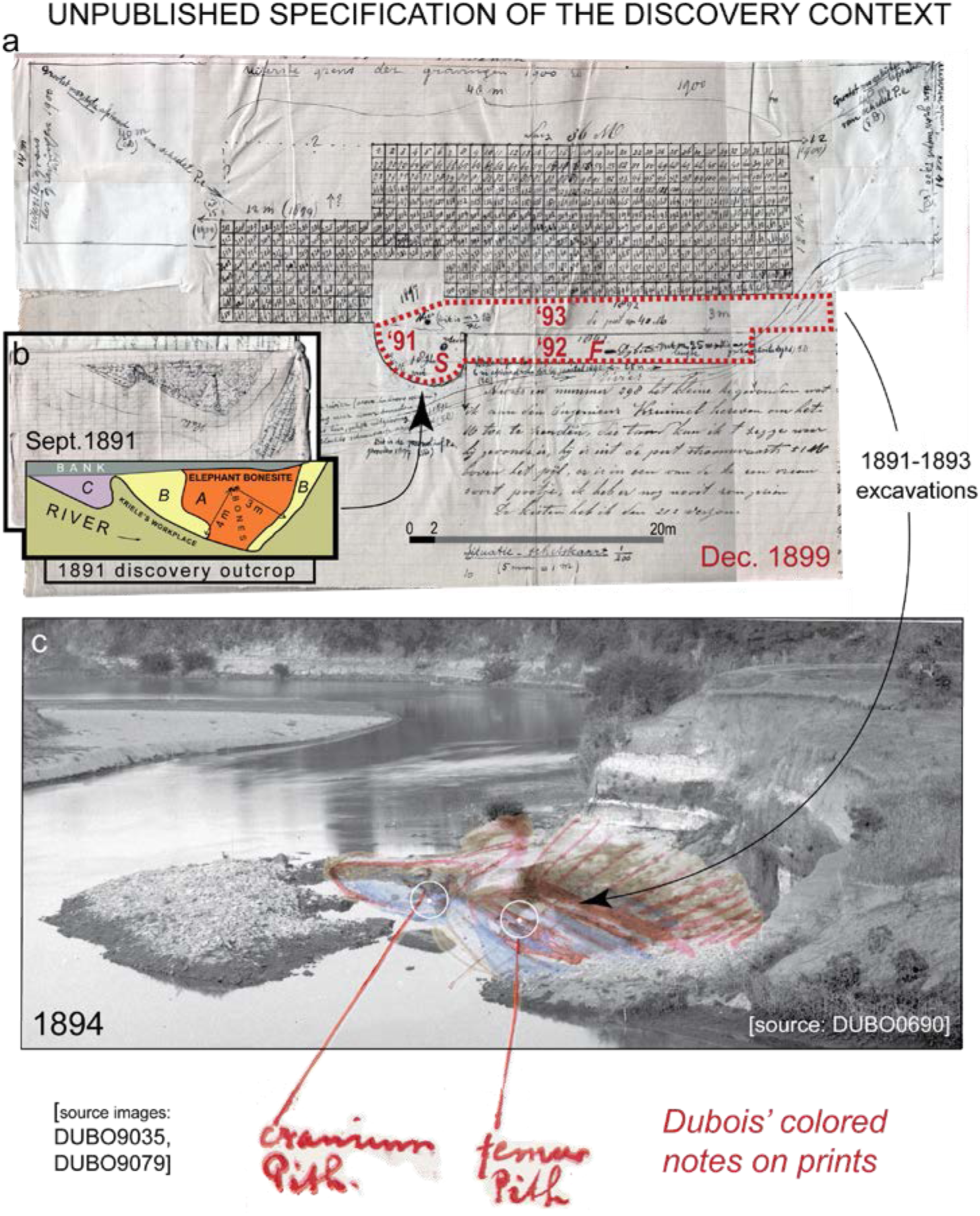
Among the most important unpublished Dubois records about the left bank are two maps and an 1894 site photograph (de Vos and Aziz 1989). (**a**) The most detailed-representation of the excavations was drawn by field supervisor G. Kriele in 1899 (south is up) and annotated by Dubois in apparent approval (S II-A4m), although the layout of excavations is partially inconsistent with other records (e.g., S II-A4r). Shown with evident accuracy are the discovery points for the Skullcap (**S**), Femur I (**F**) and 1891 Molar (*“this is m3/P.e*.” located south of the Skullcap; the 1892 Molar is not shown). Note the small size of the 1891-1893 pit and trenches (∼200m^2^) relative to the 1895-1900 excavations of ∼1650m^2^ (including the 1899 Trench, which is gridded, and the outline of the proposed 1900 Trench; S II-A4r). (**b**) This September 18, 1891, map of the site (south is up) includes an inset redisplay of the Skullcap site on the left bank (original penciled patterns are colorized). A *Stegodon* tusk and mandible had been exposed in a ledge of **LB** sandstone (S II-A1b). (**c**) This 1894 photograph was taken on September 5 during low water on the Solo River, when the 1891-1893 pit and trenches was all that had been dug. The eight-to-nine-meter high embankment to the south (right) was removed in 1896-1900 to mine the **LB** near the low-water level. Dubois annotated prints of the image specifying the hominin-discovery spots relative to the nearby geology (Huffman et al. 2015, 2018; S I Figures 3 and 4). The photographic image is from high-resolution scanning of a negative.

**Figure 4.**
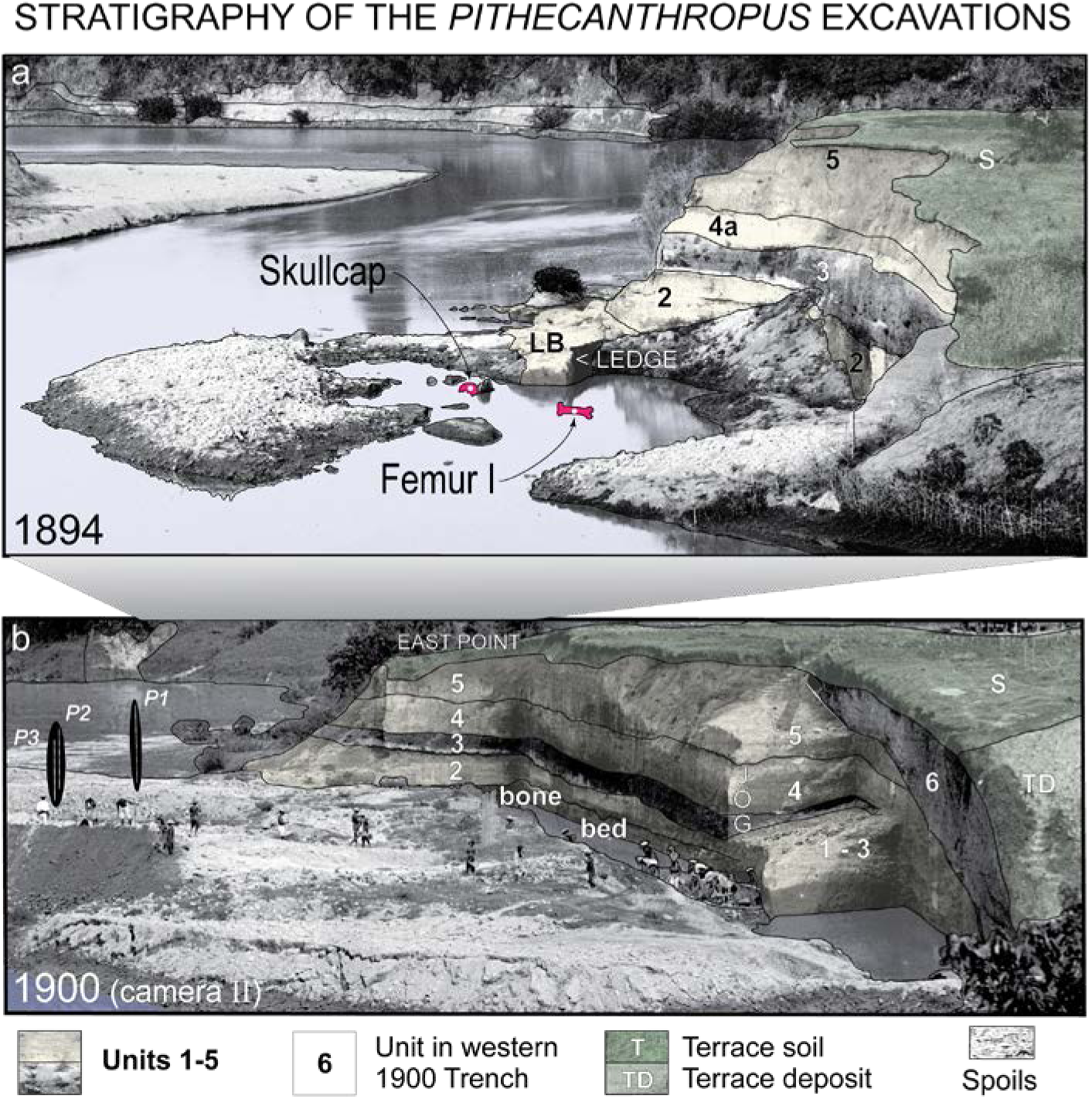
Site photographs from 1894, 1900 and 1907 permit geological analysis of the rock removed by excavation, as illustrated in Figures 4 to 9 (Huffman et al. 2015, 2018). Both views which are presented here look generally eastward at the left-bank excavations. (**a**) The hominin discovery stratum **LB** is seen in the 1894 image to have underlain the rocky stratigraphic remnants of 1893 excavation back wall, where we divide the sequence photographically into stratal units **2**-**5** (also, S I Figure 3). Only soil mantled the terrace surface (**S**), which capped the sequence. The geological circumstances match Dubois’ 1895-1896 site cross sections (Figure 2a). (**b**) The same units are recognized tens of meters away at about the same elevations in the high-standing back walls of Dubois’ 1900 Trench (this photograph was taken from camera station II in Figure 6a; also, Figure 5 and S I Figures 4 to 7) and Selenka Expedition’s 1907 Pit II (Figures 7 and 8). Unit **3** is clearest in revealing the site-wide stratigraphic continuity and horizontal structural attitude of the bedding (especially notable around East Point and the JOG in the 1900 back wall; also, S I Figures 6). In the western parts of the 1900 Trench (and also in Selenka Pit II), a superjacent stratigraphic unit with foreset bedforms (**6**) overlay an erosional surface cut into **2**-**5** (also, Figures 5, 7 and 9). Formal geographic surveying of the Trinil area was done in November 1900 and the results made into the 1900 Site Map (Figures 6a; S I Figure 7a). The terrace upland surface above the upper lip of the 1900 Trench rose in elevation southeastward to >12.5m above the river at low level and reached ∼20m farther from the 1900 Trench (e.g., higher terrace in S I Figure 3c). A bar-form feature, apparently a terrace deposit (**TD**), is visible above **6** in the 1900 image. The 1894 and 1900 photographs, as well as the 1900 Site Map, have been published widely without geological analysis (e.g., Albers and de Vos 2010, Alink et al. 2016, de Vos and Sondaar 1982, de Vos and Aziz 1989, Leakey and Slikkerveer 1993, Reader 1981, Shipman 2001, Theunissen 1989).

**Figure 5.**
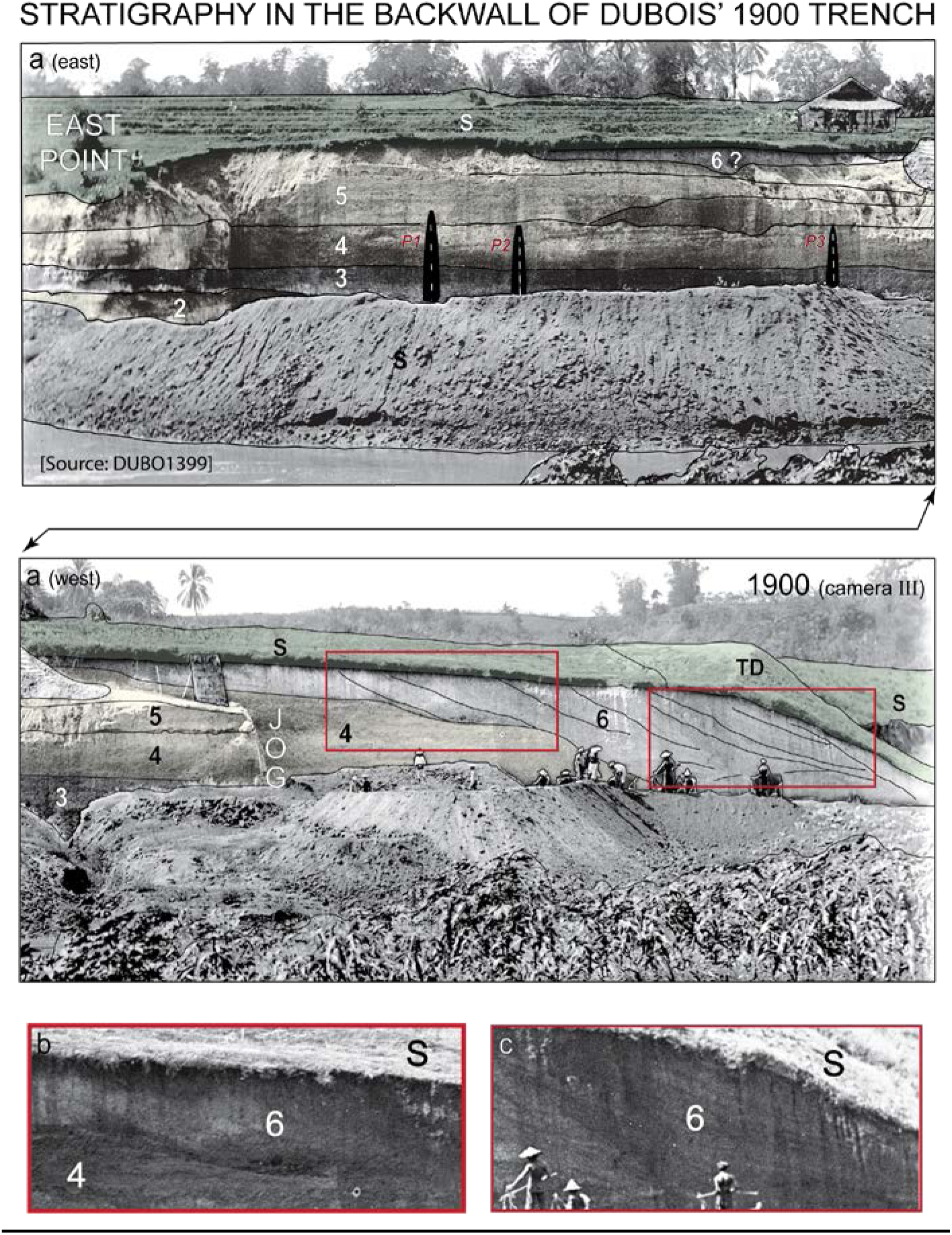
(**a**) A second November 1900 image of Dubois’ left-bank 1900 Trench confirms the lateral continuity and approximate flat-lying structural attitude of units **2**-**5**, as well as their stratal relationship to the terrace soil **S** and unit **6** (**b** and **c**). The photograph looks South and was taken along a sight line perpendicular to the ∼100m-long Trench (from camera station III in Figure 6a and S I Figure 7; S I Figure 6 has a third 1900 photograph). Unit **6** has large-scale inclined depositional layering that in part seems to truncate at terrace surface (**S**). Units **1**-**5** and **6** had been well-enough lithified to stand in near-vertical walls (Figure 7). Duyfjes (1936) included all of the beds seen here, **6** included, in his bedrock Kabuh Formation (also, Figure 6d). Excavation spoils deeply covered the 1891-1899 excavations baulks and trenches. Three scaled poles (P1-P3) were staked out in the spoil pile, presumably by Dubois’ on-site supervisor G. Kriele (also, Figure 4). They apparently were intended them to mark where the *Pithecanthropus erectus* find spots were under the spoil pile. The poles were one of several errant attempts made to relocate the find spots (S I Figure 7b). Albers and de Vos (2010: 75) has the whole unmarked image.

**Figure 6.**
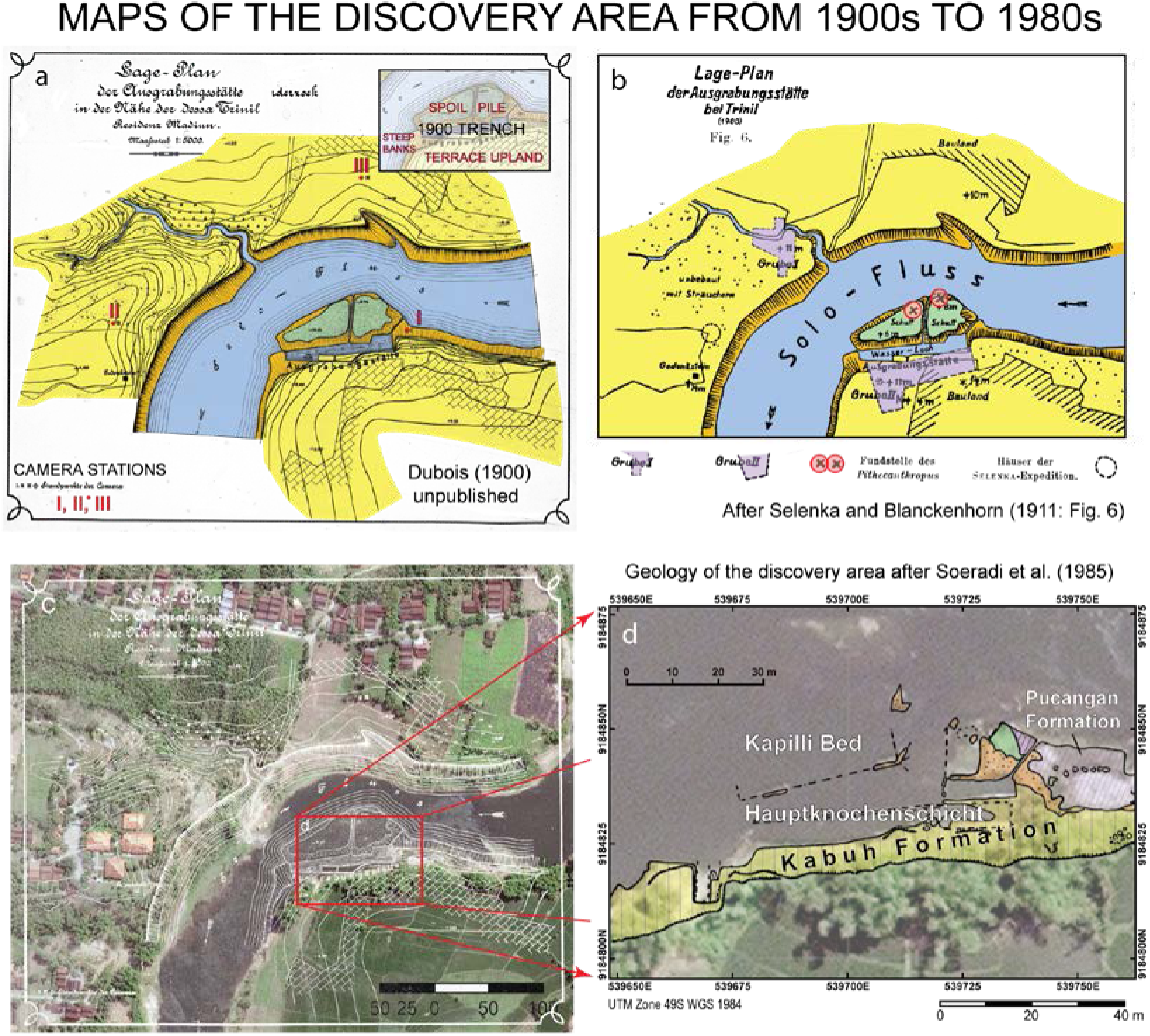
Maps and a satellite imagery help relate the *Pithecanthropus erectus* site to the modern landscape (Huffman et al. 2018). North is up. (**a**) Dubois’ 1900 Site Map (north is up) shows a partially inundated 1900 Trench lying between an extensive spoil pile and terrace upland (highlighted in inset) where the topography is defined by 1m contours. This version of the Map has been colorized and annotated (also, S I Figure 7a). Unannotated versions are in de Vos and Sondaar (1982), Leakey and Slikkerveer (1993), Shipman (2001) and others. The scales shown on the archival copies of the Map reflect the size at which they were intended to be published. (**b**) The Selenka Trinil Expedition re-use of Dubois’ map includes their attempts to relocate the Skullcap and Femur I findspots (Selenka and Blanckenhorn 1911: xiii, Plate I, Fig. 1 and 2, Berkhout and Huffman 2021). However, the spots are separated by a distance that is greater than Dubois reported (S I Figure 7b). The Selenka map greatly exaggerates the size of their left-bank excavations (Pit II) at the main bonebed, **HK**, level (Figure 7b). (**c**) Dubois’ 1900 Site Map (white lines) is here superimposed approximately on an ortho-rectified satellite image (portion of WGS_1984_UTM_zone_49S purchased in 2013). Dubois Collection prints of the Map have scales reflecting their intended sizes in reproduction, not the scale of the original map. (**d**) The 1:250 geological mapping of the left bank by Soeradi et al. (1985) has been adjusted here to fit onto the satellite image (S I Figure 18) and annotated. The Soeradi team identified outcrops of the main bonebed in the areas of Dubois’ and Selenka’s excavations (Lapilli bed and Hauptknochenschicht, respectively), and recognized the exposed strata thereabouts as being the Kabuh and Pucangan Formations (Duyfjes 1936; also, Datun et al. 1992, I.J.J.S.T. 1992, van Es 1927, 1929, 1931).

**Figure 7.**
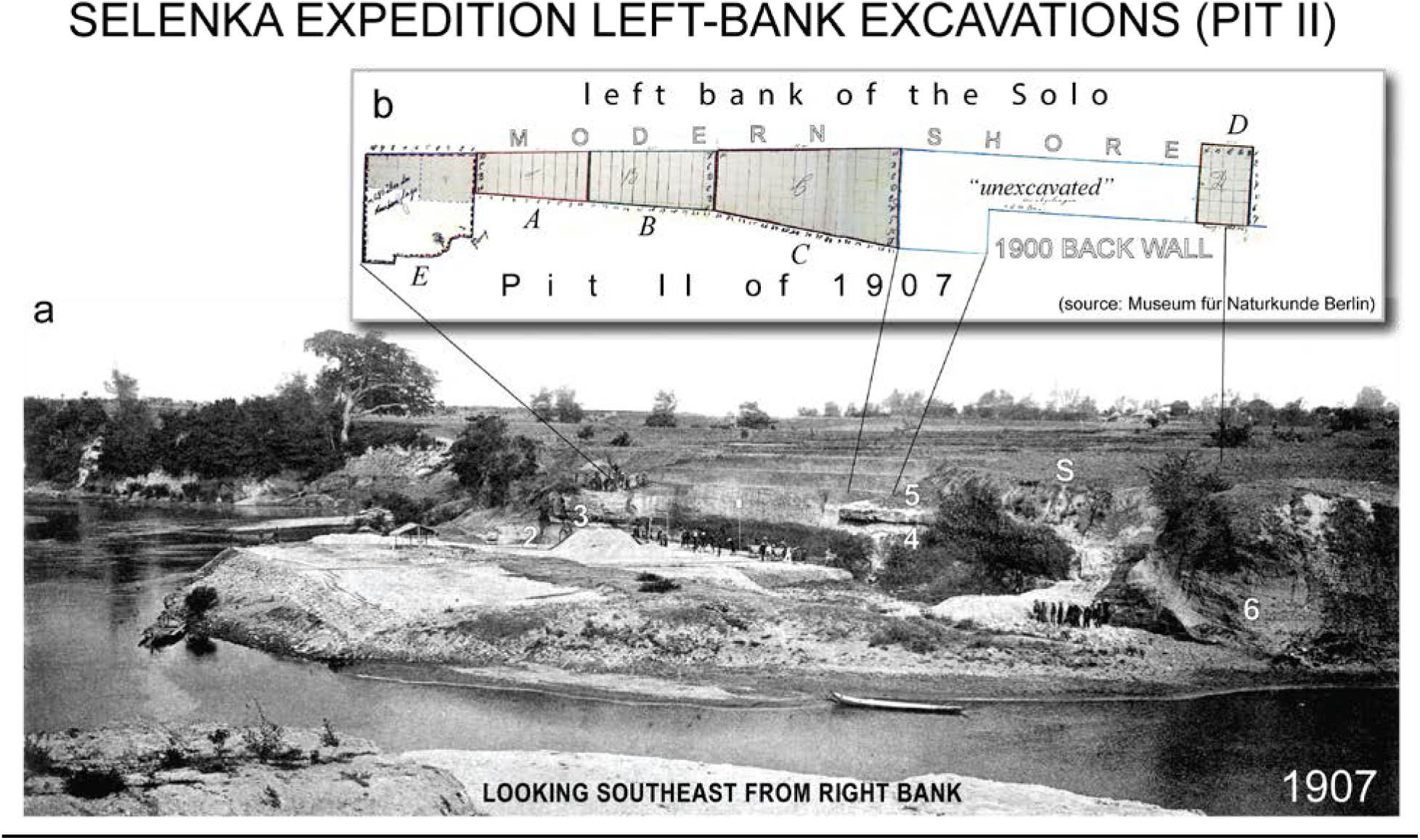
Later geological work in the left-bank area confirmed the discovery stratigraphy of Dubois. An August 1907 photograph (**a**) and excavation plat (**b**) shows the Selenka Expedition 1907 Pit II. It had been dug into the back walls of Dubois’ 1900 Trench, and was expanded in 1908. Today, Pit II and Dubois’ 1900 Trench lie adjacent to the bank of the Solo River (Figure 9b; S I Figures 2 and 7). Pit II encountered units **2**-**5** above their Hauptknochenschicht (**HK**; not noted on the photograph). Much of **6** was removed at the west end of the Pit, where the erosional base of **6** reached river level (also, S I Figure 11). The plat shows the 1-m excavation grid in use (Oppenoorth 1908). Spoils from Dubois’ 1900 trenching deeply covered his 1891-1899 excavations, including *Pithecanthropus erectus* discovery site, so that the Expedition team never saw the scars and baulks of **LB** around that location. The ‘a’ image is the majority of a figure published by Selenka and Blanckenhorn (1911: Fig. 2; also, Selenka and Blanckenhorn 1911: Fig. 1, 8, and Oppenoorth 1911: Fig. 17, 18 and 19, Berkhout and Huffman 2021). The ‘b’ map is from a photograph (PM_B_IX_148.tif) of the original drawing, provided courtesy of the Museum für Naturkunde, Berlin (MNB).

Today, the 1891-1908 excavation are delineated by low-lying baulks, but they are only small remnants of the great volume of strata removed, and vastly more fossils from the **LB**, **LB**-**HK** and **HK** are stored in European museums than will be unearthed at the site in the future. Fortunately, old photographs provide a reliable means of visualizing the strata dug away (Huffman et al. 2015). Only one conformable horizontal indurated stratal series is evident in the images (Figures 4, 5, 7 and 8). The photographic circumstances coincide in fundamental ways to Dubois’ site cross section (Figures 2a; S II-F, Foreword). This encourages confidence in re-reading original reporting on the operations and fossil discoveries; these records in turn improve geological characterization of the site and its paleontology (e.g., Figures 4, 9 and 10; Huffman et al. 2018).

**Figure 8.**
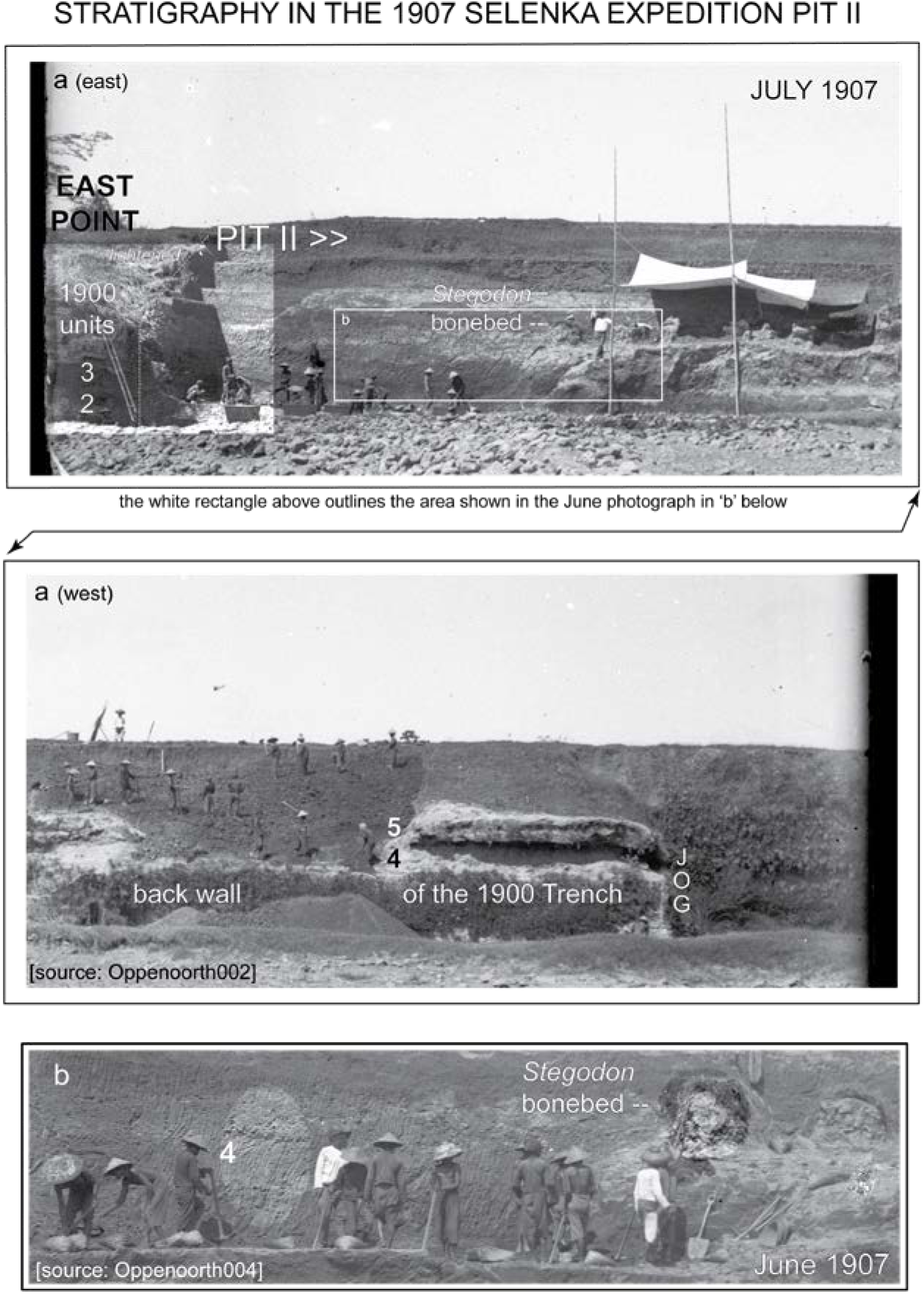
Selenka Expedition geologist W.F.F. Oppenoorth photographed the left bank multiple times in 1907 (this view is looking south is similar to the view in Figure 5). He reported that *“the bonebed … washed free”* at the end of April; the top of the bonebed was visible above low-river level in late May near East Point, where Dubois’ crew had exploited the fossil concentration in 1900 (Oppenoorth 1911: xxxi, xxxiii and Figure 20, xxxiv, Berkhout and Huffman 2021; S I Figure 8). By July 1907, Selenka’s men were digging into the Dubois 1900 back wall, which stood as much as eight-to-nine meters high (Oppenoorth 1911: xxxiii; S I Figure 9, Berkhout and Huffman 2021). The fossils dug from Pit II originated from points just a few meters away from the modern south shore of the river. (**a**, **east**) Later in July, the Hauptknochenschicht (**HK**) was reached in the easternmost Pit II (left in the image). To the west, the dig was at a stratigraphic level ∼5m higher (our basal unit **5**), where the associated elements of a *Stegodon trigonocephalus* skeleton lay covered in straw on pedestals (*Stegodon* bonebed, **SB**; S I Figures 9 and 10). (**a**, **west**) West of the **SB** exposure, units **4** and basal **5** were still visible in the 1900 backwall. (**b**) An enlarged portion of a June photograph of eastern Pit II (bottom) shows the excavators using pickaxes to penetrate units **4** and **5**, where excavation faces also show pick scars (as highlighted in the oval; also, S I Figure 9).

**Figure 9.**
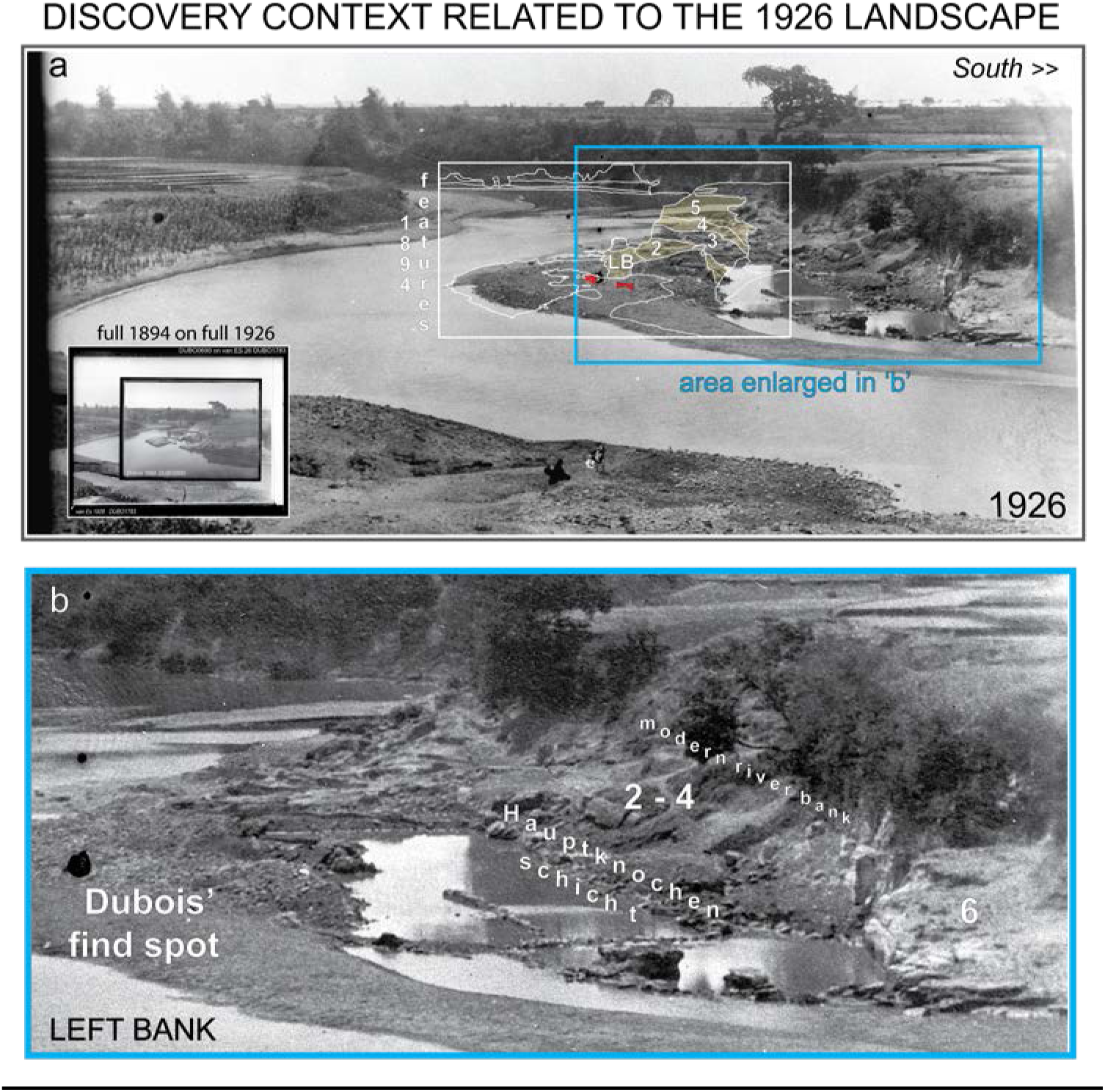
(**a**) The stratigraphic features of the *Pithecanthropus erectus* site, as they are seen in the 1894 photograph (Figure 4a), are superimposed here (as annotated white outlines) on a 1926 site photograph taken from near the same camera station (the inset, lower left, has the two images superimposed). The Dubois archive’s print of the 1926 image is annotated in his handwriting and includes an ink dot that coincides with the Skullcap discovery point that he marked on prints of the 1894 image (Figure 3c; the ink dot is absent from van Stein Callenfels’ 1929 published version of the 1926 photograph). (**b**) An enlarged portion of ‘a’ highlights the indurated condition of baulks and river-margin outcrops. They consist of the flat-lying Hauptknochenschicht and superjacent units **2**-**4** (which are not individually recognizable in the image). A cross-bedded remnant of **6** is still visible on the far west, but no **6** occurred at low-river level closer to the *Pithecanthropus erectus* discovery area. West of Femur I discovery point, the **LB**-**HK** was encountered in the 1892 20-m Trench, 1893 40-m Trench, 1896 Left-bank Pit and 1897 Downstream Pit (for 1896 and 1897, see Endnote D(iii)-(vi) and (ix), S II-A4c, -A4d, -A4f and -A4r). The 1926 photograph and another one from 1932 indicate that the spoils which had covered the discovery site in 1907-1908 were largely gone by the 1920s, giving geologists the first opportunity to examine the 1891-1893 excavation remnants since 1895. The source for the 1926 image is Naturalis scan DUBO1783.

**Figure 10.**
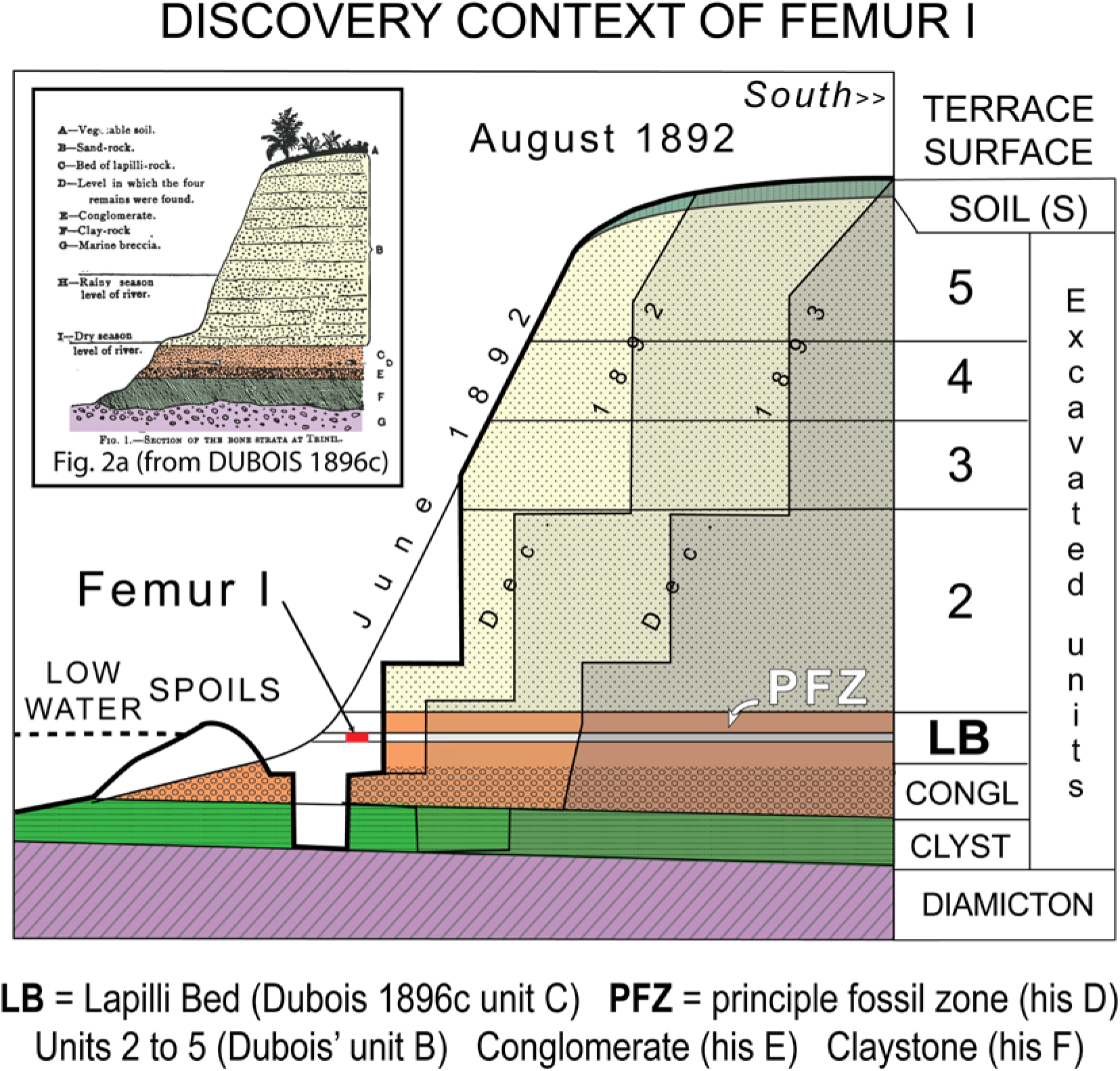
A diagrammatic cross section illustrates our reconstruction of the Femur I discovery context, based on Dubois’ records (also, Huffman et al. 2018), including his published 1895-1896 cross sections (inset) and unpublished materials analyzed in this paper (Figures 2a and 4a related our stratigraphic terminology to Dubois’). The site is depicted as it was during the August 1892 month of discovery. Faint additional indicators of the extent of excavations are shown for June 1892 through December 1893. The shape of the excavation profiles over time and thicknesses of individual stratal units are schematic (the strata above the **LB** were eight-to-nine meters thick). When the embankment was excavated in late 1892 and 1893 to the south of the August Femur I discovery point, the main bonebed was unearthed beneath hardened rock, confirming the stratigraphic placement of the Femur I. The situation at the end of 1893 was closely similar to that seen in the 1894 photograph (Figures 3c and 4a; S I Figure 3). Excavations in 1895 to 1907 dug farther southward into the embankment and encountered the same stratigraphic sequence (Figures 3-8). The right edge of this cross section is several tens of meters north of the present-day shoreline, judging from the 1899 map (Figure 3a) and 1926 photograph (Figure 9).

The basics are straightforward. Since the strata were nearly horizontal, the field supervisors’ provenience specifications were reported by elevation, and this gave Dubois the information about the stratigraphic origins of the finds that he needed. Today, the reporting allows us to follow the progress of the field operations, often week by week. Eyewitness accounts compensate substantially for the maps, profiles and other geological displays that Dubois and Selenka failed to make. The firsthand narrations commonly include the field taxonomic identity of prominent finds (Tables 1 and 2, Endnotes). This links well-known elements of Trinil fauna to the *Pithecanthropus erectus* Skullcap, and validates previous characterizations of the fauna (de Vos and Sondaar 1982).

The primary purposes of this paper are, first, to clarify the left-bank geology, and, second, to more fully document the stratigraphic attribution of the fossil discovered. These topics are addressed in two major text sections, LEFT-BANK GEOLOGY and DISCOVERY RECORD. Subsections include commentaries on related issues such as the site stratigraphy and Femur I provenience. As a result of our analysis, as detailed below, we have come to consider the discovery of the Skullcap, Femur I, and thousands of associated fossils to be the results of the rational actions of individuals who were skillful in excavating a well-understood stratigraphy under difficult operational circumstances. Their records justify presumptive acceptance of their geological and provenience conclusions. In DISCUSSION we briefly address a broader issue that intrigued Dubois and has interested us for several decades, the relation of Trinil to *Homo erectus* paleogeography in southern Sundaland.

## MATERIALS AND METHODS

This article contains a fine-grained analysis of primary, largely unpublished, documentary sources and 130 years of literature on the discovery, stratigraphic framework and paleontology of Trinil. The literature is anchored by Dubois’ own publications (1892-1908), as well as Selenka and Blanckenhorn (1911), and includes a number of useful analytical and summary works (Albers and de Vos 2010, de Vos 1985b, 1989, 2004, 2014, de Vos and Aziz 1989, Hooijer 1946a-1974, Joordens et al. 2015, Shipman 2001, Storm 2012, Theunissen 1985, 1990).

Primary narrative sources and much of the literature were composed in Dutch and German. English translation of these materials is provided in Endnotes and Supplementary Materials II (abbreviated ‘S II’), together with unpublished translations of Berkhout and Huffman (2020, 2021) and Huffman (2020). The geological evaluation of the 1894-1932 photographs of the left bank, which expands on our preliminary site evaluations (de Vos and Aziz 1989, Huffman et al. 2015, 2018), includes extensive presentation of interpreted images in Figures 1 to 14 and Supplementary Materials I, Figures 1 to 24 (abbreviated ‘S I Figures’).

Central to our evaluation is the premise that contemporaneous firsthand accounts of field observations, naïve of future events, are reliably interpretable in terms of the stratigraphic units and fossil species excavated (Table 1). Individual source documents often include observations relevant to both the stratigraphy and fossil recovery of excavated materials that have been removed completely. Crucial in this regard has been having a single compilation of translated Dubois materials (S II) around which multiple issues relative to *Pithecanthropus erectus* could be assessed concurrently and to which we could direct readers.

For example, the *Pithecanthropus erectus*-relevant stratigraphic entities that we term the **PFZ** and **LB** (Figure 2) are recognizable in many accounts that have been translations include: In letters that Dubois’ on-site field supervisors sent to him during 1891-1893 (S II-A1e); Dubois’ September 1892 memorandum and Third-quarter report submitted to the Indies government (S II-B4b; S II-B5b); his 1893 Second-quarter and Third-quarter reports and August-, September- and November memoranda (S II-B6c to -B6f and -B6h); and labels on several museum specimens (de Vos and Sondaar 1982, de Vos 1989). The **PFZ** and **LB** entities are also recognizable in Dubois’ (1894a; S II-B6j) *Pithecanthropus erectus* monograph (finished several months after excavation ended at Trinil), multiple versions of his 1895-1896 site cross section (e.g., Figure 2a; Dubois 1895b-1896g; S II-F1 to -F5), and a letter Dubois wrote about Femur I provenience the year he died (S II-E4).

Extensive analysis of the discovery record is feasible because the Naturalis Biodiversity Center, The Netherlands, preserves Dubois’ notebooks, diaries, fossil inventories, maps, reports to superiors or drafts thereof, academic papers, photographs, correspondence received, and drafts or handwritten copies of letters sent (known as the Dubois Archief or Dubois Collection). Overall ∼30,000 pages of paper, and >2,500 photographic glass negatives, film negatives and prints have been scanned by Naturalis, and each has a unique identifier, which we reference.

Special scans of particularly valuable photographic negatives, many 10-15 cm by 15-20 cm in size, were done at 4800 dots-per-inch for our project, so that we could resolve geological details at the *Pithecanthropus erectus* site (e.g., Figures 3 to 5, S I Figures 4 to 6, abbreviated S I Figures 4 to 6; Huffman et al. 2015, 2018; also, Albers and de Vos 2010). The earliest such photograph preserved at Naturalis was taken under Dubois’ direction on September 5, 1894, less than a year after the 1893 excavation ended (Figure 3c).

There are several significant limitations to the written materials. For example, the Dubois Collection contains the letters Dubois’ excavation on-site supervisors sent to him during 1891-1893 and 1895-1900, but no letters that he wrote to the field men. Similarly, while Dubois’ regular memoranda and reports to the Netherland Indies government are preserved in draft form (Dubois 1894d) and the reports were published (Dubois 1891a-d, 1892b-d, 1893a-b, 1894b-c), no communications from government officials about these submissions (if any were written) are part of the records available to us. Additionally, Dubois appears to have taken few field notes during his many visits to Trinil.

Vital information on the stratigraphy and paleontology of Trinil comes from the 1906-1908 Selenka Trinil Expedition (e.g., Selenka and Blanckenhorn 1911). Most of this material, originally written in German, is now available in English translation (Berkhout and Huffman 2021). Expedition geologist W.F.F. Oppenoorth took many valuable photographs in 1907, and most of them were never published (see Huffman et al. 2010a for biographical notes on Oppenoorth). However, his family saved negatives and prints, and generously donated many of them to Naturalis (J.M. Oppenoorth, personal communications, 2010 and 2015; also, Huffman et al. 2010a). Naturalis made high-resolution scans of key images, at our request, for the stratigraphic and provenience analysis we present here (e.g., Figure 8, and S I, Figures 8 to 10).

Our research also brought to light unpublished documentation in the Museum für Naturkunde, Berlin (MNB), from the Selenka Expedition. Among the informative records is a 1907 enumeration of field identifications of fossils, and the entries often give the stratigraphic origin of the finds. We refer to this document as the ‘1907 Listing.’ Most of the identified finds are attributed to **HK** layers (Table 2). Moreover, field numbers can still be read on many fossils retained by the MNB (L. Todd, pers. comm., 2016). This has opened a new era of paleontological study of the **HK** (e.g., Hill et al. 2015, Janssen 2017, Janssen et al. 2016). As for definition of the Pleistocene, we use the time scale approved by IUGS wherein the base of the Pleistocene is 2.58 Ma (Gibbard et al. 2010).

## LEFT-BANK GEOLOGY

### FOREWORD

The circumstances behind the discovery of *Pithecanthropus erectus* (*P.e.*) are difficult to verify in part because the find spots lie in the middle of the Solo River (Figure 1c). Visitors typically view the discovery area from a high bluff at the Trinil Museum, looking eastward up a broad river loop. Nothing then is visible of the Skullcap and Femur I pits and trenches, and trees obscure the former high-standing left-side river embankment to the south (Figures 2a,b). When the river drops towards dry-season low levels, former excavations begin to appear as excavation baulks, spoil piles and a scarred bedrock platform, as we have observed personally (Figure 6d; also, Alink et al. 2016).

The bedrock platform lies adjacent to former Selenka Trinil Expedition excavations, and contains remnants of their ‘Hauptknochenschicht’ (**HK)**. The HK at the platform is flat-lying, well-lithified pebbly volcaniclastic sandstone, which is prominently cross bedded, very poorly sorted, and locally contains large vertebrate fossils (Aimi and Aziz 1985) and lithic clasts (Huffman 2016; also, S I Figure 2). The **HK** qualifies as a bonebed because of the density of bioclastic materials embedded. In the late 1970s, the **HK** was mapped as a ‘KBGI’ unit, named in reference to Duyfjes’ (1936) Kabuh Formation (Soeradi et al. 1985; Figure 6d, S I Figure 18).

The crossbedding, coarse gravel and large bioclasts reflect bed-load transport of an ancient flood moments before deposition (Huffman et al. 2010a-b, 2012b). A lens underlying the **HK** includes laminated siltstone and matrix-supported sandy pebble-cobble conglomerate (the clayey ‘KBC’ of Soeradi et al. 1985), which potentially represent the same depositional events as the **HK**. Farther east, the KBC’ and ‘KBG1’ wedge out against a boulder-lahar unit (the ‘Pucangan lahar’ of Soeradi et al.; S I Figure 18). This is Dubois’ ‘breccia’ of 1895-1896 (Figure 2a; Huffman 2016). He evidently first recognized the stratigraphic relationship of the breccia to the bonebed in 1890 (S II-C5). In modern terminology, the ‘breccia’ is a volcanic diamicton; that is, an indurated boulder-bearing, matrix-supported conglomerate.

During lowest water levels, **LB** appears to be exposed northwest of the platform, towards the middle of river and close to the Dubois’ *Pithecanthropus erectus* discovery spots. To visualize the eight-to-nine meters of consolidated stratigraphic sequence that once held up high backwalls of the excavations, and to place bonebed outcrops into full stratigraphic context, we turn to an extraordinary trove of unpublished photographs, maps and eyewitness accounts dating from 1891 to the 1970s (Figures and S I Figures). The accounts mostly have an operational focus, but scrutiny of them discloses clue after clue about the site geology. Some of the same records are used in the DISCOVERY RECORD section, where the emphasis shifts toward the provenience of Femur I, other fossil species, and the origin of the bonebed.

### STRATIGRAPHY, 1891-1894

#### Firsthand reporting

Dubois was in Java from 1890 to 1895 and saw the 1891-1893 *Pithecanthropus erectus* discovery excavation firsthand. His 1895-1896 published accounts of the stratigraphic context (e.g., Figure 2a, b) were based on his own fieldwork, and the reporting of field supervisors G. Kriele and A. de Winter (KdW), as well an 1894 photograph Dubois had taken of the left bank (Figure 3c and 4a). The photograph constitutes an independent resource for assessing the geological framework of the discovery site from firsthand- and published-accounts (Huffman et al. 2015, 2018). KdW worked in the discovery excavations from 1891-1893, and Kriele continued alone in Dubois’ excavation until completed in 1900.

#### 1891 Skullcap Pit

The 1891 Skullcap Pit was dug into an ∼40m^2^ natural outcrop that seasonal low-water levels (LWL) on the Solo had been exposed. Fossils were *“chiseled out of the flat rocky ledge that reaches out … from the foot of the steep bank”* (Dubois 1896b: 251). When operations got underway in early September 1891, the outcrop contained a *Stegodon* tusk and cranium *in situ* (Figure 3b, S I Figures 3 and 4, S II-A1b). A particular concentration of fossils was encountered when the pit was deepened below the LWL into a subunit of the **LB** that we name the Principal Fossil Zone (**PFZ**). The **PFZ** became the *“level in which the four* [hominin] *remains were found”* in a Dubois’ 1895 site cross section (Figure 2a).

The Skullcap was found *“among hundreds of other skeleton remains, in the lapilli bed on the left bank”* (Dubois 1896c: 2, S II-F3i; also, -B2b and -B2d). The endocranial space of the Skullcap, which was unearthed in October 1891, was filled with indurated volcanic conglomerate (Figure 2c and 2d, S II-B2g). As reported around then, **LB** in the Skullcap Pit included bioclasts that are now attributable to well-known Trinil fauna species, plus fossil Testudine, Mollusca, wood and leaves (described more fully in the DISCOVERY RECORD; also, Table 1, Endnote A, S II-A1a, -A1b, -A1c, -A1d, -A1j, -B2b, -B2d, -B2e, -B2f and -B2g). Molluscan- and aquatic reptile fossils immediately indicated that the **LB** was a *“fresh water”* deposit (Endnote A(iii), S II-B2b).

#### Mid-1892 25-m Trench

By the end of 1891, the Dubois field team established an excavation protocol that all later operations followed: Strip off the fossil-poor bedrock making up the precipitously incised bank of the Solo in order to mine the fossil-rich **LB** near river level. To do this during the low-water season of 1892, KdW had to move westward from the Skullcap Pit along the shoreline (Figures 2b, 3a and 3c) and dig through the overlying beds (S II-B5a). As a result, the local stratigraphy and paleontology of the **LB** became even clearer during the 1892 excavation than it had been in 1891. The left-bank pit and trenches of 1891-1893 ultimately formed a narrow band ∼50m long. Annual excavations in those years were ∼30m^2^, 50m^2^ and 120m^2^ (Figure 3a).

When the river fell enough to expose the **LB** in late June 1892, the crew unearthed a *“harvest of bones …* [as] *plentiful as last year’s”* (Endnote B(i), S II-B3d). The finds noted in mid 1892 included key Trinil fauna species (Tables 1 and 3). In later publication, Dubois (1896c: 4) recounted, a *“new cutting was now made in the left rocky bank ….* [and] *bones were again found in great numbers, especially … in the same level of the lapilli bed, which had contained the skull-cap and the molar tooth, the left femur was found.”* He meant that Femur I (Trinil 3) was found at the same **PFZ** level that produced the 1891 Molar (Trinil 1) and Skullcap (Trinil 2) the year before.

**Table 3.**
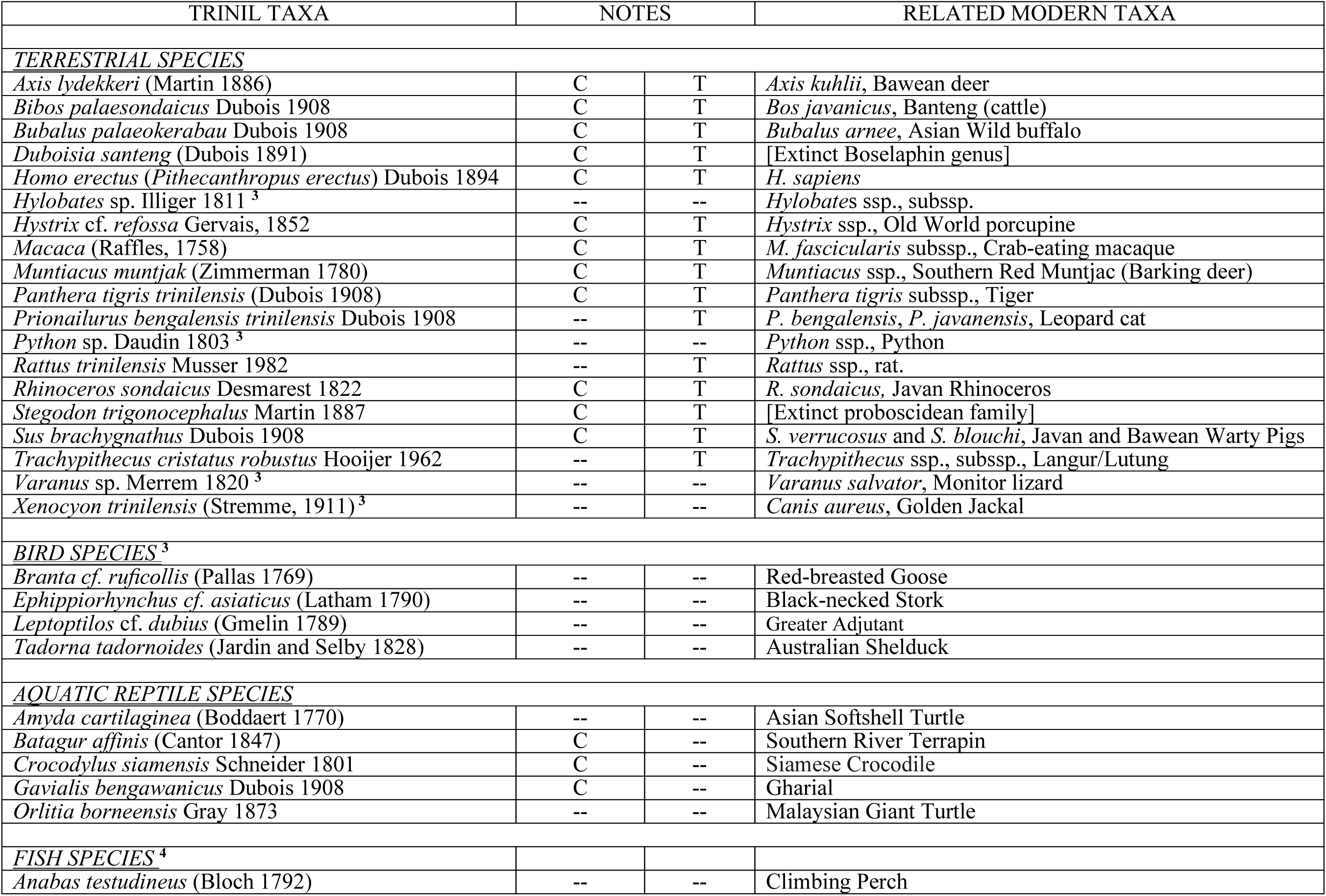
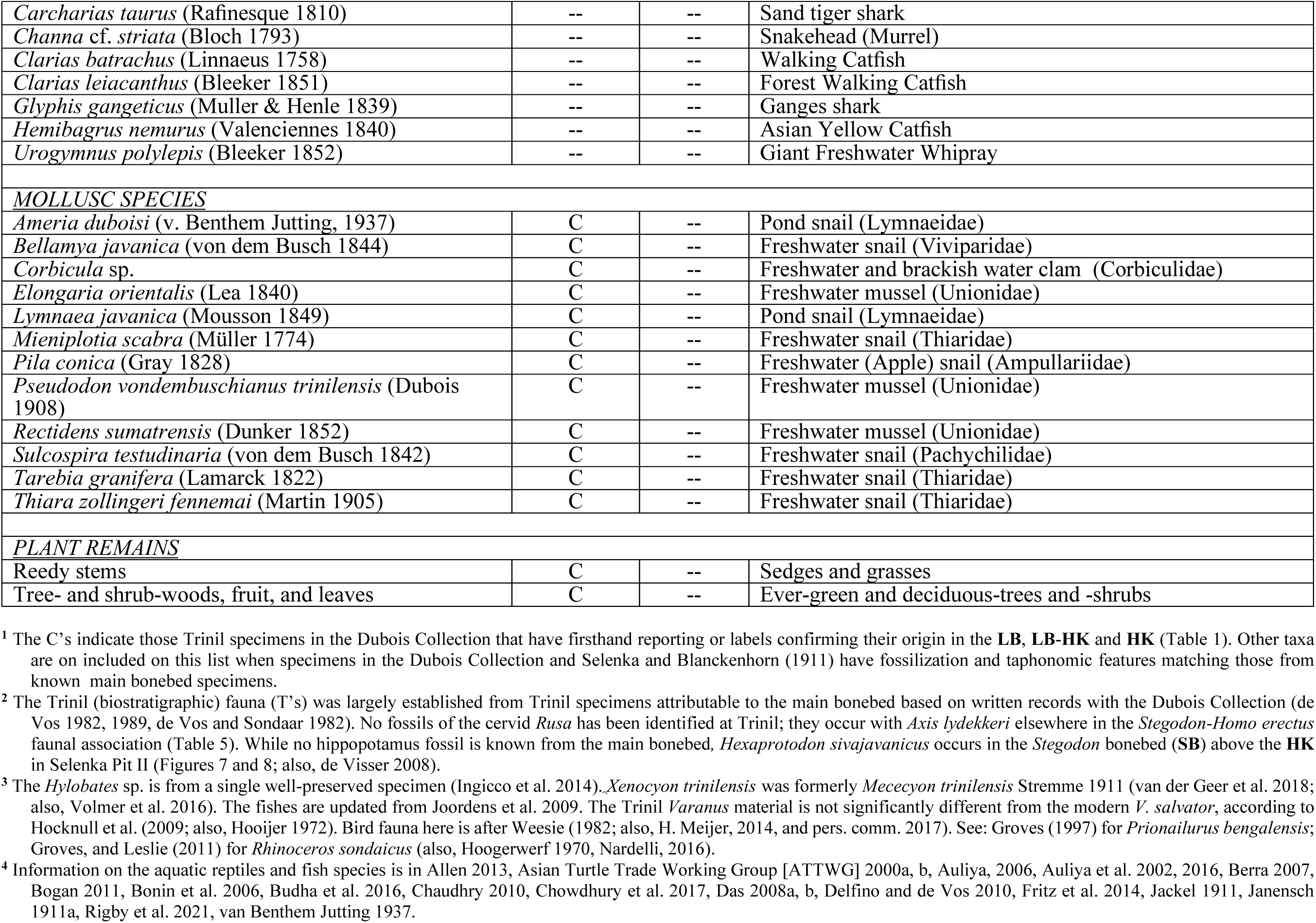
Terrestrial- and aquatic-species from Trinil. The C’s are confirmed main bonebed species; the T’s highlights Trinil fauna taxa. **^1, 2^**

KdW had given Dubois this firsthand provenience and stratigraphic specification for Femur I shortly after its discovery. Their September 7, 1892, letter reported: *“That bone* [Femur I] *was found on the same side* [of the Solo River] *as the skull* [Skullcap] *and also at approximately the same depth and even with the previous low-water level* [LWL]*, separated from each other by about 12 meters”* (S II-A2k). KdW were able to track the bonebed westward from the Skullcap Pit into the 1892 excavation (part of the 25-m Trench; Figure 3). Their wording, “same depth and … level,” referred to the **PFZ** subunit of the **LB**, as their later reporting makes clear.

KdW’s August 31 letter (S II-A2j), which was delayed in delivery to Dubois, added “if de Winter remembers it correctly the following bones were found nearby [to Femur I]: a mandible and tusk of an elephant” (Stegodon trigonocephalus). The abundance of the **LB** bioclasts at the Femur I site is further evident from the twenty-four crates of fossils shipped to Dubois by the end of August 1892; this volume was nearly equal to the whole 1891 output (S II-A2f, -A2g and –A2i). “The most abundant species continues to be the small Axis deer” during mid 1892, as had been the case in the Skullcap Pit (Endnote B(ii), S II-B4b). The Femur I context is analyzed further in DISCOVERY RECORD.

#### Mid-1892 field studies

The archival record reveals how Dubois and KdW informed themselves about the sedimentary features being excavated in 1892. Dubois spent 14-full and 8-partial field days in the Trinil area before Femur I was discovered (June 7-9, June 24-29, July 5-14 and July 18-20). He closely observed the fluvial bedding expressed in the modern sand- and gravel-bars along the meandering Solo River, and inferred corresponding paleocurrent patterns in the ancient sandy and gravelly formations exposed in its banks, including those under excavation at Trinil (S II-B3c and -C3 to- C5). He (1894a) made paleocurrent measurements from crossbedding to assess whether the **LB** was a product of the modern Solo River or an ancient watercourse.

Concerning the stratigraphy and sedimentology, Dubois’ June 1892 submission to the government reads, “at about 1 meter below the lowest water level of the river near Trinil, a blue-gray clay[stone] variety was found, immediately below the sandstone-like tuff,” which might refer to the **LB** itself (S II-B3c). The claystone “indicates a time of stagnant or very slowly flowing water,” while the tuffaceous facies “must have been deposited in faster flowing water” (S II-B3c; also, S II-C4). He noted elsewhere, “at Trinil bonebeds on both sides of the river (separated by a distance of 70m) have truncated thin [cross] beds dipping 30 degrees from west” (S II-C3). Two years later, Dubois (1894a: 1; S II-B6j) explained the conclusions he drew from observations such as these: The “left femur [Femur I] was excavated … at the same level [as the Skullcap] …. about 15 meters upstream in the direction of the current that [existed] during Pleistocene time;” that is, paleocurrent analysis indicated that the paleo-river flowed in a direction opposite to that of the modern Solo River.

G. Kriele understood Dubois’ innovative field methods for measuring paleocurrents. Before Femur I was found, Dubois was forced to return to his home at Tulungagung in East Java on July 20, 1892. He had a relapse of malaria contracted in Sumatra before the Java field project began. Dubois asked Kriele to go upstream of Trinil to record cross-bedding in strata that Dubois had not been able to examine (S II-A2f). To have success in this field assignment, Kriele must have been instructed on how to recognize lithostratigraphic formations, lithofacies differences within them, and bed sets and bedding particulars from never-before-seen outcrops of strata that matched those in the Trinil excavations. Once Kriele located appropriate upstream outcrops, he had to measure the direction of inclination of foreset laminations, having comprehended Dubois’ use of crossbedding as proxies for paleocurrent directions.

Kriele’s August 15 synoptic presentation of the results shows that the two men shared essential understandings about sedimentological matters, and had sophisticated geological field skills. Kriele presented small, annotated cross sections for the three localities, and each one showed the directions of internal cross-laminations within bed sets bounded by horizontal stratigraphic layers (S II-A2g). Kriele’s paleocurrent investigation fosters confidence that in their late August and early September exchange of letters KdW and Dubois established a precise mutual agreement on the sedimentary co-occurrence of the Skullcap and Femur I.

Cross-bed analysis seems to have been a core field practice for Dubois and his field supervisors, and serves as an indicator of a high level of sedimentary knowledge they all brought to the Trinil excavation. Dubois’ first cross-bed observations were made as soon as he started working the Kendeng Hills 1890. He had de Winter take note of crossbedding at Sangiran Dome in 1893, when *“very nice ‘oblique lamination’ … within horizontally bedded structure ….* [allowed] *the direction of* [the paleo] *current”* to be deduced (S II-B6e; also, S II-B6f). The orientation of cross-laminations continued to be on Dubois mind and among Kriele’s geological competencies until the end of excavation in 1900, when Kriele wrote: *“The thin slanted beds* [cross lamination sets] *about which you … asked me have been observed in 4 different places. They all dip approximately in this indicated direction S.W.* ////////// *N.E. but also N.N.E. …”* (S II-A4o).

Dubois’ strategy for the use of fluvial paleocurrents to reconstruct river-valley paleogeography was more than a half-century in advance of sedimentologists’ application of this approach (Potter and Pettijohn 1963/1977). KdW had sufficient geological training to put this strategy into practice.

#### Late-1892 excavation

Following the femur discovery, KdW expanded the excavation to the south, digging downward from the top of the embankment. Dubois expressed confidence in the expected stratigraphy based on their mid-1892 trenching, and mentioned the hard rock they anticipated:

> *This stretch of embankment* [destined to be the 25-m Trench] … *will have to be continued* [downward from the surface of the embankment] *to what we now estimate* [to be] *a depth of 9 meters, since … other remains* [of hominin] *are expected* [to occur in the **PFZ**] *below the water level during the East Monsoon* [low-water dry season]. *The sedimentary material to be removed is only soft enough for actual digging near the surface* [e.g., the soil of the terrace upland atop the embankment], *but for the most part we have a fairly hard sandstone-like andesitic tuff which* [is so well indurated that the material] *can only be removed with pickaxes and crow bars. …* [and] *very few other finds were made* [in the strata lying for at least 2.75m above the **LB**]. (S II-B5a)

Soon, KdW wrote to Dubois that they removed the upper 6.25m of the embankment and *“not a lot of bones have been found”* (S II-A2o). Evidently, ∼70% of the eight-or-nine-meter thick section overlying the **LB** had few vertebrate fossils, and was so lithified that the beds were tough to dig.

No more *“Chimpanzee”* remains had been seen by November 9 when KdW reported that *“the corners of the pit* [25-m Trench] … *are about 20 cm into the target bone layer* [**PFZ**]*”* (S II-A2p). At that time, the men must have stood before a tall excavated face ∼25m long. One point at the base of the scarp was just a meter or so away from the Femur I find spot (Figures 3a). This was the field situation when Dubois paid his last visit for 1892 on November 10-11.

A week later, the **PFZ** was inundated, ending the excavation season (S II-A2s). Dubois reported, *“no other* [skeletal] *parts of the* Anthropopithecus *…* [were found before] *rising water … forced us to finally abandon the work …, after having only excavated about 1/5th of the level of interest* [the **PFZ**]*”* (S II-B5c). Thus, the digging season ended with the **PFZ** exposed across the entire 25-m trench (∼50m^2^) but nearly 80% of the **PFZ** volume was still in place.

#### 1893 40-m Trench

River levels were anticipated to subside during the next excavation season in June and July, when Kriele’s crew began a 40-m-long excavation immediately to the south and southwest of the 25-m Trench (Figure 3a). Again, the men started by removing the soil on the terrace upland at the top of the embankment. The **LB** and the conglomerate beneath it were encountered in stratigraphic order during the course of digging down by horizontal increments. The penetration *“progressed extremely slowly because of the severe hardness of the rocks,”* a condition Dubois witnessed himself on June 6-8 and 26-28 (S II-A3a to –A3i and -B6c; Huffman et al. 2015).

The **LB**-**PFZ** in the 1893 40-m Trench was just as rich in large vertebrate bioclasts as the Skullcap Pit had been in 1891 (fully addressed in DISCOVERY RECORD). Included were remains referable to *Axis lydekkeri*, *Crocodylus siamensis*, *Duboisia santeng*, *Stegodon trigonocephalus*, and Testudines (Table 1; also, Endnote C(iv) and C(v), S II-B6b and -A3c/i). The *“the target layer”* (**PFZ**) not only had *“rather many bones”* but also a lot of *“wood”* and *“shells”* (Endnote C(vi), S II-A3cii), the fossil biota seen in 1891 and 1892.

In 1893, KdW gathered new stratigraphic information from their excavation. Below the **LB** *“a different layer emerges”* (Dubois’ dark-colored claystone of Figure 2b); its top contact *“slopes downstream”* to the west (Endnote C(viii), S II-A3e). Elsewhere in the Trench, the **LB** was *“becoming a little coarser downward”* with *“almost nothing in it”* by way of fossils (Endnote C(xi), S II-A3i). Five days later, the majority of the dig was deep enough to expose*“almost entirely coarse gravel”* with *“very little* [in it]*”* (Endnote C(xiii), S II-A3k). Shortly thereafter, they were *“more than a meter below the level of last year”* (which stopped in the upper **LB**), and digging *“almost entirely coarse gravel”* with few bioclasts (Endnote C(xiii) and C(xiv), S II-A3k and - A3l). Evidently, the sandstone and fine gravel of the **LB** graded downward into modestly fossiliferous conglomerate, as Dubois’ 1895-1896 cross section later showed; the conglomerate rested on the claystone which Dubois called “clay rock” in an English-language version of his cross section (Figure 2a).

Dubois’ notes for October 21, 1893, described the stratigraphic changes he observed during an on-site examination of the near-final 25-m and 40-m Trenches (S II-C7). Close to the Skullcap Pit, the *“top of black claystone* [was] *about 1 meter deeper than the Chimp skull”* which had been found in the stratigraphic middle of a *“lapilli bed about 2 meters thick.”* Along the backwall of the 40-m Trench, near the Femur I discovery spot 18m to the east of the first place he described, the crew had *“excavated 1.30m deeper into conglomerate after* [penetrating the] *alternating lapilli and sand”* in the **LB**. The conglomerate above the *“black clay*[stone] *layer”* was 0.5m thick. When Dubois made this inspection, a high excavated wall 40-m long loomed over the Femur I find spot (Figure 3b). His notes have essentially no comments about the strata exposed above the **LB**.

The apparent dip at the top of the claystone was ∼6° westward, not the near-horizontal attitude of the upper **LB**. The “*deepest spots* [in the Trench] *are about 3 meters below the current river level (East Monsoon level)”* (S II-C7). Above the claystone, the crew apparently had exposed as much as 3.7m of sandstone, lapilli-rich sandstone and conglomerate. The upper portion of this interval included the **LB** later described in publications (Figure 2b). Dubois’ notes do not address the distribution of other fossils within the lapilli-bearing sandstone. There is nothing in his reporting to suggest that the sandy and conglomeratic unit had an internal stratigraphic boundary denoting a substantial cessation of accumulation.

Dubois’ notes about his inspection of the trench are consistent with KdW’s previous stratigraphic reporting on the **LB**, but partially inconsistent with his own 1895-1896 published accounts (S II-F, Foreword). For example, he (1896b: 251, S II-F1 by) seemingly glossed over field observations when he published that lapilli *“predominate in the …* [a **LB**] *about 1-meter thick, which in turn transitions downward into a 1/2-meter-thick* conglomerate *bed that primarily consists of about walnut-sized rock fragments.”* Dubois (1896e: 4, S II-F4) also wrote that *“the rocky slopes on the banks of … Solo. … consist here primarily of … not very consolidated sandstone,”* rather than the hard rock described vividly in 1892 and 1893. Moreover, his cross sections showed the low-water well above the **PFZ**, despite field reporting to the contrary, and most of Dubois’ cross-section versions portray strata dipping slightly to the north, which conflicts with other evidence (S II-E4 and -F).

By year’s end in 1893, Kriele had shipped 25 crates of fossils and 6 crates of wood to Dubois (S II-A1h). The area of **PFZ** excavated in 1893 had been ∼170m^2^ (adding the unfinished 80% of the 25-m Trench to the 40-m Trench). Evidently, twice as much of the **PFZ** had been unearthed in 1893 as had been taken in 1891-1892, and most of the fossils Dubois recovered from the left bank over the three years came from the 1893 effort. The continuity of the fossil bone concentration across the three-years of excavations signaled that the **LB** was widely present below strata of the unexcavated embankment, setting the geological predicate for the excavations in 1895-1908.

#### 1894 photograph

Dubois documented the *Pithecanthropus erectus* discovery area photographically (Figure 3c), during a seven-day boat journey down the Solo (a trip noted in his diary). The September 1894 image and Dubois’ annotations on prints of it depict the 1891-1892 *P.e.* Skullcap Pit and the 25-m Trench in relation to the stratigraphic sequence visible in a degraded 1893 backwall (Figures 4a and 10, S I Figures 3 and 4; Huffman et al. 2015, 2018). Dubois never published the 1894 photograph. But his unpublished annotations depict the Skullcap Pit next to a sandstone ‘Ledge’ that lay just above river level (Figures 3c and 4a, S I Figures 3 and 4). The 1894 backwall was ∼35m north of the present-day shoreline, according to G. Kriele’s later mapping (Figure 6d, S I Figure 7). The Ledge produced vertebrate fossils in 1895 (Endnote D(i), S II-A4b, 15 October; also, S II-A4d).

To help track the local stratigraphy, we divide the beds identifiable in the 1894 photograph into informal lithostratigraphic units: Lapilli Bed (**LB**; unit **1**) is at the base of the scarp (e.g., at the Ledge); **2** through **5** make up the embankment; and **5** lies just below the terrace surface soil, **S**. Units **2** to **5** correspond to the *‘B-sand rock’* in the version of Dubois cross section that we present; **S** is his *‘vegetable soil’* (Figure 2a, S II-F3). The soil lay atop the embankment where the expanded 25-m Trench started in late 1892 and the 40-m Trench began in 1893.

The embankment contains no fluvial-terrace inset in either the 1894 photograph or Dubois’ cross section (Huffman et al. 2015, 2018). The soil-covered upland behind the lip of the embankment rose gently southeastward (Figures 4a, 6c and 7a, S I, Figure 2a). When Dubois (1896b) was asked at an 1895 public lecture about the presence of geologically younger deposits at the *Pithecanthropus erectus* discovery site, he described the Solo River valley as *“more of an eroding one than of alluvial deposits* [filling it]*”* (S II-F2). He apparently thought that the geomorphology in the discovery area originated by the Solo River downcutting through eight-to-nine meters of bedrock strata lying between the terrace soil (**S**) and the **LB** (also, S II-B6j).

The 1894 photograph provides independent evidence about the structural attitude of the strata dug during 1891-1893. The **1**-**5** sequence in the 1894 embankment exhibits little- or no-apparent dip in the photograph (Huffman et al. 2015). Any inclination having an east-west component of more than ∼3° would have appeared strongly accentuated in the photograph because it was taken at a shallow oblique angle to the excavation face. The photographic indication of horizontality matches Kriele’s September 1891 sketch showing **LB** outcrops lining both shorelines of the river (Figure 3b) and KdW’s explicit eyewitness accounts about the **LB** being horizontal in the 25-m and 40-m Trenches (S II-A1a, -A1e, -A1p, and -A1q; also, S II-B2h, -B2j, -B6e and -B7c).

The annotated 1894 photograph and Dubois’ and KdW’s written records of 1891-1893 confirm that the geological circumstances at the *P.e.* sites were straightforward, much as Dubois showed in published representations (Figure 2a,b).

### STRATIGRAPHY, 1895-1900

The left-bank stratigraphy, as described in records from 1893-1894, is confirmed by photographs of Dubois’1900 Trench. After he returned to the Netherlands in mid 1895, he sponsored a great expansion of the left-bank excavations (Figure 3a, S II-A4e), relying upon G. Kriele correspondence and sketch maps for firsthand information on the nature and provenience of the paleontological materials (S II-A4b to -A4l).

As the largest and last of these excavations was coming to an end in November 1900 (S II- A4n page 32), Dubois had three high-quality (large format) images taken of the left-bank site (Figures 4b and 5; S II-A4o, pages 6, 11, 25 and 32; de Vos and Aziz 1989). Surveyors spotted the camera stations on what became the 1900 Site Map (Figure 6a). Because the location of the camera stations is known and key landscape features are visible in multiple images, the photographic fields of view of the three photographs have been worked out (S I Figure 7a).

Dubois apparently did not write down the insights he gained from the 1900 photographs, but today, they are eminently interpretable geologically (Huffman et al. 2015, 2018). The sequence excavated in 1900 had the same horizontal structural attitude and regular sedimentary order as is visible in the 1894 photograph, even though nearly two-thousand square meters of the embankment separated the 1893 40-m Trench from the backwalls of the 1900 Trench (Figures 3a and 4, S I Figures 3-5, S II-A4m to -A4r).

The eastern end of the 1900 Trench (de Vos and Aziz 1989) had a prominence that we refer to as ‘East Point.’ The beds exposed there provide the best stratigraphic tie between the sequences seen in the 1894 and 1900 photographs (Huffman et al. 2015, 2018). Units **1**-**5** lie at approximately the same elevation relative to low water in the 1894 and 1900 photographs. In one 1900 image, several stratal markers at East Point are seen to have had little- or no-dip on orthogonal excavation faces (Figure 4b, S I Figure 5 camera station II, S I Figure 7a). The locale was ∼40m east-southeast of the Skullcap Pit, according to Dubois’ notes on an 1899 map (Figure 3a, S II-A4m).

Flat-lying strata is even clearer in a 1900 photograph taken from across the Solo north of the 1900 Trench (Figure 5, S I Figure 7a camera station III). That the strata had stood in near-vertical walls behind >100m of excavation in 1900 is a testament to the cohension of the sedimentary materials, if not their lithification.

The contact between unit **5** and terrace soil occurs at lower elevation at East Point than in the western 1900 Trench (Figure 4b), apparently due to shallower erosion there during the formation of the terrace. A sixth indurated stratigraphic unit, **6**, is identifiable in the western 1900 Trench. The unit overlays **4** and **5** across an erosional surface (Figures 4b, 5 and 7; also, S I Figure 11). Unit **6** has prominent large-scale internal crossbedding, and backfills against and oversteps onto the erosional base. The upper portion of **6** is truncated at the soil S developed on the terrace upland, indicating that the terrace was a product of erosion rather than substantial fluvial accumulation.

The lithofacies evident in the 1894 and 1900 images appear to range from mudstone to conglomerate and diamicton, and seemingly reflect varied depositional conditions, even within individual stratal units (S I Figures 3-6 and 9 to 11a). Judging from the photographs, the use of modern geological methodologies in Dubois’ excavations would have led to the through-going tracing of ithostratigraphic bed sets across the 1900 Trench (Figures 4 and 5).

Taken together, the 1894 and 1900 photographs show that the embankment removed by the expansive 1895-1900 excavations south of the *Pithecanthropus erectus* discovery points consisted of a uniform, indurated, flat-lying stratigraphic sequence (units **1**-**5**) that included a younger unit **6** in the western 1900 Trench (Figures 4 and 5).

### STRATIGRAPHY, 1907-1908

The Selenka Trinil Expedition photographs and records confirm the persistence of a vertebrate-fossil concentration lying near the seasonal low-river level of the Solo in a stratigraphic position beneath the same eight-to-nine meters of indurated beds that Dubois’ 1900 Trench had encountered (Figures 7 and 8; also, Selenka and Blanckenhorn 1911: Fig. 2 and 20). Units **2**-**5** are securely recognizable in the 1907 images because they show 1900 backwalls. The Expedition workmen used *“pickaxes”* to remove the hardened flat-lying strata, just as Dubois’ excavators had done in the 25-m and 40-m trenches (Figure 8b; Oppenoorth 1911: xxxiv, Berkhout and Huffman 2021: 23-24). The Selenka records are particularly valuable because they contain stratigraphic and paleontological details for the left-bank excavations nearest to the modern shoreline, including the baulks of Selenka Pit II (e.g., Figure 6b and S I Figures 2 and 7b).

During mid 1907, M. Selenka and W.F.F. Oppenoorth (the Expedition’s first supervising field geologist) readily recognized the bone-rich *Pithecanthropus erectus* bed in outcrops near the low-water level at East Point, and he documented this photographically (Endnote F(viii), S I Figure 8). He (1908a,b, 1911) took many other useful images of the new left-bank operations from boats and the opposite shore, and a number of his photographs have not been published before this (S I Figures 8-10; also, Berkhout and Huffman 2021).

E. Carthaus, who assumed Oppenoorth duties in August 1907, termed the bone-rich stratum ‘Hauptknochenschicht’ (**HK**) because it clearly had the highest concentration of vertebrate fossils of any widespread bed at Trinil (e.g., Table 2, Endnote F(viii)). C.M. Dozy (1911a: xli) located the northern termination of the **HK** in Pit I and another depositional boundary ∼200m away at the eastern end of Pit II, where the *“main bone bed had completely pinched out”* against a local paleotopographic prominence formed in breccia (Berkhout and Huffman 2021: 47). He saw enough of Pit II in 1908 to correct 1907 drafts of the Expedition’s well-known cross-section along the left bank, the *“Idealized Profile … after Carthaus”* (Selenka and Blanckenhorn 1911: 7; also, Branca 1908, Berkhout and Huffman 2021: 63).

Dozy concluded that the **HK** was a thin sedimentary lens which filled a paleotopographic low at the top of a sequence of diamictons (which later were included in the Pucangan Formation by Duyfjes 1936). Dozy (1911a: xli) also reported that survey instruments had *“proven by leveling”* that the **HK** in Pits I and II was at about the same elevation (Berkhout and Huffman 2021), thus confirming the flat-lying relationship that Kriele reported between the **LB** on the right- and left- banks in 1891 (Figure 3b). The **HK** in Pits I and II were also closely comparable in (i) stratigraphic thickness (<1m), (ii) lithofacies (sandy and conglomeratic), (iii) degree of consolidation (lithified and indurated) and (iv) abundance of large-sized vertebrate bioclasts. These similarities support the inference that the **HK** across the Trinil site was a single bioclast-rich deposition unit (Selenka and Blanckenhorn 1911, Berkhout and Huffman 2021).

There is little information on the 1908 Pit II along the left bank. Dozy (1911a: xli) explained that in 1908 they “*could not work further to the east …* [of the 1907 Pit II or] *towards the south* [into the left embankment because] *the land was privately owned ….* [so that Selenka’s work in 1908 largely] *focused on the right-bank”* Pit I (Berkhout and Huffman 2021: 47). We take Dozy’s statements to indicate that the **HK** did not pinch out southward, and the bonebed should continue beneath the modern left bank. Dozy’s emphasis on Pit I helps account for why only one 1908 photograph of the left bank is known (S I Figure 12).

The fossil assemblages in Pit I and II differed in taxonomic proportions and vertebrate-bioclasts density, but the two excavations produced much the same assemblages of species (Table 2). Furthermore, the Trinil fossils exhibit uniform stony fossilization and taphonomic parameters in the Dubois Collection at Naturalis, Leiden, and Museum für Naturkunde, Berlin (Selenka Expedition; e.g., L. Todd, pers. comm., 2015). These assemblages represent the entire span of left- and right-bank excavations from 1891 to 1908 (Table 1), and appear to represent one fauna (de Vos and Sondaar 1982), as documented elsewhere herein (Tables 1 to 2, Endnotes A to H, S II). The Selenka records indicate that the average fossil-density in the **HK** of Pit II was ∼2.7m^-2^ (Table 2, footnote 4). If this density is applied to the area of Dubois’ 1891-1893 pit and trenches (which totaled ∼54% of the size of Pit II), he collected ∼540 specimens from those early excavations.

The Selenka Expedition found vertebrate fossils at some points above the **HK** (Table 2). The most prominent shallow concentration was unearthed soon after excavation of Pit II began in 1907, when the field crew encountered a cluster of vertebrate fossils ∼5m above the **HK**. We place this deposit, the *Stegodon* bonebed (**SB**), in our lower unit **5** (Figure 8, S I Figure 9 and 10; Oppenoorth 1911: xxxii, Berkhout and Huffman 2021: 34). The **SB** overwhelmingly consisted of the disarticulated and dispersed elements of a *S. trigonocephalus* individual, which had been embedded in a clayey conglomerate with some crocodile and ‘fish ?’ remains (Table 2), which indicate aqueous deposition. The **SB** apparently formed when *Stegodon* bones accumulated in an erosional swale cut into an unconsolidated unit 4. No fossil concentrations are known to have occurred within the ∼3m of deposits and ∼1m of soil above the **SB** in Pit II.

The 1894-1907 photographs reveal sedimentary features further indicating that fluvial conditions dominated deposition of units **2**-**5**. For example, unit **4** contained inclined bedding and had a truncated top, reminiscent of bar development; the **4**-**5** contact exhibited soft-sediment deformation, as did other **2**-**5** stratigraphic levels, indicative of new accumulation on uncompacted substrates (S I Figures 4 to 6). Similar features are evident in Selenka Pit I. The bluff to the south of right-bank Pit I stood at ∼16m above low water, rising significantly higher in elevation than the upland on the left bank near Pit II (Figures 6b, 6c; S I Figure 7a). Dubois and Selenka could see, and presumably examined, prominent natural outcrops that held up the right bank below today’s Trinil Museum (S I Figure 14; also, Widiasmoro and Boedhisampurno 2001).

However, the stratigraphic successions above the **HK** in Pits I and II do not appear to have been correlated readily. In 1907, the Selenka Expedition chose separate numbering schemes for stratigraphic units in the two excavations (Table 2; Selenka and Blanckenhorn 1911). *“Blue-grey ash with … intercalated clay …* [and] *thin beds* [lenses] *… formed of leaf remnants*” (the Main leaf bed) occurred between ∼0.35 and ∼4.10m above the **HK** in Pit I, but this leaf-rich facies was rarely identified in Pit II (Schuster 1911b: 4; also, Branca 1908, Carthaus 1911a, Dozy 1909, 1911b). None of the other known records of the Expedition contain a bank-to-bank correlation of the post-**HK** units. The lithological information that the Selenka geologists did provide contains indications that the sequences on both banks on the two sides of the river are consistent in representing long periods of post-**HK** lowland accumulation which was impacted by volcanic eruption and periodic lahar deposition (Berkhout and Huffman 2021).

In summary, the photographs and records of the Selenka Expedition confirm the persistence of the main bonebed (‘Hauptknochenschicht,’ **HK**) on left bank, where the bonebed continued to occur below the same eight-to-nine meters of indurated strata that Dubois’ 1900 Trench had exposed (Figures 7 and 8; also, Selenka and Blanckenhorn, 1911: Fig. 2 and 20). This sequence should be present beneath the south-shore embankment today. The Selenka left-bank excavation (Pit II) encountered a localized *Stegodon* bonebed ∼5m above the **HK** (Figure 8). The **SB** stratigraphic level might also be identified in the modern embankment, even if **SB** fossils are missing. The Selenka record establishes that the **HK** was flat-lying between Pits I and II, just as the **LB** was seen to be on the left- and right-banks when excavations began in 1891 (Figure 3b).

### STRATIGRAPHY, 1920S AND 1930S

#### 1926 photograph

The Dubois collection has a 1926 image that shows the *Pithecanthropus erectus* (*P.e.*) site on the left bank at a critical point in time. The photograph helps to relate units **1**-**6** to the rocks cropping out along the south shore today (Figure 9). Dubois attributed the image to L.J.C. van Es, an Oppenoorth colleague at the Geological Survey of the Netherland Indies (Huffman et al. 2005 and 2001b). Van Es (1927, 1929, 1931) was making the first proper geological map of the Trinil area (along with colleague M.Th. Wiessner; Berkhout and Huffman 2020). The 1926 photograph makes at least four significant contributions to understanding the left-bank geology.

First, while annotating his print of the 1926 image, Dubois put an ink dot in the gravelly bank north of the left-bank baulks and trenches (S I Figure 3d). Because the camera station in 1926 was at nearly the same high point on the right-bank bluff that the photographer had stood in 1894 (near the current Trinil museum), the excavated features and stratigraphic units evident in the 1894 photograph (Figure 3c) can be set into the 1926 landscape (Figure 9a). Moreover, when the 1894 and 1926 annotated images are overlain, the Dubois’ ink dot coincides with the Skullcap discovery point that he marked on the 1894 image (Figure 9b). The ink dot evidently records Dubois’ effort to situate the *Pithecanthropus* discovery into the landscape visible on the left bank 35 years after he last visited Trinil.

Second, the 1926 photograph makes clear that van Es had extraordinarily good rock outcrop when he mapped the left bank. As he understood the stratigraphy there, a vertebrate-bearing sandstone unit rested on a black-clay map unit, which in turn overlay a boulder-tuff (volcanic diamicton) unit. Van Es saw the sequence as dipping southward. He did not recognize a terrace-deposit unit atop the dipping strata. By the time he published his final map of Trinil, van Es (1931) had mapped much of the Kendeng Hills geologically, improving upon the maps prepared by Verbeek and Fennema (1896; also, S II-C and S II-F) who worked in Dubois’ day.

Third, the 1926 photograph permits a new geological interpretation of the left bank. Erosion-resistant **HK** (if not also the **LB**-**HK**) sat just above low water, and blocky outcrops of indurated rock underlay the river bank where we project units **2**-**5** occur (Figure 9b). The rocky nature of the baulks and outcrops convincingly support the inference, drawn from older site photographs, that the strata removed by excavation along the left bank were strongly lithified, although by 1926 the former high-standing excavation faces of 1900-1907 had been reduced to irregular river-bank outcrops. Units **2**-**5** cannot be identified specifically in the image.

Fourth, unit **6** is seen in 1926 to form a prominent remnant near low water in the far southwest of the old excavation area, but is not discernible near low-water river elevations where the 1899 Trench and most of the 1900 Dubois Trench were dug (Figure 9b). Unit **6** therefore did not appear to occur farther north and close to the **LB** level in the 1892-1893 pits and trenches. This is consistent with firsthand accounts that the **LB** occurred throughout the 25-m and 40-m Trenches, and the **LB**-**HK** was widely present in later trenches, as described below. Thus, there is no reason to expect that unit **6** was at **LB** level anywhere near the *P.e.* discovery points.

#### 1930s excavation and mapping

Following van Es’ work, the Survey conducted excavations at Trinil in 1931-1932 (von Koenigswald 1934/1935). A large Survey dig is seen on the right bank in a W.F.F. Oppenoorth photograph (S I Figure 15a). Evidently, large-scale trenching was not re-initiated on the left bank then, and Dubois’ 1900 Trench and Selenka Pit II has bordered the left shore of the Solo ever since (S I Figure 2).

In May and June 1933, the Survey charged geologist J. Duyfjes (1933, 1936) with the task of tying the geology of the greater Trinil area into a regional lithostratigraphic framework that he and his Survey colleagues, such as van Es, had developed for the Kendeng Hills (S I Figures 16 and 17). Duyfjes (1936: 147) attributed the strata on the left bank near the *P.e.* site to his self-defined Kabuh Formation. The main bonebed was placed near the base of the Kabuh, and soil topped the section of the Formation there (S I Figure 17b). His mapping is consistent with Dubois cross section in this regard (Figure 2a).

Over a broader area around Trinil, Duyfjes mapped substantial thicknesses of both flat-lying terrace deposits and tilted-bedrock formations, the latter including the Kabuh Formation (S I Figure 16). He (1933: 13) characterized the Kabuh here as *“primarily … andesitic sandstones and tuff sandstones…* [which] *mostly contain clearly rounded grains and often show some crossbedding. …* [They] *sometimes alternate with conglomeratic beds …* [and] *ash tuff….* [and] *often contain fossil bones and fresh water mollusks,”* reflecting fluvial transportation and deposition (Berkhout and Huffman 2020: 9). Duyfjes attributed diamictons underlying the Kabuh to his Pucangan Formation (a unit to which Dubois’ “breccia” unit, Figure 2a, would belong). The larger clasts in the Pucangan were embedded in very poorly sorted sedimentary matrix typical of lahar deposits (a facies which van Es termed “boulder tuff”).

By mid 1933 Duyfjes had observed and mapped outcrops of the structurally folded Kabuh and Pucangan Formations, as well as older carbonate-bearing and younger volcaniclastic formations, along the southern Kendeng Hills for an east-west distance of ∼175km (Figures 1b and 13). He ultimately produced a map that extended from a point ∼7km west of Trinil through the *P.e.* site to and Kedungbrubus where Dubois made the first *Homo erectus* find in 1890 and the Perning site where the fossilized Mojokerto child’s skull was unearthed in 1936 to the shorelines of the Madura Strait (for Kedungbrubus, see Duyfjes 1933, 1934, 1935, 1936, 1938a-d; also, van Es 1931, Huffman 2001b, 2016, 2020, and Huffman et al. 2005; see S I Figure 21 and S II-D2 regarding Kedungbrubus).

Based largely on Duyfjes’ mapping, the acknowledged stratigraphic sequence of the greater Trinil area thereafter became (from older to younger) Kalibeng Formation, Pucangan Formation, Kabuh Formation and Notopuro Formation, all of which are unconformably overlain along the Solo River valley by terrace materials (S I Figure 16).

#### 1920s-1930s, key impacts

Excavation and geological mapping by the Geological Survey in the 1920s and 1930s impact understanding of the *P.e.* discovery site in three principal ways. First, the 1931-1932 Survey excavations were evidently done along on the right bank, and the Dubois 1900 Trench and Selenka Expedition Pit II were the last large excavations dug along the modern left-bank. Second, erosion-resistant remnants of the bonebed and units **2**-**6** were still visible in 1926 along the south side of the Solo near the *P.e.* site and these units undoubtedly occur beneath the left bank today. Third, Survey mapping did not recognize terrace deposits on the left bank between the Kabuh Formation and the soil on the terrace upland adjacent to the old excavations. This is consistent with Dubois’ representation of the geologic situation at the 1891-1893 excavation site (Figure 2a).

### LATER STRATIGRAPHIC STUDIES

#### 1:250 map of 1977

The left bank was investigated rigorously in 1977 by the Indonesian-Japanese project “Quaternary Geology of the Hominid Fossil Bearing Formations in Java….,” a multi-year effort involving multiple field teams who analyzed fossil sites in Kabuh and Pucangan Formations from Sangiran Dome to Perning (Watanabe and Kadar 1985). The Trinil project team made a 1:250 geological map of the left bank and measured sections near the *Pithecanthropus erectus* site. The team agreed with Duyfjes that the main bonebed belongs in the lower Kabuh Formation (I.J.R.C.P. 1979, Soeradi et al. 1985). The outcrops that they assessed included the erosion-resistant baulks and other exposures that we refer to as the **LB**, **LB**-**HK** and **HK** (Figure 6d, S I Figure 18). Three measured stratigraphic sections are near the 1891-1908 excavations. The stratal units in these sections portray little of the stratigraphic continuity widely seen in the 1900-1907 photographs, perhaps because outcrops in the 1970s were too limited. When a 2010 team from the Sangiran archaeological museum excavated flat-lying, cross-bedded sandstones in the area of the Soeradi et al. map on the left bank (∼100m east of the 1891-1908 excavation), they attributed the strata to the Kabuh Formation (Widianto 2012).

#### Later mapping in area

The geological efforts of the Indonesian-Japanese project, credible as they are, have not settled stratigraphic matters at Trinil, broadly considered. First, various geological teams mapped the Kabuh Formation in different outcrop patterns and handled terracing in the Solo River valley in different ways (S I Figure 19; Berghuis et al. 2021, Datun et al. 1996, de Genevraye and Samuel 1972, de Terra 1943, I.J.J.S.T. 1992, Sartono 1976, Soeradi et al. 1985, Susanto et al. 1995, van Bemmelen 1949, Widiasmoro and Boedhisampurno 2001; Berkhout and Huffman 2020, 2021, and Huffman 2020 have translations of key works).

Second, none of the stratigraphic formulations resolved a conundrum that Duyfjes’ work first revealed (Huffman 2016). His mapping of the left bank, near the *Pithecanthropus erectus* discovery point, does not have a terrace-deposit formation lying between the Kabuh Formation and the soil zone atop the terrace upland, as noted above. However, on the right bank, he mapped flat-lying terrace deposits around the former Survey and Selenka trenches, knowing that a year before, the Survey had exposed eight-to-nine meters of strata overlying the **HK** in excavation (S I Figure 17). He also mapped a substantial covering of a terrace-deposit formation to the south of the Survey trench along the right bank, where he found terrace deposits resting on top of the Kabuh Formation (S I Figure 17; also, S I Figure 14). His terrace-deposit formation overlies the dipping strata of the Kabuh to Kalibeng Formations across much of the right bank (S I Figure 16).

Third, Duyfjes illustrated the key stratigraphic relationships at Trinil in a north-south cross section which passes through the left-bank excavation area, but he erroneously put ∼9° south dip on the Kabuh-Pucangan contact there (S I Figure 17b). This would equate to >5m of stratigraphic elevation across the 1891-1907 left-bank excavations where photographic evidence demonstrates near horizontality (Figures 4, 5, 7 and 8, and S I Figures 2 to 5, 7 and 8). Moreover, the Selenka Expedition had shown that the **HK** was flat-lying over the ∼200m distance between the right-bank Pit I and left-bank Pit II (Figure 6b; S I Figure 7). Duyfjes (1936) admitted that the Kabuh and terrace deposits *“cannot be readily distinguished”* lithologically (Huffman 2020: 13).

### TERRACE CONFUSION, 1980s and 2021

Uncertainty about the stratigraphy of the left bank has centered on the contention that Dubois, Duyfjes and Soeradi et al. missed terrace fill there. G.-J. Bartstra asserted that geologically young fluvial fill (presumably flat lying) sat atop the Kabuh in the bluff *“at the spot where the skull-cap and thigh-bones … were found,”* so that, *“Dubois not only excavated in the Kabuh Formation, but in terraces of the Solo River,”* by which he meant terrace deposits; therefore the Trinil *“fossil remains … come from two stratigraphic units that differ considerably in age,”* making the collections *“a mixture of fossils from the Middle Pleistocene Kabuh beds and the Upper Pleistocene and sub-Holocene Solo Terrace sediments”* (Bartstra 1982: 97, 1983: 330, 335, 336).

Despite the offer of provenience documentation (de Vos et al. 1982, de Vos 1989, Sondaar et al. 1983), Bartstra’s contentions have had a long-lasting influence on opinions about the Trinil hominin discoveries (e.g., Bartsiokas and Day 1983, Berghuis et al. 2021, Cartmill and Smith 2008, Day 1984, Dennell 2008, C. Groves in Bellwood 2017, Hooijer and Kurten 1984, Kennedy 1983, Klein 1989, 1999, 2009, Lubenow 2004, Ruff et al. 2013, 2015, van der Geer et al. 2018; also, Day and Molleson 1973).

Bartstra had no direct evidence that terrace deposits had been encountered in the Dubois or Selenka excavations on the left bank. Remnants of the main bonebed lay in the middle of the river where strata which originally overlay it were long gone. He missed the 1977 geological mapping of the existing remants (S I Figure 18; I.J.R.C.P. 1979). Judging from the site photographs of 1900 and 1907, the terraced upland near the *Pithecanthropus erectus* site formed largely by erosion of bedrock units 5 and 6, rather than accumulation of valley fill (Figures 2a, 3c and 4).

Bartstra admits he did not find a break in the stratigraphic column between the underlying Kabuh and supposed terraces fills at the Trinil site (1983: 333 and 334). All he could know for sure is that elsewhere terrace deposits and loose gravels occurred above the incised escarpments on the terrace treads along the Solo River, and elsewhere some terrace fill was present in side banks of the river and its tributaries (Bartstra 1977, 1982; de Terra 1943, Lehmann 1936, Oppenoorth 1936, Sartono 1976, ter Haar 1931, 1934a,b), as later confirmed (I.J.J.S.T. 1992, Rizal 1998a,b, Rizal et al. 2020, Sidarto and Morwood 2004, Saefudin et al. 1995, Suminto et al. 2004, Susanto et al. 1995).

Moreover, Bartstra’s assertion about the Trinil fossils representing mixed geological ages runs counter to the faunal and taphonomic uniformity in the Dubois and Selenka fossil collections (e.g., Endnotes). The bony elements in the museums include massive *Stegodon* and bovid specimens, and consistently exhibit little indication of subaerial exposure, fluvial abrasion and mixing of taphonomic- or faunal-components (Hill et al. 2015, Huffman et al. 2018). The matrix adhering to vertebrate specimens, and their consistent stony fossilization, indicate the discoveries had been made in indurated conglomeratic sandstone (e.g., Figure 2c,d, S I Figure 20). The lithic materials seem to represent a single depositional unit, judging from hand-specimen examination.

#### Berghuis 2021

Bartstra’s miscalculations did not end confusion about terraces on the left bank. Berghuis et al. (2021) conclude the Pucangan and Kabuh Formations, together with older dipping formations around Trinil, are overlain unconformably by horizontal strath terraces at higher elevations and three Late Pleistocene valley fills at lower elevations. The seven terrace units span 27m in elevation (T2 is 13m above the Solo riverbed; T3 is 17m above; T4 is 18m; T5 is 23 m; T6 is 25m; and T7 is 27m above). The highest-elevation surface in the vicinity is twice the height of the terrace surface immediately south of the 1900-1907 left-bank excavations.

Berghuis et al. largely follows the terrace interpretations of Lehmann (1936) and several later publications (summarized in Berkhout and Huffman 2020; also, S I Figure 19). But Berghuis et al. introduce the idea that lower-elevation terrace fills followed three regional incision events which cut the Solo River valley nearly as deeply as it flows today. One of their fills (T2) is postulated to hold up the left bank lying south of the strata excavated by Dubois and Selenka. Berghuis et al. correlate this inferred fill (and apparently including the **HK** below) to the early Late Pleistocene strath terrace containing the *Homo erectus* bonebed at Ngandong, ∼10km away in the Solo River gap (Berghuis et al. Figures 5 and 10 illustrate this correlation, but do not show the **HK** inside T2 on the left-bank, obscuring their intended stratigraphic placement of the main bonebed).

The conclusions of Berghuis et al. face considerable geological and paleontological obstacles. First, the extent of lithification in units **2**-**6**, which is clear from site photographs and reports of “hard” digging in the 1892-1907 excavations, is not in keeping with a Late Pleistocene valley fill. Generally, thin sedimentary deposits of Late Pleistocene age do not indurate to the observed degree without substantial depositional overburden, and there would have been none according to the Berghuis et al. interpretation.

Second, the Ngandong strath deposit and the Perning bonebed (Table 6-B and 6-E) were distinctly less lithified than units **2**-**6** appear to have been. In digging the Ngandong *Homo erectus* site, traditional short-handled, flat-bladed agricultural mattocks, called ‘patjols’ in Java, readily cut through the ∼3m of strath-terrace sands; patjols and flat-tipped pits were able to dig vertical faces in the lithified Perning fossil sandstone, which is part of the structurally folded Pucangan Formation at the relocated discovery site of the fossil Mojokerto *Homo erectus* child skull (OFH pers. observation).

Third, taphonomic and faunal evidence favors a bedrock having been dug in the 1891-1908 left-bank excavations. The Trinil fossils are more petrified than those from the Ngandong *Homo erectus* bonebed and the Perning fossil-skull bonebed generally are (OFH pers. observation). The Trinil fauna, which is defined on the basis of specimens from main bonebed (Table 2), occurs in ∼0.9 Ma Kabuh Formation strata at Sangiran Dome (Table 6-G), is older than the fauna in the Ngandong strath terrace (Table 6-B). These considerations strongly suggest that Berghuis et al. errored in identifying the beds in the modern embankment along the left shore as geologically young valley fill (and hence were wrong to suggest the that 1891-1893 discovery excavation contained fill).

Other unsettled geological issues form barriers to accepting key conclusions Berghuis et al. draw about the geology at Trinil. Their illustrations show 5-8° southward dip in the Kabuh Formation and Pucangan Formation on the right bank around Selenka Pit I (see Berghuis et al. Figure 4, 15 and Supplement 2A and 5), despite the horizontal structural attitude that the Selenka Expedition and Duyfjes observed thereabouts. Berghuis et al. also show exposures of dipping Kabuh and Pucangan strata in the right- and left-banks of other parts of the greater Trinil area where meters of horizontal terrace fills are said to cover these bedrock formations.

Conflicting interpretations might be expected to arise from Berghuis et al. approaches to mapping. Their geomorphology portrayal comes from digital elevation modelling of satellite data (seen in their Figure 2A with the potential effects of elevation-error unspecified). And Berghuis et al. map of the dipping bedrock formations is largely a revision of Duyfjes’ work with his superjacent terrace deposit formation eliminated (compare their Supplement 5 to Duyfjes’ maps in S I Figure 16).

Also troubling is Berghuis et al. reliance on poorly illustrated and complex stratigraphic inferences in the area to the south of the Trinil Museum (S I Figure 16, ‘1’). There, they report ∼9m of fill (T4) atop inclined strata 10m above the river, and also, 11m of fill (T2) on dipping beds at 2.5m above the river. In these relations, Berghuis et al. (2021: Figures 2 and 3) see the multiple incision-infill cycles critical to their interpretation. Each of their fills (T4-T2) is specified to have a horizontal base, scarp-like sides and an aggradational top, but no field illustrations of such bases and scarps are provided. T4 and T2 appear to lack diagnostic scarp-like morphologies along their upland edges throughout the area, and T4-T2 bases and thick fills were rarely observed.

Finally, relations which are seemingly contradictory to Duyfjes’ (1936) mapping remain unexplained. His (1936: 19) terrace-deposit unit covered most of the right bank with substantial thicknesses of *“loose sandstone, gravels with fossils”* (S I Figure 16). No bedrock protrudes through in the right-bank upland, and the terrace-deposit unit locally descends below the river, conditions which are inconsistent with the thin flat-lying terrace fills of Berghuis et al. Our own experience fits Duyfjes: A quarry ∼500m west of the Trinil Museum has >10m of flat-lying sand and gravel underlying a high-elevation terrace surface (a “T5”); a quarry- and other-exposures farther north reveal >4m of horizontal sand and gravel where Berghuis (but less so Duyfjes) map bedrock (‘1’ and ‘3’ on S I, Figure 16; Huffman 2016). The deposits we observed are less consolidated than those in the 1890s-1900s excavation of the left bank appear to have been.

The Berghuis et al. interpretations require much substantiation. This should include a single, detailed, geological map of the Trinil area showing both terrace- and pre-terrace units (to replace the mapping techniques described above). Substantiation would include compelling evidence for terrace-fill south of the left-bank Selenka and Dubois excavations, in light of their firsthand reporting, and an explanation of the thorough lithification of strata dug there in 1891-1908. Important also would be full illustration of the bedrock-fill-bedrock-fill sequence that Berghuis et al. envision south of the Trinil Museum, and reconciliation of their interpretation north of the Museum with the results of the Selenka and Survey excavations there. Several other studies of strata along the Solo River to the west and east of Trinil, which bear on the terrace history at Trinil, should be assessed by Berghuis et al. (e.g., S I Figure 19).

#### Summary of the conflicts

Duyfjes mapped horizontal terrace deposits on the right bank surrounding Selenka Expedition Pit I and the Survey 1932 excavations. Old photographs and Selenka’s cross sections confirm eight-to-nine meters of flat-lying post-**HK** strata occur there. Duyfjes gave the strata ∼9° of dip, contrary to the structural attitude Selenka reported. Numerous experienced field teams have later concurred with Duyfjes’ placement of post-**HK** beds on the left bank in the Kabuh Formation, having presumably given due weight to how the lithification of the strata corresponds to that of folded Kabuh and Pucangan sandstones and mudstones elsewhere in the Kendeng Hills.

Regarding the right bank around the Selenka and Survey excavations, Berghuis et al. (2021) see prominent south-dip in the Kabuh-Pucangan (their Trinil and Batu Gajah Formations) as opposed to the flat-lying beds reported. Berghuis et al. correctly recognize the flat-structural attitude of the post-**HK** sequence on the left bank, but do not give substantial evidence for interpreting the sequence there as Late Pleistocene valley fill. They do not address indications in site photographs that the terrace upland in this vicinity is floored by eroded bedrock (Huffman et al. 2015, 2018). They do not explain how the biostratigraphic differences between the Trinil and Ngandong faunas would impact assigning the **HK** to the Late Pleistocene.

The most-plausible provisional reconciliation of geological relations around Trinil combines elements from several past interpretations: The **LB**-**HK** and younger strata (that is, those removed by excavation and projected to still be present along the left bank) are the Kabuh Formation in flat-lying structural attitude. So are the **HK** and eight-to-nine meters of superjacent beds around the former Selenka and Survey excavations on the right bank. This structural situation involves flattening of bedrock formations that locally interrupts the normal southward monoclinal dip of bedrock strata along the Solo River and in the adjacent southern Kendeng Hills. The flattened attitudes make it especially difficult to separate Kabuh and terrace deposits strata near the present-day Trinil Museum on the right bank. Confirmation is needed there for the T2-T4 valley fills that Berghuis et al. propose exist, especially their scarp-like upland edges and flat broad bases. The geological age of the Trinil fauna should continue to be taken as ∼0.9 Ma, unless field- and radiometric-age determinations at Trinil firmly establish something younger.

### STRATIGRAPHIC FRAMEWORK, LEFT BANK

Roughly 2200m^2^ of the left bank were removed in 1891-1908 for the purpose of mining the *Pithecanthropus erectus* (*P.e.*) bonebed near the seasonal low-water level of the Solo River (S II-A4r). Site photographs (1894-1926), eyewitness excavation accounts, and later geological investigations generally establish the stratigraphy that these excavations encountered: Eight-to-nine meters of flat-lying, well-indurated strata lay between the bonebed and soil near the surface of the terraced embankment.

In our view, Dubois and his field supervisors amply recorded the placement of vertebrate fossils in the Lapilli Bed (**LB**) of the 1891-1893 left-bank excavations, which spanned ∼200m^2^ (a tenth of the total left-bank excavations). The **LB** had an internal bioclast concentration, our Principal Fossil Zone (**PFZ**), which was the reported source of the *Pithecanthropus erectus* finds (Figure 2 and 3). The Selenka Expedition applied the name Hauptknochenschicht (**HK**) to the main bonebed in the ∼370m^2^ 1907-1908 left-bank Pit II, which was several-tens of meters away from Dubois’ hominin discovery points (Figure 7).

There is only sparse geological information on the bonebed encountered in the large intervening 1895-1900 excavations (S I Figure 7). However, site photographs from 1894, 1900 and 1907 allow the sedimentary sequence overlying the main bonebed to be divided into stratal units, our units **2**-**5**, which once extended across the 1895-1900 excavation area (Figures 3 to 5 and 7 and 8, S I Figures 3 to 6 and 8 to 10). A soil developed on unit **5** across most of the excavation area at the base of a terrace level near the top of the incised river embankment (e.g., Figures 4; S I Figure 4). The basal part of unit **5**, ∼5m above the **HK**, contained the localized *Stegodon* bonebed (**SB**), but generally the strata above the main bonebed had relatively few vertebrate fossils. A unit **6** rests on strata of **2** to **5** across an erosional surface in the western parts of the Dubois 1900 Trench and Selenka Pit II (Figures 4, 5 and 7, S I Figure 11a).

Unit **6** is not known to have been fossiliferous. The unit is not evident near the **LB** level in the 1891-1893 discovery excavation area, judging from the 1926 photograph (Figure 9), and accounts which indicate that the **LB** and **PFZ** were present throughout the 40-m Trench and the 1896-1900 excavations which lay between this trench and the known occurrences of unit **6**. Since 1908, remnants of the **HK** and **2**-**6** appear to have been available for re-examination by geological field teams working along the left bank (e.g., Figures 6d and 9, S I Figure 18). From 1936 onwards, geologists have attributed the left-bank remnants to the bedrock Kabuh and Pucangan Formations. 1:250 mapping in 1977 confirmed this attribution (Figure 6d, S I Figures 16 to 19).

The suggestion that Solo River valley fill makes up some or all of rock volume excavated along the left bank in1891-1908 awaits substantiation from investigations that closely integrate the archival record with modern field study, including evidence of terrace deposits occur at or immediately near the 1891-1893 *Pithecanthropus erectus* discovery excavations. New geological mapping of the greater Trinil area also is required to clarify conflicts in reporting on the distribution and stratigraphic relationships of dipping- and horizontal-sedimentary strata. In particular, resolution is needed of the apparent stratigraphic differences between the left-bank discovery area and the right bank near the 1907-1908 Selenka Pit I and present-day Trinil Museum.

## DISCOVERY RECORD

### FOREWORD

Old photographs, maps and firsthand accounts contain critically valuable documentation on the paleontology of the main bonebed, including the left-bank where the largest area of the bonebed was removed and the greatest number of fossils now in museum was collected. The same archival sources provide information on the sedimentary facies excavated. In addressing these topics, we focus first on re-evaluating Dubois’ central provenience conclusion, the *Pithecanthropus erectus “remains … were found at exactly the same level ….* [and thus were] *deposited at the same time,”* together with many hundreds of other fossils of broad biotic composition (Dubois 1895b: 4, S II-F4; de Vos and Aziz 1989). Key to evaluating this issue is the continuity and consistency in reporting from excavation supervisors to Dubois, Dubois’ accounts to government sponsors, and ultimately, Dubois’ publications. We next investigate the origin of the main bonebed (Figure 12), starting with the observations and inferences that Dubois and Selenka scientists made. Our DISCUSSION, which follows, explores several key implications that Trinil has for *Homo erectus* paleogeography in southern Sundaland (Tables 5 and 6, Figures 13 and 14, S I Figures 22 to 24).

### EXCAVATED FOSSILS, LEFT-BANK

When Dubois saw Trinil on September 6-7, 1891, he considered the site was his *“best find of all”* (Endnote A(v), S II-B2e; also, S II-B1c, -C1 and -C2). He was an experienced fossil hunter by that date. He had mapped the central Sumatra highlands geologically during 1888 to locate fossiliferous caves, one of which now is significant as the source of 63-73 ka Anatomically Modern Human remains (de Vos 1983, Dubois 1888, 1892a, Westaway et al. 2017). Starting in June 1890, Dubois conducted a pioneering geological and paleontological reconnaissance of the vertebrate-bearing deposits of eastern Java, investigating both cave deposits near the southern coast of Java and fossiliferous volcaniclastic bedrock formations in Kendeng Hills (Figure 1b; e.g., S II-B1b). His 1890 efforts in the Kendeng Hills produced a partial mandible of a *Homo erectus* (S I Figure 21). Dubois reasonably expressed high praise for Trinil because that site offered the opportunity to excavate a single well-lithified deposit with large fossils that represented multiple extinct vertebrate species.

#### September 1891, Skullcap Pit

G. Kriele began excavating the **LB** along the left bank about September 1, 1891 (Figure 3b; Endnote A(v), S II A1a). Kriele was a military corporal assigned by the government of the Netherland Indies to assist Dubois, a government employee himself. As Dubois described the 1891 Trinil site in September, the *“exposed dry shallow sandstone ledges in the riverbed were excavated below water level on both sides”* of the Solo; the *“bone remains* [were] *about 0.20 meters below”* (Endnote A(viii), S II-B2d and -B2e). He summoned A. de Winter, his second military assistant, from Patiayam (Figure 1b) to exploit the **LB** along the right shoreline at Trinil (Figure 3b; S II- B2d).

On September 3, before Dubois returned to Trinil himself on September 6-7, the left-bank **LB** produced specimens now attributable to *Bubalus palaeokerabau*, *Duboisia santeng* and *Stegodon trigonocephalus* (Table 1, Endnote A(i), S II-A1a) from the excavation that we name the Skullcap Pit (Figures 2b and 3a). The fossils, *“found all together”* stratigraphically, included deer (*Axis lydekkeri*), turtle and mussel, and *“tree trunks”* and *“leaf imprints;”* the bony elements were *“often fractured”* but disclosed no evidence of human or carnivore activity (Endnote A(iii), S II- B2b, -B2d; also, S II-B2e). On September 18, Kriele sent a sketch map of the ledges showing a *Stegodon* mandible and tusk *in situ* at the Skullcap Pit location (Figure 3b). He wrote that the fossils being transmitted were *“all either from a depth of around 0.20 m below the water level or even with it”* (S II-A1b), just as Dubois notified the government that month (Endnote A(iii)). The lower parts of the exposed Proboscidean mandible in the Skullcap Pit presumably were at this elevation (Figure 3b, inset).

The hemimandible of a *“fossil cat … about the size of an average royal tiger* [*Panthera tigris*]*”* and specimens of crocodile (*Crocodylus siamensis*) and gharial (*Gavialis bengawanicus*) were among 28 crates of fossils collected in 1891 on both river ledges (S II-B2e, -B2j; also, -A1k and –A1m). The **LB** material found in a September shipment, which had *“a considerable number of* fossils … [from] *the two sand*[stone] *ledges,”* included *“the upper-right third molar … of a chimpanzee,”* the 1891 Molar (Endnote A(v), S II-B2e; also, Endnote A(viii), S II-B2d). Dubois is not known to have visited the Molar discovery spot before the surrounding rock was removed.

#### Skullcap discovery, October 1891

Dubois’ October 1891 memorandum to the Director of the Department of Religion and Trade, Netherland Indies (S II-B6j) reported that *“close to the place where the molar was found in volcanic tuff on the left bank … a magnificent skullcap was excavated* [in Skullcap Pit **LB**]*”* (Figures 2 and 3b, Endnote A(ix), S II-B2f). The cranial find must have been in the one October shipment, transmitted on the eleventh. The Skullcap discovery was made about 16 months after Dubois started in Java and six weeks after digging began in the **LB** on the left bank.

Dubois visited Trinil on October 21-24, but his October memorandum did not describe what he saw in the Skullcap Pit. No extant record contains the date of the Skullcap finding, names of the discoverer(s) and his (their) discovery-day experiences, or other fossil remains or lithologies observed nearby. Dubois also did not prepare a map of the Pit or illustrate the context of the Skullcap in profile form.

He (1894a: 1) did stipulate in his monograph that the 1891 Molar and Skullcap had been found a meter apart (S II-B6j; also, Figure 3a). In 1895, Dubois (1896d: 241; S II-F3(ii)) specified that the Skullcap and 1891 Molar were discovered *“in the lapilli bed on the left bank* [Skullcap Pit] *… amidst among hundreds of other skeleton remains.”* M. Selenka interviewed Kriele in 1902 about Trinil (S II-A4s) and took de Winter to Java as a field consultant in 1907. Selenka understood that both men were *“present during the excavation of the skull”* (Selenka and Blanckenhorn 1911: i, xiii, Berkhout and Huffman 2021: 5, 13).

In October 1891, Dubois transmitted to government sponsors two photographs (now lost) of the specimen, and reported, “the upper part of the occipital portion is covered by a stony mass …. so hard and strongly adhering to the bone that it cannot be removed for the time being without causing damage to this precious fossil” (S II-B2g). Dubois (1894a) cleaned off the hardened exterior agglutinate before he departed Java in 1895.

This external mass was “mainly consisting of the same lime concretion found at many places in the lowlands of central Java on bones extracted from the black clay” (S II-B2g). Dubois clarified the geological distribution of such concretions (S II-C5), writing “many bones from Trinil, which in sandstone-like tuffs at the surface [exposures of the **LB** are] partially covered with lime [CaCO_3_] concretions, that adhere tightly to its surface. This is especially so with some [Axis lydekkeri] deer antlers … and a large buffalo skull [Bubalus palaeokerabau].” He further noted, “limestone concretions [occur] near Trinil at various levels (even up to 10 meters below the surface [in the incised embankment]) between thin beds of 1-, 2- to 3-cm thickness” (S II-C6).

Pebble conglomerate filled the Skullcap endocranial cavity, further making *in situ* discovery clear (Figure 2b). Dubois did not remove the cranial fill until after September of 1895. The endocranial surface which emerged is finely preserved, but full preparation of the exterior revealed it to be pitted much beyond the area affected by the stony mass (Dubois 1924b: Plate II: Fig. 1-3).

The public announcement of the Skullcap and molar discoveries appeared in Dubois’ Fourth Quarter report, which was drafted January 20, 1892, and published anonymously (S II-B2j). His October memorandum to government officials had stated *“of all known living and fossil anthropoids … the new Java chimpanzee undoubtedly ranks the highest”* in an evolutionary scale (S II-B2f; also, -B2h and -E1c). By December 23 (eight months before he had Femur I in hand), Dubois was referring to the Skullcap as *Pithecanthropus erectus* (S II-B2h; also, S II-B4b). He soon wrote to a colleague, the fossil was *“truly a new and closer link in the largely buried chain connecting us to the ‘lower’ mammals”* (S II-B2h; also, S II-E1c). To the government, he asserted, because Trinil “*has become so important to science”* and transformational in anthropology, continued excavation was *“essential,”* even though the *“major part”* of the original exposures of *“sandstone-like tuff in the riverbed … had already been excavated”* (S II-B2h to -B2j).

Besides *Homo erectus*, the mammalian assemblage in the Skullcap Pit, known from contemporaneous reporting, included *Axis lydekkeri*, *Bubalus palaeokerabau*, *Duboisia santeng*, *Stegodon trigonocephalus*, Testudinoidea (?), fresh-water Mollusca shells, fossil wood and leaf imprints (Table 1, Endnote A). At least two craniums of the *Stegodon* and one of *Axis* were recovered.

#### Femur I discovery, August 1892

While uncertainty surrounds the August date of Femur I discovery, critical information on the provenience of the specimen and the actions of the discoverers are in the archival record. In June, Kriele and de Winter (KdW) started *“excavations … into the river bank done from above,”* so that once the dig was deepened and the river dropped *“a larger surface area of the fossil-bearing layer”* (**LB**) would be exposed and could be removed in entirety at about the same time (S II-B3b; de Vos and Sondaar 1892). During the last days of June, *“harvest of bones,”* including *Axis lydekkeri “buried here in large numbers,*” indeed was removed from the fresh upper portion of the **LB** (Endnote Bi, S II-B3d; also, SII-B3b). The excavation was the first segment of what would become the 1892 25-m Trench (Figure 3a). It had produced fossils referable to *Bibos palaesondaicus*, *Crocodylus siamensis, Duboisia santeng* and *Stegodon trigonocephalus* (Table 1, Endnote B(i) and B(ii), S II-B3d and -B4b; also, S II-A2f, -A2g and –A2i).

On September 7, 1892, KdW explained: “the bone of the chimpanzee [Femur I] …. was found on the same side [of the river] as the skull[cap] and also at approximately the same depth [as the cranium] and even with the previous low-water level [LWL within the **PFZ**], [with the Skullcap and Femur I having been] separated from each other by about 12 meters” in the excavations (S II-A2k). Their August 31 letter about the find had been delayed in the mail (S II-A2j; also S II- B3b). When KdW drew this conclusion, the **PFZ** was exposed over a limited area, but the upper **LB** was broadly penentrated (described below), so that KdW must have relied on upper **LB** to gain confidence in the presence of the **PFZ** subunit at the Femur I discovery location.

There are other critical implications of KdW’s August 31 and September 7, 1892, letters. They contain information that clarifies both the situation in the femoral-discovery excavation on the day of find and what Dubois actions were when he received Femur I. We highlight five implications.

First, judging from the two letters, KdW had paid close attention to the internal stratigraphic of the “fossil-bearing layer” when it was being exploited mid 1892. Dubois’ (1896c: 3-4, S II- F3c(i)) was clear later that he had understood KdW’s 1892 reporting in this matter. For instance, he (1896d: 241-243; S II-F3(ii)) wrote in English:*“In the lapilli layer on the left bank of the river, amidst hundreds of other skeletal remains, … the third-molar tooth was found …. on exactly the same level in that layer* [in which] *the cranium was discovered at a point one meter distant …. The mammalian remains found in the same layer …. are …. the bones of …* [*Axis lydekkeri*] *…. Stegodon* [*trigoncephalus*]*, besides … Bubalus* [*palaesondaicus*] …. *rhinoceros, Sus, felis, hyena, … gavial and crocodile”* [Endnote B(i), S II-B3d; also, S II-F3 (i)).

Second, KdW’s August 31 letter states de Winter recalled that *“a mandible and tusk of an elephant”* had been found nearby Femur I (S II-A2j). The provenience letters were written by Kriele on behalf of the two men, per protocol for Trinil letters to Dubois (S II-A). The specification that de Winter was the one who remembered the *Stegodon* reflects the fact that he discovered Femur I (Kriele appears to have had been less directly involved). KdW later affirmed that *“de Winter found the leg bone”* (S II-A2q). M. Selenka (1911: xiii, xiv) learned from him that he *“personally dug up the* Pithecanthropus *femur”* (Berkhout and Huffman 2021: 13).

Third, the reporting about the mandible and tusk underscored the paleontological match between the Skullcap Pit and the portion of the 25-m Trench where Femur I was found. All three men would have known that the two large *Stegodon* fossils fit **PFZ** provenience. Most plausibly, the tusk lay parallel to the horizontal stratification and the mandible took up nearly the full thickness of a bed, making these fossils prominent both in the field and de Winter’s memory. Dubois could compare Femur I taphonomically with plenty of **LB** fossils he already had. Presumably he identified the mandible and tusk in the shipment he received in August.

Fourth, the August 31 letter clarifies the actions the men took around that time. After examining Femur I, Dubois sought explanation from the men on how certain chips of bone had been lost off the end of the specimen. KdW responded: “*About the bone* [Femur I] *de Winter tells me that the pieces that are missing* [from it] *were blown away while on a djati* [teak] *leaf by heavy winds during the process of gluing and we cannot find them again* [right now]*”* (S II-A2j).

High-water prevented access to the discovery excavation so that KdW could not recover the pieces immediately (S II-E1c). The location of the loss and condition of the trench must have been remembered clearly because KdW promised, “*as soon as the water level subsides* [such] *that we can work on the opposite* [left] *side where the bone* [Femur I] *was found….* [we will be] *searching carefully for the pieces that had been knocked from the bones”* (S II-A2j). Therefore, de Winter’s preparation of the specimen must have taken place below an elevation subject to the flooding, most plausibly within or next to the Femur I excavation. His recollection about the missing chips, like the nearby *Stegodon* fossils, stems from the events he experienced on the day of discovery.

Finally, by asking KdW about the lost fragments, Dubois divulged that he had examined the Femur I closely. The losses were from a hole in the popliteal surface and *foss intercondyloidea* (intercondylar notch) of the posterior epiphysis, losses still notable on the specimen. He (1926a) eventually stated that holes had been *“caused by excavation”* (S II-F9i; also, Dubois 1926b). Perhaps he drew this conclusion before writing KdW in August 1892. Dubois also might have had observed by then, as he (1895b: 3, SII-F4) reported three years after the discovery, that *“the marrow canal has been partly filled with a stony mass,”* which helped makes the specimen *“more than twice as heavy as a recent human femur of the same size,”* and tended to substantiate the reported provenience in the **LB**.

There is no record about the nature of the material surrounding Femur I when it was *in situ*, or even when the fossil was in de Winter’s or Dubois’ hands, only that preparation of the specimen was completed long before Dubois (1894a) left Java. The greatest challenge of preparation presumably would have been cleaning rock from around the exostosis (S I Figure 1) but this too was completed by the end of 1893.

In early September 1892, Dubois still might have been unaware just how little of the discovery excavation had fully penetrated the **PFZ**. The trench still was far short of its ultimate 25-m length and evidently just ∼2-m wide (Figure 3a). Even at the end of 1892, when the top of the **LB** had been exposed all along the full 25m length, only 20% of the **PFZ** and basal **LB** had been removed (S II-B5c). When Femur I was found, the area of full penetration of the **LB** was only ∼10m^2^ out of the final ∼50m^2^ of the 25-m Trench.

#### 25-m Trench, late September to November 1892

In late 1892, the Femur I discovery excavation was expanded into the bank southward of the find spot (Figure 3a), as explained in LEFT-BANK GEOLOGY. After digging downward from the embankment top for eight-to-nine meters and passing through hard volcaniclastic sandstone and other facies with few fossils, the crew again struck the fossil-rich **LB**. By year end, the **PFZ** level was reached across the full span of the 25-m Trench. These milestones form a framework for evaluating Dubois’ claim of co-deposition of the Skullcap, Femur I and other large fossils.

On September 23, 1892, before he is known to have gone to Trinil that month, Dubois reported to the Indies government that the three fossils of the future *Pithecanthropus erectus* originated from one *“level of the sediments”* (**PFZ**), just as he claimed in 1895 publications (S II-B4b; also, S II-F3ci, quoted above). He inexplicably started saying that Femur I was 10m from the Skullcap, not KdW’s 12m. The day after composing the 1892 memorandum, Dubois went to Trinil. He did not report the results of his field checking, any more than he had for the Skullcap.

In early September, Dubois asked KdW how they might find more parts of the recently discovered Femur I individual. The response reflects their keen awareness of the site stratigraphy. First, they proposed deepening the pit near the discovery spot, since the **PFZ** had not been fully penetrated and the lower **LB** might be fossiliferous there; second, they proposed cutting down the high embankment south of the find spot to expose more **PFZ** (S II-A2k).

KdW letters and Dubois government reporting later in 1892 are explicit about the stratigraphy they encountered while enlarging the 25-m Trench (as described in LEFT-BANK GEOLOGY; also, Figures 3 and 4): The upper two-thirds of the eight-to-nine meters above the **LB** lacked *“a lot of bones”* (S II-A2o) and was “*a fairly hard sandstone-like andesitic tuff which can only be removed with pickaxes and crow bars”* (S II-B5a). On October 28, KdW wrote, *“the corners* [of the 25-m Trench] *… progressed to about the depth* [elevation] *at which the leg bone* [Femur I] *and skull* [Skullcap] *were found”* (**PFZ**; S II-A2p). On November 7, they reported that *“the corners of the pit* [25-m Trench were] … *about 20cm into the target bone layer* [**PFZ**]*”* (S II-A2q).

By then KdW had stopped reporting prominent fossils being encountered, plausibly because Dubois already knew the paleontology of the upper **LB**. The most important new information which came from the field was that the Skullcap and Femur I **PFZ** stratigraphic level could be followed from an edge of the Skullcap Pit through and beyond the spot where the femur was found. In short, the late 1892 work had confirmed the continuity of the **PFZ** across ∼50m^2^ of excavation and revealed the **LB**/**PFZ** to be below about nine meters of indurated, generally fossil-poor strata. Even though the **LB** in this much enlarged Trench was *“just as rich in remains of vertebrates … as it usually is”* (S II-B5d), Dubois had to report to the government sponsors that *“rising water … forced us to finally abandon the work on November 16th, after having only excavated about 1/5th of the level of interest* [the **PFZ**]*”* (S II-B5c).

Records about three individual fossils support provenience conclusions drawn from letters and reports: (i) A macaque tooth which is present in the Dubois Collection has a label, most likely written by Dubois, that reads, *“trench of 25 m of 1892, lowest level, ½ m below pe”* (DC no. 3789; de Vos 1989: 227). The “pe” evidently refers to the *Pithecanthropus erectus* stratigraphic level so that the macaque tooth came from 0.5m below the **PFZ** in the lower **LB**. (ii) A molar of the Asian porcupine *Hystrix lagrelli* was recovered *“at the lowest level”* of the river, according to a label (DC no. 1482a; de Vos and Sondaar 1982: 47, 49; also, S II-B5c). (iii) A fourth *Pithecanthropus erectus* specimen, the 1892 Molar, was found during October close to the easter end of the 25-m Trench (Figure 2b; S II-F2 and –F3) *“in exactly the same* [stratigraphic] *plane”* as other *P.e.* remains (**PFZ**), according to a later Dubois (1924a: 277; S II-F8) specification that most plausibly came from an 1892 KdW label, now lost.

#### 40-m Trench, 1893

The firsthand reporting about the 1893 40-m Trench recounts events closely similar to those of late 1892, and more importantly, both document the fossil assemblage that the **LB** contained and the indurated poorly fossiliferous nature of the overlying strata (Table 1, Endnote C).

The 1893 field crew again proceeded from the terrace tread downward in horizontal increments to about eleven meters. Dubois’ experiences from 1892 motivated him to have the crew remove this *“fossil-poor rock mass as quickly as possible”* to reach *“the deeper … rich bonebed”* (Endnote B(i), S II-B3d). These desires notwithstanding, after digging a soft soil capping in the 40-m Trench, the excavators encountered strata of *“severe hardness,”* which Dubois witnessed (Endnote B(i) and B(ii), SII-B6b and -B6c; also, S II-A3a to –A3i). The laborious removal of rock continued for six-to-eight weeks. KdW wrote that the rock was *“so terribly hard that it is almost impossible to get through,”* and repeated the complaint when digging the *“lower part”* (July 7 and 14; S II-A3a-vi, -A3a-ix) of the sequence (units **1** and **2**) that was photographed the next year (Figures 3c and 4a). Dubois’ government memorandum concerning this period of excavation stressed the *“relative paucity of fossils”* above the **LB** (Endnote B(iii), S II-B6c).

When Dubois visited on August 17-19, 1893, the crew was poised to greatly expand horizontal exposure of the **LB**. Part of this new excavation was only several meters away from the Femur I discovery point (Figure 3a). Dubois saw fossils attributable to *Axis lydekkeri*, *Crocodylus siamensis*, *Duboisia santeng* and *Stegodon trigonocephalus* (Endnote C(iv), S II-B6d). Near the end of the month, the **LB** Kriele reported finding *“1 nice elephant tusk* [*Stegodon trigonocephalus*]*, 1 crocodile skull* [*Crocodylus siamensis*]*, 1 antelope skull* [*Duboisia santeng*]*, 1 turtle* [Testudine]*, a few leg bones, deer antlers* [*Axis lydekkeri*]*, some ribs and vertebrae”* (Endnote C (v), S II-A3c/i).

The *“the target layer”* (**PFZ**) was reached by September 1, and had *“rather many bones”* and a lot of *“wood”* and *“shells,”* according to KdW (Endnote C(vi), S II-A3cii). They found another oversized fossil, the cranium of *Stegodon* with *“the molars still in it,”* and the specimen would not fit any crate on hand (Endnote C(vii), S II-A3d). The abundance of fossil recovery from the **LB** continued for much of September (Endnote C(ix), S II-A3f).

Soon, however, coarser, conglomeratic fossil-poor sandstone was evident in the lower **LB** (Endnote C(xi), S II-A3i), although it did produce one skull of a buffalo (*Bubalus palaesondaicus*) and two skulls of antelope (*Duboisia santeng*; Endnote C(xii), S II-A3j). Kriele recommended halting work in this *“the hard coarse* [conglomeratic] *layer ….* [which contained] *nothing* [in the way of important fossils]*,”* only a few antlers and isolated finds (Endnote C(xiv), S II-A3i). The sedimentary materials in the lower **LB** clearly had substantial amounts of large gravel but few fossils, compared to the **PFZ** and upper **LB**.

Dubois gave additional information on fossil recovery in 1893: A “black coaly clay bed …. 11 to 12 meters below ground level …. proves to be the underlying formation to the bone-bearing volcanic tuffs …. [the **LB**, which produced] “many antlers of the small Axis-like deer species [A. lydekkeri] and also remains of Stegodon [trigonocephalus], [Duboisia santeng], Bubalus [palaesondaicus], …. [together with] the first almost complete skull of a crocodile [Crocodylus siamensis]” (Endnote C(xv), S II-B6f; also, LEFT-BANK GEOLOGY).

In both “last year’s excavation [25-m Trench] …, only partially worked” and 1893 40-m Trench (S II-B6e), the crews “were able to essentially dig away the entire bone-bearing bed … before the 26^th^ [of November] when the work [site] became hopelessly inundated” (S II-B6h). Kriele had shipped 25 crates of fossils and 6 crates of wood from Trinil, but Dubois concluded that among the “fossils … none were the hoped-for additional parts of the curious Anthropopithecus …. [, the other remains of which must have] washed away during the formation of the [incised modern Solo] river bed, together with a large portion of the bone-rich tuff [**LB**]” (S II-B6i). Given this prospects, Dubois gave up the Trinil operations, and KdW install“a small pillar,” the now-famous Dubois “P.e.” monument (at the current Trinil Museum; S II-B6h; also, -A3a/ix).

#### 1894, Dubois’ last visit

Early the next year, Dubois (1894a, 1895a) turned attention to completing his *Pithecanthropus erectus* monograph (Endnote G, S II-F7). It arrived at the publishers on February 8 and was published on August 25, 1894, three years after vertebrate-fossil collecting had commenced from the sandstone ledges on the shores of the Solo (S II-B6j). Neither the monograph nor Dubois’ final 1893 periodic submissions to the Indies government enumerated the taxa he recognized during his scrutiny of the Trinil fossils. On September 5, Dubois had the left-bank excavation site photographed from across the Solo River (Figure 3c). This was done in conjunction with his final boat trip along the waterway. In November, Dubois wrote to a colleague about the toll success in Java had cost him. *“I have sacrificed my whole career, my health and my good humor,”* and even the wellbeing of *“my wife and children”* (S II-E1d). He spent much of 1894 following up on his early 1890s study of the geology and paleontology near Kedungbrubus, Butak and other areas of the Kendeng Hills (S I Figure 21, S II-B7; also, S II-D2 and Albers and de Vos 2010). The fossils he found near Kedungbrubus are the basis for Kedung Brubus fauna, while fossils he collected at Butak contribute to our recognizing Butak bonebed (Tables 5, 6-C and 6-D, S II-B7a).

### PROVENIENCE OF FEMUR I

The foregoing foundation puts us in a better position to reassess the provenience of Femur I. We illustrate our results in a cross section (Figure 10), and for the following reasons, endorse KdW’s and Dubois’ stratigraphic placement of Femur I in the **LB**/ **PFZ** in preference to conjectural possibilities lacking strong foundation in the archival record.

The on-the-ground circumstances along the left bank in 1891-1893 were straightforward geologically, particularly with regard to the flat-lying attitude of erosion-resistant strata (Figures 3 and 4a) and the strong concentration of large-sized vertebrate fossils in one stratum. At the end of June 1892, a month before Femur I was found, excavation of the embankment at the site neared the seasonal low-water level, and *“a large surface area of the fossil-bearing layer”* (**LB**) produced a *“harvest of bones”* which included the species and types of bioclasts removed from the **LB** in the Skullcap Pit (Endnotes A and B). No fossil concentration was reported above the **LB**.

Before the discovery, Dubois developed special expertise to bring to his analysis of the paleontology and stratigraphy of the Femur I excavation. First, he evaluated the taphonomic characteristics of the **LB** fossils found there, and second, he studied the fluvial sedimentary processes visible in the Solo River and evident from deposits exposed in its banks (e.g. Endnote B(i), S II-B3d, and -C3 to -C5). KdW’s sharp geological awareness at the time is evident in Kriele’s own early August 1892 field analysis of cross bedded strata (S II-A2g), and KdW’s accurate September 7 assessment about where more fossiliferous **LB** could be unearthed near the Femur I discovery spot (S II-A2k).

KdW’s description of the provenience of Femur I unambiguously placed the find in the **PFZ** portion of the **LB** (S II-A2k and S II-A2j). Dubois had other **PFZ** and **LB** fossils at hand in August 1892 with which to compare the taphonomic condition and adhering sediment on Femur I. He promptly reported the provenience specification to the Indies government (S II-B4b and -B5b). His confidence in the provenience did not alter after visiting Trinil in September 1892.

Late that year and again in 1893, KdW and Dubois gained confirmatory evidence on the stratigraphic context of the **LB** when the crew dug downward by horizontal increments through sparsely fossiliferous indurated beds of the high embankment of the Solo toward the fossil concentration near low water level. The **PFZ** was reported to be traceable from the edge of the Skullcap Pit to points near the Femur I discovery spot and beyond in the 25-m Trench. Again, no shallower fossil concentrations were reported.

As the excavation neared completion in both 1892 and 1893, the men stood before eight-to-nine meters of strata displayed across 25-m and 40-m-long exposures. Dubois documented the stratigraphy with a high-quality photograph taken from across the Solo River in 1894 (Figure 3c, S I Figure 4), knowing the strata seen were the same as excavated in 1891-1893 (Figure 10). The same sequence is traceable photographically across several thousand square meters of excavation from the 1894 exposures to 1900-1907 trenches (Figure 4 and 5). Dubois’ 1895-1896 published accounts unequivocally attributed the *Pithecanthropus erectus* fossils to the same stratigraphic level (**PFZ**) in one fluvial deposit, the **LB** (Figure 2a; S II-F1 to -F5). He kept to this view throughout his life (S II-E2).

Firsthand reporting contradicts various alternative provenience speculations for Femur I. It could not have come from below the **LB** because KdW’s September 7, 1892, letter indicates more **LB** remained under the discovery level at the spot where the Femur I was found (S II-A2k). No shallow fossil concentration was reported as having occurred anywhere in the 1892 and 1893 excavations, nor later adjacent ones (described below). No terrace deposits or slumped beds were recognized by KdW or Dubois; rather, their contemporaneous accounts indicate only indurated strata arrayed in regular depositional order, necessarily excavated by persistence because of the hardness of the rock.

No large-sized, well-preserved fossil, such as Femur I, was reported as having been introduced into the excavations by flooding of the Solo River during the years of Dubois’ work at Trinil, leaving no support for the proposition that Femur I was an extraneous clast carried into the 25-m Trench by high water (the trenches did fill with mud during the 1893 wet season; S II-A3a).

If Femur I was found close to the *Stegodon* fossils, as de Winter remembered, the context seemingly would have been difficult to mistake, because large-size proboscidean bioclasts were known by then to characterize the **LB**. Moreover, the mandible quite likely either extended into the upper **LB** above the **PFZ**. The specimens presumably was remembered because it was identified before the digging reached the Femur I level, had to be unearthed laboriously after Femur II was found, or extended below the **PFZ**. *Stegodon* fossils occurred ∼5 m above the **HK** in Selenka Pit II (Figure 8), but KdW and Dubois never reported fossils comparable to this in the 1891-1893 trenches. In any event, a provenience error off stratigraphically by five meters is not credible.

It is further implausible to suppose de Winter lost track of who was working in what stratum in August 1892, given the consistent operational procedures and straightforward stratigraphy of the excavations. If the Femur I was collected before de Winter saw it *in situ*, he surely would have noticed that a crew member had been digging at a large, well-preserved fossil significantly above the well-known **PFZ** context in which de Winter and the rest of the men were working. If de Winter had been handed a well-preserved fossil that he did not see in place, but thought might originate from above the **LB**, KdW surely would have searched in stratigraphically higher beds in the hope of finding more high-quality specimens.

If KdW or Dubois subsequently suspected that important fossils might occur above the **LB**, they surely would have searched diligently for higher-level finds while re-excavating the embankment in late 1892 and 1893. KdW letters about work during these times would have reported that more fossils had been found at a higher level, or that no such fossiliferous stratum had been encountered. The letters mention neither.

In sum, narrative and photographic records should lead to the strong presumption that the Skullcap and Femur I, together with other Trinil fauna species, originated from one fluvial accumulation, as the discoverers asserted. It is probable, perhaps highly so, that the Skullcap and Femur I were embedded contemporaneously and are of one geological age. No substantial provenience, sedimentary or paleontological alternative has been published. Such an alternative should include evidence from museum specimens of paleontological mixing (e.g., two faunas) and field identification in immediate vicinity of the Skullcap Pit and 25-m Trench of two deposits containing large, well-preserved, vertebrate fossils that taphonomic, sedimentary and geochronological criteria show to be meaningfully different in geological age. As it stands, the records from both 1891-1893 and later excavations support Dubois.

### 1895-1900 FOSSILS

Kriele’s letters to Dubois about 1895-1900 excavations (Figure 3a) repeatedly describe finding large skeletal fossils of Trinil fauna species near the seasonal low-water level of the Solo River, and Kriele sometimes specifically refers to the bonebed (Tables 1, Endnotes D and E). He also mentions the rarity of vertebrate remains in the overlying strata (units **2**-**5**). While the fossils that Kriele refers to in his letters have not been tied to individual Dubois Collection (DC) specimens, the assemblage reported matches the taxonomic and taphonomic characteristics of Trinil fossils in the DC (e.g., Table 1, Endnote D). Highlights follow.

#### 1895-1897

When the level of the Solo in 1895 dropped *“as low as it was in the first years”* of 1891-1893, the **LB**-**HK** Ledge next to the Skullcap Pit (Figure 4a) became exposed and yielded *“an incomplete antelope skull* [*Duboisia santeng*] *… with one horn, an elephant molar* [*Stegodon trigonocephalus*]*, as well as several other pieces of bone”* (Table 1, Endnote D(i), S II-A4b ‘page 30’ to 32’). Beginning in 1896, new pits and trenches were dug south of the 1891-1893 Skullcap Pit, 25-m Trench and 40-m Trench (Figure 3a). The provenience information and sketch maps that Kriele provided concerning these new works evidently were specific enough for Dubois to maintain confidence that the fossils originated from the same fossil-rich stratum that he had known personally in 1891-1893 and described in 1895-1896 publications.

The 1896 Left-bank Pit was dug west of the 40-m Trench and *“as deep as the bone bed …. more than 2 meters below the water level,”* where the stratum produced a *Sus brachygnathus* mandible and *“1 complete turtle”* (Table 1, Endnote D(iii), S II-A4c, p. 29 and -A4c, p. 35). The 1897 Upstream Pit, located immediately south of the Skullcap Pit, was *“brought to the depth on which no more bones are to be expected”* and *one complete* [*Axis lydekkeri*] *deer skull with antlers”* was found, apparently while digging the **LB**-**HK** (Endnote D(v) and (vi), S II-A4d). The 1897 Downstream Pit, which was situated southwest of the 40-m Trench, found *“nothing special”* as deep as “*8 meters down”* (units **2**-**5**, Figure 4, evidently were poorly fossiliferous).

But by twelve meters in depth, after passing through the **LB**-**HK**, Kriele had large bioclasts referable to the Trinil fauna species: a complete *Stegodon trigonocephalus* tusk 1.55 m long, an incomplete *Bibos palaesondaicus* cranium with complete horn cores, and an incomplete *Bubalus palaeokerabau* cranium with one full horn core (Table 1, Endnote D(ix), S II-A4d, Nov. 11). Additionally, a *Sus brachygnathus* mandible in the Dubois Collection (no. 502) has an original label indicating *“that the specimen has been found … 1.25 m below the lowest water level”* in the 1897 Downstream Pit, and *“a right upper first molar (M^1^ dext., Coll. Dubois no. 317)”* of *Rhinoceros sondaicus* is from *“0.75 m below the lowest level of the river”* (de Vos and Sondaar 1982: 48; S II-A4d).

Dubois added notes to Kriele’s 1897 drawing which shows that the 1897 Upstream Pit was south of the Skullcap Pit; Dubois seemed to have approved this representation of the 1892-1893 left-bank excavations, which differed in dimensions from his own published version of them (Figure 2b; S II-A4e; also, S II-A4f and -A4i and -A4k for other plats prepared in 1898 and 1899). The drawing also shows the 1897 Premolar originated in the 1897 Upstream Pit (also, S II-A4i and –A4k). This left inferior premolar (P_2_ sin.; Trinil 5) was attributed to *Pithecanthropus erectus* by Dubois (1899) at the 1898 Fourth International Congress of Zoology in Cambridge (de Vos and Sondaar 1982, Theunissen 1989, Smith et al. 2009).

#### 1899

In December 1899, Kriele submitted his most elaborate-ever illustration of the left bank excavations (Figure 3a). By Dubois’ annotations on the document, he is seen agreeing with Kriele on the relative locations of the Skullcap Pit, 25-m Trench, 40-m Trench, 1896 Left-bank Pit, 1897 Upstream Pit, 1897 Downstream Pit, and 1899 Trench, together with plans for the 1900 Trench (de Vos and Aziz 1989: Fig. 5: 414; S II-A4m). The 1899 Trench lay directly south of the 1893 40-m Trench, so that strata which were removed in 1889 held up the embankment seen in the 1894 photograph (Figures 3c and 4a). The Trench was divided into 450 one-meter squares (and was 3.75 times the size of the 40-m Trench).

Once Kriele *“reached the bone bed”* (**LB**-**HK**), he highlighted the recovery of massive fossils that are now attributable to the Trinil fauna: a *Stegodon trigonocephalus* tusk two meters long; a partial *Bubalus palaeokerabau* cranium; a complete upper *Rhinoceros sondaicus* cranium; an incomplete *Panthera tigris* mandible; some complete *Axis* antlers; and what *“turned out to be an ape’s tooth”* that was *“found about 0.5 meters above the lowest water level”* (a non-hominin catarrhine; Table 1, Endnote D(xi), S II-A4k). By December 21, 1899, Kriele had 7 crates with 1069 finds, apparently all from the 1899 Trench (Endnote D(xii), S II-A4l).

The average frequency appears to have been 2.4 fossils per m^2^ including 850 teeth, molars and diverse other specimens. Large bioclasts were rare in 1899 compared to the smaller bony remains and teeth among the fossils.

In all 1896-1899 excavations totaled ∼720m^2^ (S II-A4r) and produced many large-sized bioclasts from elevations near seasonal low-water levels. In terms of known DC species, the finds included: three complete *Axis lydekkeri* craniums with antlers; two partial *Axis* craniums; 15 *Axis* antlers; one *Bibos palaesondaicus* cranium with complete horns; a *Bubalus palaeokerabau* cranium with horn core; one *Stegodon trigonocephalus* cranium with tusks and molars; another *S. trigonocephalus* tusk, ∼2m long; a third *S. trigonocephalus* tusk, ∼1.10m long; a fourth *S. trigonocephalus* tusk, ∼1.55m long; one smaller tusk and molar of *Stegodon*; the partial cranium and two horns of *Duboisia*; one *Rhinoceros* calotte and molar; one partial *Sus brachygnathus* mandible; and *Sus* tusks and molars; as well as two complete sets Testudine remains; two partial *Crocodylus* craniums; wood and Mollusca (Table 1, Endnote D, S II-A4e to –A4q). The depths spanned by the labeled DC specimens from 1897 and 1899 is 1.75m.

#### 1900

Dubois (1899) launched an even-larger left-bank excavation in 1900, having been encouraged to do so by the 1898 Fourth International Zoological Congress (Figures 3a, 4b, 4c; Shipman 2001). As diagrammed by Kriele in advance and again during excavation, the 1900 Trench consisted of 900 one-meter squares (Figures 3a and 4b; S II-A4m and -A4o). It extended ∼100m east to west and was 6-to-19m north to south (Figures 3a, S I Figure 7, S II-A4m; also, SII-A4r).

By *“a depth of 2 meters”* below the top of the river embankment in the 1900 Trench, the crew had found little, and when about half of the Trench was dug to *“an average depth of around 4 meters,”* they had only *“some insignificant specimens”* in hand (Endnote D(ii to iii), S II-A4o). Our unit **5** evidently was near devoid of sizeable fossils. However, late in 1900, Kriele reported large-sized finds referable to Trinil fauna species that presumably are from the **LB**-**HK**: a *Stegodon trigonocephalus* cranium with complete tusks and molars; a *S. trigonocephalus* mandible with complete molars; a complete *Bubalus palaeokerabau* cranium with horn cores and molars; a partial *Bibos palaesondaicus* cranium; two partial *Axis* craniums with antlers; and the incomplete remains of several turtles (Table 1, S II-A4o).

Kriele had *“not been able to get anything of the ape-human”* (S II-A4m and -A4n), but failed to identify four *Pithecanthropus erectus* femora that Dubois (1932; also, 1934, 1935) concluded in June 1932 came from the 1900 workings (below). At least 850 Trinil fossils are recorded from 16 crates shipped to Leiden in 1900 (S II-A4p and –A4q). Considering the size of the 1900 Trench, Kriele seems to have been instructed to leave many finds in Java.

One DC specimen has labeling of a type that Kriele might have prepared for many of the finds: *“no. 536, a lower jaw of* Bubalus palaeokerabau *Dubois, on which a label was stuck with* [the detail that the fossil was in] *… fine sand, 1.25 m above the lowest level of the river”* in the western part of the 1900 Trench (de Vos and Sondaar 1982: 43; S II–A4m). When considered with labeled specimens from 1897 and 1899, the buffalo mandible would seem to indicate a span of discovery depths of 2.5m.

Kriele’s letters in 1900 have essentially no information on the **LB**-**HK** lithofacies, except the cross-lamination patterns that he reported (mentioned in LEFT-BANK GEOLOGY). However, Dubois had an abundance of data on the discovery bed(s) lithology from the sandstone and conglomerate adhering to museum fossils. Lithic matrix is still visible on the DC specimens, even after extensive cleaning over the course of a century (S I Figure 20).

Dubois recognized four additional hominin femora in his 1900 assemblage (Femur II to IV; Trinil 6 to 9). They have the dark color and stony fossilization characteristic of other Trinil specimens in the DC. Kriele had written *“Trinil”* on Femur II and Femur V, suggesting that while he did not identify the specimens as anthropoid, he recognized them as more consequential than the most post-cranial fossils transmitted to Dubois; when first seen by Dubois, Femur II was partially encased in hard pyrite-bearing rock, a lithology known to have been common in the **LB** and **HK** (S II-A4n, -F7 to –F9; also, Carthaus 1911, Berkhout and Huffman 2021: 73). Very coarse-grained sandstone still fills the medullary space of this specimen, as it does in Femur I (S II-F1, -F2, -F4 and –F9i; also, Ruff et al. 2013, 2015, 2021).

Although Dubois (1907, 1908) published little on the geology of the 1895-1900 excavations, his unpublished materials reveal a continuation of both the presence of the fossil-rich concentration lying near the low-water levels (Table 1) and the poorly fossiliferous condition of the superjacent eight-to-nine-meters of strata (our units **2**-**5**; Figures 4 to 5, S I Figures 4 to 6). Dubois made a full-body standing reconstruction of *Pithecanthropus erectus* for the 1900 Paris International Exposition (Shipman 2001). The model was based on Skullcap and Femur I, of course, but a recent rendering has been done using Femur II in place of Femur I (https://www.kenniskennis.com/homo-erectus/).

### 1907-1908 FOSSILS

Information on the Selenka Expedition’s left-bank Pit II gives a higher level of detail on the fossil content of the main bonebed than Dubois’ records do. This permits the spatial density of fossils in the **HK** to be estimated better, and the Expedition documented one localized fossil concentration in the overlying section, the *Stegodon* bonebed (**SB**). Together the Dubois and Selenka records allow for a reliable characterization of the unusually diverse main bonebed biota (Tables 1 and 2).

#### Hauptknochenschicht (HK)

Oppenoorth readily identified the *Pithecanthropus erectus* discovery deposit on the left bank in 1907 (S I Figure 8a, Endnote F(iv) and F(viii)). The final Pit II that year expanded modestly upon Dubois’ 1900 Trench, and was ∼37m east-to-west and ∼4m to ∼9m north-south (Figure 7a, S I Figure 7). Oppenoorth (1908a: 181) mentions that *“about 700”* fossils originated from this excavation in 1907 (Endnote F(ii) and F(ix)). The 1907 Listing enumerates 506 **HK** finds (from field layers 3 and 4, Table 2). Sixty percent have been attributed to taxonomic categories (243 of 405; 36% of total Pit II entries). Terrestrial species comprise ∼93% of the identified **HK** finds (225 of 243). Cervid and large-bovid specimens make up ∼86% (61% cervid plus 24% large bovids). *Stegodon trigonocephalus*, *Sus brachygnathus* and *Duboisia santeng* account for 1-5% each of the 243 identified specimens from 1907 Pit II (Table 2). *Rhinoceros*, hippopotamus and primate occur as one or two entries apiece. The remaining finds are fish, Crocodylia and bird fossils. The assemblage is substantially the same as the one in the DC (Table 1; de Vos and Sondaar 1982), although total frequency of cervids and large bovid is somewhat greater than the 68% sum of these taxa in the DC (Table 1).

#### Spatial variations of bioclasts in the HK

The density of vertebrate fossils in the **HK** was approximately three bioclasts per cubic meter, although the density differed between Pits I and II and varied vertically within the **HK** of Pit I.

Just a month of leaving the field during 1907, Oppenoorth wrote that in “the actual bone bed…. fossils are distributed rather randomly and one can find parts of a Stegodon, deer, buffalo, predator, crocodile etc.” (Endnote F(i)). He soon specified that the **HK** of 1907 Pit II had “produced about 700 fossils over a surface area of 250 square meters … mostly smaller bones, teeth, vertebrae, hand [fore-] and foot [hind-limb] bones, etc.,” making Pit II average ∼2.7 m^-2^ (Endnote F(viii)). Because the **HK** was no more than about one-meter thick, the average volumetric density in the bonebed exceeded ∼2.7. But Oppenoorth was clear about the varying spatial distribution of bioclasts. “Many times, the number of specimens found per square meter was larger (or smaller) [than the average],” and “sometimes more than 100 specimens had been deposited within a few square meters” (Endnote F(ii)).

The bioclasts from the 1907 Pit II were generally smaller than those in the 1891-1893 pit and trenches from which the Skullcap and Femur I came. The **HK** assemblage from the 1907 Pit II also differed in several ways from the fossils in Pit I, which was located on the other side of the Solo River (Figure 6b). In Pit I, *“about 1224 pieces …. were spread over a surface area of about 350 square meters”* in the **HK** giving *“an average* [fossil density] *of 3.5 pieces per square meter …. mainly* [comprised of] *the large bones like skulls, pelvis, vertebrate, ribs, etc.,”* as opposed to the smaller bioclasts and less-dense fossil occurrence of Pit II (Endnote F (ii)).

The Pit I fossils were concentrated in the upper and lower levels of three **HK** stratigraphic subunits (Table 2b, Endnote F(ix)). This vertical bioclast distribution was essentially the opposite of that in the 1891-1893 left bank excavations, where **PFZ** fossil concentration was in the middle of the **LB** (Figure 2a). The 1907 Listing also makes plain that in Pit I the number of finds (per subunit) ranges spatially from zero to ten in individual meter-sized squares (S I Figure 11c).

Despite the variations of bioclast-density in the Pit I **HK**, the fossil assemblages in the three subunits were consistent in taxonomic composition. Cervids, large bovids and *Stegodon* fossils comprised 90%, 93% and 92% of the finds in 15, 16 and 17, respectively; cervids were 63%, 74% and 62% of the finds in the three subunits (Table 2b). The consistency provides strong evidence that the subunits derive from the same precursor taphonomic events, and thus that the differences in spatial frequency were sedimentological consequences.

Antlers were especially common in the **HK** of 1907 Selenka Pit I (Table 2b) but infrequent in Pit II, where most of the deer were in the western portion. KdW had noted the large number of antlers in the right-bank **LB** in 1891 (Endnote A(ii), S II-A1b; also, S II–A1d). The 1892 25-m Trench apparently produced more antlers from the **LB** than the Skullcap Pit. By the end of 1893, Dubois (1896f, S II-F4: 725) had seen *“hundreds of complete antler beams and fragments”* (also, Endnote C(iv), S II-B6d).

#### Vertebrate fossils from beds overlying the HK

The 1907 Listing enumerates two fossil concentrations from above the **HK** in Pit II (Table 2). Seventeen finds are listed for Pit II layer 1, the *Stegodon* Bonebed, **SB** (Figure 8). The fossils overwhelmingly were the disarticulated remains of a *Stegodon trigonocephalus* individual, including a partial cranium with dentition, a maxilla with dentition and tusks, a half pelvis, some ribs and other post crania (S I Figures 9 and 10; also, van den Bergh 1999). Other **SB** fossils were crocodile, hippopotamus, fish and mollusc. No cervid- and bovid-remains, which were common in the **HK**, were reported being in the **SB**.

Pit II layer 2, which was situated at an unspecified level between the **HK** and the **SB**, had only 67 finds, compared to the 405 from the **HK** layers, and the taxonomic composition of the reported layer 2 species differed from that in the **HK** (Table 2a). Across the site as a whole, vertebrate fossils occurred *“sporadically”* above the **HK** (Endnote F(iii)), and consisted of *“incidental bone remains … here and there”* (Endnote F(vi)); these finds were mostly large bovid and *Stegodon*, rather than the deer which commonly occurred in the **HK** (Table 2).

Pit I occasionally encountered shallow bonebeds in 1908. A thin “red bone-bearing” lens was 4-5 meters above the **HK**; “2 thin lapilli beds (2 and 5 meters above the main bonebed) [were] on average 0.20 m thick” and “in appearance are identical to the main bonebed … [notably being as] rich in skeletal remains” and molluscs; these were “primarily Unio [a fresh-water mussel] and Melania [an aquatic gastropod],” which also were well-known in the **HK** (Dozy 1909: 609, Selenka and Blanckenhorn 1911 Tafel X; Berkhout and Huffman 2021: 49). The Selenka reporting about shallow right-bank bonebeds matches Kriele’s encounter of “a complete thigh bone, a complete tibia and a tusk, all of an elephant [Stegodon trigonocephalus], in addition to a few vertebrae and ribs” in the top 6m of the 1895 Dubois excavation, which probably was dug on the right-bank (S II-A4b, page 11).

One of the Selenka’s shallow bone-bearing lenses also might have been the *“richest fresh-water mollusc bed”* recognized in 1907 Pit I (Selenka and Blanckenhorn 1911, Tafel VI, Profil 2, Berkhout and Huffman 2021). The shallow *“thinner bonebeds … originated from … heavy eruptions”* of volcanoes in addition to having fluvial deposition like the **HK** (Dozy 1909: 611; also, Dozy 1911b: 35, Berkhout and Huffman 2021: 88). On the whole, the lithofacies *“overlying the main bone bed is highly variable”* at the site (Dozy 1909: 608). As noted above, plant-rich layers (without reported vertebrate fossils) were prominent above the **HK** in Pit I but rare in Pit II (Endnote H(viii)), and no plant beds described at all for the large left-bank excavations of Dubois, except to the extent the wood and leaves occurred in the **LB**.

### MAIN BONEBED FACIES

#### Foreword

Dubois’ and Selenka’s geological descriptions are irreplaceable resources for understanding the sedimentary and paleontological origin of the main bonebed, and are thereby valuable in drawing broader-scale biogeographic inferences about *Homo erectus*. As we document here, the main bonebed was poorly sorted, pebbly volcaniclastic sandstone studded with thousands of gravel-sized skeletal bioclasts. The fossils mainly were the remains of terrestrial species but included those of riverine vertebrates, fresh-water molluscs and plants (Figure 3). The vertebrate bioclasts ranged from isolated teeth to the craniums of cattle, buffalo and proboscideans (Endnotes A to F). Nearly all of the vertebrate remains were disarticulated and disassociated. The bony surfaces were not notably abraded by fluvial transport. The central question arising from these features is how thousands of large, disarticulated and little-worn vertebrate bioclasts accumulated in the company of numerous other biotic remains in a thin, localized, poorly sorted pebbly sand along a lowland river. No other *Homo erectus*-bearing deposit in Java has such broad biotic diversity and only one other hominin-fossil bed developed in a similar way sedimentologically (e.g., Tables 5 and 6). The following analysis evaluates the bioclastic and lithological features of the main bonebed in the light of paleogeographic context around Trinil and ecological information on the species that comprise the Trinil fossils assemblage. The DISCUSSION thereafter addresses the broad paleogeographic setting of the *H. erectus* and its associated fauna.

#### Main bonebed bioclastic features

The terrestrial-vertebrate skeletal material from the main bonebed consists overwhelmingly of broken disarticulated elements, and has uniform fossilization, fine-preservation of bony surfaces, low levels of weathering and abrasion attributable to fluvial transport, and lacks evidence of hominin- or terrestrial-carnivore involvement in the ungulate deaths. There is a great size range of both vertebrate and woody fossils (as large and long as proboscidean craniums and tusks and tree trunks). They were dispersed and apparently matrix-supported *in situ* with both spot-to-spot and vertical changes of density. The longest remains had bed-parallel orientations, and the largest fossils would have taken up most or all of the thickness of the bonebed.

These features are evident from firsthand discovery accounts and associated museum materials, both relating to thousands of specimens. The **LB** fossils that Dubois saw were *“generally isolated and widely distributed and usually broken”* condition and *“pieces that fit together are sometimes lying 20 to 30m apart;”* however, *“several vertebrae of a small ruminant were preserved in natural articulation,”* and *“a number of vertebrae and ribs of a large ox in their natural relative position and a piece of the great artery of a relatively small ruminant (probably a deer), entirely filled with andesite tuff”* (Endnote A(iii), S II-B2b, -B3b and -B3d), a specimen that turns out to be the bone of a bird rather than fossilized soft tissue. Carthaus remarked that *“articulated whole skeletons are absent”* in the **HK**, and the remains must have been *“transported more or less already decomposed animal corpses* [, and only] *a number of days, weeks or even months must have passed after these animals had been killed”* and then *“they ended up in the lahar flow,”* which carried them to Trinil (Endnote F(vi)).

Dubois and Carthaus both remarked on the low degree of fluvial abrasion on the hundreds of skeletal remains that they examined from the main bonebed. Dubois (1895b: 157-158) had noted of the **LB** that the *“bones do not appear to be rounded,”* despite coming from a *“fluvial”* deposit (S II-F2). After examining the 1891-1900 collection, he (1908: 1242-1243) thought that the bones *“were deposited in fresh condition”* (Endnote G, S II-F7). The common occurrence of long bones that are nearly whole anatomically is notable in the Dubois Collection (DC).

Carthaus (1911b: 26, 28) likewise concluded that “the main bonebed [**HK**] … is characterized by undamaged animal bones, without traces of rubbing against stones in flowing water” or other signs of “long distance transportation” (Endnote F(vi)). Dubois (1908: 1242-1243) thought that the occurrence of “hundreds of antlers of the same deer species … [was] explained by the simultaneous extermination of the entire herd of these Axis-like deer” (Endnote G, S II-F7). Oppenoorth (1911: xxxiv) observed in 1907 that “the bones were mostly embedded in broken condition … and in a few [instances, the breakage of **HK** bones clearly occurred] before fossilization [resulting in] … many bone fractures … filled with tuff” (Endnote F(iv)).

The broad-size range, broken shapes, and good surface preservations of the vertebrate fossils from the main bonebed also are clear in paleontological illustrations (e.g., Selenka and Blanckenhorn 1911, Hooijer 1958a) and museum collections. Hill et al. (2015) tested Dubois’ and the Selenka geologists’ impressions quantitatively, using 3736 Trinil vertebrate fossils in the DC and Selenka material at the Museum für Naturkunde, Berlin (MNB; the study sample was 68% of the assemblages). The material examined consisted dominantly of cervid and large-bovid specimens.

Overall, a *“limited amount of pre-burial weathering and transportation damage”* is evident (Hill et al. 2015). Comprehensive taphonomic evaluation of 234 humeri, representing 4.4% of the Trinil assemblage in DC and MNB, revealed that ∼95% of the specimens lacked notable signs of abrasion rounding (codes 2 and 3 of Fiorillo 1988) and weathering (stages 0 and 1 of Behrensmeyer 1978; M. Hill and L. Todd, pers. comm. 2015). Fractures are often sediment-coated or -filled (Hill et al. 2015, M. Hill and L. Todd, pers. comm. 2015, pers. observation).

The DC vertebrate specimens have strikingly uniform fossilization, and exhibit a higher level of preservation than we have seen at other sites in the *Homo erectus*-bearing formations of eastern Java. The taphonomic condition of the terrestrial vertebrate fossils is consistent with deaths a few months before burial, as Carthaus thought.

Dubois further observed that *“in no case were the usually recognizable signs of the teeth of land predators undoubtedly observed”* on the ungulate fossils of the **LB** (Endnote B(i), S II- B3d). Hill et al. (2015) verified this 125-year-old conclusion after having examined the 1891-1907 finds in the museum collections (also, Choi 2003). Dubois focused on the taphonomic role that crocodiles played in the **LB** formation. He maintained that they served to *“break …. and distribute”* the bones after carcasses of terrestrial animals arrived at Trinil, their *“soft tissue”* theretofore having *“protected* [the bones] *against wear at the bottom of the current”* (Endnote A(i), S II-B3b, -B3d).

On the other hand, H. Stremme (1911) noted that only one well-preserved **HK** specimen in the Selenka materials had punctures that were probably referable to crocodile predation (MNB MB.R.1959; in Selenka and Blanckenhorn 1911: 146). Dubois had had special anthropological reason for interest in the crocodiles. He (1926a, S II-F9) reported that Femur I was one of the remains that had crocodile damage: The *“caput femoris, preserved for the most part, presents however extensive defects on the margin of the globular articular surface, which were probably caused by crocodiles.”* Hill et al. (2015) confirmed that circular compression fractures are present on certain Trinil specimens (e.g., DC 1860 and MNB MB.Ma22309 at left; MB.Ho.476.1 above), but the marks on the proximal end of Femur I are less conclusive (M. Hill and L. Todd, pers. comm. 2015).

Unsurprisingly, no porcupine gnaw marks have been reported from the bony remains or teeth in the MB, although porcupine fossils are present in the DC materials from Trinil (*Hystrix lagrelli*, NISP = 2). Gnawed teeth occur in geologically younger fossils in eastern Java cave deposits (e.g., at the Punung and Gunung Dawung sites; Storm and de Vos 2006, Storm et al. 2005, Storm 2012). In terms of the mortality events that led to development of the **HK**, Carthaus (1911b: 28-29) surmised that the *“animals* [whose remains were embedded as fossils in the deposit] *had been killed during the initial explosive eruption”* at a distant volcano (Endnote F(vi)). A similar scenario made sense to Dozy (1911b: 36, Berkhout and Huffman 2021: 91):

> *when huge rains fell on the slope of the Lawu and Wilis volcanoes, enormous quantities of loose tuff and lapilli … were swept into the plains … destroying everything. This same process presently still takes place (this year there was a very similar catastrophe … near the volcano Semeru* [Figure 1b]). *Skeletal parts …, which lay on the ground, were swept along and embedded in the tuff layer* (**HK**).

Several additional features of the embedded condition of the main bonebed bioclasts are of interest in considering the depositional mechanisms by which it accumulated.

First, elongate skeletal fossils, particularly tusks one-or-two meters long, must have lain parallel to the boundaries of the main bonebed, which was often less than a meter thick. In 1891, *“the tree trunks and leaves are always found horizontally*” (Endnote A(v), S II-B2e; also, S II- B2d). Segments of trees 1-3m long had a *“random”* attitude within the **HK**; that is, they were horizontal without any preference azimuth orientations (Carthaus 1911b: 14, Berkhout and Huffman 2021: 67). The long-bones and ribs of large ungulates, complete deer antlers and cattle-buffalo horn cores presumably also had bed-parallel attitudes in the main bonebed.

Second, cranial fossils of *Stegodon*, *Bibos*, *Bubalus* and *Rhinoceros* (when the remains were largely complete) must have taken up the full thickness of the main bonebed. For example, large cranial fossils in the 0.20 m-thick **PFZ** would have extended into or sometimes through the upper **LB**, as for instance f the *Stegodon “tusk and skull”* visible in the initial left-bank **LB** outcrop, and *“skull of a banteng almost complete”* and *“elephant skull”* found in **LB** on the right bank in 1891 (Endnote A(i) and A(iv), S II-A1b and -A1c). Likewise, the *“mandible and tusk”* of *Stegodon* that de Winter remembered being in the **PFZ** *“nearby”* Femur I would have taken up the whole **PFZ** and extended into levels above it (also, Endnote A(xii), S II-A1j).

Remains which took up the full thickness of the main bonebed would have included the *“complete turtle”* carapace found in 1896, *“skull of a cow with complete horns”* and *“skull with 1 complete horn of a water buffalo”* of 1897, *“elephant skull with complete tusks”* in 1899 and *“complete buffalo skull with complete horns”* of 1900 (Endnotes D and E, S II-A4c, -A4d, -A4h, -A4l and -A4o).

#### Main bonebed lithofacies

Archival accounts contain less information on the lithofacies of the main bonebed than they do on its bioclastic content. However, Dubois and the Selenka geologists offered some key lithological observations, and most helpfully, identified modern depositional analogs to the main bonebed. In 1895, Dubois summarized the lithologies excavated in 1891-1893:

> *Bones are present within beds of tight and hardened volcanic tuffs, consisting of clay, sand and lapilli rocks. These tuffs suggest a fluvial origin, especially indicated so by a strong general presence of fresh water animals* [such as molluscs] *and by certain fluvial structures that English geologists call current bedding* [cross-bedding or -lamination]. (Dubois 1896b: 251, S II-F1)

Evidently, the gravel in the river mixed with volcanic ash and sand, and included skeletal, shelly and vegetal clasts (e.g., Endnotes A to C; S II-B6c, -B2b, -B2d, -B2h, -B3f, -B4b and -B5b). Rather than volcanic ejecta, the **LB** lapilli were epiclastic granules and pebbles of volcanic lithologies, based on the matrix on museum fossils.

The “hardened” rock fits the firsthand reporting of substantial induration (e.g., the strata were *“so terribly hard that it is almost impossible to get through,”* even in the *“lower part;”* S II-A3a-vi, -A3a-ix). The endocranial space of the Skullcap itself was filled with the indurated pebble conglomerate, which consisted of fresh, very poorly sorted, volcanic minerals and rock fragments within a fine-grained matrix (Figure 2c, d; also, S I Figure 20).

The clastic materials adhering to Trinil bioclasts in the DC range from granule-pebble conglomerate to very fine-grained sandstone (S Figure 20; Huffman et al. 2018). Long bones that are anatomically complete tend to have finer-grained- and partial-infills of clastic material, whereas bones that were broken *in situ* tend to have coarser-grained fills and adhering sediment (Hill et al. 2015, M. Hill and L. Todd, pers. comm., 2015).

Based on the broad size range of DC fossils, they represent the concurrent deposition of hydrodynamically different elements of Voorhies (1969) dispersal groups (e.g., craniums with long bones). Labile minerals in the sandstone affixed to the specimens include plagioclase, hornblende and pyroxene, indicating that volcanoes in the Trinil paleo-river uplands were shedding fresh rocks, crystals and glass.

The competence of the flow responsible for the **HK** was evident in its cobbles and boulders of hard andesite, pumiceous gravel, rip ups of *“clayey marl cobbles”* and clots of entwinned antlers (e.g., Oppenoorth 1908a, Endnote F(ii), and Oppenoorth 1911: xxxiv and Carthaus 1911b: 14; also, Berkhout and Huffman 2021: 38 and 73). The clasts were

> *ash, very-small as well as somewhat-larger lapilli* [pebbles], *which in part show transition* [in internal fabric] *to a pumice*[ous] *structure, and also pieces of pumice*. [Furthermore] *here and there, rounded cobbles of dense solidly crystallized andesite and andesitic lava are found, even up to large size* [boulders] …. [along with] *cross-wise lying pieces of tree trunks and branches up to 1- or 3-meters long* [and other bioclasts of varying densities]. (Carthaus 1911b: 14, Berkhout and Huffman 2021: 67).

The tusks, tree trunks and very large bioclasts of more equant shapes appear to have been rafted into place amidst a dense fluvial load of mud, sand, pebbles, gravel and smaller bioclasts.

The lithofacies of the bonebed fits transport *en masse* under suspended- and traction-movement attained during a sediment-heavy flooding, as opposed to common modes of episodic river-channel sedimentary transport and deposition. The main bonebed might be thought of as a bioclast diamicton wherein biotic- and lithic-materials were emplaced simultaneously (see Pantin 1967 for types of diamictons). The cross-bedded **HK** seen today along the left bank (e.g., S I Figure 2b), and the cross-laminated **LB**-**HK** that Kriele observed in 1900, must have formed during traction transport, but the oversized vertebrate bioclasts probably had been carried to Trinil by hyperconcentrated flow.

Not all of the main bonebed was conglomeratic sandstone, judging from Oppenoorth’s reports on 1907. He (1908b) saw fossiliferous *“bluish volcanic tuff, which reminds me of a very soft sandstone,”* and reported that the **HK** *“consists of three portions …* [in which] *finer-grained grades into a coarser-grained layer”* containing cobbles and perhaps small boulders *“a few decimeters”* in diameter. *“Fine blue clay with harder clay concretions”* made up the upper **HK** (Oppenoorth 1911: xxxiv, Berkhout and Huffman 2021: 35). The **HK** in Selenka Pit I did have three vertical subunits with spatially varying bioclast densities (Table 2, S I Figure 11c).

Although the sedimentary and paleontological differences in the main bonebed suggest that variable flow conditions occurred from moment-to-moment and place-to-place during the responsible depositional events, firsthand reporting lack evidence that a long cessation of accumulation took place during the formation of the main bonebed.

#### Modern depositional analogies

Many comments that Dubois and the Selenka’s geologists made about the origin of the main bonebed draw links to historic fluvial deposition of volcaniclastic sediments observed around the active volcanoes of Java. Even without modern descriptive-and analytical-skills, these men could make visual comparisons of the bonebed to historic laharic sediments and see how sedimentary processes in modern lahar-prone drainage basins might apply to the Trinil stratigraphic sequence. Their inferences further benefitted from leading-edge levels of vulcanological knowledge during that turn-of-the-century era (e.g., Neuman van Padang 1983, Voight et al. 2000).

In mid 1892, when Dubois first-considered the origin of the **LB** vertebrate fossils deeply, he wrote, “only a catastrophe, and a volcanic catastrophe at that, comparable to, but on a larger scale than, those that accompanied eruptions of the Salak (1699), Galungung (1822) and Kelud (1848) [stratovolcanoes in Java] can explain … these accumulations” of the lapilli-bearing fossil beds (S II-B3d). He earlier concluded that:

> *The fossil-bearing sediments* [of the Kendeng Hills] …. *appear to have been deposited in the same manner* [as the Recent sedimentary] … *rocks of the lowlands. Historical eruptions of the Kelud that delivered products to the Kediri lowlands* [along the Brantas River drainage], *consisting of sands, sometimes hardened to sandstone, tuffs and breccias,* [which are] *indistinguishable from the Pleistocene on the Kendeng slopes. Sedimentary rock material that encloses the remains of the Pleistocene Java fauna has undoubtedly similarly been carried to its destination by an eruption. This would have been partly in dry condition as volcanic sand, lapilli and bombs etc., but especially during heavy rains that mixed with them in the form of heavy slurry flowing down the slope* [lahars]. *The animals would have perished in the same manner that the inhabitants of the Kelud slope can now tell us about during the historical eruptions of this volcano. After the last eruption, many cadavers of pigs, kidangs, deer, bantengs, tigers and other forest animals were found on and within the volcanic sand etc*. (S II-B1h; also, Dubois 1892a)

Carthaus also drew attention to 19^th^ century events in the Kediri lowlands of the Brantas River (East Java), which drains the active Kelud volcano in East Java (Figure 1b) For him (1911b: 27-28, Berkhout and Huffman 2021: 86), *“portrayals* [of flood conditions on the Brantas] *are quite appropriate to explain the rich occurrence of animal bones and pieces of wood in the main bonebed of Trinil.”* He (1911b: 27, Berkhout and Huffman 2021: 86) gave an example wherein the Brantas River carried a man by *“lahar sand flow during the last eruption”* of Kelud volcano, but *“did not suffer any hard knocks from the rocks … in a flowing mass of very thick slurry.”*

Carthaus concluded that “the main bonebed of Trinil is the … product of an extraordinary large lahar flow, which originated from [an emptying of] an erstwhile western crater of the Wilis,” analogous to the historically active volcanoes at both Kelud and Semeru (Figure 1b; Carthaus 1911b: 16, Berkhout and Huffman 2021: 76). Dozy (1911b; 21, Berkhout and Huffman 2021: 80) imagined the slurry “loose volcanic material, mainly ash and lapilli.” Carthaus referred to “sand flows” to emphasize the dense sandy flux in lahar flooding which he envisioned responsible for the **HK** (Carthaus 1911b: 14, 27, Berkhout and Huffman 2021: 75, 86).

Carthaus’ (1911b: 29) thinking was influenced by catastrophic deaths near the Semeru stratovolcano, where in 1909 *“more than 500 people lost their life”* in *“an immense tuff mudflow which,* [took place] *during an enormous rainfall in the upper regions …* [concurrent with] *incessant ash, lapilli, pumice and volcanic bombs … from the volcano”* (also, Cool 1909). Semeru regularly has produced deadly lahars since that year (Lavigne and Suwa 2004). In 1919, a deadly lahar flooding again struck a Kelud drainage, killing 5110 humans and 1571 livestock; the event illustrates the potential for lahars to produce mass death when large-mammal populations are concentrated geographically (B. Voight in Huffman et al. 2010b).

Carthaus (1911b: 21; Berkhout and Huffman 2021: 80, 91) further surmised: “The Trinil conglomerate [underlying the **HK**] may thus possibly be the product of the first outpouring of enormous tuff mudflows from the giant western crater of the Wilis that was probably filled by a huge lake [such as the one in the Kelud caldera of 1909].” Carthaus carried forward the analogy when stating, “the overlaying main bonebed … arrived in the vicinity of Trinil during continuation of the eruption either through the same channel as the prior tuff mudflow or in a different one created by damming.”

Dubois and the Selenka geologists evidently saw the main bonebed, and perhaps overlying strata, as a continuation of the laharic paleogeographic regime which is so prominent in diamictons of the underlying formation (Pucangan Formation of Duyfjes 1936). These men were perceptive to focus on long-run out lahar flows as a mechanism for transportation and accumulation of the main bonebed (Huffman et al. 2012b). Subsequent research on lahars amplifies the broad spectrum of geological conclusions one might draw from the identification of lahar deposits within a sedimentary sequence.

In the Pleistocene of eastern Java, the implications of lahar include insights into: (i) Regional paleogeography (stratovolcanoes existed in the hinterland, and the watersheds were subject to fluvial processes that started as lahars). (ii) Paleo-vulcanology (lahars and coarse-grained sandy deposits of volcaniclastic materials often reflected penecontemporaneous volcanic eruptions, while little- or no-such volcaniclastic accumulation might imply protracted dormancy of the volcanoes in the watershed). (iii) Paleo-sedimentological setting (a high-rate volcaniclastic deposition occurred in the lowlands and adjacent water bodies; Figure 11). (iv) Depositional mechanisms (modes of fluvial transport related to lahars contributed to sedimentary accumulation down-drainage, particularly when debris flows evolved into floods).

**Figure 11.**
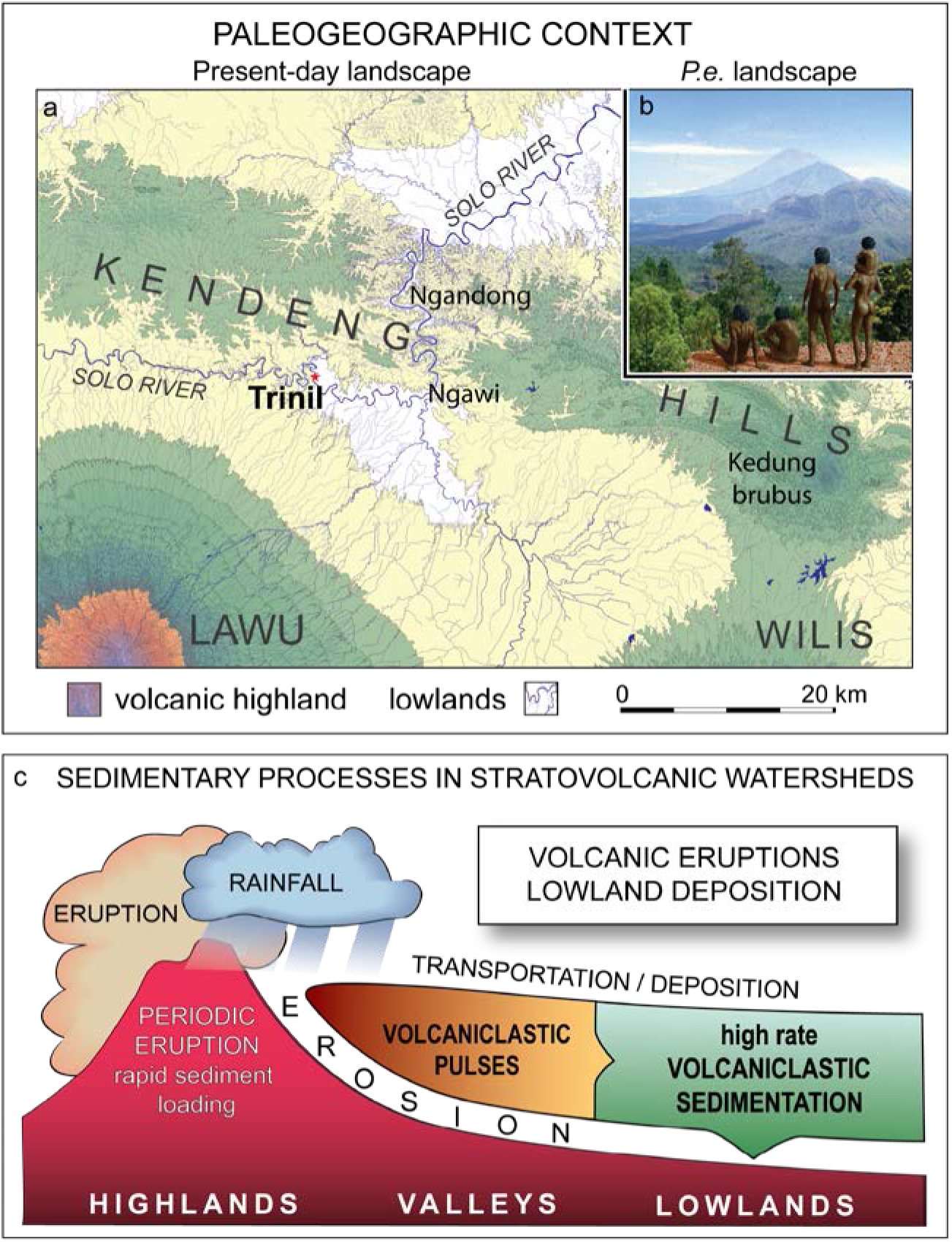
(**a** and **b**) When the *Pithecanthropus erectus* lived in the Trinil paleo-river watershed and the main bonebed formed, the sedimentary processes were similar then to those in the historic Trinil stratovolcanic watershed. The lowland near Trinil today is less than 50m above sea level; to the south, Lawu stratovolcano rises above 1000m (green) along a dense parallel network of drainages (presumably having complex biotopes), and passes upwards through forested highlands (red) to a crest at 3118m. The Solo River flows from west to east along the southern edge of the Kendeng Hills, and shifts course sharply at Ngawi to traverse the Hills through a deeply incised valley (Solo River gap), where the Ngandong *Homo erectus* site is located (Table 6-B). The modern Solo watershed upriver of Ngawi covers 9827 km^2^ and the Madiun River portion is 3765 km^2^ (∼7% and ∼3%, respectively, of Java as a whole). (**c**) Other *H. erectus* sites in eastern Java also formed in stratovolcanic watersheds similar to modern ones (Figures 12 and 13), as did archaic fossil hominin sites in western Java, Flores, Sulawesi and Luzon. Stratovolcanoes dominated the terrane encountered by all hominin populations known to have lived in the region (Huffman et al. 2010b, 2012b). Part of the (b) image is reinterpreted from a Turner and Antón (2004) illustration.

**Figure 12.**
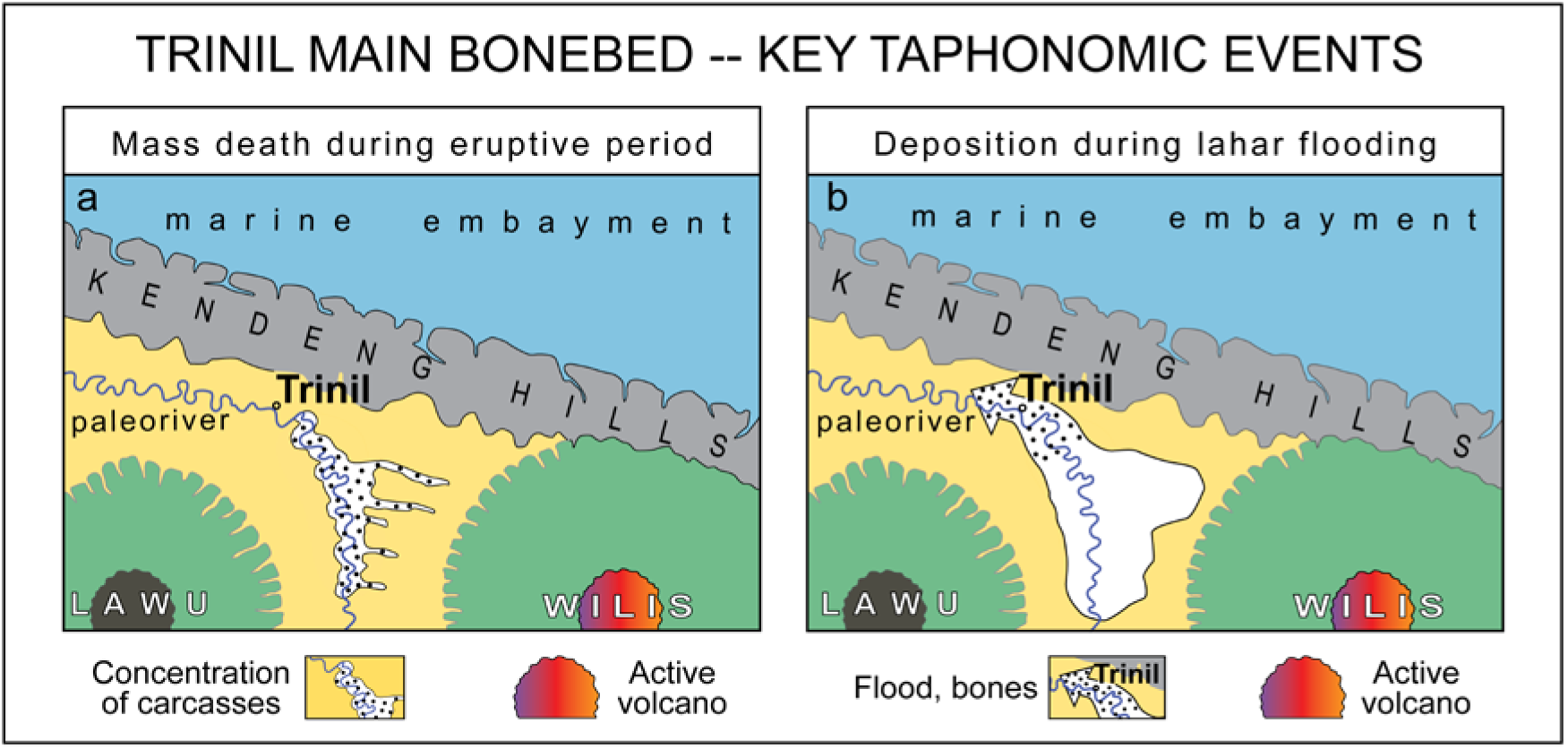
The Trinil main bonebed evidently developed in two stages (partially after Huffman et al. 2010a,b, 2012b, 2018). (**a**) The accumulation of the main bonebed was preceded by a mass death of ungulates along a flood plain up river from Trinil, and the resulting carcasses were skeletonized. (**b**) Lahar flooding entrained the bone remnants and carried them (with aquatic reptiles and molluscs) to a point of accumulation at Trinil. This portrayal has the Trinil paleo-river flowing westward, and eruptions occurring at paleo-Wilis, a large stratovolcano that existed near modern-day Wilis during deposition of the Pucangan Formation (Figure 13; Huffman 2001a,b, 2020). Had Lawu been the primary source for clastic materials in the main bonebed, the flooded part of the drainage system might have had a different configuration and the trunk river might have flowed eastward (e.g., Sartono 1976, Berghuis et al. 2021), but the sequence of events in the formation of the main bonebed would have been largely the same. Similar taphonomic benchmarks (e.g., mass death of ungulates) and sedimentary development (e.g., lahar-flood accumulation) led to the deposition of the Ngandong *Homo erectus* bonebed (Ngandong site, Tables 5 and 6-B; Huffman et al. 2010a, b, Rizal et al. 2020).

**Figure 13.**
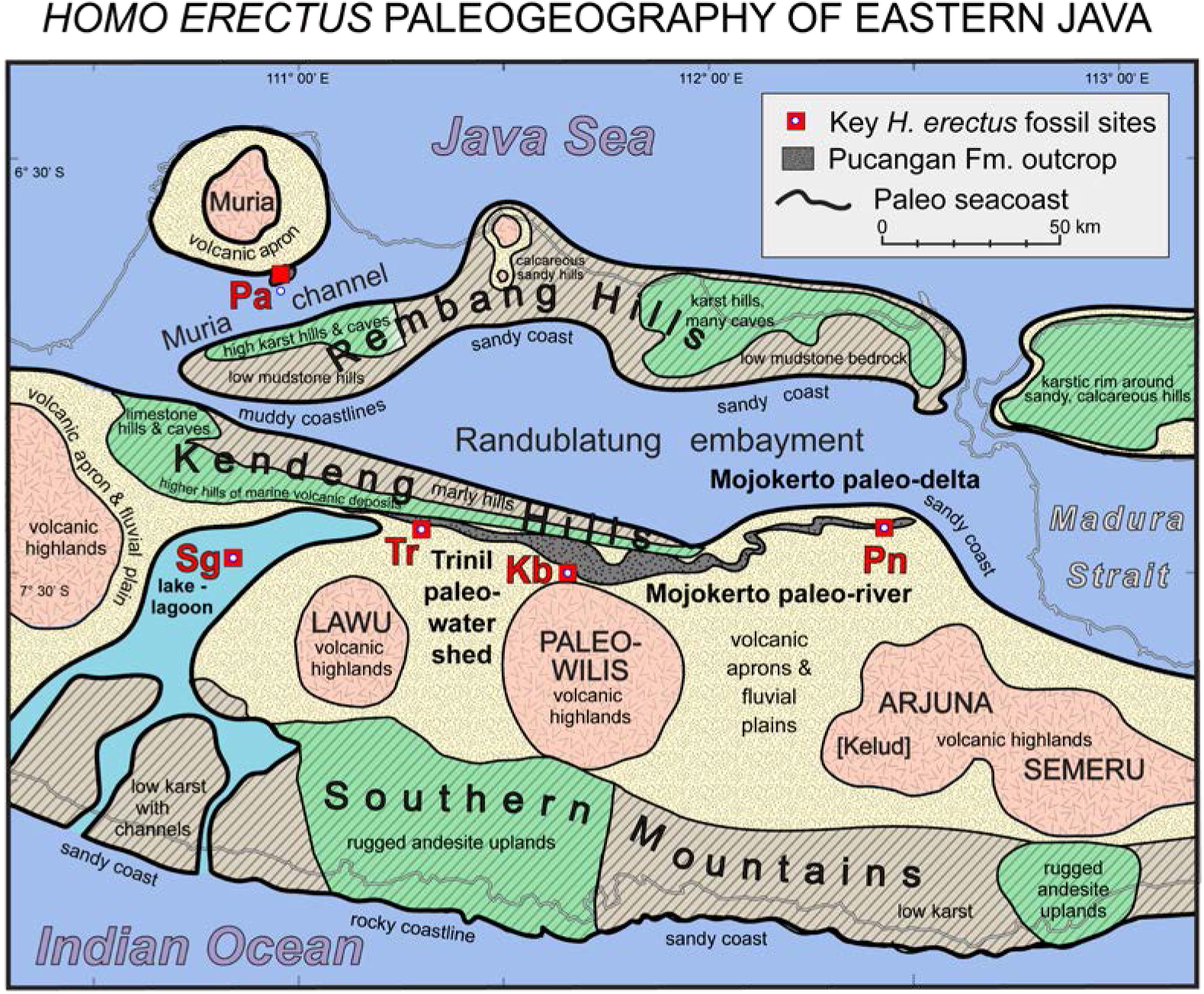
A broad variety of potential habitats were available to the *Pithecanthropus erectus* population in eastern Java, due to the presence of high-standing stratovolcanoes, hilly pre-Pleistocene carbonate- and volcanic-bedrock terranes, large- and small-rivers, various seacoasts, and islands reachable by short sea crossings (after Huffman 1997, 1999a, b, 2001a, 2020, Huffman et al. 2000). Figure 1b has the same key *H. erectus sites* displayed on a modern physiographic map with geological overlay. Figure 14 has a graphical summary of the range of *H. erectus* potential habitats. Table 6 provides information on the sites and key geological relations underlying this paleogeographic mapping. Prior to the Pleistocene, eastern Java had had a complex geological history of marine and volcaniclastic periods, but few large terrestrial vertebrates (e.g., Lunt 2013). By the time *H. erectus* arrived (Figure 15a), most large-scale physiographic features seen today were recognizable (compare to Figure 1b). This particular map is a generalized portrayal of the Early-Middle Pleistocene conditions under which the upper Pucangan Formation and lithostratigraphically correlative strata accumulated. Included for orientation is the outcrop area of the Pucangan Formation, which is distinctive in its abundance of gravelly volcanic diamicton. When the Pucangan was traced by field mapping from the Trinil- to Mojokerto-areas, the Formation was found to be thickest north of Wilis-Liman volcano (the Kedungbrubus-Butak area, ‘Kb’), indicating that the principal source of the lahar flows responsible for the diamictons was Pleistocene paleo-Wilis (Table 6-C and 6-D; van Es 1931 and Duyfjes 1936, Duyfjes 1938a-d in Huffman 2020: 23-35 and 41; also, De Genevraye and Samuel 1972, Huffman 2001a,b, Huffman et al. 2006, I.J.J.S.T. 1992, Lunt 2013: Fig. 135, Watanabe and Kadar 1985, and, multiple unpublished geological studies conducted for petroleum exploration, such as those used by Huffman et al. 2000). Other paleogeographic representations of the region are in Berghuis et al. (2021), Djubiantono (1992), Djubiantono and Sémah (1993), Lunt (2013), Rizal et al. (2020), Sartono (1976) and Zaim (1989, 2010).

**Figure 14.**
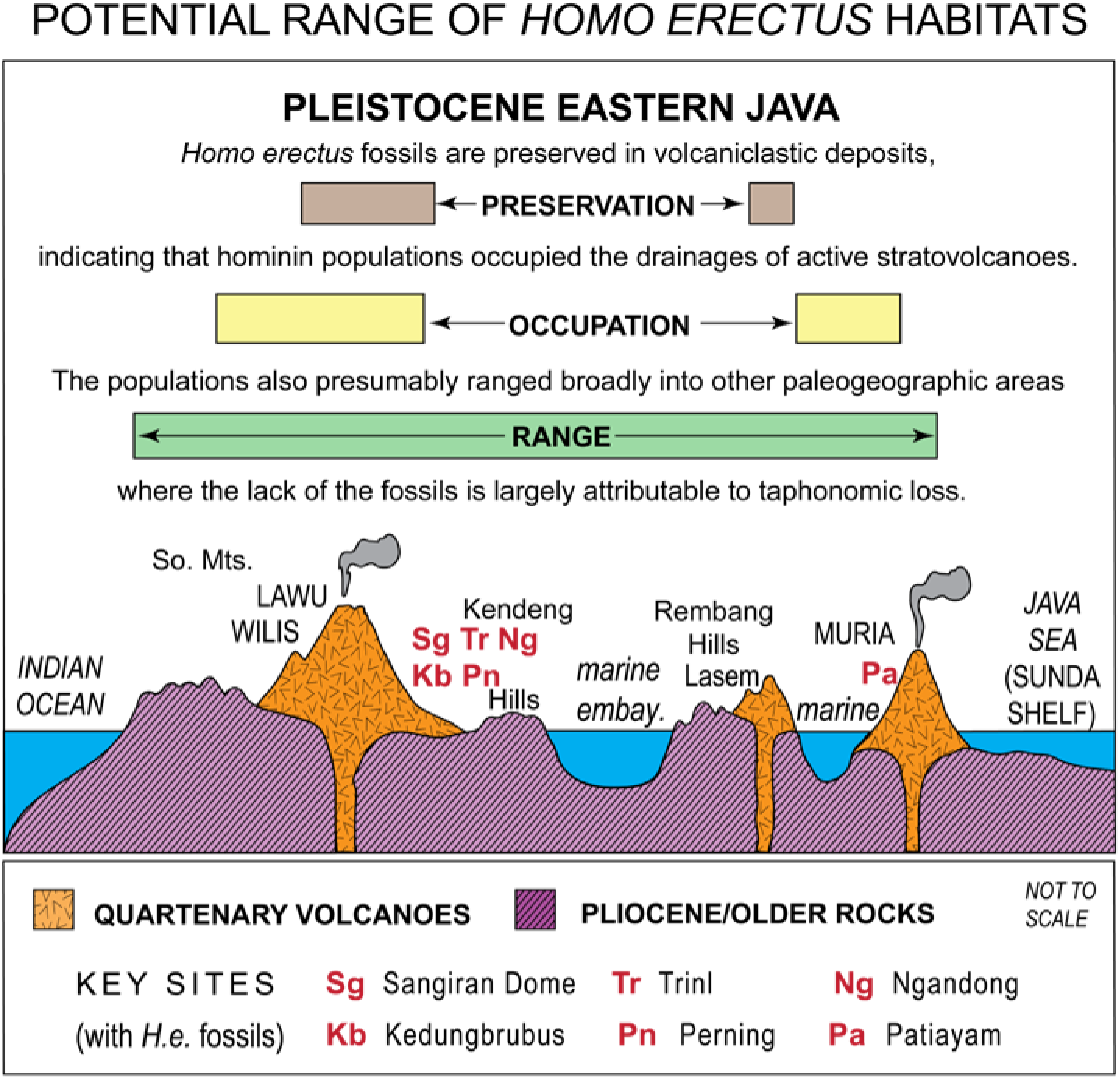
*Homo erectus* fossil occurrences in drainages of stratovolcanoes (Kb to Tr, Figures 1b and 13) strongly suggest that archaic hominins occupied or frequented parts of eastern Java where none of their fossils have been discovered, including the Kendeng Hills, Rembang Hills, Southern Mountains (Huffman 1999b, 2001a; also, S I Figure 22). There appears to have been insufficient volcaniclastic accumulation in these uplands to preserve the hominin skeletal remains (after Huffman et al. 2012a, b).

Additional insights might be on: (v) Paleoclimate at a basin-wide scale (stratovolcanoes capable of producing lahars are often sufficiently lofty to concentrate rainfall orographically and create rain-shadow effects). (vi) Paleo-hydrology (the rivers delivering lahars to the lowlands are part of an integrated drainage system which originated at volcanic peaks and varied in hydrological features from place to place; e.g., Figure 11). (vii) Montane paleo-vegetation (the highlands of the stratovolcanoes, if not also on their flanks, are forested). (viii) Mass death (both lahar floods and associated volcanism had the capacity to produce high rates of biotic mortality, particularly notable when thousands of large animals concentrate in the path of large volcanic- and fluvial-events).

The influence of lahar deposits in the Pleistocene hominin record of Java is well established (e.g., Bettis et al. 2004, 2009, Huffman 2001a, b, Huffman et al. 2006, 2010a, Rizal et al. 2020, Zaim 2010). Dubois had this insight in the 1890s.

### FORMATION OF THE MAIN BONEBED

#### Death of hundreds

Museum collections of Trinil fossils provide evidence that hundreds of individuals lived as a single paleofaunal community and died penecontemporaneously, much as Dubois and Carthaus envisioned when stating: *“The animals would have perished in the same manner that the inhabitants of the Kelud slope”* did during eruptions (Dubois 1892a, S II-B1h); the main bonebed resulted from a *“simultaneous extermination of the entire herd”* (Endnote G, Dubois 1908: 1242-1243, S II-F7); and the *“animals* [now **HK** fossils] *had been killed during the initial explosive eruption”* at a distant stratovolcano (Endnote F(vi), Carthaus 1911b: 28-29, Berkhout and Huffman 2021: 87).

Paleontological observations confirm that a single paleofaunal community was involved (Tables 1 to 3, Endnotes) with the Minimum Number of Individuals (MNI) in the in the Dubois Collection (DC; Storm 2012) indicating the death of hundreds of ungulates (Table 4). Among 3478 museum specimens, the MNIs for *Axis*, *Bibos*, *Bubalus*, *Duboisia*, *Stegodon*, *Sus* and *Rhinoceros* total 164 individuals (94% of all non-hominin taxa in terms of MNI). *Axis* specimens are about as common as each of the two large bovids (Table 4, Endnote F(i)).

**Table 4.**
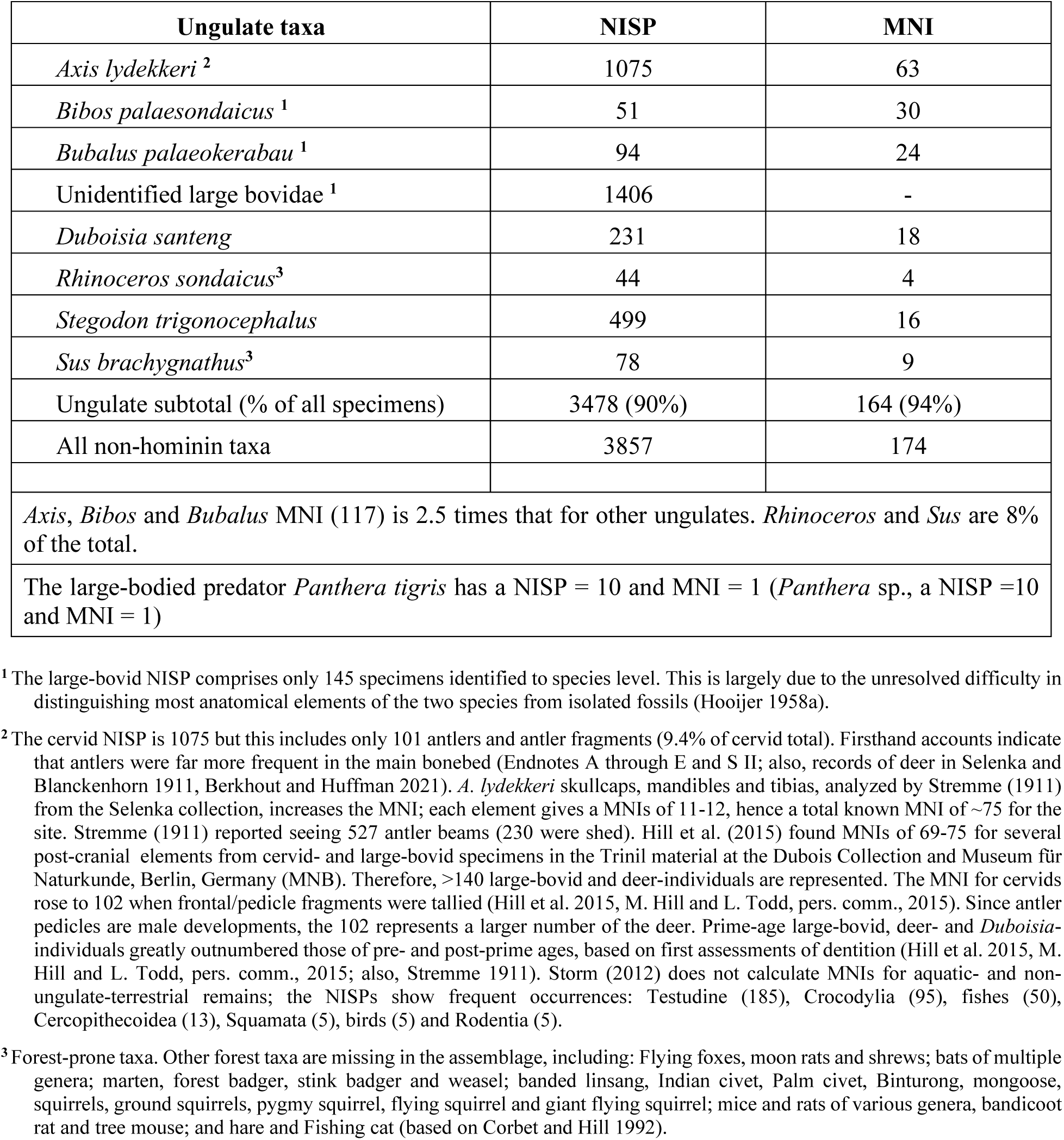
Minimum number of individuals (MNI) among the Trinil ungulate materials in the Dubois Collection (after Storm 2012).

| Ungulate taxa | NISP | MNI |
| --- | --- | --- |
| *Axis lydekkeri*<sup>2</sup> | 1075 | 63 |
| *Bibos palaeosondaicus*<sup>1</sup> | 51 | 30 |
| *Bubalus palaeokerabau*<sup>1</sup> | 94 | 24 |
| Unidentified large bovidae<sup>1</sup> | 1406 | - |
| *Duboisia santeng* | 231 | 18 |
| *Rhinoceros sondaicus*<sup>3</sup> | 44 | 4 |
| *Stegodon trigonocephalus* | 499 | 16 |
| *Sus brachygnathus*<sup>3</sup> | 78 | 9 |
| Ungulate subtotal (% of all specimens) | 3478 (90%) | 164 (94%) |
| All non-hominin taxa | 3857 | 174 |
| *Axis*, *Bibos* and *Bubalus* MNI (117) is 2.5 times that for other ungulates. *Rhinoceros* and *Sus* are 8% of the total. |  |  |
| The large-bodied predator *Panthera tigris* has a NISP = 10 and MNI = 1 (*Panthera* sp., a NISP = 10 and MNI = 1) |  |  |

<sup>1</sup> The large-bovid NISP comprises only 145 specimens identified to species level. This is largely due to the unresolved difficulty in distinguishing most anatomical elements of the two species from isolated fossils (Hooijer 1958a).

**Table 5.**
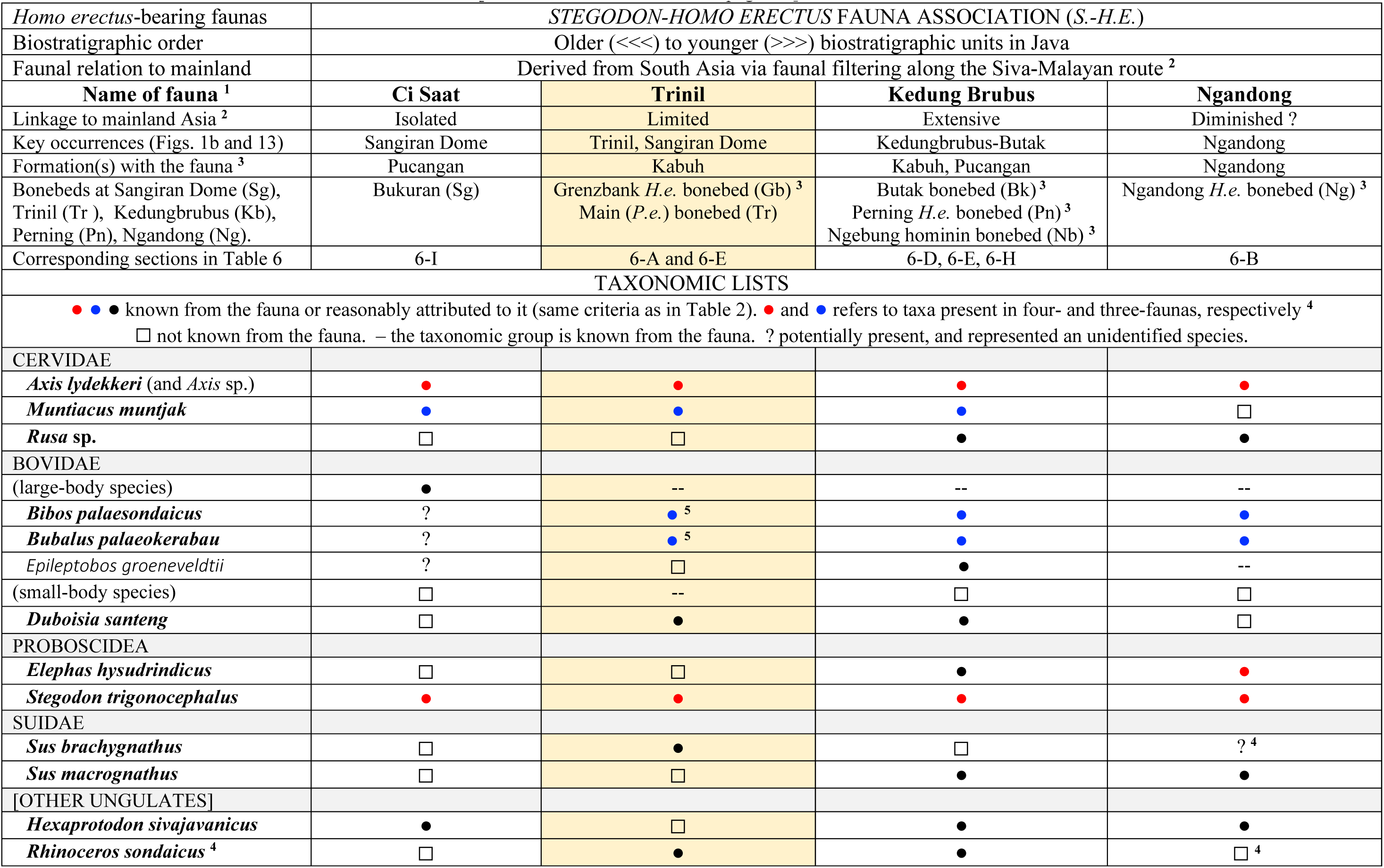
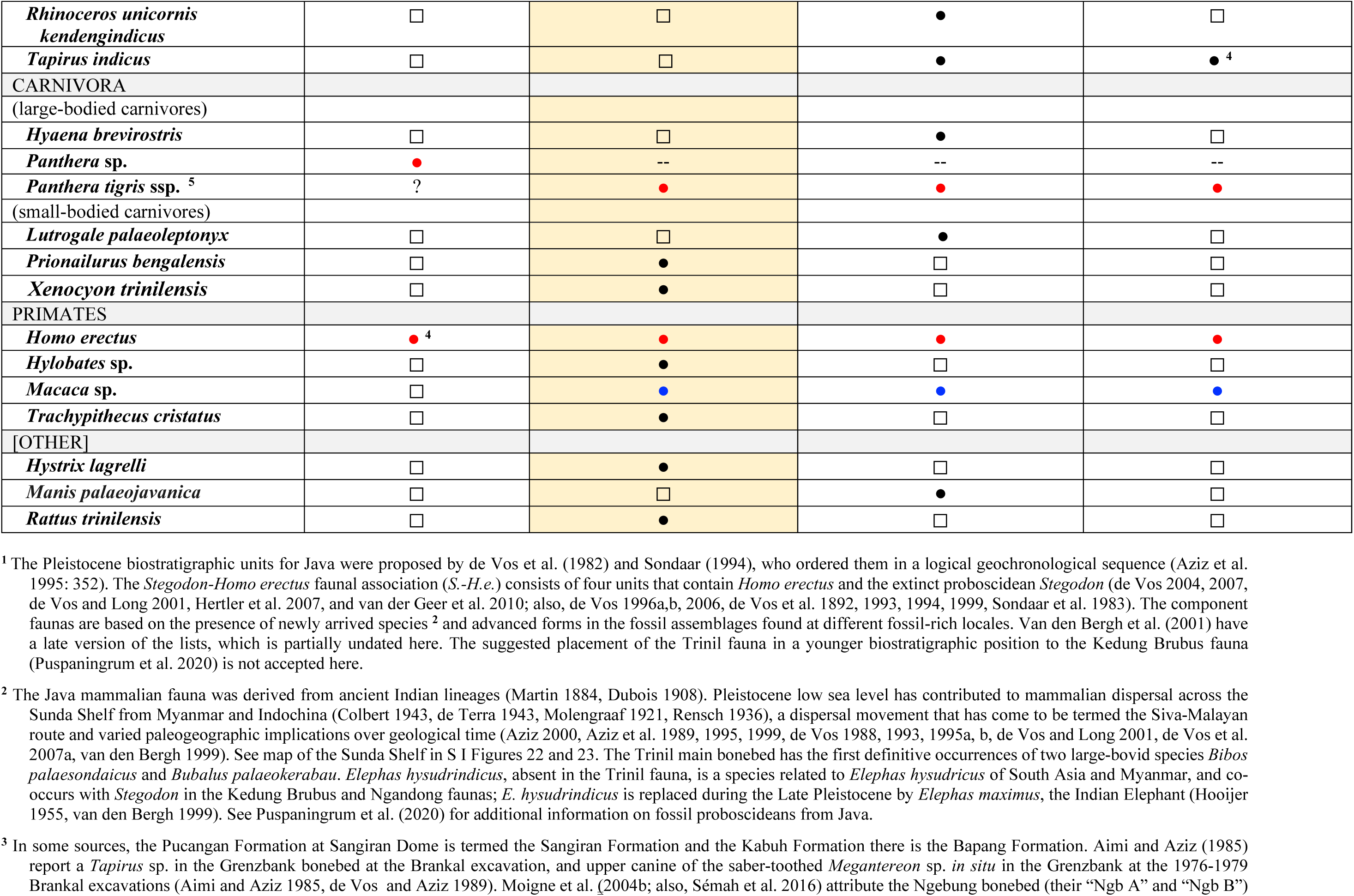
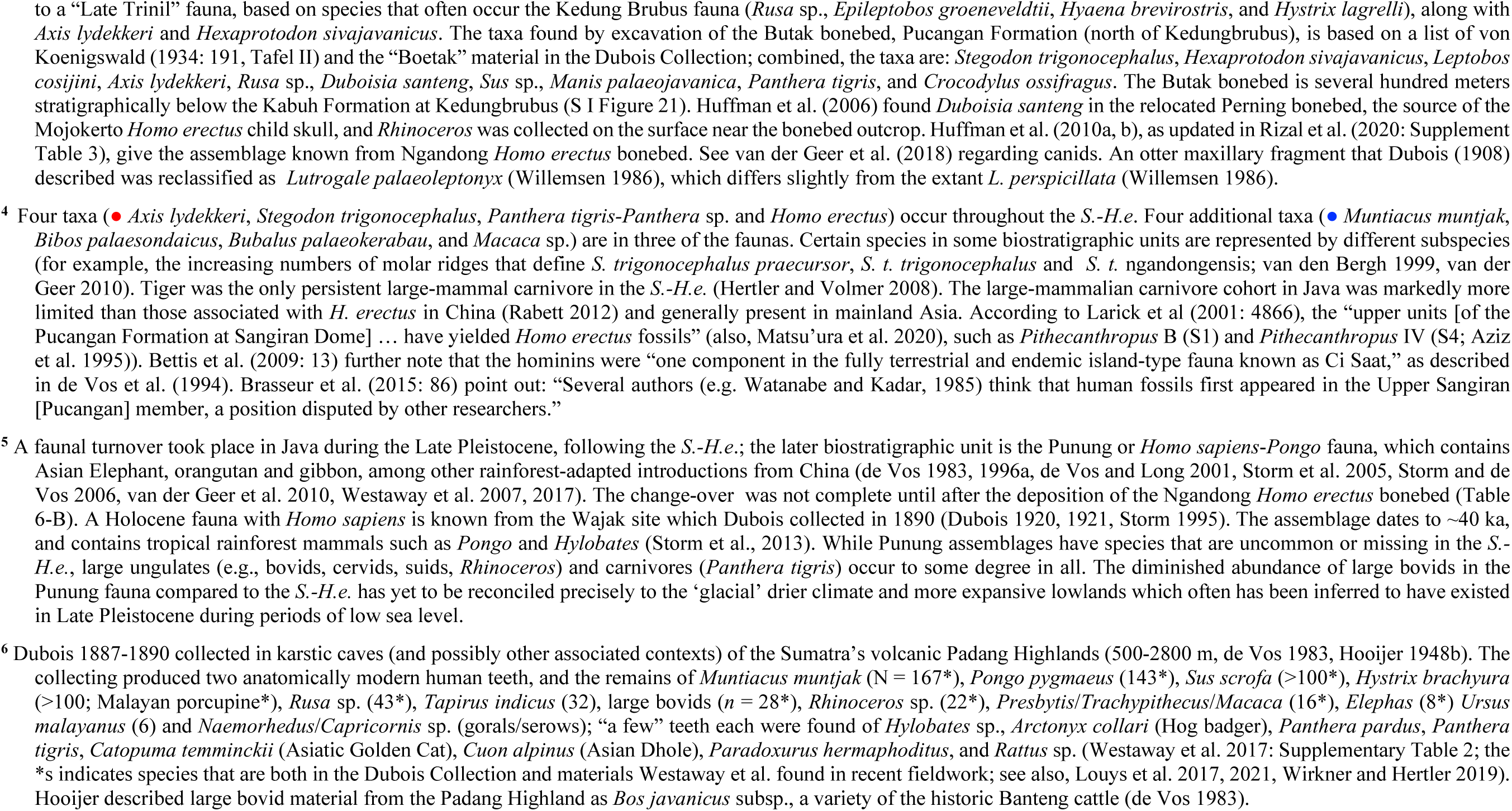
Trinil fauna (Table 3) compared to other biostratigraphic entities of the *Stegodon-Homo erectus* fauna association**^1^** (also, Table 6).

No taphonomic distinction has yet been recognized that would indicate that any large number of specimens in the assemblage is inconsistent with a single death event. Rather, Hill et al. (2015, pers. comm.) saw isotaphonomy among 3736 large-bodied vertebrate bioclasts examined in the Trinil collections of Dubois and Selenka (Museum für Naturkunde, Berlin).

Among the deer and *Duboisia*, prime-age individuals had greatly outnumbered those of pre- and post-prime ages, based on preliminary assessments of dentition, and the deaths involved appear to have been catastrophic rather than attritional (M. Hill and L. Todd, pers. comm., 2015). Van den Bergh (1999: 362) concluded that the *Stegodon* individuals from Trinil in the DC appear to have been healthy when they met *“catastrophic death.”* The *Stegodon* specimens from the main bonebed in DC and Museum für Naturkunde Berlin give an MNI of 32 (van den Bergh 1999: 353).

#### Terrestrial fauna

The flood plain of the paleo-river upstream of Trinil must have had sufficient herbaceous open-terrain or forest-understory to support herd-sized populations of ungulates during the months or few years prior to the catastrophic death*. “The high number of large bovids* [in the Trinil fauna] *means a drier biotope”* and *“a more open woodland”* existed in the Trinil paleo-river floodplain than in the Southern Mountains of Java during the Late Pleistocene and Holocene when orangutan lived in a *“humid forest”* of Java and Sumatra (Punung fauna; de Vos 1985a: 216, de Vos 1989, de Vos et al. 1994: 131; also, Aziz et al. 1995).

The paleo-Trinil terrain envisioned was e similar to the modern grassland-forest mosaic in northeast India’s Brahmaputra River lowlands, an area with high annual levels of strongly monsoonal rainfall. The lowlands carry a large population of *Axis* and cattle, water buffalo and elephant herds; the deer inhabit tall- and short-grasslands, wetlands and mixed-deciduous forests, and periodically drown in floods (Endnote I(i)).

Detailed analysis of Trinil large bovids has yet to settle upon their paleoecology (Endnote I(ii)). *Axis*, large bovids, *Panthera,* and *Stegodon* fossils are widespread and long-lasting in the Pleistocene of Java (Table 5), and hence appear to have been sufficiently ecologically flexible to inhabit varying ecological setting over the course of hundreds of thousands of years. Such flexibility is consistent with the wide distributions of the historical cattle, deer and tiger populations in Southeast Asia (Table 3 and Endnote I).

There are specific indications that the *Stegodon-Homo erectus* faunas (*S.-H.e.*) had the adaptability to live under strongly contrasting climates. At Sangiran Dome, paleo-pedological and palynological studies of the vertebrate-fossil-bearing Pucangan and Kabuh Formations reveal shifts in rainfall regimes over geological time; periods of severe dry seasons were identified, as were localized evapotranspiration differences between wetlands and interfluves (Table 6-F to -H). Pleistocene populations of all large mammals found suitable habitats in river valleys of the Mojokerto paleo-delta lowland (represented by the Perning *Homo erectus* bonebed), Ngandong paleo-drainage (Ngandong *Homo erectus* bonebed), Solo paleo-watershed (multiple bonebeds and fossil deposits at Sangiran Dome), and Trinil paleo-river valley (*Pithecanthropus erectus* main bonebed; Tables 5 and 6, Endnote A to H).

Forests must have been present to at least moderate extent in the portion of the Trinil paleo-river valley where the main bonebed ungulates died. Low- but significant-frequencies of several forest-prone ungulates occur in the Trinil fauna. The rhinoceros and pig combined NISP is 122 in the DC. Javan Rhinoceros had wide forest distribution in the historic past, and was present in Java approximately as long as *Homo erectus* was (Table 4, Endnote I(v)). Most historic *Sus* species inhabit forests, and the extinct Trinil Pig (*Sus brachygnathus*) appears to have been part of a long lasting and geographically widespread Southeast Asia complex of lineages (*Sus* spp. probably originated there; e.g., Melleti and Meijaard 2017). The δ^13^C results for Trinil *S. brachygnathus* enamel are consistent with C3 vegetation and omnivorous behaviors normal to *Sus* (Janssen et al. 2016, Jansen 2017). Judging from anatomical and genetic studies, *S. brachygnathus* is most closely related phylogenetically to the present-day Javan Warty Pig of Java and Sumatra, and the Bawean Warty Pig lives on the same forested island in the Java Sea that *Axis* deer do (e.g., Endnote I(v)).

The Trinil *Panthera tigris* supports an inference of woody undergrowth in the forests of the Trinil paleo-river watershed. Holocene tiger populations occupied coastal forests, lowland rainforest and moist-deciduous forests, and montane forests of the greater Indomalaya realm (e.g., Endnote I(v)). Other forest-prone species are represented (muntjac, macaque, porcupine, leopard cat, langur and gibbon; Table 3). Fossils of rat, python and monitor lizard, the modern representatives of which commonly live in covered settings, are components of the Trinil assemblage (Table 3, Endnote I(vi)). Hundreds of small- to medium-sized arboreal and ground-dwelling species, which have not been identified among Trinil fossils, presumably also inhabited the watershed.

#### Plant bioclasts and paleovegetation

Plant macrofossils and palynological data indicate that the Trinil paleo-river lowland was a reed- and grass-dominated arboreally open terrain interspersed with wet forests (e.g., swamp forest and riparian forest) and probably had montane rainforests up drainage. The main bonebed commonly contained bioclasts of wood, leaves and reedy grasses (Cyperaceae), according to firsthand reports (Endnote H(i) to (iv)). The main leaf bed, which had its lowest expression just above the **HK** in Selenka right-bank Pit I, had leaves from lowland (evergreen) rainforests (Endnote H(iv), as discussed extensively by Selenka associates (e.g., Selenka and Blanckenhorn 1911, Schuster 1911a, b, Berkhout and Huffman 2021; also, Flenley 1979, Matthew 1928). Palynological analyses of the Pucangan and Kabuh Formations at Trinil strongly suggest that the paleo-lowland had herbaceous vegetation with forested portions, while the uplands of the drainages had montane rainforest (Endnote H(v)). One palynological sample collected near the **HK** stratigraphic level on the left bank contained more pollen of woody taxa than did samples from the underlying Pucangan and less dryland-tree signal than samples from the overlying Kabuh Formation, leading to a suggested up-section shift toward wetter paleo-landscapes through the period of **HK** accumulation (Polhaupessy 1990, 2002, 2006).

#### Riverine fauna

Because nearly all of the Trinil aquatic species are extant (Table 3), the conditions in the Trinil paleo-river can reliably be inferred from the present-day ecologies of these taxa. Based on its aquatic fauna, therefore, the main bonebed accumulated along a distal lowland segment of a large perennial river which was linked to long-standing lakes and ponds, and had crocodiles, turtle and certain fishes living on its banks during the monsoonal dry-seasons.

*Crocodylus siamensis* (Siamese Crocodile), a key main-bonebed species, was historically widespread from Java, Borneo (Mahakam River) and Indochina (Endnote I(vii)). The last record in Java was from a large freshwater swamp lying at ∼100m elevation, substantially inland of the west coast (Whitten et al. 1996). Modern analogs of the Trinil Testudines indicate that the Trinil paleo-river flood plain had extensive perennial water bodies. Fish closely related to those found at Trinil are distributed widely in Indomalaya rivers. The most-numerous Trinil species have adaptations favoring dry-season survival (Endnote I(vii)).

Mussels and gastropods in the main bonebed further establish the biotic diversity of the Trinil paleo-river (Table 3). The molluscs include species that today inhabit both sizeable perennial rivers and still-water settings, including lakes or freshwater swamps (Joordens et al. 2009, 2015). Joordens et al. (2015: Supplementary Information 2) report dimensional difference in paired valves (MNI of 60) of the mussel *Pseudodon vondembuschianus trinilensis* that might reflect *“several different environmental settings along the* [paleo] *river.”* Some specimens of this mussel species are known to have originated from the main bonebed, including the type specimen (Dubois 1908), but evidently, none of the extant shells which have patterns of breakage that are potentially attributable to hominin action are known to have been originated in the bonebed.

While the Trinil aquatic assemblage came to rest in a lowland portion of the watershed, some elements of the assemblage could have originated from river ways and floodplains tens of kilometers up river of Trinil and several hundred in meters elevation. Mussels analogous to those in the main bonebed occur along the modern Brantas River, Java’s second largest following the Solo (Affandi et al. 2013, 2017). The Brantas is subject to marked seasonal variations in flow volume, and lahar floods emanating from the active Kelud volcano impact the river as far down as its delta at Madura Strait (Figure 1b). One Trinil mussel species (*Elongaria orientalis*) is still widespread in Java’s rivers and commonly occurs around lakes (Whitten et al. 1996). Several viviparid snails in the bonebed today prefer clear, still waters with abundant plants in Java (e.g., around a freshwater swamp in Central Java). Other bonebed gastropods are a species that presently inhabit vegetated stagnant water bodies or freshwater-dominated tidal zones.

The Trinil portion of the paleo-river was not necessarily *“near-coastal”* (Joordens et al. 2009: 664). No purely marine deposits are reported from the upper Pucangan and Kabuh formations around Trinil, and no mangrove indicators are seen in palynological reporting. The ocean outlet could have been tens of kilometers down river from Trinil. An oceanic coast existed ∼20 km across the Kendeng Hills directly north of Trinil (Figure 12), but the deposits filling the marine embayment lack the voluminous volcaniclastics materials that would be implicated by major drainage crossing the Kendeng upland (Figure 13). Brackish-water incursions might have come up the Trinil paleo-river from an estuary, delta or lagoon, since the paleo-river experienced strong seasonal variations in water levels due to temporal changes in precipitation within the drainage.

The Trinil paleo-river floodplain seems to have had segments similar to Cambodia’s Sre Ambel river. About 54km of the Sre Ambel is navigable. The “*floodplain extends 5–7 km on either side of the main river channel and is characterized by wetlands and evergreen riparian forest …. subject to backwater flooding,”* while *“adjacent to the floodplain …* [are] *mixed deciduous forest and open savanna* [and] *all … observations of living crocodiles … occurred at wetlands adjacent to the river”* (Platt et al. 2006: 183).

#### Regional paleogeography

The Pucangan Formation, which underlies the main bonebed, contains voluminous lahar deposits. Diamictic lithofacies are typical of that Formation on the south side of the Kendeng Hills from the greater Trinil area through Kedungbrubus to Mojokerto (Figure 13). Two Pleistocene stratovolcanoes, which were the primary or lone source for the lahars, lay south of the *Pithecanthropus erectus* site in Trinil paleo-river valley. Smaller non-volcanic watersheds were situated within the Kendeng Hills to the north of the valley (Figures 11).

The evidence for the presence of Wilis Pleistocene stratovolcano, located southeast of Trinil, is particularly relevant to the regional paleogeography (Table 6-B to 6-E, Figure 11). Laharic breccia comprise most of a 275m-thickness of the Pucangan Formation near Kedungbrubus, where the Pucangan and Kabuh combined have 765m of volcaniclastic deposits (Table 6-D). The exposed relationships make for an open-air cross-section of the cone-shaped northern flank of the immense Pleistocene paleo-Wilis stratovolcano (Figure 13). Volcaniclastic sands and lahars originating from the center evidently spanned 150km east-west and reached both Trinil and Mojokerto. The west-flowing drainage apparently was part of the Trinil paleo-watershed (Figure 12).

The lithofacies of the east-directed drainage emanating from paleo-Wilis is on display in the eastern Kendeng Hills where volcaniclastic non-marine facies of the Pucangan Formation transition eastward and northward across the Mojokerto paleo-delta to marine mudstones (Figure 13). The deltaic deposits contain the relocated discovery bed of the Mojokerto *Homo erectus* child-skull fossil (Table 6-E). The pollen spectra and grass-phytolith assemblages in the stratigraphic section enclosing the child-skull bed indicate that dry grasslands were abundant in the Mojokerto paleo-river valley at the time, along with mangroves, swamps, delta- and riparian-forests and distant montane vegetation (Table 6-E). The grasslands appear to reflect an aridity similar to modern conditions hundreds of kilometers to the east in one of the driest parts of Indonesia, Nusa Tenggara (Morley et al. 2020).

Despite the dry paleo-hydroclimate of the Mojokerto paleo-delta, the large mammal fossil fauna from the vicinity (e.g., the Perning and Jetis districts) is similar in taxonomic expression to the Kedung Brubus fauna and thus has elements in common with the Trinil fauna (Tables 2, 4 and 6-E). From Mojokerto westward to Sangiran Dome, the upper Pucangan and Kabuh Formations regularly have fossil crocodilians, turtles and fresh-water mollusks, indicating that perennial rivers often existed at the core of the Pleistocene stratovolcanic watersheds in which *Homo erectus* lived. Ongoing flow in the rivers presumably was sustained by runoff from the volcanic highlands (Figure 11). Keystone resources in the lowlands near these rivers might have contributed to the long-term persistence of *Axis*, *Homo,* large bovids, *Panthera*, *Rhinoceros* and *Stegodon* regionally (Table 5).

#### A central challenge regarding the main bonebed

A central issue in assessing the origin of the main bonebed is explaining how thousands of large, disarticulated and little-abraded vertebrate bioclasts and numerous other biotic remains became concentrated (with certain internal irregularities) within a thin, localized, poorly sorted gravelly volcaniclastic sand along a lowland section of a large perennial river. The biofacies and lithofacies, as evaluated above, lead to a plausible answer.

Several characteristics of the biofacies are particularly significant. When bonebeds are considered worldwide, they are seen to *“form through complex combination of biotic and abiotic mechanisms”* and *“multitaxic bonebeds are frequently a source of paleocommunity data”* (Brinkman et al. 2007: 221, 223), as is the case at Trinil. Leverage is enhanced further when: (i) evidence of *“hypothetical standing population”* comes from the *“catastrophic mortality of a gregarious group”* (Eberth et al. 2007b: 283, 290), as is true of the main bonebed assemblage; (ii) three- or more-taxa dominate in near-equal shares of the mass-death population, such as Trinil’s *Bibos*, *Bubalus* and *Axis* did; (iii) *“paraphyletic”* aquatic- and terrestrial-components in close association contribute to interpreting causality (Eberth et al. 2007a: 104), which is true with the aquatic- and forest-prone animals in the main bonebed; and (iv) key taxa are *“isotaphonomic”* and have *“similar general taphonomic histories”* (Blob and Badgley 2007: 341), as is seen in taphonomic analysis of main bonebed ungulate species.

On the other hand, the Trinil main bonebed departs significantly from global norms for fluvial bonebeds. First, the Trinil bonebed is an unusual lowland-river deposit in having so many disarticulated vertebrate elements that also exhibit fine surface preservation. Exposure of carcasses tends to weather bones substantially, and rivers generally damage bioclasts and disperse them along their courses, rather than preserve and concentrate them locally (Behrensmeyer 1991, 2007; J. Rogers, pers. comm. 2018), as clearly happened in forming the main bonebed.

Second, there is no evidence that the main bonebed fossils were reworked from an older sedimentary formation, which distinguishes Trinil globally from most bonebeds with multi-individual multi-dominant bioclast concentrations. Third, the bioclasts of terrestrial species in the main bonebed at Trinil do not exhibit typical size- or shape-sorting, nor have strong evidence of fluvial abrasion, normal for fluvial bonebeds deposited in a paleo-drainage lowland. Fourth, even though it was deposited in the lower reaches of the drainage, the Trinil bonebed would not be classified with many others as either a normal *“channel-fill”* or *“channel-lag”* concentration (Rogers and Kidwell 2007: 6-7), because the bioclasts in these types of deposits have variable abrasion levels, and are shape- and size-sorted by their selective movement in river currents (Behrensmeyer 1991, 2007, Blob and Badgley 2007, Eberth et al. 2007a,b, Lyman 1994).

Fifth, lahar-flood transport, as Dubois and Selenka’s geologists thought occurred in the formation of the Trinil bonebed, is rare globally in several statistical senses. Bonebeds caused by flooding and drowning were only 18% of 185 bonebed sites studied in one worldwide sample (all outside Indonesia) and the closest volcanic categories, *“debris flows”* and *“ash falls,”* are just 5% of the cases (and lahar flood cases are not mentioned; Behrensmeyer 2007: 84). Finally, Rogers and Kidwell (2007: 24), while not specifically considering the Trinil main bonebed in their global review of bonebeds, *“find it conceptually difficult to accept … that disarticulated bones and teeth of numerous animals delivered from widely separated point sources at different times would travel downstream … and collectively accumulate.”*

#### Origin of the main bonebed

Our proposals for the origin of the main bonebed avoid these general conceptual difficulties by marshalling evidence favoring simultaneous death of hundreds of ungulates in one section of the Trinil paleo-river drainage (specifically, the flood zone where the animals had concentrated) and *en masse* transport of the skeletonized, little-weathered remains by lahar flooding, which resulted in little bioclast-damage and -sorting (Figure 11).

Ungulate populations in the Trinil hinterland doubtless inhabited wide areas across the flanks and lowlands of the stratovolcanoes (see Butak bonebed, Table 6-D). Large eruptions occasionally affected the paleo-drainages, as is typical for stratovolcanic terranes. Periods of intense eruption or drought would be capable of rendering uninhabitable large portions of the landscape surrounding an active volcano in the Trinil paleo-watershed, and severe resource limitation could force hundreds of ungulates towards areas with plentiful water and forage.

One such refuge of *Axis*, cattle, water buffalo, *Stegodon*, *Duboisia*, *Sus* and *Rhinoceros* (Table 1 and 2) formed in the floodplain of the trunk Trinil paleo-river or lowland tributary (Figure 12a). Tigers and a few dogs presumably followed the ungulates into the refuge, as perhaps did hominin groups. Grass and forbs were sufficiently widespread there to sustain the deer herd. Reed grounds or the woody forest undergrowth presumably offered safe harbor. The Trinil deer were in the riverine refuge long enough for many males to shed antlers.

The forest seemingly had fewer ground-living and arboreal animals than did tree-covered areas elsewhere in the watershed. Denser forests were likely to have been widespread around the hilly areas of the stratovolcanoes and in the Kendeng Hills and Southern Mountains (Figure 1b). The water courses passing through the refuge had the same diverse suite of riverine reptiles, fishes and molluscs as existed in other perennial tributaries and standing water bodies of the watershed.

Catastrophic mortality decimated the ungulates in the refuge. The individuals were largely in the prime of life. Their deaths most plausibly resulted from particularly devasting volcanic events (ash falls, pyroclastic-surge eruptions or lahars) or intense drought. The kill-off might have come from a worsening of the conditions that caused the ungulates to flee into the refuge, or an independent calamitous event. Hundreds of ungulate individuals could have perished over hours to weeks (particularly in the case of volcanism) or weeks to months (if drought led to the deaths). Perhaps, the animal aggregation and mass mortality might have been far up river and the cadavers transported part way down stream by flooding to a location where the bodies deteriorated.

Almost all of the carcasses decayed to the point of skeletonization, creating a scatter of the tens-of-thousands of bony elements on the flood plain. Exposure was too short for the bones to weather to ruin. Bioerosion did not prominently affect them. The bony elements might have been broken by the trampling of the large ungulates that survived. Avian scavengers and other consumers of carcasses left no record of substantial contribution to the skeletonization. Tiger, dog and hominin populations played little-if-any role in the death of the ungulates. Ash falls possibly inhibited scavenging and preserved bone surfaces. The hominin population in Trinil paleo-lowland used lithic tools so sparingly that no flakes or artifacts came to be embedded with the voluminous amounts of granules and pebbles excavated from the bonebed.

A lahar flood inundated the paleo-river floodplain, sweeping up thousands of decomposed skeletal remains. When the waters surged through, the flood had sufficient hydrodynamic competence to suspend and carry nearly whole *Stegodon* and large-bovid crania. The bone field might have been situated a kilometer from Trinil or several tens of kilometers up river, but the lahar flood water largely originated in the volcanic uplands, as flooding in the region normally does today (Figure 11c). Most of the sand and gravel in the lahar flood had entered the flood before it reached the skeletal scatter. The lithic materials gave the flood waters the density sufficient to move the largest skeletal pieces and lithic boulders downriver. As the bone-rich flood proceeded, the bones were not shape- and size-sorted because the surge moved in hyperconcentrated flow regimes with little interruption. The flooding incorporated logs, reeds and leaves, and river-living reptiles, fishes and molluscs from the paleo-river floodplain or other points along the drainage.

Multiple sectors of the watershed might have contributed water to the flood, but they did not furnish substantial numbers of vertebrate bioclasts with taphonomic histories notably different than those in the principal area of bone scatter. The lahar flooding presumably completed the disarticulation of bony sections of remains which had not been fully disjoined during exposure. A few skeleton elements stayed together and retained vestiges of soft tissues. Crocodiles left impressions on some bones, perhaps anticipating access to fleshy tissues.

Stratovolcanic landscapes in Java offer multiple opportunities for lahar flooding. In the main bonebed case, a crater lake at the center of an active volcano might have emptied catastrophically; a sector of the volcano might have collapsed into a debris-flow which evolved into a lahar flood after dilution by discharge from other tributaries of the watershed; or a part of the highlands might have been struck so heavily by rainfall during a wet-season storm that the lahar flood arose when rivers bulked up with sand and fine gravel.

The river bottom at Trinil had been lined with dark-colored mud and gastropod banks before the lahar flooding arrived. Even earlier, the river carved older consolidated diamicton into bedrock edges of the channel. The flood waters might have backed up behind bedrock constrictions, inducing accumulation of the main bonebed in a matter of hours. Bed-load traction movement deposited pebbly sand with large-scale crossbedding as it passed the present-day left bank. Over the course of hours or days, internal streams and pulses segregated bioclasts and lithic materials sufficiently to create internal lithic and bioclastic facies within the main bonebed in toto.

A pulse carrying the *Pithecanthropus erectus* (*P.e.*) materials and other large-mammal bioclasts followed an initial surge which carried fewer bioclasts and more cobbly gravel. The Skullcap and Femur I arrived at Trinil shortly thereafter with hundreds of Trinil fauna bones. The hominin femora from the 1900 excavation might have accumulated at Trinil within minutes, hours or several days earlier or later than the original *P.e.* remains.

The main-bonebed flood also might have deposited bioclast concentrations down stream of Trinil, but more likely, as the flooding continued, the high-rate flow dispersed skeletal fragments along the trunk river and deposited them in low densities along various sections of its floodplain, estuary, delta or immediate offshore. High-rate river flows potentially continued to pass Trinil for days or several months after the main bioclast-rich flood had surged down valley, due to continuing rain and landscape disturbance in the watershed.

Several meters of sandy volcaniclastic deposits containing vertebrate- and molluscan-bioclasts which represent the same perimortem events as those expressed in the main bonebed might have accumulated on top of it. Although differing geological age from the main bonebed, the Ngandong *Homo erectus* bonebed is a bony concentration in the lower portion of the ∼3m thick terrace deposit that formed by ongoing lahar flooding (Tables 5 and 6-B).

Fluvial deposition continued for millennia after the main bonebed formed, leading to the accumulation of the overlying Kabuh Formation. Occasionally the remains of a few individuals were embedded, most notably in the *Stegodon* bonebed ∼5m above the main bonebed. Few bony materials were otherwise embedded, since mortality in the paleo-drainage almost always decomposed large-mammal remains before they entered the river systems, and the surviving bones were reworked repeatedly by periods of exposure and burial.

Uplift of the Kendeng Hills tilted the Kabuh and Pucangan Formation gently southward, but in the immediate vicinity of Trinil, the strata were spared the regional monoclinal movement by a structural anomaly. Long afterwards, the Solo River formed its present main valley and became Java’s largest river (Figures 1b and 11c). The Solo River ultimately flowed in a strongly meandering course and then incised. At the left-bank *Pithecanthropus erectus* site, the incision penetrated in excess of eight-to-nine meters of the Kabuh Formation, exposing the **LB**.

More than a century ago, Dubois and Carthaus contextualized *Pithecanthropus erectus* by analyzing the paleontological and sedimentological features and paleogeographic setting of the main bonebed. As we address in DISCUSSION, next, this same approach helps to elucidate the conditions that sustained *Homo erectus* in eastern Java for around a million years, and concentrated hominin-fossils in this one portion of the southern Sundaland.

## DISCUSSION

Eugène Dubois considered *Pithecanthropus erectus* and associated Trinil fossils in broad paleobiogeographic terms. Here, we stress roles that the main bonebed plays in *Homo erectus* paleogeography of southern Sundaland (Java, Java Sea portion of the Sunda Shelf, southern Sumatra and southern Borneo). Tables 5 and 6 have briefs on key hominin-fossil bonebeds and sites in eastern Java. Figures 13 and 14 and S I Figures 22 to 24, present paleogeographic context.

Volcaniclastic materials form the discovery deposits at the consequential hominin-fossil sites of Ngandong, Kedungbrubus, Mojokerto, Sangiran Dome, and Trinil, and indeed dominate the lithofacies of the hominin-fossil bearing formations. All *Homo ere*ctus fossils so-far discovered in the region were embedded in paleo-watersheds that drained high-standing stratovolcanoes, much like the circumstances in which *P. ere*ctus originated (Huffman 2017).

The large-mammal species in the main bonebed defines the Trinil fauna, and it anchors the *Stegodon-Homo erectus* faunal association (Tables 3, 5 and 6). The *S.-H.e.* embodies a paleobiogeographic link between *Homo erectus* and certain lineages of large bovids, cervids, proboscideans rhinoceros, suids, and tiger (de Vos 1995b). *Axis lydekkeri*, *Panthera tigris* (or *P.* sp.) and *Stegodon trigonocephalus* were present during the whole period of known *H. erectus* occupation. So were inhabitants of large river systems (*Crocodylus*, Testudines and molluscs). Both terrestrial and aquatic fossils occur widely in the *H. erectus*-bearing formations (Table 5), placing the faunas in the same volcanic watersheds as the hominins. Seemingly, *H. erectus* populations were as omnipresent as those of the most-persistent other faunal lineages.

Radio-isotopically dating at Ngandong and Sangiran Dome provides a geochronological framework for the life and death of *Homo erectus* in the paleo-watersheds of stratovolcanoes. The Ngandong *Homo erectus* bonebed is early Late Pleistocene in age, modelled to be ∼0.1 Ma (117 to 108 ka; Table 6-B; the Pleistocene sub-epochs are used in the sense shown in S I Figure 22a). The oldest-known hominin fossils at Sangiran Dome are late Early Pleistocene; that is, ≥0.9 Ma and more likely <1.3 Ma than <1.5 Ma (Table 6-G). Thus, archaic hominin populations inhabited eastern Java for >0.8 Ma and perhaps ∼1.4 Ma.

Comparing Ngandong to the youngest Sangiran *Homo erectus* leaves an apparent lacunae of ∼0.8 Ma. Radio-isotopic dating at Trinil is taken by some to indicate that the main bonebed is ∼0.5 Ma (Joordens et al. 2015) or even ∼0.1 Ma (about the same age as the Ngandong terrace deposit in Berghuis et al. 2021). A more established logic is based on radiometric results and biostratigraphic data from Sangiran Dome. The large-mammal assemblage in the Grenzbank *H. erectus* bonebed at the Dome represents the Trinil fauna (Tables 5 and 6-G). Since the Grenzbank is >0.9 Ma, the Trinil fauna evidently was present in eastern Java by the end of the Early Pleistocene (Table 6-G). This inference is supported by the ∼0.8 Ma age of Ngebung hominin bonebed, which overlies the Grenzbank and has the late Trinil fauna (Table 6-H). The Trinil fauna is older in biostratigraphic terms than the Ngandong fauna (Tables 5 and 6).

Another paleontological inference arises from comparing the Grenzbank and Trinil bonebeds. Terrestrial vertebrate fossils in the former are reworked bioclasts and produced over a protracted period of geological time, in contrast to relatively short-term taphonomic development of the Trinil main bonebed (Table 6-A and 6-G). The faunal similarity between the two bonebeds, one having reworked bioclasts and the other not, suggests that the quicker-formed Trinil main bonebed, and also the palimpsest-like Grenzbank, had fossils of most of the large mammals present then in stratovolcanic watersheds of eastern Java, if not across the region’s non-volcanic uplands as well (e.g., Figure 14).

Even though the Ngandong and Trinil bonebed are distinguishable on the basis of their large-mammal faunas and have different geological ages (Tables 2 and 5), the two bonebeds had similar taphonomic and sedimentological histories. Both bonebeds formed after (i) an aggregation of ungulates in a floodplain of a major stratovolcanic drainage, (ii) decimation of populations, (iii) skeletonization of remains without severe bone destruction, (iv) transport of tens of thousands of bones downriver by flood, and (v) local concentration of skeletal materials in a river (Table 6-A and 6-B, Figures 11 and 12; Huffman et al. 2010a, 2012b). Each set of events took place over a matter of a few years, and both terrestrial-fossil assemblages appear to closely approximate the large-mammal fauna present in the respective stratovolcanic paleo-watersheds (Figure 11).

The stratovolcanic paleo-drainages in which both the Ngandong and Trinil bonebeds formed were dominated by the same large-scale topographic features; that is, Pleistocene versions of the Kendeng Hills, Lawu volcano and Wilis volcano (Figures 1b, 11 to 13). Paleo-Wilis was a massive- and long-lasting stratovolcano. Geological evidence of this is clear from its northern flank in the Kedungbrubus-Butak area where hundreds of meters of gravelly laharic diamicton accumulated in the Pucangan Formation (Table 6-C and 6-D, Figure 13, S I Figure 21). Large-mammals are evident early in the era of volcanism and lahar flows there (Table 6-D).

The watersheds on the west side of paleo-Wilis fed the Solo River where the Ngandong bonebed accumulated ∼0.1 Ma (Figure 11c). Discharge from paleo-Wilis headwaters might have previously drained past Trinil (Figure 12). The watershed on the east of paleo-Wilis clearly fed the Mojokerto paleo-delta where the Mojokerto *Homo erectus* child skull was deposited near a deltaic shoreline (Table 6-E, Figure 13). Viewed broadly, the Mojokerto, Ngandong, Sangiran Dome and Trinil discovery sites represent different proximal-to-distal positions along Pleistocene stratovolcanic drainages (Figure 14).

There were profound temporal environmental and geographic changes during the *Homo erectus* period of occupation in eastern Java. The youngest-documented hominin-fossil at Sangiran Dome is ∼0.8 Ma (Table 6-H), making the minimum span of inhabitation in the paleo-Solo Basin >0.5 Ma. This period of inhabitation corresponds approximately to the global Mid-Pleistocene Transition (MPT). Episodes of glacio-eustatic change intensified during the MPT compared to older Pleistocene patterns (S I Figure 22a). The first hominins in the Sangiran area probably arrived before the MPT or during its earliest phases, when the large-mammal fauna of Java (Ci Saat fauna) had less taxonomic diversity than the Trinil fauna (Tables 5 and 6-F). Periods of lower global levels during the MPT doubtless created vase land areas between Java and mainland Asia (Sunda Shelf), and afforded the Trinil fauna terrestrial lineages greater access to southern Sundaland (S I Figure 22a and 23b).

Little is known about the impact that glacio-eustatic fluctuations had on the populations of *Homo erectus* and associated *S.*-*H.e*. species, but the youngest well-dated *S.*-*H.e*. deposit, the Ngandong *Homo erectus* bonebed, provides a useful reference point. Ngandong *H. erectus* and associated Ngandong fauna flourished in the Javan interior ∼113 ka (Table 6-B). Ten- to fifteen-thousand years earlier, ancestor populations evidently lived in relative geographic isolation during a global sea-level high-stand of the last interglacial period (LIP, Marine Isotopic Stage, MIS, 5e). During the preceding penultimate glacial period (PGP, MIS 6; S I Figures 22a and 23), older *S.*- *H.e*. populations would have had the opportunity for expansion across the Sunda Shelf.

Thus, late in their occupation of Java, the *Homo erectus* and certain non-hominin *S.*-*H.e.* lineages adjusted successfully to profound glacio-eustatic changes in landscape and climate. Perhaps the replacement of the Kedung Brubus fauna by the Ngandong fauna (Table 5) reflects the extreme paleoenvironmental conditions of the LIP (highstand) or PGP (lowstand), but Middle Pleistocene sea-level and paleoenvironmental fluctuations before the PGP might also have led to the faunal changes (S I Figure 22a and 23b; also, S I Figure 24). The continuity of large mammal lineages in the *S.*-*H.e.* (Table 5), *H. erectus* included, seems to indicate that glacio-eustatic fluctuations neither displaced the *S.*-*H.e.*-mammal populations from southern Sundaland, nor led to hominin speciation there (S I Figure 23a).

The capacity *Homo erectus* and other long-lasting *S.-H.e.* species had to persist through a range of hydroclimates, including glacio-eustatic extremes, is evident in paleoclimatic information from the key fossil sites in eastern Java. The Mojokerto-child *H. erectus* population appears from paleobotanical information to have lived in a drier climate than was present in the Trinil paleo-river valley during the time of *P. erectus* (Table 6-E). Paleo-pedological and palynological studies of the *H. erectus*-section at Sangiran Dome reveals fine-grained hydro-climatic and vegetation variations, some involving severely dry conditions (Table 6-F to 6-H).

Recent climatic patterns suggest a potential mechanism underlying the faunal continuity. Historic Java varied from dry-monsoonal to everwet-climate from east to west, and also from south-to north-coasts, lowlands to mountains, and watershed to watershed (S I Figure 23a). When the *S.-H.e.* lineages inhabited Java. there presumably were similar geographic variations in climate, and associated vegetation- and mammalian-biotopes. Quite plausibly therefore *H. erectus* persisted in southern Sundaland because of its capacity move from one lowland-, coastal- or montane- biotope to another as they shifted across the region over time (Figures 13 and 14; Huffman 1999a,b, 2001a). Habitat flexibility in *H. erectus* is reasonably supposed to have played a critical role in its >0.8 Ma occupation of southern Sundaland.

The *S.*-*H.e.* failed to survive into the second half of the Late Pleistocene. A faunal turnover is evident from teeth of *Homo sapiens* and mountain-forest vertebrate species in the Sumatran cave that Dubois discovered (Table 5, note 5). Recently, fossil-bearing breccia at his Lida Ajer cave has been dated to 63-73 ka (Westaway et al. 2017). This places a fauna with *H. sapiens* in a mountainous peripheral sector of Sundaland during MIS 4 (71-50ka). MIS 4 included a pre-PGP episode of very low-sea-level (e.g., de Deckker et al. 2019, Schneider et al. 2013), when the modern humans might have dispersed widely across the Sunda Shelf and Sunda islands, and replaced all *H. erectus* populations (S I Figure 23b). *H. sapiens* teeth are also reported to be among the 128 ka +/- 15 ka rain-forest faunal remains recovered from Punung rock shelter in the Southern Mountains of eastern Java (Table 6-J). If so, *H. sapiens* took over some sectors of southern Sundaland, such as these Mountains, while for millennia afterwards *H. erectus* and other *S.*-*H.e.* large mammals inhabited other sectors.

*S.-H.e*. occurrence outside of the stratovolcanic watersheds of mid-island eastern Java supports the inference that *Homo erectus* ranged broadly across in southern Sundaland. Hominin fossils occur with abundant remains of *S.-H.e.* species ∼100km north of Trinil at Patiayam (Table 6-I, Figure 1b). This collection area, which was on the flank of an Early to Middle Pleistocene stratovolcanic island, lay across the Kendeng Hills and Randublatung marine embayment from the middle of the Java area of *H. erectus* discovery (Figure 13). *S.-H.e.* mammals evidently crossed the seaways that separated paleo-island from the Kendeng Hills and Rembang Hills. Archaic hominins reached into the Southern Mountains, where artefacts at the Song Terus cave record a presence during the Middle Pleistocene (Table 6-J).

The lack of *Homo erectus* skeletal fossils in the Southern Mountains, Rembang Hills and Kendeng Hills (except for Kedung Brubus and Ngandong terrace deposit) should not be taken to mean that Pleistocene populations were absent across the broad extent of these uplands, just that destructive taphonomic conditions generally prevailed in them (Figure 14). Erosional paleo-landscapes in these uplands apparently accumulated too little volcaniclastic material to preserve bony remains (Figure 13; Huffman et al. 2012b). The skeletal materials of hominins and other large-mammals might have been destroyed systematically on the ground surface or in corrosive subsurface settings in these mountainous terranes.

The Patiayam and Song Terus localities are not the only sites that indicate broad paleogeographic distribution of *S.-H.e.* species. Several sites west of Sangiran Dome and Patiayam in western Java have archaic hominin skeletal or dental remains (S I Figure 22b). The type area for the oldest fauna of the *S.-H.e.*, the Ci Saat fauna (Table 5), is in Central Java, ∼200km west of Sangiran. The western-most known *S.-H.e.* occurrence is a Trinil fauna pig jawbone in a non-marine sequence with marine intervals recovered by coring near the Java Sea coast at Jakarta, ∼450km west of Patiayam (S I Figure 22, Marks 1956, Yulianto et al., date unknown). Indirect biological evidence suggests that many *S.-H.e.* species, such as its bovids, cervids, suids, rhinoceros and tiger, had sufficient ecological flexibility to attain wide distribution in southern Sundaland.

The potential for *S.-H.e.* dispersal north of Java also is evident from the paleo-landscapes that sub-bottom seismic data show beneath the present-day Java Sea. Modelling Pleistocene paleogeography (e.g., Salles et al. 2021) benefit critically from close attention to marine-geophysical resources (Alqahtani et al. 2015, Darmadi et al. 2007, Huffman et al. 2012a, 2013, 2018). For example, directly north of the eastern Java *Homo erectus* discovery area (Figure 1b) seismic profiles reveal widespread Pleistocene paleo-landscape features (S I Figure 24a), as do ‘3-D’ data volumes in both the westernmost and easternmost Java Sea (S I 24b-d). The data demonstrate immense river-valley systems developed across the Sunda Shelf during multiple periods of the Pleistocene. Valley systems are clearest seismically for low-stand episodes when the Shelf terrane was the largest. However, marine beds in Java attest to the likelihood that portions of Sunda Shelf continued to be inundated during most of the Early and Middle Pleistocene.

The 3-D data, in particular, allow for spatiotemporal (four-dimensional) analysis of the paleogeomorphology, greatly increasing confidence in environmental interpretation of the landscapes that formed under lower-than-present sea-level conditions (S I Figure 23). In Central Java, many north-draining Pleistocene watersheds fed directly into the low-sea level river valleys of the Sunda Shelf, and some onshore watersheds have *S.-H.e.* fossils (S I Figures 22b and 23). Large-mammal populations potentially expanded down the former valleys and interfluves during periods of depressed sea level and contracted back into the Javan core as sea level rose.

In the western portion of Java, Pleistocene highlands drained into the Sunda paleo-watershed that emptied into the Indian Ocean via Sunda Strait. The northern headwaters of this watershed abutted low-sea-level drainage divides and headwaters now under the western Java Sea between Sumatra, west Borneo and the Malay Peninsula (S I Figure 23). During lowered sea level, these Pleistocene territories would have given *S.-H.e.* populations pathways into the Sunda uplands of Borneo, Sumatra and Malaysia, and account for the arrival of new Asian mainland lineages in the Trinil fauna and Kedung Brubus fauna (Table 5), among other taxa.

At the same time, hominin and other terrestrial mammals might well have lived in peripheral parts of the Sunda Shelf in southeastern-most Sundaland. For instance, the stratovolcanic lowland which produced the Perning *H. erectus* continued for ∼250km eastward to the end of Java. The Pleistocene upland of the Rembang Hills extended for hundreds of kilometers through Madura Island towards the Kangean archipelago at a southeastern corner of Sundaland. Between Madura and Kangean islands, Pleistocene rivers, which had flowed for hundreds of kilometers across the Sunda Shelf, found exit into the deep-water Bali-Flores Sea or areas north of the Kangean archipelago (S I Figure 24d). At times, such as when sea-level lay off the continental shelf edge, large braided-river systems carried voluminous clastic materials for hundreds of kilometers from headwaters in central Borneo across the Sunda lowlands towards the coasts that lay on the eastern edge of the continental shelf (S I Figure 24c).

In sum, *Pithecanthropus erectus* of Trinil, and other archaic hominin occurrences in Java, are profitably viewed as samples of hominin populations that broadly inhabited southern Sundaland. The Trinil fauna anchors the *Stegodon-Homo erectus* large-mammal faunal association (*S.-H.e.*), which often occurs at *Homo erectus* fossil sites in eastern Java (Table 5, Figures 1b), and links *S.- H.e.* paleogeographically to watersheds of stratovolcanoes (Figure 11). Radiometric dating at Sangiran Dome and Ngandong establishes hominin residency in this setting from 0.9 Ma to 0.1 Ma, if not over a longer geological time span. Glacio-eustatic fluctuations of sea level and climate, which were prominent then, presumably impacted the distribution of suitable hominin habitats in Java and the Sunda Shelf, and led to interchange of terrestrial biota with other Sunda islands and mainland Asia (S I Figures 23 and 24). *H. erectus* survived the harshest apparent conditions until the Late Pleistocene.

Eugène Dubois ventured to Sumatra and Java thinking of paleo-biogeography in expansive ways. Once in the Indies, he applied geology and paleontology to rock outcrops and caves in search of fossil specimens of early ancestor species. The foregoing discussion illustrates that the premier product of Dubois’ efforts, the discovery of *Pithecanthropus erectus* at Trinil, continues to offer avenues for explication of regional Pleistocene paleobiogeography, when Trinil is viewed in conjunction with archaic hominin discoveries that followed Dubois’.

## SUMMARY CONCLUSIONS

Archival materials, museum specimens and associated literature persuasively support Eugène Dubois’ reporting that the Skullcap, Femur I and the vast majority of non-hominin fossils excavated in 1891-1900 along the left bank of the Solo River at Trinil came from a main bonebed (Tables 1 to 3, Figure 2 and 10). These sources of information further indicate that deposition of the Trinil bonebed followed a mass death of ungulates in a paleo-watershed of a stratovolcano (Table 4, Figures 11 and 12). Trinil is representative of all archaic hominin fossils discovered in eastern Java, given that they also were embedded in the sediments of Pleistocene stratovolcanic watersheds (Table 5, Figures 1b, 13 and 14).

When supervisors G. Kriele and A. de Winter (KdW) wrote to Dubois on September 7, 1892, that Femur I came from “approximately the same depth [elevation]” as the Skullcap and other vertebrate bioclasts unearthed in 1891 (S II-A2k), they were well positioned to assess the discovery provenience (e.g., SII-A2g, -B3b), and knew that the strata under excavation were flat lying (Figure 3b, S II-A1b, -B2d). The discovery Trench was large enough by November 9, 1892, to allow them to follow the discovery subunit (**PFZ**) from the immediate vicinity of the Skullcap Pit to the Femur I discovery spot and beyond (Figure 3a; S II-A2q). Then, field crew twice exposed the **LB**/ **PFZ** at the base of the high-standing backwalls in the 25-m and 40-m Trenches (Figure 3c), obtaining confirmation of the stratigraphic context the Skullcap, Femur I and other fossils.

This sedimentary sequence in a remnant of the 1893 backwall was photographed from across the Solo in 1894 (Figures 4 and 10). The context was so familiar to Dubois that he later marked discovery points on prints of the image (Figures 3c and 4a). When Dubois published that the Skullcap and Femur I came from a thin bioclast-rich Lapilli Bed (**LB**; Figures 2a and 10), this attribution was backed up by numerous recorded observations of his field supervisors and his own. The stratigraphy of the 1891-1893 excavations had fortuitously been straightforward. The **LB** bonebed was strongly lithified and near-horizontal, and underlay eight-to-nine meters of indurated near-horizontal strata. At the top of the excavation section, a soil had developed along a largely erosional the river-terrace upland (Figures 2a and 10).

Contemporaneous reporting also substantiates the stratigraphic association of the Trinil fauna with *Pithecanthropus erectus* (Tables 1 to 3). Besides the *Pithecanthropus erectus* Skullcap, the Skullcap Pit (Figures 2b and 3a) also produced large bioclasts now attributable to *Axis lydekkeri*, *Bubalus palaeokerabau*, *Duboisia santeng*, *Stegodon trigonocephalus*, Testudine, and “tree trunks,” together with freshwater mussel shells and “leaf imprints” (Endnote A).

Femur I originated from a portion of the 25-m Trench in which the **LB** yielded numerous *A. lydekkeri* fossils, and specimens referable to *Bibos palaesondaicus*, *Crocodylus siamensis, D. santeng* and *S. trigonocephalus* (Endnote B). The **LB** in the 40-m-Trench, which was dug beside the 25-m Trench, contained *A. lydekkeri*, *C. siamensis*, *D. santeng*, *S. trigonocephalus*, shells and wood (Endnote C).

In the excavations of 1895-1900, large skeletal remains of the same ungulate taxa occurred near the seasonal low-water level (Endnote D). The new pits and trenches, portions of which were adjacent to those of 1891-1893 (Figures 2 to 4), penetrated the same sequence that was encountered during the earlier years. The strata dug in 1895-1900 and we term units **2** to **5** correspond to Dubois’ 1895 ‘sand rock’ formation (Figures 2b, 4, 5 and 10).

Firsthand reporting and 1907 photographs of the Selenka Expedition confirm the persistence of the main bonebed (their ‘Hauptknochenschicht,’ **HK**) near the seasonal low-river level, while at higher elevations in their excavation (Pit II), they encountered the same eight-to-nine meters of hardened strata that Dubois’ crews had (Figures 7 and 8). In total roughly 2200 m^2^ of the Trinil left-bank had been removed during 1891-1908 to mine the *Pithecanthropus erectus* bonebed near the low-water level of the Solo River (S II-A4r).

A 1926 site photograph substantiates the indurated condition of the main bonebed and the superjacent beds (Figure 9). The 1894 photograph can be overlaid directly on the younger one. The superposition relates the *Pithecanthropus erectus* discovery context, as Dubois had noted it on the older photograph, to the 1926 setting. Seen then were the scars and baulks of the bonebed which spoils had covered when the 1900 and 1907-1908 excavations were underway.

Remnants of unit **6**, which formed the topmost stratigraphic unit in the western parts of Dubois’ 1900 Trench and Selenka Pit II, was visible in 1926. The unit is not evident at **LB** level close to the 1891-1893 excavation area. For decades after 1926, geologists mapped the remnants on the left bank as bedrock. Since 1936, it has been assigned to the Kabuh Formation (Figures 6d and 11; S I Figures 16 to 19). Erosion-resistant remnants of the main bonebed occur today along the south side of the Solo at Trinil, and some-or-all of units **2**-**6** must be beneath soil and vegetation along the left bank.

Besides substantiating the site stratigraphy and taxonomic content of the main bonebed, Dubois’ and Selenka geologists’ descriptions and photographs offer indispensable observations on the taphonomic and sedimentary circumstances surrounding the origin of the main bonebed. As the contemporaneous sources portray, thousands of large, disarticulated and irregularly distributed bioclasts of terrestrial vertebrate species were concentrated in a thin, poorly sorted, gravelly volcaniclastic stratum, which also contain the remains of freshwater molluscs and reptile. Most of the bioclasts evidently were matrix supported, with the longest bioclasts laid out parallel to the bedding and the biggest bioclasts occupying the whole main bonebed. No clear evidence of a long-term cessation of fluvial accumulation was reported.

The vertebrate- and plant-fossils ranged in size from small teeth and leaves to large craniums and logs, as is indicated by both original descriptions and museum collections. While the skeletal specimens were commonly broken, levels of abrasion attributable to fluvial transport were unwaveringly low. Fine surface preservation and uniform fossilization of bony materials, including Femur I, add to a picture of an isotaphonomic assemblage of terrestrial remains. Because the bioclast density varied from place-to-place and vertically within the main bonebed, this isotaphonomic assemblage accumulated during multiple surges of river flow, which followed a set of taphonomic events that the remains experienced in common.

Dubois’ and Selenka geologists’ further analyses of main bonebed origin were grounded in analogies with historic lahar deposition surrounding active volcanoes in Java. The men rightly focused on long-run out lahar flows as a mechanism for transport and accumulation in the case of the main bonebed. The large ungulates and other animals and plants had been living in Pleistocene stratovolcanic watershed up river of Trinil when they were decimated catastrophically.

After the ungulate carcasses had been skeletonized, flooding transported the remains to the Trinil discovery site (Table 6-A, Figures 10 and 11, Endnotes A to F). As Dubois and Carthaus suspected a century ago and is evident now in museum collections, formation of the main bonebed involved the penecontemporaneous deaths of more than one hundred ungulate individuals (Table 4). The very poor size sorting of both lithic- and biotic-clasts in the main bonebed fits transport *en masse* by a sediment-heavy flood.

When the deaths of ungulates occurred, the Trinil paleo-river floodplain had sufficient herbaceous open-terrain or forest-understory to support herd-sized populations of *Axis lydekkeri*, *Bibos palaesondaicus*, *Bubalus palaeokerabau* and *Stegodon trigonocephalus.* A mosaic of forested and open vegetation seems likely. The main bonebed contained bioclasts of wood, leaves and reedy grasses, and nine forest-prone taxa are in the vertebrate assemblage (Table 3). Palynological sampling further strongly suggest that the paleo-lowland had herbaceous vegetation with forested portions, while the uplands of the drainages had montane rainforest. The aquatic species in the Trinil faunal assemblage indicate that the main bonebed formed in a perennial river linked to long-standing lakes and ponds. The drainage doubtless was affected by strong seasonal variations in water level, but water was seemingly not in short supply.

We advance proposals to explain the taphonomic and sedimentological events evident in the main bonebed. Intense volcanism or drought made much of the Trinil paleo-watershed uninhabitable, driving ungulates into refuges, one of which formed in the floodplain of the trunk Trinil paleo-river or a lowland tributary (Figure 12a). Catastrophic mortality decimated the ungulates. Almost all of the carcasses in the floodplain decomposed to the point of skeletonization, creating a scatter of the tens-of-thousands of bony elements. A lahar flood inundated the plain, sweeping up thousands of decomposed skeletal remains and transporting them to Trinil.

Hyperconcentrated flow incorporated logs, reeds and leaves, and river-living reptiles, fishes and molluscs. At Trinil, a pulse carried the *Pithecanthropus erectus* (*P.e.*) Skullcap and Femur I along with the bones and teeth of well-known Trinil fauna species. This followed a surge containing cobbly gravel and few large-sized bioclasts. The flood-related accumulation presumably was finished at Trinil within days or a few weeks, when the remaining *Pithecanthropus erectus* femurs were deposited.

Proposals such as these are testable by field-, museum-, archival- and analytical-research. Trinil fossils at Naturalis (Leiden, the Netherlands) and Museum für Naturkunde (Berlin, Germany) continue to be incompletely exploited as a paleontological resource for evaluating the formation of the bonebed and the paleoecology of the watershed from which it originated. New geological field investigations of the 1891-1908 excavation face considerable hurdles. Remnants of the main bonebed comprise a small portion what existed prior to excavation. This calls for a close linking of the features observable in these remnants to archival accounts. Stratigraphic relations between the left- and right-banks and terrace-versus-bedrock formations in the vicinity, which lack resolution, must be determined.

The record of the left-bank excavations at Trinil, as presented here, contradicts Berghuis et al. (2021) suggestion that strata excavated on the left bank were valley fill as young as the twenty-meter ∼0.1 Ma terrace remnant at Ngandong. Additionally, the main bonebed Trinil fauna (Table 3) correlates on biostratigraphic criteria with ∼0.8-0.9 Ma beds at Sangiran Dome (e.g., Grenzbank and Ngebung bonebeds, Trinil and Ngandong faunas Tables 5 and 6-G and 6-H), not the early Late Pleistocene Ngandong fauna (Tables 4 and 5-A and 5-B). The cranial form of the *Homo erectus* in the Ngandong bonebed is more derived anatomically than the *Pithecanthropus erectus* calvaria.

The lithofacies and biofacies of the main bonebed are essential to evaluating the >0.8 Ma of archaic hominin prehistory in the stratovolcanic drainages of eastern Java and southern Sundaland more broadly (Table 5, Figure 13; S Figures 21 to 24). By way of examples, we relate the geological age range of the Trinil fauna to *Homo erectus* at Sangiran Dome and Ngandong discovery sites, highlight *Stegodon-Homo erectus* large-mammal faunal association occurrences in western Java, and show seismic evidence for Pleistocene fluvial valleys under the Java Sea portion of the Sunda Shelf (S Figures 22 to 24). Investigating topics such as these is closely aligned with Dubois’ broad paleobiogeographic and paleoanthropological goals for his Sumatra and Java efforts over 130 years ago. Dubois’ efforts placed the Skullcap and Femur I in one sedimentary deposit along with thousands of other fossils, establishing that *Pithecanthropus erectus* lived in a biotically rich stratovolcanic watershed.

## ACKNOWLEDGEMENTS

This paper is an outgrowth of research that F. Aziz, J. de Vos and O.F. Huffman have done with colleagues over the course of decades, as the References largely reflect. We thank Dr. Joke M. Oppenoorth for making W.F.F. Oppenoorth’s personal photographs and lantern slides of Trinil available to us for scanning at Naturalis Biodiversity Center and donating many of them to the institution. Todd Green, Jeffrey Horowitz and Chris Huffman helped with graphics, and L.R. Todd with sections of text. From 2016 to 2019, OFH (Huffman 2016) and PCHA participated in fieldwork at Trinil and follow-up studies of the “Studying Human Origin in East Java” co-operative between ARKENAS (Pusat Penelitian Arkeologi Nasional; Sekretariat Perizinan Penelitian, Asing Kementerian Riset dan Teknologi), Indonesia, and the University of Leiden (Joordens et al. 2017).

---

**Table 6.** Characterizations of the bonebeds, Kedungbrubus-Butak area, and selected other sites in eastern Java (Table 5; also, Figures 1b and 13). [**tab6** Use ‘reto6’ to return to page 70]

**Table 6-A**. <u>TRINIL</u> (Tr in Table 5): *Pithecanthropus erectus* (*P.e.*) main bonebed at Trinil (**LB**, **LB**-**HK** and **HK**) accumulated in a Pleistocene stratovolcanic river drainage setting that exemplifies the paleo-watersheds in which all *Homo erectus* fossils so far discovered in eastern Java were found (Figure 14). The relationship is exemplified here with reference to the bonebeds and collection areas presented in Table 5 described below. **^1^** The Trinil main bonebed formed as a lens spanning ∼200 m of a paleo-river channel, and was generally about a meter thick. Based on firsthand accounts, most of the vertebrate bioclasts were broken, disarticulated, disassociated, well-preserved remains of terrestrial ungulates of the Trinil fauna (Table 3). The remains were mixed with bioclasts of aquatic vertebrates, freshwater molluscs and plants, including sedges and logs, giving the main bonebed a degree biotic diversity exceeding that in any other *H. erectus*-fossil-bearing deposit in eastern Java (Table 4). The assemblage included many craniums of cattle, buffalo and proboscidean, among thousands of smaller skeletal elements and isolated teeth (Endnotes A to F, S II). The matrix surrounding the isolated gravel-sized skeletal bioclasts consisted of fluvially transported volcaniclastic materials, and was dominantly well-indurated, very poorly sorted conglomeratic sandstone. This material filled in the endocranial space of *P.e.* Skullcap (Figure 2c). As is explained in the ‘Origin of the main bonebed’ section (DISCOVERY RECORD), terrestrial vertebrate fossils in the main bonebed (Tables 1 to 3) most plausibly originated from the catastrophic mortality of ungulate- and hominin- populations living in the flood zone of the Trinil paleo-river valley (Table 4, Figure 12). The valley lay between the Kendeng Hills and the foot of Lawu volcano and also might have had headwaters on Wilis volcano (Figures 11 and 12). After the vertebrate carcasses were skeletonized, the bones were transported by lahar flooding to Trinil along with aquatic fauna (Figure 12). As reviewed in DISCUSSION, the Trinil fauna also occurs in the Grenzbank bonebed (**G**, below). It contains reworked bioclasts and lithic materials derived from drainages in the Kendeng Hills and perhaps Southern Mountains, as well as stratovolcanoes (Figure 13). The faunal similarity between the main bonebed and Grenzbank suggests that the Trinil fauna is a good representation of the large vertebrate populations present in the region at that time. On the other hand, the taphonomic and sedimentary history of the main bonebed is closely similar to that of the Ngandong *Homo erectus* bonebed (**-B**, below).

**Table 6-B.** <u>NGANDONG</u> (Ng in Table 5): Excavation at Ngandong produced 14 *Homo erectus* specimens together with ∼25,000 other vertebrate bones from a ∼3m-thick terrace remnant which rests unconformably on deformed marly bedrock ∼20m above the Solo River in the Kendeng Hills.**^1^** The Ngandong *Homo erectus* bonebed (Ng of Table 5) is a thin, individual member of the terrace remnant, and consists of fossil-rich, coarse-grained, gravelly, very poorly sorted, partially diamictic, volcaniclastic bar sands (locally >5 fossils m^-3^). This facies was succeeded by >2m of river-bar sands and muddy laharic diamicton in a single depositional sequence that indicates a lahar-flood origin for the terrace accumulation.**^2^** The volcanic materials in this deposit originated from the Wilis and Lawu watersheds (Figure 11). The biotic materials in the bonebed resulted from mass-mortality and lahar-flooding events that were similar to those from which the Trinil main bonebed formed (Figure 12).**^1^** Since their discovery in 1931-1933, the Ngandong *Homo erectus* specimens generally have been considered to be among the geologically youngest representatives of the species (due to their advanced anatomical features and occurrence in a terrace remnant perched above the modern Solo River). Radiometric dates now indicate that the hominin specimens and Ngandong fauna at the site are early Late Pleistocene age (117 to 108ka in modelled results).**^2^** Ancestor *Stegodon-Homo erectus* (*S.-H.e*.) populations had lived through maximum glacio-eustatic paleogeographic changes during the preceding 10-15ka (S I Figure 22a). Combined with radiometric and paleomagnetic studies at Sangiran Dome (-**G**, below), Ngandong discoveries establish that of *Homo erectus* occupied the stratovolcanic drainages of medial eastern Java for >0.8 Ma. One Ngandong cervid skull reportedly has cut marks, **^3^** but the ungulate fossils are otherwise notable for their lack of indications of human action. **^1, 2^**

**Table 6-C.** <u>KEDUNGBRUBUS FOSSIL COLLECTION AREA</u> (Kedung Brubus in Table 5): The Kedungbrubus area was source for the *Homo erectus* mandibular fragment that Dubois found in November 1890 (S I Figure 21). The find is one of three key *Homo erectus* discoveries made from the bedrock formations of the Kendeng Hills (-A and -E described the others). The hominin specimen is a partial mandibular corpus.**^1^** Dubois recognized the specimen as *Pithecanthropus erectus* after his discoveries at Trinil.**^2^** The hominin find very likely came from the outcrop area of the (later-designated) Kabuh Formation (S II-B7a). The Kabuh commonly produced freshwater molluscan- and vertebrate-fossils. The latter includes the Dubois Collection used to define the Kedung Brubus fauna (Table 5). The Kabuh Formation around Kedungbrubus is “225 meters of … coarse [-grained], andesitic sandstones,” which are “often cross-bedded,” and contain conglomerate lenses, “occasional ash-tuff layers” and marly beds of reworked marine materials from the older formations in the Kendeng Hills.**^3^** While fluvial accumulation appears to dominate the Kabuh Formation in the Kedungbrubus area, and its situation on the north flank of paleo-Wilis is clear, the paleogeographic configuration of drainage system during Kabuh deposition remains uncertain.**^4^**

**Table 6-D.** <u>BUTAK BONEBED</u> (Bk in Table 5): The Butak bonebed crops out at Butak peak northeast of the Kedungbrubus area. The Kabuh Formation in the area is underlain by 425m of the Pucangan Formation; 275-m of this formation consists of gravely diamicton-rich members and the rest is rich andesitic sandstone; the lithofacies represent the onset and continuation of volcanic activity at paleo-Wilis.**^1^** The thick stratigraphic mass of diamictic-and interbedded-sandy volcaniclastic rocks extends laterally in outcrop for >35km east-west.**^2^** The exposed relationships make for an open-air cross-section of the cone-shaped northern flank of the immense Pleistocene paleo-Wilis stratovolcano, the full paleogeographic effects of which extended from Trinil to Mojokerto along major river drainages (Figure 11 to 13).**^4^** The west-flowing paleo-drainage was part of the Trinil paleo-watershed (Figure 12). The east-directed drainage is clearly crop out in the eastern Kendeng Hills of the greater Mojokerto area, far to the east of Kedungbrubus (Figure 13). The Butak bonebed contained *Stegodon,* large bovid, *Axis* deer, *Rusa* deer, *Duboisia*, pig, hippopotamus, anteater, tiger and crocodiles (Table 5, note). The first-known vertebrate fossils in the section, the bonebed is just above the lower boulder-diamicton member, which gave a 1.87 Ma K-Ar isochron (whole-rock analysis of an andesite clast).**^3^** The Butak bonebed indicates that lahar-prone slopes of Early Pleistocene Wilis strato-volcano were inhabited by a diverse large-mammal fauna.

**Table 6-E.** <u>PERNING *HOMO ERECTUS* BONEBED, MOJOKERTO AREA</u> (Pn in Table 5): The Kendeng Hills north of Mojokerto affords good exposures of the Pucangan Formation. Facies changes within its members represent the long-standing Mojokerto paleo-delta.**^1^** Generally, volcaniclastic non-marine facies on the southwest, which were derived substantially from paleo-Wilis, transition episodically toward the east and north into muddy-marine strata of the former Madura Strait (Randublatung embayment, Figure 13). The Perning *Homo erectus* bonebed is a thin lithified conglomeratic sandstone lens deposited in a small lobe of the paleo-delta; the lobe is represented in outcrop by a ∼70-m-thick deltaic sequence of the upper Pucangan Formation.**^2^** Pollen spectra and grass-phytoliths from the ∼70-m-thick sequence show that the paleo-delta lowlands were characterized by dry grasslands when the Perning *Homo erectus* population lived, but also indicate the area had mangroves, swamps, delta- and riparian forests and distant montane vegetation.**^3^** The grasslands indicate Pleistocene aridity similar to modern climates hundreds of kilometers to the east in one of the driest parts of Indonesia (Nusa Tenggara; e.g., Flores).**^4^** The large mammal species in the upper Pucangan at Perning and Jetis is the Kedung Brubus fauna and many taxonomic elements also occur in the Trinil fauna (Tables 2 and 5).**^5^** Even forest-prone *Rhinoceros sondaicus* and *Panthera tigris* had habitats in the paleo-delta around the time that the Perning discovery bed accumulated. The paleogeographic setting of the Mojokerto child skull discovery at Perning indicates that *Homo erectus* and the Kedung Brubus (large-mammal) fauna inhabited the Mojokerto paleo-delta and -river valley(s) under particularly dry climatic conditions.

**Table 6-F**. <u>BUKURAN BONEBED, SANGIRAN DOME</u> (Sg in Table 5): The Sangiran Dome is a gently dipping diapiric outcrop of Plio-Pleistocene beds exposed over ∼5 by 8-km uplift in the midst of the Solo Basin. The Bukuran bonebed biofacies and lithofacies suggest that early hominin populations encountered complex peri-lacustrine habitats and variable climates in the paleo-Basin (Figure 13). Where excavated, the Bukuran bonebed was ∼0.4m of sandy silt with seed fossils and skeletal remains. The vertebrate assemblage (NISP = 394) consists of 60% aquatic species (∼55% fish, 39% turtle, 6% crocodile and 2% bird fossils); the mammalian specimens are 29% cervid, 9% bovid, and 4% hippopotamus, and notably lack the *Stegodon*, *Duboisia* and *Panthera* which occur at other fossils sites at the Dome.**^1^** The Bukuran bonebed, which is in the upper Pucangan Formation (see Table 5, note 3), contains remains of a biota that lived within and surrounding a paleo-lake: The ∼10m of strata that contains the bonebed includes a layer of foraminiferal sand derived from the erosion of distant marine bedrock, thin tuffs which represent far-off pyroclastic eruptions, an ∼0.75m-thick peaty-shell deposit that reflects an invertebrate-rich lacustrine fauna, and palynological indications of a *“thick growth of grass … Cyperaceae* [sedges and] *ferns”* covering a lowland that varied over time from swamp to savanna.**^2^** The upper Pucangan is thought to contain the earliest *Homo erectus* fossils at Sangiran Dome (Table 5, note 4). The ”radius … of a … *Bos* sp. [from Bukuran] …. revealed two clusters of cut marks …. [indicating] hominids’ intentional defleshing” of a carcass (this taphonomic reporting needs confirmation, and discovery circumstances and stratigraphic level of the *Bos* specimens are not specified).**^3^** Paleosols in the upper Pucangan record protracted periods of lowland exposure under “strongly seasonal climate with a dry season oscillating between short[er] and long[er] durations.” **^4^**

**Table 6-G.** <u>GRENZBANK BONEBED, SANGIRAN DOME</u> (Gb in Table 5): The Grenzbank (“boundary bed”) is the lowest member of the Kabuh Formation and unconformably overlies the Pucangan Formation (see Table 5, note 3).**^1^** The Grenzbank has a Trinil-like mammalian assemblage that is substantially more diverse taxonomically than vertebrate collections from the Pucangan Formation. Sandstone in the Grenzbank is often cross bedded and streaked with dark heavy mineral laminae representing volcanic provenance. Carbonate-cemented pebbly sandstone lenses in the bonebed indicate that erosion of Kendeng Hills and Southern Mountains contributed sediment to the paleo-drainage in the Sangiran Dome area.**^2^** Most vertebrate remains in the bonebed had experienced “multiple reworking events that generally resulted in the … selective removal of …. low-density skeletal elements.”**^3^** For example, individual teeth make up 86% of 215 ungulate-, carnivore- and rodent-specimens that were excavated at in the Brankal-site from ∼2.25- m-thick Grenzbank “granule and pebble gravel [conglomerate] with mammalian and molluscan fossil[s]”; cervids, *Duboisia* and *Sus* were consistently found in six Brankal trenches, as were *Crocodylus*, Testudinoidea and fish; proboscidean and forest-prone species *Hystrix, Rhinoceros*, *Tapirus* and *Tragulus* also were present (but large-bovid remains were rare).**^4.^** The first hominin fossil discovered at Sangiran Dome was found in 1937. Since then, >80 skeletal hominin fossils have been discovered with ∼15% of them originating in the Grenzbank bonebed unit; this discovery density is higher than from any other stratigraphic level at the Dome **^5^** (but not as dense as the 14 *Homo erectus* finds that the Ngandong bonebed is reported to have contained, -**B**, above). The Grenzbank is ≥0.9 Ma, more likely <1.3 than <1.5 Ma.**^6^** The *Duboisia*, *Rhinoceros* and *Sus* in the Grenzbank indicates a Trinil-fauna (Table 5).**^7^** Since the Grenzbank accumulated at an unconformity, the deposit sampled the living fauna over a longer time span than was the case in the formation of the Trinil main bonebed. Thus, the terrestrial vertebrate assemblage of the Trinil fauna (Table 2) appears to have included all of the larger-mammal species then present in the intervolcanic watersheds of eastern Java.

**Table 6-H.** <u>NGEBUNG HOMININ BONEBED, SANGIRAN DOME</u> (Nb in Table 5): The Ngebung bonebed lies above the Grenzbank bonebed in the lower Kabuh Formation. The bonebed contains the only *in situ* evidence of tool used to consume large-mammal remains in the *Homo erectus*-bearing formations of Java. “Whole rock and single grain argon dating of volcanic effluents, ESR dating of volcanic quartz, combined U-series and ESR dating of enamel from herbivorous fossil teeth converge to assign the site an age of 0.8 Ma.” **^1^** The anthropological contents occur within <2.5m of claystone which also contains sandstone and conglomerate. The beds reveal traces of hominin inhabitation, partially disturbed by sedimentary movement.**^2^** The hominin occupation was situated along the clayey bank “of an ancient river.” **^3^** The clay accumulated at low elevation in a floodplain with poorly drained soils.**^2^** The non-aquatic vertebrate fossils (NISP = 246) are 81% *Axis*, *Bubalus* and *Stegodon*, representing a well-developed mid- *S.-H.e.* assemblage, a Trinil fauna with Kedung Brubus elements (e.g., *Epileptobos groeneveldtii*).**^4^** The bonebed included “an almost undisturbed [hominin] occupation floor” with pebbles that “appear to have been used or worked by man” but there is no evidence of a “workshop.” **^5^** A molar with *H. erectus* morphology (NG91-G10 no.1) was recovered. **^4, 6^** As further reported by the excavators, the ungulate “skeletons were already dismembered, the skulls opened, and the mandibles broken before deposition;” and, “impact and cut marks are visible on long bones from adult bovids as well as *Stegodon* tusks.” **^7^** The Kabuh Formation in the Ngebung portion of the Dome has three fining-upwards depositional cycles; paleosols (vertisols and protosols) and soil-carbon isotopes from the floodplain facies indicate water-tolerant soils supporting grasses, shrubs and trees (lower cycle) and well-drained soils formed under mixed vegetation and climate with annual dry seasons (middle and upper cycles).**^8^** The dating of the Ngebung bonebed (∼0.8 Ma) supports the Early Pleistocene dating of the underlying Grenzbank *Homo erectus* bonebed and upper Pucangan Formation’s Bukuran bonebed. The evidence of hominin activity in the Ngebung bonebed draws attention to the general dearth of *in situ* lithic artefacts in the *H. erectus* formations of Java, and therefore represents a break in the common association of stone-tools with archaic hominin activities. **^8^** Artefacts are common with archaic hominin-fossil areas of Flores, Sulawesi and Luzon to the east of Sundaland in Southeast Asia.

**Table 6-I.** <u>PATIAYAM, NORTH CENTRAL JAVA:</u> The Patiayam lies north of the Kendeng Hills and the mid-island volcanic belt of Java (Figure 1b), and is a window of strata surrounded by deposits of the historic Muria volcano (1625 m crest and ∼30 km across). The *S.-H.e.* fossils at Patiayam come from deposits of a Pleistocene strato-volcano that evidently was a paleo-island separated from the Hills by the Randublatung marine embayment (Figure 13).**^1^** The volcaniclastic strata were laid down near a lake or swamp, and have produced isolated fossils of *Stegodon trigonocephalus*, large bovids, cervid species, *Sus brachygnathus* and *Rhinoceros* sondaicus, together with *Crocodylus* and Testudines.**^1^** Species that occur are typical of the *S.-H.e*. and include both Trinil fauna and Kedung Brubus fauna elements. Two isolated premolars and six parietal fragments of an archaic hominin species attributed to *Homo erectus* were unearthed from tuffaceous siltstone within the Pleistocene volcaniclastic sequence.**^2^** It includes an older eruptive series dated (by K-Ar methods, mid 1980s) to 0.64-1.11 Ma and a younger one dated to 0.41-0.78 Ma.**^3^** Vertebrate fossils were collected at Patiayam as early as 1850 by Franz Junghuhn; the area was a target of Dubois’ 1891 field program before de Winter was called to Trinil to supervise initial excavations on the right bank.**^1^** The Patiayam collections indicate that *S.-H.e.* species ranged across most or all of the eastern Java and resided along the margin of southern Sunda Shelf (Figure 14). Lasem volcano (806 m) lies ∼60 km east of Muria in the Rembang Hills (Figure 13). No vertebrate fossils are reported from Lasem but a mudflow tuff there contained a paleo-flora. Bawean Island, which lies 160km north-east of Lasem in the Java Sea, evidently was a Pleistocene volcanic center, but seismic data between Bawean and Java reveals no other centers. No archaic-hominin sites or Paleolithic artefacts have been reported from the Rembang-Tuban Hills and Madura Island. In both areas, pre-Pleistocene carbonate formations are widely exposed, including karstic development in the Rembang range (Figure 13); these terrains were uplifted and deformed before deposition of the *H. erectus*-bearing formations.**^4^** Some caves have young archaeological materials but none appear to be contemporaneous with the Middle Pleistocene lithic assemblage of Song Terus, Southern Mountains. (**-J**, below).

**Table 6-J.** <u>SONG TERUS, SOUTH CENTRAL</u>: Song Terus cave lies in a karstic upland, Gunung Sewu (Thousand Mountains), lying at ∼50-500m in elevation and covering a 10-by-80-km area where annual rainfall is high for eastern Java (1500-3000mm). The upland is part of the older Southern Mountains volcanic and limestone terrain, which is trace intemittently from southeastern Sumatra to Bali along portions of the Indian Ocean margin of Sundaland.**^1^** At Song Terus, “a large number of rolled and fresh artefacts occur …. in certain layers” within Pleistocene “fluviatile … [sedimentary] remnants” in the deeper portions of the cave **^2^** “Retouched flakes” were unearthed in the hundreds with fragmentary skeletal fossils, including those of the forest-prone species of rhinoceros and tapir and are c. 300ky, [based on] combined U-series and ESR dating of enamel.” **^3^** The Southern Mountains differed from the stratovolcanic watersheds to the north in topography, soils, climate due to a proximity to the Indian Ocean coast (e.g., the Southern Mountains historically have a wetter November-May monsoon seasons than do parts of eastern Java to the north). The G. Sewu karst formed before ∼500 ka,**^4^** and the karstic upland was probably present throughout the hominin occupation of Java (Figure 13). The absence of archaic hominin skeletal fossils in the Southern Mountains is most likely due to systematic taphonomic decomposition of remains, rather than a lack of older Pleistocene large-mammal occupation, given the widespread preservation of bony fossils in the *Homo erectus*-bearing formations north of the Mountains (Figure 14).**^5^** Other Cave sediments in the Punung area produced the remains of forest species, such as siamang and orangutan. These species characterize the Punung fauna, in which *Homo sapiens* appears and no extinct species occur (Table 4, note 6).**^6^** Immigration of *H. sapiens* into Southeast Asia occurred before 63-73 ka, based on widely accepted studies considering genetic and archaeological evidence in Eurasia and Australasia.

A. **^1^** Huffman, O.F. 2017, Huffman et al. 2010a, 2012b, 2018, ter Haar 1931, 1934a, b; the Ng fossil concentration is in unit 2/II of Oppenoorth 1932 and unit II of ter Haar 1934b (Huffman et al. 2010a: Table 3 and Figures 3A, and 7, respectively), and Facies C of Rizal et al. 2020: (e.g., Fig. 2, and Supplementary Information: section 2, part 3).
B. **^1^** Huffman et al. 2008a,b, 2010a,b, 2012b; also, Sidarto and Morwood 2004, Susanto et al. 2004 . **^2^** Rizal et al. 2020: especially 384, Figure 2, and Supplementary Information 2(3): 4-7, OFH pers. observation. **^3^** Choi 2003.
C. **^1^** de Vos and Sondaar 1982, Duyfjes 1936, Storm 2012, van Es 1931. **^2^** de Vos 2014, Dubois 1907, 1924a, Tobias 1966. **^3^** Duyfjes 1934, 1935, 1936: 143, Hidayat et al. 1995, Huffman 2020: 9; also, van Es 1931, Kimura et al. 1995. **^4^** Sartono 1976.
D. **^1^** Duyfjes 1936, Huffman 2020: 7, 20-35, Itihara et al. 1985b. **^2^** van Es 1931, Huffman 2020: 23. **^3^** Bandet et al. 1989, Duyfjes 1934, 1936, Huffman 2020: 25. **^4^** Duyfjes 1934, 1936, 1938a-d (and unpublished mapping from the late 1930s), Huffman 2020: 41; also, Shibasaki 1995, Yamamoto and Suminto 1995.
E. **^1^** Duyfjes 1935, 1938a-d, Huffman 2020: 41, sheets 110 and 116. **^2^** Huffman and Zaim 2003, Huffman et al. 2005, 2006; also, Morwood et al. 2003 regarding dating. **^3^** Morley et al. 2020. **^4^** See Monk et al. 1987. **^4^** Aziz et al. 1995, Cosijn 1931, 1932, Huffman et al. 2006, 2007, von Koenigswald 1934; also, de Vos et al. 2007b.
F. **^1^** Aimi and Aziz 1985, Kadar et al. 1985. **^2^** Brasseur et al. 2015, Kadar et al. 1985, Tokunaga et al. 1985: 202-203, Yoshikawa and Suminto 1985; also, Sémah et al. 2016. **^3^** Choi and Driwantoro 2007: 51 and 55. **^4^** Bettis et al. 2009, Brasseur et al. 2015: 97.
G. **^1^** Brasseur 2009, Itihara et al. 1985c, Sudijono 1985, Zaim et al. 2011. **^2^** Zaim et al. 2011: 366-367; also, Bouteaux 2005, 2008, Bouteaux et al. 2007. **^3^** Aimi and Aziz 1985 (Trenches III and IV), Sudijono et al. 1985: 75. **^4^** Zaim et al. 2011; also, Brasseur et al. 2015, Itihara et al. 1985a, Matsu’ura et al. 2020, Widianto et al. 2001 (regarding artefactrs). **^5^** Brasseur 2020, Matsu’ura et al. 2020. **^6^** Sondaar 1984: 232; also, Bettis et al. 2009, de Vos 1994, Watanabe and Kadar 1985.
H. **^1^** Falguères et al. 2016; also, Brasseur 2020, Matsu’ura et al. 2020, Saleki et al. 1998. **^2^** Brasseur 2009, Ingicco et el. 2022, Sémah 2001. **^3^** Simanjunkak et al. 2010: 419; also, Sémah et al. 2003 (figures). **^4^** Bouteaux and Moigne 2010**. ^5^** Sémah et al. 1992: 443, also, Moigne et al. 2004a,b. **^6^** Zanolli 2013, Martín-Francés et al. 2018. **^7^** Sémah et al. 1992: 443, also, Moigne et al. 2004a. **^7^** Bettis et al. 2009, Brasseur et al. 2015. **^8^** E.g., Ingicco et al. 2022, Sémah, A.-M., et al. 2016; Sémah, F., et al 1992; Simanjuntak et al. 2010.
I. **^1^** de Vos 2014, Huffman 2001a, Huffman et al. 2000, Saléh 1867, Sartono et al. 1978, Siswanto and Noerwidi 2016, Soeria-Atmadja et al. 1988, van Bemmelen 1949, van der Geer et al. 2010, van Es 1931, Zaim 1989, 2010; also, Lunt 2013. **^2^** Zaim 1989, 2010; Y. Zaim, pers. comm. 2011. **^3^** Bandet et al. 1986, Bellon et al., 1989, Maury et al. 1987, Schuster 1911a, Zaim 1989; also, Lunt 2013, Mulyaningsih et al. 2008. **^4^** Nurani et al. 2008, Lunt 2013.
J. **^1^** Haryono and Suratman 2010. **^2^** Sémah et al. 2004: 54. **^3^** Ansyori 2010, Falguères et al, 2016: 9, Hameau et al. 2007, Sémah et al. 2004, Simanjuntak et al. 2010: 419. **^4^** Rizal et al. 2020, Simanjuntak 2002, van Bemmelen 1949, Westaway et al. 2007. **^5^** Huffman et al. 2012b. **^6^** Storm and de Vos 2006, Storm et al. 2005, Westaway et al. 2007; also, Kaifu et al. 2022.

## ENDNOTES

Annotated translated excerpts (A to H) of source documents concerning the paleontology of **LB**, **LB**-**HK** and **HK** in the 1891- to 1907-excavations with ecological notes on selected taxonomic lineages (I). In Endnotes A-H, we have <u>double underlined</u> the finds from the left-bank excavations, <u>single underlined</u> the right-bank finds, and used no underlying when the river side is unspecified. “Oversized” bioclasts have been given **bold** fonts. The letters from Kriele and KdW (Kriele and de Winter) were written to Dubois. Dubois memoranda and reports were written to sponsors in the Indies government. Longer excerpts are in Supplementary Materials II (S II-…), as cited below.

### Endnote A. *1891*

i. [Kriele’s letter, September 3, 1891 (S II-A1a)] Tinil [= Trinil] … I have been working here already for 3 days [Figure 3b, insert; Skullcap Pit] … and … found … different species. For instance, some small jaws of goats [*<u>Duboisia santeng</u>*] or kidang [a small cervid] as well as … horns …. elephant [*<u>Stegodon</u> <u>trigonocephalus</u>*] and buffalo [*<u>Bubalus palaeokerabau</u>*] …
ii. [Kriele’s letter, September 18, 1891, with annotations on a sketch (Figure 3b; S II-A1b)] Tinil … work site of Kriele [in the Skullcap Pit included] a bone-bearing spot which contained that **tusk and skull of the elephant** [*<u>Stegodon trigonocephalus</u>*, a find that Dubois apparently saw during a visit ten days earlier.] [At the] de Winter work site [right-bank]. He has found bones as well: deer horns [antlers] with a piece of that skull [*<u>Axis lydekkeri</u>* the small-bodied deer].
iii. [Dubois’ August 1891 memorandum, dated September 18 (S II-B2b)] An even more important discovery spot … was one in the currently dry portion of the Solo [River] bed. Primarily, found all together [in the left-bank LB, which Dubois saw on September 7-9], were the remains of *Cervus axis* [*<u>Axis</u> <u>lydekkeri</u>*], *Anoa* [*<u>Duboisia santeng</u>*], *Bubalus palaeoindicus* [*<u>Bubalus palaeokerabau</u>* and], *Stegodon* [*<u>S.</u> <u>trigonocephalus</u>*] in andesitic sandstone at Tinil [Trinil] as well as of the fossil land turtle [<u>Testundinoidea</u>] … besides *Unio* shells [<u>Mollusca</u>] and fossilized wood …. The numerous shells indicate that the bones settled here in fresh water. Once again the [the vertebrate fossils] are often fractured but no indication of violence was found [that is, no indication of predation by terrestrial animals]. Also, several vertebrae of a small ruminant were preserved in natural articulation …. The **rear skull half** of a fossil *Anoa* [*<u>Duboisia</u> <u>santeng</u>*] with horn was found …. The fossil wood appears similar to wood from a European bog….
iv. [Kriele and de Winter (KdW’s) letter, September 26, 1891 (S II-A1c)] Tinil …. Herewith I am sending six crates of bones [including] really beautiful specimens … amongst those found at my [Skullcap Pit was] …. a goat **skull with 2 horns** [*<u>Duboisia santeng</u>*] and a **deer skull with two horns** [a cranium with antlers of *<u>Axis lydekkeri</u>*] …. Nice specimens have also been found at de Winter’s site [on the right bank], for instance the **skull of a banteng almost complete** [the nearly whole cranium of *<u>Bibos</u> <u>palaesondaicus</u>*, an ancestor species to the native cattle of Java, called bantengs] and **deer horns** [*<u>Axis</u> <u>lydekkeri</u>* antlers].
v. [Dubois’ third-quarter government report 1891 (S II-B2e)] …. near Tinil …. A considerable number of fossils have already been excavated in two shallow sand[stone] ledges that were exposed on both sides by low water level in the bed of the Bengawan [Solo]. These fossils are generally better preserved and are distinguished in that they are more complete. Among them some nice pieces of skulls and jaws were found of a curious small ruminant [*<u>Dubosia santeng</u>*] …. more or less complete skulls of ***<u>Bubalus</u>*** *paleoindicus* [***B. <u>palaeokerabau</u>***], and molars, a tusk and two incomplete skulls of ***<u>Stegodon</u>*** [***<u>trigonocephalus</u>***]. In addition, beautiful leg shields of a terrestrial Turtle (probably *<u>Batagur</u>*) and the presence were noted of *<u>Bibos</u>* [*<u>palaesondaicus</u>*], *<u>Rhinoceros</u>* [*<u>sondaicus</u>*], *Sus* [*<u>brachygnathus</u>*], felis and [*<u>Panthera tigris</u>*] a small rodent (*Mus*? [Muridae, presumably the Trinil rat, *<u>Rattus trinilensis</u>* Musser 1982; see Musser 1982]) And one or two lower apes that cannot be further identified with certainty [<u>Primate</u>]. Among the reptiles: *<u>Crocodylus</u>* [*<u>siamensis</u>*], *<u>Garialis</u>* [*<u>bengawanicus</u>*] and *Trinonyx* [<u>Testudinoidea</u>]. However, the most important find [from the LB] was a molar (the upper right third molar) of a chimpanzee (*<u>Anthropopithecus</u>*). … Fossil wood and leaves and numerous river fossils clearly identify the formation [LB] as a fresh-water deposit. The **tree trunks** and leaves are always found horizontally and some bones have their natural articulation still preserved [in the LB]. Hence, the animal and plant remains are found at their original place of deposition and the beds have not subsequently been deformed.
vi. [KdW’s letter, October 5, 1891 (S II-A1d)] Tinil …. If a piece is missing from an excavated [fossil] bone, it is mostly because it was hit with a pickaxe or a crowbar [which were needed to excavate the hard lithology of the LB,] and after the bone has been hit, it is almost impossible to find the matching pieces [because the bone is so thoroughly fossilized and brittle that it shatters where struck a hard-enough blow].
vii. [KdW’s letter, October 19, 1891 (S II-A1j)] Tinil … I have found [in the left-bank Skullcap Pit] a **lower jaw** of a hippopotamus with 2 molars attached [no *Hippopotamus* occurs in the Dubois Collection from Trinil, Table 2].
viii. [Dubois’ September 1891 memorandum (S II-B2d)] [The] exposed … [LB] sandstone ledges in the river bed … [had been] excavated to below water level [into the PFZ] on both sides of the river …. The sediments [LB] were identified as fresh-water deposits by the occurrence of fossil wood and imprints of leaves, as well as numerous freshwater shells. A number of often well-preserved remains of vertebrates were found [in the LB]. *Anoa*, the small ruminant [*Duboisia santeng*]; many **antlers** and a **skull** of the fossil *Axis* deer [*A. lydekkeri*], buffalo [*Bubalus palaeokerabau*], **giant skulls**, *Stegodon* [*Stegodon trigonocephalus*] molars, and some incomplete **skulls** connected to extant species of *Batagur* [sp. Testudine], *Crocodylus* [*siamensis*] and *Garialas* [*Gavialis bengawanicus*]. However, the most important find of all was a molar (the upper right third molar) of a chimpanzee (*Anthropopithecus*).
ix. [Dubois’ October 1891 memorandum (S II-B2f and –B2g)] After it was discovered in September that a chimpanzee was also part of the Pleistocene fauna of Java through the find of a singular molar [1891 Molar; Trinil 1], we were now able to judge this species even better. This month, close to the place where the molar was found in volcanic tuff on the left bank of the Bengawan [the LB, Skullcap Pit], a magnificent **skullcap** was excavated [Skullcap; Trinil 2] that, at first glance, undoubtedly must be ascribed to the species *<u>Anthropopithecus</u>* (*Troglodytes*), just like the molar was. …. Fossils of other animals were excavated, amongst which a **skull** of *… Leptobos* [a large bovid, most likely *<u>Bibos palaesondaicus</u>*]. The unexcavated surface area of fossil-bearing strata along the Bengawan near Tinil, in which we have now dug to 1.50m below the water level [probably basal LB or through it]… [no “*Leptobos*” {*Epileptobos groeneveldtii* (Dubois1908)} is found today in the Trinil material of the Dubois Collection (de Vos and Sondaar 1982).]
x. [KdW letter, November 18, 1891 (S II-A1g)] Tinil …. At de Winter’s site [on the right bank] a small pig [*<u>Sus brachygnathus</u>*] **jaw** has been found still containing a canine tooth.
xi. [KdW’s letter, November 30, 1891 (S II-A1h)] Tinil …. I am sending … 6 crates [of finds] including one crate containing wood [that co-occurred with the vertebrate fossils] …. At the top where de Winter works [right bank], they have found many horns [probably antlers of *<u>Axis lydekkeri</u>*] in the side [of the natural embankment, perhaps in the upper LB].
xii. [KdW’s letter, December 8, 1891 (S II-A1j)] Tinil …. At de Winter’s site [right bank] they have found today more bones than ever …. [and in the Skullcap Pit] today I found there an **elephant skull** [*<u>Stegodon trigonocephalus</u>* cranium found] with even larger molars than I have ever seen before ….
xiii. (**xii**) [Twenty-eight Dubois Collection specimens had *“Tinil”* written on their surfaces in the field (S II-A1m). This spelling of the locality was only used in 1891. Among these specimens are those (per the museum catalog) of *Axis lydekkeri* (**partial mandible and cranium**), *Bibos palaesondaicus (***mandibles***), Chitra* sp. (**carapace**), *Duboisia santeng* (**craniums**), *Garialis bengawanicus* (*Gavialis* cranial fragment) and *Rhinoceros sondaicus.* The finds might have come from excavations on either bank of the river.]

### Endnote B. *1892*

i. [Dubois’ second-quarter report 1892 (S II-B3d)] Then the river banks were dug into so that after the water sufficiently lowered, a large surface area of the fossil-rich deeper bed [LB] would be obtained. Because of this preparatory work [on the left bank], the harvest of bones was similarly plentiful as last year’s only during the last days of June. Amongst these [fossils] were those of a deer species [*<u>Axis lydekkeri</u>*] that has been buried here in large numbers and the small ox-like antelope [*<u>Duboisia santeng</u>*], as well as *Stegodon* [*<u>Stegodon trigonocephalus</u>*], important specimens to better determine the species. Several vertebrae and ribs of a large ox [*<u>Bibos palaesondaicus</u>*] found in their natural relative position…. However … the bones are generally isolated and widely distributed and usually broken. Separation [disarticulation and breakage] by mechanical action of flowing water is not likely because they [the skeletal elements] look well-preserved (except for the fractures). … Only animal violence [predation] could … break them and distribute them, while they were still surrounded by soft tissue and articulated (and thus be protected against wear at the bottom of the current while it pushed them along). In addition, proof is provided by a fossilized piece of aorta filled with sandstone. However, in no case were the usually recognizable signs of the teeth of land predators observed without doubt, instead many breaks of the long bones were seen caused by dentitions larger than those of the strongest carnivores. The bones are in fresh water deposits and it is logical to hold large crocodiles [*Crocodylus siamensis*] responsible, the remains of which are found so frequently amongst the other animal remains (primarily teeth). Round holes in some softer bones appear to actually correspond to the cone shaped teeth of these gluttonous water monsters. But these reptiles cannot be the cause of death of such large numbers of animals of many different species of which we now find the remains accumulated in fresh-water volcanic sand and lapilli, presently hardened to tuff, and spread out over such large distances. At Trinil alone remains of many hundreds of deer were excavated over a few hundreds of square meters of surface area.
ii. [Dubois’ August memorandum, dated September 23 (S II-B4b)] …. The most abundant species continues to be the small *Axis* [*A. lydekkeri*] deer of which the antlers of as many as 50 individuals were collected this month, as well as several complete **skulls** [These finds might have come from the Femur I discovery trench, or excavations on the right bank] …. However, the most important find was a femur of the chimpanzee previously identified from a molar and skullcap, which evidently are from the same individual. …. This femur [<u>Femur I</u>; Trinil 3] possesses … such complete human characteristics … that there can be no doubt that our *Anthropopithecus* … had already adopted an erect posture…. [and] represents a line that brings humans in closer relation to their living relatives amongst the mammals. … evidently the entire skeleton was buried in the volcanic tuff [LB], although somewhat spread around. The skull was located about 10 meters … from the femur and the molar was very close to the skull … at the same level in the sediments …. At this time (September), a new section of bank wall is being removed so as to expose [in the 25-m Trench] another portion of the fossil-bearing deeper bed in which these precious remains are to be expected.
iii. [Dubois’ October memorandum dated November 25 (S II-B5c; also, -B5e).] The desired level with the bonebed [LB] was finally encountered at both ends of the elongated pit [the full 25-m Trench; Figure 3a]. A number of fossils were dug out [from the newly exposed upper LB], amongst which some very nice ones, but still no other parts of the *Anthropopithecus*. A small but important find was that of a molar of a lower ape, a M.3 [*<u>Macaca</u>* sp.] [Among seven Trinil macaque teeth in the Dubois Collection, one (no. 3789) is this *Macaca* M_3_ dextra (Hooijer 1962: 50, Figs. 4 and 5) “taken in 1892 from the trench of 25 m, indicated by a note in the box containing the tooth” (de Vos and Sondaar, 1982: 49). No. 3789 came from the “lowest level, ½ m below pe,” presumably the level at which the *Pithecanthropus erectus* fossils were found (de Vos 1989: 227). The molar of the Asian Porcupine *Hystrix lagrelli* (Dubois Collection no. 1482a, formerly assigned to *Acanthion brachyurus*) also originated “at the lowest level in the trench of 25 m, 1892” (de Vos and Sondaar 1982: 47).]

### Endnote C. *1893*

i. [Dubois’ June 1893 memorandum, dated July 21 (S II-B6b)] We started a new stretch of the bank, 40 meters in length and 5 meters in width [the 40-m Trench; Figure 3a] adjacent to and further landwards of it [the 1892 25-m Trench, was inundated on this date]. Soon [after digging through the soil at the top surface of the bank], this work progressed extremely slowly because of the severe hardness of the rocks and the relatively low number (40) of available forced laborers. So as to remove this shallower fossil-poor rock mass as quickly as possible, an additional 10 to 20 paid coolies were temporarily employed.
ii. [Dubois’ second-quarter report 1893 (S II-B6c)] Because of the high-water level… we could not yet access the deeper exposed rich bonebed [LB in the 25-m Trench] …. [and we] instead initiated excavation in a new strip of the riverbank more landward and adjacent to it [the 1892 Trench] with the dimensions of 40 m in length and 5 meters wide [40-m Trench; Figures 2b and 3a]. The work progressed extremely slowly because of the extraordinarily severe hardness of the rocks, with the result that during this quarter [April-June] the aforementioned rich bonebed did not yet become exposed.
iii. [Dubois’ July 1893 memorandum (S II-B6c)] During this month [of July], excavations at Trinil again progressed extremely slowly because of the severe hardness of the rock …. and … the relative paucity of fossils in the beds encountered [above the LB].
iv. [Dubois’ August 1893 memorandum (S II-B6d)] At Trinil, the deeper fossil-bearing layer [LB] was partially exposed [in the 40-m Trench]. But the severe hardness of the rocks [in the LB] continued to be a handicap towards speedy progress of the work and also seriously impeded proper retrieval of the fossils by keeping them intact. … Nevertheless, some attractive specimens were obtained, most of them of a small deer species [*<u>Axis lydekkeri</u>*] of which many hundreds are buried at Trinil and also of *Boselapahus* [*<u>Duboisia santeng</u>*]*, <u>Stegodon</u>* [*<u>trigonocephalus</u>*] etc. In addition, the first almost complete **skull** of the previously discovered crocodile was found. It was now possible at this time to assign this species to a position in the system as far as available comparative material permitted; the species cannot be distinguished from *Crocodylus porosus* [later attributed to a new species *<u>C. siamensis</u>*].
v. [Kriele’s letter, August 27 (S II-A3c)] After you departed [on August 19], we found 1 nice elephant **tusk** [*<u>Stegodon trigonocephalus</u>*], 1 crocodile **skull** [*<u>Crocodylus siamensis</u>*], 1 antelope **skull** [*<u>Duboisia</u> <u>santeng</u>*], 1 turtle [<u>Testudinoidea</u>], a few leg bones, deer antlers and a few ribs and vertebrae [*<u>Axis</u> <u>lydekkeri</u>*]. The work is progressing very well, but we have not advanced as deep as the target layer [PFZ] in the middle of the [40-m] trench [which was near the Femur I discovery spot in the 25-m Trench]. It will take at least 4 to 5 days before we are at that depth. As we agreed, I will dispatch the bones on the first of next month.
vi. [Kriele’s letter, September 1 (S II-A3c)] [I am sending] herewith 14 crates, one of which contains [fossil] wood. … We are now finding rather many bones [in the 40-m Trench]. At this moment we have encountered the target layer [PFZ] in which much of that **wood** occurs and, by the way, also these shells [<u>Mollusca</u>].
vii. [Kriele’s letter, September 4 (S II-A3d)] Saturday we found an elephant’s **head** [*<u>Stegodon</u> <u>trigonocephalus</u>*] with the molars still in it.
viii. [Kriele’s letter, September 12, 1893 (S II-A3e)] We are working now at such a depth that on one side of the pit [east end of the 40-m Trench that] a different layer [below the LB] emerges, by the way, because the other end of the pit is already 0.25m deeper, and we are still in the same sandstone [LB there]. We will be obliged to keep bailing out water at the one end as the black soil [dark-colored claystone, the different layer] slopes down downstream. The work is going very well and quite a number of bones are being found, although nothing special lately.
ix. [Kriele’s letter, September 24 (S II-A3f)] Since you have been here [nine days ago] the water has risen 1 meter twice and is currently still at that level…. As to the finding of bones: still the same amounts as in the previous days but nothing special.
x. [Kriele’s letter, September 29 (S II-A3g)] because of high water, [even with] 5 people working [bailing] through the night can hardly keep up …. Hardly any bones, a few deer antlers [*<u>Axis lydekkeri</u>*] and some vertebrae [evidently they had passed through the PFZ and were in fossil-poor sandstone].
xi. [Kriele’s letter, October 4 (S II-A3i)] I could not send you the elephant’s **head** [with molars reported on September 4], as I did not have a crate [big enough] for it. … The river has subsided so much during the last 4 days that it is again at its low[est] level [∼LWL]. I have now dug the bail pit about 0.75 meters deeper [than before] but I am still in the same sandstone formation except that it is becoming a little coarser downward. Finding of bones is starting to diminish quite a bit, almost nothing has been found over the last few days. Because of the fact that [the base of] this sandstone formation [a lower conglomeratic LB] is dipping very steeply in the downstream direction. The river … is again at its low[est] level [∼LWL]. I have now dug the bail pit about 0.75 meters deeper … in the same sandstone … [which] is becoming a little coarser downward [below the PFZ, and the] finding of bones is starting to diminish quite a bit.
xii. [Kriele’s letter, October 7, 1893 (S II-A3j)] Work is progressing very well now [in the 40-m Trench], because in the morning we start with a dry trench [perhaps a portion in which the PFZ had not been yet dug ?]. We have found a buffalo **head** [*<u>Bubalus palaesondaicus</u>*], two small antelope **heads** [*<u>Duboisia santeng</u>*], as well as a leg and some ribs and vertebrae that will be in our next shipment.
xiii. [Kriele’s letter, October 9 (S II-A3k)] I believe that it would be wise to discontinue digging in this coarse layer [the conglomerate below the PFZ], since I do not believe that we will find any bones, and since the only thing that is being produced is almost entirely coarse gravel. In general, very little [in fossils] is being found in this [conglomerate] during the last few days.
xiv. [Kriele’s letter October 14 (S II-A3l)] I have now progressed so fast with our [excavation] work that at this moment we are more than a meter below the level of last year’s pit [that is, probably ∼ 1m below the PFZ in the 25-m Trench], and I would like you to come over and have a look, before I have them do pointless work. In about the whole pit [25-m and 40-m Trenches, combined] we are at the hard coarse [conglomeratic] layer, on some spots already through it a bit [into the underlying claystone]. As nothing [in the way of important fossils] has been found in it [the conglomeratic portion], I wanted to ask whether we should finish out [excavating] this hard, coarse layer. It’s thickness up to the black ground [claystone] is about 0.75m, and it [the conglomerate] also contains a lot of big stones, half a meter in diameter or even larger. The last few days hardly any bones were found. They become fewer by the day, yesterday, the 13^th^, e.g. 2 deer antlers [*<u>Axis lydekkeri</u>*], a small leg with 1 [?] and 2 vertebrae.
xv. [Dubois’ third-quarter report, November 8 (S II-B6f)] The new excavation at Trinil on the left river bank of the Bengawan was continued up to a depth of 1 to 1.75 meters below the level where previously the skull and femur of *Anthropopithecus erectus* had been found and where the black coaly clay bed was reached. This [contact with the clay bed] is 11 to 12 meters below ground level [atop the terrace embankment] and 3 meters below river level during the East Monsoon [∼LWL]. This black clay bed proves to be the underlying formation to the bone-bearing volcanic tuffs here [LB including lower conglomerate], but also elsewhere along the Bengawan. Up to the level of this clay bed, skeletal remains of already known Kendeng vertebrates were found in these tuffs, first of all many antlers of the small *Axis-*like deer species [*<u>A. lydekkeri</u>*] and also remains of *<u>Stegodon</u>* [*<u>trigonocephalu</u>s*]*, Boselaphus* [*<u>Duboisia santeng</u>*]*, <u>Bubalus</u>* [*<u>palaesondaicus</u>*]*, Hyaena* etc. Also [we found] the first almost complete **skull** of a crocodile [*<u>Crocodylus</u> <u>siamensis</u>*] …. However, among these remains were none of the Java *Anthropopithecus.* After this [40-m Trench was completed], work was resumed in the old excavation [25-m Trench] that had to be abandoned last year because of the commencement of the West Monsoon, and in which the skull and femur of the above anthropoid had been found [the 25-m Trench].
xvi. [Dubois’ November 1893 memorandum (S II-B6h)] … we were able to essentially dig away the entire bone-bearing bed [LB] in the excavation pit [25-m and 40-m Trenches] before the 26^th^ when the work [site] became hopelessly inundated. The collected fossils from this period were very carefully analyzed, but it turned out that none among them were the hoped-for additional parts of the curious *Anthropopithecus*. Although they almost certainly must once have been present [near the Skullcap and Femur I find spots in the LB], it seems that they have been washed away during formation of the river bed, together with a large portion of the bone-rich tuff [in which they were embedded]. Further searching for this transitional form therefore appears to be fruitless.
xvii. [Dubois’ Fourth-quarter report 1893 (S II-B6i)] The collected fossils were carefully examined, but it turned out that none of the still anticipated parts of the curious anthropoid (for which the excavations at that location were primarily instigated) were amongst them. Having once almost certainly been present although quite spread out [within the LB], as is generally the case with other species, these bones must already have been washed away during formation of the current river bed, together with a large portion of the bone rich tuff. It appears therefore to be fruitless to continue searching for this transitional form.

### Endnote D. *1895-1899*

i. [Kriele November 1895 (S II-A4b)] … we were able to find [in the LB-HK Ledge adjacent to the Skullcap Pit (Figure 4a), an] … incomplete **antelope skull** [*<u>Duboisia santeng</u>*, and a] … **elephant molar** [*<u>Stegodon trigonocephalus</u>*] …
ii. [Kriele’s letter, September 7, 1896 (S II-A4c, page 29)] I am now digging already 13 days on the opposite side of the river [left bank] in a sector of 10x10 meters [1896 Left-bank Pit], which [was dug downward from the top of the bluff, and] …. we have to dig more than 2 meters below the low water level [to fully penetrate the LB-HK] …
iii. [‘Kriele’s letter, September 16 1896 (S II-A4c, page 33)] … I have now progressed [down in the 1896 Left-bank Pit] … as deep as the bone bed [LB-HK], but nothing … except for a **deer skull with complete antlers** [*<u>Axis lydekkeri</u>*], and an incomplete **crocodile skull** [*<u>Crocodylus siamensis</u>*], as well as several incomplete and complete deer antlers, leg bones, phalanges etc. [*<u>Axis lydekkeri</u>*] Anything special about the man-ape has not been discovered yet…
iv. [Kriele’s letter, October 1 1896 (S II -A4c, page 35)]. The latest sector [of the 1896 Left-bank Pit] has just about been worked to its proper depth [through the LB-HK]. In total, we have now excavated to a depth of 11.50 meters [below the top of the embankment] and we are already 1 meter below the lowest water level [∼LWL]. We found [LB-HK to contain] 1 **complete turtle** [<u>Testudinoidea</u>], 1 incomplete] **crocodile skull** [*<u>Crocodylus siamensis</u>*, 1 incomplete deer **skull** [*<u>Axis lydekkeri</u>*], 1 incomplete lower jaw of a pig [*<u>Sus brachygnathus</u>*] and also several incomplete and complete deer antlers [*<u>Axis lydekkeri</u>*], leg bones, vertebrae etc. etc. I have also started on a small sector upstream … left behind in the past [the Ledge; Figure 4a] and …. we find wood, shells and bones [from the upper LB-HK there], but the bone is mostly not very useful [for taxonomic identification].
v. [Kriele’s letter, August 31, 1897 (S II-A4d)] …. This [left-bank 1897 Upstream Pit] is a very nice trench [which is located] upstream and adjacent to the trench of previous years [the 25-m and 40-m Trenches]… . Up till now, 7.5 meters [in depth] has been excavated, measured from the surface [atop of the embankment] and … [just a few] small bones have been found but nothing very special. …
vi. [Kriele’s letter, September 9, 1897 (S II-A4d)] .… the [1897 Upstream Pit] pit has now been dug 9 meters [downward from the embankment-top surface, and thus just above or within the upper LB-HK] … one complete **deer skull with antlers** [*<u>Axis lydekkeri</u>*], a small kind of **tusk** of an elephant [*<u>Stegodon</u> <u>trigonocephalus</u>*], and several diverse vertebrae, carp/tarsals, teeth and molars but … we are not yet deep enough [into the LB-HK] to find larger amounts…
vii. [Kriele’s letter, November 11, 1897 (S II-A4d)].… [the left bank 1897 Downstream Pit of] 10×8 meters … already brought 8 meters down [from the surface and digging just above LB-HK], but nothing [except] some deer [*<u>Axis lydekkeri</u>*] antlers, vertebrae, an elephant’s molar [*<u>Stegodon trigonocephalus</u>*], a rhinoceros [*<u>Rhinoceros sondaicus</u>*] molar, some legs, teeth and molars [have been unearthed].
viii. [Kriele’s letter, January 12, 1897 (S II-A4d)] … The [1897 Downstream Pit] was dug [through the LB-HK] to a depth of 12 meters. … we found [in it] 1 complete **tusk** of an elephant [*<u>Stegodon</u> <u>trigonocephalus</u>*] **of 1.55 meters** [length], 2 incomplete lower jaws with complete molars, 2 very incomplete upper jaws with complete molars, 1 incomplete **skull of a cow with complete horns** [*<u>Bibos</u> <u>palaesondaicus</u>*], 1 incomplete **skull with** 1 complete **horn of a water buffalo** [*<u>Bubalus palaeokerabau</u>*] as well as some deer antlers [*<u>Axis lydekkeri</u>*], leg bones, vertebrae and a variety of teeth and molars. …
ix. [Kriele letter, September 16, 1899 (S II-A4h)] … I have started … a trench of 24 meter in length and 6 meter in width [first-dug was the central part of the 450 m^2^ 1899 Trench] …. [We] reached the bonebed [LB-HK] in which so far we have not found anything special. Still, we did find … an elephant [*<u>Stegodon</u> <u>trigonocephalus</u>*] **tusk** with a **length** of about **2 meters** in 4 pieces and a complete upper **skull** portion of a **rhinoceros** [*<u>Rhinoceros sondaicus</u>*], as well as some nice small leg bones, vertebrae, [*<u>Axis lydekkeri</u>*] deer antlers and various teeth and molars.
x. [Kriele’s letter, December 8, 1899 (S II-A4k)] … recently found 1 incomplete **buffalo** [*<u>Bubalus</u> <u>palaeokerabau</u>*] **skull** and 1 incomplete mandible of a tiger [*<u>Panthera tigris</u>*] … also a molar [a lower premolar, according to Dubois’ annotation] as well as a tooth [an ape or monkey tooth, according to Dubois’ annotation] …. [that was] found about 0.5 meter above the lowest water [upper LB-HK; these teeth, which apparently were non-human <u>Catarrhines</u>, were found southwest and southeast of the Skullcap Pit of 1891].
xi. [Kriele December 21, 1899, list (S II-A4l)] … I am sending herewith a list of the contents of 7 full crates [from the 1899 Trench, including in part an] incomplete Elephant [*<u>S. trigonocephalus</u>*] **skull with complete tusks and molars**[,] a complete **buffalo skull with horns** [*<u>Bubalus palaeokerabau</u>*] and molars[,] an incomplete **buffalo** [*<u>B. palaeokerabau</u>*] **skull**[,] a small crate [with] 40 various small vertebrae, 2 antelope [*<u>Duboisia santeng</u>*] horns[,] 1 tiger [*<u>Panthera tigris</u>*] mandible[, and] 2 pig [*<u>Sus brachygnathus</u>*] tusks and 5 molars[,] 3 incomplete [*<u>A. lydekkeri</u>*] deer skulls[,] 1 incomplete antelope [*<u>Duboisia santeng</u>*] skull[,] 15 [*<u>A. lydekkeri</u>*] deer antlers[,] 1 elephant [*<u>S. trigonocephalus</u>*] **tusk** in 5 pieces with a total **length** of **1.10 meter**[s, and] a small crate [with] 665 various teeth and molars. [The incomplete *Panthera tigris* mandible from the 1899 Trench is the second occurrence of the species reported from the excavations, and the first known to have come from a left-bank pit or trench].
xii. [A specimen of *Hystrix lagrelli* (Dubois Collection no. 1487) and the holotype of *Rattus trinilensis* (no. 1478) bear labels indicating they were found in 1899, presumably from the LB-HK, and are identified in the museum catalog (de Vos and Sondaar 1892; de Vos 1989; Muser 1982; S II-A4g). Several ape or monkey teeth were found in the 1899 Trench, most likely from the LB-HK. One of the two non-hominin Catarrhines in the might be the isolated silvery langur *Trachypithecus cristatus robustus* molar that is only specimen of this species from Trinil (no. 3738, Hooijer 1962; S II-A4k).]

### Endnote E. *1900*

i. [Kriele’s letter, February 22, 1900 (S II-A4o)] We have now worked down to a depth of more than 2 meters across the entire sector [of the first-dug portion of the ∼950m^2^ 1900 Trench]. There are no bones in this except for some insignificant pieces.
ii. [Kriele’s letter, April 6, 1900 (S II-A4o)] For the entire work site [a large portion of the 1900 Trench] we have now reached an average total depth of around 4 meters. We have not yet extracted any bones from this, except for some insignificant specimens.
iii. [Kriele’s letter, August 24, 1900 (S II-A4o] …. We have not found many bones this month because we had to work on top most of the time.
iv. [Kriele’s letter, September 15, 1900 (S II-A4o) [In the first-dug section of the 1900 Trench] we have excavated … to a depth where no longer bones are present …. [below the LB-HK and while penetrating it] found: a complete **buffalo skull with complete horns** [*<u>Bubalus palaeokerabau</u>*] and molars; a complete **elephant skull with complete tusks** and molars [*<u>Stegodon trigonocephalus</u>*] and further an assortment of specimens. I regret to write that nothing has yet been found of the ape-human. …
v. [Kriele’s letter, October 10, 1900 (S II-A4o)] We are now working in the last trench upstream [part of the 1900 Trench] and very close to the spot where recently the molar of the ape-human was found [Skullcap Pit]. But we have not yet found anything of him [the *Pithecanthropus erectus*]. We did find an incomplete turtle [<u>Testudinoidea</u>]; a complete lower **jaw** of an **elephant** with molars [*<u>Stegodon</u> <u>trigonocephalus</u>*] and some assorted bones.
vi. [Kriele’s letter, October 21, 1900 (S II-A4o)] During the last few days we found two incomplete turtles [<u>Testudinoidea</u>] and an assortment of several leg bones, an elephant [*<u>Stegodon trigonocephalus</u>*] molar and an incomplete **skull of a cow** [*<u>Bibos palaesondaicus</u>*]. These were all found [in the LB-HK] … next to where last year the two molars [a molar and a tooth] of the ape-human were found [in the 1891 Skullcap Pit and 1892 25-m Trench].
vii. [Kriele’s letter, November 1, 1900 (S II--A4o)] We recently found turtle [<u>Testudinoidea</u>], 2 incomplete deer **skulls with several horns** [*<u>Axis lydekkeri</u>*] and some assorted leg bones, vertebrae, ribs, teeth, molars and small jaws. I regret to say that we have not been able to get anything of the ape-human.
viii. [Dubois (1926a,b, 1932, 1934, 1935) recognized *<u>Pithecanthropus erectus</u>* femora II, III and IV and attributed them to the 1900 Trench (S II-A4m).]
ix. [One specimen of *<u>Prionailurus bengalensis</u>*, present in the Dubois Collection (no. 1484), was found in the trench of 1900, based on a note with the specimen (de Vos and Sondaar 1892; de Vos 1989).]
x. [Kriele’s 1900 letters mention numerous “oversized” ^2^ finds: A *<u>Bubalus palaeokerabau</u>* **cranium** complete with horn cores; a *<u>Stegodon trigonocephalus</u>* **cranium with tusks**; partial turtle remains; a *<u>Stegodon trigonocephalus</u>* **mandible**; two partial *<u>Axis lydekkeri</u>* craniums; and an incomplete *<u>Bibos</u> <u>palaesondaicus</u>* **cranium**; etc. (S II-A##). A list made when unpacking crates has discoveries attributable to the LB-HK with characterizations of the lithology of the embedding deposit (S II-A4p and –A4q): An *<u>Axis lydekkeri</u>* **cranium** without horns in fine-grained blue sandstone, 1.0-1.5m below the lowest river level; a metacarpal in fine blue sandstone, 1.0-1.5m below the lowest level; *<u>Axis lydekkeri</u>* **cranium** in coarse sandstone, even with lowest river level; *<u>Bibos palaesondaicus</u>* occipital from section 5, unit 157, in coarse sandstone, 25 cm below the river level; *<u>Bibos palaesondaicus</u>* **calotte with two horns** from 1.50m below lowest river level; *<u>Stegodon trigonocephalus</u>* **supra-occipital bones** from trench section I, unit 149, in coarse sandstone 1.0m below the river level; *<u>Sus brachygnathus</u>* **hemimandible** in fine sandstone, 0.28m above the river level; <u>Testudinoidea</u> in fine sandstone, 1.0-1.5m below the lowest level; 4 tibial fragments, a partial rib, a vertebral fragment, 1.0-1.5m below the lowest level in fine blue sandstone.]

### Endnote F. 1907-1908

i. [Oppenoorth’s 1907 newspaper article published shortly after he left Trinil; Oppenoorth 1907] …. in the actual [main] bonebed [HK]. … all fossils are distributed rather randomly and one can find parts of a stegodon [*Stegodon trigonocephalus*], deer, buffalo [*Bubalus palaeokerabau*], predator, crocodile [*Crocodylus siamensis*] etc., sometimes more than 100 specimens had been deposited within a few square meters. … And it was in exactly this bed that the parts of *Pithecanthropus erectus* were found. … The major portion of these are [*Axis lydekkeri*] deer antlers, almost all being **six pointers** as well as a few smaller antlers mixed in …, crocodiles and deer teeth, which however are more specifically restricted to Pit II (on the left bank; Figure 7).
ii. [Oppenoorth (1908a] …. the bonebed was … easy to observe since the various layers [in the excavated sequence] show clear and sharp boundaries …. [The HK was] tuffstone …. [in which there] were hard clumps of clayey marl and lava bombs … distributed here and there, and by preference up against the [vertebrate] fossils …. Often, deer [*Axis lydekkeri*] **antlers** were found in between [the marl and lava cobbles], maybe **3 or 4** [of antlers occurring] **twisted together**. A test was conducted in Pit II with sieving of the material, as a large number of very small bones and teeth (sometimes 0.5 cm in size) were found there. The excavated material was mixed with water and poured onto a sieve with a mesh of 1cm^2^. However, …. no single small fossil was found in this manner. … More than 2000 specimen were collected [by hand excavation], as well as a large number of crocodiles [*Crocodylus siamensis*] and deer [*Axis lydekkeri*] teeth, which were not numbered separately [in the 1907 Listing]. About 1224 pieces came from Pit I spread out over a surface of about 350 square meters, hence on average 3.5 pieces per square meter. These were mainly the **large bones** like skulls, pelvis, vertebrae, ribs etc. of *Stegodon* [*trigonocephalus*], Bos [*<u>Bibos</u> <u>palaesondaicus</u>*] etc. Pit II produced about 700 fossils over a surface area of 250 square meters, hence about 2.7 specimen per sq. m, mostly smaller bones, teeth, vertebrae, hand- and foot-bones etc. Many times, the number of specimens found per square meter was larger (or smaller); for instance (gridpoint) II A 3 [in the left-bank Pit II], where primate teeth [<u>Primate</u>] were found yielded 7 specimens. Most groups from the animal world were represented. Mollusks were encountered in various layers, in the bonebed as well as in the underlying black clayey marl, which in some locations formed a bank of Melania [most likely *Melanoides tuberculata*, a fresh- to brackish-water gastropod]….
iii. [Oppenoorth (1908b: 145, 148)] From June to September [1907] the bonebed [**HK**] itself was worked on. …. The … so-called bonebed, consists of a bluish volcanic tuff, which reminds one of a very soft sandstone. The layer consists of three portions; the upper and lower contain the majority of the fossils, while the layer in between only contains a few of them. From top to bottom the fine-grained layer grades into a coarser-grained layer [sandy] which also contains several large lava bombs of a few decimeters in size. Sporadically, fossils occur in shallower layers, mainly of Bos [*Bibos palaesondaicus*] and *Stegodon* [*trigonocephalus*]. … Overlying the bonebed [HK] (or where the clay layer with leaf imprints is present [above it on the right bank Pit I]) is a brown- to bluish-clay which grades upward into a sandier composition, with thin sand intercalations and which eventually grades into a soft yellowish sandstone, called lahar sandstone by Carthaus. … Below the bonebed [HK] lies a much harder, coarse-grained, red–brown lapilli layer, which in places forms broad shoals in the river at low water levels [Dubois’ “Marine breccia,” Figure 2a]. Several fossils were found in this layer, but due to the hardness of the rock they could not be recovered, besides the fact that they always consisted of broken fragments. However, curious was the unusual fossilization that always resulted in [specimens that are] lighter colors than those in the bonebed. Overlying this [red-brown ‘lapilli’ conglomerate and directly below the HK] is a black–gray marly clay that is very brittle, with conchoidal fracturing and rich in Melania. This layer also varied greatly in thickness, from 0.2 to 2 meters.
iv. [Oppenoorth (1911: xxxi-xxxiv; Berkhout and Huffman 2021: 31-37)] At the end of April when the water level had become somewhat lower, [xxxii] work was started on excavating the left bank of the Solo [Pit II], where the bonebed had partly been washed free and a beautiful **skull** of *Bos* had been found [based upon other records this was *<u>Bubalus palaeokerabau</u>*]. … Pit II…. adjoined … the Dubois excavations had dug landward from near the edge where the discovery of *Pithecanthropus* had been made. …. [xxxiv] Once the bonebed [HK] was reached… further work was done with much care, and the layer was scratched away with patjols (hoes) until bones were reached [also, Figure 8]. {Footnote: Incidentally, the bones were mostly embedded in broken condition anyway, and in a few, it could be seen that they had definitely had broken before fossilization had taken place. Next to the **skull** of an immature *Stegodon* [*trigonocephalus*] … [for example] was a broken-off **tusk** of 30 to 40cm [in length] which was cemented to the skull in the wrong direction [presumably in the **HK**]. Also, in many other bone fractures could be observed that were filled with tuff}. Most of the bones were thoroughly silicified …. Extensive work was required with the remains of a huge *Stegodon* [trigonocephalus] … found in the upper beds of Pit II in light colored clay, about 5 meters above the bonebed (Table 2, Figure 8, S I Figures 9 and 10). They consisted of a skull with upper jaw and tusks (2.10 meters long …), thigh bone (over 1m. long), pelvis and ribs, while at about a distance of 5 meters from there, the associated lower jaw ….. Also, a few hippopotamus molars were found. Unfortunately, all these bones were poorly preserved, because they were not silicified like the deeper ones. The *Stegodon* [*trigonocephalus*] material is of course the most attractive of the finds. This species was found in nearly all beds, even at the surface in the top soil.
v. [Branca (1908: 270) about Carthaus’ work] The main bonebed [Hauptknochenschicht], as Carthaus names it … has a thickness of 0.40 to 1 meter [and] consists of finer masses of ash and lapilli within which only occasionally larger andesite pieces can be found. Among the numerous bones in this bed are also some mollusk shells….
vi. [Carthaus (1911b: 26-30; Berkhout and Huffman 2021: 85-88)] [26] The main bonebed [HK] … is characterized by undamaged animal bones, without traces of rubbing against stones in flowing water. They might easily have been transported over large distances by a tuff[aceous] mudflow, which is what is assumed for all material in the main bonebed. … [28] The bones in the [HK] main bonebed permit us to recognize two notable phenomena: first there are no traces on them of long-distance transportation; secondly articulated whole skeletons are absent. It can therefore hardly be assumed that still-articulated cadavers were swept along with the lahar flow into the main bonebed. From these circumstances it can be concluded that the lahar flow … transported more or less already decomposed animal corpses, which fell apart at the slightest friction, or a few isolated skeletal parts that were still partially articulated. Accordingly, a number of days, weeks or even months must have passed after these animals had been killed during the initial explosive eruption, before they ended up [29] in the lahar flow. With respect to the degree of fossilization: this can vary for the animal bones in the [HK] main bonebed, as well as for the wood remains, and was not as a result of the varying geological age of the bones, but visibly due to the variable nature of the lahar material surrounding the organic remains. The chemical properties are even more important in this regard than the mechanical conditions. [30] Moreover, it could be observed that fossilization had already advanced further in those that had large amounts of small pyrite crystals. … In addition, I want to mention that of the incidental bone remains that are found here and there in the beds overlying the [HK] main bonebed mostly appear to be much more weathered and leached, and thus tend to fall apart when exposed to air, than those found in the main [bone] bed. … [wherein the HK fossils] are heavier and more severely impregnated with mineral substances …. as a consequence of being [continuously] saturated [with ground water] and mineral salts effecting the enclosed organic remains longer and more strongly.
vii. [Dozy (1911b: 35; Berkhout and Huffman 2021: 93)] The [HK] main bonebed immediately overlies the conglomeratic tuff in both pits [on top of the Pucangan Formation of Duyfjes (1936)]…. The tuff conglomerate layers [of the Pucangan] …. were exposed to erosion for a long time, so that many terrain irregularities were formed at its surface [before HK deposition]. The bones, which had been laid down on the ground nearby, were then swept [off the erosional surface, and accumulated] together into these flat depressions. The distance of transportation was mostly very short, so that those bones did not undergo any rounding; the main bonebed was thus formed in this manner. … The presence of mollusks indicates that already the river flowed through the plains, probably with a small gradient and very large width.
viii. [Examples of HK entries in Pit II from the 1907 Listing (Museum für Naturkunde, Berlin, MNB document PM_S_II_ Selenka_FB_1-78), giving year, field number and character of the fossils (Table 2), including MNB catalog descriptions in {…} brackets] 1906 [sequential number entries] #2 [to] #55 [came from the] *Pithe*[*canthropus* bed of Pit] II [and include finds characterized as] Cervus antlers [*<u>Axis lydekkeri</u>*] … vertebra[e] … leg [bones] and rib [and] … foot bone[s]. 1907 [nos.] #29 [to] #65 [includng finds from the] *Pithe*[*canthropus* bed, and] vertebra[e] … leg bone[s] … lower jaw … [and] teeth. [Then on] June 12 [and] 13 [nos.] #161 [to] #187 … All [came] out of layer 3/blue-sandstone bed. Ent[ered on] 18-8-1907 [by] Opp[enoorth] … [the finds included an] elephant [*<u>Stegodon trigonocephalus</u>*] **tooth** #161 … deer [<u>Axis</u> <u>lydekkeri</u>] **mandible** #162 … foot bone … (pig?) thigh bone #164 … **Bovid skull** #165 … deer thigh bone #168 {MB.Ma.22553} … [several] pig **jaw**[s; *<u>Sus brachygnathus</u>*] … [*<u>Axis lydekkeri</u>*] deer antlers #178 … foot [hindlimb] bone [etc.]. [From] July 16 [to] October 4, #439 [to] #1813 [there are 201 from layers 3 and 4 in Pit II, nearly half matched with Museum specimens (L. Todd, pers. comm., 2015); examples include] vertebra #439 {MB.Ma.22316} … footbone #444 {astragalus MB.Ma.22509} … vertebra #469 {cervical vertebra of cervid MB.Ma.22329} … Cervus skull #471 {occipital fragment MB.Ma.22280} … deer jaw #478 {partial **mandible** MB.Ma.22115} … shoulder blade #490 {cervid scapula MB.Ma.22690} … foot bone #495 {complete cervid phalange MB.Ma.22636} … rib #564 {partial cervid rib MB.Ma.22383} … head of thigh bone #568 {partial femur MB.Ma.39320} … vertebra #630 {partial large-bovid thoracic **vertebra** MB.Ma.23381} … tooth (crocodile ?) {crocodile MB.R.4617.7} [*<u>Crocodylus siamensis</u>*] … vertebra #684 {complete cervid lumbar vertebra MB.Ma.22351} … deer jaw #755 {MB.Ma.22113} … vertebra #803 {partial thoracic **vertebra** of a large bovid MB.Ma.23379}… foot bone #806 {complete proboscidean metacarpal MB.Ma.17017} … deer antlers #1059 {complete right cervid **antler** MB.Ma.22453} … vertebra #1158 {partial large bovid lumbar **vertebra** MB.Ma.23395}… vertebra #1319 {complete axial **vertebra** of a large bovid MB.Ma.23424} … foot bone Calcaneus Elephas 1377 {complete Stegodon **metacarpal** MB.Ma.17014} [*<u>Stegodon trigonocephalus</u>*] … piece of (leg) bone #1572 {complete cervid phalanx MB.Ma.22625} … vertebra #1665 {complete cervid thoracic vertebra MB.Ma.22340} …vertebra #1690 {complete axial **vertebra** of large bovid MB.Ma.23412} … vertebra #1703 {partial vertebra of large bovid MB.Ma.23452}.
ix. [Further notes about the Pit II HK entries in the 1907 Listing: The HK finds reported from the *Pithecanthropus* stratum include (1) those collected in May, (2) layer “3” excavated during July and August (299 entries during the two months) and (3) layer “4” dug in September and October (106 entries). Most of the “3” finds have unit-grid locations in blocks A, B, C or E (Figure 7a). Bed “4” finds are from the western block C. Beds “3” and “4” might have been lateral facies in the HK or the same bed designated differently when a new team assumed Oppenoorth’s duties in August. Entries in the Listing indicate fossil density of ∼2m^-2^ over the central part of Pit II (405 entries for layers “3” and “4” in ∼210m^2^ of blocks A through C). Seventeen Pit II entries originated from field stratigraphic layers “5” to “9,” which evidently were within or below the HK. Some HK finds can be identified taxonomically because the 1907 Listing has diagnostic anatomical information on them (such as tusks or antlers) or records a recognizable animal name. In other situations, the sequential numbers can be linked to specimens that Selenka and Blanckenhorn (1911) identify, or the fossils still have a field number on them in the Museum für Naturkunde, Berlin (MNB) where the Museum catalog identifies the fossil taxonomically (L. Todd and M. Hill, pers. comm., 2016). A similar taxonomic profile is recognizable in the 1907 Pit II entries that lack bed designations. The excavated material was screened with water through centimeter-sized meshing, because Pit II produced “a large number of very small bones and teeth (sometimes 0.5cm in size);” however, “no single small fossil was found” in this way (Oppenoorth 1908a). Thorough recovery was due to examining the excavated material “twice” so that even “bone splinters” and “many small crocodile teeth and the minutest shark teeth … were found” (Oppenoorth 1911: xxxv; Berkhout and Huffman 2021: 38). The screens tended to become clogged with lithic granules of the HK. The bioclast density of the HK fits within the range normal for vertebrate bonebeds globally (Rogers et al. 2007).
x. [Summary of HK entries from Pit I in the 1907 Listing: Of 1136 entries which have layer designations from Pit I in the 1907 listing, 1060 (93%) came from the HK (layers “15” to “17” in the stratigraphic scheme of 1907), giving an average of 2.5 fossils m^-2^. Since the HK was 0.4 to 1.0 m thick in Pit I, the volumetric fossil density was higher than the spatial one; if a 0.7-m average thickness is used, for example, the volumetric density would be ∼3.5m^-3^. In 1907 Pit I, “the [HK] layer consists of three portions” with the upper and lower parts having more fossils than the middle subunit (Oppenoorth 1908b: 181). The 1907 Listing shows Oppenoorth’s three HK subdivisions in Pit I. “15” and “17” had 42% and 40% of the total HK finds (respectively), while the intermediate subunit had only 8% of the total. The HK also varied in bioclast occurrence from place to place in Pit I (S I Figure 11c). There were 99 squares with subunit-15- finds alone; 31 squares had layer 16 fossils alone, 132 subunit-17-only squares, 48 squares had both 15 plus 17 finds, 8 had fossils in all three layers, and 48 squares lacked any finds at all. Thus, across the 350 m^2^ 1907 Pit I, HK fossils were scattered laterally and vertically. The bioclasts were widely dispersed within a lithic matrix.

### Endnote G. Dubois’ 1908 paper on “The geologic age of the Kendeng- or Trinil fauna”

[Dubois’ 1908 paper expanded upon the taphonomic remarks that he gave in his 1892 2^nd^ quarter report, B(i) above, which had been written just before the Femur I discovery. The 1908 paper listed the taxa found in the 1891-1900 excavations (on pages 1239, 1241-1242, 1244-1245) but did not describe individual finds (S II-F7).]

[1240] It is true that many of the [fossilized] bones that have been dug up [in excavations] near Trinil (among them the skullcap of the *Pithecanthropus* [which] at its outer [anatomical] surface) have suffered from corrosion caused by groundwater containing sulfuric acid. The tuffs in this location [Trinil] happen to contain many sulfur compounds; pyrite has been deposited on the bones and on lignites, …. Fortunately, most of the Trinil [fossilized] bones have not suffered from corrosion, the effect of which on the bones, it should be noted, is completely different than that of [the abrasion which results from] transportation in a river bed. … [1242] The facts force us to accept that the bones are still at the very same place where they were deposited in fresh condition, before their fossilization and that they are located in their primary resting place [that is, the fossils are not reworked] ….

I imagine the events to have taken place about in the following manner … see my Note in … 1892, 2^nd^ Quarter … [section B(i), above]. From the nature of the bedding [likely referring to small-scale facies changes within the LB (e.g., S II-C7)], it is logical that the animals perished during volcanic eruptions in a manner similar to, but even more violently than, they took place often during ancient [historic] times in Java. The eruptions, to which we indirectly can attribute [the deaths and injuries represented by] the fossil bones, must at times have repeated themselves again and again, even if they [the deaths] all occurred in the same geologic period. Most of the bones in the richest discovery places [along the southern Kendeng Hills] such as near Trinil, Kedungbrubus and Bangle [east of Kedungbrubus], occur in a lapilli bed, which rests on a gray-black claystone, which is calcium-rich, and when dry, very crumbly, and contains fresh water mollusks …. One of such clay layers, which must have been deposited in very quietly flowing water, reaches a thickness of about 35 meters near Kedungbrubus, while near Trinil it [the claystone below the LB] is on average only 1 meter thick.

The lapilli bed signifies then, the beginning of the volcanic break-through [when the lake in the crater of an active volcano drains catastrophically] and animal carcasses [have been killed in the preceding volcanic events] must have been washed together exactly at those quiet places (where under normal circumstances clay was deposited). The main bonebed [Dubois used the word Hauptknochen-schicht, here] near Trinil was deposited in this manner. During times of diminishing volcanic activity, or after most animals had perished, the animal remains in the tuffs [volcaniclastic strata] at those times must have become less abundant. This explains the relative paucity of remains in the other beds of Trinil [such as Dubois’ “Sand-rock” strata; our units **2**-**5**, Figures 2a and 4]. The cadavers in those quiet places fell apart to some extent through decay, but much more likely they must have been ripped to pieces by crocodiles which often live in such places, preying on dead and living animals. The bones were then separated from each other and were broken many times. … In many cases I noticed tooth marks of these reptiles on the less fossilized bone fragments. Colossal crocodile teeth are, by the way, the most common fossils of Trinil.

The fact that hundreds of antlers of the same deer species … have been found [1243] near Trinil, is simply explained by the simultaneous extermination of the entire herd of these *Axis*-like [*Axis lydekkeri*] deer (It is well known that the currently living *Axis* species also builds large herds), and were swept together at these tranquil places. Precisely these antlers which are not covered by flesh are left mostly untouched by the crocodiles, while the other bones were almost always broken, as they are in fact encountered. It must be pointed out here in particular, that many shed antlers were also found. This can be explained by the fact that these quiet locations in the river were also used as watering places.

### Endnote H. Paleobotantical materials from Trinil

i. [Dubois’ August 1891 memorandum, (Endnote A(ii) and A(iii), above, S II-B2b, -B2d and -B2e] … fossilized wood [and] imprints of leaves [occurred in the **LB** of the Skullcap Pit, where] the tree **trunks** and leaves are always found horizontally [and the remains were] similar to wood from a European bog.
ii. [Kriele’s August 27, 1893, letter, Endnote A(v), above, S II-A3c] … much wood [was present the **PFZ** in the 40-m Trench, as it was in the **LB** of 1896 Left-bank Pit [Endnote A(iv), above, S II -A4c].
iii. [Carthaus (1911a: 14, Berkhout and Huffman 2021: 72-73) observed that there were] woody remains … in the main bonebed [included] tree **trunks and branches** up to 1- or 3-meters long. [Dozy (1908: 609) noted that] … wood, fossilized in various manners, and carbonized wood occur … in both Pits [presumably including the **HK** of Pit II, where the Main Leaf Bed was not well represented; see viii, below].
iv. [Schuster (1911a: 243, 244, 246) reported that] the fossilized reed grass of Cyperus … [was] abundant in the Hauptknochenschicht [**HK**, which also had some remains of a Sterculiacean tree and an evergreen wood-apple tree, as well as (1911b) silicified wood of another evergreen tree and a tree-like evergreen shrub]. [Regarding herbaceous remains, he (1911a: 244; Berkhout and Huffman 2021: 205) further reported that] the fossilized reed grass of *Cyperus* is not only abundant in the Hauptknochenschicht and the main leaf bed, but is also present in the intervening clay and ash layer [the Cyperaceae (sedge-) grounds implicated by these remains might have been in fully open vegetation or forest understories].
v. [Schuster (1911a: 243) saw the fossils in the main leaf bed, encountered primarily in the right-bank Pit I, as representing evergreen trees (such as those growing historically in the region), and shrubs, deciduous leaf fragments, some epiphytes and Cyperaceae/*Cyperus* sp., and concluded that] the plants of Trinil belonged to an open tropical forest almost devoid of liana. [Flenley (1979: 94) affirmed that “the [leaf] assemblage … are mainly lowland rain forest species,” [probably having drawn this conclusion from Shuster’s illustrations and descriptions; see also, Morley et al. 2020: 577, “wet lowland forests”].
vi. [Sémah (1986: 121; also, 1984, and Sémah et al. 2016) inferred a] open [as opposed to a closed canopy forest since] the only quantitatively important taxa [in her palynological samples] are Poaceae [grasses], Cyperaceae [sedges] and ferns.
vii. [Polhaupessy (1990, 2002, 2006) analyzed three claystone samples in the Trinil discovery area and one sample was evidently collected just below the **HK** along the left bank. Herbaceous taxa dominated her recovery with Poaceae (= Gramineae) being the principal component and Cyperaceae a prominent second. Polhaupessy also identified 22 arboreal taxa, including nine of large-size forest trees and montane elements, such as *Podocarpus* (a prominent component in the “oak-laurel” lower-montane forests of Sundaland and Indochina of Morley 2018: 478). Samples from 3.5-4.5 m above the **HK** (at approximately the level of the *Stegodon* bonebed in Selenka Pit II and the main leaf bed in Pit I) had more dryland arboreal constituents than did in claystone immediately below the HK). Five claystone samples from the Pucangan Formation, recovered Poaceae pollen more frequently than was the case stratigraphically higher. She (2002: 91-92) inferred a “widespread occurrence of grass-dominated swamps within a fluvial setting, and also possibly the occurrence of savanna grassland” in “a markedly seasonal climate,” suggesting a shift in paleoclimate from drier conditions during Pucangan deposition towards wetter conditions at the time of the HK.]
viii. [Notes about the lithofacies of the Main Leaf Bed: As digging progressed in 1907 Pit I, there were “thin clay layers … here and there with leaf imprints and partially carbonized wood …. [inferred to be in] a fluvial deposit” (F(iii), above); the plant-rich interval in 1908 was a “complex [intercalation] of blue-grey ash … with … clay[stone] … and many plant remains” spanning ∼3.75m of stratigraphic thickness and lying as low as ∼0.35 m above the **HK** (Selenka and Blanckenhorn 1911: Tafel X, Berkhout and Huffman 2021: 49). In total, the “main Leaf Bed…. [contained] rapidly thinning beds [lenses] … of plant material … most … in …. augite-andesite tuff with mainly green- and brown-hornblende [phenocrysts]” (Schuster 1911a: 4).

### Endnote I. Ecological notes on selected vertebrate species from the main bonebed

(i) *<u>Axis lydekkeri</u>*: The main bonebed had largest assemblage of *Axis lydekkeri* skeletal remains known from a single deposit. Stable carbon isotope data from six isolated *A. lydekkeri* teeth from the current museum collection fall tightly within a δ13C range generally attributable to a “pure C4 diet” and reportedly showed minimal seasonal variability (Janssen 2017, Janssen et al. 2016: 150). *A. lydekkeri* appeared first in Java before the earliest-known *Homo erectus* fossils and then continued as descendent subspecies through the time that the Ngandong *Homo erectus* lived (Table 5), when the deer had somewhat different antler morphology and a larger body size (de Vos 1996a, Gruwier et al. 2015, Leinders et al. 1985, Sondaar 1984, van den Bergh et al. 1999, 2001). *Axis* is also the most numerous identified taxa of the Perning *Homo erectus* discovery bed (Tables 4 and 5-E), so that Lydekker’s deer inhabited the Mojokerto paleo-delta when dry grasslands with few dryland trees dominated delta plain and associated river valley (Morley et al. 2020). The deer was the most common ungulate fossil in swamp deposits that formed under wet climatic conditions at Sangiran Dome (Bukuran bonebed, Table 5-F).

*Axis lydekkeri* is probably phylogenetically related to extant hyelaphid deer (Gruwier et al. 2015; also, Gruwoer 2019). The hyelaphid *A. porcinus* (Hog Deer) lived historically in Pakistan, India, Sri Lanka, Indochina and southwestern Yunnan, China (e.g., Duckworth et al. 2015, Lekagul and McNeely 1988). “Hog Deer has long been known in Myanmar … and occurred throughout the country wherever there were grassy plains …. In the southwestern coastal lowlands of Cambodia, where apparently the species was once common …, the species appears to use an open habitat mosaic including brackish … sedge marshes and ‘upland’ tall … grasslands, and areas of scrubby open secondary woodland interspersed with ‘dry’ short stature grasslands …. Hog Deer is a primarily a grazer of young grasses…; it also takes herbs, flowers, fruits and browse” (Timmins et al. 2015: 8, 10, 11).

The largest and most-thoroughly investigated native populations of *Axis porcinus* live along foot of the Himalaya, most notably in the Kaziranga National Park (KNP), 430 km^2^ located in northeastern India along the Brahmaputra River, its plains and nearby hills. The prime area of deer occupation is dominated by tall- and short-grasslands, wetlands and mixed-deciduous forests (e.g., Bradley-Martin and Vigne 1989, Choudhury 2010, Rahmani et al. 2016, Bajariu 2016). KNP averages ∼ 2200mm of precipitation per year, but has a severe dry season marked by months of <100mm in rainfall. In recent decades, KNP has been known to support 5045 *A. porcinis* (1999 census; and perhaps triple this number), and 1666 *Bubalus arnee* (in 2002; Wild Asian Water Buffalo), 1206 *Elephas maximus* (in 2005), 468 *Cervus duvaucelli* (in 2002; Swamp Deer), 431 *Sus scrofa* (in 1999), 100 *Muntjacs muntjac* (in 1972), and 86 *Panthera tigris* (in 2000), together with many other large- and small-terrestrial animals, birds and aquatic species, including Gharial. Floods periodically inundate 70-80% of KNP, drowning as many as a thousand Hog Deer in each event.

The living hyelaphid species nearest geographically to Java’s former Lydekker’s deer population is *Axis kulhii*, which potentially was a descendant of *A. lydekkeri. A. kulhii* inhabits the still-forested Bawean Island (∼192km^2^, 0-646 m elevation) in the middle of the Java Sea ∼150km north of eastern Java (Blouch and Atmosoedirdjo 1978, 1987, Hoogerwerf 1966, 1967; also, Rademaker et al. 2016). *A. kuhlii*, which is primarily a nocturnal feeder, “grazes on herbs and grasses, but also browses young leaves and twigs” (Blouch and Atmosoedirdjo 1978, Semiadi et al. 2015: 4). The second related extant island species, *A. calamianensis*, lived in parts of the Palawan archipelago, southeastern Philippines (Widmann and Lastica 2015). *Axis* population evidently passed through Borneo at points in the geological past. No *Axis* populations are known elsewhere in Sundaland or farther eastward into the Philippine archipelago, Sulawesi or Nusa Tenggara, all of which are all separated from Sundaland by deep oceans. Modern hyelaphid appear to generally prefer mosaic landscapes with water, grass and other palatable plants, as well as safety in forests, scrub or tall grasslands.

(ii) <u>Large-body bovids, *Bibos palaesondaicus* and *Bubalus palaeokerab*</u>*au*: A grass-prone diet is seen in the δ13C results from several dozen isolated large-bovid teeth from Trinil (reported as *Bubalus palaeokerabau* and species undetermined; Janssen 2017, Janssen et al. 2016; modern Southeast Asian bovids were not sampled). However, when Weinand (2005: v and 81; also, 2007) analyzed the astragali metrics for various Southeast Asian bovids, including 81 large-body specimens from Trinil, he concluded that the bovids were anatomically adapted more for *“heavy cover”* of *“densely vegetated river valleys and upland forests, broken by open grasslands.”* The large-sized bovid species from the main bonebed are anatomically similar to (and often inferred as having been the ancestors of) extant *Bibos javanicus* (Banteng cattle) and *Bubalus arnee* (Wild Asian Water Buffalo), respectively (Dubois 1908, Hooijer 1958a). Amano et al. (2016b: 158; also, 2016a) observed mesowear- and microwear-patterns on the molars of *Bibos javanicus* that reflect both grazing (*“feed on grass”*) and browsing (*“feed on dicotyledonous plants”*).

Present-day and historic natural populations of Banteng inhabit open- and mixed-forests, from coasts to mountain highlands, and in some cases, rely upon the browsing on herbs and bark. The same is quite plausibly inferred for *Bibos palaesondaicus*. In Java, human settlement drove Bantengs into remote areas, forced them into suboptimal habitats, gave rise to domestication (Bali Cattle), and led to Banteng introductions into non-native habitats, far and wide (in part from S. Hedges, pers. comm., 2018). In the historic past, Banteng lived *“very frequently* [from] *… mountain forests* [to] *… flat-lying jungles along the coasts, especially near marshy lakes, gently streaming rivers or … widening of mountain valleys”* (Encyclopaedie van Nederlandsch-Indie 1895-1905). *Bos javanicus* inhabited Java’s ever-wet westernmost tip to its easternmost seasonally dry end. Banteng are well known from Ujung Kulon National Park with ∼3000-mm rainfall per year and Baluran National Park with ∼1600-mm annually, on opposite end points of Java (Whitten et al. 1996). The essentials for Banteng were forests, grazing land, fresh water and sources of salt. No *B. javanicus* is known to have dispersed eastward across narrow seaways leading from Bali towards the drier islands of Nusa Tenggara archipelago or the wetter landscapes of Sulawesi and the Philippines.

Banteng herds native to historic Java, Borneo and Indochina are sometimes assigned to separate geographic subspecies, such as *Bos javanicus birmanicus* in Cambodia (Matsubayashi et al. 2014). Banteng were common historically in south and east Borneo, grazing and browsing in forested landscapes (for example, Banteng occur in 5 of 15 orangutan protected areas; Gardner et al. 2014, 2016, 2021, Matsubayashi et al. 2014, McKinnon et al. 1996). At Niah Cave, Sarwak, northern Borneo, Pleistocene cave deposits yielded *Bos* remains along with those *Pongo* and other forest-dwelling genera (also, Table 5, footnote 6, concerning *Bos* in cave assemblages from the Sumatran highland).

Banteng were formerly widespread in wet central Thailand, including its forested western and northern mountains (Corbet and Hill 1992, Humphrey and Bain 1990, Lekagul and McNeely 1988). Around 250 of these cattle still reside in the forests of one mountain sanctuary in Thailand, where the animals browse as needed on shrubs, herbs and tree bark in the dry season, for example, and co-exist with the Gaur (an oxen), Asian Wild Buffalo, Sambar (*Rusa deer*), Sumatran Serow, muntjacs, Clouded Leopard, Sun Bear, and Tiger, among other forest species (Prayurasiddhi 1997). Banteng is one of two large-bodied prey species of the robust tiger population there (Pakpien et al. 2017). Thousands of wild *Bos javanicus* also thrive primarily on grasses in the dry-deciduous forests in eastern Cambodia (Duckworth et al. 1999, Gray and Phan 2011, Gray et al. 2012, 2016, Nguyen 2009, Steinmetz 2004).

An inadvertent experiment in Banteng-foraging adaptability was conducted in NW Australia. Over 6000 free-ranging pure-strain Banteng inhabit open eucalypt-forest on the Cobourg Peninsula (11.3°S, 132.2°E), an area with ∼1300mm average annual rainfall. Set free there 170 years ago in the absence of large terrestrial predators, the small introduced population expanded numerically to overspread ∼70km^-3^ (1988) by shifting their diet to primarily sedges, trees and shrubs, with peak grass consumption now being ∼40% in the late wet season (Bowman et al. 2010, Calaby 1975, Choquenot 1993, Corbett 1995, De Konnick 2014).

*<u>Bubalus palaeok</u>erabau*, like *<u>Bibos palaeson</u>daicus* are Bovidae-Bovini, probably of the subtribe Bubalina (e.g., Castelló 2016, Grubb 2005, Hassanin 2014). *Bubalus arnee* and the domesticate *Bubalus bubalis* (Asian Water Buffalo) have habitat tolerances exceeding what one might envision for them, although one must go farther afield from Java to demonstrate this. The largest present-day Wild Asian Water Buffalo population occurs in Assam, India, (Choudhury 2010, 2017), particularly in Kaziranga National Park (KNP) along the Brahmaputra River (Mahanta et al. 2016, Bajaru 2016). Present-day herds there are only a small remnant of broad South Asian populations that existed in the region thousands of years ago. Some 1963 buffalo were counted at KNP in 2008, when the Buffalo primarily frequented dense grasslands and open-seasonal forests that are subject to high-levels of wet-season rainfall and annual floods, as well as severe dry months. With densities of 6.45km^-2^ the Buffalo are closely tied to ample grass forage and freshwater bodies, which the buffalo use frequently for wallowing (sources cited under *Axis*, above).

During the Holocene, Wild Asian Water Buffalo (*Bubalus arnee*) were not restricted to grassy lowlands but inhabited both major river valleys and the adjacent mountains in Indochina (e.g., Bacon et al. 2018, Hedges et al. 2008; also, Kaul et al. 2019). A montane remnant numbering >40 individuals is still to be found along the grassy river banks and in adjacent deciduous forests in ∼70km^2^ of the same Thai sanctuary with the Banteng mentioned above (Chaiyarat 2002, Chaiyarat et al. 2004). Feral Asian Water Buffaloes share the Baluran National Park, Java, with Banteng, and appear to be less tolerant of browsing there than are the wild cattle (S. Hedges, pers. comm., 2018). Feral buffalo also inhabit savanna lands of very dry Komodo islands (∼800 mm per year rainfall), near Flores in the Nusa Tenggara archipelago.

However, the most remarkable feral *Bubalus bubalis* occurrence is in NW Australia, as was the case with Banteng, where Asian Water Buffaloes introduced in the mid-19^th^ Century now number hundreds of thousands. Grazing and subsidiary browsing (they do consume relatively more grass than do the feral cattle) and in the absence of large terrestrial predators, the Asian Water Buffaloes have spread over hundreds of thousands of square kilometers, inhabiting a far broader range than the Banteng, and exhibiting astounding ecological flexibility in that dry climate (Bowman et al. 2010, Corbett 1995, Freeland and Choquenot 1990, Petty et al. 2007, Saalfeld 2014).

Broad Pleistocene dispersal and adaptation by speciation in *Bubalus* is evident in forest *Bubalus* spp. known from islands east of Sundaland. *Bubalus palaeokerabau* is full-size when compared to modern *B. arnee*, but dwarf buffaloes occur on islands beyond the limits of Sundaland (Burton et al. 2005, 2016, Rozzi 2017). *Bubalus mindorensis* (tamaraw) was formerly widespread on the 9,735 km^2^ Philippine island of Mindoro, where the species ranged from “sea level to the high peaks (to over 1,800 m), inhabiting open grassland or forest glades, thick bamboo-jungle, marshy river valleys, and low to mid-elevation forests” (Boyles et al. 2016: 4, Cebrian et al. 2014, Custodio et al. 1996.; also, Croft et al.2006, Huffman, B., 2016).

Smaller miniaturized forest buffalo occur on Sulawesi (181,000km^2^), *Bubalus depressicornis* (Lowland Anoa) and *B. quarlesi* (Mountain Anoa, which is the smaller, more antelope-like; Burton et al. 2005, 2016, Rozzi 2017). The dwarf species indicate Pleistocene *Bubalus* populations were present in eastern Sundaland, and unlike Banteng, succeed in crossing ocean barriers that separated Sulawesi and the Philippines from Borneo while evolving into miniaturized species. The dispersal small-statured buffalo to islands peripheral to Sundaland, together with the ecological range of modern *Bubalus arnee*, support the notion that *Bubalus* spp. were widely distributed between mainland Indochina and Java during the Pleistocene. This might indicate that *Bubalus palaeokerabau* potentially was more adaptable to varied paleoenvironmental conditions than was *Bibos palaesondaicus*.

(iii) *<u>Duboisia santeng</u>* (Dubois, 1891): Dubois’ Antelope, is a boselaphin notable for its small size, unique horn-core anatomy, and concentration in the *S.-H.e.*, especially in the main bonebed (e.g., van den Bergh 1988). In reviewing various evidence concerning the *D. santeng* from Trinil, Rozzi et al. (2013; also, 2014) related the mesowear properties of the teeth to those of “a forest dweller living in close canopy settings.” Janssen et al. (2016), on the other hand, obtained a C4 signal from three Trinil samples of *Duboisia* enamel, similar to the results obtained from more numerous specimens of other ungulate species from the site. One fossil calotte with intact horn cores and cranial dimensions similar to the Java specimens has been described as *Duboisia* aff. *D. santeng* from a fluvial site in the Khorat Plateau, Thailand (Nishioka and Vidthayanono 2018). *D. santeng* was only slightly more massive than diminutive deer, *Axis lydekkeri* (Hooijer 1958, van den Bergh 1988). The species’ estimated body mass of ∼54kg is an ∼70% size reduction from *Boselaphus namadicus*, an extinct boselaphin from the Middle Pleistocene Siwalik Group (Pakistan; Rozzi and Palombo 2014, Rozzi et al. 2013). *Duboisia* is sometimes linked phylogenetically to the Indian Nilgai (Blue Cow), *Boselaphus tragocamelus*, one of the two extant Asian boselaphins. *B. tragocamelus* males weigh ∼250 kg; the species are mixed feeders in small groups across South Asian ecological zones but nowhere closer to Java (Sankar and Goyal 2004). Imported Nilgai have thrived on grass in the dry cattle rangeland of south Texas since the 1930s.

(**iv**) *<u>Stegodon trig</u>onocephalus*: The main bonebed has the greatest concentration of *Stegodon trigonocephalus* (joining *Axis lydekkeri* is this distinction known), except possibly for the Ngandong *Homo erectus* bonebed (Table 5). *S. trigonocephalus* is member of the extinct family of generally large-bodied proboscideans that were widely distributed in Southeast Asia, East Asia and South Asia during the Pleistocene, so that *Stegodon* spp. might have occurred widely in Sundaland during this time period (e.g., Zeitoun et al. 2016). *Stegodon* spp. from islands east of Sundaland are dwarfed (e.g., Powley et al. 2021, van den Bergh 1999, 2001, 2019, van den Bergh et al. 2001, 2008, van der Geer et al. 2010, 2016). Presumably, like modern elephants, *S. trigonocephalus* lived in herds of related individuals. Evidence favoring a diet of C4 plants for East Java *Stegodon* comes from the stable-carbon-isotope signals in the enamel of three teeth, all isolated surface finds, that were collected from near the discovery site of the Mojokerto child skull in an area of the upper Pucangan Formation outcrop, 2.5km east of long-known discoveries of large-sized *S. trigonocephalus* specimens (Gondang, Cosijn 1931, 1932, Dubois Collection; see Huffman and Zaim 2003). The three specimens had δ13C values of -0.73 (hominin site) and -0.07 to -0.88 (Gondang), all within the range associated with C4 vegetation (with corresponding δ18O of -6.83 and -6.68 and –7.41; T. Cerling and J. Kappelman in Huffman and Zaim 2003).

(v) <u>Rhinoceros, Trinil pig and Trinil t</u>iger: Three prominent forest-prone large vertebrates occur in the main bonebed. Javan Rhinoceros had wide forest distribution in the prehistoric past, going back in approximately as long as any species of *Homo* inhabited Java. The Javan Rhinoceros (*Rhinoceros sondaicus*) occurs in the Trinil H.K., Kedung Brubus and Ngandong faunas, most of the *Stegodon*-*Homo erectus* faunal association (Tables 5). The teeth of *R. sondaicus* also commonly occur in the Holocene and Late Pleistocene rain-forest cave assemblages of Java, Sumatra, and northern Borneo (Niah Cave; Cranbrook and Piper 2007, Dammerman 1934, Aimi and Aziz 1985, Simanjuntak 2001, Storm and de Vos 2006, Storm et al. 2005, 2013, Westaway et al., 2007, 2017). *R. sondaicus* existed as subspecies in Java, Sumatra, Borneo (at least its northern portion), the western Malay Peninsula, Indochina (western mountains of Thailand, Cambodia, the Laos uplands, and Vietnam), and eastern South Asia, notably Assam and the Sundarbans delta. The teeth of the species are abundant (NISP = 238) in the Vietnam Coc Muoi cave, along with the teeth of tapir, cattle, *Sus* and serow, among others, in a late Middle Pleistocene rainforest paleoenvironment (MIS6-5 transition; Bacon et al. 2008b; also, Bacon et al. 2004-2008a). Older fossil occurrences are known from southern China, Myanmar and Pakistan. *R. philippinensis*, a related fossil species from Luzon dating to 709ka, demonstrates the capacity of the rhinoceros to migrate east of Sundaland (Ingicco et al. 2018).

The ecology of *R. sondaicus* is best known from Java, where the species lived historically in the rain-soaked, perhumid forests of West and Central provinces (Groves and Leslie 2011, Hoogerwerf 1970, van Strien et al. 2008). While principally a lowland-forest dweller, Javan Rhinoceros inhabited coastal forests and montane forests on the upper volcanic slopes. The species *“is a generalist browser and consumes little to no grass and few herbaceous species, preferring leaves, shoots, and twigs of woody species”* and is well-suited anatomically for reaching vegetation over its head, (Groves and Leslie 2011: 199). *R. sondaicus* is less specialized anatomically than the one-horned *R. indicus* (Indian Rhinoceros) of South Asia, Indochina, southern China (as *R. sinensis*). *R. indicus* (∼ *R. kendengindicus*) also occurs in the *S.-H.e.* of Java (Table 4). Modern *R. indicus* consumes more grass than does the Javan Rhinoceros.

The extinct Trinil Pig (*Sus brachygnathus*) has a MNI of 9 in the Dubois Collection (Table 4). The δ^13^C results for Trinil *S. brachygnathus* enamel are consistent with individuals living from of C3 vegetation and having the omnivorous behaviors normal to *Sus* (Janssen et al. 2016). *Sus* spp. probably originated in Southeast Asia, where they are widely distributed, and most inhabited forests historically (e.g., Melletti and Meijaard 2017). Judging from anatomical and genetic studies, *S. brachygnathus* is most closely related phylogenetically to the present-day Javan Warty Pig of Java and Sumatra, and the Bawean Warty Pig of the forested island in the Java Sea (e.g., Badoux 1959, Hardjasamita 1987, Rademaker et al. 2016, Semiadi et al. 2015).

Janssen et al. (2016) found that the enamel of a Trinil tiger specimen has carbon- and oxygen-isotopic values *“reflecting a food chain with a mixed C_3_/C_4_ base.”* The Trinil taxon is one of numerous subspecies of *Panthera tigris*. Of the South Asian tiger, Bhattarai (2011: 1) summarized, *“the basic habitat requirements … include the thick cover of forest, proximity to water and … an abundance of large and medium-sized prey”* (also, Chanchani et al. 2014). Tigers still inhabit Sumatra, where they prefer thick understory, and prey, by surprise and stealth, on medium- to large-ungulates and ground-dwelling primates, such pigs, muntjacs, deer and macaques (e.g., Goodrich et al. 2015, Linkie et al. 2008, Wibisono et al. 2011). A robust tiger population in a Thai mountain reservation preys on Banteng and another large-bodied bovid species (Pakpien et al. 2017). The Trinil dog *<u>Xenocyon trinilensis</u>* Stremme, 1911 is the only other large carnivore known from Trinil. Stremme (1911: 83-86; Tafel XVI, Fig. 1 and 2) named a partial left mandible with P3, M1 and M2 (Museum für Naturkunde, Berlin, specimen MB.Ma.28893) as the new canid species.

(vi) Low-but seemingly significant-frequency of other forest-prone species (muntjac, macaque, porcupine, leopard cat, langur and gibbon; Table 3) indicate a modest extent of forest in the area of ungulate death associated with the formation of the main bonebed. Although the fossils of rat, python and monitor lizard are components of the Trinil assemblage, hundreds of small- to medium-sized arboreal and ground-dwelling species, which have not identified among Trinil fossils, presumably also inhabited the watershed. Paleobotantical remains support this supposition.

(vii) Riverine species are common in the main bonebed. Aquatic reptile- and fish-specimens (NISP of 330 in the DC) are equal in numbers to *Duboisia*, *Rhinoceros* and *Sus* combined (NISP of 353), and fall between those of *Duboisia* and *Stegodon* (NISP of 499 and 231, respectively). Turtle shells and crocodylian skulls, some largely complete (S I Figure 20d), are prominent in the aquatic assemblage, as they are in field accounts (e.g., S II-B2d, -B2e, and -A4l; also, S II-A1m). *Crocodylus siamensis* (Siamese Crocodile) was historically widespread from Java, Borneo (Mahakam River) and Indochina (e.g., Bezuijen et al. 2012, Cox 2004, Das 2015, Griggs and Kirshner 2015, Han et al. 2015, Platt et al. 2006). The species concentrates in *“freshwater lakes, swamps and slow-moving rivers, from near sea level to an elevation of 600 meters”* in Cambodia, where reptilian populations still exist locally; the reptiles prefer water bodies with gentle banks in both open- and shaded-areas surrounded by forest (Han et al. 2015: 154; also, Daltry et al. 2003, Sitha et al. 2005). *<u>Gavialis bengawanicus</u>* Dubois 1908 is closely similar in morphology to *G. gangeticus*, the Gharial, of present-day South Asia and westernmost Indochina, and resembles a fossil of a narrow-nosed crocodylian (cf. *Gavialis*) from Sulawesi and *Gavialis* cf. *bengawanicus* from northeastern Thailand (Delfino and de Vos 2010; also, Das 2015). *G. gangeticus* inhabits deep dry-season pools and wet-season inundated areas of large rivers (Saikia 2012; also, Choudhury et al. 2007; also, Lang et al. 2019).

