## Supplementary material for "Geology and discovery record of the Trinil *Pithecanthropus erectus* site, Java": S I

**SUPPLEMENTARY MATERIALS I FIGURES (S I Figures).** Additional Trinil photographs and maps for "Geology and discovery record of the Trinil *Pithecanthropus erectus* site, Java"

O. Frank Huffman

Department of Anthropology, University of Texas at Austin, SAC 4.102, 2201 Speedway Stop C3200, Austin TX 78712, USA.

Aart W.J. Berkhout

15706 Nedra Way, Dallas, TX 75248, USA.

Paul C.H. Albers

Naturalis Biodiversity Center, Darwinweg 2 2333 CR Leiden, The Netherlands.

John de Vos

Naturalis Biodiversity Center, Darwinweg 2 2333 CR Leiden, The Netherlands.

Fachroel Aziz

Geological Research and Development Centre, Jalan Diponegoro 57, Bandung 40122, Indonesia.

SUPPLEMENTARY MATERIALS I FIGURES

**S I Figure 1.** *Pithecanthropus erectus* Skullcap … and … Femur I
**S I Figure 2.** Excavation-scars and -baulks … along the left bank … during low-water level
**S I Figure 3.** Dubois' annotations on unpublished prints of the 1894 photograph
**S I Figure 4.** High-resolution scanning of Dubois' 1894 photograph … former back wall of the 1893
**S I Figure 5.** …photograph taken in 1900 … shows … 1900 Trench
**S I Figure 6.** … third 1900 photograph … shows the main bonebed (LB-HK)
**S I Figure 7.** The topographic features of the Trinil area in 1900
**S I Figure 8.** Selenka Expedition personnel readily identified the *Pithecanthropus erectus* bonebed
**S I Figure 9.** Selenka's excavators encountered hard-to-dig strata
**S I Figure 10.** The proboscidean remains of the *Stegodon* bonebed (SB)
**S I Figure 11.** … Pit II … channelized unit 6 …. The strata above the HK in the right-bank Pit I
**S I Figure 12.** The high back walls of the 1907 Pit II … in 1908.
**S I Figure 13.** … Pit I encountered flat-lying, well-indurated volcaniclastic and muddy strata
**S I Figure 14.** … strata above the main bonebed … on the right bank
**S I Figure 15.** W.F.F. Oppenoorth … lantern slides from 1932 … Trinil excavation
**S I Figure 16.** Duyfjes' (1936) geological map of the Trinil area
**S I Figure 17.** Three versions Duyfjes' geological mapping
**S I Figure 18.** A geological map of the left bank published at 1:250
**S I Figure 19.** Geological mapping done along the Solo River valley by multiple field teams
**S I Figure 20.** Trinil fossils in the Dubois Collection.
**S I Figure 21.** … geological illustration of the Kedungbrubus- and Butak-area
**S I Figure 22.** Southern Sundaland during the Pleistocene
**S I Figure 23.** … southern Sundaland when sea level was significantly below the present
**S I Figure 24.** Seismic geomorphologic patterns in Pleistocene strata … reveal … paleo-landscapes
REFERENCES

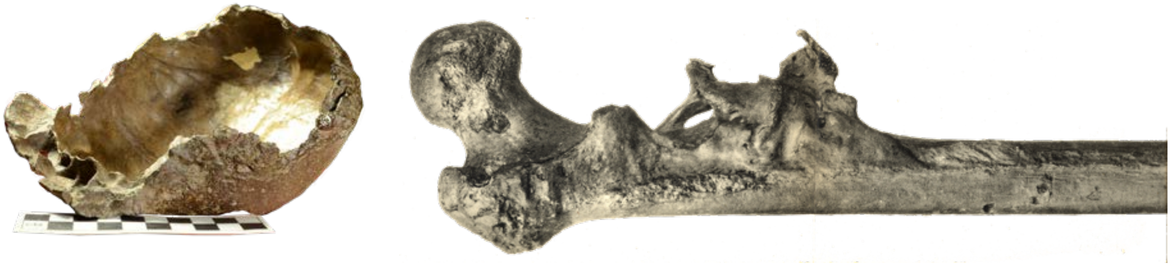

**Supplementary Materials I Figure 1 (S I Figure 1).** *Pithecanthropus erectus* Skullcap (Trinil 1, left) and proximal portion of Femur I (Trinil 3, right; Huffman et al. 2015). The Skullcap photograph was taken at the Naturalis Biodiversity Center, Leiden, by the authors. The image of the femur is colorized version from a Dubois (1926b: Figure 1) photograph (S II F9; also, S II F8).

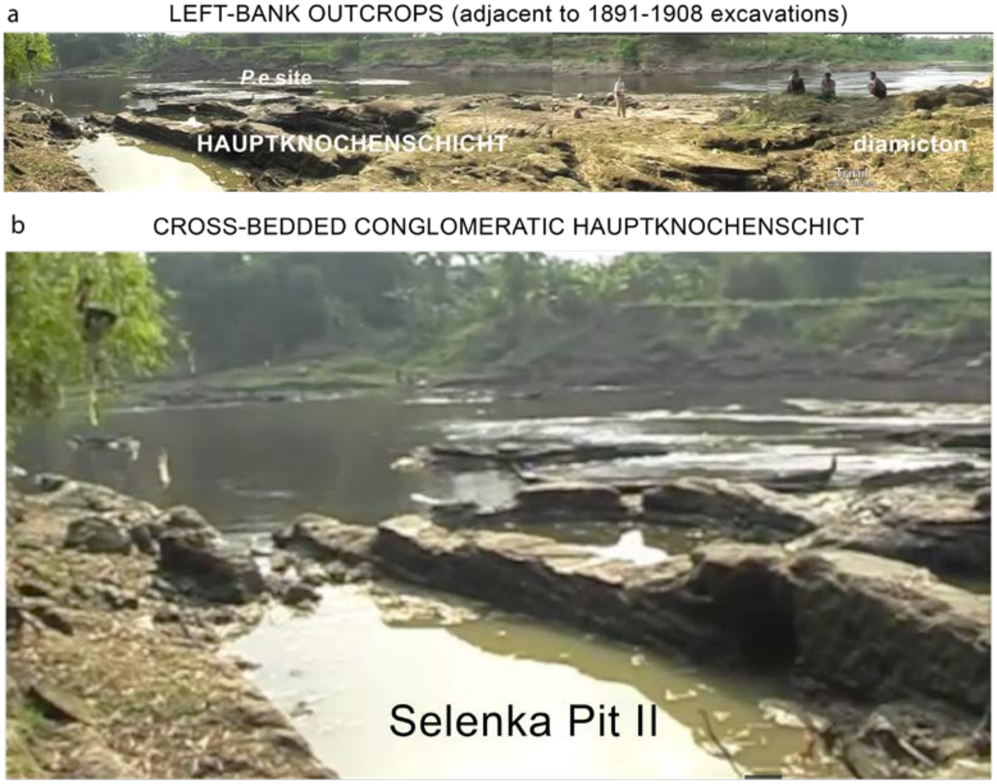


**Supplementary Materials I Figure 2 (S I Figure 2).** (**a**) Excavation-scars and -baulks are prominently exposed along the left bank of the Solo during dry season periods of low-water level. This 2008 view looks westward across the Selenka Expedition Pit II (1907-1908) towards a mid-river gravel bar which partially covers the former 1891-1892 *Pithecanthropus erectus* (*P.e.*) discovery site. (**b**) An enlarged portion of 'a' highlights cross-bedded conglomeratic sandstone in the main bonebed, the Hauptknochenschicht of Selenka and Blanckenhorn (1911). From Dubois' time in Java onwards, geological studies have treated the *P.e.* bonebed and superjacent strata as bedrock (e.g., van Es 1931, Duyfjes 1936, Soeradi et al. 1985, and I.J.J.S.T. 1992). Scores of site photographs and first-hand accounts from the period of excavation substantiate the lithified condition and flat-lying attitude of the eight-to-nine meters of volcaniclastic strata that were excavated between the main bonebed and soil of the terrace surface (de Vos and Aziz 1989; Huffman et al. 2015, 2018). This terrace lies ~20 m below the uplands of the river valley (e.g., S I Figure 7a). Source is the YouTube video "Trinil 3" of Chris Turney (posted April 28, 2008; downloaded 01/20/2014). The authors examined the site in 2016 (Huffman 2016), 2017 and 2018 (PCHA).

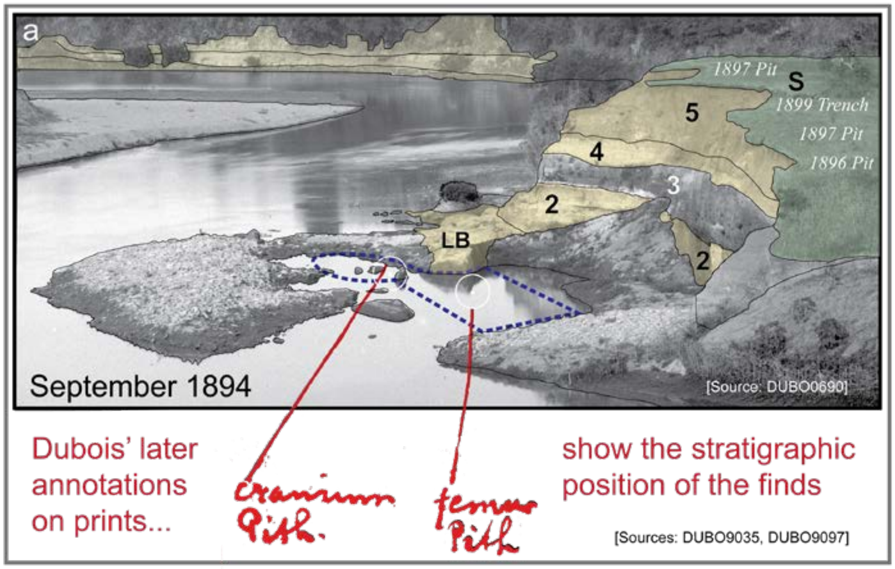


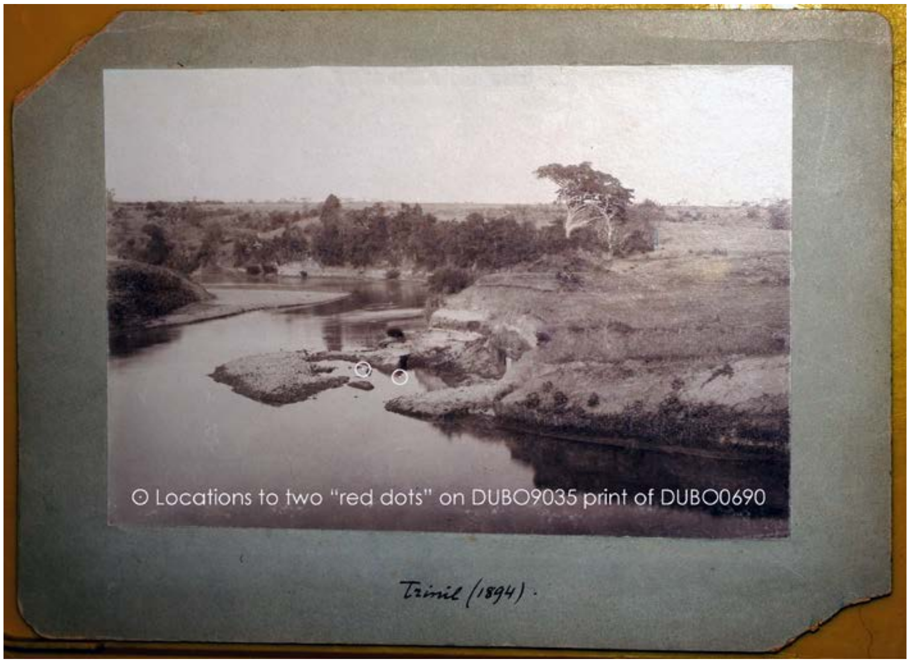


**Supplementary Materials I Figure 3 a, b (S I Figure 3a, b).** Dubois' annotations on unpublished prints of the 1894 photograph situate the *Pithecanthropus erectus* discovery location relative to nearby geography and geology (Huffman et al. 2015, 2018). (**a**) A summary of Dubois' annotations (Figures 3c and 4a, main text) and our interpretations of them tie the discovery site to the stratigraphic units we recognize in the nearby embankment and later Dubois excavations, as detailed in S I Figure 4 (also, Figures 3 and 4, main text). (**b**) One of the annotated prints has Dubois' handwritten inscription *"Trinil (1894)"* and two dots at the discovery points. Another print has markings that highlight the discovery pits and trenches, and shows the placement of the *"cranium Pith."* and *"femur Pith."* The back side of a third print has the Dubois note, *"Find spot of* P. erectus *viewed from near the pillar Trinil September 5 1894,"* the pillar being the Dubois monument at the present-day Trinil Museum (code for the annotated print is DUBO3286). (**S I Figure 3c** and **d**, next page).

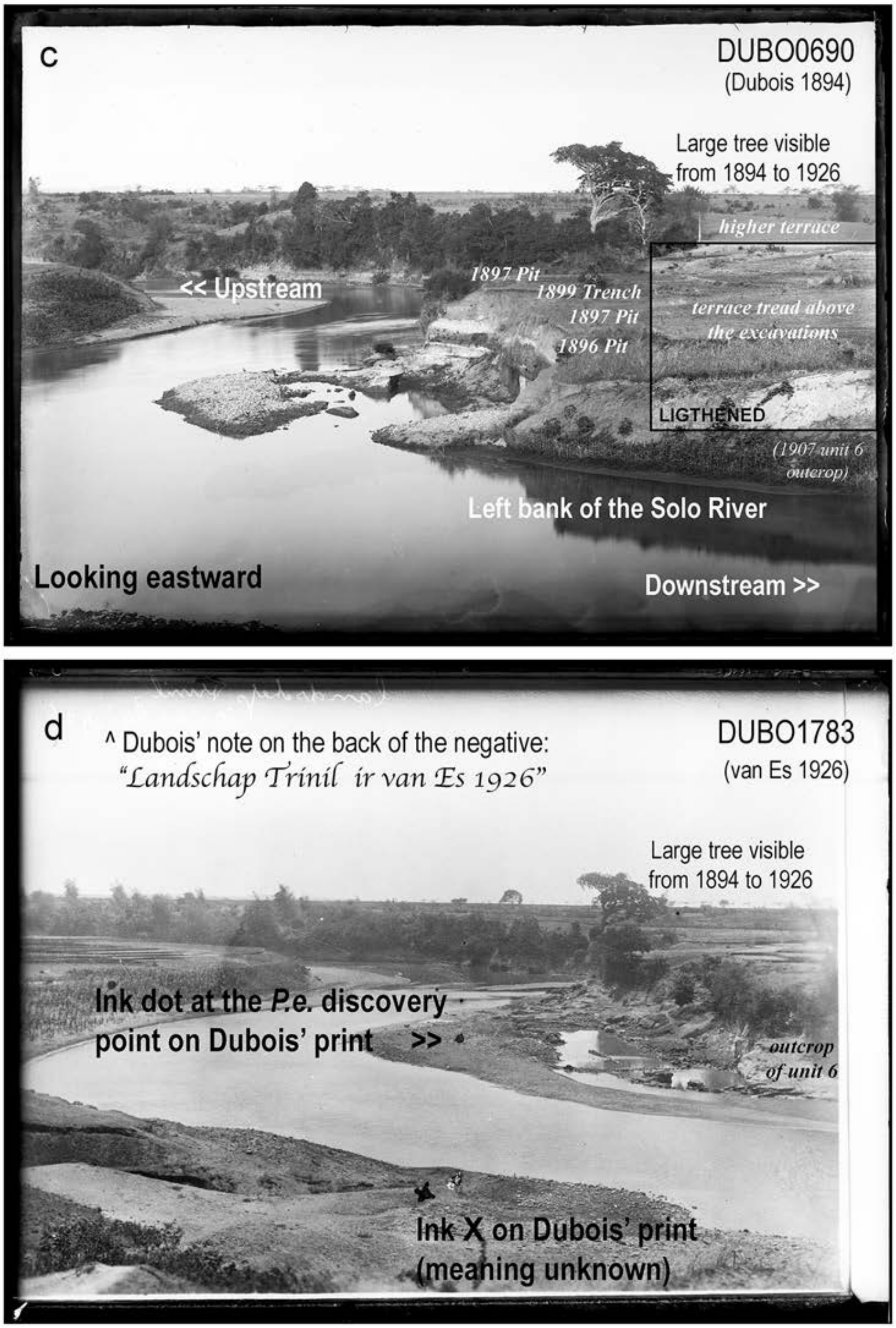


**Supplementary Materials I Figure 3 c, d (S I Figure 3c, d).** Our interpretation of the annotated 1894 image (c) and a 1926 photograph (d), both looking eastward from about the same camera station (also, Figure 9, main text).

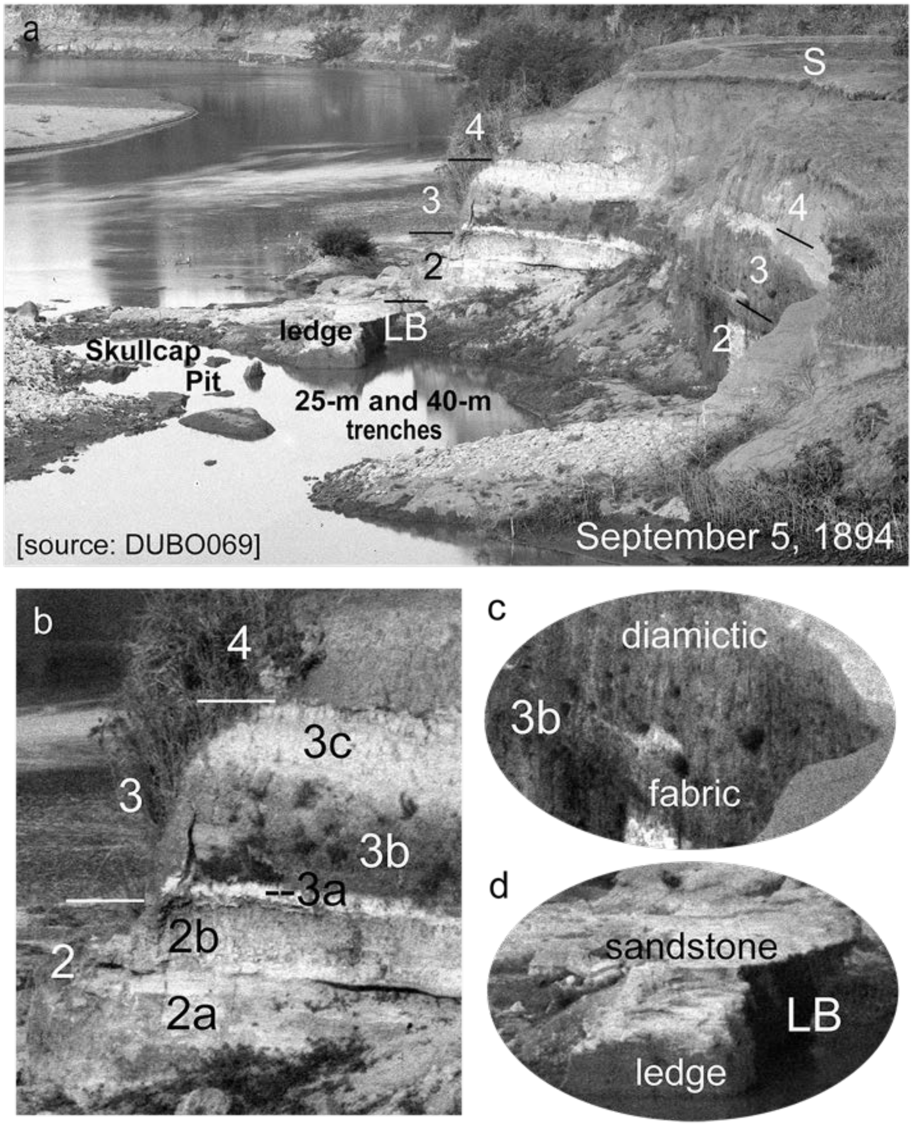


**Supplementary Materials I Figure 4 (S I Figure 4).** High-resolution scanning of Dubois' 1894 photograph allows the beds in the former back wall of the 1893 40-m Trench to be divided into useful stratigraphic units, **LB** (**1**) and **2-4**, as presented in Figures 3c and 4a, main text (also, Huffman et al. 2015, 2018). (**a**) The 1894 image looks eastward from the right bank (S I Figure 3b). The inundated excavation around the discovery points (S I Figure 3a) has a shape fitting the combination of the Skullcap Pit, 25-m trench and 40-m trench (Figure 3a, main text). (**b** and **c**) Unit **2** in the embankment includes light-colored subunits. The upper one is notable for a prominent depositional base. The middle of unit **3** consists of a dark-colored subunit exhibiting an apparent diamictic sedimentary fabric, which probably represents lahar deposition. The lighter-colored subunits above and below this diamictic unit might have been rich in volcanic ash. Volcanic diamictons are often termed "breccia," "tuff breccia," "conglomeratic tuff" or "boulder tuff" by geologists working on the Neogene in Java, and the diamictons have depositional lithologies characterized by extreme ranges in clast sizes, and often have boulders in a sand-mud matrix. (**d**) The rock ledge beside the Pit appears to consist of indurated sandstone that would have been at the top of Dubois' Bed of Lapilli (**LB**; Figure 2a, main text).

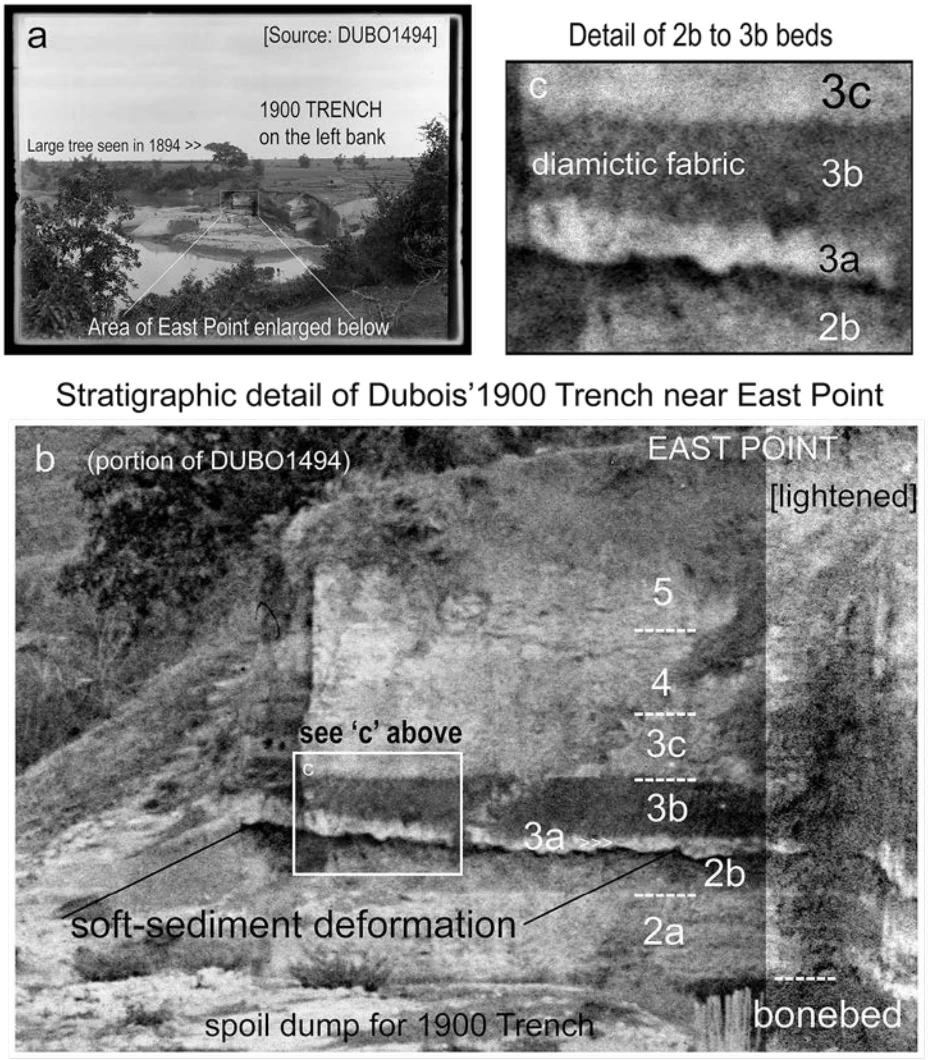


**Supplementary Materials I Figure 5 (S I Figure 5).** This photograph taken in 1900 from Camera station II (S I Figure 7) shows that the stratigraphic units recognized in the 1894 embankment extended into the 1900 Trench. Its backwalls near East Point particularly reveal stratigraphic and sedimentological detail useful to the correlation. The units in the 1900 Trench lie at about the same elevation (relative to seasonal low-water levels) as they did in the 1894 embankment, demonstrating that the units in the 1892-1900 excavations were essentially flat lying. The stratigraphic continuity between the excavated sequences is clearest for unit **3**, largely due to the distinctive diamictic fabric of its middle subunit (**3b**) and light-colored basal subunit (**3a**). The soft-sediment deformation features, such as those in **3a**, reflect the accumulation atop an unconsolidated substrate. When this photograph was taken in November 1900, the excavators were unearthing the main bonebed (**LB-HK**) in the western part of the Trench (Figure 4a, main text). The camera station was a top of the right bank north of the Dubois monument (station II in S I Figure 7). Negative of three 1900 photographs were scanned at 4800 dots-per-inch resolution for our use from negatives by Naturalis Biodiversity Center.

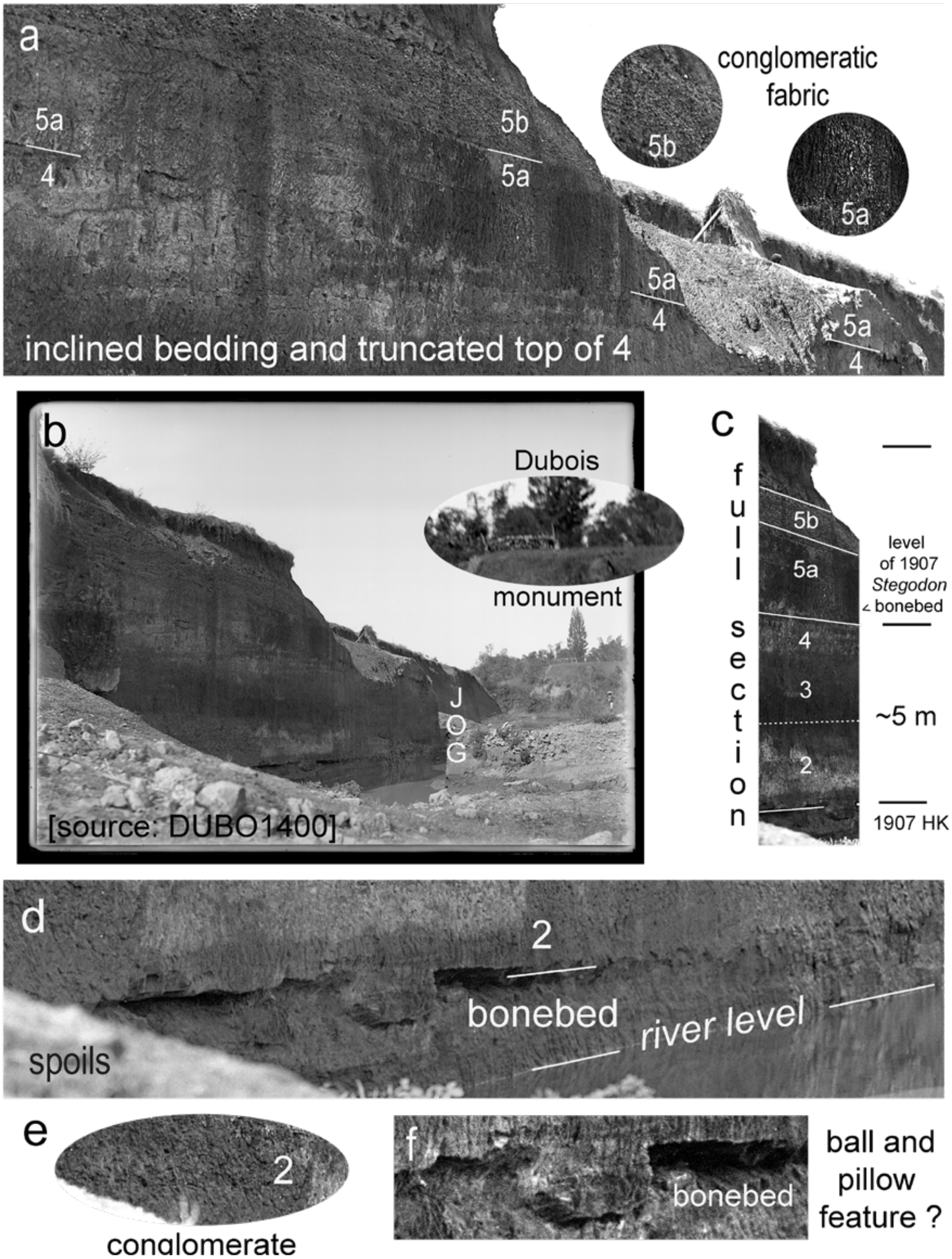


**Supplementary Materials I Figure 6 (S I Figure 6).** The third 1900 photograph (taken from Camera station I, S I Figure 7) shows the main bonebed (**LB-HK**) when freshly excavated and the common occurrence of conglomeratic fabrics in the overlying bed. The Dubois monument is seen on top of the right-bank in the distance.

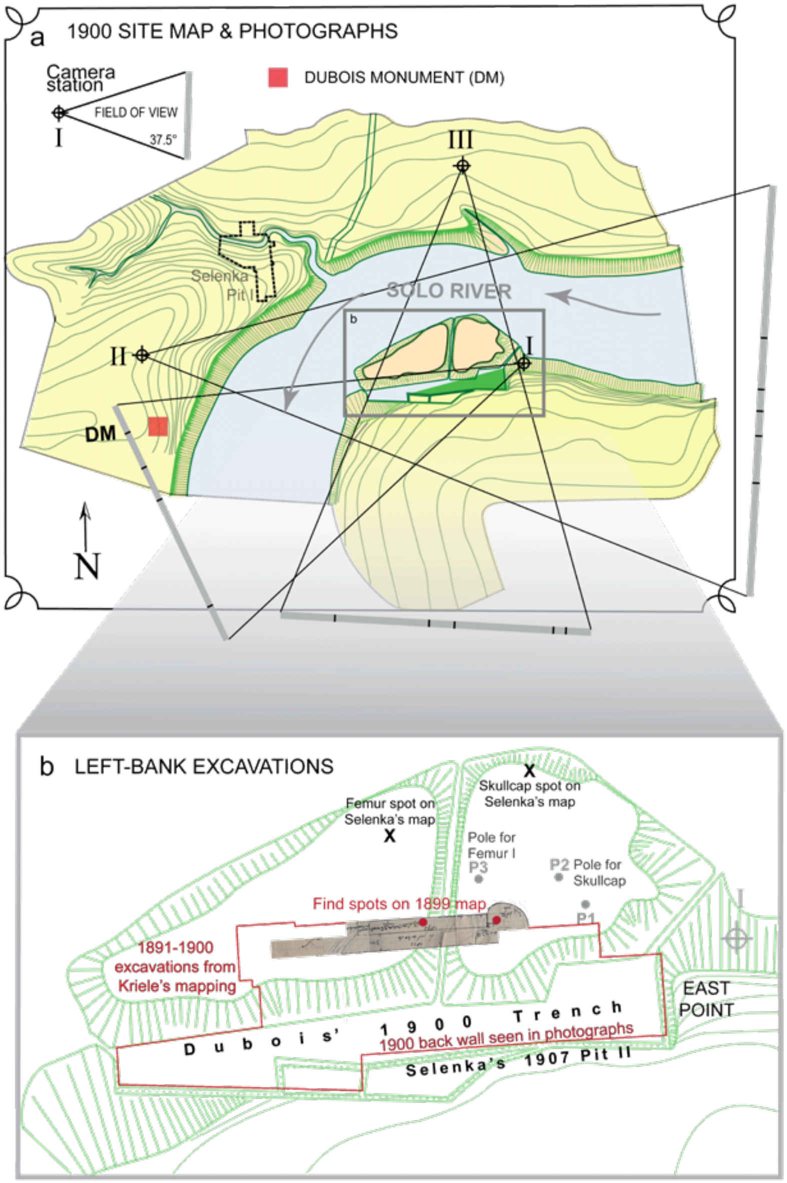


**Supplementary Materials I Figure 7 (S I Figure 7).** The topographic features of the Trinil area in 1900 is portrayed in a map that Dubois possessed but did not publish. We refer to it as the 1900 Site Map (Figure 6a, main text). (**a**) It gives the locations of the three 1900 camera-station stations (I-III) to which we added the photographic fields of view, as determined by matching features visible in multiple photographs. Note that the terrace upland on the right bank around Dubois monument (at the present-day Trinil Museum) sits at a higher elevation than the terrace south of the 1900 Trench on the left bank. (**b**) Attempts to correctly position the Skullcap and Femur I discovery points relative to the modern landscape have proved problematic. Locations in Kriele's 1899 map (red dots, here; also, Figure 3, main text), the three poles on the spoil pile visible in several 1900 photographs (Figure 4, main text) and Selenka relocation points (Figure 6b, main text) are inconsistent with other records. Moreover, the exact spatial relationship between the boundaries of Dubois' 1900 Trench and the Selenka Expedition 1907 Pit II is not fully established. This version of Dubois' 1900 Site Map was made from Naturalis Biodiversity Center scan M…059-220 (S II-E4; also, S II-E6 and -E7); the image of the 1891-1893 pits and trenches is from Figure 3a, main text, and the outline of 1891-1900 excavation is from S II-A4m and -A4r.

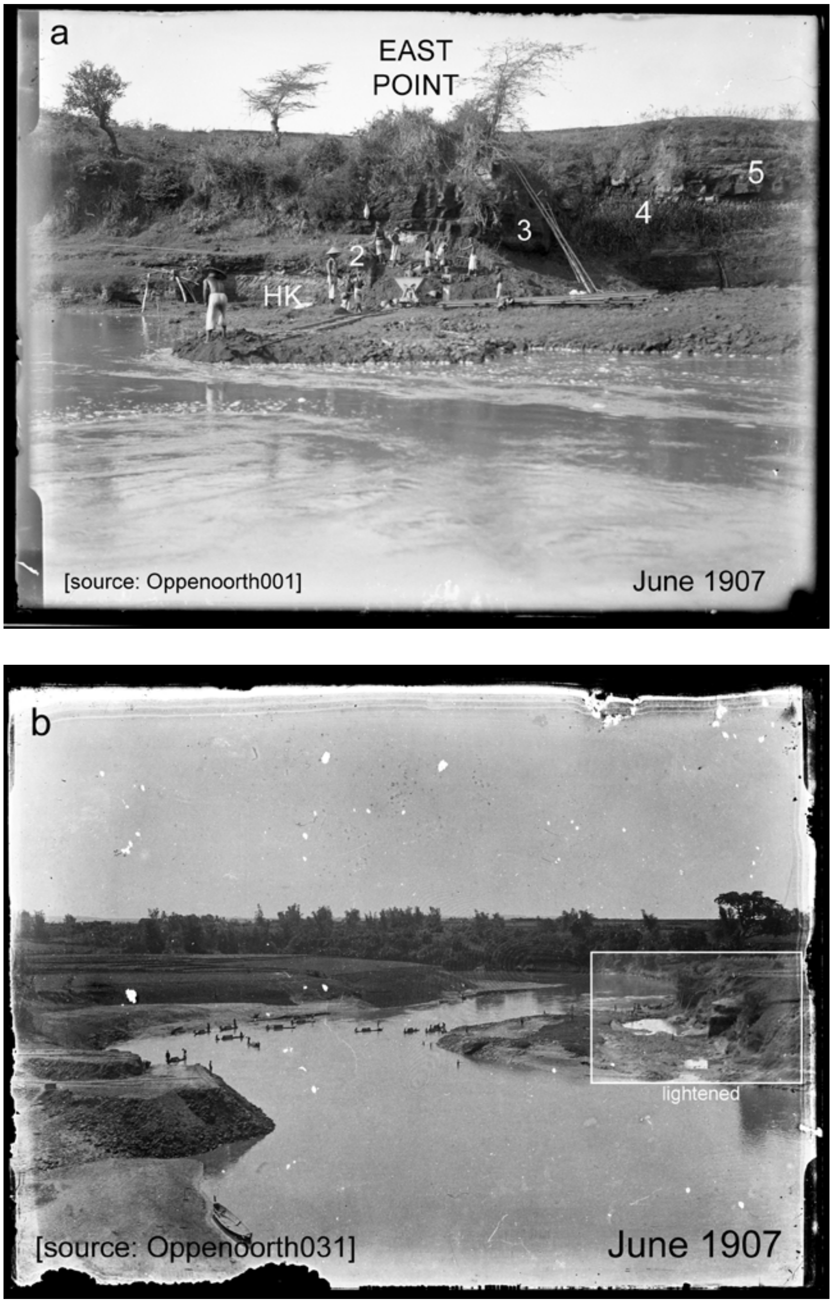


**Supplementary Materials I Figure 8 (S I Figure 8).** Selenka Expedition personnel readily identified the *Pithecanthropus erectus* bonebed on the left bank in 1907, as these photographs taken by supervising geologist W.F.F. Oppenoorth illustrate. (**a**) He began excavating near East Point by June 1907. About this time, the bonebed *"was visible from the Solo when it was exposed above water and immediately produced … bones and teeth;"* a buffalo skull had already been found at the site (Oppenoorth 1911: Fig. 20, xxxiv, Berkhout and Huffman 2021: 37). The bonebed was termed the Hauptknochenschicht (**HK**) later in 1907. Units **2** and **3** are recognizable here in the former walls of Dubois' 1900 Trench (to the right and left of the ladder, where grass and detritus still covered the walls). (**b**) More of the **HK** was unearthed as the river level fell. J.M. Oppenoorth made negatives of these images available to us from her family archives; the negatives were scanned at high resolution at the Naturalis Biodiversity Center.

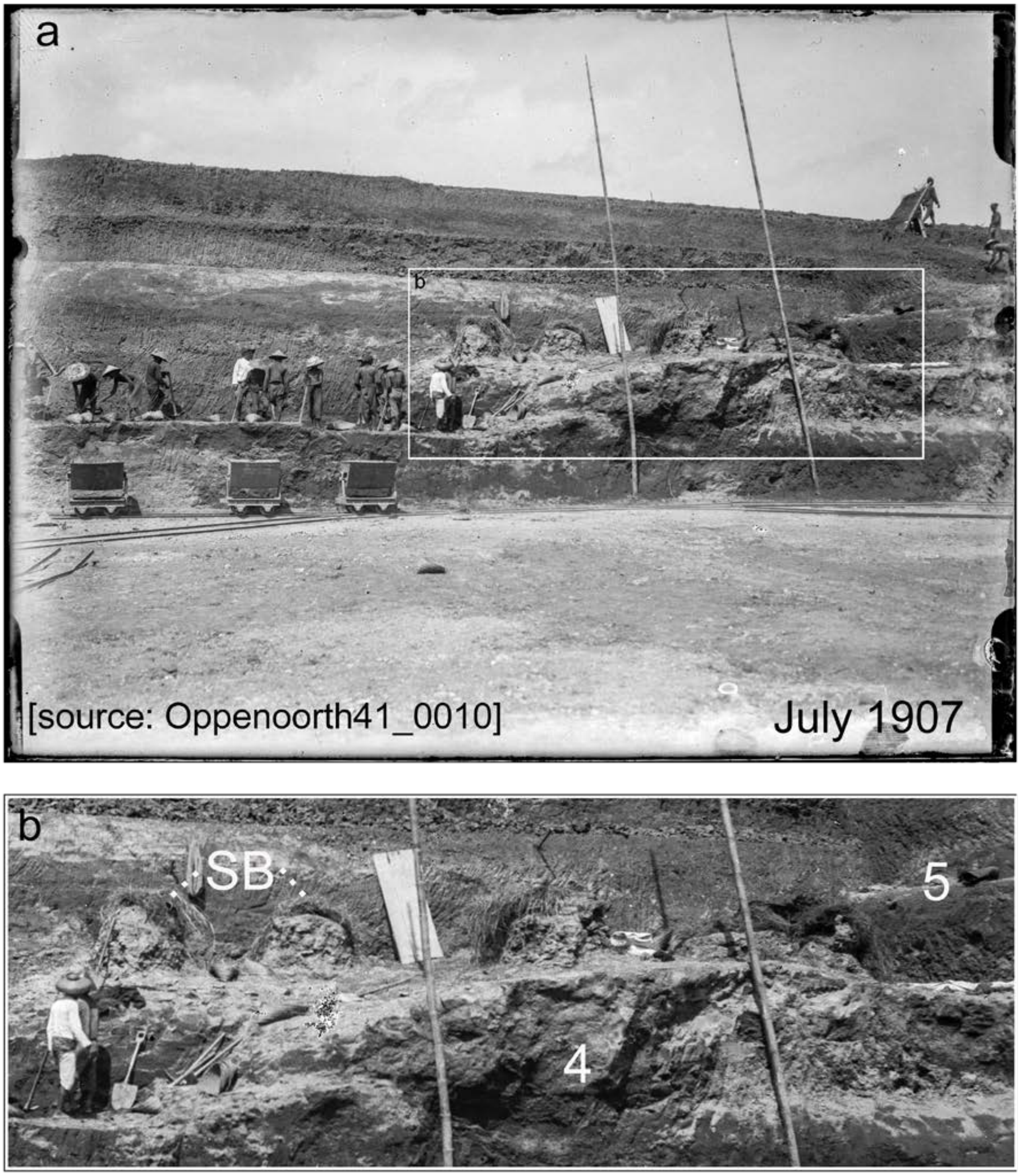


**Supplementary Materials I Figure 9 (S I Figure 9).** Selenka's excavators encountered hard-to-dig strata as they deepened the 1907 Pit II from the agricultural fields above. (**a**) *"In the beginning of July …. the steep riverbank … , landward of Dubois' excavations, was removed …. the highest wall reached 8 to 9 meters"* (Oppenoorth 1911: xxxiv, Berkhout and Huffman 2021: 37). The excavation expanded westward from East Point (Figure 7b, main text) and penetrated indurated strata (our units **5** and **4**), which included the fossil concentration in the *Stegodon* bonebed (**SB**). (**b**, a close up from 'a') The **SB** fossils had to be covered with straw while awaiting removal. The strata below the **SB** (in unit **4**) formed blocky excavation faces of well-lithified sedimentary rock, most likely sandstone. J.M. Oppenoorth made the negative of this image available to us from her family archives. The negative was scanned at 4800 dpi resolution at the Naturalis Biodiversity Center and assigned the source code shown.

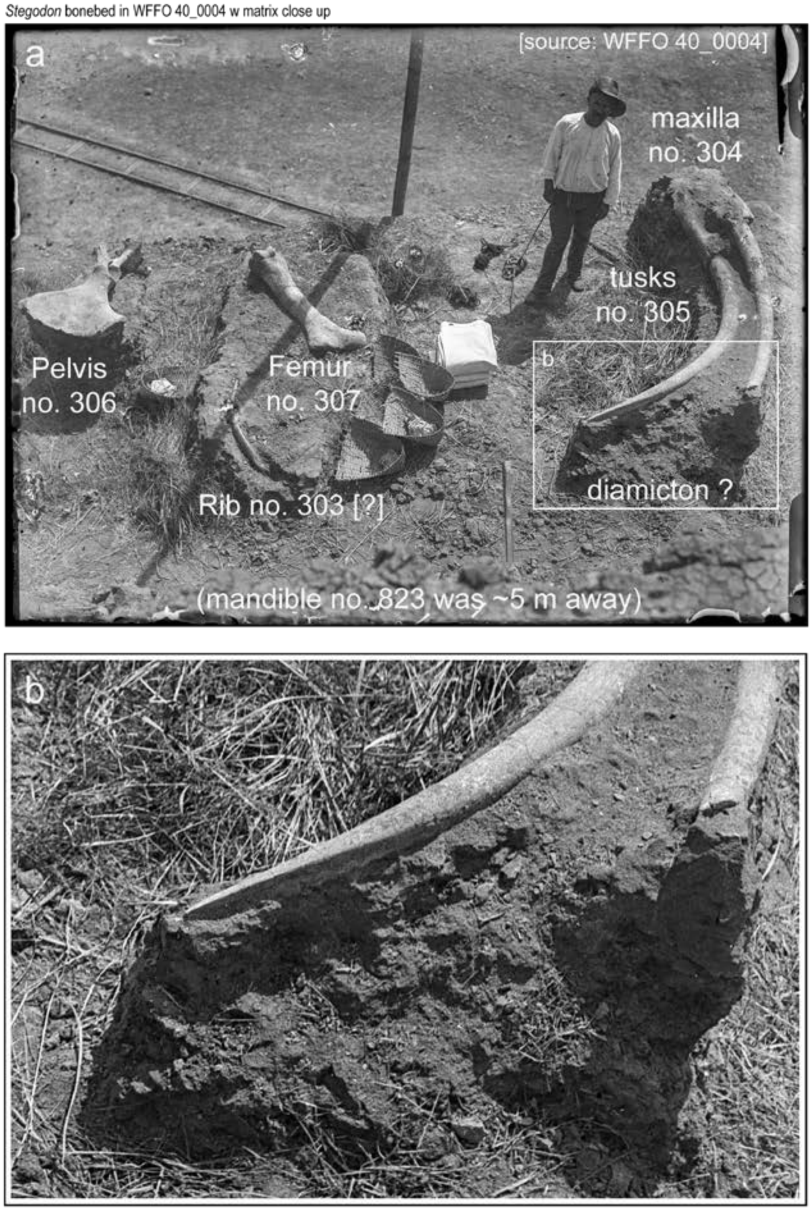


**Supplementary Materials I Figure 10 (S I Figure 10).** (**a**) The proboscidean remains in the *Stegodon* bonebed (**SB**) were predominantly disarticulated and dispersed elements of a large *Stegodon trigonocephalus* individual. The remains occurred at the top of bed with a clayey conglomeratic sedimentary fabric (also, S I Figure 9) seemingly indicative of laharic deposition (also, Oppenoorth 1911: Fig. 24, Berkhout and Huffman 2021: 42). Farther west (near the 'JOG' in the backwall of the 1900 Trench), this stratigraphic level was poorly resistant to erosion, apparently reflecting the clayey composition (Figures 4, 7 and 8, main text). (**b**) Oppenoorth (1911: xxxvi) characterized the bed as *"light grayish clay"* and described the proboscidean remains as *"a skull with upper jaw and tusks (2.10 meters long…), a thigh bone (over 1 m long), the pelvis and ribs, … associated lower jaw… vertebrae, two ribs and a femoral head (see* [his] *Fig. 23* [xxxvii]*) ….* [and] *hippopotamus molars"* (Berkhout and Huffman 2021: 40 and 41). Mandible 823 is the largest specimen of this skeletal element in the Trinil collections (van den Bergh 1996; also, Pohlich 1911). J.M. Oppenoorth made a photographic negative of this image available from her family archives, and it was scanned at 4800 dpi resolution by the Naturalis Biodiversity Center.

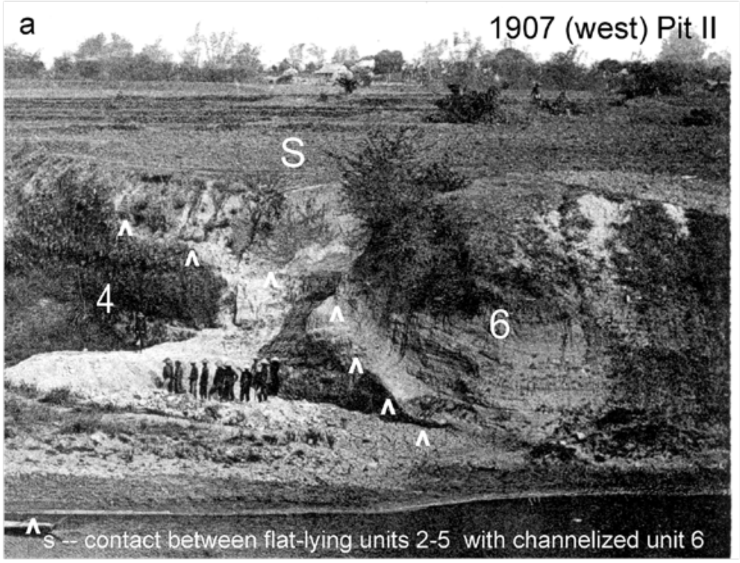


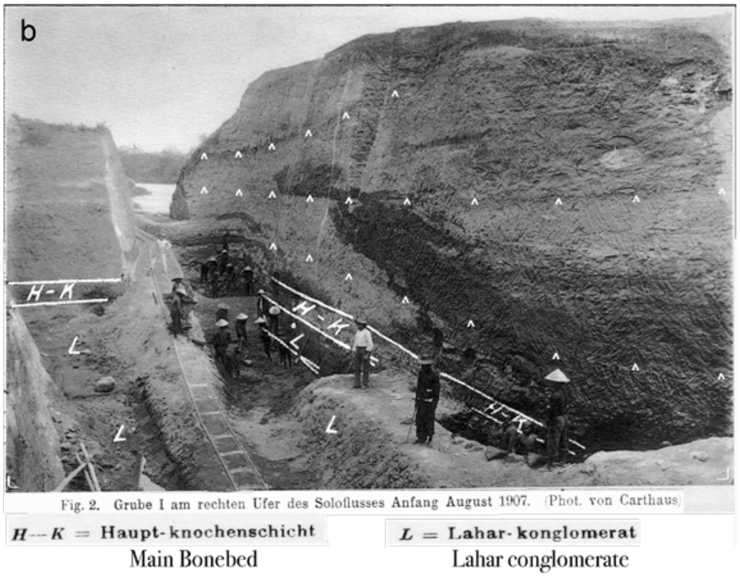


**Supplementary Materials I Figure 11 (S I Figure 11).** (**a**) In the western portion of Pit II, the base of the channelized unit **6** reached river level (from Selenka and Blanckenhorn 1911: Plate I, Fig 2, Berkhout and Huffman 2021: 5). (**b**) The strata above the **HK** in the right-bank Pit I, north of the present-day Trinil Museum (Figure 6a, main text), also contained channel-form features (Selenka and Blanckenhorn, 1911: Plate VII, Fig. 2, Berkhout and Huffman 2021: 50; the ^s have been added to highlight stratigraphic relations). Despite similarities in lithofacies, the Selenka Expedition geologists did not appear to have established a reliable stratigraphic correlation between deposits in Pit I and II (S I Figure 7a). The uncertainty continued when Duyfjes (1936) attributed the post-**HK** sequence on the right bank to terrace deposits and the post-**HK** sequence on the left bank to the bedrock Kabuh Formation (S I Figure 16). (**c**, next page) The 1907 Listing (of Selenka finds allows the irregular vertical and horizontal distribution fossils in the Hauptknochenschicht (**HK**) to be seen (north is up). The finds were reported for the three stratigraphic subunits of the **HK** and we list each level separately for each meter square in the 1907 Pit I.

### PIT I FOSSIL-ENTRY FREQUENCY IN THE 1907 LISTING
#### FOR HAUPTKNOCHENSCHICHT (SUBUNITS ) LAYERS 15, 16 AND 17

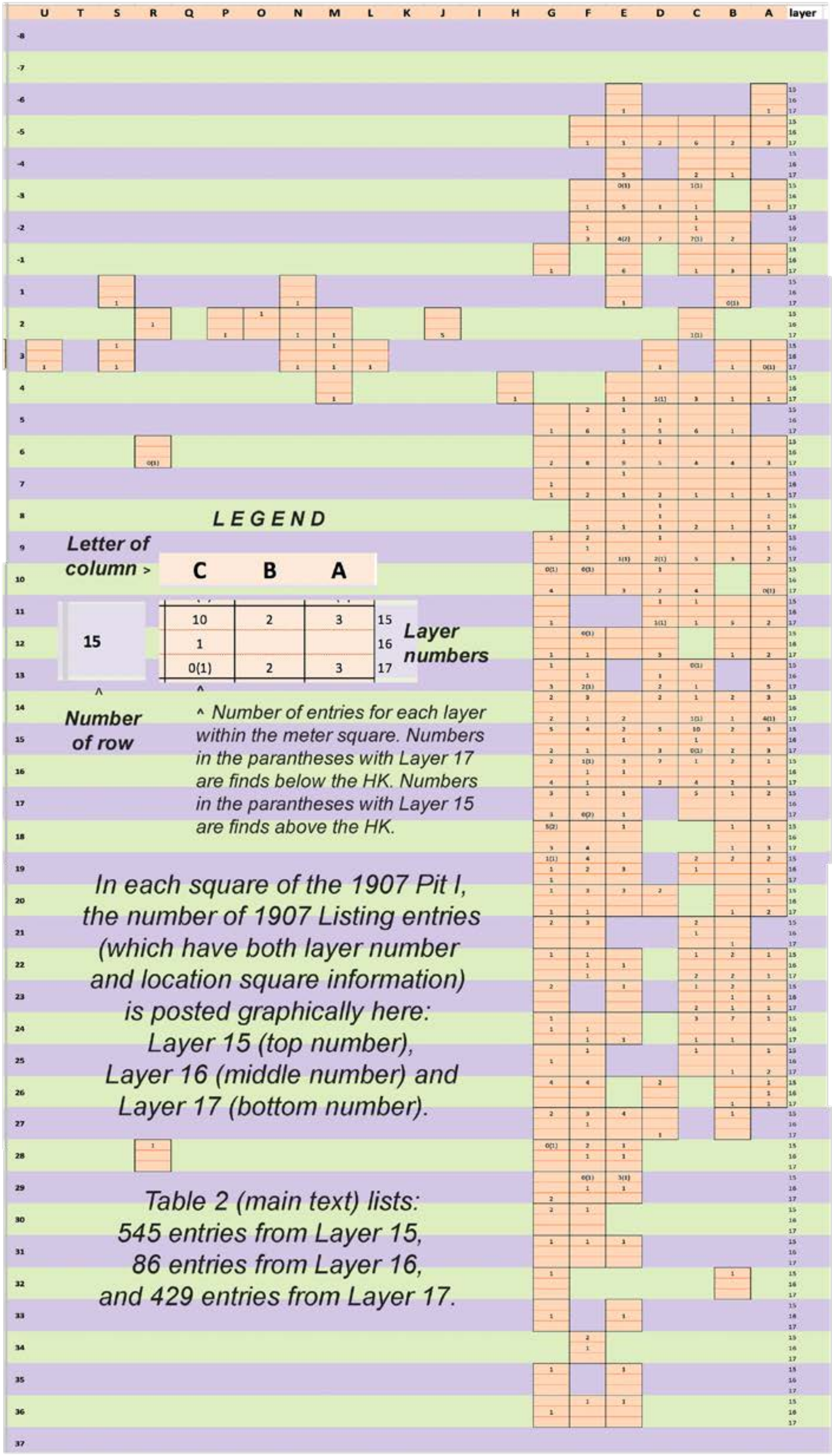

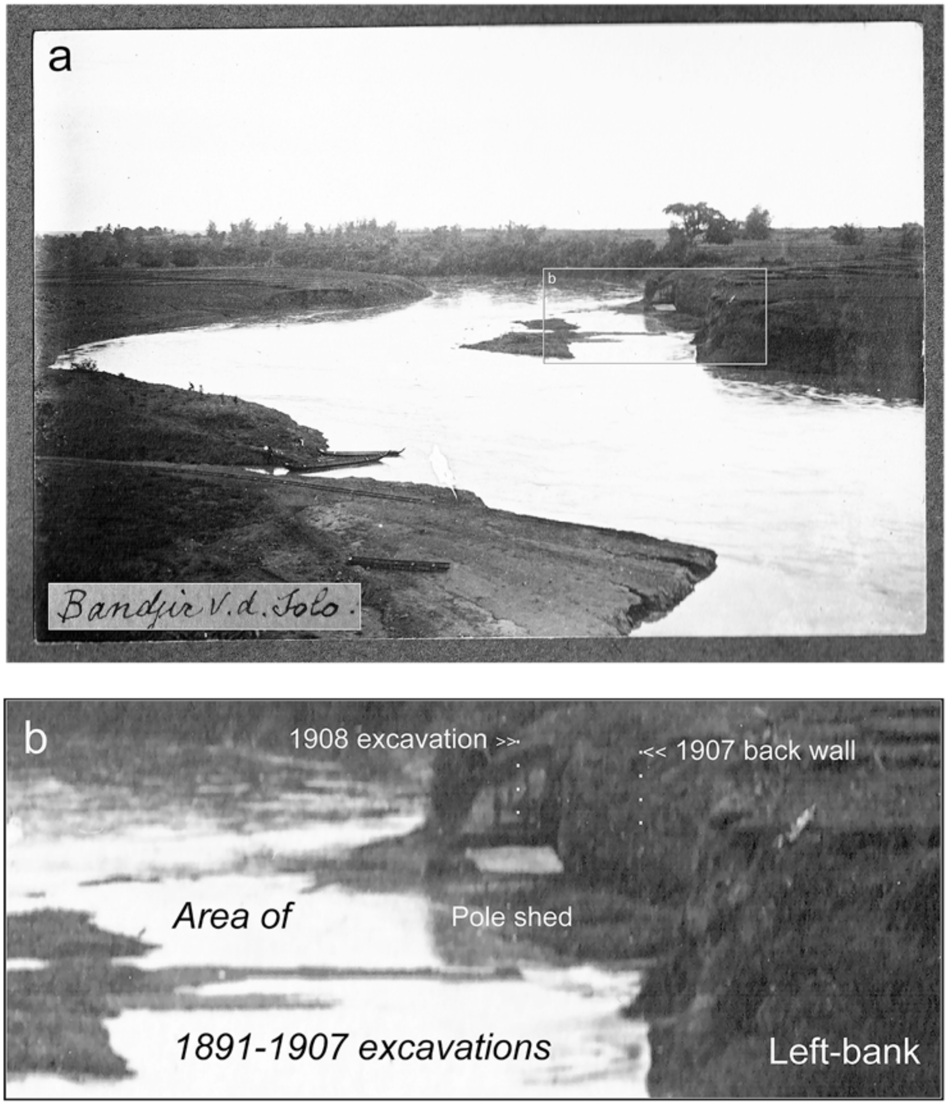


**Supplementary Materials I Figure 12 (S I Figure 12).** The high back walls of the 1907 Pit II were still standing vertically in 1908. (**a**) This October 1908 photograph, which looks upriver from a high point on the right bank, shows that most of Pit II was inundated by flooding. (**b**) However, a portion of the 1907 back wall near East Point was freshly excavated (behind the pole shed a light-colored roof). The photograph was taken (without known scientific purpose) by W.J. ('Wim') Oppenoorth, a friend of C.M. Dozy and the brother of W.F.F. Oppenoorth. The photograph is in an Oppenoorth family album and is labeled *"Banjir of the Solo"* (inset within 'a'). W.J.O.'s granddaughter has the album, and J.M. Oppenoorth, his grandniece, provided us with a scan of the photograph and its provenience history.

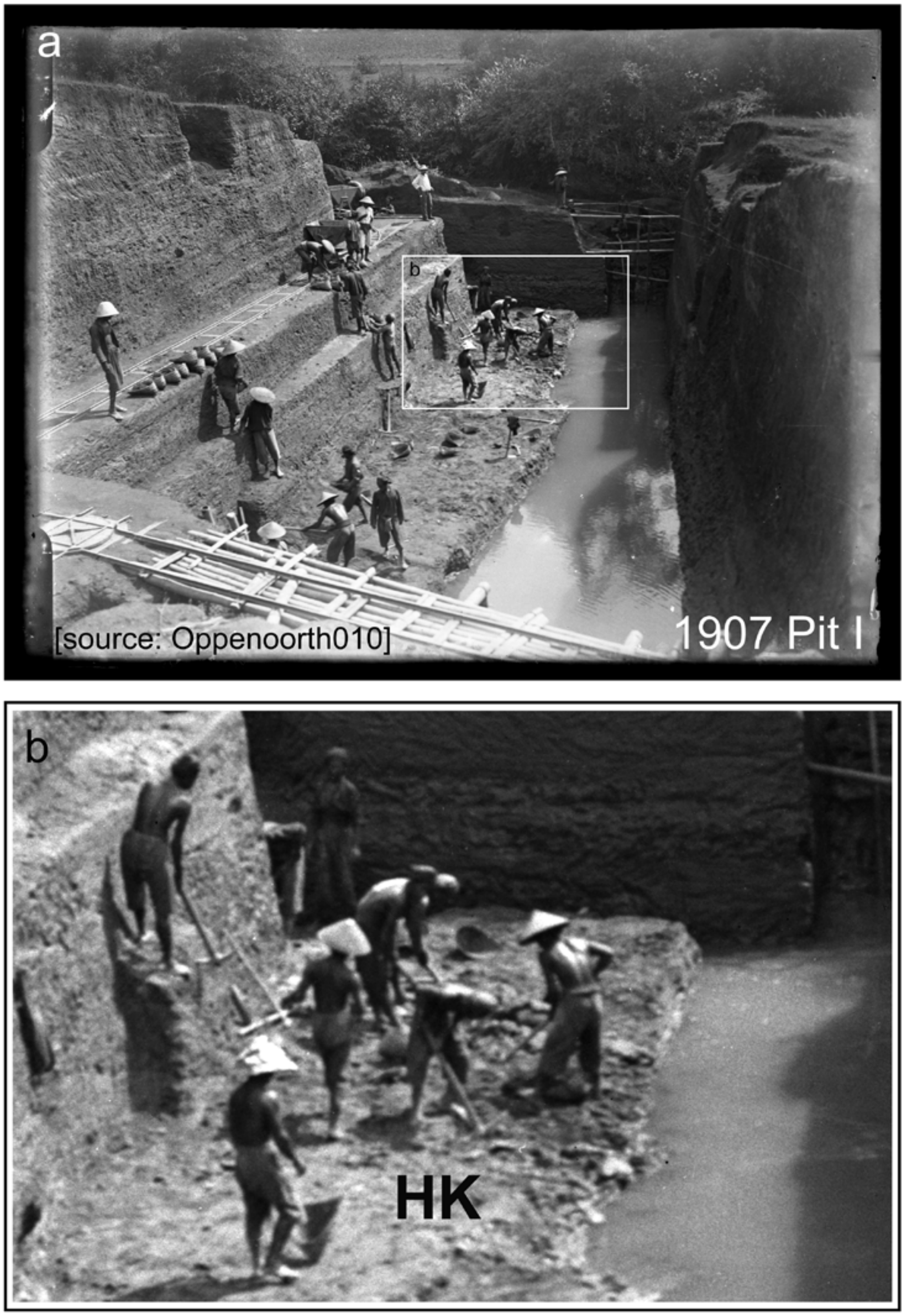


**Supplementary Materials I Figure 13 (S I Figure 13).** The Selenka Expedition Pit I encountered flat-lying, well-indurated volcaniclastic and muddy strata, which were amply described and pictured in the Expedition's published and unpublished materials. (**a**) The sedimentary consolidation of the stratal sequence is evident in this high-quality unpublished version of an image taken in June 1907 (see Selenka and Blanckenhorn 1911: Plate III, Fig. 3; W.F.F. Oppenoorth took the photograph and this version was provided courtesy of J.M. Oppenoorth; Berkhout and Huffman 2021: 36). (**b**) Near the bottom of the excavation, a group of workers used pickaxes to unearth the **HK** and superjacent beds. The **HK** had already been removed from the nearby inundated area. The horizontal structural attitude of the **HK** is evident from its contact with the water.

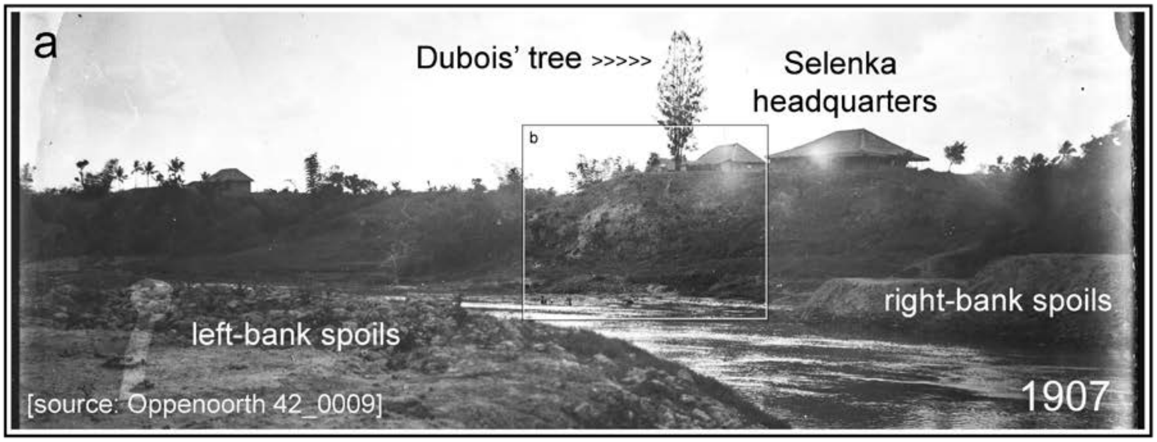


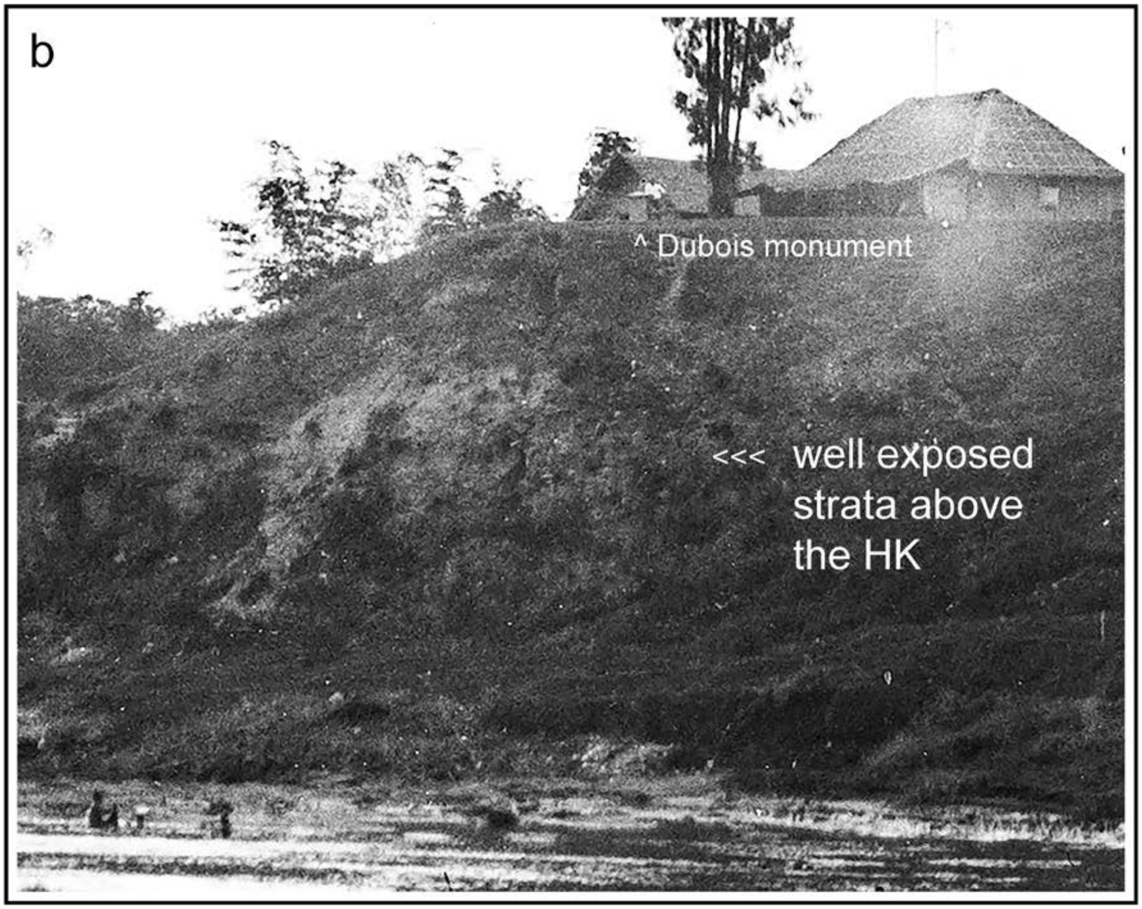


**Supplementary Materials I Figure 14 (S I Figure 14).** From Dubois' day forward, strata above the main bonebed were available for examination on the right bank, as shown in this photograph of the area below the Selenka field headquarters (now the location of the Trinil Museum). The strata in the bank appear to have a flat-lying structural attitude. J.M. Oppenoorth made a negative available from her family archives; it was scanned at high resolution at Naturalis Biodiversity Center.

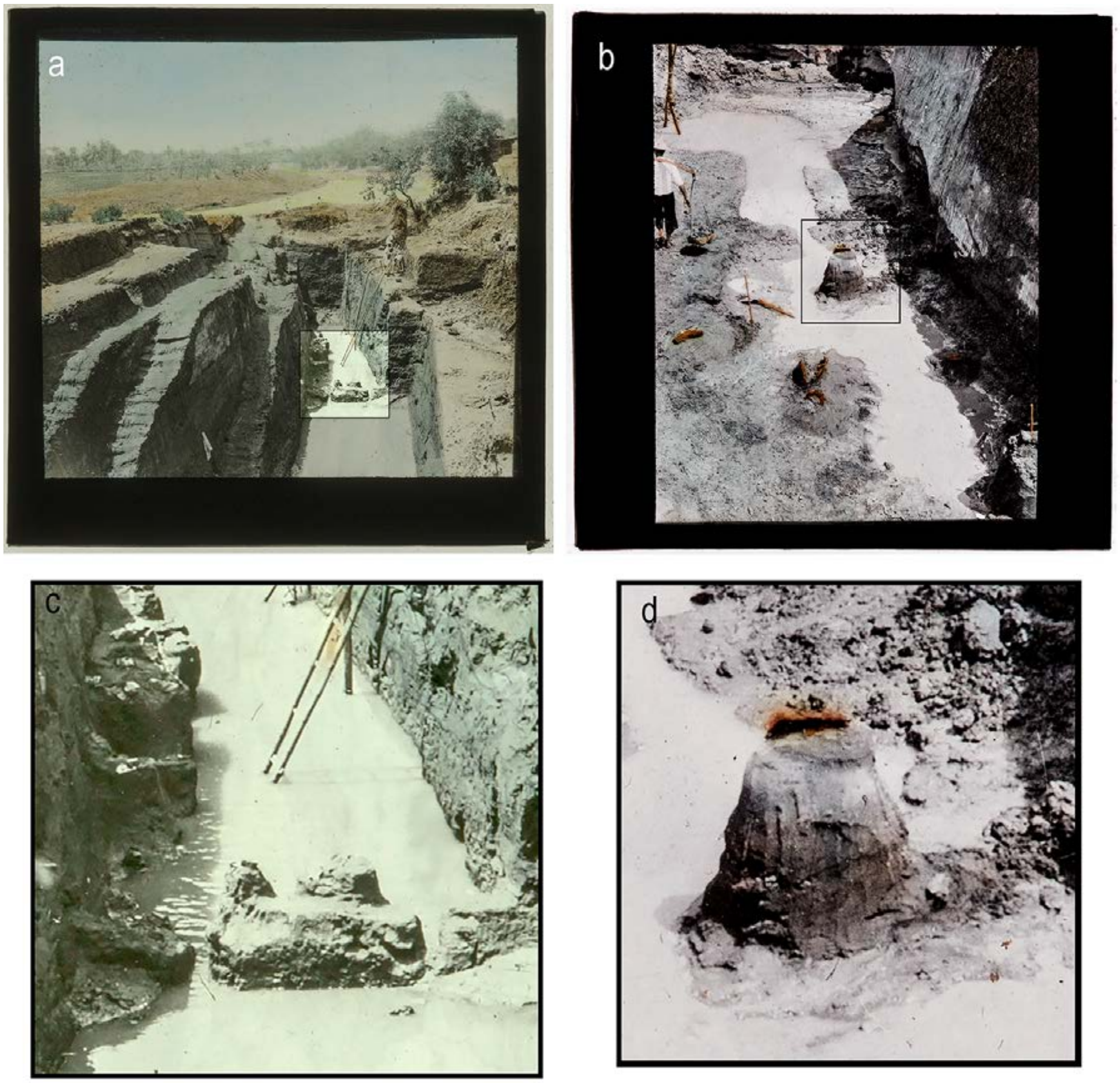


**Supplementary Materials I Figure 15 (S I Figure 15).** W.F.F. Oppenoorth made lantern slides from 1932 photographs of a Trinil excavation undertaken by the Geological Survey of the Netherland Indies. No maps or cross sections of this 1931-1932 work have been located, but the slides are labeled *"Trinil Excav. 1932"* and reveal a large right-bank excavation that was cut into and perhaps through the **HK**. Trenching of the left bank evidently had ended in 1908. The 1931-1932 Survey work produced 1306 vertebrate fossils and numerous molluscan shells, apparently most-if-not-all from the **HK** (von Koenigswald 1934, 1934/1935; Duyfjes 1936; van Benthem Jutting 1937; Joordens et al. 2009). The 1932 Trinil dig was part of a program of geological mapping and excavation that was conducted along the Solo River valley and led to the discovery of 14 *Homo erectus* specimens and ~25,000 other fossils at Ngandong (Huffman et al. 2010, Oppenoorth 1932, Rizal et al. 2021). The Trinil and Ngandong excavators followed field procedures similar to those that the Selenka Expedition used. This includes similar provenience descriptions. For example, a 1932 annotation about one Trinil find reads *"Sheet 93B* [referring to the pre-World War II topographic quadrangle] *Ingr.*[aving; excavation] *I, block C, Layer* [bed] *II. 23. VII. 1932,"* meaning Excavation I, block C, from source stratum II, the **HK**, and recorded in the fossil registry on July 23, 1932 (van Benthem Jutting 1937). J.M. Oppenoorth made these slides available to us from her family archives, and they were scanned at high resolution by the Naturalis Biodiversity Center.

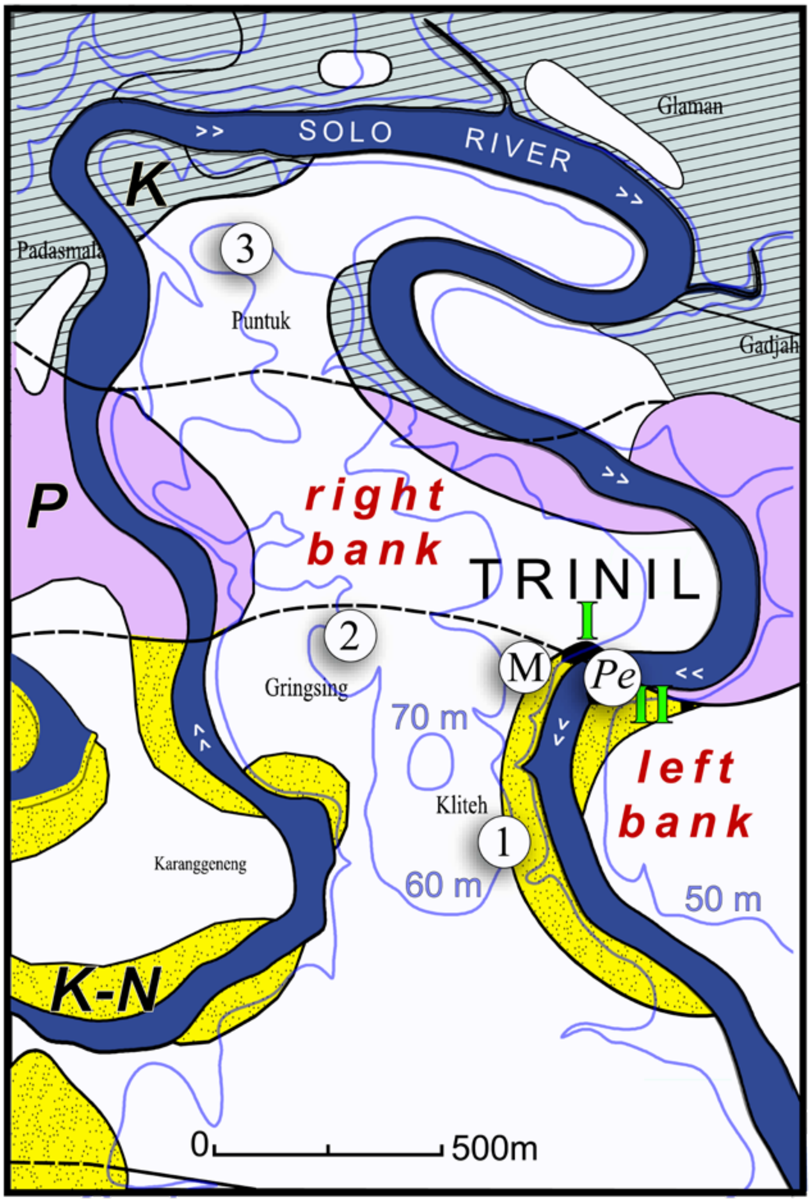


**Supplementary Materials I Figure 16 (S I Figure 16).** Duyfjes' (1936) geological map of the Trinil area (redrafted from B.W. Seubert in Huffman 2020). He recorded the presence of widespread terrace deposits (white) overlying gently south-dipping Kalibeng (***K***), Pucangan (***P***) and Kabuh-Notopuro (***K-N***) Formations. The ***P.e.*** marks the general location of Dubois' *Pithecanthropus erectus* site (see S I Figure 17, for detail). The Selenka Expedition 1907-1908 Pit II (**II**) and Pit I (**I**) were on opposite sides of the river with **I** being located northeast of the present-day Trinil Museum (**M**; S I Figure 7). Duyfjes' (1933) field map of this area showed terrace deposits entirely surrounding Pit I (S I Figure 17a), but for publication in 1936, he raised the elevation of the contact between the terrace deposits and the Kabuh Formation (S I Figure 17b). Locations '**1**'-'**3**' are four- to ten-m-tall exposures of horizontal strata that conform to Duyfjes' recognition of a thick widespread flat-lying unit atop the right bank (Huffman 2016). Duyfjes' mapping has served as a basis for stratigraphic understanding of the Trinil area since the 1930s, although uncertainty continues about the extent of terrace deposits versus the Kabuh Formations there (e.g., S I Figure 19).

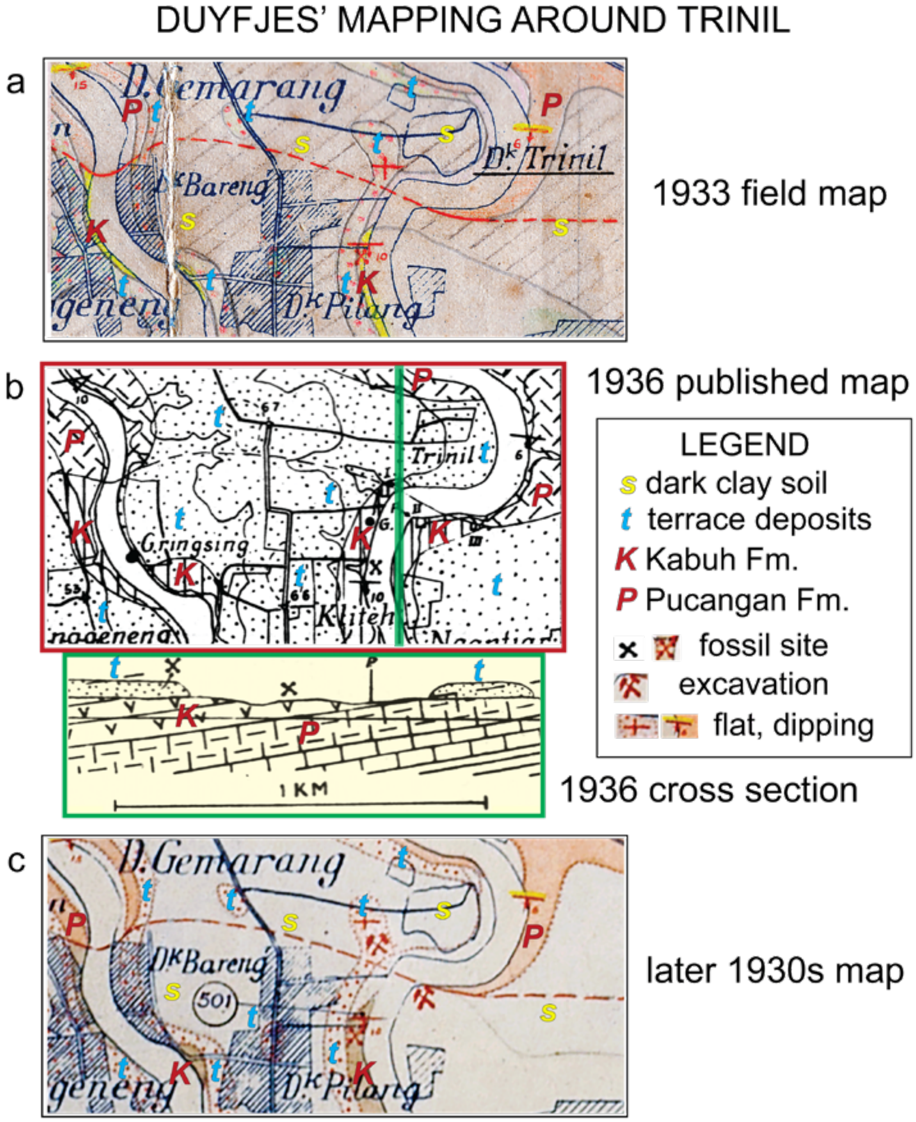


**Supplementary Materials I Figure 17 (S I Figure 17).** Three versions Duyfjes' geological mapping of the discovery area are available (cropped and annotated here): (**a**) The map with his unpublished 1933 report on the fieldwork (from the geological museum library, Bandung; Duyfjes 1933); (**b**) Duyfjes' (1936) published map with a north-south cross section; and (**c**) a later unpublished map (retained by Naturalis, Leiden, and provided by F.P. Wesselingh). Four key relations are portrayed. First, flat-lying terrace deposits surround the Survey's 1932 and Selenka's 1907-1908 right-bank excavations (clearest in 'a' and 'c'), where >9m of section were exposed above the **HK** (Selenka and Blanckenhorn 1911 and S I Figure 15). Secondly, the contact between the Kabuh and terrace deposits is stratigraphically higher in the published map than in unpublished ones. Third, Duyfjes had the Kabuh Formation beneath terrace soil on the left embankment, south of the former hominin discovery site, and full development of terrace deposits farther south along that bank (shown in cross section). Finally, he measured south dip in both the Kabuh and Pucangan formations, but saw substantial thicknesses of horizontal terrace deposits on top of these (and older) formations on the right bank. The **HK** is situated near the base of the Kabuh Formation in Duyfjes' cross section. It greatly exaggerated the thickness of the terrace deposits. Duyfjes' work is the only known geologic mapping of the right- and left-banks recorded at a scale as fine as 1:25,000 (Huffman 2020 and Berkhout and Huffman 2020 have translations of Duyfjes' publication and report on the Trinil area).

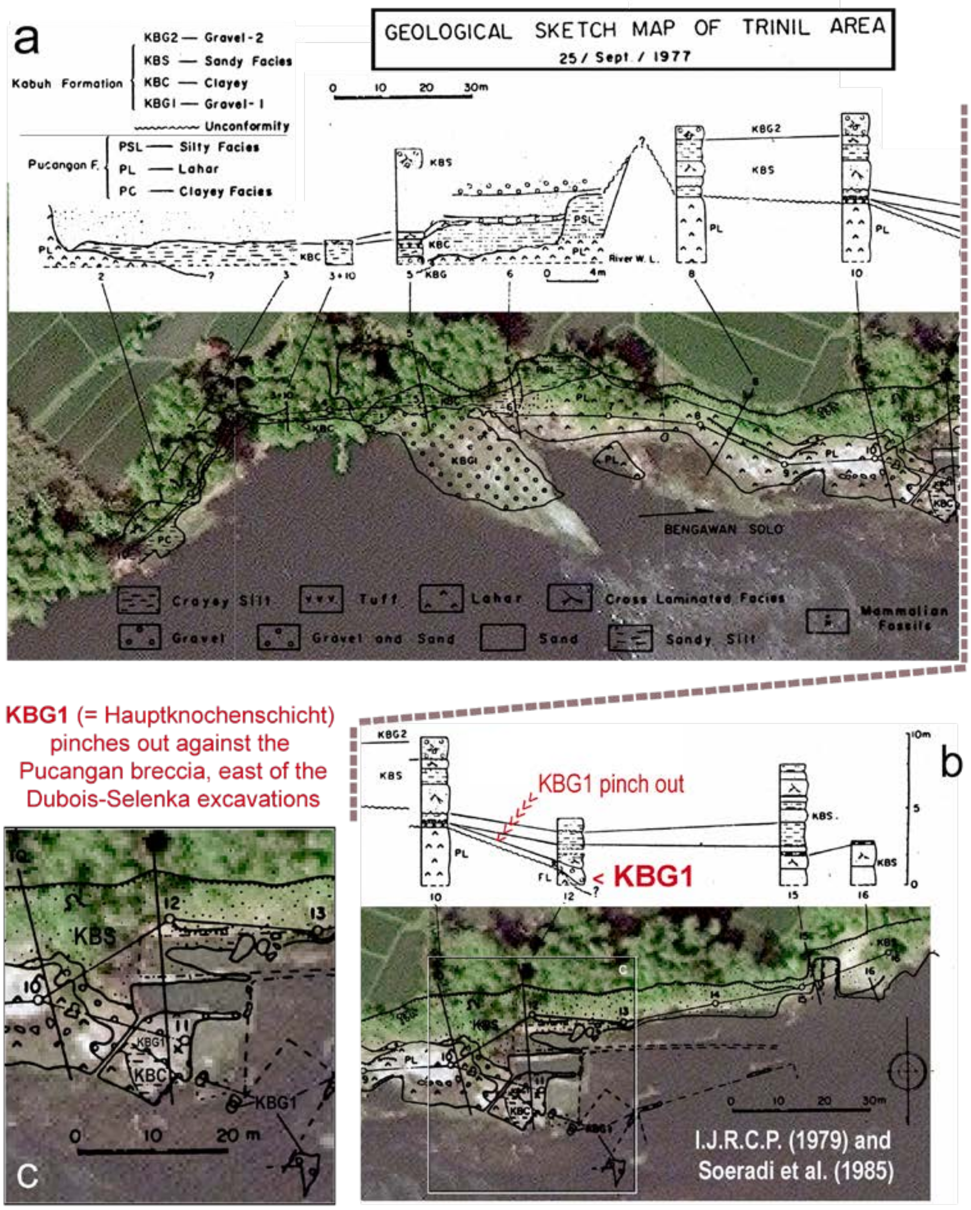


**Supplementary Materials I Figure 18 a-c (S I Figure 18 a-c).** A geological map of the left bank and several measured sections were published at 1:250 (Soeradi et al. 1985; previously published by I.J.R.C.P. 1979). The mapping is registered here onto satellite imagery (as is the case with the geological mapping in Figure 6d, main text). North is up (as it is Figure 3a, main text, but opposite to the orientation shown in Figure 6d, main text). Soeradi et al. found that the lithology and paleontology of their KBG1 map unit matched the features of main bonebed (**LB** and **HK** of Dubois and Selenka; also, Aimi and Aziz 1985). They agreed with Duyfjes' (1936) assignment of the bonebed and overlying strata to his self-defined and regionally mapped Kabuh Formation (also, S I Figure 19). Soeradi et al. did not see the ~9$^{o}$ south dip that Duyfjes attributed to the strata (S I Figure 17b) but confirmed the eastward onlap and pinch out of the bonebed onto a diamicton-bearing unit, a relationship first described by Dozy (1911a: xli, Berkhout and Huffman 2021: 47). Soeradi et al. did not relate they sandy, gravely and clayey facies they found in the superjacent 8m of strata (section 15 in 'b') to the stratigraphic sequence described in Selenka and Blanckenhorn (1911).

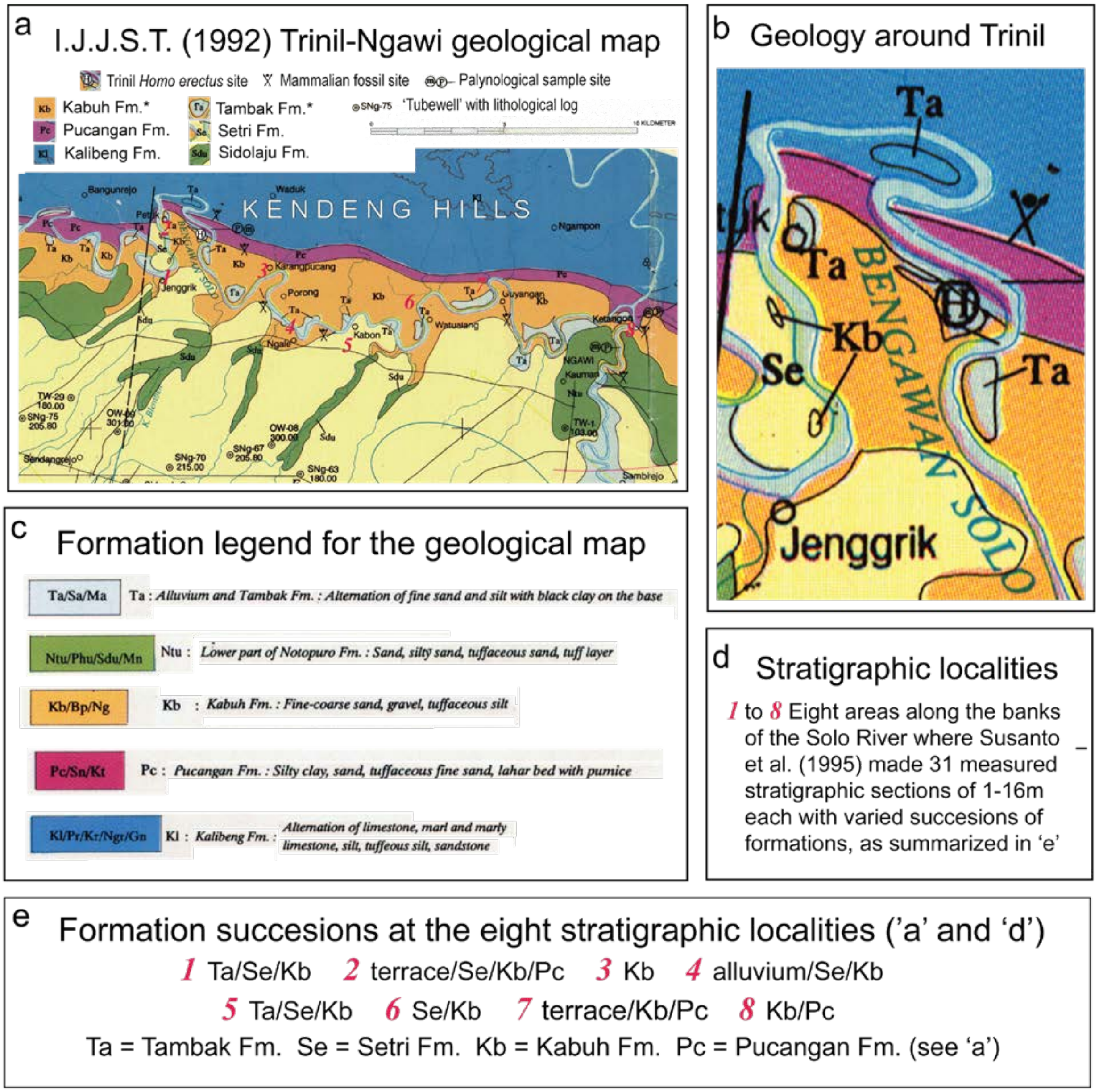


**Supplementary Materials I Figure 19 a-c (S I Figure 19 a-c).** (**a** to **c**) Geological mapping done along the Solo River valley by multiple field teams shows prominent outcrop bands of the Kabuh and Pucangan Formations. This map was prepared by Indonesian-Japan Joint Study Team (I.J.J.S.T. 1992), who had mapped the Kabuh and Pucangan from Sangiran Dome to Kedungbrubus (Figure 1b has these locations; the elements of the I.J.J.S.T. map are rearranged and annotated here). At Trinil, I.J.J.S.T. mapped alluvium and Recent deposits, combined as "Ta…", on the terrace upland south of the *Pithecanthropus erectus* discovery area. "Ta…" is not otherwise described. Few areas of "Ta…" were recognized above the Kabuh Formation on the right bank (S I Figure 16; also, Datun et al. 1996). The light-yellow unit mapped on the south side of the Solo River valley represents a thin sedimentary cover of materials derived from Lawu volcano. (**d** and **e**) In contrast to the simplicity of the I.J.J.S.T. geological interpretation of the Trinil area, others have seen complex of stratigraphic sequences along the banks Solo River, which we represented here by stratigraphic synopses of one field team at eight localities (Susanto et al.1995; also, I.J.R.C.P. 1979).

**Supplementary Materials I Figure 20 (S I Figure 20). Trinil fossils in the Dubois Collection.**

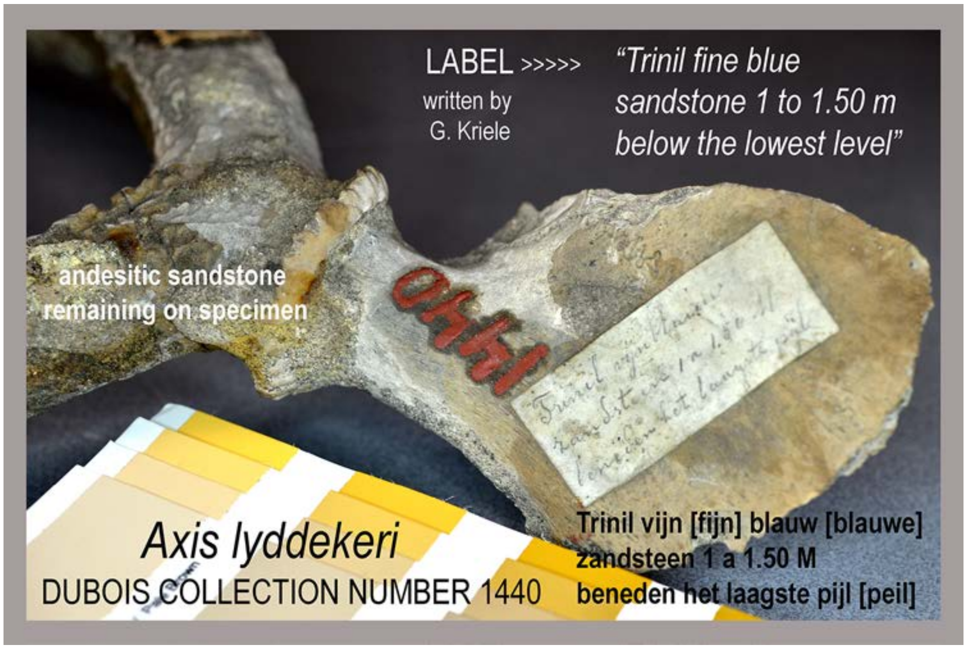


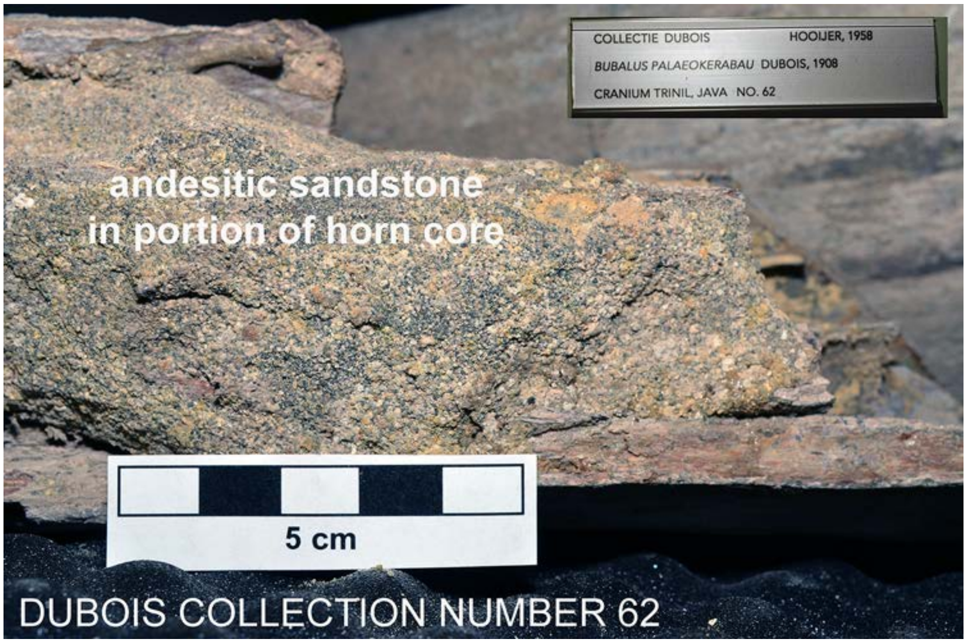


**Supplementary Materials I Figure 20a**. (top) Antler and cranial fragment of *Axis lydekkeri* with label written by G. Kriele that give provenience information. The specimen has a coating of andesitic sandstone typical of many Trinil fossils. (bottom) Coarse volcaniclastic sandstone fills part of the *Bubalus* horn core, also a common feature of the large specimens in the Dubois Collection. The photographs for S I Figure 20 were taken by OFH in 2010 and 2013.

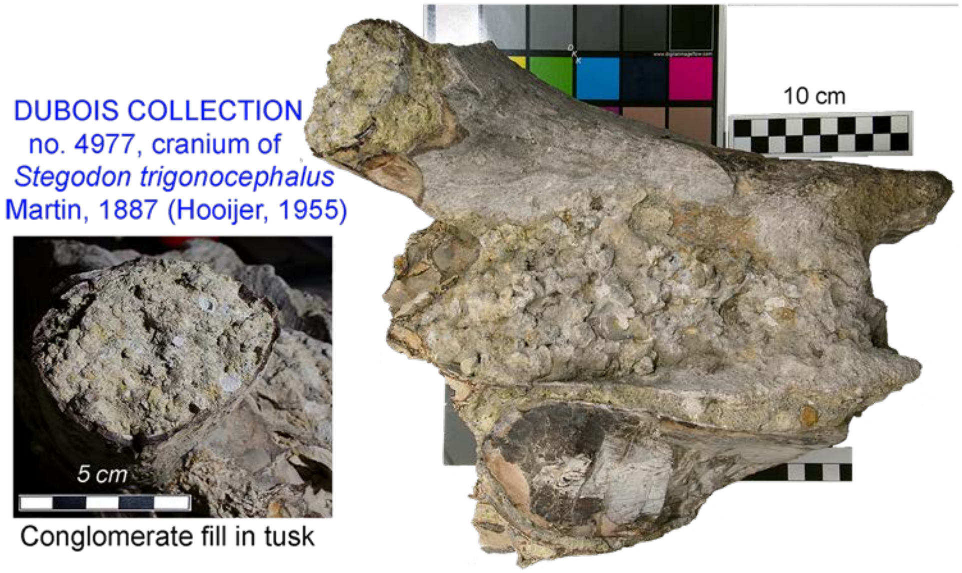


**Supplementary Materials I Figure 20b.** Conglomerate adhering to a partial cranium of a *Stegodon trigonocephalus* specimen and filling the cavity of its tusk.

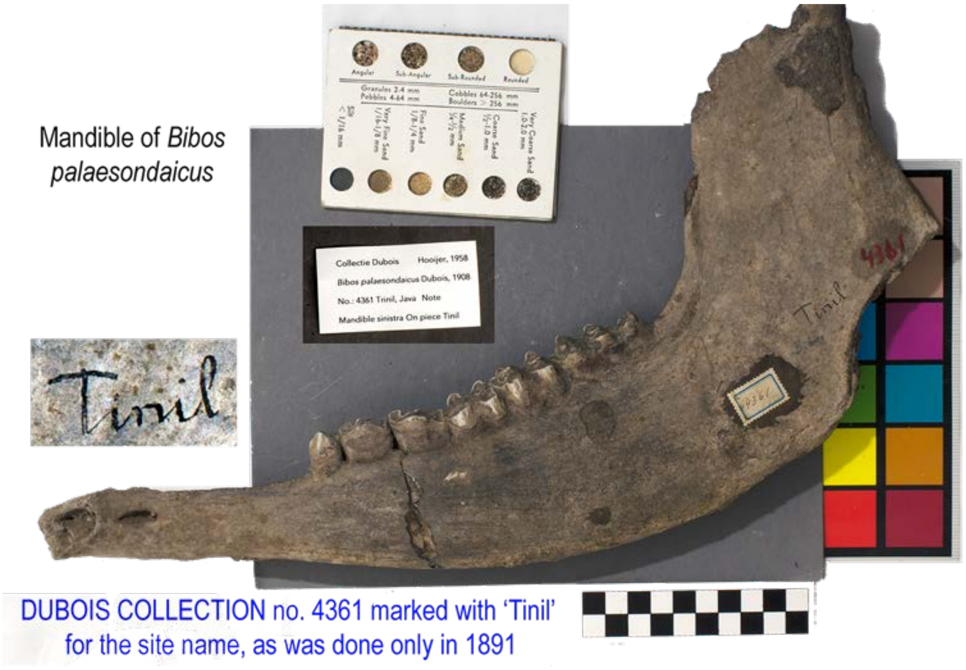


**Supplementary Materials I Figure 20c** A half mandible of *Bibos palaesondaicus* marked "Tinil," which is a spelling of Trinil that was only used by Dubois' field team in 1891. The specimen, which exhibits typical Trinil fossilization, has a dense, dark-purplish brown, porcellaneous interior, which is visible here where the specimen is chipped in the upper right (Ingicco et al. 2015 have images and an account of a Hylobatidae femur from Trinil with similar fossilization; also, Hill et al. 2015, Huffman et al. 2018, Pop et al. 2020, Ruff et al. 2015).

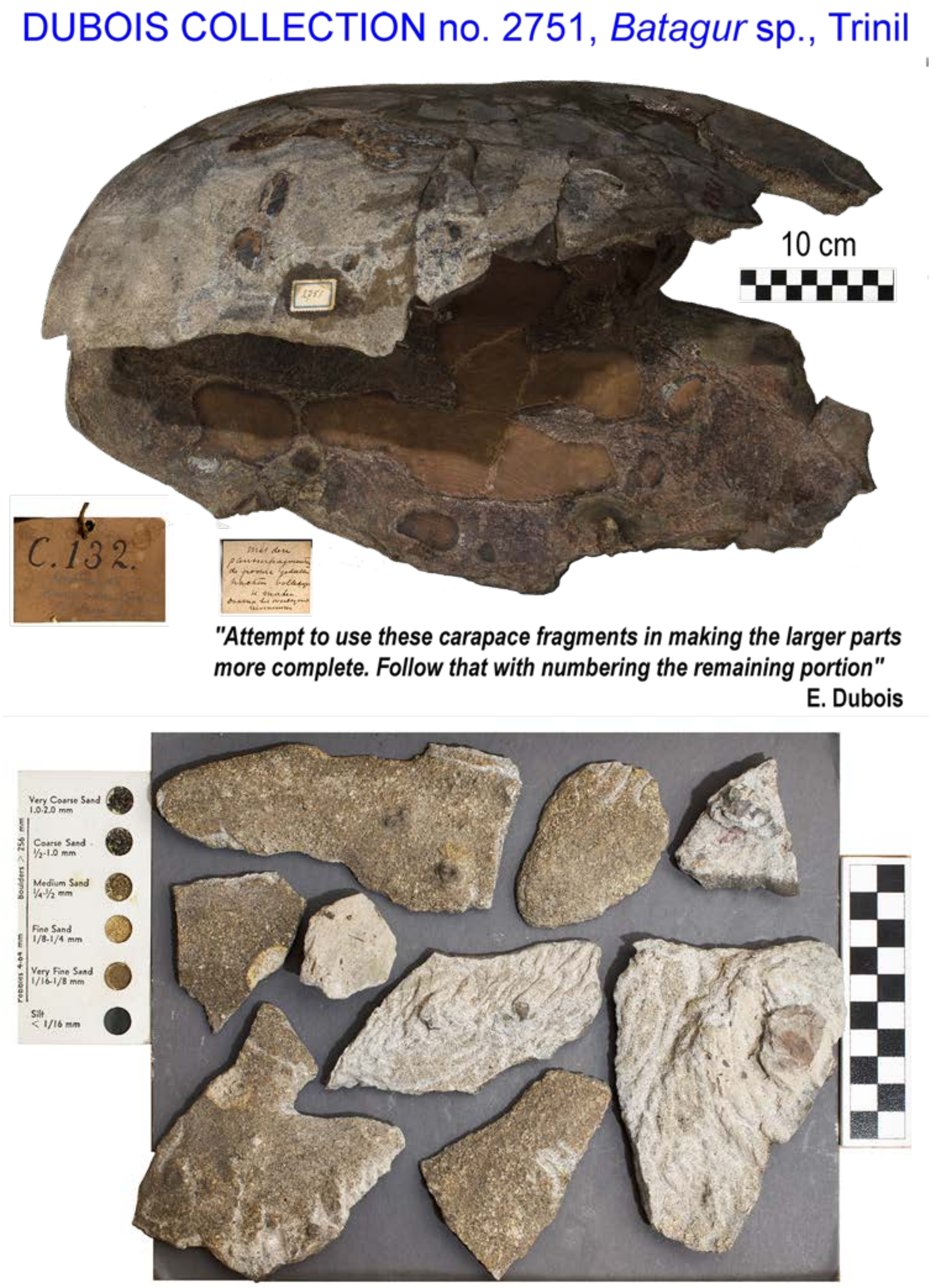


**Supplementary Materials I Figure 20d.** A *Batagur* carapace, reconstructed according to Dubois' written instructions, and fragments of sandstone matrix that had been removed from the fossil (see also Selenka and Blanckenhorn 1911: xxxv, Figure 21, of jacketed "turtle" specimens from their excavations at Trinil; Berkhout and Huffman 2021: 39).

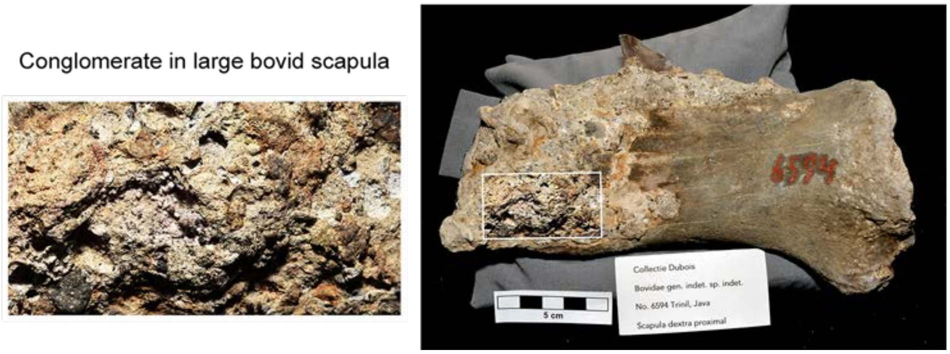


**Supplementary Materials I Figure 20e.** Conglomerate infilling a portion of a large bovid scapula with enlargement showing fresh crystals in addition to lithic clasts.

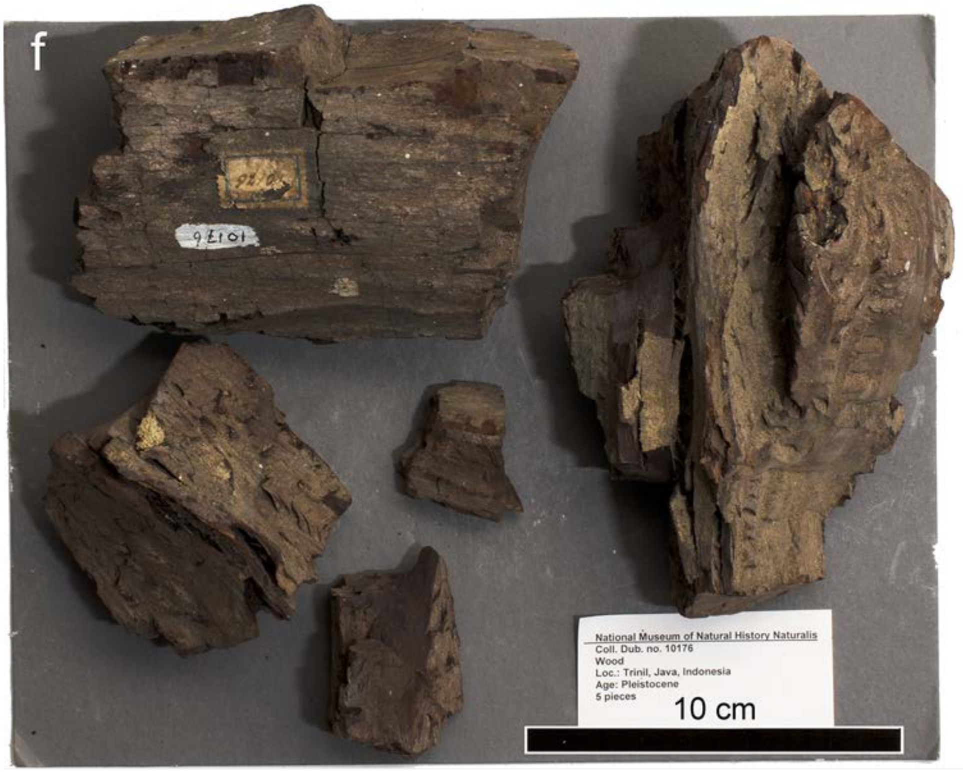


DUBOIS COLLECTION no. 10176, Trinil wood

**Supplementary Materials I Figure 20f.** The main bonebed contained large fragments of wood, according to both Dubois with respect to the **LB** (e.g., Appendix A(iii), A(iv), and C(vi)) and the Selenka Expedition personnel concerning the **HK** (e.g., Carthaus 1911b: 14; Berkhout and Huffman 2021: 72). The five pieces comprising this specimen number fit the description of the main bonebed wood, but no documentation about the stratigraphic source is included with the Naturalis records with the specimen.

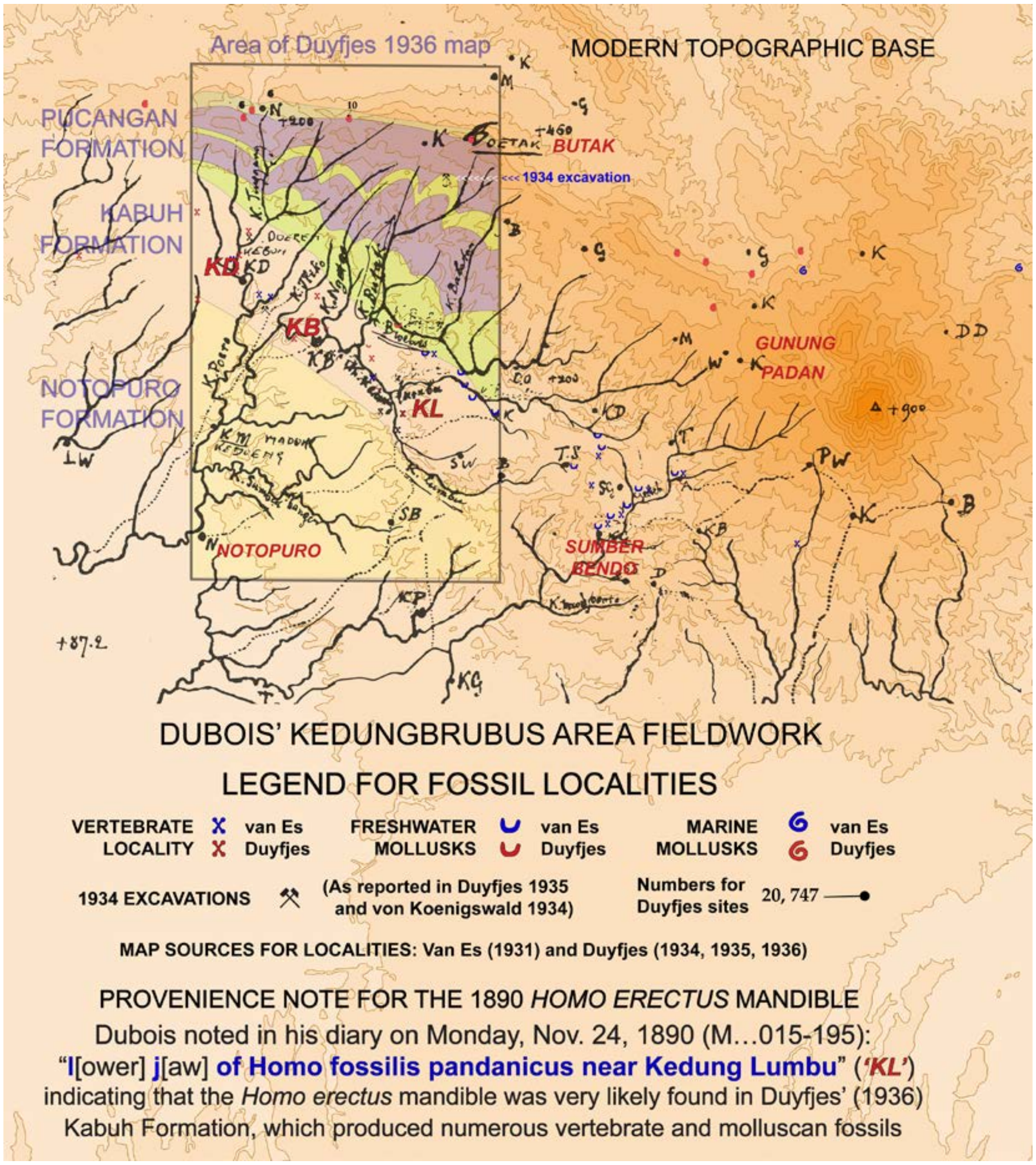


**Supplementary Materials I Figure 21 (S I Figure 21).** This composite geological illustration of the Kedungbrubus- and Butak-area (Figures 1b and 13) highlights: The Pucangan, Kabuh and Notopuro Formations as mapped by J. Duyfjes (1936); Dubois' 1890s exploration and mapping over a broader area (black); and a modern topographic base (rust-colored). The *Homo erectus* mandibular fragment that Dubois' field team discovered in 1890 probably came from the band of fossiliferous Kabuh Formation outcrop that extends from Kedung Duren (***KD***) to Kedung Brubus (***KB***) and Kedung Lumbu (***KL***). Kedung Lumbu is the former village, *"about 2 km southeast of Kedung Brubus"* (Dubois 1907: 451, S II-F6), near the discovery spot of the mandible, according to Dubois' diary (S II-B1f and -B1g). After a perceptive analysis of the local geology in 1890 (S II-D2), Dubois returned to the Kedungbrubus collection area in 1894 (S II-B7), when he completed collecting the fossils in the Dubois Collection that are the basis for the Kedung Brubus fauna (Tables 5 and 6-C, main text). Both Dubois and Duyfjes (1934) excavated fossils from the Butak bonebed (labeled "1934 excavation" on the map; Table 5 and 6-D, main text; S II-B7a); the bed was in the lower Pucangan Formation of Duyfjes (Huffman 2020: 20-21, 24-25, 32; also Suminto et al. 1995a,b). See also Albers and de Vos 2010, Shipman 2001, Storm 2012, Tobias 1966.

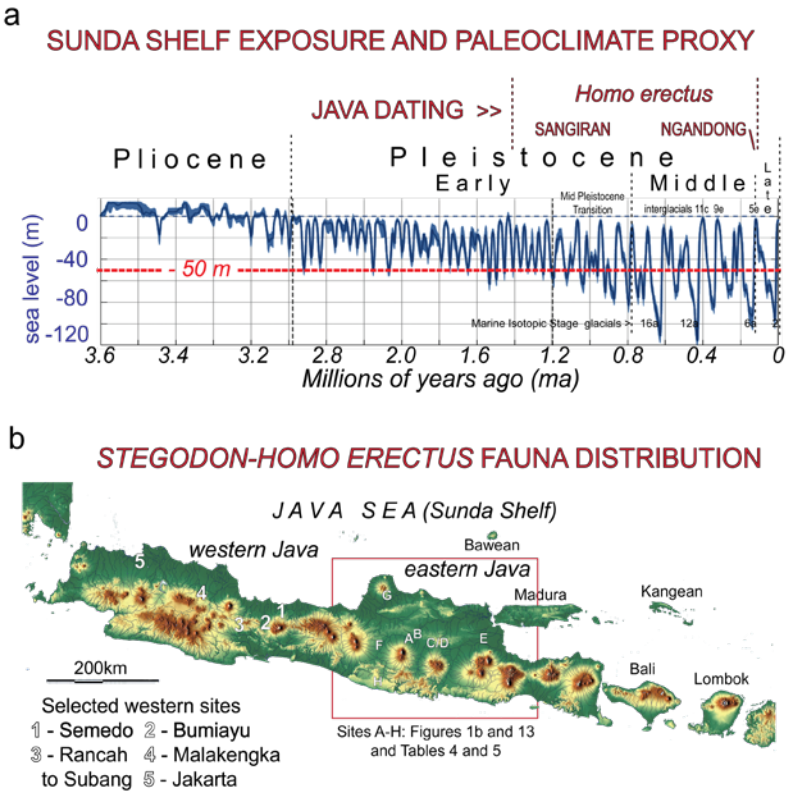


**Supplementary Materials I Figure 22 (S I Figure 22).** (**a**) Southern Sundaland experienced profound changes in landscape and climate during the Pleistocene, as illustrated by a proxy curve for global sea level (SL, blue; Berends et al. 2020). The Sunda Shelf (beneath the modern Java Sea) was exposed and inundated to varying degrees over time, and the climate changed repeatedly, while *Homo erectus* occupied the region (S Figures 23 to 24). For example, the 117-108 ka Ngandong *Homo erectus* bonebed (Table 6-B) postdates the last interglacial period (LIP, MIS 5e) at ~116-128 ka (Rohling et al. 2007; also, Shackleton et al. 2003), and the penultimate glacial maximum (PGM, MIS 6a) at ~140 ka (Cheng et al. 2006, Schneider et al. 2013; also, Railsback et al. 2015). Between lowstands and highstands, "Java Sea would have been neither [all] ocean nor dry land, but [included an ever changing] a complex … of lowlands, rivers, lakes, lagoons, shorelines, estuaries, and bays" (Huffman 2001: 242-243; also, Aziz et al. 1995). (**b**) The *Stegodon*-*Homo erectus* (*S.-H.e.*) faunas extended beyond eastern Java sites (**A-H**) to sites in western Java (also, Zaim 2010). Examples (**1-5**) are shown in relation to modern coasts, lowlands, rivers, mountains and volcanoes (green/dark tones transition to yellow/light and brown/medium at higher elevations). **1** and **3**: Archaic hominin fossils have been discovered at Semedo and Rancah >200km west of Sangiran Dome (e.g., Kramer et al. 2005, Noerwidi and Siswanto 2014, Noerwidi et al. 2016, Siwanto and Noerwidi 2014, Widianto and Noerwidi 2020). **2**: Vertebrate fossils have been collected since the 1930s from a thick Pleistocene sequence at Bumiayu, where the type section for the oldest *S.-H.e.* unit, the Ci Saat fauna, is located (e.g., Sondaar 1984, Suhiryogi et al. 2019, ter Haar 1929, 1935, van der Maarel 1932). **4**: A series of *S.-H.e.* sites have been identified from Majalengka to Subang in an outcrop belt along the north flank of the mountains in West Java (Hertler et al. 2007, Insani et al. 2015, Wibowo et al. 2019, Widianto and Noerwidi 2020, Zaim 2010, Y. Zaim, pers. comm. 2020). (**5**): Fossils representing the Trinil fauna occurred as far west as Jakarta (~450km west of Patiayam), where a core contained *"a fragment of the left side upper jaw of Sus brachygnathus"* (a Trinil fauna species); three marine intervals occurred there within a predominantly non-marine section, and palynological study identified Pleistocene mangroves to mountain forests taxa (De Neve 1950, Marks 1956, Yulianto et al. date unknown, Eko Yulianto, pers. comm., 2006).

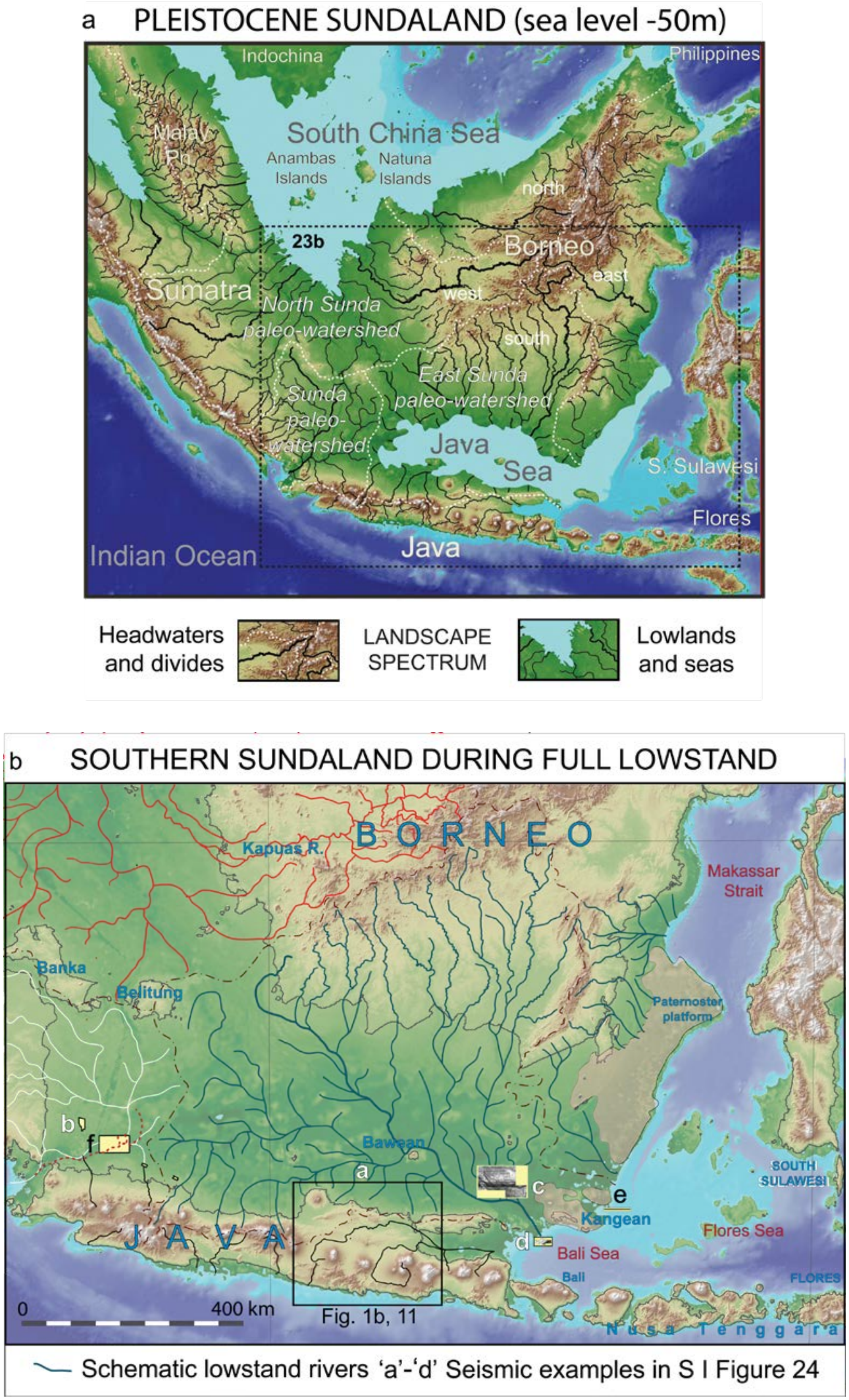
a PLEISTOCENE SUNDALAND (sea level -50m)
Indochina
Philippines
South China Sea
Malay Pn
Anambas Islands
Natuna Islands
north
23b
Borneo
east
Sumatra
North Sunda paleo-watershed
west
south
East Sunda paleo-watershed
Sunda paleo-watershed
Java Sea
S. Sulawesi
Flores
Indian Ocean
Java
Headwaters and divides
LANDSCAPE SPECTRUM
Lowlands and seas
b SOUTHERN SUNDALAND DURING FULL LOWSTAND
BORNEO
Kapuas R.
Makassar Strait
Banka
Belitung
Paternoster platform
b
f
Bawean
a
c
e
Kangean
SOUTH SULAWESI
JAVA
d
Bali Sea
Flores Sea
Bali
FLORES
0
400 km
Fig. 1b, 11
Nusa Tenggara
Schematic lowstand rivers 'a'-'d' Seismic examples in S I Figure 24

**Supplementary Materials I Figure 23 (S I Figure 23).** [preceding page] Schematic maps of southern Sundaland when sea level was significantly below the present (after Huffman et al. 2012, 2013; also, Voris 2000). The presentation is primarily applicable to the last glacial period (LGP) since the mapping is primarily based on the modern bathymetry of the Java Sea, but also should reflect conditions during the preceding glacial period (PGP), if not also earlier ones. (**a**) Southern Sundaland is divisible into four major watersheds. Three of them are especially relevant to the *Stegodon*-*Homo erectus* (*S.*-*H.e.*) faunas. The *East Sunda paleo-watershed* drained portions of Central Java and south Borneo, and flowed into the paleo-Java Sea. The *Sunda paleo-watershed* had headwaters in portions of West Java and South Sumatra, and led to Sunda Strait. The *North Sunda paleo-watershed* drained much of Sumatra and west Borneo, and emptied into the South China Sea. When Middle and Late Pleistocene sea level was below -50m, exposed areas of the Sunda Shelf would have facilitated faunal interchange between the Malay Peninsula, Borneo, Sumatra and Java via the low headwaters and divides in the present-day area of Banka, Belitung and Karimata islands. (**b**) During lowstands of ~125m below modern sea level, the Java Sea portion of the Sunda Shelf expanded to 30 times the area of modern Java (~$1x10^6$ $km^2$ for Java compared to $4x10^6km^2$ for the Java Sea from the Sunda- to Karimata- and Makassar-straits to the west, northwest and northeast of Java, respectively). The locations of examples of subsea seismic expression of the paleo-watersheds, presented in S I Figure 24, are identified on the map by the ‘a’ through ‘d’ lettering.

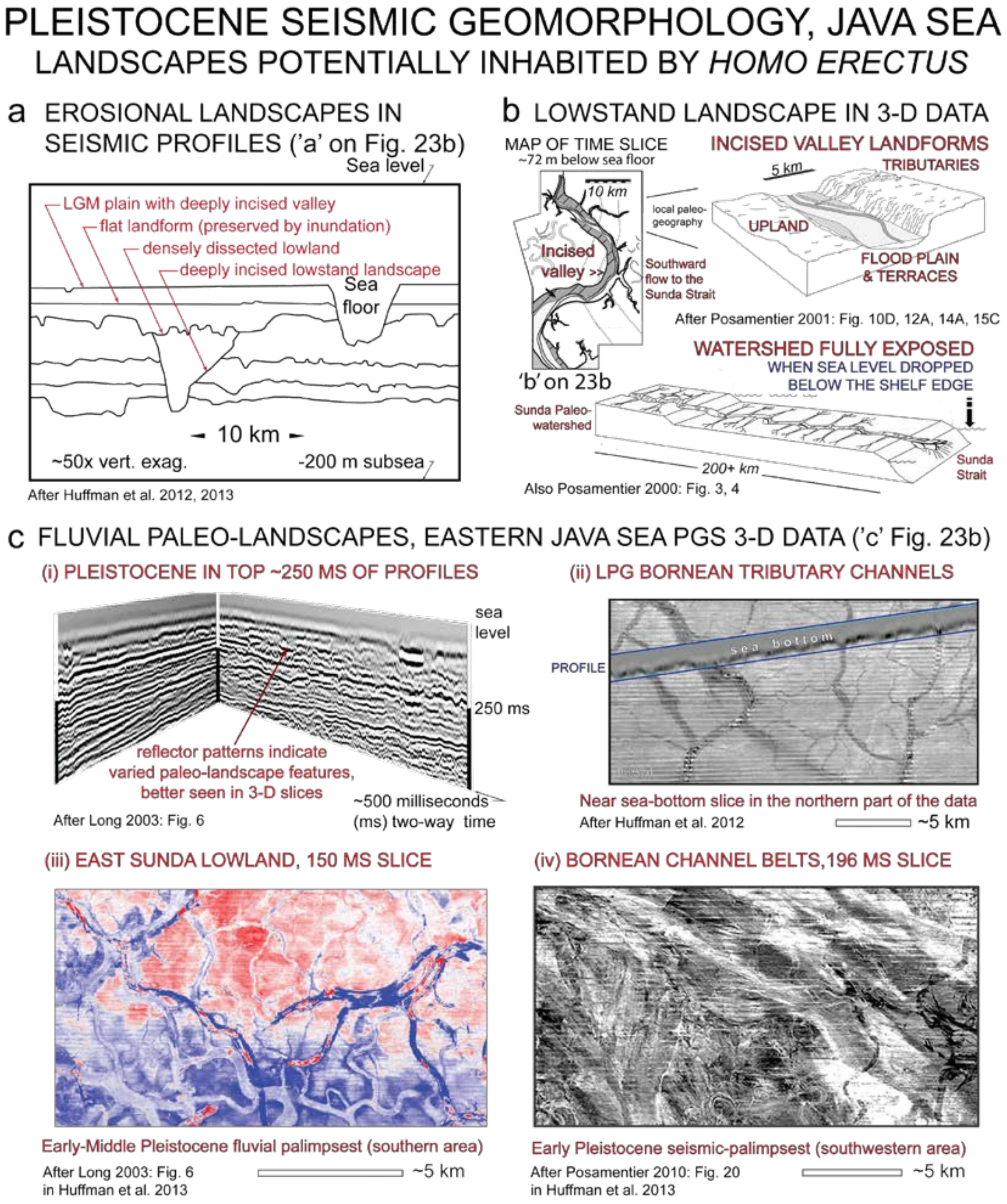


**Supplementary Materials I Figure 24 a-c (S I Figure 24 a-c)** (locations on S I Figure 23b). Seismic geomorphologic patterns in Pleistocene strata beneath the Java Sea reveal a variety of fluvial paleo-landscapes and (thereby) diverse conditions that large-mammal and hominin- populations might have inhabited. River valleys developed across the whole southern Sunda Shelf during lowstand conditions, as illustrated here in 'b' and 'c(ii).' Paleo-river systems reached the edges of the continental shelves during the Last Glacial Maximum (LGM, ~19 ka ago in MIS 2, 14-29 ka), when sea level was nearly -125m below the present (S I Figures 22a and 23a). During the Last Glacial Period (LGP) and earlier low-sea level periods seismic evidence indicates that braided river belts ('c (iv)') carried sandy and coarser-grained sediment from Borneo through the distal East Sunda paleo-watershed (for 'c;' also Posamentier et al. 2004). Seismic data from a 6500km$^2$ area southeast of 'b' ('f' in S I Figure 23b) details the drainage pattern of the Sunda paleo-watershed during the LGP (Gresko and Lowry 1996; also, Posamentier 2001, Susilohadi, 1995, Susilohadi and Soeprapto 2015).

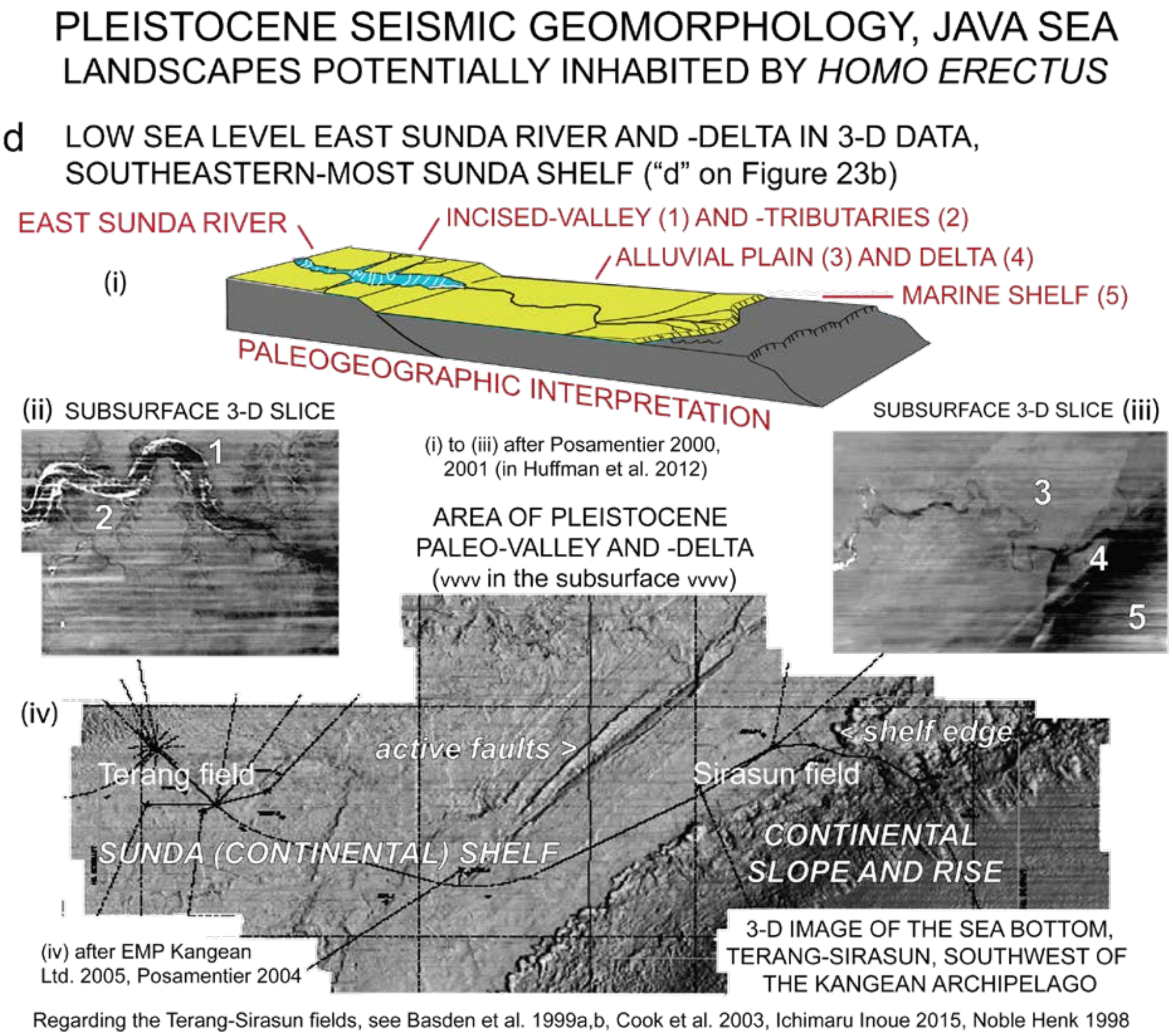


**Supplementary Materials I Figure 24d (S I Figure 24d)** (location on S I Figure 23b)**.** Subsea seismic data from the southeastern corner of the Java Sea, near the Kangean archipelago in far eastern Java, reveal a low-sea-level incised valley, prograding coastal plain and shore zone with small delta, all lying adjacent to a shallow marine platform that is west of the modern continental-shelf edge. Pleistocene physiographic conditions in the valley and plain should have been suitable for large-mammal occupation, including archaic hominin habitation (F.H. Henk, pers. comm. 2000). North of the Kangean archipelago at the southeastern corner of Sundaland ('e' in S I Figure 23b), seismic profiles reveal Plio-Pleistocene clino-forms representing repeated episodes of low sea-level shelf-slope progradation derived from the East Sunda paleo-river system (Bransden and Matthews 1992: Fig. 21, Granath et al. 2001: Fig. 9a).

*Geological Map of the Ngawi Quadrangle, Jawa, Sekala (Scale) 1:100,000*. Peta Geologi Bersistem Indonesia/Systematic Geological Map of Indonesia. Geological Research and Development Centre (Pusat Penelitian and Pengembangan Geologi), Bandung (Indonesia).

de Neve, G.A. 1950. Notitie over vertebraat-fragmenten uit de proefboring Kebajoran. *De Ingenieur in Indonesie IV. Mijnbouw en Geologie „De Mijningenieur"*. 5, 23-25.

de Vos, J., and Aziz, F., 1989. The excavations by Dubois (1891-1900), Selenka (1906-1908), and the geological survey by the Indonesian-Japanese team (1976-1977) at Trinil (Java, Indonesia). *Anthropological Society of Nippon* 97, 407-421. https://www.jstage.jst.go.jp/article/ase1911/97/3/97_3_407/_pdf

Dozy, C.M. 1911. Part III Die Arbeiten im Jahre 1908 von C.M. Dozy (mit Tafel II und IV). In *Die Pithecanthropus-Schichten auf Java: Geologische und Palaeontologische Ergebnisse der Trinil-Expedition (1907 und 1908)*, Selenka, M.L., Blanckenhorn, M., (eds.). Leipzig: Wilhelm Engelmann. [also, Berkhout and Huffman 2021]. pp. XL-XLII

Duyfjes, J. 1933. *Javakaarteering. Maandverslag over Mei/Juni 1933 een Tournee in Blad 93B Ngawi* [Java Mapping Program Monthly report for May/June 1933 a survey in Quadrangle 93B Ngawi (which include Trinil)]. [Internal unpublished type-written report of the Geologische Dienst, Bandung, Java; Reference no. E33-79 with 1:25,000 geological map]. Library of the Geololgical Research and Development Center (Bandung). 18 p.

Duyfjes, J. 1934. *Javakaarteering Maandverslag over Maart 1934 onderzoek op Blad 99b Madiun in de omgeving van Kedungbrubus, Pusat Penelitian dan Pengembangan Geologi* [Report No. 20/G/34; Ref. No.E34-62]. Library of the Geololgical Research and Development Center (Bandung). 18 p. [with handwritten note on file copy:] Bijlagen [Attachments], 1 geol. kaart omgeving Kedoeng Broeboes [1 geological map K. Brubus area]; 1 geol. Kaart Bl. 110B met blad profielen [1 geol map of Sheet 110B with cross sections for this sheet].

Duyfjes, J. 1935. *Javakaarteering: Maandverlag over Nov./Dec. 1934 van Ir. J. Duyfjes* [Java Mapping Program Monthly Report for Nov./Dec. 1934 by Ir. J. Duyfjes]. [Internal unpublished type-written report dated March 18, 1935, of the Geologische Dienst, Bandung, Java]. Reference E-34-53.] 30 p.

Duyfjes, J. 1936. Zur Geologie und Stratigraphie des Kendenggebietes zwischen Trinil und Soerabaja (Java) [On the Geology and Stratigraphy of the Kendeng Hills between Trinil and Surabaya (Java)]. *De Ingenieur in Nederlandsch-Indië, IV. Mijnbouw & Geologie, De Mijningenieur*. 3, 136-149 [translation in Huffman 2020].

Gresko, M.J., and Lowry, P.  1996. Seismic expression and channel morphology of a Recent incised-valley complex, offshore Northwest Java. In *Proceedings of the International Symposium on the sequence stratigraphy in S.E. Asia. Jakarta 1995*, C.A. Caughey et al. (eds.). Indonesian Petroleum Association. (IPA), pp. 21-33.

Granath, J.W., Christ, J.M., Emmet, P.A., and Dinkelman, M.G. 2011. Pre-Tertiary sedimentary section and structure as reflected in the JavaSPAN crustal-scale PSDM seismic survey, and its implications regarding the basement terranes in the East Java Sea. In *The SE Asian gateway: history and tectonics of Australia-Asia collision*, R. Hall et al. (eds.). Geological Society of London, Special Publication 355, pp. 53-74.

Hertler, C., Rizal, Y., and Zaim, Y. 2007. Habitat differentiation in the Pleistocene of Jawa - Introduction of the new Pleistocene fossil locality Majalengka. *Courier Forschungsinstitut Senckenberg* 2007, 165-174.

Hill, M., Huffman, F., Kappelman, J., Todd, L., and de Vos, J. 2015. *Taphonomic history of the Homo erectus site at Trinil, Java.* Paleoanthropology Society [Poster abstract]. http:/grsle.org/taphonomy/Hill_et_al_2015.pdf

Huffman, O.F. 2001. Plio-Pleistocene environmental variety in eastern Java and early *Homo erectus* paleoecology - a geological perspective. In *Sangiran: Man, Culture, and Environment in Pleistocene Times,* Simanjuntak, T., Prasetyo, B., Handini, R. (eds.). Proceedings of the International Colloquium on Sangiran Solo, Indonesia, 21st-24th September 1998, Jakarta. The National Research Centre of Archaeology/Ecole Francaise d'Extreme-Orient/Yayasan Obor Indonesia, pp. 231-256.

Huffman, O.F. 2016. Initial report of O.F. Huffman on August 9-19, 2016, fieldwork, ARKENAS-Leiden University project, Studying Human Origin in East Java at Trinil. (August 27, 2016). 41 p.

Huffman, O.F. 2020 (unpublished). *Annotated translation of J. Duyfjes' influential 1936 paper on the stratigraphy of the Kendeng area between Trinil and Surabaya, Java (by B.W. Seubert), and related materials* (from Huffman, A.W.J. Berkhout and Shimon Cohen). doi.org/10.13140/RG.2.2.28619.92964

Huffman, O.F., de Vos, J., Berkhout, A.W., Aziz, F. 2010. Provenience Reassessment of the 1931-1933 Ngandong *Homo erectus* (Java), Confirmation of the Bone-Bed Origin Reported by the Discoverers. *PaleoAnthropology* 2010:1-60. http://www.paleoanthro.org/media/journal/content/PA20100001.pdf

Huffman, O.F., A. Balzeau, J. de Vos, Y. Zaim, J. Kappelman. 2012. A 'Black Hole' in Southeast Asian *Homo erectus* Biogeography. Abstracts of the Paleoanthropology Society 2012 Meeting. *PaleoAnthropology* 2012: A0001-A0039. http://www.paleoanthro.org/posters2012/Huffman_2012poster.pdf

Huffman, O.F., Posametier, H.W., Wood, L.J., de Vos, J., Balzeau A., Berkhout A.W.J. 2013. Quaternary glacioeustatic record and hominin Paleobiogeography of the Sunda lowlands, Southeast Asia. Abstract 22st Annual Meeting, Honolulu, Hawaii, the April 2-3, 2013, Paleoanthropology Society 2013: A1-A41. https://paleoanthro.org/media/journal/content/PAS2013A_1.pdf

Huffman, O.F., Albers, P.C.H., Berkhout, A.W., and de Vos, J. 2015. *Did Trinil Femur I originate from a younger formation than the Homo erectus skullcap? Implications of early site photographs and other provenience records.* (Paleoanthropology Society Meeting Abstracts, San Francisco, CA, 14-15 April 2015; Poster). PaleoAnthropology 2015, A15. http://www.paleoanthro.org/media/meetings/files/Huffman_etal_2015.pdf http://www.paleoanthro.org/media/journal/content/PAS2015A.pdf

Huffman. O.F., de Vos, J., Albers, P.C.H., and Berkhout, A.W. 2018. *Trinil's Main Bonebed (Java) and Homo erectus paleobiogeography*. (Abstracts of the 2018 Paleoanthropology Society Meeting; Poster). PaleoAnthropology 2018:A15-A16. http://www.paleoanthro.org/media/meetings/files/Huffman_etal_2018.pdf http://www.paleoanthro.org/media/journal/content/PAS2018A.pdf

Ichimaru, Y., and Inoue, H. 2015. Exploration and development of Kangean block, East Java, Indonesia. Japanese Assoc. *Petroleum Technologists* 80, 19-26. (https://www.jstage.jst.go.jp/article/japt/80/1/80_19/_pdf/-char/en)

I.J.J.S.T. (Indonesia-Japan Joint Study Team) 1992. Peta geologi lingkungan kuarter daerah Solo-Madiun. Quaternary environmental geologic map of the Solo-Madiun area (1:100,000). Kerjasama Indonesia-Jepang pada Penelitian Lingkungan Geologi Kuarter. Indonesia - Japan Joint Study Team on Geology of the Quaternary Environment. Pusat Penelitian dan Fengembangan Geologi (Geological

Research and Development Centre). 2 sheets (with cross sections and detailed legend descriptions). [also, Appendix 1 In *Geology of Quaternary environment of the Solo-Madiun area, Central Java (GRDC-JICA Joint Study on Geology of Quaternary Environment, Bandung, April 1988 to March 1992),* Sudijono, Mano, K., and Wikarno, R. (eds.). Special Pub. 17; Geological Research and Development Centre, Bandung). 128 p.]

I.J.R.C.P. (Indonesia – Japan Research Cooperation Programme CTA-41). 1979. Progress report on the Indonesia – Japan joint research project on geology of human fossil bearing formations in Java. *Bulletin of the Geological Research and Development Centre* 1, 47-60. [also Soeradi et al. 1985].

Ingicco, T., de Vos, J., and Huffman, O.F. 2014. The Oldest Gibbon Fossil (Hylobatidae) from Insular Southeast Asia: Evidence from Trinil, (East Java, Indonesia), Lower/Middle Pleistocene. *PLoS One* 2014. 9(6). doi.org/10.1371/journal.pone.0099531

Insani, H., Wibowo, U.P., Setiyabudi, E., and Kurniawan, I. 2015. On variation of extinct Java hippopotamuses: A note from a new finding of Hippopotamidae fossil from Subang, West Java, Indonesia. *Proceedings Joint Convention Balikpapan* 2015, HAGI-IAGI-IAFMI-IATMI, 5–8 October 2015.

Joordens, J.C.A., d'Errico, F., Wesselingh, F.P., Munro, S., de Vos, J., Wallinga, J., Ankjærgaard, C., Reimann, T., Wijbrans, J.R., Kuiper, K.F., Mücher, H.J., Coqueugniot, H., Prie, V., Joosten, I., van Os, B., Schulp, A.S., Panuel, M., van der Haas, V., Lustenhouwer, W., Reijmer, J.J.G., and Roebroeks, W. 2015. *Homo erectus* at Trinil on Java used shells for tool production and engraving. *Nature* 518, 228-231.

Kramer, A., Djubiantono, T., Aziz, F., Bogard, J.S., Weeks, R.A., Weinand, D.C., Hames, W.E., Elam, J.M., Durband, A.C., and Agus. The first hominid fossil recovered from West Java, Indonesia. *Journal of Human Evolution* 48, 661-667.

Long, A. 2003. The Resolution Revolution. Technical Focus. *Petroleum Exploration Society of Australia News* 66, 51-53.

Marks, P. 1956. *Foraminifera-Ketjil dari Sumur No. 1 (Sumur 1) di Kebajoran, Djakarta. Smaller Foraminifera from Well No. 1 (Sumur 1) at Kebajoran, Djakarta.* Republik Indonesia Kementerian Perekonomian Djawatan Geologi, Bandung. Publikasi Keilmuan NR. 30, Seri Paleontologi, 25-29.

Noerwidi, S., and Siswanto. 2014. Alat batu situs Semedo: keragaman tipiologi dan distribusi spasialynya. Stone tool from Semedo site: its typology diversity and spatial distribution. *Balai Arkeologi Yogykarta* 34, 1-16.

Noerwidi, S., Siswanto, Widianto, H. 2016. Giant Primate of Java: A New Gigantopithecus Specimen from Semedo. Berkala Arkeologi 36(2), 141–160. https://doi.org/10.30883/jba.v36i2.241

Noble, R.A., and Henk, F.H. 1998. Hydrocarbon charge of a bacterial gas field by prolonged methanogenesis: an example from the East Java Sea, Indonesia. *Organic Geochemistry* 29, 301-314

Oppenoorth, W.F.F. 1911. Arbeitsbericht über die Ausgrabungen. I. Teil. Die Arbeiten des Jahres 1907 bis August [Report on the Excavation Work. Part I. Work up to August 1907]. In *Die Pithecanthropus-Schichten auf Java: Geologische und Palaeontologische Ergebnisse der Trinil-Expedition (1907 und 1908),* Selenka, M.L., and Blanckenhorn, M. (eds.). Wilhelm Engelmann, Leipzig. pp. XXVI-XXXVIII (288 p.). [also, Berkhout and Huffman 2021]

Oppenoorth, W.F.F. 1932. *Homo (Javanthropus) soloensis*, *Een Pleistoceene Mensch van Java* [*Homo (Javanthropus) soloensis*, A Pleistocene Human from Java]. Voorloopige Mededeeling [Provisional Communication]. Wetenschappelijke Mededeelingen [Scientific Communications] 20. Dienst van

den Mijnbouw in Nederlandsch-Indië, Batavia, pp. 49-79 (see also, partial re-publication of the English summary of this article, edited by Ales Hrdlicka, under the title of *"Homo (Javanthropus) soloensis"* in the 1933 *American Journal of Physical Anthropology* 17, 244-246).
Posamentier, H.W. 2000 (unpublished). *Incised valleys and fluvial systems: Insights from 3D Seismic data.* Presentation to Indonesian Petroleum Association (5 pages with 6 figures; H.W. Posamentier, personal communication, January 5, 2006).
Posamentier, H.W. 2001. Lowstand alluvial bypass systems: incised vs. unincised. *American Association Petroleum Geology Bulletin* 85, 1771-1793.
Posamentier, H.W. 2004. Seismic geomorphology: Imaging elements of depositional systems from shelf to deep basin using 3D seismic sata: Implications for exploration and development. *Geological Society, London, Memoirs* 29, 11. doi.org/10.1144/GSL.MEM.2004.029.01.02
Posamentier, H.W., Laurin, P., Warmath, A., Purnama, M., and Drajat, D. 2010. Seismic Stratigraphy and Geomorphology of Oligocene to Miocene Carbonate Buildups, Offshore Madura, Indonesia. In: Morgan, W.A., George, A.D., Harris, P.M., Kupecz. J.A., Sarg, J.F. (Eds.) Cenozoic carbonate systems of Australasia. (2010). pp.175-192. 10.2110/sepmsp.095.175. [For methods update, see Posamentier, 2020a. The application of seismic stratigraphy and seismic geomorphology to sequence stratigraphy and sedimentology. FOSI [Forum Sedimentologiwan Indonesia] Talk [Webinar] 16 May 2020 'Seismic Geomorphology and Sequence Stratigraphy by Henry Posamentier.' https://www.youtube.com/watch?v=3Col260pCdg ]
Pop, E., Adhityatama, S., Hilgen, S., Berghuis, H., Sutisna, I., Kostenko, A., Batenburg, J., and Joordens, J. 2020. *Quantitatively studying bone taphonopmy* [sic.] *of Homo erectus fossils from Trinil, Indonesia, using surface texture and CT analyses.* Nederlands Aardwetenschappelijk congres (NAC), Utrecht (NL).
Railsback, L.B., Gibbard, P.L., Head, M.J., Voarintsoa, N.R.G., and Toucanne, S. 2015. An optimized scheme of lettered marine isotope substages for the last 1.0 million years, and the climatostratigraphic nature of isotope stages and substages. *Quaternary Science Reviews* 111, 94-106..
Rizal, Y., Westaway, K.E., Zaim, Y., van den Bergh, G.D., Bettis, E.A., Morwood, M.J., Huffman, O.F., Grün, R., Joannes-Boyau, R., Bailey, R.M., Sidarto, Westaway, M.C., Kurniawan, I., Moore, M.W., Storey, M., Aziz, F., Suminto, Zhao, J., Aswan, Sipola, M.E., Larick, R., Zonneveld, J.-P., Scott, R., Putt, S., and Ciochon, R.L., 2021. Last appearance of *Homo erectus* at Ngandong, Java, 117,000–108,000 years ago. *Nature* 577, 381–385. doi.org/10.1038/s41586-019-1863-2
Rohling, E.J., Grant, K., Hemleben, C., Siddall, M., Hoogakker, B., Bolshaw, M., and Kucera, M. 2007. High rates of sea-level rise during the last interglacial period. *Nature Geoscience* 1. 38-42. 10.1038/ngeo.2007.28.
Ruff, C.B., Puymerail, L., Macchiarelli R., Sipla, J., and Ciochon, R.L. 2015. Structure and composition of the Trinil femora: Functional and taxonomic implications. *Journal of Human Evolution*. 80, 147-158.
Shackleton, N.J., Sánchez-Goñi, M.F., Pailler, D., and Lancelot, Y. 2003. Marine Isotope Substage 5e and the Eemian Interglacial. *Global and Planetary Change* 36, 151-155. doi.org/10.1016/S0921-8181(02)00181-9.
Selenka, M.L., Blanckenhorn, M., 1911. Die Pithecanthropus-Schichten auf Java: Geologische und Palaeontologische Ergebnisse der Trinil-Expedition (1907 und 1908) [The *Pithecanthropus* beds in

Java: Geological and Paleontological Results of the 1907 and 1908 Trinil Expedition], Leipzig: Wilhelm Engelmann. 288 p. http://www.biodiversitylibrary.org

Schneider, R., Schmitt, J., Köhler, P., Joos, F., Fischer, H. 2013. A reconstruction of atmospheric carbon dioxide and its stable carbon isotopic composition from the penultimate glacial maximum to the last glacial inception, Clim. Past, 9, 2507–2523, https://doi.org/10.5194/cp-9-2507-2013, 2013. https://doi.org/10.5194/cp-9-2507-2013 [with Supplement: Schneider, R., Schmitt, J., Koehler, P., Joos, F., Fischer, H. 2013. A reconstruction of atmospheric carbon dioxide and its stable carbon isotopic composition from the penultimate glacial maximum to the glacial inception. PANGAEA, https://doi.org/10.1594/PANGAEA.817041]

Shipman, P. 2001. The Man Who Found The Missing Link. Eugène Dubois and His Lifelong Quest to Prove Darwin Right. Simon & Schuster, New York.

Siswanto, Noerwidi, S. 2014. Fosil Probocidea dari Situs Semedo: Hubungannya dengan Biostratigrafi dan Kehadiran Manusia di Jawa. Berkala Arkeologi 34(2), 115–130. https://doi.org/10.30883/jba.v34i2.20

Soeradi, T., Shibasaki, T., Kadar, D., Sudijono, Itihara, M., Kumai, H., Hayashi, T., Furuyama K., Aziz, F., Siagian, H., Furutani, M., Suminto, and Yoshikawa, S. 1985. 4. Geology and stratigraphy of the Trinil area [with “Geological sketch map of the south bank of Bengawan Solo, east of Dubois’ monument, Trinil (after IJRCP, 1979)”]. In *Quaternary geology of the hominid fossil bearing formations in Java. Report of the Indonesia-Japan Joint Research Project CTA-41, 1976-1979,* Watanabe, N., and Kadar, D. (eds.). Geological Research and Development Centre. Special Publication No. 4. (378 pages), pp. 49-53.

Sondaar, P.Y. 1984. Faunal evolution and the mammalian biostratigraphy of Java. The Early Evolution of Man with Special Emphasis on Southeast Asia and Africa. *Courier Forschungsinstitut Senckenberg* 69, 219-235.

Storm, P. 2012. A carnivorous niche for Java Man? A preliminary consideration of the abundance of fossils in Middle Pleistocene Java. *Comptes Rendus Palevol* 11, 191–202.

Suharyogi, I.Y.P., Prasetyo, U., Wibowo, Insani, H., and Setiyabudi, E. 2019. *Duboisia santeng* (Bovidae, Artiodactyla) dari Bumiayu. [Universitas Padjadjaran, Fakultas Teknik Geologi] *Bulletin of Scientific Contribution: Geology* 17, 1–8.

Suminto, Utsugi, K., Kodama, K., Saefudin, I. 1995a. Geology of Mt. Lawu northern foothills. In: Sudijono, Mano, K., Wikarno, R. (Eds.), Geology of Quaternary environment of the Solo-Madiun area, Central Java (GRDC-JICA Joint Study on Geology of Quaternary Environment, Bandung, April 1988 to March 1992). (Special Publication 17; Geological Research and Development Centre, Bandung). pp. 45-52.

Suminto, Kimura, J, Hirayama, T. 1995b. Subsurface geology in Sangiran, Kedungbrubus and Ponorogo areas. In: Sudijono, Mano, K., Wikarno, R. (Eds.), Geology of Quaternary environment of the Solo-Madiun area, Central Java (GRDC-JICA Joint Study on Geology of Quaternary Environment, Bandung, April 1988 to March 1992). (Special Publication 17; Geological Research and Development Centre, Bandung). pp. 81-86.

Susanto, E.E., Mano, K., Sudijono, Sihombing, T., and Aziz, F. 1995. In *Geology of Quaternary environment of the Solo-Madiun area, Central Java (GRDC-JICA Joint Study on Geology of Quaternary Environment, Bandung, April 1988 to March 1992),* Sudijono, Mano, K., and Wikarno, R. (eds.). (Special Publication 17; Geological Research and Development Centre, Bandung). pp. 39-44.

Susilohadi, 1995. *Late Tertiary and Quaternary Geology of the East Java Basin, Indonesia.* Ph.D. thesis, unpublished, The University of Wollongong, Australia.

Susilohadi, S., and Soeprapto. T.A. 2015. Plio-Pleistocene Seismic Stratigraphy of the Java Sea between Bawean Island and East Java. *Berita Sedimentologi* 32, 1-16.

Ter Haar, C. 1929. *Boemi-Ajoe District.* Fourth Pacific Science Congress, Java 1929, Bandung, Excursion Guide E4, p. 1-15.

Ter Haar, C. 1935. *Geologische kaart van Java, 1:100,000. Toelichting bij blad 58 (Boemiajoe).* Dienst Mijnbouw Nederlandsch-Indie, Batavia. 50p.

Tobias, P.V. 1966. A re-examination of the Kedung Brubus mandible. *Zoologische Mededelingen 41, 307-320.*

van Benthem Jutting, T. 1937. Non marine Mollusca from fossil horizons in Java with special reference to the Trinil fauna. *Zoologische Mededelingen* 20, 83-180.

Van den Bergh, G.D. 1999. The Late Neogene elephantoid-bearing faunas of Indonesia and their palaeozoogeographic implications; a study of the terrestrial faunal succession of Sulawesi, Flores and Java, including evidence for early hominid dispersal east of Wallace's Line. *Scripta Geologica* 117. 419 p.

van der Maarel, F.H. 1932. Contributions to the knowledge of the fossil mammalian fauna of Java. *Wetenschappelijke Mededeelingen van de Dienst Mijnbouw van Nederlandsch-Indië* 15, 1-208.

van Es, L.J.C. 1931. *The Age of Pithecanthropus*. The Hague: Martinus Nijhoff. 142 p. (with geological maps).

von Koenigswald, G.H.R., 1934. Zur Stratigraphie des javanischen Pleistocän. *De Ingenieur in Nederlandsch-Indië , IV. Mijnbouw & Geologie, De Mijningenieur* 1, 185-201.

von Koenigswald, G.H.R. 1934/1935. *Rapport over de vindplaatsen van fossiele gewervelde dieren uit de Solo-vallei* [Report on the discovery sites of fossil vertebrate animals from the Solo valley]. Von Koenigswald Archive, Frankfurt a Mann, file number 19-01-03 (undated and unsigned copy of an unpublished report of the former Bureau of Mining [Mijnwezen] of the Netherlands Indies). 5p.

Voris, H.K. 2000. Maps of Pleistocene sea levels in Southeast Asia: shorelines, river systems, time durations. *Journal of Biogeography* 27, 1153-1167.

Wibowo, U.P., Erick Setiyabudi, E., and Kurniawan, I. 2018. A *Stegodon* Mandible from Cipanaruban, Subang, West Java; Description and its position in the Java Vertebrate Biostratigraphy. Sebuah Rahang Bawah *Stegodon* dari Cipanaruban, Subang, Jawa Barat; Deskripsi dan Posisinya dalam Biosostratigrafi Vertebrata Jawa. *Jurnal Geologi dan Sumberdaya Mineral* 19, 9 –14. ISSN 0853 - 9634, e-ISSN 2549 – 4759.

Widianto, H., and Noerwidi, S. 2020. It's time to look to the west: A new interpretation on *Homo erectus* findings distribution of Java. Saatnya menengok ke barat: Sebuah interpretasi baru tentang distribusi temuan *Homo erectus* di Jawa. *Berkala Arkeologi* 40, 153-178. doi.org/ 10.30883/jba.v40i2.598 https://berkalaarkeologi.kemdikbud.go.id

Yulianto, E., Hutasoit, L.M., Pindratno, H., Sukapti, W.S., and Hirakawa, K. Date unknown. *Plio-Pleistocene Boundary in Jakarta: Micropaleontological analysis.* Eko Yulianto, pers. comm., January 20, 2006. http://www.geocities.ws/ekoy001/foram-rev-2-web.htm

Zaim, Y. 2010. Geological Evidence for the Earliest Appearance of Hominins in Indonesia. In *Out of Africa I. Vertebrate Paleobiology and Paleoanthropology*, Fleagle J., Shea J., Grine F., Baden A., and Leakey R. (eds.). Springer, Dordrecht. Pp. 97-110 doi.org/10.1007/978-90-481-9036-2_7
