## Supplementary material for "Geology and discovery record of the Trinil *Pithecanthropus erectus* site, Java": S II

**SUPPLEMENTARY MATERIALS II NOTES (S II-…).** Author's notes for "Geology and discovery record of the Trinil *Pithecanthropus erectus* site, Java" (annotated, indexed and translated excerpts from Eugène Dubois archival documents and publications)

O. Frank Huffman
Department of Anthropology, University of Texas at Austin, SAC 4.102, 2201 Speedway Stop C3200, Austin TX 78712, USA.

Aart W.J. Berkhout
15706 Nedra Way, Dallas, TX 75248, USA.

Paul C.H. Albers
Naturalis Biodiversity Center, Darwinweg 2 2333 CR Leiden, The Netherlands.

John de Vos
Naturalis Biodiversity Center, Darwinweg 2 2333 CR Leiden, The Netherlands.

Fachroel Aziz
Geological Research and Development Centre, Jalan Diponegoro 57, Bandung 40122, Indonesia.

SECTIONS

**S II-A.** Discovery narrative in the Trinil letters of G. Kriele and A. de Winter's (KdW's), 1891-1900
**S II-B.** 1890-1893 Trinil discovery narrative recounted in Dubois' governmental submissions.
**S II-C.** 1890-1894 field notes of Dubois.
**S II-D.** Example geological exchanges between E. Dubois (ED) and R.D.M. Verbeek (RV).
**S II-E.** Other E. Dubois' documents concerning Trinil.
**S II-F.** Dubois' published accounts concerning the Lapilli bed (**LB**) from 1895-1896, 1907-1908 and 1924.

REFERENCES

CONVENTIONS: [Bracketed] entries in the following denote our comments. Where used, the underlining is our form of emphasis. Those fossil finds shown in the **red** font come from the left bank of the Solo River (Figures 1b and 7, main text); those in **blue** from the right bank; **black** font means that the side of the river of a fossil was uncertain. The following information is used in Table 1 and the Endnotes, main text.

NOTES: Dubois' whereabouts in 1890-1895 were largely deduced from entries in his diary. Naturalis scan codes for individual documents in the Dubois Collection have the form MM774-0000##-### (e.g., MM774-000007-203). The abbreviation used here is M…###-### (e.g., M…007-203). The materials were scanned in the order that they were stored physically so that related materials often have consecutive numbers.

**S II-A. . Discovery narrative in the Trinil letters of G. Kriele and A. de Winter's (KdW's), 1891-1900**. Citations in the text of the paper to these letters have the form of 'S II-A1a.' The inferred taxonomic identifications of the finds are in brackets. Those in **Red** font are from the left bank … in Blue, the right bank … in black, unspecified side of river. Salutations and closing are omitted.

### S II-A1. Letters to Dubois concerning the 1891 excavations at Trinil

**S II-A1a** [M…058-538]
Tinil 3 September 1891 [Thursday] [This letter was written by Kriele.]
I have been working here [on the left bank at Trinil] already for 3 days, as you had ordered me to do, and so far I have found various bones of different species. For instance, some small jaws of goats [***Duboisia santeng***] or kidangs [small cervid sp.], as well as a small goat horn and kidang horns. In addition, I found elephant [***Stegodon trigonocephalus***] and buffalo [***Bubalus palaeokerabau***] bones and small legs of a sort of animal that I don't recognize. The deposit of sand and rock where I am now working covers a large surface area [of outcrop]. …  [On July 19-21, Dubois made a trip to the top of Mt. Lawu (Figure 1b, main text), and evidently visited Trinil on September 7-8.] [De Winter arrived at Trinil on September 14.]

**S II-A1b** [M…058-545]
Tinil [Trinil] 18 September 1891 [Friday] [This letter was written by Kriele.]
[Original note in blue writing states] See reverse side for map [which is M…058-547; see Fig. 3b, main text, and * below] I am sending you herewith [some] crates with bones, of which 3 crates are from the other side of the [Solo] river near Kritik [right-bank, near the current Trinil Museum]. The bones that were found at Tinil [= Trinil; referring here to the left bank] are all either from a depth interval 0.20 meters below [the dry-season low-stand] water level [LWL] or even with it [an interval defined in this paper as the Principal Fossil Zone, PFZ, Figure 2, main text]. The added excavation is about 0.75 m above river level [near the top of the LB]. De Winter is very satisfied with his work here, and likes the site as well. During his excavations [on the right bank] he found bones at the place which you had pointed out to me for him to work [on September 7-8]…. [The fossil concentration at 0.00 to -0.20 m was noted by de Vos and Sondaar (1982) and de Vos and Aziz (1989).]

* {[Annotations on the map relating to the bone-bearing outcrops on the right bank are as follows; also, Figure 3b, main text] De Winter's work site. [He] has found bones as well: deer horns [***Axis lydekkeri*** antlers] with a piece of that skull. [Annotations relating to the outcrop and excavations of left bank, which is the south side of the river] work site [of] Kriele. Bones. Bone-bearing spot which contained that tusk and skull of the [an] elephant [***Stegodon trigonocephalus*** cranium with tusk]. 4m. 3m. [to the northwest- and northeast-shorelines, respectively; the sketch indicates a triangular-shaped outcrop projecting ~5m into the river, extending ~15-17m along the bank, and covering ~40m$^2$]}.

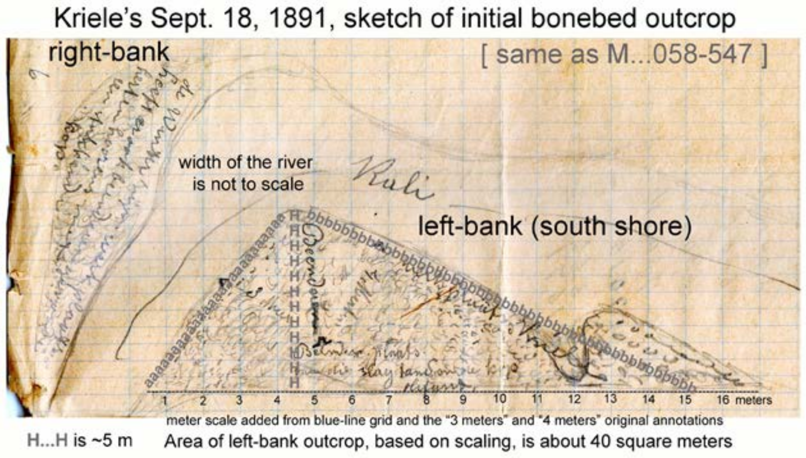


[Trinil was misspelled 'Tinil' on the map that Dubois initially had of the area (M…050-073), and some fossils in the Dubois Collection have 'Tinil' written on them in ink, linking them to the 1891 excavations (-A1m).

**S II-A1c** [M…058-540]

[This letter was written on behalf of Kriele and de Winter by Kriele, judging from the handwriting.]

Tinil [Trinil] 26 September 1891 [Saturday]

Herewith I am sending six crates with bones. If I am not mistaken, [the] really beautiful specimens are amongst those found at my site [Skullcap Pit]. For instance, there is a goat skull with 2 horns [***Duboisia santeng***] and a deer skull with two horns [***Axis lydekkeri*** with antlers] but one of them has a piece missing. Nice specimens have also been found at de Winter's site, for instance the skull of a banteng [***Bibos palaesondaicus***] almost completely preserved and deer horns [***Axis lydekkeri***]. That the bones are so fractured is not something we can do much about since we find them already broken up in the sedimentary rocks. [Kriele habitually referred to antlers as 'horens' in Dutch (horns in English), instead of 'gewei' in Dutch (antlers in English)]

[KdW's shipments were doubtless dispatched from the right-bank location that the Selenka Expedition (Selenka and Blanckenhorn 1911) later used as headquarters (Oppenoorth 1911), near the current-day Trinil Museum. A wagon track led south via Soko, and hence to the main east-west road to Ngawi, the nearest city. By packing crates on the right-bank, KdW avoided moving the boxes across the Solo by boat. The right-bank probably was where fossils were prepared and the men lived (S II- A4d '30-3-1897'), as was the case for the Selenka Expedition. Loads were probably taken to the Kedung Galar train station, which Dubois used to go to/from Trinil. Shipments could reach Dubois' home at Tangung Alugn (TA; Figure 1b, main text) within a day.]

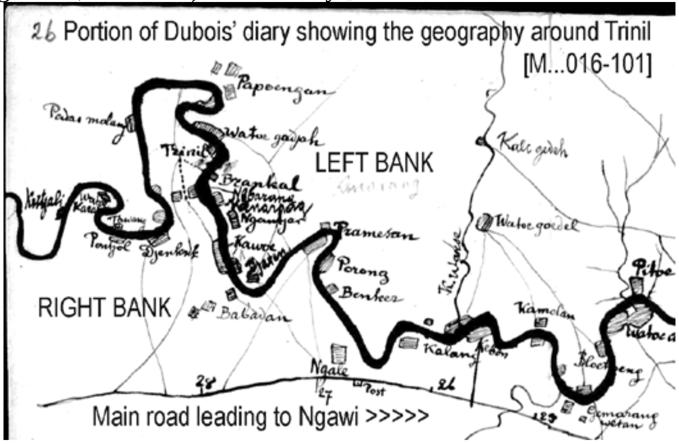


[Note that the map (color version in S II–E1, below) and other maps of Dubois' (-A4r, below), mistakenly show the section of the Solo River near his left-bank excavations as having been oriented in a SW- (downstream) -to-NE (upstream) direction. The river curved more sharply toward the west here so that the left bank at the *Pithecanthropus*-discovery site was nearly east-west in orientation (see Figures 6, main text).]

**S II-A1d** [M…058-547]

[This letter was written on behalf of Kriele and de Winter by Kriele, judging from the handwriting.]

Trinil; 5 October 1891

We collected our wages and the advances in Ngawi. You asked me if we had enough forced laborers. It would not be a bad idea if you could get more of them since work would then progress more rapidly. In my opinion it is not possible for us to start excavating the lower ledge, where the same amount of bones is being found, before the rainy season starts. De Winter is finding exclusively deer horns [***Axis lydekkeri*** antlers] and he has gathered already a sizeable collection [from his excavation on the right-bank]. … If a piece is missing from an excavated bone, it is mostly because it was hit with a pickaxe or a crowbar [which had to be used because of the induration of the LB] and after the bone has been hit, it is almost impossible to find the matching pieces.

[The brittle "bones" reflect the well-known stony fossilization of the skeletal material from the LB.]
[KdW had not recognized 1891 Molar, which the excavators had discovered in September.]

**S II-A1e** [M…058-548, right]
[Letters continue to be written by Kriele on behalf of Kriele and de Winter.]
Trinil; 11 October 1891 [Sunday]
I am sending you herewith 7 crates containing bones which have all been excavated from the same depth [that is, the same elevation relative to the river that was reported before, and thus the stratigraphic level PFZ, as defined herein]. The work is progressing very well, and we still keep finding many bones. At the site where I am working [on the left-bank excavations], the increase [of finds] is gradually lessening [perhaps because digging was below the PFZ in the lower LB] and it has become quite difficult to properly extract the bones. We have not found teeth of humans or apes, but we are now looking very carefully for them [evidently Dubois had informed KdW that the 1891 Molar had been recognized]. At de Winter's site [on the right bank], the same sand[stone] in which he had been finding bones remains [in the PFZ] about 0.20 meter below groundwater level and that is becoming a real hindrance.

[7 crates of fossils came from the PFZ, which was horizontal between the left and right bank; no fossil was mentioned that could be the Skullcap. Numerous KdW accounts contain references to the PFZ using various forms of expression (S II-A1b, -A1e, -A1i, -A1k, -A1i, -A1m, -A1o, -A1p, -A1q, -A1r, -A3ci and -A3cii).]

**S II-A1f** [M…058-549]
Trinil; 19 October, 1891 [Monday]
[No shipment was made; a variety of operational issues are described in the letter; the most relevant is:]… Due to the rainy season … water levels sometimes rise so much that it becomes impossible to build up tall enough dikes to block it. … I do not believe that we have thrown away bones or molars, since we promised a reward to the detainees for finding a human or ape molar, as well as for any other nice molar or tooth. [On the right-bank western shore], de Winter has found a human lower jaw with 4 molars and 4 teeth attached, but we believe that the specimen is not fossilized. Also, I have found [on the left-bank, south shore Skullcap Pit] a lower jaw of a hippopotamus with 2 molars attached. [Dubois (1908) later indicated that no hippopotamus fossils had been found at Trinil]. [Dubois visited Trinil October 21-24, after announcing the Skullcap find (-B2c).]

[KdW did not mention any fossil that could be the Skullcap. Dubois' assistant Dr. J.J.A. Bernsen noted in his diary on February 19, 1931, that Dubois said they thought it was the carapace of a tortoise, an idea that has subsequently appeared in several published accounts (Leakey and Slikkerveer, 1993; Shipman, 2001: 141; van Veen 2004). Dubois' October memorandum of 11-17-91 notified the Indies government of the Skullcap discovery (-B2f)]]

**S II-A1g** [M…058-552]
Tinil [Trinil]; 18 Nov. 1891 [Wednesday]
[no fossils sent, so the letter addressed operational issues, including] …. At de Winter's site [on the right-bank], a small pig jaw [***Sus brachygnathus***] has been found still containing a canine tooth. At the present time we are not finding many bones, but not a day goes by that we don't find at least something. Other than that, the work is progressing well …. [More than a month had passed with discoveries occurring at a slow pace; Dubois visited November 22-25.]

**S II-A1h** [M…058-555/556]
Tinil [Trinil] 7 Dec 1891 [Monday]
…. I regret to tell you that our house was already finished when we received your letter. We worked as we had agreed to do. When the river level had risen for weeks to the extent that we could no longer work below in the trenches, we began with 15 men on the house. All of them worked on it on the day that we went to Ngawi and that's how the job was finished in no time at all. We would very much like to stay together when working but if the river keeps on acting up it would impossible. The river rises 1 meter in one day and then subsides again the next day. Incidentally, since you left us the river level has not subsided to the previous low water level [LWL].

With respect to the finding of bones in the river embankment, we are sad to have to report that little has been found, since in a time span of 6 days no more has been collected than a bucket full. Those fossils were found at de Winter's site, about 0.75 m above the previous low water level. It turns out that this morning the river subsided again enough that we now can find bones again at 0.75 meter above the previous low water level. At our new work site nothing is found at this level. Below, we cannot get to that level. …

[see Dubois' November, 1891, memorandum (-B2h) and Fourth-quarter-report, -B2j, below).]

**S II-A1j** [M…058-558]
Tinil [Trinil]; 8 December 1891 [Tuesday] [The letter, like most from Trinil, is in Kriele's handwriting.]
We are happy … today the river subsided by about 0.50 meters and this allowed us to work down below in the side ledge [of the upper (?) LB]. At de Winter's work site [on the right bank, therefore] they have found today more bones than ever before in one day. Also, after I wrote you yesterday that nothing had been found in the new excavation trench [started on the left bank], today I found there an elephant skull [***Stegodon trigonocephalus***] with even larger molars than I have ever seen before. Also [there was] a very similar skull, but completely fractured and other bones, so it appears that this area also contains a variety of bones after all. If the river remains at this level [near the LWL] for a few days we could still extract more bones, but if the river rises 0.75 meter, finding of bones will also be interrupted [meaning most fossils encountered were below +0.75 m], your serving workmen, Kriele and de Winter [Kriele's handwriting]

[see Dubois' November memorandum (-B2h).]

**S II-A1k** [M…058-542]
Tinil [Trinil] 13 December 1891 [Sunday]
I am sending you herewith 6 crates of bones, most of which were found at about 0.75 meter above the previous water level [LWL on the left bank]. Bones are also found at de Winter's [right-bank] work site, but at his site they are from about 1.50 meter above the previous water level [LWL]. We are working now both at the top [in the side or on the top of the incised embankment] and below [near the LWL] depending on the water level. If the level is low we [can] work below and when it rises we work on top so that there is no interruption of activities. After all, the formations on top must first be removed before we can get to the bones [below] so it appears that this area [of the embankment, once dug to sufficient depth] also contains a variety of bones after all. your serving workmen, Kriele and de Winter

[de Vos (1989: Table 1) notes that a ***Panther tigris*** right mandible (Dubois Collection no. 1479) was "found in December 1891," based the labeling of the specimen, and citing this M…058-542 letter, that most of the finds then had been made between 0.75 and 1.50 above the LWL; that is, in the upper LB and perhaps immediately above it.] [see Dubois' November memorandum written on 12-23-91 (-B2h, below).]

**S II-A1l** [M…058-559]]
Tinil [Trinil] 21 December, 1891 [Monday]
... I can say that all is going quite well, since there are days that the river level rises suddenly but then also drops again at the same rate. For instance, on the 17$^{th}$ and 18$^{th}$ of this month, water level was about 1 meter above the previous level [LWL, submerging the LB]. Then on the 19$^{th}$ and 20$^{th}$ water level dropped again to 0.25 m above the previous level [LWL], and at that level, we can find more bones [in the upper LB (?)] as well as higher up on de Winter's side embankment [perhaps at +1.50]. It would benefit us if you would come and visit us and see it all for yourself. … Kriele and de Winter [The letter is in Kriele's handwriting.]

[Dubois visited on December 25-27, presumably because KdW had notified him that the LB was again accessible] [Note that generally Kriele worked at the left-bank Skullcap Pit during the 1891 season, while de Winter worked at the right-bank excavation on the northwestern shore.] [Evidently, the fossils that were excavated during September, October and most of November 1891 came overwhelmingly from the PFZ (at the LWL to 0.20 m below it), but some were from the upper LB as much as 0.75 m and even 1.50 m above the LWL (see below, especially, -A1b, -A1j and –A1l, and also, -A1a, -A1e, -A1h, -A1i and –A1k). In mid-October, de Winter also reported finding an un-fossilized jawbone from an unspecified right-bank context. In his 1894 monograph, Dubois (1894a) stated that the Skullcap and 1891 Molar were found "1 meter from each other" (-B9).]

**S II-A1m** [labels, Dubois Collection from Trinil]

[At least 28 specimens in the Dubois Collection have the word “Tinil” hand-written in ink on their surface while in the field (e.g., S I Figure 20c). The “Tinil” spelling was used during in 1891, as recorded above, so that the finds can be attributed to the excavations of 1891. The condition of the Tinil” specimens is typical of the material in the Dubois Collection from Trinil. The right mandible of ***Panther tigris*** that Dubois reported in 1891 is identifiable as collection no. 1479 (de Vos and Sondaar, 1982). Specimens bearing the “Tinil” label are:

***Bibos palaesondaicus***: No. 4361, mandible sinistra P2, P3, P4, M1*. Bovidae Gen. indet., sp. indet.: No. 2605, mandible dextra M1, M2, M3; No. 2606, mandible sinistra M1, M2, M3; No. 2682, phalange I**; No. 2763, maxillary dextra P2, P3, P4, M1, M2.

***Axis lydekkeri***: No. 12015, cranial fragment; No. 16799, antler dextra proximal; No.16782., antler sinistra proximal; No. 0669, mandible sinistra M2; No. 5262, tibia sinistra distal; No. 5378, tibia sinistra distal; No. 6279, tibia sinistra distal; No. 5562, metacarpal dextra juvenile; No. 7991, metacarpal dextra proximal.

***Duboisia santeng*** No. 2042, cranium*; No. 2043, cranium*; No. 2074, horn core sinistra; No. 2061, horn cores sinistra and dextra; No. 2506, horn core sinistra; No. 2070, horn core dextra fragment; No. 2062, maxillary dextra P2, P3, P4, M2, M3; No. 2062, maxillary dextra P4, M1, M2, M3; No. 5815, femur dextra diaphyse proximal; No. 1678, metacarpal dextra.

***Rhinoceros sondaicus****:* No. 1979, $M^3$ superior sinistra fragment***; No. 0324. *Rhinoceros* sp. indet., Molar fragment.

***Garialis bengawanicus***: No. 1622, cranium fragment.

***Chitra*** sp. indet.: No. 2745, carapax.

* see Hooijer, D.A. 1958. Fossil Bovidae from the Malay Archipelago and the Punjab. *Zool. Verhand. Mus. Leiden* 38: 1-112. Also, de Vos and Sondaar (1982: 49).

** Noted in the collection catalog as having a crocodile bite mark.

*** see Hooijer, D.A. 1946. Prehistoric and fossil Rhinoceroses from the Malay Archipelago and India. *Zool. Med. Mus. Leiden* 26: 1-138.]

### S II-A2. Letters to Dubois concerning the 1892 excavations at Trinil

**S II-A2a** [M…058-654]
Tjaruban; 13 May 1892 [Friday]
… we left Kedungbrubus on the 11th …. Tomorrow we will … head for Paron.
[Paron is a town south of Trinil. Dubois made a trip to Trinil on May 13-16]

**S II-A2b** [M…058-655]
Ngawi; 28 May, 1892 [location of Ngawi relative to Trinil is shown on Figures 1b and 11c, main text]
… total number of workers at Trinil will then be 40 …

**S II-A2c** [M…058-657]
Trinil; 29 May 1892 [Sunday]
We visited Ngawi [10 km east of Trinil] and looked for a proa [the Indonesian sailboat with a triangular sail] …. The river is subsiding nicely albeit slowly and is now about 0.50 meter above the [lower] sand ledge [upper LB, perhaps the “Ledge on the left bank, Figure 4a, main text, and S I Figure 4a]. … Kriele and de Winter [Trinil had begun to be spelled properly, not as “Tinil.”]

**S II-A2d** [M…058-659/660]
Trinil 5 June 1892 [Sunday]
…. We cannot send you good news yet about Pitu [a fossil site east of Trinil; -A1c], because the river rose again by 60cm on June 3, which caused the water level to be too high for working there. But we are able to continue work satisfactorily at the old work spot since high water does not interfere there. …. your serving workmen, Kriele and de Winter

[“The old work spot” might refer to 1891 excavations at Pitu (see also, –B4a). The next day Dubois left for Trinil via the Kedung Galar train station, which is south of the Pitu site; he spent two full days near Trinil, leaving on June 9; the infrequency of letters in late June and July was apparently due to Dubois’ visits to Trinil during this period.]

**S II-A2e** [M…058-660]
Trinil 15 June 1892 [Wednesday]
Herewith [we ship] 3 crates of bones [evidently from Pitu and Trinil]. The river rose again yesterday by about 40 cm … , your serving workmen, Kriele and de Winter [Handwriting indicates that this letter was written by Kriele.]

[Dubois left for Trinil eight days later on June 24, and spent four full days at Trinil and Pitu before returning to TA on June 29. Dubois’ June memorandum (-B3b) indicates that because the fossiliferous layers (LB) were inundated on the left bank, a new excavation was started “from above into the river bank,” beginning the excavation in which Femur I was discovered during August (Figure 10, main text). The size and shape of this June-August workings are unknown but the discovery excavation was later incorporated into the 25-m; only a fifth of the PFZ, ~10 $m^2$, was excavated in the 25-m Trench by the end of 1892 (-B5c) with the remaining ~40 $m^2$ in the upper LB.]

**S II-A2f** [M…058-661]
Trinil 26 July 1892 [Tuesday]
Herewith [we ship] 12 crates of bones one of which contains all your packed belongings [which Dubois had left at Trinil when he quit fieldwork due to illness, below]. About the finding of bones here, [I can tell you that] up to now the same volume is being extracted [as last year; the PFZ, again being excavated, was highly fossiliferous]. Incidentally we have not found any new species yet. As we agreed, I will travel upstream next week to investigate other bonebed exposures [and collect paleo-current measurements on cross bedded deposits] along the river. … Kriele and de Winter [This letter was written by Kriele.]

[Dubois had been at/near Trinil July 5-14 and 18-20, amounting to 9 full days, and making a total for June and July of 14 full- and 8 partial-days; the 12 crates was a large shipment, especially since it appears to have contained the finds that KdW had been digging in the LB for only 10 days.]

[On Tuesday, July 19, 1892, the day before he left Trinil, Dubois wrote in his diary, “*Claystone beneath andesite: also in thin layers of 5 {or 3}; ½ foot higher [shallower] in between at Trinil.*” He might have been referring to either the claystone beneath the LB (S II-B3c), claystone in the diamicton-rich upper Pucangan Formation of Duyfjes (1936).]

**S II-A2g** [M…058-662/663/664]
Trinil 15 August 1892 [Monday]
Herewith [we are sending] 6 crates with bones, one of which contains a buffalo skull with molars in place [***Bubalus palaeokerabau***]. [There is] also a antelope skull [***Duboisia santeng***] and a deer skull [***Axis lydekkeri***]. On August 9 the river rose again so much that our work site became completely inundated and we were forced to work on the opposite [or other] side of the river [probably moving from the right to the left bank*]. In fact, that [inundation] is the reason for this rather small shipment.

Enclosed are also the measurements that I made for the slanted beds [crossbedding] from places where I could observe them best [see part of M…058-662, below]. We also searched for other find spots but could not find any. We did find nice exposures that we worked on with the same kind of sand[stone] as at Trinil, but we could not find bones in them. I searched all along the river as far as dessa [village of] Galon and past dessa Sondé [Kriele had worked up stream of Trinil for several kilometers]. … [M....058-664] The last letter that I received from you was registered and I even had to collect it in Ngawi. Was that actually a mistake? Please send us also a bit of glue … Kriele and de Winter [This letter was written by Kriele.]

[The six crates and large skulls suggest that finds KdW reported came from the PFZ; they made no mention of a fossil that could be Femur I.]

[M…058-662; Drawings (in Kriele's handwriting) of steeply dipping cross-bed sets with foreset faces inclined in the direction of paleocurrents]

North of Dessa Djengkrik  N to SW  [cross laminations dip N]
N.W. of Dessa Kritjak  SE to NW [cross laminations dip to the SE]
N.E. of Watu karas E to W [cross laminations dip to the E]

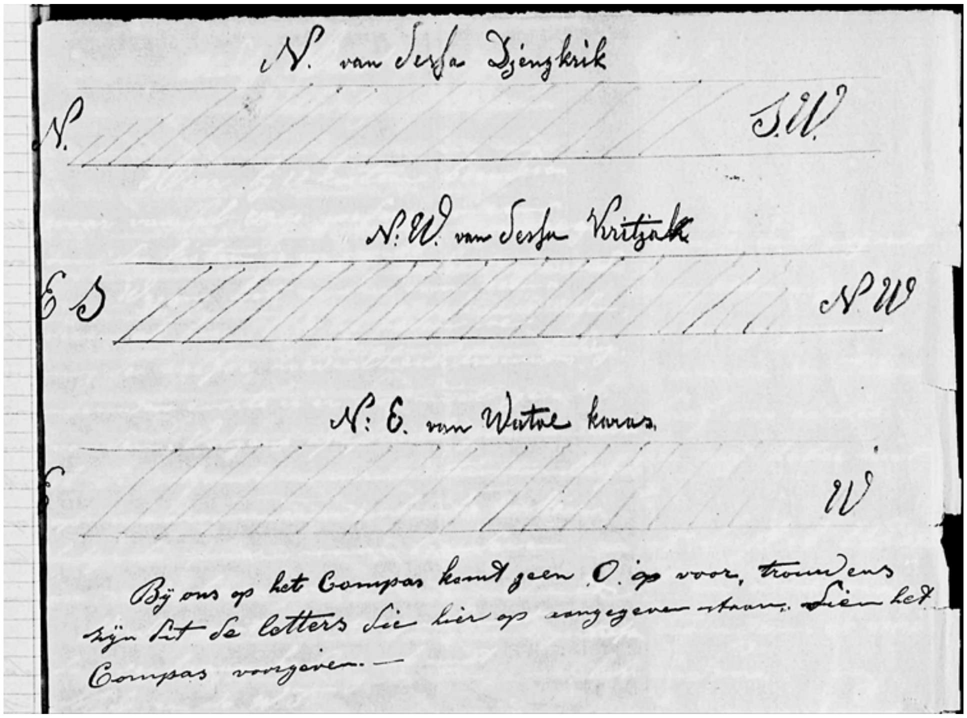


[The note at the bottom states] There is no letter O [Oost = East] on our compass and by the way these are the letters that are indicated on the compass. [Kriele evidently did not realize that the E in English on the compass under use was East, which would be Dutch Oost.] [The fieldwork mentioned in the July 26th letter as being planned for the "following week" would have taken place on Sunday July 31 to Saturday August 6.]

[* **The phrase "the opposite (or other) side of the river" [Dutch: "...aan de andere zijde van de rivier..."] is taken to mean the south-shore (left-bank) because:**

1. The phrase "work on the other side of the river" was used during 1892 (–A2j, and –A2m below) to mean the left bank, most importantly in the August 31 letter regarding Femur I, wherein the phrase "opposite side" is specified to mean "where the bone {the femur} was found" on the left bank (see also, -A1l and –A2n).
2. In April 1893, as KdW started the 40-m Trench, they called the left bank the "other side" of the river from the place where de Winter's crew was building "bone sheds" on the right bank (-A3a i-iii).
3. In 1896 and 1897, moreover, Kriele clearly used the phrase "opposite side of the river" to refer to the left bank (-A4c, especially pages 23, 25 and 29; -A4d).
4. The "opposite" side of the river evidently was across the Solo from the fossil shipping point on the right bank. This is where a cart track headed south to primary roads without crossing the Solo River, as explained in–A1c.

Inundation of one excavation and not the other could have occurred, for example, if higher elevation pits were being worked on one side than the other, or even if the the dike system along one shore was higher than on the other.]

**S II-A2h** [M…058-664]
Trinil 22 August 1892 [Monday]
[We] received the crates [that you returned to us], one of which contained the medicines as you promised us. We could not find any glue inside, though. One of the mandurs … was punished on the 9th of this month …. In our opinion, we again found attractive specimens: a pig [***Sus brachygnathus***] jaw with tusks and two small teeth in the front. There was also a small head [partially hidden word] that we think is of a fish, but because we are not sure, we are sending it to you immediately. The previous [finds of] bones have all been put together with the glue that you gave us [but would not be transmitted until a later shipment]. The river has been subsiding every day [and so I am asking you] should we search any further [for new sites] along the river? … Kriele and de Winter [This letter was written by Kriele.]

[No mention was made of a fossil that could be Femur I, and the most "attractive specimens" were the ***Sus brachygnathus*** mandible and "fish head. It is unclear whether the crates Dubois returned to Trinil were accompanied by a mid-August letter from him.]

**S II-A2i** [M…058-665]
Trinil; 30 August 1892 [Tuesday]
[We are sending] herewith 8 crates with bones, all from this side [right-bank *] of the river. During the last days of the month we were not able to glue the bones, because we were too busy extracting them. That is the reason that no glued bones are in this shipment. In the crate with this letter, you will find 6 deer skulls [***Axis lydekkeri***] and in a different crate there is also a buffalo skull with a row of molars [***Bubalus palaeokerabau***] still attached. All these bones were found in the new excavation work site on this [right-*] side of the river. The feet of K.L. have now healed and otherwise all is going very well with the work activities and the people, but there are a few who suffer from fever. your serving workmen, Kriele and de Winter [This letter was written by Kriele.]

[Again, no mention was made of fossil that might have been Femur I; KdW must have gotten Dubois' letter inquiring about Femur I after this letter was mailed on the 30th.] [* "This side" probably refers to the right bank by the reasoning described in –A2g, above]

**S II-A2j** [M…058-666/667]

Trinil, 31 August 1892 [Wednesday]

Most respected Sir

We received your letter and the [fossil] gluing process here will now be halted.

About the bone [**Femur I**], de Winter tells me that the pieces that are missing were blown away while on a teak leaf by heavy winds during the process of gluing and we cannot find them again.

The bone [femur] in question was found on the same side of the river where earlier the chimpanzee [***Pithecanthropus erectus*** **Skullcap**] was, and if de Winter remembers it correctly the following bones were found nearby: a mandible and tusk of an elephant [***Stegodon trigonocephalus***]. The bone [**Femur I**] was found at approximately the same depth [elevation] as the chimpanzee skull [**Skullcap**] even with the previous [dry-season] low-water level [LWL], and separated from it by about 12 meters.

Because of high water, we have done little further work on that [left-bank, south] side [since Femur I was discovered]. Between the 29$^{th}$ and 30$^{th}$ of August the water rose again by about one meter [which would have inundated the whole LB]. As soon as the water level subsides [we expect] that we can work on the opposite [southern, left bank] side where the bone [Femur I] was found. We will be searching carefully for the pieces that had been knocked from the bones.

Because of the fact that the water level is so variable, it is very difficult to find new ledges [perhaps referring to other sites along the river, mentioned in the August 22 letter].

Faithfully yours,

Kriele and de Winter [This letter was written by Kriele.]

[Hand written in red pencil above the salutation is:]

*"Femur P.e."*

[Hand written in the left-hand margin by an unidentified person is:]
*"This is the reporting letter that was thought to have been lost but which simply arrived late"*

[De Winter must have found Femur I because KdW referred to de Winter's memory in reporting about some aspects of the discovery and cleaning of the specimen. KdW letter of November 9, 1892, confirms that de Winter discovered Femur I (S II-A2q). High water prevented KdW from returning to the left bank excavation to search for the missing bone fragments that de Winter had lost from Femur I.]

[In stating Femur I came from "approximately the same depth as the chimpanzee skull" KdW put the find in the PFZ, the LB having been nearly horizontal in the excavation (S II-A1b, -A1e, -A2p, and -A2q). KdW's report about lost "pieces" indicates that de Winter gave the specimen some preparation. Their promise to search for the pieces on the left-bank when the river subsides indicates that de Winter's work on the fossil took place at the discovery excavation on or about the day of discovery. The lost material refers to fragments that Dubois (1926a) described as missing on the posterior epiphysis, losses that are still notable on the specimen (S II-F9, below).]

[MM774C-000058-667, right; A TRANSCRIPTION OF THE 31 AUGUST LETTER FOLLOWS]

*"Dit is de verloren gewaande, doch alleen te laat ontvangen dienstbrief*"

*Femur P.e.*

Trinil, 31 Augustus 1892

Weledele Heer!

U brief ontvangen en het lijmen zal alhier gestaakt worden.

Wegens dat been, verteld mij de Winter als dat die stukjes die eraan ontbreken door een zwaren wind op een djatie blad onder het lijme zijn weggewaaid en niet meer terug kunnen vinden.

Dat bedoelde been is gevonden aan dezelfde kant van de rivier waar dat vroeger die chimpansekop is gevonden. Zo de Winter zich zelven niet vergist dan lagen er deze beenderen in de buurt: een kaak en slachtand van een olifant. Het been is zoo wat ongeveert gevonden op die diepte van de chimpansekop, gelijk met de vroegere lagen waterspiegel, van elkaar zoowat 12 meter.

Aan die zelfde kant hebben wij te met niet meer gewerkt door als dat de waterstand aldaar te hoog is geweest. Tusschen 29 en 30 Augustus is de waterstand weer ongeveerd een meter gewaschen. Zoodra de waterstand weder gedaald is als dat aan de overkant of waar dat been gevonden is [MM774C-000058-667, left] kunnen werken zal men er zoo oplettend gelijk zijn om de stukjes die van de beenderen w[aren?] afgeslagen op te zoeken.

Door dat de waterstand zoo ongestadig [is], is het zeer lasting om nieuwe bankken te vinden.

U dienede werklieden

Kriele en de Winter

[MM774C-000058-666; SCANNED FIRST PAGE OF THE AUGUST 31st LETTER]

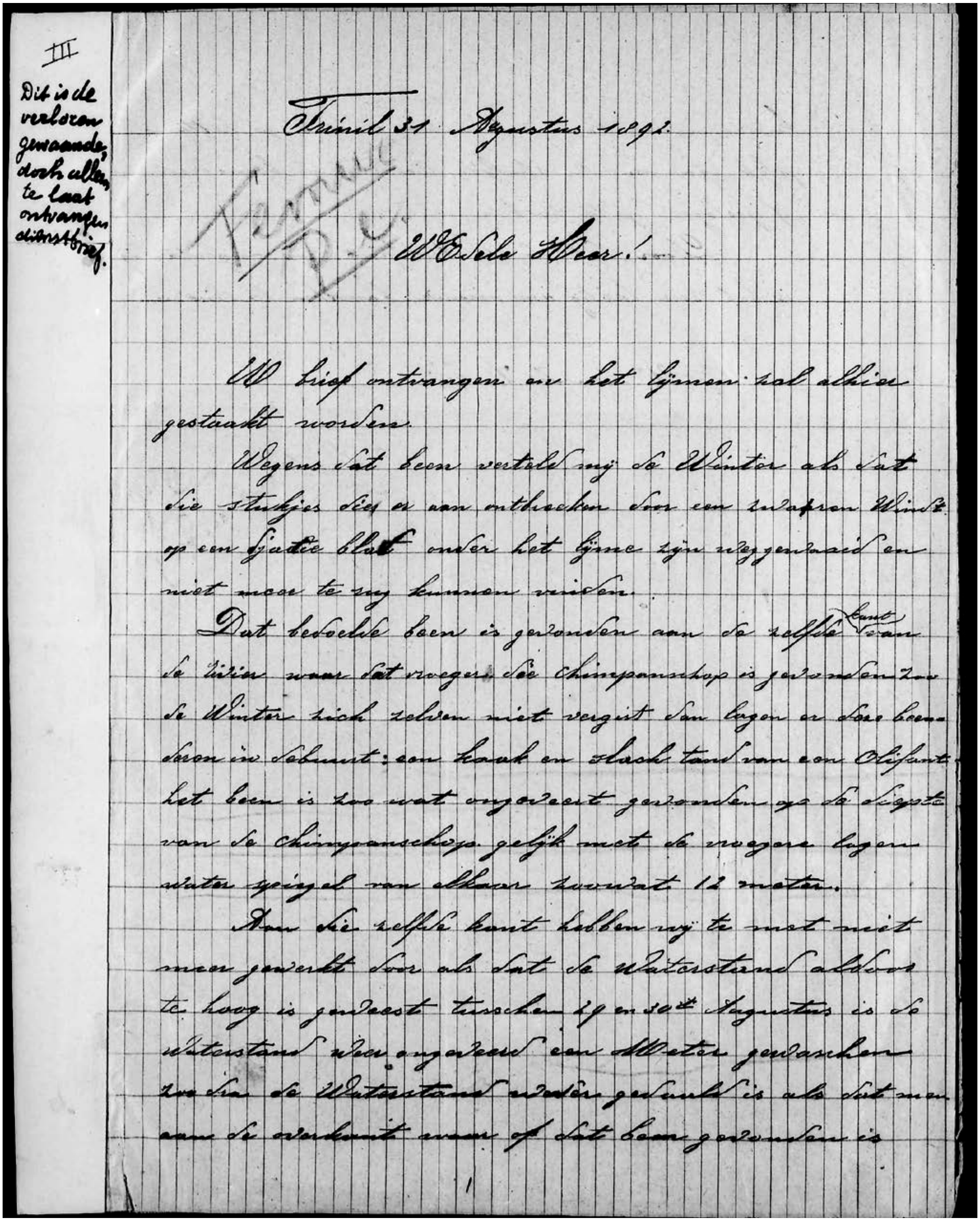

III

Dit is de verloren gewaande, doch alleen te laat ontvangen dienstbrief.

Trinil 31 Augustus 1892.

WEdele Heer!

UW brief ontvangen en het lijmen zal alhier gestaakt worden.

Wegens dat been vertelde mij de Winter als dat die stukjes die er aan ontbreken door een zwaren Wind op een djatie blad onder het lijme zijn weggewaaid en niet meer te zij kunnen vinden.

Dat bedoelde been is gevonden aan de zelfde kant van de rivier waar dat vroeger die chimpanschop is gevonden zoo de Winter zich zelven niet vergist dan lagen er daar beenderen in de buurt: een kaak en slachtand van een Olifant het been is zoo wat ongeveer gevonden op de diepte van de chimpanschop gelijk met de vroegere lagen water spiegel van elkaar zoowat 12 meter.

Aan die zelfde kant hebben wij te met niet meer gewerkt door als dat de waterstand aldoor te hoog is geweest tusschen 29 en 30e Augustus is de waterstand weer ongeveer een Meter gezakken zoo dra de Waterstand weder gedaald is als dat men aan de overkant waar op dat been gevonden is

1

**S II-A2k** [M…058-667/668]

Trinil 7 September 1892 [Wednesday]
Most respected Sir!

I cannot understand that you have not yet received the reporting letter that I sent already 7 days ago now. In that letter I wrote about where the [femoral] bone of the chimpanzee was found. That bone [Femur I] was found on the same side [of the Solo River] as the skull [Skullcap] and also at approximately the same depth and even with the previous low-water level [LWL], separated from each other by about 12 meters.

You are asking if we might find more of it [the chimpanzee skeleton]. To attempt that, it would be best to dig 3 to 4 meters deeper into the face [of the embankment to the south]. Incidentally, the spot where that bone [femur] was found has not been fully excavated. That will probably take another month or a little less.

Bones of this chimpanzee might possibly be found deeper. It would then be best to work that place first somewhat deeper, and after that dig farther into the face. This site would have been long finished but we have had too much hindrance of the water. We have been unable to work at it now for 10 days, but if the river continues to subside the way it has done for several days, we think that we might possibly return to work there again on Friday.

I have a polite request for some eye drops. My feet have healed. Incidentally I have had infected eyes for 4 days now and that gives me a lot of trouble when I am in the sun. Also, some quinine since we always have people with fever and our quinine supply is just about finished.

I am sending this letter registered since I am afraid that my [previous] reporting letter has been lost.
Faithfully yours, Kriele and de Winter [This letter was written by Kriele.]

[Hand written in pencil below the date on the letter is:] The letter of 31 August is the letter that was thought to have been lost. [The letter implies that: (i) KdW expected to encounter more fossiliferous rock below the level in which Femur I occurred; and (ii) KdW were sure that the PFZ extended beneath the strata in the discovery excavation back wall, so that expanding removing more of the embankment would access more PFZ. The "10 days" of flooded excavations were from August 29 to about September 11).]

[MM774C-000058-667, right; TRANSCRIPTION OF THE 7 SEPTEMBER LETTER]

Trinil 7 September 1892
[in Dubois' handwriting] (*De brief van 31 Augustus is de verloren gewaande dienstbrief*)
Weledelg. Heer! [faintly written by hand: "*femur*"]

Ik kan niet begrijpen als dat U die dienstbrief, dien ik voor 7 dagen al verzonden heb, waar in vermeld stond op welken plaats dat been van de bedoelde chimpan gevonden is, noch niet ontvangen hebt. Dat been is gevonden aan dezelfde kant waar dat de schedel is gevonden ook zoo ongeveerd op dezelfde diepte gelijk met de lagen water spiegel van vroeger, van elkander ongeveerd 12 meters. U vraagd hoe men er nog meer van zouden kunnen vinden, dan zou het besten en er het meeste kans toe zijn, een meter 3 a 4 dieper het wand in te graven. Trouwens is die plaats waarof dat been in gevonden is, nog niet afgewerk. Daar zal nog wel een kleine maand voor noodig zijn. Mogelijk zitten er nog wel beenderen van die chimpan lager, dus het is het besten eerst die plaats op zijn diepte te werken, en daarna dieper de wandt in [MM774C-000058-668] (te) graven. Die plaats had al lang afgewerkt geweest, maar te veel last van het water gehad. Nu hebben wij er al in geen 10 dagen in kunnen werken. Als de rivier zoo door blijft dalen, als dat hij nu een paar dagen doet, dan zullen wij er mogelijk Vrijdag er weder aan kunnen beginnen.

Een beleefd verzoek of ik niet een weinig oogwater kun krijgen. Mijnen voeten zijn genezen. Trouwens heb ik nu al reeds 4 dagen zieken oogen en daar ik (er) veel last van heb als wanneer ik in de zon kom, ook een weinig kinine, daar men aldoor koors leiders hebben en de kinine te met op is.

Ik zendt U deze brief maar als bestel als dat ik bang ben dat die dienstbrief verloren is gegaan.

Uw dienende werklieden

Kriele en de Winter

**S II-A2l** [M…058-608 with M…058-609, left, back of this letter, has drawing of a tooth or two]

Trinil; 21 September, 1892 [Thursday]
Things are currently very bad with the prison workers …. We are now 8 men short and if no one is ill, we can work with 29 …. Almost no bones are being found [because the excavations on the left bank were near the top of the embankment].

[Dubois visited Trinil on September 24-25; see –B4a and –E1b for his actions on the 23rd.]

**S II-A2m** [M…058-609]

Trinil; 4 October, 1892 [Tuesday]
The number of prison laborers has now expanded to 40 …. Work on the opposite side of the river [left-bank; see –A2g] progresses so well that we can expect to reach the top of the bone layers [LB in the 25-m Trench expansion] within 6 days [that is, October 10]. From today onward, 7 prison laborers are working on this side [right bank] and the rest on the opposite side of the river [that is, 33 men worked in the 25-m Trench on the left bank, and 7 were working at an excavation at an unknown location on the right bank]. De Winter has been complaining for days about stomach pains …. Kriele and de Winters

**S II-A2n** [M…058-610]

Trinil; 11 October, 1892 [Tuesday]
…. Of the 40 prison workers, we are missing 5 again … [Still] work progresses very well, although bones are not being found [and] the shipment of the 15th will be very small [because the LB had not yet been reached in the expanded trench]…. Kriele and de Winter

**S II-A2o**

(i) Trinil; 15 October, 1892 [Saturday] [M…058-612]
Hereby [we transmit] 4 crates [of fossils] …. we now are still working with just 35 men. Work progresses very well. We think we will reach the target bone layer in 6/7 days [that is, reach the PFZ in the 25-m Trench on October 21-22]. From the 13th on we have been working as a joint force again [both crews were on the left bank] since we have already started excavating in the uppermost bone layer [the upper LB]…. Kriele and de Winter [This letter was written by Kriele.]

(ii) Trinil; 21 October, 1892; Friday [M…058-468]
Herewith [we are sending] a smaller kind of thigh bone. We are not entirely sure, but it looks very much like the bone that has been drawn on the sketch. We are getting here already for 12 days now such a terrible amount of rain that [as a consequence] our new work site [on the left bank] is below water and [since] at this side of the river [right bank] we have possibly about 5 days of work left, so that when that has been completed and the river will not subside [we are asking] what we should then continue doing? At our new work site and counting from the top [of the embankment ~9 m above the LB in the 25-m Trench; *S II-B5a*], we have gone down already 6.25 meters and not many bones have been found in it. On the 20th our prisoner-worker labor force was increased to a total number of 40. Your serving workers, Kriele and de Winter

[Dubois came to Trinil on October 22-24, perhaps anticipating KdW's penetration of the PFZ; the 25-m-long, 11-m-high exposure presumably gave the all a clear view of the left-bank stratigraphy.]

**S II-A2p**

(i) Trinil; 28 October, 1892 [Friday] [M…058-615]
…. In regard to the river … its level is still at the same height as when you left here [four days ago] …. We think that the river will soon subside. With respect to the digging, that is progressing very well, although we are hindered a lot by ground water [seepage due an excavation depth below river level], but we can work in spite of that. For that reason we miss as many as 5 men during the day because they need to work at night [bailing and maintaining dikes], and we can do little to help that problem. Every day bones are being found, but nothing [more] of the chimpanzee [skeleton]. In the corners [of the 25-m Trench], we have progressed [below the top of the LB] to about the depth at which the leg bone and the skull were found [in the PFZ]. Your serving workmen, Kriele and de Winter [This letter was written by Kriele.]

[The foregoing statement indicates that KdW had a clear idea where the "leg bone and skull" level (the PFZ) was situated within the LB.] [Dubois' October 1892 memorandum (-B5c, below) notes the discovery of the ***Macaca*** sp. molar.]

(ii) Trinil; 5 November 1892 [Saturday] [M…058-616]
…. It is now starting to look quite depressing with respect to the river. From the 1st to the 2nd of November the river level rose so rapidly that our work site [apparently referring to the 25-m Trench] became fully inundated …. and on the morning of the 3rd we started bailing water and continued until a point when we could keep it [the trench] dry.

On the 4th about 5 A.M., the river started rising again to the extent that we had to work hard to keep the barrier dikes [e.g., baulks] above water. In the end our efforts were in vain and then water flowed across them. We even had raised the retaining dams [spoil dikes] so high they were even with the top of the baulks [muurtjes in Dutch].

From the 2nd to the 4th we were unable to carry out any work [that is, excavation] and at this moment our site is completely inundated to the extent that nothing is visible [all is submerged] and for that reason we cannot do anything on the 5th and 6th. It would take at least 3 or 4 days to bail out the water and remove sand that was deposited in the [excavation] site. What should we do now, considering that the work on this side [right bank] has also become inundated. Your serving workmen, Kriele and de Winter

[Note in Dubois' hand-writing on letter >>>>>> gives his instructions to KdW] Answered [on] 7 Nov (so they can be expected to receive it on the 8th): 'Keep on working as long as is possible. Send me news every two to three days. Ship the bones as soon as you have a day when you cannot work and your are able to pack them. If you cannot work for 5 consecutive days, stop all work after first breaching the barrier dams in such a manner that the river cannot deposit sand during the West Monsoon.'

**S II-A2q** M…058-617/618

Trinil; 9 November, 1892 [Wednesday]
…. Our worksite [on the left bank] has been under water from the 3$^{rd}$ to the 7$^{th}$ [of November]. We did resume work on the 8$^{th}$ and think that we need to work at least to the evening of the 9$^{th}$ to bail out the trench so that, if the river remains at this level, we probably can continue our excavations on the 10$^{th}$.

To complete the entire work would likely take 12 to 14 days without interruptions, since we are missing a lot of workers [men that can be put to excavating] because of the need to keep the trench dry [from ground-water seepage]. For instance we need 6 men at night and 6 during the day which amounts to 12 men who do nothing else but keep the pit dry.

In the corners of the [25-m] trench, we are about 20 cm into the target bone layer [PFZ], having found nothing [more of the] Chimpanzee [skeleton]. Also [as previously reported] de Winter found that bone [the femur] at this level. We are watching very carefully and not a single piece of bone that we don't recognize is being thrown away. … Kriele and de Winter [This letter was written by Kriele.]

[This letter confirms that de Winter discovered Femur I, and KdW both thought that the specimen came from the PFZ. Dubois visited Trinil on November 10-11, perhaps prompted by the November 9th letter informing him that the PFZ was again being excavated. During this visit, he should have been able to examine exposures of the upper LB and overlying strata at a location that was immediately adjacent to the portion of the 25-m Trench from which Femur I originated. Dubois should have been able to follow the upper LB from the Ledge near the Skullcap Pit (Figure 3c, main text) westward along the wall of the 25-m Trench past a point near the Femur I discovery location to the western end of that Trench, and seen much of the strata above the LB in nine-meter high back wall.]

**S II-A2r** [M…058-622]

Trinil; 14 November, 1892 [Monday]

Yesterday's order including F25 [25 guilders] … has been received in good order. The river now has subsided already to as low as 25 cm above the solid ledge [the left-bank feature referred to as the Ledge, Figure 4a, main text], so that if we see that the river at this level [just above the LB] for another 10 days, we think we can finish our work [that is, protect the 25-m Trench from inundation and dig out the LB]. We are still finding bones there, and by the way, we have not discovered anything related to the Chimpanzee. … Kriele and de Winter

**S II-A2s** [M…058-618/619; also, -620]

Trinil; 18 November, 1892 [Friday]

…. This night our work site became completely inundated. It has been raining here from around 3 o'clock in the afternoon on the 15$^{th}$ till about 7 o'clock at night and this continued on the 16$^{th}$ as well as on the 17$^{th}$ till 8:30.

For that reason we also could not get any free coolies which we had planned to hire for work during the night and so we had a tremendous amount of work in the morning to get the trench dry. You can thus imagine that we did not do much [productive] work during these three days. Added to this is that this morning the river is still rising.

In the case that we have to leave from here, it would be best to do it now in our opinion since we fear that the river will not subside very quickly. …. We will send the bones … this afternoon, your serving workmen, Kriele and de Winter [This letter was written by Kriele.]

PS After you left us [on November 11] we have progressed about 0.75 meter deeper [in the LB than the PFZ] in the corners of the trench, but there still is no noticeable change in the formation [that is, they were still in the LB, rather than the underlying claystone].

[Apparently based upon what he observed during his November 10-11 visit, his last of 1892, Dubois reported in his October 1892 memorandum (-B5c, below) that only 20% of the LB, which had been exposed in the 25-m Trench, was fully excavated.]

**S II-A2t** [M…058-620; also, M…058-619]

Trinil; 20 November, 1892 [Sunday]

Yesterday we have received the money in good order. The river is still rising. We started packing up this morning. Kriele and de Winter [This postcard was written and signed by Kriele, judging from handwriting.The 1892 letters from November 23 onward are written from Banglee, east of Ngawi.]

[Based upon a label in the box with the specimen, de Vos and Sondaar (1982) and de Vos (1989: Table 1) reports that a *Macaca* $M_3$ dext. (no. 3789) was found in the "the trench of 25 m of 1892, lowest level, ½ m below pe.", meaning that the discovery level was one-half meter stratigraphically below the one from which the *Pithecanthropus erectus* Skullcap and Femur I were discovered. Based upon a label with the specimen, de Vos and Sondaar (1982) and de Vos (1989: Table 1) reports that a *Hystrix lagrelli* molar (termed *Acanthion brachyurus* at the time), Dubois Collection no. 1482a, originated "at the lowest level in the trench of 25 m, 1892," and hence from the LB of the 25-m Trench (although not necessarily in the PFZ).]

### S II-A3. Letters to Dubois concerning the 1893 excavations at Trinil

**S II-A3a** [Kriele wrote the following letters.]

(i) Kedung pingit, April 8, 1893 [Saturday] [M…059-022, right]

De Winter has been to Trinil. We drew straws for it and he pulled the winning one. From what he said it sounds like your house is still in good shape except that the top needs to be replaced. The walls of the prisoner house are fine but the top has blown away .... Our house is standing fine ... The shed and kitchen have been completely blown over.

As far as the work site is concerned, nothing is visible on this side of the river [right bank], but on the other side [left bank] a lot of black material [soil] has been washed away at the top [of the former excavation]. The trench is full of mud but on the outside of the baulk everything [all spoil] has been washed away [the wet-season flooding damaged the 25-m trench site]. According to what the lura tells me, de Winter has made some depth sounding but he couldn't reach it very well without a proa and there was none available.

The small tree that you planted is at least 3 meter high. …. The tools kept by the assistant Wedono are still in good shape…. your serving workmen, Kriele and de Winter

[Other letters came from Pingit on April 15, May 22, and May 28. On May 30 Kriele telegrammed Dubois, "received 20 detainees" [to restart work at Trinil] (M…059-055).]

(ii) [M…059-056 and -057] Trinil; 4 June 1893 [Sunday]

We have been here at Trinil from May 30th onwards but we have not received yet a supplemental group of prisoner workers. You can thus imagine that work on the house is not moving along too quickly. Although the river is still at a high level, the small baulk of the work site on the opposite side [the LB Ledge, Figure 4a, main text, surrounding the 1892 25-m Trench on the left bank] can just about be seen and it does not appear that there is much sediment inside the trench [despite the severity of the wet-season flooding, the 25-m Trench had not been filled with river sediment]. … Your serving workmen, Kriele and de Winter ….

(iii) [M…059-057, right, and M…059-058, left] Trinil, 14 June 1893 [Wednesday]

After you left here, the river has lowered to about 0.15 meter below the large stone [the Ledge]. The trench is about 41 meters in length and the work is progressing very well…. We received the proa yesterday in good shape. The total number of prisoner workers is still the same at 40. … The houses are finished except for the bone sheds on which 4 men are still working with de Winter [on the right bank]. The rest are now all working on the opposite side of the river [left bank] …. Kriele and de Winter

(iv) [M…059-058] Trinil, 18 June 1893 [Sunday]

We received the letter …. We also received the provisions. … In regard to the measurements we made from the marsh [bank top (?)] to the water level [of the river], we measured 5.75 meter from the highest point that you had indicated to me. To measure now the elevation [difference] at the current level is very difficult since the distance is too great from above starting at the wall to the surface of the river. We actually think that the river is still 1 meter above the lowest level [LWL]. … The houses are all finished and all workers are now busy along the river, …. Kriele and de Winter

(v) [M…059-059] Trinil, 23 June 1893 [Friday]

We received our provisions this morning in good order. … we think that in 2 or 3 days we will have removed the [easily dug] top layers up to the solid padas [indurated strata]. The river has risen again in two days to the same height as it was when we measured the 5.75 meter [elevation difference]. Should we keep the free coolies on after we have completed removal of the top layers? …. your serving workmen, Kriele and de Winter

(vi) [M…059-060/061] Trinil, 7 July 1893 [Friday]

Today we received at no cost to us, the following materials from the foreman in Ngawi: Portland cement, lime masonry stone and a tile made of hard rock. The monument pillar [Dubois monument] is now almost completed. The work is progressing very well but I am afraid that we may not have bones by the 15th since we are finding virtually nothing. We are continuing to work with free coolies and we think that we won't finish the upper padas [strata] because it is so terribly hard that it is almost impossible to get through with a pick axe. …. your serving workmen, Kriele and de Winter

(vii) [M…059-023 and -024] Trinil, 13 July 1893 [Thursday; postcard]

I mailed you a letter on the 7th in which I wrote that the monument pillar [Dubois monument] was just about finished as well as the fact that we received the provision orders for de Winter and also about the work activities. Just this morning I have shipped some of the provisions, your serving workman, Kriele

(ix) [M…059-026] Trinil; 14 July, 1893 [Friday]

Yesterday we received our order in good shape. [Since starting in early June] more than 18 m [of the 40-m Trench] have been worked [excavated downward] from the topsoil to about 0.50m above the small baulk [*muurtje* in Dutch; perhaps the Ledge, Figure 4a, main text]. And so there is 22m still to be removed [of the 40-m Trench]. That will take 12 to 14 days, as the lower part is so terribly hard that it is almost impossible to get through. [And the rocks removed had so few fossils in them that by mid July only one crate was filled]. The following sketch shows what the small pillar looks like [the Dubois monument]. Its roof is extended a bit so that rain cannot run along its sides. your serving workmen, Kriele and de Winter

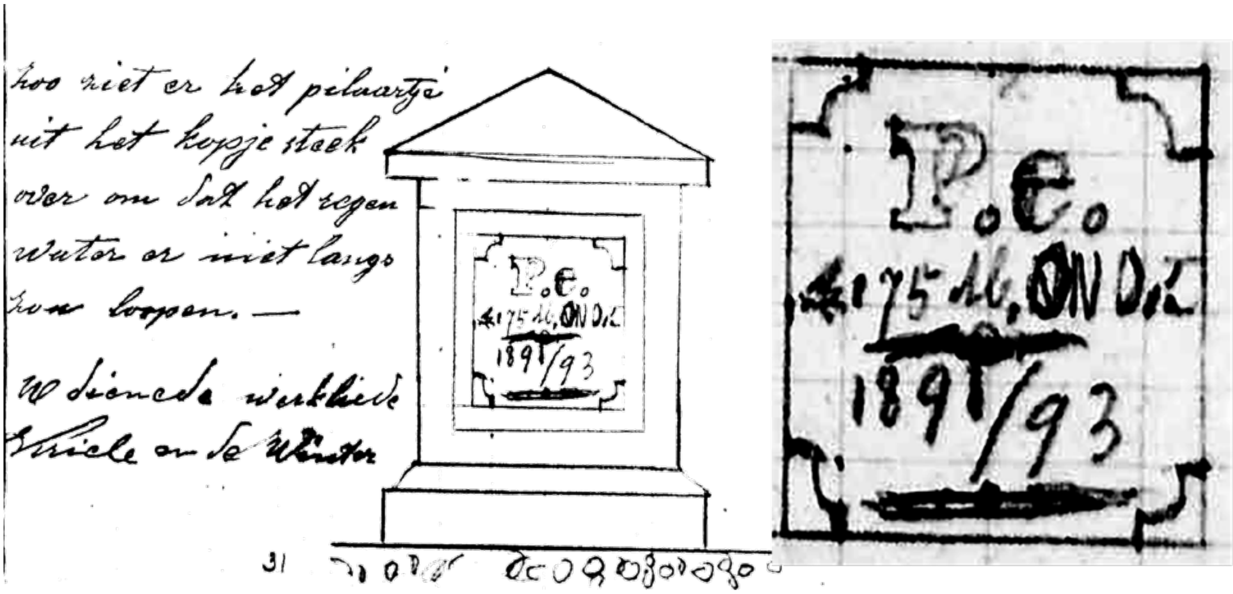


[**"P.e.< 175 M. ONO {ENE} 1891/93."** Kriele's letter of 4 Sept. 1893 (M… 059-030/032) has a final rendering.]

[Dubois' June memorandum described the start of a 40-m long trench south of and adjacent to the 25-m Trench (Figure 3a, main text). The 80% of the 25-m Trench that had been left unexcavated in 1892 was removed also (e.g., -B6b, below). KdW tracked the stratigraphy above the LB in the 40-m Trench and anticipated the depth at which they would encounter the LB/PFZ. In mid-July, they found the "lower part [of the sequence above the LB was] so terribly hard." Dubois visited on July 18-19, before the LB was reached, and fresh excavation exposures would have been visible along back walls 40-m-long, 9-m-high (see Figure 10, main text).]

**S II-A3b**

[M…059-027]
Trinil; 8 August 1893 [Thursday]
Most respected Sir! [I have] a very friendly request, if it is at all possible, to separate me as soon as possible from de Winter. This is because yesterday we had such a severe argument that [I think] it will not remain restricted to that and I fear that others [arguments] will become even more intense. It seems to me then that it is best to warn you ahead of time and before it comes to the point that accidents are going to result from them. Your serving workman, the corporal Kriele

**S II-A3c**

(i) [M…059-028] Trinil; 27 August, 1893 [Sunday] [Kriele wrote his and subsequent letters.]

…. After you departed [on August 19], we found 1 nice elephant [***Stegodon trigonocephalus***] tusk, 1 crocodile skull [***Crocodylus siamensis***], 1 antelope skull [***Duboisia santeng***], 1 turtle [**Testudinoidea**], a few leg bones, deer antlers [***Axis lydekkeri***] and a few ribs and vertebrae. The work is progressing very well, but we have not advanced as deep as the target layer [the PFZ] in the middle of the [40-m] trench [near the Femur I discovery spot]. It will take at least 4 to 5 days before we are at that depth. As we agreed, I will dispatch the bones on the first of next month ….

[Dubois had visited August 17-19 when the LB was being excavated; the fossils reported on August 27 were from the PFZ near the Skullcap and femur findspots (–A3cii). During the rest of 1893, Kriele was responsible for the excavations. De Winter was sent to Sangiran Dome, and wrote to Dubois from Bapang on August 27, September 15, 20 and 24 (M…059-0101; M…059-011; M…059-012; M…059-013); in the last of these, he noted "searching and digging … without finding even one piece of bone and … waiting … for you to arrive" (M…059-012 and 013).]

(ii) [M…059-029] Trinil, 1 September, 1893 [Friday]

…. [I am sending] herewith 14 crates, one of which contains [fossil] wood. The work is progressing very well. We are now finding rather many bones. At this moment we have encountered the target layer [PFZ] in which much of that wood occurs and, by the way, also these shells [**Mollusca**; presumably the species found in the LB during 1891-1982]. We have much hindrance from upwelling ground water [because the excavation was below the river level] and almost always have to designate 3 men to do bailing work so that the trench can remain dry. Other than that, all is going well with the prisoner laborers except that they are suffering much from sore feet, your serving workman, Kriele

**S II-A3d**

[M… 059-030/031] Trinil; 4 September, 1893 [Monday]

…. Yesterday evening at about eleven o'clock the shed of the prisoners has burned down …. Saturday we found an [***Stegodon trigonocephalus***] elephant's head with the molars still in it. The work is going well, but the welling [ground] water is a lot of bother, also a lot among the punished are ill with fever and sore feet…

**S II-A3e**

[M…059-033]Trinil; 12 September 1893 [Tuesday]

…. We are working now at such a depth that on one side of the pit [east end of the 40-m Trench that] a different layer emerges, by the way, because the other end of the pit is already 0.25 m deeper, and we are still in the same sandstone [there]. We will be obliged to keep bailing out water at the one end as that black layer of material [dark-colored claystone] slopes down downstream. The work is going very well and quite a number of bones are being found, although nothing special lately. Kriele

[Evidently, while the top of the LB and PFZ were horizontal (see main text), the top of the underlying claystone dipped slightly; Dubois confirmed the dipping top of the claystone during fieldwork on October 21, 1893 (see S II-C6)]

**S II-A3f**

[M…059-034/035] Trinil; 24 [perhaps 29] September1893 [Sunday]

....We have had a lot of rain last week. Since you have been here [on September 15] the water has risen 1 meter twice and is currently still at that level. I therefore need more people to bail out the water .... As to the finding of bones: still the same amounts as in the previous days but nothing special. [The dikes around the 40-m Trench evidently were >1m above the LWL.] [handwritten side note:] answered 27[th].

### S II-A4. KdW and Kriele letters, maps and summary accounts concerning later excavations

**S II-A4a.**

[M…059-095/096] Tingang [= Tinggang] 16 November, 1895
Most respected Sir !

[M…059-095, right] We received your letter of September 30 here on November 3 [which had been in transit from the Netherlands]. Since you mention in that letter that you would very much like to know the slant dip [of cross beds] at Tawang [~3 km west of Trinil, which Dubois first encountered in1890] and also get a report on how the bones at Trinil were found, [I must tell you] that is nearly impossible for me to answer properly, since I was already in Tingang when I received your letter. The [water level of the] river has now risen so much that the work site is completely inundated.

If I had received the letter perhaps 3 months earlier, I would have been able to immediately attach a label to the bones after excavation at Trinil. Since the bones that were found above and below [M…059-096, left)] are now mixed up and it will be impossible to sort the bones and accurately identify the locations where they were found.

I have also noticed that we previously have made a big mistake in measuring the depth of the excavations below the lowest river level. I measured on the shore side of Ngantjar [Ngancar is located east of Trinil on the left bank] in the 25 meter trench of 1892, in excess of 2 meter depth when the river was at its lowest level. Other than that, all is well with my health.

Otherwise, all is well with my health.

Faithfully yours, Kriele

[Kriele started excavating at Trinil on July 1 and sent reports of his fossil finds to Dubois on July 7, July 21, August 4, August 18, September 2 and September 15, which then evidently prompted Dubois to ask him in his letter of September 30 to sort out the fossils from high and low in the section.]

**S II-A4b.**

[With regard to **1895** work at Trinil, de Vos & Sondaar (1982: 42) state: "…Excavation…. started in July under the direction of Kriele (Kriele 21-7-1895 [letter]). A trench measuring 20x5m was dug. Few fossils {most notably ***Stegodon trigonocephalus*** remains} were found down to a depth of 6 m … (Kriele 21-7-1895).… The left bank was stopped in September. In October again, a small trench was dug on the right bank. As the water level was as low as in the first year of investigation [LWL], there was a dry surface of 6m$^2$ in the rich layer of fossils (Kriele 15-10-1895). Here the skull of an antelope (***Duboisia santeng***), a ***Stegodon*** {***trigonocephalus***} molar and a few bones were found. This work terminated at the end of November (Kriele November 1895)."

De Vos and Aziz (1989: 412-413) add, "The 1895 excavations lasted five months." Probably however, the 1895 Trench was situated on the right bank while the 6m$^2$ outcrop was the left-bank Ledge (Figure 4a, main text), because: None of Kriele's maps of the left-bank show a suitable 20-25x5-m trench, nor is one labeled 1895 on his maps (see Figure 3a, main text, and 1895 pages 11, 21, and 30, below, and maps in -A4e, -A4f and -A4i).

Moreover, Kriele normally used the term "opposite bank" to refer to the left bank, which would be across the river from the 1895 Trench on the right bank. Kriele used "opposite bank" in this way in 1892, 1893, 1896 and 1897 (-A2g; –A4c, especially pages 23, 27, 29 and 33; and –A4d, page 7).

The “big rock at the other side” (page 11, below) and “ledge measuring 6 square meters” (page 30, below) mentioned by Kriele in 1895 was probably the upper LB outcrop, the Ledge, that is seen in the 1894 photograph of the left bank (Figure 4a, main text), an area that was dug away during the 1897 season (Figure 3a, main text; –A4d), worked on during1895 and of interest in 1896.

Dubois presumably knew which sides of the river were being discussed for these reasons and because he had discussed plans for work at Trinil with Kriele in late June before leaving Java (see pages 3 and 4, below).]

[Page] 3 [numbers such as this were written later on the original documents] [M…059-071, right side]
Tinggang, June 10 1895.
Most distinguished Sir!
After you left us, we have not found anything except for what was brought to us: two broken molars of an elephant derived from the upper jaw and a broken molar from the lower jaw. We have not heard a thing about the tools but as soon as they arrive, Kriele will go to Trinil because these kinds of tools are not available here and it takes too much effort to have them made. Should we also excavate [4] [M…059-072, left side**]** above Trinil at the place whose name I now have forgotten? Otherwise all is fine here and Kriele and de Winter wish you and your family a pleasant journey. your humble workmen, Kriele and de Winter.

[Page] 5 [M…059-072, right]
Tinggang, June 11 1895.
Most distinguished Sir! ….
We politely request that you send us a short message about how we should handle shipment of bones that we [hope to] find at Trinil. [6] [M…059-073] Up till now nothing has been found in the Solo valley and the elephant skull will be sent today. Your serving workmen, Kriele and de Winter

[Note: Tinggang is located along the Solo River far downstream of Trinil and north of the Kendeng Hills. An irrigation (canal) project was being conducted near Tinggang using a mechanical digger, and Kriele and de Winter were securing fossils that showed up in the operation (M…059-185)].

**[**Page] 7 [M…059-074]
[Stationary head: “Bevloeiingswerken in de Solo Vallei,” the Solo Valley Irrigation Project]
Tinggang July 7, 1895.
Most distinguished Sir!
The 5 crates with goods have been received in good order. Up till now I have stored everything in the closets at Tinggang since Kriele had already left for Trinil. … The bones that were sent to you before from Ngluwak, were not found in the valley but at the other side of the Solo river and that site has now been entirely washed in by mud and so it will take a lot of effort to find something again. For that reason Kriele decided to go directly to Trinil. … your serving workman no. 19237, de Winter

[Page] 9 [M…059-076] [Kriele started at Trinil on July 1]
Trinil, July 7 1895.
Most distinguished Sir!
Since nothing can be found at Ngluwak, I left there on the 1st of July for Trinil and I have worked here now already for 6 days with 20 free workers. Nothing will come yet of going to Tawang [3 km west of Trinil] since the river level is too high, but as soon as it lowers and it is possible to work there, I will immediately leave for there. During the last period that I still was at Tinggang, nothing was found and I am afraid that [10] [M…059-077, left] it will still be a long time before anything is found. …I request that you, distinguished Sir, let me know as soon as news is received about the extension, preferably before January 12 in view of my service period since it would be very problematic for me to add service time. Also let me know where the bones that I am finding here should be shipped to. Otherwise the excavation here is progressing very well and we found two broken shoulder blades and vertebrae but nothing much of great value. … Kriele

[Page] 11 [M…059-077, right]
Trinil, July 21 1895.
Most distinguished Sir!
During the time that I have been working here, virtually nothing has been found. I am afraid that we have not dug deep enough. The only items that were found are a complete thigh bone, a complete tibia and a tusk, all of an elephant [***Stegodon trigonocephalus***], in addition to a few vertebrae and ribs [apparently a partially articulated skeleton]. I am working on a trench of a length of 20 meters and a width of 5 meters, and we have excavated to about a depth of 6 meters. As far as the water level is concerned, it does not look very encouraging to us, since the big rock at the other side of the river [the upper LB Ledge, Figure 4a, main text] has still not become visible. Therefore [12] [M…059-078, left] to travel now to Tawang would be an effort in vain and I will therefore wait long enough till I am sure that the ledge at Tawang has been exposed. The tree that you, distinguished Sir, planted here at Trinil is looking very much alive and it is at least 7 meter tall. The same good condition can be reported for the small pillar and the small stone [referring to the Dubois monument installed in 1893]. Otherwise all is going well with my health …. Your humble worker, Kriele

[Excavation of 20x5m on right bank found “elephant” remains at 6m in depth or less; “the big rock at the other side of the river” was still submerged (Figure 3c, main text, Ledge).]

[Page] 13 [M..059-078, right]

[{Stationary heading} Bevloeiingswerken in de Solo valley {Irrigation works in the Solo valley}]
Tinggang July 31, 1895.
Most distinguished Sir!
The work activities here [near Tinggang, not Trinil] are up till now not progressing very well and not very much is being found because little digging is taking place at the spots where bones might be found. … The river has not seen any lower level. After you left [on June 27], we have not had even 8 days of dry weather. I don’t know how Kriele is doing at Trinil but I don’t believe that he will be able to continue working there this year. Mr Duijvetter has told me that only F 850 had been allotted for us for the year and Kriele spends F 150 per month on coolies alone. Would that mean then that there is a danger that we may not stay next year? I am a bit worried about that because I would be very pleased to continue working for you next year. There is talk that the Solo Valley [project] will be amended if there is a shortage, but would that really be true if it becomes that much? I am only asking how you judge this to be and if you don’t see a danger then we should also not worry. Otherwise all is going very well with me and my health. In my next letter I hope to be able to send better news about the work activities. Did you and your family arrive safely in Europe? Your serving workman, de Winter

[Page] 14 [M…059-079]
Trinil August 4 1895.
Most distinguished Sir!
In the last two weeks we have found [in the 1895 Trench at Trinil]: 1 elephant [***Stegodon trigonocephalus***] tusk with thigh bone and tibia [which originated above 6 m in depth, (noted above)]; 1 occipital portion of a banteng [***Bibos palaesondaicus***] skull with one complete and one incomplete horn; 1 half lower jaw of a small crocodile [***Crocodylus siamensis***] and further several [***Axis lydekkeri***] deer antlers, vertebrae leg bones of smaller species etc. etc. In recent days we have had much hindrance from high water to the extent that we were not able to work. I am now working with 20 coolies and I am afraid that if we get again such poor weather once or twice, we will not be able to [15] [M…059-080, left] finish the trench. I am not inclined to add more workers to our force because I would then be spending a lot of money and de Winter has written me that he heard that we may not spend more than exactly that sum of F 1,500 from which our bonuses must be deducted. And so with that sum of money we cannot possibly get to December since I spend about F 150 per month on the work force alone. At the present time the water is low again and the

skies are nice and clear and according to the Javanese it will now stay dry for some time. If I should have to pack the bones here so that they can be transported directly to Europe, I will need to hire a carpenter to manufacture the packing crates during the time [16] [M…059-080, right] that I remain at the work site and so the sooner I know where they should be sent, the better. I mean the bones from Trinil. Otherwise, my health is very good. Your humble worker, Kriele

[Large fossils were reported from the 1895 Trench.]

[Page] 17 [M…059-081]
Trinil August 18 1895.
Most distinguished Sir!
During the last two weeks we have found [in the 1895 Trench of 20-m length at Trinil]: 5 complete skulls of which one still has all dentition but has no horns and others with complete horns [perhaps referring to deer antlers]. In addition many short horns were found of which some are very nice. We are not finding as many bones of small species as we did in the past, for instance small jaws, teeth or molars of deer [***Axis lydekkeri***] or other kinds of animals. Also very few teeth and molars of large animals are being found and so far no jaws at all. Added to this is still the find of an elephant [***Stegodon trigoncephalus***] ulna and some [18] [M..059-082] forelimb bones of animals that I don't recognize and in addition a quantity of different sort of bones for instance vertebrae, ribs etc. etc. If we keep having this nice weather and the river water continues to subside as it is doing now, I believe that I will be able to go to Tawang in mid September and start excavating there. … Your humble workman, Kriele

[Five complete skulls, but not many small fossils, found in 1895 Trench, presumably all from the LB/HK]

[Page] 19 [M…059-083]
Tinggang August 21 1895.
Most distinguished Sir! ….
We did not find a thing this month in the valley…. From what Kriele is saying, he must have gathered now already a nice collection since this month no rain has fallen yet and the river has subsided nicely. Would you allow Kriele to send me some bones? In that way I can still do some work if we are not digging for bones in the valley and I am not very occupied and so prepare the Trinil bones. I could for instance do the gluing and careful packing etc. I have [20] [M…059-084] asked Kriele about this already but he told me that your orders are to keep the bones in Trinil until further notice…. Your serving workman, de Winter

[Page] 21 [M…059-085]
Trinil September 2 1895.
Most distinguished Sir!
From August 14 to the end of the month, we have not found many bones [in the 1895 Trench at Trinil]. The reason for this is that in the sector [*vak*] of 25 meters in length, I finished the first 17 meters of it and that was completed on August 20 [the 1895 Trench had been lengthened to 25 m; 17 m of it had been fully excavated]. During that time [when the 17 m was dug through the LB-HK] we did find some [***Axis lydekkeri***] deer antlers, a thigh bone of an elephant [***Stegodon trigonocephalus***] as well as some vertebrae, phalanges, ribs etc. etc. We did not find anything very special [usually meaning no potentially hominin material]. As soon as I finish the remaining piece [the last 8 m of the 25-m trench] I will leave for Tawang. My prediction is that I will complete it [the 1895 Trench] on the 20$^{th}$ of this month. If nothing is recovered at Tawang, we could work a bit on the opposite side of [22] [M…059-086] the river [at Trinil] until December so that we then will also have time to pack the bones. Would you agree that I can send the bones from Trinil to Tinggang? De Winter would then be able to get going on manufacturing the crates, and here in Ngawi I am not able to obtain very good packing crates. …. Kriele

[1895 Trench lengthened to 25m with (17m completed); work planned on "the opposite side," the left bank.]

[Page] 30 [M…059-093]
Trinil 15 October 1895.
Most distinguished Sir !
Since almost nothing was being found at Tawang, I returned to Trinil on the 12th. At Tawang I did find an incomplete lower jaw of an elephant with nice molars still attached. In addition two loose elephant molars and two loose molars of a rhinoceros derived from its mandible, as well as several others pieces of bone. On the 14th I started [excavating] again at Trinil on the opposite side of the river [left bank], since the water level of the river is now as low as it was in the first years when we started digging and the dike has just about been washed away and an additional piece of the ledge [the LB Ledge, Figure 4a, main text] has become exposed measuring about 6 square meters. I do [31] [M…059-094, left] think that I will still be able to finish this before the river rises again. The rock formations at Tawang are exactly the same as those at Trinil [Duyfjes, 1933, later mapped the beds at two localities as the Kabuh Formation (-*G6a*), validating Kriele's remark]. They contain also much petrified wood which is as hard as can be [at Tawang], [whereas] at Trinil this wood is soft. The clay layer occurs at a depth of about one meter. And, by the way, I also tried two other spots but nothing much resulted from that either. Other than that, all is well with my health. …
Kriele

[Fossils found in the left-bank Ledge: "On the 14th [of October] I started … on the opposite side [left bank] … since the water level … is now as low as it was in the first years [LWL] …. and a … ledge [the Ledge] … has become exposed measuring about 6m square meters." Excavation on the "on the other side … is inundated," after excavating ***Duboisia santeng*** and ***Stegodon trigonocephalus*** fossils from the LB-HK Ledge (Figure 4a, main text).]

[Page] 32 [M…059-094, right].
Tinggang, [early?] November 1895
Most distinguished Sir!
During the last days that we were able to work at Trinil, we were still able to find the following: an incomplete antelope skull [***Duboisia santeng***] together with one horn, an elephant [***Stegodon trigonocephalus***] molar as well as several other pieces of bone, but incidentally nothing special was amongst this [usually meaning no potentially hominin material]. Since the water is now rising so rapidly, to the extent that the work site on the other side of the river [left bank at the Ledge] is inundated, I have been forced to stop the work here, which I did on the 21st of October. [33] [M…059-095, left]
I then started packing the bones immediately, so that I could take them with me in a proa to Tinggang. Once there, I began making crates and packed them inside those, to be prepared to send them to the address which you promised to relay to us.
Otherwise all is well with my health. … Kriele.

[Page] 34 [M…059-095, right] Tinggang 16 November 1895. [See -A4a, above, for this letter, which concerns a mistake in measurement within the 1892 excavations.]

[Page] 36 [M…059-096, right)] Tinggang 16 November 1895. Most distinguished Sir! Since the work near the bone discovery spots in the Solo valley had been halted, I have started manufacturing crates for shipment to Europe after [learning of] Kriele's writing to you. … We have just started and made good progress.

Page] 37 [M…059-097] with packing of the bones and I hope to send these bones to you before the New Year. …. Your serving workman, de Winter

[Page] 38 [M…059-098] Tinggang, November 30, 1895. Most distinguished Sir! The bones destined for Europe have been packed for several days now …. No work is being done … in these surroundings …. Kriele and de Winter

[Page] 40 [M…059-099] Tinggang, December 15, 1895. We are still waiting on word where to send the crates with destination for Europe …. We have searched here … but found nothing …. Kriele and de Winter

[Page] 1 [M…059-105] [Written by hand in margin *"1896"]*
Tinggang January 1, 1896.
Most distinguished Sir!
Enclosed herewith are the specifics and contents of the crates [containing the 1895 finds]. The most attractive bones were packed in the first 7 crates with a label stuck to each specimen. Actually it is impossible to indicate the exact size [of the specimens ?]. These 7 crates contain the small bones, which you requested to be packed and sent to Europe immediately. However, we received news on the 25th [of December 1895] that all bones should immediately be packed which we then did and they had to be sent to Bodjonegoro and from there onwards. In the last 9 crates, no labels were stuck to the Trinil bones, only to those that were found at [2] [M…059-106, left] other locations. We wouldn't have had the time to stick labels on everything knowing that for example we might have been transferred all of a sudden at the end of December since we still had heard nothing about an extension**.** Also enclosed are the dip at the black formation [in the irrigation canal] which is now in accordance with what they reported to me at Picket Nr. 135. This black formation is currently found at a depth of about 9 meter. Other than that, the work is progressing very poorly. But our health is very good, your serving workmen, Kriele and de Winter

[Labels put on fossils: "Labels [were] stuck to each specimen" but not on any of them from Trinil.]

[Additional records on the fossils shipped: The fossils described in the foregoing letters were packed in December 1895 and shipped in early 1896; the following list mentions many fossils from Trinil (noted by our underlining), but gives no stratigraphic information for them; most must have come from the LB-HK in the 1895 Right-bank Trench (we have omitted our inferred species names in the following). This list, written by Oosthout, differs slightly from one that Kriele gave in M…040-357, which follows.]

[Page] 167 [M…040-356] [Stationary head:] "Bevloeiingswerken in de Solo Vallei" [Irrigation works in the Solo Valley] 20614: Packing list of fossils, destined for the Geological Mineralogical Museum in Leiden. Sent per prao from Boojonegoro to Surabaya.
Columns: Sequence Number; Kind of packing; Content; Dimensions (length/width/height); Volume (Cubic M); Weight (in KG)

1. Crate; Content: 33 complete deer antlers from Trinil. [Length] 0.7 M [Width] 0.40 M [Height] 0.42 M [Volume] 0.118 Cu M [Weight] 37 KG [*specimens had provenience labels*]

2. Crate, Content 22 deer antlers from Trinil and 9 deer skulls from Trinil [Length] 0.65 M [Width] 0.46 M [Height] 0.41 M [Volume] 0.123 Cu M [Weight] 37 KG. [*specimens had provenience labels*]

3. Crate [Content] 1small chest with 39 assorted fossil leg bones and small jaws etc., 1 small chest with an antelope skull, 2 small bone legs and 17 assorted small teeth from Trinil; 1 small chest with 3 crocodile teeth, 4 tips of elephant tusks and 9 molars of ruminants from Trinil; and 1 small chest with 2 crocodile jaws and 2 rhinoceros molars (from Trinil); 1 small chest with 17 assorted small bones, small jaws etc (Trinil); 1 small chest with 11 assorted bones (from Trinil), 3 deer skulls and 1 banteng skull with horns Trinil). [Length] 0.99 M [Width] 0.43 M [Height] 0.38 M [Volume] 0.162 Cu M [Weight] 55 KG [*specimens had provenience labels*]

4. Crate [Content] Bones from Tinggang; 2 elephant tibia; 2 flat and 1 small elephant molars, 1 upper jaw with 2 molars; 1 half lower jaw with 1 molar (all from Tinggang); 1 package with ruminant molars; 1 half lower jaw of a rhinoceros without molars; 1 small package of sea shells from Ngambon and 1 Banteng horn from Ngluwak. [Length] 0.71 M [Width] 0.58 M [Height] 0.46 M [Volume] 0.189 Cu

M [Weight] 90 KG Carry over Total: Volume 0.592 Cu M; Weight 219 KG [*specimens had provenience labels*]

[168] [M…040-357, left] 5. Crate [Content] 1 upper jaw (elephant) from Trinil. [Length] 0.63 M [Width] 0.63 M Height 0.48 M [Volume] 0.191 Cu M [Weight] 87 KG [*specimens had provenience labels*]

6. Crate [Content] 24 leg bones (in part incomplete); 1 buffalo jaw; 3 elephant molars and 2 ribs; 1 piece of a tusk; 2 broken turtle pieces; 1 half of a rhinoceros jaw; 1 banteng horn; 1 broken buffalo horn and 3 phalanges (all from Trinil) [Oosthout omits 3 vertebrae, see Kriele's list, below] [Length] 0.99 M [Width] 0.33 M [Height] 0.41 M [Volume] 0.134 Cu M [Weight] 64 KG [*specimens had provenience labels*]

7. Crate [Content] 1 lower jaw (elephant); 14 phalanges; 1 broken piece of a turtle and 2 rhinoceros thigh bones [Kriele includes leg bones, see his list below]. [Length] 0.67 M [Width] 0.41 M [Height] 0.42 M [Volume] 0.115 Cu M [Weight] 59 KG [*specimens had provenience labels*]

8. Crate [Content] 130 incomplete deer antlers, from Trinil. [Length] 0.88 M [Width] 0.47 M [Height] 0.51 M [Volume] 0.211 Cu M [Weight] 82 KG [*specimens in Crates 8-16 had no provenience labels*]

9. Crate [Content] 1 elephant tusk 2 M long in 3 pieces (from Trinil); 1 elephant tusk 1.10 M long in 2 pieces (from Tinggang); 26 deer antlers and 1 ulna (Trinil). [Length] 0.95 M [Width] 0.55 M [Height] 0.41 M [Volume] 0.214 Cu M [Weight] 96 KG

10. Crate [Content] 1 tibia (from Trinil); 2 elephant molars (from Tinggang) and 15 broken bones of smaller animals. [Length] 0.94 M [Width] 0.38 M [Height] 0.30 M [Volume] 0.139 Cu M [Weight] 71 KG

11. Crate [Content] 1 elephant thigh bone (Trinil); 1 lower elephant jaw (from Tinggang) and 11 assorted bones from Trinil. [Length] 0.62 M [Width] 0.56 M [Height] 0.37 M [Volume] 0.128 Cu M [Weight] 66 KG

12. Crate [Content] 1 sacrum bone (Trinil); 1 elephant thigh bone (Tinggang) and 1 elephant thigh bone from Trinil [Kriele's list does not specify that they are elephant specimens]; 9 assorted bones (from Trinil). [Length] 1.02 M [Width] 0.52 M [Height] 0.53 M [Volume] 0.281 Cu M [Weight] 156 KG

13. Crate [Content] 12 assorted bones from Trinil. [Length] 0.96 M [Width] 0.51 M [Height] 0.43 M [Volume] 0.211 Cu M [Weight] 83 KG

14. Crate [Content] 20 assorted ribs from Trinil. [Length] 0.69 M [Width] 0.35 M [Height] 0.37 M [Volume] 0.089 Cu M [Weight] 32 KG

15. Crate [Content] 19 assorted bones from Trinil. [Length] 0.59 M [Width] 0.49 M [Height] 0.47 M [Volume] 0.136 Cu M [Weight] 58 KG

16. Crate [Content] 29 assorted bones from Trinil. [Length] 0.87 M [Width] 0.50 M [Height] 0.41 M [Volume] 0.178 Cu M [Weight] 73 KG

. Total [Volume] 2.619 Cu M [Weight] 1,146 KG

PS: All crates are marked: Government Goods, Ministry of Colonial Affairs, 's Gravenhage, Destined for the Rijks Geologisch-Mineralogisch Museum, Leiden

Bodjonegoro, January 3, 1896; The Chief Warehouse Master;
per The First Clerk of the Warehouse Service
[signed] L. Oosthout

[169] [M…040-357, right; a stratigraphic sketch with explanatory notes concerning outcrops (probably) near Tinggang in the Solo valley has been omitted]

[The following appears to be the contents of the 16 crates listed by Oosthout, made as they were unpacked in Leiden. Trinil fossils are noted by our underlining, but our inferred species names are omitted in the following. The identity of the person who wrote this second list is not known.]

Crate No. 1: 33 complete deer antlers (Trinil). [*Oosthout lists '33 complete deer antlers from Trinil' in Crate 1.*]

Crate No. 2: 22 deer antlers (Trinil) and 9 deer skulls from Trinil. [*Oosthout lists '22 deer antlers from Trinil and 9 deer skulls from Trinil' in Crate 2.]*

Crate No. 3: 1 small chest with 39 assorted fossil leg bones, small jaws etc., 1 small chest with an antelope skull, 2 small bone legs and 17 assorted small teeth (Trinil); 1 small chest with 3 crocodile teeth, 4 tips of elephant tusks and 9 molars of ruminants (Trinil); 1 small chest with 2 crocodile jaws and 2 rhinoceros molars (from Trinil); 1 small chest with an assortment of 17 small bones, small jaws etc (Trinil); 1 small chest with an assortment of 11 small bones (from Trinil), 3 deer skulls and 1 banteng skull with horns (Trinil).

Crate No. 4: Bones from Tinggang; 2 elephant tibia; 2 flat and 1 small elephant molars, 1 upper elephant jaw with 2 molars; 1 half lower elephant jaw with 1 molar; 1 small package with ruminant molars; 1 half lower jaw of a rhinoceros without molars; 1 small package of sea shells from Ngambon and 1 banteng horn from Ngluwak.

[170] [M…040-358] [there are differences with Oosthout for crates 6 and 7] Crate Nr. 5: 1 upper elephant jaw (Trinil).

Crate No. 6: 24 complete and incomplete leg bones; 1 buffalo jaw; 3 vertebrae [*missing on Oosthout's list*] ; 3 elephant molars, 2 ribs; 1 piece of a tusk; 2 pieces of turtle; 1 half of a rhinoceros jaw [from Trinil]; 1 banteng horn; 1 broken buffalo horn and 3 phalanges (Trinil).

Crate No. 7: 1 lower jaw (elephant); 14 phalangess; 1 broken piece of a turtle and 2 rhinoceros leg bones. [*Oosthout wrote "thigh bones"*]

Crate No. 8: 130 incomplete deer antlers (Trinil).

Crate No. 9: 1 elephant tusk 2 meter long, in 3 pieces (Trinil); 1 elephant tusk 1.10 meter long in 2 pieces (Tinggang); 26 deer antlers and 1 ulna (Trinil).

Crate No. 10: 1 tibia (from Trinil); 2 elephant molars (from Tinggang) and 15 broken bones of smaller animals (Trinil).

Crate No. 11: 1 elephant thigh bone (Trinil); 1 lower jaw of elephant (Tinggang) and 11 assorted bones from Trinil.

Crate No. 12: 1 sacrum bone (Trinil); 1 thigh bone (Tinggang) and 1 thigh bone (Trinil) 9 assorted bones (from Trinil).

Crate No. 13: 12 assorted bones (Trinil).

Crate No. 14: 20 assorted ribs (Trinil).
[169] [M…040-357, right; a stratigraphic sketch with explanatory notes concerning outcrops (probably) near Tinggang in the Solo valley has been omitted]

[The following appears to be the CONTENTS OF THE 16 CRATES listed by Oosthout, made as they were unpacked in Leiden; the identity of the person who wrote this second list is not known [our inferred species names are omitted]

Crate No. 1: 33 complete deer antlers (Trinil). [*Oosthout lists '33 complete deer antlers from Trinil' in Crate 1.*]

Crate No. 2: 22 deer antlers (Trinil) and 9 deer skulls from Trinil. [*Oosthout lists '22 deer antlers from Trinil and 9 deer skulls from Trinil' in Crate 2.]*

Crate No. 3: 1 small chest with 39 assorted fossil leg bones, small jaws etc., 1 small chest with an antelope skull, 2 small bone legs and 17 assorted small teeth (Trinil); 1 small chest with 3 crocodile teeth, 4 tips of elephant tusks and 9 molars of ruminants (Trinil); 1 small chest with 2 crocodile jaws and 2 rhinoceros molars (from Trinil); 1 small chest with an assortment of 17 small bones, small jaws etc (Trinil); 1 small chest with an assortment of 11 small bones (from Trinil), 3 deer skulls and 1 banteng skull with horns (Trinil).

Crate No. 4: Bones from Tinggang; 2 elephant tibia; 2 flat and 1 small elephant molars, 1 upper elephant jaw with 2 molars; 1 half lower elephant jaw with 1 molar; 1 small package with ruminant molars; 1 half lower jaw of a rhinoceros without molars; 1 small package of sea shells from Ngambon and 1 banteng horn from Ngluwak.

[170] [M…040-358] [there are differences with Oosthout for crates 6 and 7] Crate Nr. 5: 1 upper elephant jaw (Trinil).

Crate No. 6: 24 complete and incomplete leg bones; 1 buffalo jaw; 3 vertebrae [*missing on Oosthout's list*] ; 3 elephant molars, 2 ribs; 1 piece of a tusk; 2 pieces of turtle; 1 half of a rhinoceros jaw [from Trinil]; 1 banteng horn; 1 broken buffalo horn and 3 phalanges (Trinil).

Crate No. 7: 1 lower jaw (elephant); 14 phalangess; 1 broken piece of a turtle and 2 rhinoceros leg bones. [*Oosthout wrote "thigh bones"*]

Crate No. 8: 130 incomplete deer antlers (Trinil).

Crate No. 9: 1 elephant tusk 2 meter long, in 3 pieces (Trinil); 1 elephant tusk 1.10 meter long in 2 pieces (Tinggang); 26 deer antlers and 1 ulna (Trinil).

Crate No. 10: 1 tibia (from Trinil); 2 elephant molars (from Tinggang) and 15 broken bones of smaller animals (Trinil).

Crate No. 11: 1 elephant thigh bone (Trinil); 1 lower jaw of elephant (Tinggang) and 11 assorted bones from Trinil.

Crate No. 12: 1 sacrum bone (Trinil); 1 thigh bone (Tinggang) and 1 thigh bone (Trinil) 9 assorted bones (from Trinil).

Crate No. 13: 12 assorted bones (Trinil).

Crate No. 14: 20 assorted ribs (Trinil).

Crate No. 15: 19 assorted bones (Trinil).

Crate No. 16: 29 assorted bones (Trinil).

[177] [M…040-365] [same handwriting as in …364] (5) large and small fragments of the shell of Hardella (?)

[SUMMARY: 457 fossils were attributed to Trinil in 1895. None had labels. With the exception of the ***Stegodon trigonocephalus*** molar and ***Duboisia santeng*** skull from the left-bank Ledge, which was adjacent to the Skullcap Pit (Figure 4a, main text), the finds most likely came from the 1895 Right-bank Trench, and overwhelmingly these appear to have been from a **LB-HK** level, judging from Kriele's letters and the composition of the 1895 collection.

211 (46%) of the "Trinil" specimens were *Axis* antlers, 184 (40%) assorted bones, 31 (7%) teeth), accounting for 93% of the total. The dental specimens included two *Rhinoceros sondaicus* molars and three *Crocodylus siamensis* teeth. 31 other fossils included

- "an antelope [*Duboisia santeng*] skull," four tips of [*Stegodon trigonocephalus*] tusks,
- two crocodile [*Crocodylus siamensis*] mandibles, 12 *Axis* [*lydekkeri*] craniums,
- a *Bibos palaesondaicus* skull with horn cores preserved,
- a *Stegodon trigonocephalus* tusk that was 2 m long, several tibia,
- a *Stegodon trigonocephalus* femur, and a sacrum.

Kriele also reported that on July 21 a complete femur and tibia of an "elephant" (*Stegodon trigonocephalus*) occurred with a tusk and a few vertebrae and ribs at a depth of 6m or less, well above the **LB-HK**.].

## S II-A4c.

[With regard to **1896**, de Vos and Sondaar (1982: 42) state: "The excavations started in June and a trench measuring 10x5 m was dug on the left bank … (Kriele 30-6-1896), and at the end of the month the trench was lengthened to 14m (Kriele 7-9-1896). In August work was again being done on the right bank of the river (Kriele 15-8-1896). Here a trench of 100m was dug (Kriele 12-10-1896) to a depth of 11.50m, which was 1m below the lowest water-level. In October the work was stopped, due to lack of money (Kriele November 1896)." De Vos and Aziz (1989: 413) provided a copy of Kriele's map of the 1896 excavations. Reformatted versions of the maps are provided below with annotations added.]

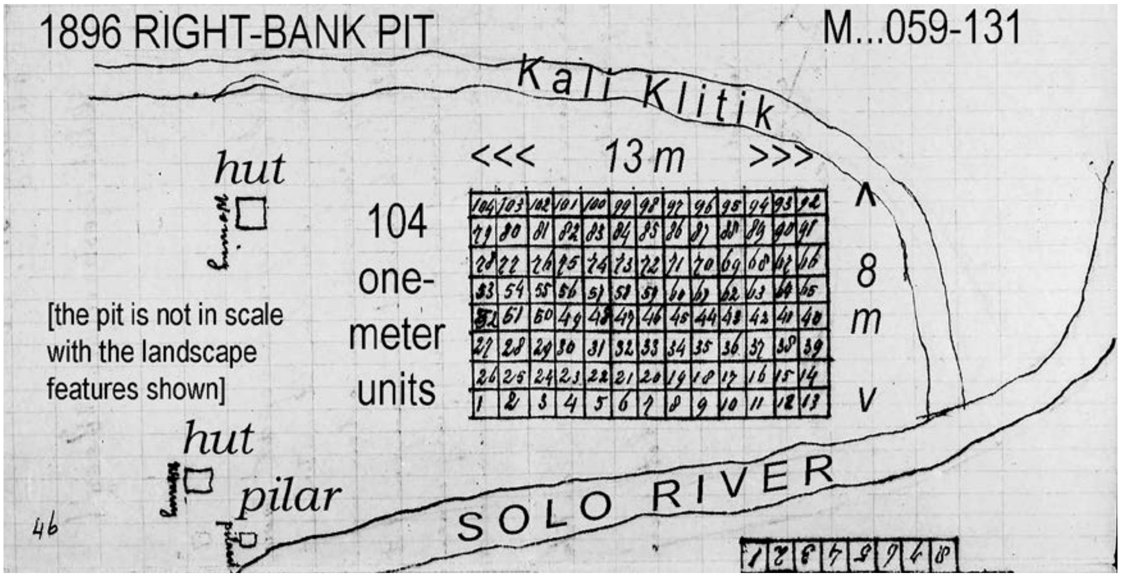

[The 1896 Right-bank Pit, as Kriele sketch (above) was dug first (see letters below). Where the 1895 Trench was relative to the 1896 Right-bank Pit is unknown. The Selenka Expedition Pit I was located adjacent to one of these Dubois' excavations (-G1d). The 1896 Left-bank Pit is shown below in Kriele's sketch.]

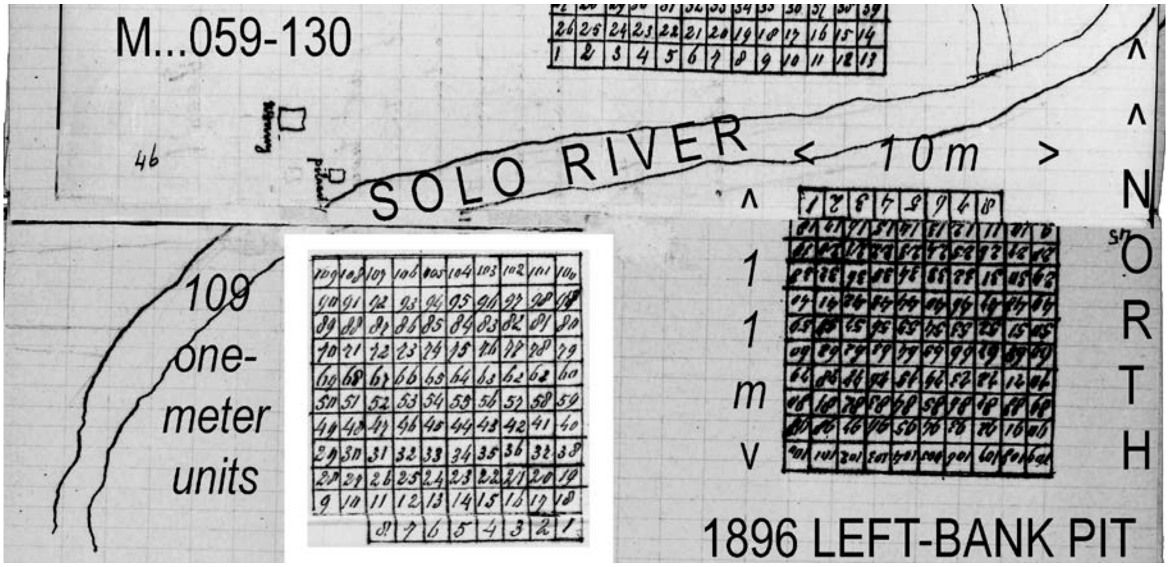


[The 1896 Left-bank Pit was located at the western end of the 1891-1893 excavations (-A4f and –A4o, below). As the following Kriele letters reveal, the Left-bank Pit was dug downward through the high embankment, which is seen in the 1894 photograph to have been made up of the units **2** to **5** (Figures 3c and 4a, main text). Kriele had to dig 11.50m below surface and 1.0m below lowest river level to fully penetrate the **LB-HK** (page 25, below), repeating the experience that KdW had had in 1891-1893 and recovery of large fossils from the 6m$^2$ Ledge in 1895.]

[Kriele's letters about 1896 excavations are below.]

[Page] 12 [M…059-111, right]
Tinggang, April 21 1896
Most distinguished Sir!
For some days now I have been a private citizen [out of the Army], and I have been assigned to the Solo valley project at a salary of F 1.50 per day plus F 40 bonus, which is paid from the funds by Paleontology. From this, distinguished Sir, you can conclude that I will remain available for your research work, but keep in mind that I also have to serve the Solo valley project. … I don't know if I am allowed to go to Trinil, but I did tell them that if it is allowed I could easily start there on the 1st of May. In regard to my salary, the amount is still the same as before, except that I am now receiving my pension in addition to it. I don't think I will write now to Mr. Delprat about when the research for you will be done. Perhaps you, distinguished Sir, will you be so good as to write a letter to Mr. Delprat about a job. [14] [M… [059-112, right] Other than that, all is well with me and my wife, as well as with de Winter your humble workman, Kriele.

[Page] 17 [M…059-114, right]
Tinggang, May 15 1896
Most distinguished Sir!
Since we have to report to you once again that nothing has been found we may note that from Picket 140 through 136, fewer than 100 men are at work. So you will understand how much ground is being displaced and in addition they are digging mostly in the black formations, Things are looking bleak in terms of progress of the work. You asked me recently if I would be allowed to go to Trinil, so our local chief intended to request that, and promised us that we would be warned immediately after it was approved that one of us would [18] [M…059-115, left] be able to go there. Other than that, all is going well with our health, your serving workmen, Kriele and de Winter.
[Kriele is permitted to work at Trinil again.]

Page] 19 [M…059-115, right]

Trinil, June 15 1896.
Most distinguished Sir!
I left Tinggang on the 8th for Trinil and started work with 15 coolies since I was not allowed to employ more than those. I have undertaken excavation of a sector of [only] 10 meters in length and 5 meters [on the right bank; see below] in width since I am afraid, according to what I hear, that we may not be allowed to work more than 2 months and thus I did not dare to undertake a larger sector. Yesterday, I found [20] [M…059-116, left] in the red padas [probably not at Trinil; Padas is a locality east of Trinil along the Solo River], 6 vertebrae grouped together. In regard to the documents for my marriage, I don't have to get them sent from Europe. … Beyond that I hope to be able to report better news to you at the end of the month about this excavation sector. Other than that, all is going very well with my health, your serving workman, Kriele
[Kriele started the 1896 Right-bank Pit at Trinil on June 8.]

[page 21] [M…059-116, right]

Trinil June 30 1896.
Most distinguished Sir!
I have enlarged the sector of 10 meters to 14 meters [on the right bank; on his later plat, above, it is 8x13 m], since I realized that I could finish that even in the case that we would be allowed to work no more than 2 months. This sector is located [east of the Dubois monument and] immediately next to the gully through which water runs from a spring [Kali Klitih on -A4q]. During the 20 days that we have been at work here, we have found the following [presumably from the LB]: 1 complete turtle [**Testudine**], 3 elephant [***Stegodon trigonocephalus***] molars, 1 complete lower jaw of a pig [***Sus brachygnathus***] as well as several molars. Of the ruminants we found leg bones, phalanges, [page 22] [M…059-117, left] vertebrae etc. Other than that, all is going very well … Kriele

[page 23] [M…059-117, right]

Trinil, July 16 1896.
Most distinguished Sir!
During this month, we found [in the 1896 Right-bank Pit, and probably in the LB]: 1 elephant [***Stegodon trigonocephalus***] tusk, 1 turtle, several [***Axis lydekkeri***] deer antlers, 1 [***Duboisia santeng***] antelope skull with complete horns, 2 rhinoceros [***Rhinoceros sondaicus***] molars in addition to several complete and incomplete leg bones, ruminant jaws and 2 [***Stegodon trigonocephalus***] elephant molars as well as some vertebrae, phalanges etc. This sector of 14 meters will be finished this month. I don't know yet whether or not we will be allowed to continue working. [24] [M..,059-118, left]. It is starting to become a problem to dispose of our water and the spoils. In order to be able to continue working it would be best to start again at the opposite side of the river [left bank], since the loose spoil piles [adjacent to the 1891-1893 excavations] have now been washed away. About measuring the thin slanted beds [cross beds] locally, this is very problematic in those places where I am now not excavating since all these slanted beds are running right through each other [dipping in varying directions]. Other than that, all is very fine with my health, your serving workman, Kriele

[page 25] [MM774C-000059-118, right]

Trinil July 31 1896.
Most distinguished Sir!
During the last half a month we have found the following [in the 1896 Right-bank Pit]: 1 exceptionally beautiful [***Axis lydekkeri***] deer skull with complete antlers, 1 incisor of a small species and in addition several incomplete leg bones, vertebrae, deer antlers, lower and upper jaws of ruminants etc. etc. but nothing very special. For several days now we have been working on a new sector, which is 10 meter long and 4 meter wide. I have not received any orders that [26] [M…059-119, left] I have to stop working here at Trinil by the middle of August. The river is still too high to finish the sector on the other side [left bank; and referring to the excavation of the Ledge (Figure 4a, main text), where work began in 1895]. Other than that, all is going very well with my health, your serving workman, Kriele

[The 1896 Left-bank Pit was started “on the opposite side” of the river from the 1896 Right-bank Pit (sketches, above), while the LB-HK Ledge, which had been worked “last year,” was still submerged.]

[Page] 29 [M…059-120, right] Trinil, September 7, 1896. Most distinguished Sir! Please be not mad at me for receiving my letter of August 15 with [only] a postage stamp of 10 cents. …. I am now digging already 13 days on the opposite side of the river [left bank] in a [page 31] [M…059-121, left] sector of 10x10 meters, which [is being dug from the top of the embankment downward, and] will be enlarged below [nearer to the objective bonebed]. I did not dare to make the sector larger since I am afraid that I may not finish that. As you know yourself, we have to dig more than 2 meter below the low water level [to fully penetrate the LB-HK], and that gave us much trouble in the past with the penetration of ground water. Since no more than 15 coolies may now be employed, the work downward will progress very slowly. During the last half month we have found nothing of value because we are still working too shallow [to have reached fossiliferous beds in the left-bank 10x10 m trench]. Your letter of July 27 arrived [31] [M…059-121, right] here yesterday... My plan is now to remain forever in the Indies…. If next year there would [32] [M…059-122, left] not be an extension granted [to work at Trinil], I would very much like to be employed with the Railways ….You would be assisting me a lot for that with for instance Mr. Delprat ... But if the work for you is to be extended, I will continue to work for you. Other than that, all is well with my health, your serving workman, Kriele

[Excavation was still “too shallow” in the 1896 Left-bank Pit to find fossils, and Kriele exhibited confidence in his stratigraphic knowledge of the left bank in stating that his crew would “have to dig more than 2 meters below the low water” to fully penetrate the LB-HK.]

[page 33] [M…059-122, right] Trinil, September 16 1896. Most distinguished Sir! This month I was allowed to employ 5 coolies more per day so that means 20 per day. I have now progressed with the trench on the opposite side of the river [the 10x10 m 1896 Left-bank Pit, first reported in the September 7th letter] as deep as the bone bed [LB-HK], but nothing special has been found yet except for a deer skull with complete antlers [***Axis lydekkeri***], and an incomplete crocodile skull [***Crocodylus siamensis***], as well as several incomplete and complete deer antlers, leg bones, phalanges etc. Anything special about the man [34] [M…059-123, left] -ape has not been discovered yet, but I am saving all pieces of bone that I am unsure of recognizing. De Winter signed up for one additional year on September 6. Would you Sir, be able to give me some idea if an extension will be permitted for next year so that I can plan for that accordingly? Other than that, all is well with my health and also with the work activities, …. Kriele

[Page] 35 [M…059-123, right]
Trinil, October 1 1896.
Most distinguished Sir!
The latest sector [in the left bank 10x10 m pit] has just about been worked to its proper depth [through the LB]. In total, we have now excavated to a depth of 11.50 meters and we are already 1 meter below the lowest water level [that is, the top of the embankment had been 10.50 m above the low-water level]. We found [assuredly in the bonebed] 1 complete turtle [**Testudinoidea**], 1 incomplete [***Crocodylus siamensis***] crocodile skull, 1 incomplete [***Axis lydekkeri***] deer skull, 1 incomplete lower jaw of a pig [***Sus brachygnathus***] and also several incomplete and complete deer antlers, leg bones, vertebrae etc. etc. I have also started on a [36] [M…059-124, left] small sector upstream with a few coolies. This was left behind in the past [near the Ledge] and we were not able to finish it last year [1895]. In this small sector we find wood, shells [**Mollusca**] and bones [in the LB-HK], but the bone is mostly not very useful. So far, nothing special has been found. … your serving workman, Kriele

[The 1896 Left-bank Pit had penetrated the LB-HK when at an excavation depth of 11.50m below the terrace surface and 1.0m below lowest river level, much as had been the case three years when Kriele’s crew dug the 40-m Trench into the LB farther east.]

[Page] 37 [M…059-124, right] Trinil, October 12 1896. Most distinguished Sir! It appears that your request addressed to Mr. Pierson about which you informed me in your letter of July 27, asking to let me work several more months in Trinil or as long as it would be possible considering the water level, has not been favorably received. It appears that way because I have been given orders to discontinue working at the beginning of this month and cease all further work. I don't know exactly what the cause for this is, but according to [38] [M…059-125, left] statements by the chief in Tinggang it is all a question of funding. After the first of October, I have not accomplished very much except for packing the bones for shipment to Tinggang and I am now traveling with them. … I would very much like to know before January if an extension will be granted. If the answer is yes, I will continue to work for you, if it is no then I will leave the military service …. if [39] [M…059-125, right] you would help me in this matter and I am promising you that I will apply myself to the fullest at that work. …. your serving workman, Kriele

[Work in 1896 ended at Trinil.]

[Page] 40 [M…059-126]
Tinggang November 2 1896.
Most distinguished Sir!
Since I am back again in Tinggang, I am working with de Winter in manufacturing crates for shipment. The shipment is not very large this year since not very much was found at Trinil compared to last year. We found virtually nothing at Tinggang and also not much is being found now. As far as my health goes, I am suffering from fever but de Winter is doing fine, your serving workmen, Kriele and de Winter [November 15th letter about Tinggang omitted [41] [M…059-127].

[Page] 43 [M…059-128, right]
Tinggang December 15 1896.
Most distinguished Sir!
Several days ago an upper skull portion of a hippopotamus was exposed by the excavator (Picket 112), but the highly damaged excavator buckets have raked its teeth and these are now completely gone. In addition, the two tusks were broken off and we have not been able to find these either. Yet it is still a very nice skull and complete with the occipital portion attached and I don't believe that you have as nice a specimen as this one [Crate 9 '96 at Picket 110]**.** Also, we have now completed the job of packing all the bones. [44] [M…059-129] We have heard nothing about an extension. Other than that, we both are in very good health, Your serving workmen, Kriele and de Winter

[Page] 45 [M…059-130] [the sketch with unit grids on both banks of the river at Trinil, given above]

[Page] 50 [M…059-134] Tinggang 31 December 1896. Most distinguished Sir! [text is partially hidden at the right margin] We have finished the job of packing the bones but we have not yet received information as to when they should be dispatched. We did receive notice that an extension of 6 months has been requested [for us] but we don't know yet if it will be permitted. Other than that, we must report that recently nothing has been found here at Tinggang. Both of us are in good health, your serving workmen, Kriele and de Winter

[Two lists of fossils put into shipping crates are available, but most of the listed entries have no site indication, and Trinil finds were mixed with those from other sites. The specimens explicitly attributed to Trinil are "40 complete and incomplete deer antlers" [***Axis lydekkeri***] and "32 assorted leg bones." Kriele's letters mentioned other finds (above). The incomplete ***Crocodylus siamensis*** skulls in his September 11 and October 1 letters are not listed as Trinil specimens (our inferred species names are often omitted below.)]

[Page] 47 [M…059-132] Dubois wrote at the top of this page:

[This list] “goes with the letter of 31 December 1896 (and the sketch of the grid layout (at Trinil)”

[M…059-130, shown above at the beginning of –A4c] [The 10 crates were not shipped until March 1897, see official list below]

Complete list (by Kriele) of the contents of the 10 full crates [*listed without site provenience*]:

Crate 1: 1 incomplete turtle; 19 assorted bones

Crate 2: 1 complete turtle; 10 assorted bones

Crate 3: 37 assorted bones

Crate 4: 1 incomplete buffalo skull; 22 assorted bones

Crate 5: 1 incomplete tusk with a length of 1.50 meter; 1 incomplete turtle; 25 assorted bones

Crate 6: 39 assorted bones

Crate 7: 1 tusk with a length of 0.50 meter; 1 tusk with a length of 0.25 meter; 1 complete deer skull with incomplete antlers; 1 incomplete crocodile skull; 1 small chest with an assortment of 13 small leg bones; 1 small chest with an assortment of small phalanges and jaws; 1 small chest with an assortment of 10 small jaws; 1 small chest with an assortment of 45 molars and teeth; 35 complete and incomplete deer antlers; 1 incomplete lower jaw of a pig

Crate 8: an assortment of 8 bones [49] [M…059-133, right]; 1 small chest with an assortment of 175 molars and teeth; 1 small chest with an assortment of 18 small jaws; 1 small chest with an assortment of 14 small vertebrae; 1 small chest with an assortment of 19 small leg bones; 5 incomplete and complete elephant molars; 2 rhinoceros molars; 7 small phalanges; 1 incomplete deer skull with complete antlers; 1 small chest with an assortment of 10 leg bones; 1 incomplete antelope skull; 5 incomplete jaws of ruminants; 40 complete and incomplete deer antlers

Crate 9: 1 incomplete lower jaw of a hippopotamus (Picket 110 Tinggang, Solo valley); an assortment of 32 bones (Trinil)

Crate 10: 1 incomplete upper skull of a hippopotamus from picket 112 Tinggang, Solo valley; 15 ruminant molars; 1 incomplete lower jaw of a buffalo; assortment of 5 broken leg bones.

[Page] 171 [M…040-359] [Stationary head] Bevloeiingswerken in de “Solo valley”

March 24, 1897

Packing List of fossils, destined for the Geological/ Mineralogical Museum in Leiden
Sent by proa Tjunia, master Kasiman, from Tinggang to Bodjonegoro, for further shipping by the Resident of Surabaya

[Columns] Sequence Number – Type of Packing – Content – Measurements (Length, Width, Height) – Volume of Container in Cu M – Weight in KG

1 Crate [Content] 1 incomplete turtle; an assortment of 19 leg bones, phalanges (etc) from Lgandjar. [Length] 0.65 M, [Width] 0.60 M, [Height] 0.40 M; [Volume] 0.150 Cu M [Weight] 75 KG

2 Crate [Content] 1 complete turtle; an assortment of 10 leg phalanges (etc.) from Lgandjar [Length] 0.62 M [Width] 0.53 M [Height] 0.39 M [Volume] 0.128 Cu M [Weight54 KG

3 Crate [Content] An assortment of 37 leg bones, phalanges (etc) from Lgandjar. [Length] 0.71 M [Width] 0.52 M [Height] 0.43 M [Volume] 0.159 Cu M [Weight] 62 KG

4 Crate [Content] An incomplete [water] buffalo skull; 22 assorted vertebrae, ribs (etc) [Length] 0.80 M [Width] 0.51 M [Height] 0.45 M [Volume] 0.184 Cu M [Weight] 79 KG

5 Crate [Content] 1 complete elephant tusk with a length of 1.50 M in 10 pieces; 1 incomplete turtle; 25 leg bones, vertebrae etc. (Lgandjar) [Length] 0.72 M [Width] 0.58 M [Height] 0.59 M [Volume] 0.246 Cu M [Weight] 100 KG

6 Crate [Content] 39 assorted leg bones, vertebrae etc. (Lgandjar) [Length] 0.95 M [Width] 0.39 M [Height] 0.42 M [Volume] 0.156 Cu M [Weight] 67 KG

7 Crate [Content] 1 elephant tusk of 0.50 M length in 2 pieces; 1 0.25 M-long elephant tusk in 2 pieces; 1 complete deer skull with incomplete antlers; 1 incomplete crocodile skull; a small chest with 13 assorted leg bones; a small chest with 12 assorted small phalanges; a small chest with 10 small jaws and a small chest with 45 assorted molars and teeth; 35 complete and incomplete deer antlers; an incomplete lower jaw of a pig (Lgandjar) [Length] 0.71 M [Width] 0.60 M [Height ]0.45 M [Volume] 0.192 Cu M [Weight] 73 KG

8 Crate [Content] An assortment of 8 small leg bones and small phalanges; a small chest with an assortment of molars and teeth [175]; a small chest with an assortment of 18 small jaws; a small chest with 14 assorted small vertebrae; a small chest with 19 assorted small leg bones; 5 incomplete...

Carry over figures Transport Volume 1,221 Cu M; Weight 534 KG

[172] [M..040-360]… and complete elephant molars; 2 rhinoceros molars; 1 incomplete upper jaw of a pig; 7 small phalanges; 1 incomplete deer skull with complete antlers; a small chest with an assortment of [10] leg bones; 1 incomplete antelope skull; 5 incomplete jaws of ruminants; 40 complete and incomplete deer [***Axis lydekkeri***] antlers (Trinil) [Length] 0.71 M [Width] 0.50 M [Height] 0.41 M [Volume] 0.146 Cu M [Weight] 61 KG (55 KG)

9 Crate [Content] 1 incomplete hippopotamus lower jaw at Picket 110 near Tinggang, Solo valley; 32 assorted 32 leg bones, vertebrae (etc.) (Trinil) Length 0.75 M, widt 0.58 M, height 0.43 M; Volume 0.187 Cu M; Weight 80 (79) KG

10 Crate [Content] 1 incomplete hippopotamus upper skull [Picket 112]; 15 molars of ruminants; 1 incomplete lower jaw of a [water] buffalo and 5 pieces of broken leg bones (Solo valley, Tinggang) [Length] 0.73 M [Width] 0.51 M [Height] 0.36 M [Volume] 0.184 Cu M [Weight] 61 KG

Total : Volume 1.688 Cu M; Weight 736 KG

All crates have been marked as: A Government Goods, To the National Geological Mineralogical Museum in Leiden and B. are numbered 1 to 10.

Tinggang, March 24 1897 [February is crossed out]

Chief of 1st Department

[signed] T Vrijbergen

## S II-A4c.

[All excavations at Trinil during 1897 were on the left bank, simplifying interpretation of Kriele's letters about the work. Regarding 1897, de Vos and Sondaar (1982: 42) state: "the excavations started in August (Kriele 2-8-1897) and two trenches were dug with a grid system of square meters. One trench was downstream and measured 10x8 m, the other one was upstream and measured 12x17 m (Kriele 16-10-1897) [see illustrations below]. In the upstream trench in the month of September a left lower second premolar ($P_2$ sin.) was found; it was attributed to ***Pithecanthropus erectus*** by Dubois. The excavation came to an end in November." The premolar, the 1897 Premolar, is D.C. No. 11621., De Vos and Sondaar (1982: 48) also note that specimen of ***Sus brachygnathus***, a mandible] "Coll. Dub. No. 502," had an original label indicating "that the specimen has been found in grid square no. 23, 1.25 m below the lowest water level, in the downstream trench." Likewise, "a right upper first molar ($M^1$ dext., Coll. Dubois no. 317)" )" of ***Rhinoceros sondaicus*** has a label indicating that it came from "0.75 m below the lowest level of the river in grid square 32, of the downstream trench of 1897."]

## S II-A4d.

[Regarding 1897, de Vos and Sondaar (1982: 42) state: "the excavations started in August (Kriele 2-8-1897) and two trenches were dug with a grid system of square meters. One trench was downstream and measured 10x8 m, the other one was upstream and measured 12x17 m (Kriele 16-10-1897) [see illustrations below]. In the upstream trench in the month of September a left lower second premolar ($P_2$ sin.) was found; it was attributed to ***Pithecanthropus erectus*** by Dubois. The excavation came to an end in November." The premolar, the 1897 Premolar, is D.C. No. 11621. De Vos and Sondaar (1982: 48) also note that ***Sus brachygnathus***, a mandible] "Coll. Dub. No. 502," had an original label indicating "that the specimen has been found in grid square no. 23, 1.25 m below the lowest water level, in the downstream trench." Likewise, "a right upper first molar ($M^1$ dext., Coll. Dubois no. 317)" )" of ***Rhinoceros sondaicus*** has a label indicating that it came from "0.75 m below the lowest level of the river in grid square 32, of the downstream trench of 1897." ] [see records in -A4e and –A4f, below].

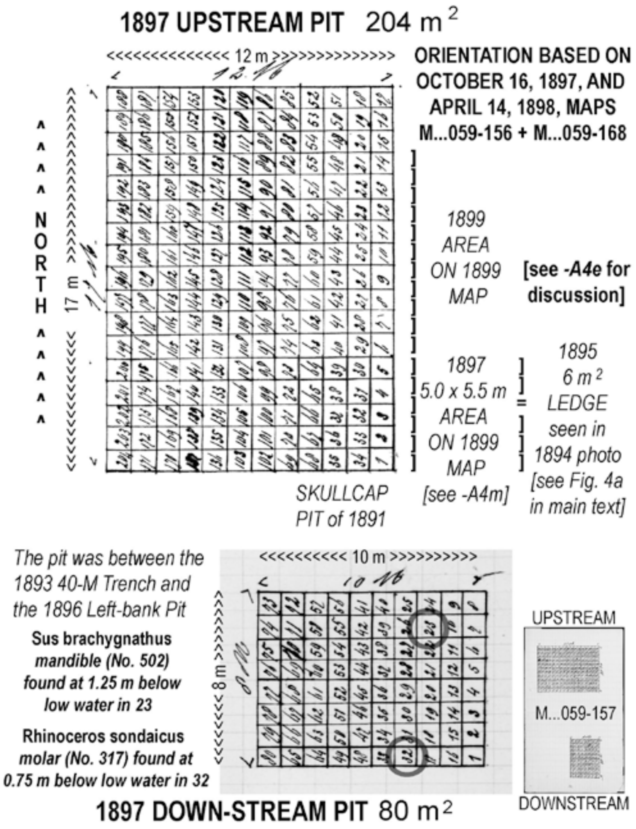

(i) [M…059-141/142]

Tingang 30-3-1897
…. About Trinil it would be best to start at the other side of the River [left bank] as there you could even start a block downstream or upstream of 10×10 m that could be worked by 25 men. … best to get prisoner workers …. I don't believe that much is to be gained by digging on the side where the houses stood [on the right bank]. By the way, it is also easier to carry away the spoils at the opposite side of the river [left bank] since previous spoils are being washed away each year [10] [M…059-142, right] which is not the case at all on the side of the houses [right bank]. The opposite side of the river also gives us the best chance of finding bones…. your serving workmen, Kriele and de Winter

[Work was planned "at the other side of the river [Dutch "andere kant"] …. [across from] the houses on the right bank.]

(ii) [M…059-143, 144]

[Stationary head: "Bevloeingswerken in de Solo vallei" = "Irrigation works in the Solo valley"]

Tinggang, May 16 1897
Most distinguished Sir!

As far as finding bones here at Tinggang is concerned, the situation remains the same. We have found nothing this month except at Ngluwak, consisting of a very incomplete tusk of an elephant. I have heard nothing yet about a move to Trinil and the local Department Chief cannot give me a definite time for that, notwithstanding the fact that I told Mr. van Rossum several days ago that it certainly would be possible to work at Trinil at the beginning of May.

I think that de Winter will leave for Europe when his service time has ended. He told me about it this morning.

I have been placed at the Solo valley [project] and I have started working on construction of new houses. I enjoy my work very much which by the way turned out to be better than I expected.

I have met a very nice and sweet woman and we like each other very much and if I am not mistaken I may also become a father.

Otherwise all is well with us and we hope from the bottom of our hearts that you have overcome your illness, your serving workmen, Kriele and de Winter

(iii) [M…059-144]

Tinggang June 4, 1897
…. In addition one of us will go to Trinil but the date has not been determined yet. We could have easily been working there for one month and a half now. I also have [M…059-145, left] told Mr. van Rossum that it would certainly be possible to work there at the beginning of May, but it seems that they are waiting to see if an extension will be granted again after these 6 months. Otherwise all is well with my wife and I. De Winter is currently a bit confused, since he does not know what he wants to do during the last part of his service time. … [M…059-145, right] Tinggang 16-5-1897 …. We also have not heard a word about Trinil and when one of us can go there. Also, nothing is being found at Tinggang. As we have previously written to you, we hear that it could easily be a few months …. [M…059-146, right; M…059-147, left] Tingang 13-7-1897 …. In two months time, de Winter will travel to Europe…. We are continuing to wait for Trinil. … Would you be able to arrange for us to go there next year (if indeed an extension is granted) in the month of June? That is the time when we can easily already start working and a lot of formation material has to be removed before we reach the bone bed. We would likely have to displace a huge amount of sediments in order to have a better chance of finding bones.… . your serving workmen, Kriele and de Winter

(iv) [M…059-148/149]

Trinil 2-8-1897
… de Winter has opened a square [on the left bank] of 10×10 m [and is now digging] at 2.5 m depth with 20 forced labourers … too few people ….

(v) [M…059-150]

Trinil 31-8-1897
…. I have the honor of letting you know that I relieved de Winter yesterday since he is leaving for Europe after his service is ending. This is a very nice trench upstream and adjacent to the [left-bank] trench of previous years. It is a pity that we can't get more workers. Up till now, 7.5 meter [in depth] has been excavated, measured from the [embankment] surface and the trench measures 12x12 meters. Also, some small bones have been found but nothing very special. … I also heard from Mr. van Rossum that this year will most likely be the last year that an extension is granted. …Kriele

(vi) [M…059-152]

Trinil 17-9-1897
…. the [expanded 12x17 meters 1897 Upstream] pit has now been dug 9 meters [downward from the terrace surface] and has yielded, since the relief [of de Winter], one complete [***Axis lydekkeri***] deer skull with antlers, a small kind of tusk of an elephant [***Stegodon trigonocephalus***], and several diverse vertebrae, carp/tarsals, teeth and molars but all in all nothing special and we are not yet deep enough to find larger amounts… [apparently, still above the PFZ].

[At 9 m depth from top, as the 1897 Upstream Pit was expanded to 12x17m, an ***Axis lydekkeri*** skull, a ***Stegodon trigonocheplus*** tusk and small fossils were collected.]

(vii) [M…059-153]

Trinil 16-10-1897
…. the pit of 12 × 17 meters has been brought to the depth on which no more bones are to be expected [that is, it had penetrated the LB]. I have started a second pit 10×8 metres [down stream of the 40-m Trench] that I hope to be able to finish before the river rises … lately found are 2 complete elephant molars, together with several other types of molars and teeth, but nothing special, no larger bones at all, and smaller bones a few. ...
[See S II-A4e, above] [The LB-HK was penetrated in 1897 Upstream Pit.]

(viii) [M…059-154]

Trinil 4-11-1897
…. the pit of 10×8 meters we have already brought 8 meters down, but nothing special has come out yet, some [***Axis lydekkeri***] deer antlers, vertebrae, an elephant's molar [***Stegodon trigonocephalus***], a rhinoceros molar [***Rhinoceros sondaicus***], some legs, teeth and molars. About the ape [**1897 Premolar**] molar that we found, it appears that the most respected Mr. Pierson is very interested in this, since I received a message to make a small sketch of the find spot where in the past the skull and the femur have been found and how far apart from each other they were found [see maps -A4e and –A4f, above]. I heard from mr Vrijbergen that he has sent you a drawing of that molar and that he will also send you the molar …. Kriele

[Fossils in the upper 8m of the 1897 Downstream Pit were "nothing special." A "primate molar" was found in the 1897 Upstream Pit.]

(ix) [M…059-158/159]
Tingang 1-12-1897.
I have the honor of letting you know that the work at Trinil has now been finished. The final trench was dug to a depth of 12 meters. During the last half month we found 1 complete tusk of an elephant with a length of 1.55 meter [***Stegodon trigonocephalus***] in 5 pieces; see Crate 2 below], 2 incomplete lower jaws with complete molars, 2 very incomplete upper jaws with complete molars [see Crate 3 below], 1 incomplete skull of a cow with complete horns [***Bibos palaesondaicus***], 1 incomplete skull with 1 complete horn of a water buffalo [***Bubalus palaesondaicus***; see Crate 2 below] as well as some deer antlers [***Axis lydekkeri***], leg bones, vertebrae, pasterns etc. [also, Crate 8 below]. …. The chief here at Tingang says that for this year the research is finished …. Kriele

[1897 Downstream Pit was dug to 12m in depth and found a ***Stegodon*** tusk, a *Bibos* skull, a ***Bubalus*** skull and ***Axis*** antlers in LB-HK.]

(x) [35] [M…059-159, right]

Tinggang December 17, 1897.

Most distinguished Sir!
I received your letter dated November 10th in good order. You (or better said you, respected Sir) are asking me in that letter, whether I have given up on the job with the National Railways. No Sir that is absolutely not the case. I hope to work there and I am sure that I then will be able to advance in my work with all that I have learned before, judging from what I can tell by the many folks that are now working in the Solo valley. But, nevertheless, I will never abandon my work for you, even if it will last many more years. You, respected Sir can count on me, but it is only unfortunate that I do not make any advancement in salary and you Sir, cannot do much about that. [36] [M…059-160, left]. About the job of packing the bones, I can tell you that I have finished that this afternoon, exactly on my wife's birthday as she turned 20 years of age. In this letter, I have enclosed a small sketch of our work arranged in numerical order in which the bones were found [M…059-157, above], the content of the crates as well as the quantities. According to what I heard from Mr. Vrijbergen, he will send you a sketch [showing] at which level the last molar was found [the **1897 Premolar**]. I don't know yet when I will be able to send the crates. In the case that I can continue exclusively working on collecting bones, it would be a great advantage since I then could travel from one place to the other to check on the presence of bones. It would then not be necessary to have you call me up unless I would just by chance happen to be there. Nowadays, it is very easy to move from one place [37] [M…059-160, right] to the next since the workers train travels along the entire Solo valley. The only thing is that I then will have greater expenses because of the cost of my meals which I would have to arrange for myself during my travels and I would thus have to support two households. So please forgive me for having asked in the beginning of this letter an allowance of a small increase in salary, if possible, since traveling is so expensive. Otherwise, my wife and I are doing very well. Incidentally she still has no child. She had an accident on the way to Banglee, but notwithstanding that, she is now again in that state [of pregnancy]. She asked me to humbly thank you Sir for the concern you have shown for her. Serving you respected Sir, your workman, Kriele

[The sketch that showed the level at which the last molar was found is not present in the Dubois Collection archive. The following shipping list of 1897 fossils, prepared by Mr. Vrijbergen on December 24, 1897, gives no stratigraphic data, but attributes 283 specimens to Trinil, all of which must have come from the left-bank excavations (the 1897 Upstream- and Downstream-Pits).

At least 56 of the 1897 Trinil specimens (>20%) were vertebral elements. Notable also were:

"an incomplete [**Bubalus palaeokerabau**] buffalo skull with one incomplete horn" [also in 12-01-87 letter];
"an incomplete [**Stegodon trigonocephalus**] elephant tusk … length 1.15 m" [not noted in letters];
"an incomplete [**Stegodon trigonocephalus**] elephant tusk … length 1.55 m" [also in 12-01-87 letter];
"1 incomplete [**Stegodon trigonocephalus**] elephant upper jaw with complete molars;" and
"1 upper and 2 lower jaws of a pig," [**Sus brachygnathus**, which is not mentioned in letters, but could be Collection Dubois No. 502, referred to above].

(x) [A second list (on the following page) has "1 molar of Rhinoceros [**Rhinoceros sondaicus**] Trinil" [noted in the 04-11-97 letter; Dubois Collection No. 317 which was found at 0.75 m below the low water. Our inferred species names are omitted below]
[Page] 178 [M…040-366] [Stationary head] Bevloeiingswerken in de "Solo vallei" 508/19 1898
Copy 1st Section 1st Department

Packing List of fossils destined for the Geological, Mineralogical Museum in Leiden, shipped by S.S. Gede.

Columns: Sequence number; Packing type; Content; Measurements (Length, Width, Height); Volume in Cu. M; Gross Weight in K.G.

1 Crate [Content] An incomplete elephant skull (Grebehan – Tenggor) [Length] 0.89 M [Width] 0.82 M [Height] 0.82 M [Volume] 0.598 Cu M [Weight] [?? KG]
2 Crate [Content] An incomplete [water] buffalo skull with one incomplete horn; an incomplete elephant tusk in 4 pieces, length 1.15 m; a complete elephant tusk in 5 pieces, length 1.55 m at Trinil [not mentioned in the letters]. [Length] 0.67 M [Width] 0.68 M [Height] 0.52 M [Volume] 0.237 Cu M [Weight] 112 KG
3 Crate [Content] 2 incomplete elephant upper jaws with complete molars; 1 elephant molar and 7 vertebrae at Trinil; a piece of skull and molar of an elephant belonging to crate # 1 (Grebekan – Tenggor) [Length] 0.76 M [Width] 0.73 M [Height] 0.48 M [Volume] 0.266 Cu M [Weight] 124 KG
4 Crate [Content] 1 elephant tusk, length 35 cm; Two elephant tusks, length 30 cm; an elephant lower jaw with molars; 1 incomplete skull with complete horns of a cow; 1 incomplete upper jaw with complete molars of an elephant; 22 assorted vertebrae at Trinil [Length] 0.71 M [Width] 0.68 M [Height] 0.42 M [Volume] 0.203 Cu M [Wt] 88 KG Carry over Total [Volume] 1.304 Cu M [Weight] 324 KG

[Page] 179 [M…040-367]

Sequence number; Packaging type; Content; Measurements (length, width, height); Volume per package in Cu M; Gross weight in KG

5 Crate [Content] 1 incomplete [water] buffalo skull from Ngluwak; 3 elephant molars and a rhinoceros leg from Tinggang; 1 incomplete elephant upper jaw with complete molars and 27 assorted vertebrae from Trinil. [Length] 0.81 M [Width] 0.61 M [Height] 0.53 M [Volume] 0.262 Cu M [Weight] 106 KG
6 Crate [Content] 1 small chest with 66 teeth and molars of ruminants; 20 small deer jaws; 370 teeth and molars of deer; 1 small chest with 135 assorted teeth and molars; 22 assorted small leg bones; 103 assorted small vertebrae, knuckle bones (etc) from Trinil. [Length] 0.67 M [Width] 0.62 M [Height] 0.58 M [Volume] 0.241 Cu M [Weight] 83 KG
7 Crate [Content] 80 bones from Trinil. [Length] 0.71 M [Width] 0.63 M [Height] 0.62 M [Volume] 0.277 Cu M [Weight] 107 KG
8 Crate [Content] 1 incomplete deer skull with complete antlers; 1 incomplete deer skull without horns; 6 rhinoceros molars; 7 elephant molars; 21 shells; 1 upper and 2 lower jaws of a pig; 36 assorted bones from Trinil and 7 sample rocks from Banglee (Kertorono) [These samples were evidently the result of Kriele's trip to investigate an oil seep near Banglee on which he reported in his letter of August 2 1897]

[Length] 0.62 M [Width] 0.56 M [Height] 0.56 M [Volume] 0.194 Cu M [Weight] 79 KG
Total [for this shipment] [Volume] 2.278 Cu M [Weight] 699 KG

All crates have been marked as:

A. Government Goods to the National Geological Mineralogical Museum in Leiden and B. Numbered from 1 to 8.

Tinggang 24 December 1897, The Chief of the First Department (signed by) Vrijbergen

For an identical copy by: Secretary Bookkeeper [signature] H.C. Bruine [?]

(x) [The following listing appears to have been made in Leiden when 1897 fossils were unpacked; the handwriting is not that of Dubois, but some corrections are his. Only one fossil is identified as coming from Trinil, and most others have no provenience information. [our inferred species names are often omitted below]

[Page] 175 [M…040-363]

1898 [arrival year]

1. elephant molars; deer antlers, bones, ruminant vertebrae
3. Elephant tusks
   2 Lower jaw and molars; Thigh bone elephant; Shells
   2 rows of stegodon molars

Columns -10 10 20 30 40 [*only entries are in Column10, but they are all crossed out*]

[Page] 176 [MM774C-000040-364] [Continuation of list prepared in Leiden] 1898 [arrival year]
1 mol[ar] Rhinoceros Trinil [presumably the ***Rhinoceros sondaicus*** mentioned in the 04-11-87 letter]
molar young Stegodon (6 pl.) 5 M above lowest level upstream. No. 169 below Kaverly?
Same 5 pl. mo 58. ~ 1.25 M below lowest level downstream also No. 169 below Kaverly?
Tusks elephant; Molar *Stegodon* (upper jaw) and small fragments of molars; Fragment of a turtle
Tusk in many pieces; Deer antlers; Bones and vertebrae ruminants; Vertebrae and ribs of a Banteng; Metacarpus;
Molar, heel bone, small tusk *Elephas*; Large shells; Lower jaw fragment with 5 molars;
(2) Lower jaw stegodon; 2 large shells; thigh bone elephas; lower jaw stegodon; ~10 fragments of molars with skull portions *Steg Trig*?; Shells; incomplete right lower jaw with chin portion hippopotamus;
(2 and 3) tusk *Elephas*; occipital skull portion of hippopotamus with 2 molars; piece of lower jaw of *Elephas* with molar.; (1 and 3) tusks *Elephas*;
(3) skull *Stegodon* with 2 complete molars Trinil; 2 shells *Ostria*; (no numbers) 2 rows of *Stegodon* molars; 3 lower jaw fragments Hippopotamus [crossed out].

**S II-A4e.**

[Kriele's first sketch map of the left-bank excavations since September 18, 1891, drawing dates to 1897 and has the discovery locations of the ***Pithecanthropus erectus* Skullcap** and **Femur I**, as well as the **1897 Premolar**. We have added our own annotation in bold font, such as "1893 40-m Trench, to those of Dubois' in pencil.]

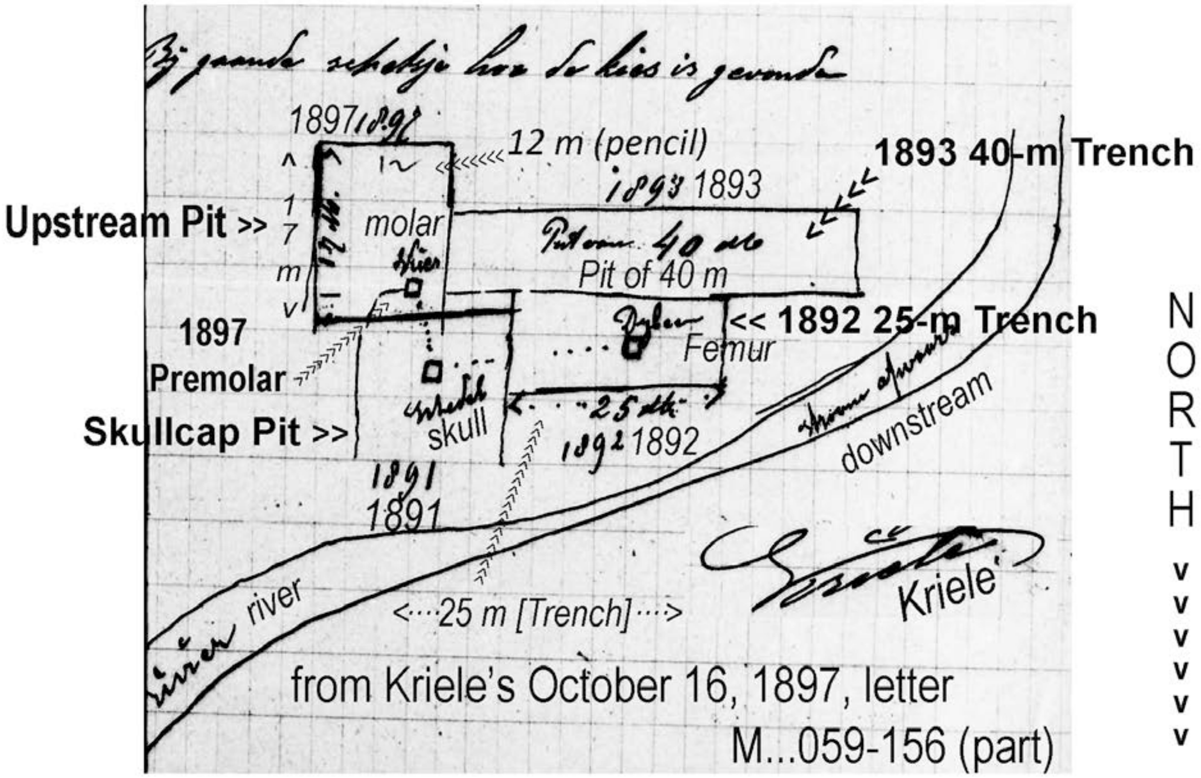


[The sketch appears to show the find spot for the 1897 Premolar (also, -A4i and –A4k). This fossil, a left inferior premolar ($P_2$ sin.), was announced as a *Pithecanthropus erectus* find by Dubois (1899) at the 1898 Fourth International Congress of Zoology in Cambridge, U.K. (de Vos and Sondaar 1982; Theunissen 1989; Smith et al. 2009). The specimen is sometimes known as Trinil 5 (Jacob 1975; Schwartz and Tattersall 2005, who published a high-quality photograph of the specimen), and sometimes is mistakenly said to have been found in 1898 (Indriati 2004).

The mapping shows that the 1897 Upstream Pit was immediately south of the 1891 Skullcap Pit and east of the 1893 40-m Trench. Thus, the 1897 Upstream Pit finished removing the sandstone Ledge, which we see in the 1894 photograph was located just south of the inundated 1891 excavation (Figure 4a, main text). The Ledge outcrop is surely the "big rock at the other side" (-A4b 'page 11' above) and "ledge measuring 6 square meters" (-A4b 'page 30' above) mentioned by Kriele in 1895, when he collected fossils there, and the place that interested him in 1896. Despite reporting a 17x12 m dimension and providing a gridded plat of a pit of this size (-A4d, above), Kriele showed the 1897 area of excavation to be only 5.0x5.5 m (~27.5 $m^2$) in his scaled map of December 21, 1899 (-A4m; Figure 3a, main text). The 5.0x5.5m seems to be the 6$m^2$ ledge that he mentioned in 1895. This 1899 map indicates that the area to the north was excavated in 1899 and 1900. See –A4o.]

**S II-A4f.**

[With respect to 1898 operations, de Vos and Sondaar (1982: 42) state "Kriele (14-4-1898) made proposals for excavations to be carried out at Trinil in **1898**, but nothing came of this. In a sketch accompanying the proposals the trenches of previous years … are clearly seen" (see next page; also de Vos and Aziz, 1989: 14, Figure 5). The proposal was from Dubois to Kriele. The sketch is given below. The relevant letters of Kriele are:]

(i) [M…059-161 and -162 contain labels for the year 1898]
Page 1 [M…059-163] [handwritten in margin by Dubois: *"1898"]*
Tinggang, January 31, 1898.
Most distinguished Sir! I received your postcard of the 17th in good order in which you, respected Sir, ask for a sketch of the find spot of the tooth [1897 Molar; see –A4e, above], but I [already] sent you a letter at the end of December [1897] in which everything is explained. And from what Mr. Vrijbergen told me, he was going to send you a sketch as well. He copied that from the small sketch that I have sent you. In regard to the work activities in Trinil, in the event that I would be called upon to work there, it would be best to start in the month of May and then begin on a trench upstream [to the East] with a large group of workers. The crates [containing 1897 fossils] are going to be shipped tomorrow or [Page] 2 [M…059-164, left] the day thereafter to Surabaya and from there onwards [to Europe]. ... From December 30 till January 17, I have been in the hospital in Batavia [now part of Jakarta] because I was bitten by a suspected rabid dog, but in spite of that I have now recovered again. Other than that, all is well with my health and also with my wife, your serving workman, Kriele

(ii) Page 3 [M…059-164, right]
Tinggang, February 28, 1898.
Most distinguished Sir!
During this month we found below Tinggang in the Solo valley at Picket 96 an incomplete skull of a cow and at Pickets 114 -116 an incomplete lower jaw of a hippopotamus [Crate 2 '98] as well as 2 incomplete elephant molars [Crate 1 '98]. Other than that, all is still the same here. Work is progressing very slowly because so few free workers (coolies) are showing up. I have not heard anything about going to Trinil and it would be best that you insist that we start as soon as possible. If we [Page] 4 [M…059-165, left] were to begin too late it would not be worth the effort and there also would be much less chance of finding something [(?) as deep in the excavation as the main bonebed]. Other than that, all is very well with me and my wife, your serving workman, Kriele

(iii) Page 5 [M…059-165, right]
Tinggang, March 22, 1898.
Most distinguished Sir!
This morning I asked Mr. Vrijbergen if anything had happened in regard to my departure for Trinil, since this has become urgent if we wish to find something. Mr. Vrijbergen tells me that he does not know anything yet and he also does not know yet if an extension permit will be granted. Would you respected Sir, be so kind then to send me a letter which I could show to Mr. Delprat and if you have no problem with it also a certificate for the time that I have worked for you. In the event that your work is finished or an extension permit is not granted, my preference would be to work at the Railways. I am convinced that if I am as conscientious at the Railways as I have been here [M…059-166, left] at Tinggang, I will make very rapid progress, thus I humbly request from you, respected Sir, help with what I was writing about above. And as you know, I am now married and I have to take care not only of myself. At Pickets Nr. 116 and 117, the mechanical excavator has removed the second cut, [we found] 2 thigh bones [Crate 2 '98], 2 upper front leg bones [Crate 2 '98], 1 elephant tusk of about 1-meter length [Crate 1 '98], 1 incomplete skull with a few molars belonging to a hippopotamus [Crate 2 '98]. Other than that all is still the same here and my wife and I are doing very well, your serving worker, Kriele
Page 7 [M…059-166, right]

(iv) Tinggang 14 April 1898.
Most distinguished Sir!
The situation here is still the same. I have not heard a thing about an extension nor have I heard anything about Trinil. In regard to finding bones in this second cut with which the mechanical excavator has now been busy for two months, nothing has been found, except for what I mentioned to you in my last letters. This second cut is being dug to a depth of about 5 meters of which approximately 1 meter is below clay (or marl) and thus [in] about 4 [meter] of sand. The same as what you have seen in the past from above, where previously the [varkensvoet? = pig's foot] was found in light [Page] 8 [M…059-167] [colored] rocks. It is exactly the same as when you observed the bones there. That is to say, that nothing has been excavated yet at that spot. Please don't hold it against me that my letters are now arriving so intermittently. I am always thinking each day let me wait a little longer, since it is possible that I still could find something or that I will hear good news about an extension or a move to Trinil, but so far I have not heard a thing about that. Have you received the shipment of bones for the year of [18] 97? Other than that, all is well with my health and that of my wife, your serving worker, Kriele

(v) [M…059-169] [Blank page, then page] 10 [M…059-170]
Tinggang, April 17 1898.
Most distinguished Sir!
I still have heard nothing about an extension. Several days ago I found 2 elephant tusks [Crates 1 '98], but you will notice that for both the points are missing which is a real pity because they measure now about 2.50 meters in length. If the points had been preserved they likely would have reached 3 meters. We also found several pieces of a turtle but small bones or small teeth have not been seen. Other than that, all is still the same here and my wife and I cannot complain about our lives, your serving workman, Kriele

(vi) [Page] 11 [M…059-171]
Tinggang May 20 1898 [*"received June 20", as written by Dubois*].
Most distinguished Sir!
I received your letter dated April 14 in good order and I learned from it that I will have to go to Trinil at the end of May. It will be my pleasure to work there on the sectored trenches [vakken in Dutch] I, III and IV in the sketch [M…059-168, below]. These are nice trenches but number II has been completely excavated since there is no sandstone left to be dug any longer. That is unless the river would lower enormously but I don't believe that the water will get that low. Mr. Vrijbergen has not heard a thing about the extension and if we still have to wait some time [Page] 12 [M…059-172, left] for instance till the end of May just like last year, then I am afraid that we will not be able to complete the 4 sectored trenches indicated on the sketch. Nothing has been found lately at Tinggang but the mechanical excavators work now day and night. Other than that, all is well with my health and that of my wife, your serving workman, Kriele

(vii) [Page] 9 [M…059-168]

[The sketch map of April 14, 1898 (next page), shows the proposed sectioned trenches (vakken) I–V juxtaposed to 1897 and 1893 trenches, although Trench II was fully worked over according to Kriele. Kriele drew the outlines of the 1897 Upstream Pit (writing *Put 1897*), 1893 40-m Trench (writing *de put van 40 m*), 1897 Downstream Pit (writing *Put 1897 stroom opwaarts*), 1896 Left-bank Pit (writing *1896*); he noted the "molar" (*kies*), skull (*schedel*), femur 1892 (*dijbeen 1892*); Dubois added a proposal for trenches I to V. This sketch is the only known map showing the location of 1896 Left-bank Pit. It was immediately west of the 1897 Downstream Pit, and must have removed part of the bank west of the 1891-1893 excavation area, as seen in the 1894 photograph (Figure 4a, main text).]

[De Vos and Sondaar (1982: 42) state that "Kriele (14-4-1898) made proposals for excavations to be carried out at Trinil in 1898, but nothing came of this. In a sketch accompanying the proposals the trenches of previous years … are clearly seen." Proposed excavations I-V were Dubois' ideas.]

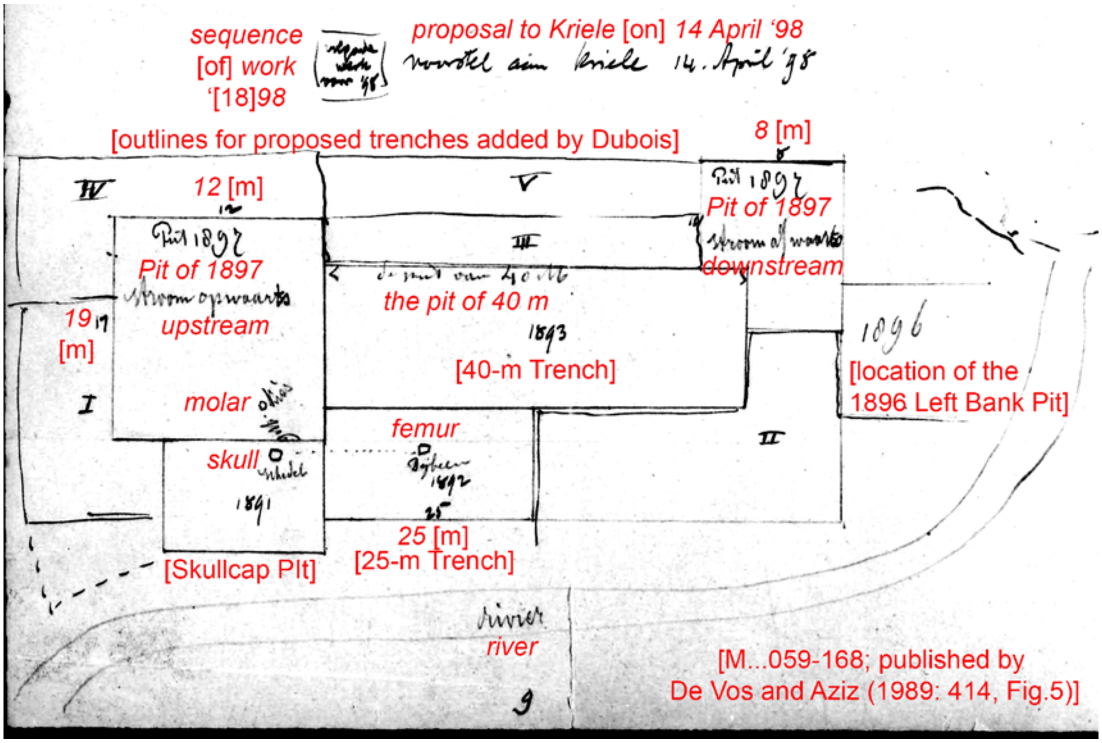


[Handwritten by Dubois] *proposed to Kriele* [on] *14 April 1898* [was a] *sequence of work for* '[18]*98*

(viii) [Page] 13 [M…059-172, right]

Tinggang, July 1 1898.
Most distinguished Sir!
I asked Mr. Vrijbergen two days ago when I can go to Trinil and he told me that Mr. van Rossum, the Chief in Padangan knows nothing about when the time comes to go to Trinil. If we wait much longer and no adequate numbers of prisoner workers or free coolies are assigned to us, it will not be worth our while to even start on this. It would be best that you respected Sir, request as soon as possible sufficient coolies [Page] 14 [M…059-173] since we would then have the best chance to find something. Here in Tinggang, we recently found yet a beautiful very complete elephant skull with complete molars [Crate 3 '98]. Since the mechanical excavator is now down for repairs, nothing is being found. Other than that, my health is very good and my wife is doing fine as well, I am happy to be serving you, respected Sir, Kriele
(ix) [Page] 15 [M…059-174] Drawing of a bone:

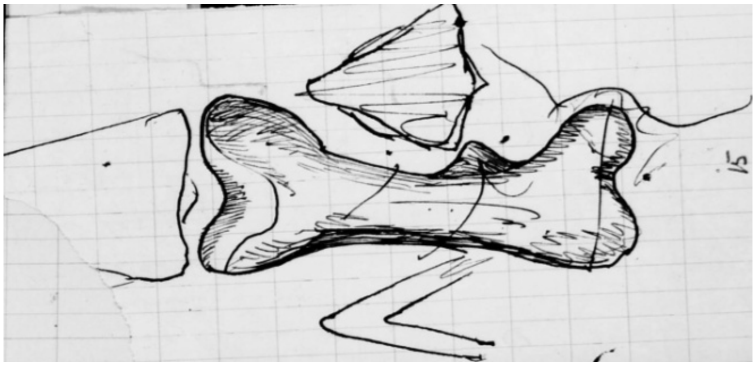

(x) [Page] 16 [M…059-175] [Slip of paper with the following note, probably pertaining to shipment of fossil drawn above]: "Sturdy wooden box … packed as registered and postage stamped with 3…. sea mail (per 15 gram or parts of 15 gram 10 cents"

[Page] 21 [M…059-178, right]
Tinggang, June 10 1898.
The situation here at Tinggang is still the same. There were a few pieces of bone that surfaced near Picket 196 but not much of great value and no digging is taking place at other locations. From what I hear, it looks like Mr. Pierson will retire to Europe. Should the bones that have been found here be sent off? Since it is not very pleasant to work here at the Solo Valley [project] and because personnel will be drastically reduced, I restate my request [22] [M…059-179, left] to be placed with Mr. Delprat at the Railways and for you to send me a letter with what is needed for me to approach this esteemed man. All is well with my wife and with my health and since no news can be reported from here, I remain with great respect for you, your humble serving workman, Kriele

(xi) [Page] 19 [M…059-177, right]
Tinggang, June 19 1898.
The situation here is still the same. Nothing is being found since the mechanical excavator is out of order most of the time. I did ask if I could go to Trinil in connection with the photographic work [none from 1898 is otherwise known], but I did not get an answer to that. Mr. Vrijbergen has been transferred to Makassar and he left already 5 days ago for his new assignment. As far as myself is concerned, I would very much like to be placed at the Railways, since here at the SV [Solo Valley Project] things are getting more discouraging by the day. The only trend I see is one of cut backs and I read [20] [M…059-178, left] into it that in a few months time many folks will have to look for another job. So I hope and wish that you, respected Sir can assist me with that and before this will happen provide me with a written statement for the respected Mr. Delprat, if possible in the form of a certificate. There is a good possibility that I will be one of the first [to be fired] since no work is being done that requires my specialty. In the case that I could remain here I would, because of the cut backs, be placed on a salary of F 45 which is what is given to most and so I am asking for your help. All is very well with my wife and my health. I am happy to serve you, respected Sir, Kriele.

(xii) [Page] 17 [M…059-176, left] repeat of …059-175 on top of 174 (pages 16 on top of 15) [17] [M…059-176, right]
Tinggang July 31 1898.
Most distinguished Sir! I still have not heard a thing about going to Trinil. I don't believe that anything will come of it since I recently asked Mr. Vrijbergen when it might happen but he still did not know anything. He did tell me that Mr. Pierson did not dare make the decision himself and he must have written about it to Batavia. Nothing can be found here at Tinggang, since the work has virtually come [18] [M…059-177, left] to a halt. One of the mechanical excavators does not function and the other one works in the clay formation. Other than that, there is no news but my health is fine and also that of my wife. I am happy to be serving you, respected Sir, Kriele

(xiii) [Page] 23 [M…059-179, right]
Tinggang 15 November 1898.
Most distinguished Sir! You must certainly have heard about the great setback we had with the stoppage of work in the Solo valley. It looks very hopeless here for the hundreds of Europeans who will therefore all be without income and that includes me as well since I asked Mr. van Rossum yesterday if the work for you was included in that and his answer was yes! I am allowed to pack the bones and these will be sent to you. Dear respected Sir please be so good to help me get [24] [M…059-180, left] that promised job with Mr. Delprat since otherwise I will suddenly be in trouble and if I would be without work here for a long time or would fall into poverty, I have already made a decision in that case to immediately depart for Europe

and make ends meet on my small pension. The situation regarding the workers here looks very discouraging and it might well be possible that unrest will start to happen. Other than that, no news from here except that my wife and I are in good health and are hoping for the best with your helping hand. And so I remain as always, your humble and serving workman Kriele.

(xiv) [Page] 25 [M…059-178, right]
Tinggang, December 29 1898.
Most distinguished Sir!
I have the honor of letting you know that the day after tomorrow I will leave here for Padang [Sumatra] and will be placed under Mr. Delprat in the coal mines. Since the work here in the Solo valley has been stopped and I was told to find a job, I went ahead and wrote to Mr. Delprat and included the letter that I received from you in the past. As a result I have been appointed for which I remain very grateful to you. I have recently read in the newspaper that your research [26] [M…059-181] will continue as well as the associated work activities. In the case that you will need me, you will thus know where I am and if I could be of service to you, just give me a warning. It will be a pleasure to again do such work for you no matter where it is. The [3] crates with bones have now been sent but nothing very special is amongst the content. [see document M…040-361 Other than that, there is no further news and so we wish you and your family a happy new year, your humble workman and his wife, Kriele
[Kriele worked from December 1898 to September 1899 in the Sumatran coal mines.]

**S II-A4g.**

[All excavation in 1899 was done on the left bank with most of that excavation being south of the 1893 40-m Trench. Regarding **1899** excavations, de Vos and Sondaar (1982: 43) state: “The excavations started in September (Kriele 16-9-1899). Figure 3 [copy in –A4h, below] shows the work plan for that year. The [1899] trench is divided into a square meter grid system." De Vos and Aziz (1989: 414) add that “from the figure we can deduce that the surface of the trench is 510 $m^2$. Totally there was excavated during four months.” De Vos and Sondaar (1982:47) report that among “two porcupine [***Hystrix lagrelli***] molars ... in the Dubois Collection … no. 1482b [was] found in 1899,” but it may not have come from the LB-HK.]

(i) [Page] 1 [M…059-184] [letter from van Rossum in Padangan to Dubois in Amsterdam]
Dr. E Dubois. Amsterdam. Padangan, April 29 1899.
Most respected Sir, I was pleasantly surprised upon receiving a copy of your presentation given in Amsterdam …. your “Kringloop der stof op de aarde” [Circulation of basic compounds on earth]. … [Page] 2 [M…059-185, left] …. Because as rewarding as it may be to displace a few million cubic meters of material rock [in operations of the irrigation works project in the Solo Valley], it is not especially mentally stimulating. This displacement of sediment has also not been as beneficial for paleontological research as one would wish. It is indeed a pity that Trinil [Dubois initialed in the margin: *ED*] has been inactive this year. On top of that Kriele has departed to Chief Engineer Delprat in Sumatra because of the lack of work resulting from the discontinuation [of the Solo Valley project]. Whether or not the work activities in the Solo valley will be resumed will especially depend on the views in this matter of the political representatives. From a general economical point of view, it would certainly be a pity if this peaceful exploitation of such a rich soil would not materialize. …. [Page] 3 [M…059-185, right] as well as my kindest regards also of my wife. With the highest regard for you, respectfully, van Rossum

[Trinil was inactive during most of 1899 while Kriele was in Sumatra. His letters started in September 1899, after completing time in the coal mines there.]

(ii) [M…058-474]
Trinil, 16 August 1999
Most learned Sir:

It is my honor to let you know that I arrived at Trinil on the 14th of this month and I have already started my work. As you certainly will realize that in the short amount of time that is left for us before the West monsoon begins, not much can be accomplished any more. [And] as you once suggested that we work through the rainy monsoon period and work the terrain as much as possible on top when spoils would wash away during banjirs [floods], [it] would leave us facing a nice number of meters to work on during the East monsoon. That is after all [the low-water levels that occur during the dry season are] where the bones are expected to come from and we would then have a good chance of finding the intended objects as well as other nice ones [in the LB-HK main bonebed].

As I have heard, there is also a great chance that as soon as the rains start in December, I would have to return to Sawah Lunto [Sumatra] (since I have been loaned out to that place), [and] in which I no longer have much interest. It is there very dangerous working and it is unhealthy. I will not say [M…058-475] that there is not a good future there, but you are playing with your life. I was there for 6 months and have already been buried under the coal and experienced a hydrocele [watery fluid collection] as a result, which will bother me for the rest of my life. These are matters that are not very appealing, and on top of that, there are many other incidents that you hear about and see every day. Would you perhaps know of a different job for me, it doesn't matter where it is as long as it is not in the coal mines, since I have developed a terrible aversion for that and also am a bit too old for that?

I will do my best to take care of the work here as well as I can and the bones that we find, should they be sent to the same address?

Other than that everything is going well with my wife and I.

Your respectful and humble servant,
Kriele

**S II-A4h. -A4f.**

(i) [4] [M…059-186, right]
Trinil, 16 September 1899. Most distinguished Sir!
I have started staking out a trench of 24 meters in length and 6 meters in width [the 1899 24-m Trench; the long dimension lay in an east-west direction, while the 6-meter width was dug southward into the bank from the former edge of the 1893 40-m Trench (1894 photograph, Figure 4a, main text, and Kriele's April 14, 1898, sketch, -A4f, p. 9, above)]. [A small sketch, South upward, is in the September 1899 letter.]

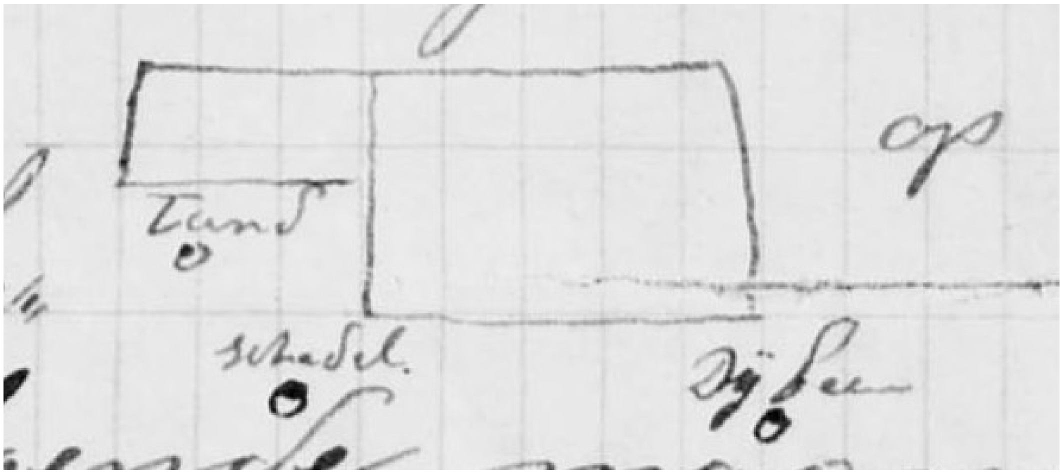


[ tand = tooth    schedel = skull (**Skullcap**)    Dijbeen = Femur (**Femur I**) ]
[The tooth (*tand*) might be the ape-tooth Dubois referred in a note on Kriele's 12-8-1895 letter (-A4k).]

In this manner, I think I will finish the outlined trench early next month. After that, I will leave this and stake out a second trench upstream [farther east]. We have already reached the bone bed [LB-HK] in which so far we have not found anything special. Still, we did find several pieces of bone, for instance an elephant [***Stegodon trigonocephalus***] tusk with a length of about 2 meters in 4 pieces [see Crate 1 '99 shipment, -A4l, below] and a complete upper skull portion of a rhinoceros [***Rhinoceros sondaicus***; Crate 5 '99], as well as some nice small leg bones, vertebrae [Crate 3 '99], deer [***Axis lydekkeri***] antlers [Crate 7 '99] and various teeth and molars [Crate 7 '99]. Would you respected Sir, be willing to request for me to be placed [again] in the Solo valley, since then I would [5] [M…059-187, left] be present during the excavations just as before in case something is found. I would be very happy if you could assure the Chief of the works that I received a nice Certificate from Engineer van Rossum related to the work I did in the past during my time in the Solo valley. Mr. Delprat is eager for me return [to Sumatra], but my wife is very much against it because the work there is quite dangerous and she is now several months pregnant, so that I must be with her and stay in Java. Other than that, no news and both my wife and I are doing fine. In the hope of receiving a favorable message soon, and with great respect, I remain, faithfully yours, Kriele

**S II-A4i.**

[Page] 6 [M…059-188]
Trinil, November 7, 1899.
Most distinguished Sir!
[I] received your letter of October 5, from which I understand that you are thinking of continuing work through the rainy season so that when the dry season approaches we will most surely have exposed an excellent trench. I experienced a slide in the trench of 24m at a depth of about 11 meters [while digging in the LB-HK] and so I have now started one trench upstream [referred to as the 1899 5-m Upstream Pit, as shown in a copy of Kriele's sketch at the end of this letter] and another one downstream [which was 10m long and 6m wide, and is referred to as the 1899 10-m Extension]. As soon as the downstream trench has been dug to the same depth as the one of 24m [that is, to ~11m below the terrace surface], both trenches will be finished at the same time [combining the 1899 10-m Extension and 1899 24-m Trench to form the 1899 34-m Trench]. We have recently found an incomplete skull of a cow [***Bibos palaesondaicus***, see Crate 1 or 5 '99 shipment, -A4l, below] and an assortment of leg bones [see Crate 2 '99], vertebrae [Crate 3 '99], teeth and molars [see Crate 7 '99]. In addition, there are various complete and incomplete deer [***Axis lydekkeri***] antlers [Crate 7 '99]. Also, we found nicely preserved small leg bones as well as a very small jaw with a molar and a small canine tooth [all in Crate 7 '99]. [Page] 7 [M…059-189, left] I have also received notification of a decision made by Buitenzorg that I have to complete my work no later than December or [preferably] earlier, including packing and shipment. So I plan to dispatch this shipment near the end of December from either Kedung Aller [?] or the Paron station. Other than that, we are in very good health and I remain in the happy hope of not working again in the [Sumatran] coal mine, I am happy to be serving you, respected Sir, your humble workman, Kriele.

[The Sketch, below, shows the 1899 24-m Trench and its 10-m Extension. Together they comprise a 34-m Trench, which sat south of 1893 40-m Trench and the 1897 Upstream Pit. East of the 1897 Upstream Pit, the map shows the 1899 5-m Upstream Pit. Refer also to Kriele's April 14, 1898, sketch, -A4f, p. 9, above).]

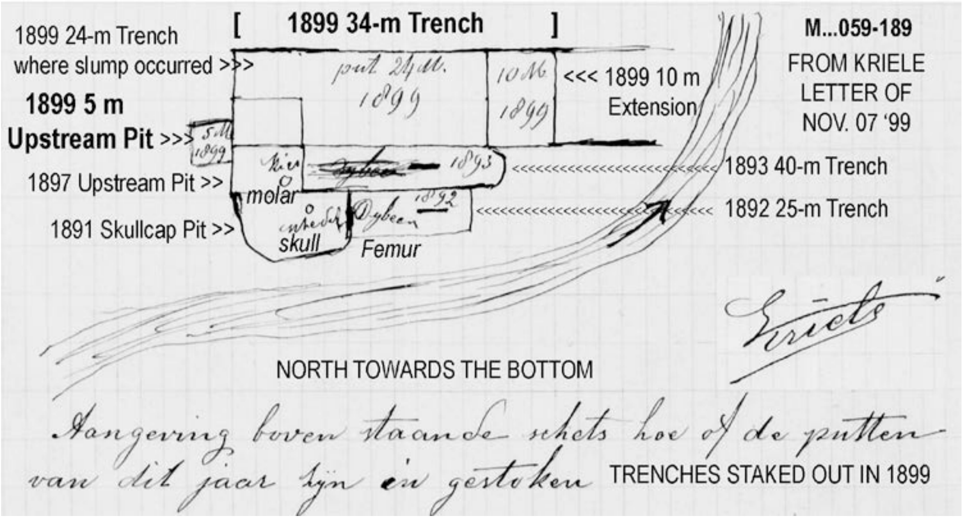


**S II-A4j.**

[Page] 8 [M…059-189, right] Trinil, 8 December 1899. Most distinguished Sir! Please excuse me for having waited so long in writing, but the reason for it is that I was called up by Engineer [*Ir.*] Kruimel to appear at Ngluwak so I could tell him how we would work during the West Monsoon period. I did explain my plan to him in great detail and he fully approved of my explanation. I have not received any word yet about continuation of the work. [In the LB-HK of 1899 34-m Trench (shown in the 1-11-99 sketch, above)] We recently found 1 incomplete buffalo skull [***Bubalus palaesondaicus***; see also, Crate 5 '99, -A4l, below] and 1 incomplete mandible of a tiger [***Panthera tigris***; Crate 7 '99] as well as an elephant tusk [***Stegodon trigonocephalus***; Crate 7 '99] and an assortment of other bones. There was also a molar [*laaste kies* on sketch] of a ape [*aap* in Dutch] or a human ["or a human" was bracketed and crossed out by Dubois on the letter; hence the molar was a non-hominin Catarrhini], as well as a tooth [*tand*] if I am not mistaken [in the margin: "*It turned out to be an ape's tooth- ED,*" likewise a non-hominin Catarrhini (see comment below)]. I have given both of them to Mr. Kruimel with a request to send them to you, which he promised to do. I am not sure if it is indeed a tooth, but it was found about 0.5 meter above the lowest water level and [Page] 9 [M…059-190, left] it is slightly damaged. There is also 1 small jaw with canine tooth and molar, which I also gave to Mr. Kruimel so that he could send it to you. Other than that, all is going as we wish and my health is also good. I wish you and your family a happy New Year, yours faithfully, Kriele

[Kriele provided another small sketch in the margin of the letter, showing the following: The tooth (*tand*) was from the 1899 5-m Upstream Pit ~0.5m above the lowest water, hence in the LB. The molar (*laatste kies*) was from the eastern part of a 24-m section of the 1899 dig. Dubois wrote inside of an oval on the letter that the find was a "*pm*" (lower premolar). Kriele also showed the previous find of a molar (*kies*), presumably the 1897 Premolar, in the 1897 Upstream Pit. One of these teeth might be the isolated molar of a robust silvery langur, *Trachypithecus cristatus robustus* later recognized in the D.C. by Hooijer (1962; also, de Vos and Sondaar, 1982).]

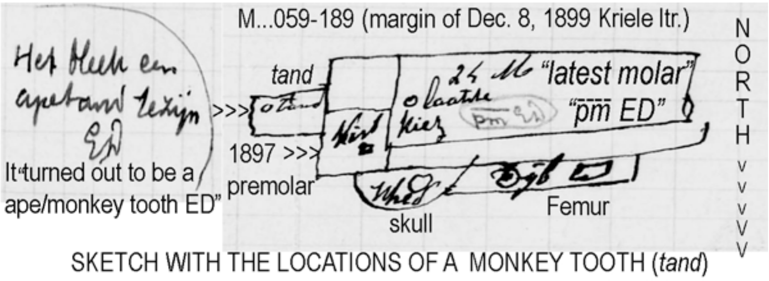


SKETCH WITH THE LOCATIONS OF A MONKEY TOOTH (*tand*)
AND A LOWER PREMOLAR ("*latest molar*") FOUND IN 1899

**SII-A4l.**

[Kriele listed the fossils from the 1899 Trench, and mentioned staking out the 1900 Trench. To judge from Kriele's letters (above) all of the fossils reported came from the 1899 Trench and most from the LB-HK. 1069 finds were listed from Trinil, but only 35 (3%) have information enough to identify the species found.]

I am sending herewith a list of the contents of 7 full crates.

No. 1 An incomplete Elephant skull with complete tusks and molars. A complete tusk of an elephant, in four pieces with (total) length of ~ 2 M [from the LB-HK, see Kriele's September 16, 1899, letter].

No. 2 A complete buffalo skull with horns and molars
An upper arm bone of the (an) elephant
A complete shoulder blade (scapula)
An assortment of 11 broken pieces

No. 3 An incomplete turtle
An incomplete shoulder blade (scapula)
An assortment of 25 vertebrae

No. 4 An assortment of vertebrae (62)

No. 5 An incomplete rhinoceros skull [*R. sondaicus;* from the LB-HK, based on the 09-16-89 letter]
An incomplete buffalo skull

No. 6 An assortment of [broken] pieces, 6 in total

No. 7 A small crate [with] 665 various teeth and molars
A small crate [with] 33 various small jaws
A small crate [with] 40 various small vertebrae, 2 antelope horns, 1 tiger mandible
2 pig tusks and 5 molars.
[12] [M…059-193] 1 small chest with an assortment of 185 diverse specimens;
3 incomplete deer skulls;
1 incomplete antelope skull;
15 deer antlers [at least some ***Axis lydekkeri*** were from the LB-HK, based on Kriele's 9-16-99 letter];
1 elephant tusk in 5 pieces with a total length of 1.10 meter;
3 elephant molars.

The single frontal molar of the Rhinoceros skull [***R. sondaicus***] that is missing is packed in crate Nr. 7 and is lying on top, tied to a rib. This month, we have found an elephant tusk [***Stegodon trigonocephalus***] and an assortment of other pieces. The trenches have been completed, and [because they were dug through the LB-HK near the LWL] from the 15th onward, they have been inundated [since] by water. Since I have not received notice that we must continue working next year, I have decided to keep the prisoner workers here and I have already made a beginning in staking out on top a trench for 1900 and that is progressing excellently. As soon as I receive notice, I will let you know immediately. … I remain in the hope to be able to continue working here, Serving you, most distinguished Sir, your workman Kriele

### SII-A4m.

[The folder with Kriele’s December 21, 1899, letter has the most detailed available map of the left bank (Figure 3a, main text). The plat shows the 1900 excavation, divided into sectors (*vakken* in Dutch; *vak*, singular) (M…059-194; see also, de Vos and Aziz, 1989: 415, Fig. 7). The map must have been drawn after the excavation was mostly complete but before it was final (see -A2q, below).]

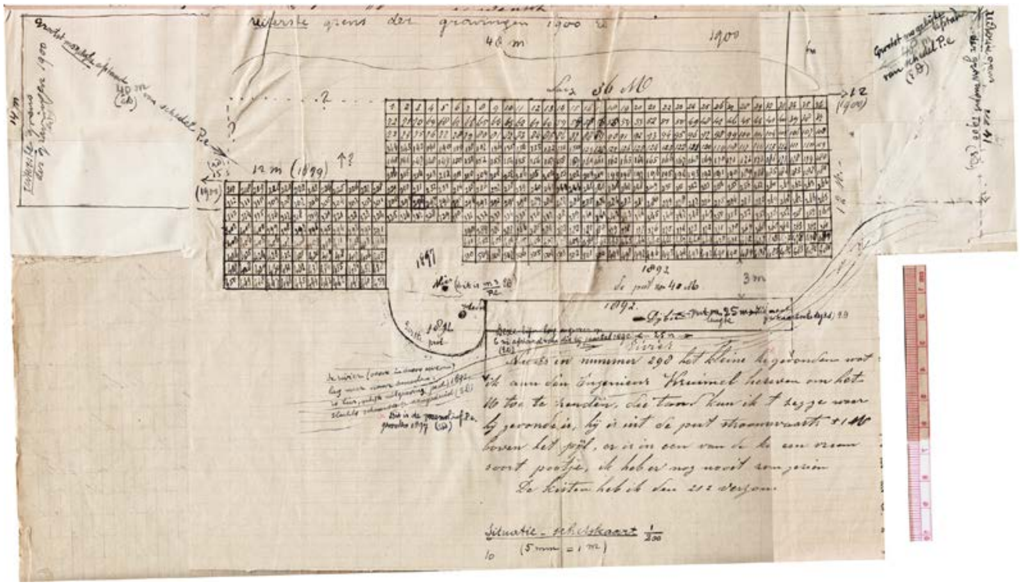

**[SOUTH IS UP. Kriele (GK) and Dubois (ED) annotations on the map are**:]

[Left edge note by Dubois] “14 meters Extreme limit of the excavations 1900 (ED)”

[Diagonal note in the upper left] “Largest possible distance 40m from the skull P.e. (ED)”

[Note by GK along the upper edge of the 1899 area, divided into one-meter squares 1 to 36] “length of 36m” [see –A4i]

[Area of 1899 excavations, divided by GK into about 450 one-meter squares] “1 … 36” [top row]; “72 … 37” [next row down]; “216 … 187” [lowest row in 36 m area]; “217 … 264” [highest row of wider area of excavation]; “313 … 360” [lowest row of the wider area of excavation]; “402 … 361” [highest row where excavation is on both sides of 1897 pit]; “423 … 415” and “451 … 480” [lowest row where excavation is on both sides of the 1897 pit]; “427 … 438” and “450 … 439” [at lower left of the 1899 excavation]. [The grid system must have been created after the 1899 excavations were expanded in November, by which time the original 24x12-m trench was largely completed; the 36-m wide back edge is two meter wider than indicated on Kriele’s 11-07-99 map, -A4i.]

[Note by Dubois at upper left corner of the 1899 Trench] “(ED) 15 [m] (1900)”

[Note by Dubois along upper left edge of 1899 Trench] “12m (1900)”

[Dashed lines above the foregoing notes, added by Dubois] “---- ? -----“

[Line at the upper edge added by Dubois] “Extreme limit of the [proposed] excavations 1900 (ED)”

[Diagonal line on upper right and note by Dubois] “Largest possible distance from skull P.e. (48 meters ED)”

[Note by Dubois at upper right corner of 1899 excavation] “→ 12 [m] (1900)”

[Note by Dubois in upper right edge] “14 meters Extreme limit of the excavations 1900 (ED)”

[Bracket near upper edge, added by Dubois] “48m 1900”

[Annotations in the areas of the 1897, 1891, 1892 and 1893 excavations, drawn by GK]

In the “1891” or “1897” [area], “kies” [molar] by GK with “(this is m3 /P.e. ED)” by Dubois [1891 Molar]

In the “1891” [area], “schedel” [skull; Skullcap] by GK, and “First pit,” and “skull” added by Dubois

In “1892” [area], “Dijbeen” [Thighbone] by GK; “pit of 25 meter in length” “see also the sketch maps that are not to scale ED”

In “1893” [area], “du put van 40m” [Pit of 40m] by GK, and < 3m > as north-south dimension of the 40-m Trench, added by Dubois [correcting the width he gave in 1895, Figure 2b, main text]

[Notes by Dubois along lines representing the river] “This line was located at about a distance of 6 meters from the one that goes with the year 1892 (ED),” and “The river (shore in the dry season) was further down. It is here only indicated schematically, like the excavation (pit) 1892 (ED)” [Long note by GK] Now in number 298, the small[x] [faint pencil] [specimen] was found, which I gave to Ingenieur Kruimel to forward to you [see –A4d ‘4-11-1897’ above]. I cannot say where the tooth was found [not necessarily referring to the “small [specimen”]. It is from the pit down [or up)] stream +/- 1 meter above the [river ? ] level. In one of the crates [?] there is a strange kind of small animal leg. I have never seen one like this before. [Note by ED to the left: **X** [in faint pencil, similar to the X at “small,” above] “This is the pre molar inf P.e. found in 1897 (ED).”

[Scale] Situation Sketch Map 5mm = 1m

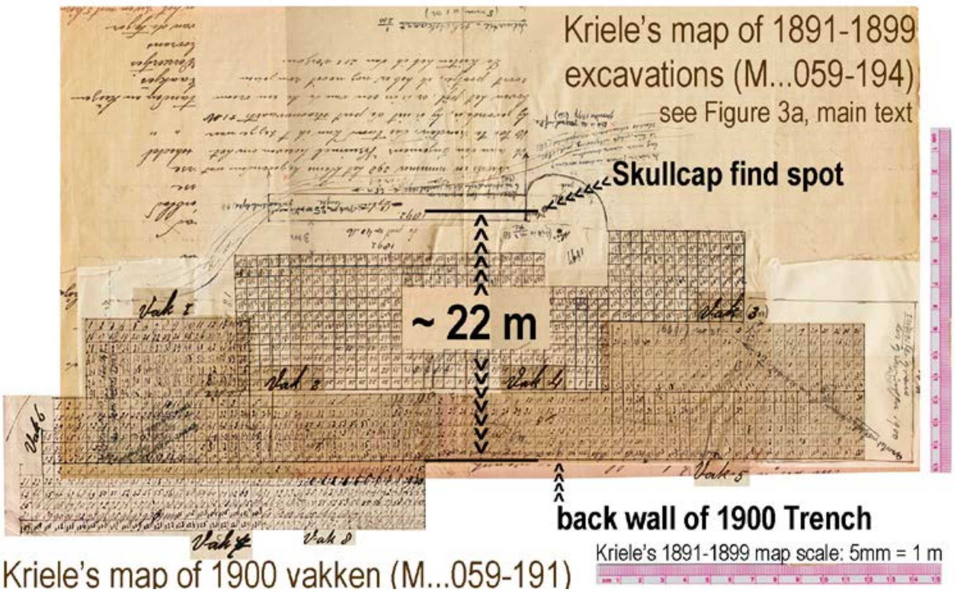


[The folder containing the December 21, 1899, letter includes a Kriele plat of the final 1900 Trench with North facing up (an annotated version is below; the plat is mentioned by Kriele in his December 28, 1900, letter, -A4o). The 1900 Trench was divided into sectors (*vakken* in Dutch; *vak*, singular). Kriele's mapping must have done after most of it had been dug but before completion (see -A2q, below). When his mapping of the 1900 Trench is overlain (as best as possible) on the sketch from 1899 (previous page), the Skullcap discovery point falls ~22m north of the back wall of the Trench, as illustrated above. This positioning was essentially Dubois' conclusion. He annotated Kriele's December 21, 1899, map to indicate that the southwest and southeast corners of the proposed 1900 excavation were expected to be (or had been) 48m and 40m from the Skullcap discovery point, respectively. He noted along the east side of the proposed dig that 1899 unit 313 was 14m north of the proposed 1900 back edge. These dimensions place the Skullcap find spot 16m from the closest portion of the 1900 vak 3 and 22m from the proposed 1900 back wall.]

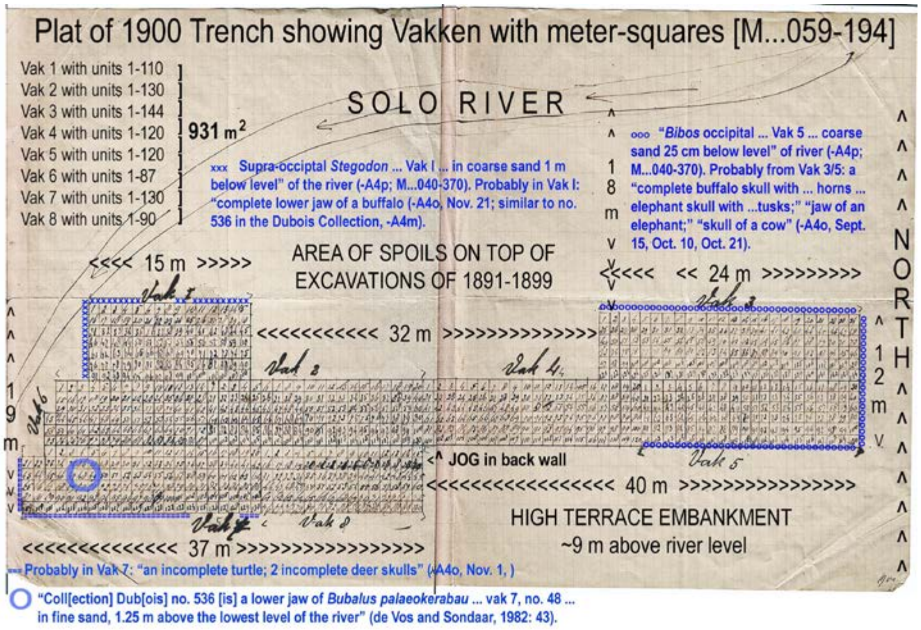


[See -A4r, below, for an assembly of Dubois 1891-1900 excavations, based on Kriele's maps.]

**S II-A4n.**

[With regard to the 1900 excavations, De Vos and Sondaar (1982: 43) state "The excavations lasted from January until May and the assistance was given by 25 local laborers; and from May until November 50 local labourers were employed. …. A very large excavation was made, measuring 75 by 6 to 14m (see Kriele's plat in -A4m, above). It surrounded on three sides all the previous excavations on the left bank …. (Figure 3a, main text; S II–A4l and –F10). De Vos and Aziz, 1989: 415, make further reference to "Dubois, 1932: 719" and state: "Dubois added the outline of the 1900 excavations to a sketch by Kriele of the 1899 trench," as shown in their Fig. 7, and "from this figure we can deduce that the surface of the trench was about 916 $m^2$ (consisting 931 units in -A4m). In November 1900, three photographs (see photographs 2 and 3) were taken of the excavations on the left bank (Kriele, 12-11-1900; the positions from where these were taken, are indicated in Figure 8. The excavations lasted eleven months."

De Vos and Sondaar (1982: 43) further recounted the recognition during the 1930s of new *Homo erectus* femora from among Trinil fossils found in 1900: "In this large pit four incomplete femora of *Pithecanthropus erectus* were obtained (femora II, III, IV and V). The exact spots where they were collected is not known, as at the time they were not recognized as being of any importance. In the Dubois Collection at Leiden they were found among fragments of ribs and other fossils of little value (Dubois, 1932: 719). …. The [1900] trench is divided into eight sections [*vak* in Dutch] each of which was divided by a square-meter grid sectors. That Kriele aimed at providing the fossils with detailed labels is shown, i.e., by Coll. Dub, no. 536, a lower jaw of ***Bubalus palaeokerabau*** Dubois, on which a label was stuck with the information_'vak 7, no. 48, vijn {properly fijn} zand, 1.25m boven het laagste pijl' (i.e., the specimen was found in section 7, square 48, in fine sand, 1.2 m above the lowest level of the river [shown on plat in -A4m; also, Kriele letter of Nov. 21, below].... In November 1900, three photographs [five images from three camera stations] were taken of the excavations on the left bank (Kriele 21-11-1900) …." (S I Figure 7a).]

[Dubois (1926a,b, 1932, 1934, 1935) published four papers on the femora from his 1900 excavations. The two excerpts given here are the parts that focus on the provenience of the specimens:]

> (i) [Dubois, 1932: 716-719, written in English]: Forty years ago the two principal skeletal remains, the skullcap and the femur, of *Pithecanthropus erectus* were excavated at Trinil …. [and] possibly organically associated … in that they were found in exactly the same layer of fluvial deposit of volcanic ashes, the one from the other at only about 12 m distance. …
>
> This, assuming the two skeletal remains were organically associated, implies the supposition that in the anthropogenesis the acquisition of the human, exclusively upright gait preceded the evolution of the brain and skull …. [see –B4b, below, for Dubois' first expression of this conclusion, 40 years earlier]
>
> Now, as to know a species well, one single individual is insufficient …. Will another femur present the same differentiating character? ….
>
> The answer … has been long in coming …. In these forty years no remains of another pithecanthrope came to light [from Trinil] …… till the first day of this month [June 1932] eight overlooked pieces of three thigh bones of the described species.
>
> On that day, at Leiden, in my Java collection, from a lot of inconsiderable fragments of ribs from different Trinil mammals … were separated some dissimilar fragments not belonging to ribs. Amongst them a bone a foot long, still partially covered with rock, which my diligent assistant in the arranging of the Collection, Dr. [J.J.A.] Bernsen … had put aside for my inspection …. Then, searching further between the rib fragments for similar pieces, that might possibly fit, I found the defective upper extremity to that thigh bone shaft, … and 6 other pieces of different sizes, which enabled me to compose two more shafts of *Pithecanthropus* thigh bones. …
>
> From the first and the least incomplete bone [**Femur II**] I have now removed nearly the whole of the hard strongly adhering rock, a pyritous impregnation of andesite [mostly likely andesitic sandstone with authigenic pyrite]. The two other bones [**Femur III** and **IV**] were nearly bare of rock but more broken.

The surface of the three new femora is more of less strongly corroded … most so III and IV, the least II …. The four thigh bones [found in 1892 and 1900] are from four different individuals …. II and IV are from the right side, I and III from the left side….

The new femora of *Pithecanthropus erectus* were found in a collection of fossil bones acquired from excavations executed at Trinil in 1900,****** conducted under my direction by my former assistant in Java, the later sergeant-engineer Kriele. The collection was sent to Leyden in the same year …. The exact site of [discovery of] these fossil remains in the [1900] excavation is not known, because they were not remarked as uncommonly important and mixed up with fragments of different ribs and some other fossils of small value. It is, however, possible to ascertain that the distance of the three new thigh bones from the site of the skullcap of 1891 was between 16 and 48 m. [see –A4m, above; in giving 16 m, Dubois seems to have been referring to the distance to the closest part of vak 3] Now considering that the skullcap and femur I were found at only about 12 m the one from the other (perhaps only 10 m, certainly not 15 m, as appears from my notes [publications] at the time, in a part of the excavations where corrosion of the bones was generally less {footnote: Femur I is quite unaffected by corrosion, the skullcap yet less than the new femora}, whereas the surface of the three new thigh bones is intensively corroded, considering, further, that very likely the skullcap and the femur I are both feminine, it appears very probably that none of the three new thigh bones was individually associated with the skullcap."

(ii) [Dubois (1934: 139-140)]: In the meeting of June 25, 1932, I exhibited and briefly described three incomplete new femora ([designated] II, III and IV), which, on the first day of that month, I had recognized amongst a great many seemingly more or less worthless fragments of ribs of large Trinil mammals, in my Java collection…. on December 15, 1932, I recognized, in the same collection, a fifth thigh-bone of *Pithecanthropus* …. One having a length of 20 ½ cm [which] clearly exhibits the extraordinary internal structure of the thigh-bone…. Concerning the provenience of this specimen I may remark, that, the same as two fragments of the first mentioned new thigh-bones [**Femur II** and **IV**], this one bears the inscription *Trinil*, in the hand-writing of my former technical assistant at the explorations and excavations, the late sergeant-sapper G. Kriele [footnote states in part, 'I did not recognize the femur fragments in the year 1900']. Presumably he signed these three fragments, because he suspected them more than the multitude of the 'more or less straight and nearly cylindrical fragments, resembling the pithecanthropus thigh-bone also in thickness,' which according to my request [was] sent along with a good collection of unquestionably valuable fossils from his Trinil excavations in the year 1900, in order that I should examine them and select what might indeed be parts of those much desired limb bones.

[Kriele' lists transmitting the crates carrying the 1900 finds do not mention a box containing ribs, nor does the partial list that possibly was done in The Netherlands at the time of the shipment's arrival (-A4p, below).]

**<u>S II-A4o.</u>**

[M…059-195 = Label on a bundle of Kriele letters from 1900]

(i) [page 5] [M…059-197] ['*Kriele 1900'* written in the left margin] [written in the right margin*: 'ED: This decision was a result of the Fourth International Zoological Congress (Cambridge, August 1898)* which had asked the Netherlands Government to continue the excavations at Trinil'*] Trinil, January 10, 1900. Most distinguished Sir! I have the honor of letting you know that I have received a decision from Batavia to continue my work this year with 25 men from January 1st through May and with 50 men during the remaining months. I hope now that we may find once again something very nice [a hominin fossil]. I have staked out a sizeable trench upstream as well as one downstream where I am now busy dumping the [easily dug] top ground into the river. The middle sector [*vak*] will come later since the soil [along the terrace embankment] in that sector can be washed away with sawah water [drainage] after the river flow decreases and the rice fields are flowering. Further no news and all is very well with my health and I have great hope for the future. With great respect for you, distinguished Sir, your humble servant, Kriele

[*see the account in Shipman, 2001: 305-316, especially the resolution described on pages 315-316. See also http://biodiversitylibrary.org/item/76163#page/14/mode/1up Dubois is listed as attending (Dubois, E., Sweelinckplein, 12, 's Gravenhage. Jesus College, p. 15) (p. 32) Friday, August 26th, General Meeting of the Congress to hear an address by Professor Haeckel, "On our present knowledge of the Descent of Man." Etc.]

(ii) [page 6] [M…059-198] Trinil, January 31 1900. Most distinguished Sir! The work is progressing very well, but we are not finding bones. Upstream I have staked out a trench of 12 x 6 meters, which down below will be enlarged by 6 x 6 [meters, becoming a 18x6m, 108 $m^2$, eastern section]. Also a trench downstream of 10 x 5 meters [a 50$m^2$ western section] and one trench in between which measures ~ 50 meters in length and 6 meters wide [a 300$m^2$ middle section]. Down below we will thus get a large area of more than 400 [square] meters [that is, an excavation of 78x6 meters, 468$m^2$ at the LB-HK level] and my hope is that I will finish it and find something worthwhile [that is, a hominin fossil]. … [7] [M…059-199, left] … Besides that, I can't complain because I was allowed to have a house built for myself along the shore of our work. Our work is moving along very well. However if you, distinguished Sir, would have some medications for me that would be really useful right now, because traveling constantly to Ngawi generally cost me money. My wife is doing very well and so I remain in the hope that I may soon recover and remain with great respect for you, distinguished Sir, your humble workman, Kriele

[Note at bottom] Wrote to Kriele on 25 March 1900: smoke colored glasses; dig around former trenches; photo from the monument pillar; and from really close by (or [from the] opposite side); in September or October. ED [This includes instructions about the locating the camera stations from which the 1900 photographs should be taken.]

(iii) [Page] 8 [M…059-199, right] Trinil, 22 February 1900. Most learned Sir! I have the pleasure of letting you know that the work is moving along very well. We have now worked down to a depth of more than 2 meters across the entire sector [*vak* in Dutch; evidently referring to the 78x6 [m] area described in the January 31 letter]. There are no bones in this [upper portion of the embankment below the terrace surface] except for some insignificant pieces. My headache is still just as bad as before. I couldn't sleep during several nights and that went along with fever [the causes were not explained]. I am still being treated by a doctor but it does not indicate a cause of the headache. Otherwise all is well. Serving you, most learned Sir, your humble workman, Kriele

(iv) [Page] 9 [M…059-200, right] Trinil 17 March 1900. Most learned Sir! The work is progressing very well. I have widened the trench upstream by about 6 meters and I will also have the trench of 50x6 meters widened to a width of 10 meters if the water level stays where it is for another two months [the excavation was 50x10 m, 500 $m^2$]. In my opinion we can likely complete this work with 50 coolies per day. My headaches are still the same and I am still being treated by a doctor. I have not learned what the cause is but it is extremely debilitating. Otherwise all is going as well as we can wish. With great respect for you, most learned Sir, your humble servant, Kriele

(v) [Page] 10 [M…059-201] Trinil 6 April 1900. Most distinguished Sir! The work activities here are progressing very well. The upstream pit [? the 18x6 m eastern section] has been dug 6 meters deeper. For the entire work site [evidently referring to >78 by 6 m or >468 $m^2$] we have now reached an average total depth of around 4 meters. We have not yet extracted any bones from this [top four meters of rock below the top surface of the terrace embankment], except for some insignificant specimens. It appears that the rain showers are now diminishing and the water level is currently relatively low. My headache has not improved much and I have been treated by a doctor now already for 3 months. Other than that, esteemed Sir all is going as we wish. Yours Faithfully, Kriele

(vi) [Page] 11 [M…059-202, right, -203, left] Trinil, 28 April 1900. Most distinguished Sir! I received your letter dated 25 March [1900] in good order. My work is progressing smoothly and, if the river level continues to fall at the present rate, I think I will get to the bone bed in a portion of my trench by next month. Downstream, the trenches have been staked out as indicated in the sketch below [on the next page, showing the 1900 Trench was 75m long and 6-16m wide]. It would be very difficult to uncover the trenches of the years 1891, 1892 and 1893, unless we were given a lot of time, since at least 4 to 5 meters of sediment is covering them plus whatever [sediment] is inside the trenches. The water would be an additional problem since in the past we removed the small dikes as much as possible. I will uncover the 1898 [sic; 1898 was mistakenly overwritten on the correct year of 1897] trench together with the one of 1899 and then check to see if anything has been left standing. Other than that, as you can see from the enclosed sketch, the latest trench that I staked out is about where you had indicated it should be. I will try to work in all trenches as much as possible to the depth of the clay [stone] bed.

About your request for photographic work [in you March 25 letter, above], it would be best that you ask this yourself from Mr. Kruimel, since I cannot speak to him in person and my writing could possibly be misinterpreted. If you receive a positive response, it would be helpful to alert me in advance so that I can adjust the work activities to have a good spot for the photographer. There is no local photographer here in Ngawi and I don't know if there is one in Madiun, but I will inquire about it and if indeed there is one available, should I then arrange for him to come if you couldn't get it done through Mr. Kruimel? …. [Page] 13 [M..059-203, right] ….

My wife is doing quite well and she is always happy, but she still fears that we will have to return to Sumatra after the work here has been completed. She is afraid of the coal mine just as much as I am since it is unhealthy and extremely dangerous work. During the 4 months that I worked there, I had just about 3 different serious accidents one of which was a water rupture [?] due to a fall and another time I got buried under the coal. So I hope that you will request a different assignment for me since I am afraid that Mr. Delprat will become angry because he cannot retain people for that work. I remain in great hope of your help in getting me a different working environment and that you will let me know before the work that I am doing now for you has ended, Your humble servant, Kriele

[Sketch of the 1900 Trench included with the foregoing April 28, 1900, letter is on the next page.]

(vii) [Page] 22 [M…059-212] Trinil August 24 1900. Most learned Sir! We have had some days of delay due to the fact that the trench became inundated at high water levels and caused large amount of mud. The water level rose so quickly that within ~3 hours my work site filled up completely and that lasted so for 4 days (while the total increase in water level was ~5 meter). After that it slowly subsided again so that tomorrow I can again resume work in the old trench and we can clean it up. We have not had so much rain since the latter part of last month and the beginning of this one. We have not found many bones this month because we had to work on top most of the time. Otherwise all is going as wished with the work activities and also with my wife and my health. With great respect towards you, most learned Sir, your very humble workman, Kriele

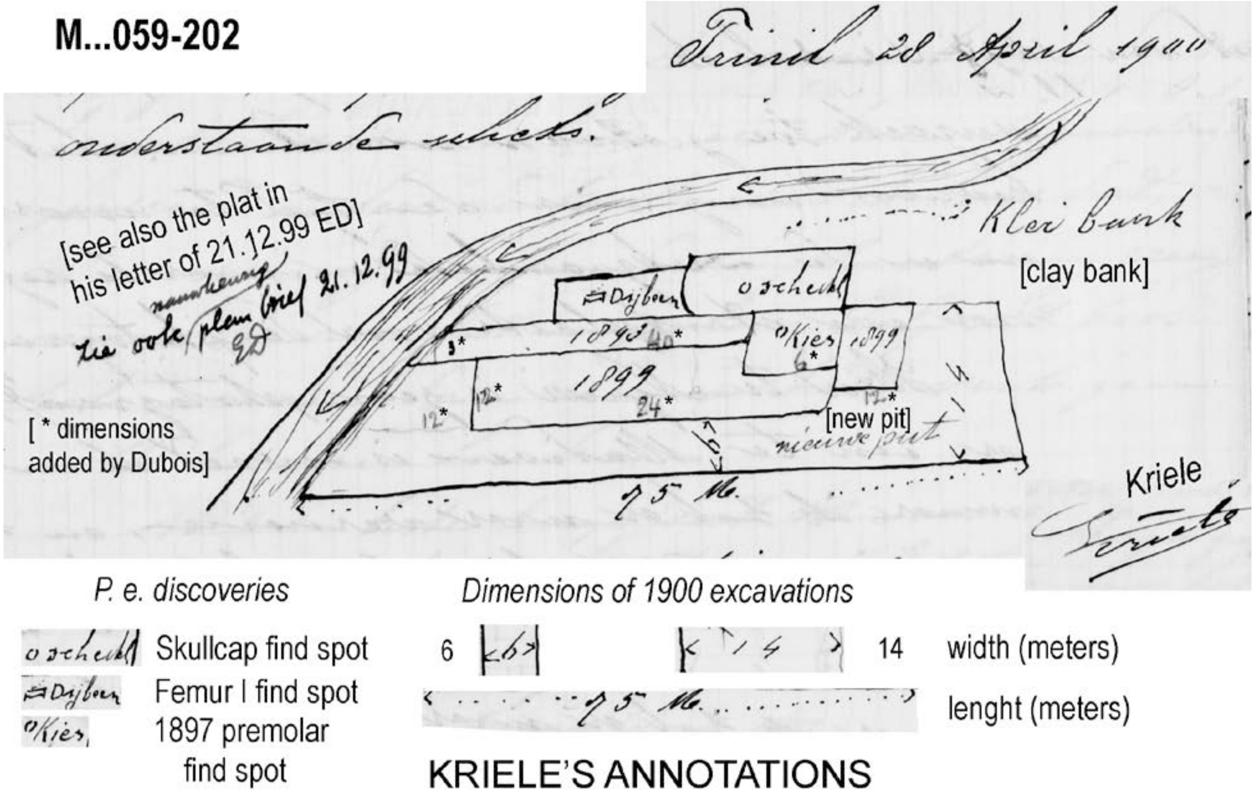


[This map, which is oriented with North up and has been annotated and reorganized here for further clarity, shows the 1900 Trench long before it reached the LB-HK level (April 6 letter, above). Judging from Kriele's account, the scars of the 1899 Trench, 1897 Upstream Pit, 40-m Trench, 25-m Trench, and the Skullcap Pit, all of which are shown in his, might have been covered by spoils and river sediment (specifically, in his 28 April 1900 letter, he noted a "4 to 5 meters … covering" of the 1891-1893 pit and trenches). In November Kriele added a 23m extension onto the southwestern end of the 1900 Trench (described below). See -A4r, below, for an assembly of the 1891-1900 excavations, based on Kriele's maps.]

(ix) [Page] 23 [M…059-213] Trinil, September 15 1900. Most learned Sir! We have excavated the trench of ~ 25 meters in length to a depth where no longer bones are present [presumably because the LB-HK had been fully penetrated; the 25-m trench was probably 24-m long vak 3 shown in –A4m, above]. In this trench, we found: a complete buffalo skull with complete horns and molars [***Bubalus palaeokerabau***]; a complete elephant skull with complete tusks and molars [***Stegodon trigonocephalus***] and further an assortment of specimens. I regret to write that nothing has yet been found of the ape-human [including the femora Dubois recognized in the early 1930s]. Since we are slowly approaching the end of the year, I would very much like to know what kind of work I may expect, or would you, distinguished Sir, think that another year's extension may be granted? [I am asking] because it saddens me not to have any certainty and I would object very much to return to the coal mines because of the accidents and unhealthy conditions that go along with it [Page] 24 [M…059-214] The thin slanted beds [cross lamination sets] about which you, distinguished Sir, asked me have been observed in 4 different places. They all dip approximately in this indicated direction S.W. ////////// N.E.  but also N.N.E. … In the great hope of receiving an encouraging response, I remain as always, with great respect for you, most learned Sir! Your humble workman, Kriele

(x) [Page] 25 [M…059-215] Trinil, October 10 1900 Most learned Sir! Since I have suffered from severe illness again for several days, I was unable to answer your writing dated August 12. Since I now feel a little bit better, I consider it my duty to answer you as to who has visited me during this time. I had a visit from Prof. Boehm, who observed all the work here and I also accompanied him to … where the sea shells occur [perhaps the Sonde locality]. … The professor also asked for a [26] [M…059-216, left] collection of those sea shells that occur there, which I have already instructed my coolies to search for on his behalf. But if you find it not permissible I will not hand them over to him. In regard to the work activities all is going very well. We are now working in the last trench upstream and very close to the spot where recently the molar of the ape-human was found [1897 Premolar; Vak 3, 4 or 5 on Kriele's plat of the 1900 excavations, -*A4m*, above]. But we have not yet found anything of him [more of the *Pithecanthropus erectus* skeleton]. We did find an incomplete turtle; a complete lower jaw of an elephant (***Stegodon*** [***trigonocephalus***]) with molars [possibly Dubois Collection no. 1657] and some assorted bones. I am also very much looking forward to a visit by the photographer [who took the 1900 photographs in November], since I have to stake out a new trench on top within 14 days and that would lead to filling in of the old trench which would spoil the photographs. I also read in your letter that this will be the last year that we work here [Dubois had decided by August 12 to end the Trinil excavations] and that you would consult with Mr. Delprat about my work assignment. I would be extremely thankful to you [Page] 27 [M…059-216, right] if I could be assigned to a location other than the mine, for instance as supervisor at the Railways, it does not matter where else. With great respect for you, most learned Sir, I remain Your hard working workman, Kriele

(xi) [Page] 28 [M…059-217] Trinil October 21 1900. Most learned Sir! Yesterday, I received your letter of September 12 in good order. I have not heard a thing from these two professors about which you wrote. If they would arrive here I would handle it entirely in the manner that you wish. And if the two gentlemen by chance would like to continue excavating next year and if they would make me an offer to work for them if your work were to be finished, I still wouldn't do it if it were against your wishes. This is also in view of my job which I hope will be permanent and work for these two gentlemen would be only temporary. So I like to ask a very friendly request from you to let me know yet before the work has been completed and how things will go then with my work after it ends. If Professor G. Boehm had not had a letter of recommendation from Mr. Delprat, I would in that case not have told him anything. Incidentally he also did not come to [Page] 29 [M…059-218] get bones but sea shells. The way he presented himself, I told him and the control person that I am not allowed to do it. By the way, he was here only half a day that is to say at Padas Malem [Padasmalang] and Sundeh [Sonde] where the shells occur. During the last few days we found two incomplete turtles [**Testudinoidea**] and an assortment of several leg bones, an elephant molar [***Stegodon trigonocephalus***] and an incomplete skull of a cow [***Bibos palaesondaicus***]. These were all found in the upstream trench immediately next to where last year the two molars [a molar and a tooth] of the ape-human were found. That trench is as good as finished and I am still waiting for the photographer before I finish the small dam in between [as seen in the photograph of Figure 4b, main text; possibly Vak 5 on, -A4m, above]. If he does not show up this month, I will start on that before it is too late. Otherwise all is going well with the work activities as well as with my health. In anticipation of favorable news from you, I remain as always, to you most learned Sir, Your very humble workman, Kriele

(xii) [Page] 30 [M…059-219, left] Trinil November 1 1900. Most learned Sir! Up till now I have not heard from the two gentlemen professors [of undetermined identity]. I also have not heard a thing from the photographer. I did hear from the Chief Engineer Kruimel that he was transferred to Tegal but I don't know who replaced him as the new Chief for that sad work. I did hear that the Solo Valley [project] will be completely halted which is really too bad for a lot of folks. I had hoped myself for a continuation of that work. The large trench has been completed except for a few portions upstream. We have staked out a new trench downstream with a length of 23 meter and a width of 6.5 meters [probably the 22x6-m Vak 7 on -A4m, above] and that one has already been excavated to a depth of ~ 7 meters [it was a western extension to the 75-m trench dug; see Figure 4a, main text]. We recently found [presumably from the LB-HK in other portions of the 1900 Trench] an incomplete [Page] 31 [M…059-219, right] turtle [**Testudinoidea**]; 2 incomplete [***Axis lydekkeri***] deer skulls with several horns and some assorted leg bones, vertebrae, ribs, teeth, molars and small jaws. I regret to say that we have not been able to get anything of the ape-human [including the femora Dubois recognized in the early 1930s]. Further, all is going well with the work activities and also with my wife and my health. In the hope to still find something really attractive, I remain as ever to you most learned Sir, Your willing and humble workman, Kriele

(xiii) [Page] 32 [M…059-220] Trinil November 21 1900. Most learned Sir! Several days ago I was visited by the photographer and he has taken photos of the work from three different stand points [1900 site photographs] with indications of the spots where the skull, femur and molar have been found [P1, P2 and P3 in Figures 4b and 4c, and S I Figure 7; the surveying for the 1900 site map has the camera stations, and therefore was underway in late November or done after that time]. The work has also been documented in profile and also even the small monument pillar and in a square around the work [the profile does not appear to be in the Dubois Collection]. I also received a message from the new chief of the Solo Valley [project] Engineer Elenbaas that if the two professors would arrive here, I should immediately send a telegram to him so that an engineer of the Solo Valley [project] could come over. But I have not heard a thing from those two gentlemen. I regret to inform you that the work is not progressing very well this month, since ten days ago my trench got inundated after a heavy banjer [flood] [Page] 33 [M…059-221] and the water has steadily remained at the same level. I have now staked out a new trench of 15x6 meters so that if the water subsides soon we can still finish that [Vak I on the plat, -A4m, an area previously covered by spoils, as seen in 1900 photographs]. We have not found many bones lately, just a complete lower jaw of a buffalo [***Bubalus palaeokerabau***; see –A4m and –A4n] and several assorted leg bones, phalangeal bones (etc) as well as teeth and molars. Otherwise, all is going well with the work …. Your most humble workman, Kriele

**S II-A4o.**

(i) [Page] 186 [M…040-373] [written in December 1900 by Kriele] [our inferred species names are often omitted below]

Copy. “Bevloeiingswerken in de “Solo vallei” [handwritten Stationary head]

Complete content of 16 crates filled with fossils. [*dimensions added below from M…040-375*]

Crate 1. 1 complete elephant skull with molars and tusks [*Length 1.12 M, Width 0.80 M, Height 0.69 M*]
Crate 2. 1 incomplete elephant skull with molars and tusks [*Length 0.98 M, Width 0.70 M, Height 0.62 M*]
Crate 3. 2 incomplete turtles; a skull of a cow; 18 assorted specimens [bones] [*Length 0.90 M, Width 0.84 M, Ht 0.56 M*]
Crate 4. 45 assorted specimens [*Length 0.71 M, Width 0.54 M, Height 0.60*]
Crate 5. 58 assorted specimens [*Length 0.70 M, Width 0.62 M, Height 0.54 M*]
Crate 6. 36 assorted specimens [*Length 0.89 M, Width 0.79 M, Height 0.40 M*]
Crate 7. 2 incomplete buffalo skulls [*Length 0.78 M, Width 0.64 M, Height 0.59 M*]
Crate 8. 59 assorted specimens [*Length 0.93 M, Width 0.79 M, Height 0.43 M*]
Crate 9. 35 assorted leg bones [*Length 0.85 M, Width 0.62 M, Height 0.50 M*]
Crate 10. 53 assorted specimens [*Length 0.72 M, Width 0.65 M, Height 0.43 M*]
Crate 11. 45 assorted specimens [*Length 0.80 M, Width 0.51 M, Height 0.50 M*]
Crate 12. 45 complete and incomplete deer antlers and 59 vertebrae. [*Length 0.80 M, Width 0.66 M, Height 0.35 M*]

Crate 13. 1 complete buffalo skull with horns and molars; 2 elephant leg bones; 1 complete lower jaw of an elephant; 1 small chest with 51 molars of buffalos and cows; 1 small chest with 192 deer molars; 1 small chest with 140 deer molars; 1 small chest with 39 assorted small jaws; 1 small chest with 25 assorted small vertebrae; 1 small chest with 24 assorted small leg bones; 1 small chest with 37 alligator (caiman) and crocodile teeth; 1 small chest with 62 assorted specimens. [*Length 1.36 M, Width 0.86 M, height 0.60 M*]

Crate 14. 1 complete skull of a cow with horns; 2 complete elephant leg bones; 2 complete elephant molars; 14 assorted specimens [*Length 0.96 M, Width 0.72 M, Height 0.56 M*]

Crate 15. 1 small chest containing an incomplete antelope skull with 2 horns; [187] [M…040-374] 1 small chest containing a complete pig skull with lower jaw; 1 small chest with 2 deer skulls; 1 small chest with 3 deer skulls; 3 elephant molars; 2 elephant tusks; 1 incomplete buffalo skull; 17 assorted specimens. [*Length 0.85 M, Width 0.72 M, Height 0.55 M*]

Crate 16. 1 incomplete lower jaw of an elephant; 1 small chest with assorted teeth and molars and several assorted specimens. [*Length 0.98 M, Width 0.71 M, Height 0.46 M*]

Trinil, 31 December 1900. The temporary mining foreman (signed by) Kriele.

For identical copy: the 2nd commissary (signed by) illegible. [signed for copying] P.C. Blommendaal

(ii) [188] [M…040-375] [handwritten Stationary Head]

Bevloeiingswerken in de "Solo valley".

Copy. Contents in Cubic meter and Gross weight [KG] of 16 crates filled with fossils destined for the National Geological Mineralogical Mueum in Leiden.

[details on crate sizes included with Crate list above]

Trinil December – 1900
The temporary mining foreman (signed by ) Kriele

For identical copy, The 2nd Commissary (signed by) [illegible] [signed for copying] P.C. Blommendaal

[180] [M…040-368] **1900** [The handwriting of the entries is not that of Dubois, but the corrections made to the list are probably his. The details on the provenience of various fossils evidently came from information written by Kriele on tags attached to the fossils. The list does not exclusively refer to fossils found at Trinil in 1900. Many entries are attributed to Tawang (~3km west of Trinil on the Solo River), Tinggang (far downstream of Trinil), elsewhere in the Solo valley, Patiayam and Kendungbrubus, where no fossil collecting was known to have occurred in 1900. These specimens might have been found during previous years. Specimens from Trinil are underlined]

(1) lower jaw of a young *Stegodon* with 4 molars [***Stegodon trigonocephalus***] (Trinil); banteng horn; incomplete stegodon skull (Trinil) with 2 molars and fragments of tusks [***Stegodon trigonocephalus***] [Kriele's list has '1 complete elephant skull with molars and tusks' in Crate 1]

(2) piece of crocodile jaw without teeth; banteng horns 1.5 m above lowest level, fine yellow sandstone; lower jaw Banteng; small deer [***Axis lydekkeri***] skull without horns Trinil (fine blue sand at 1 to 1.5 m below low water level) [***Stegodon trigonocephalus***]; bones of extremities (in fragments); fragmented jaw; deer antlers and skull pieces; stegodon skull Trinil with 2 molars and small fragments of tusks [***Stegodon trigonocephalus***]; metacarpal bone (Trinil) in fine blue sandstone 1 to 1.5 m below lowest level; lowest portion of a tibia (Tawang), coarse blue sandstone; 4 pieces of tibia and a piece of rib as well as occipital deer skull portion [***Axis lydekkeri***] and a piece of vertebra and bone fragments (all from Trinil), fine blue sandstone 1 to 1.5 m below lowest level Upper leg bone deer at Tawang Picket 135 along the river; stegodon molar 8½ pl [plooien = folds?] in 3D Tawang, 1.5 M above the lowest level; pieces of bone extremities (Tawang) along the river; buffalo horn Tawang 1.5 M above

the lowest level in fine yellow sandstone; pieces of turtle [**Testudinoidea**] from Trinil in fine blue sandstone 1 to 1.5 M below lowest level; stegodon molar 6 ½ pl [folds?] Tawang in coarse blue sandstone; fragmentary tusk; heel bone and vertebra; piece of deer [***Axis lydekkeri***] skull portion (Trinil); complete and incomplete deer antlers Trinil; right half of a stegodon lower jaw with 2 molars (Trinil) [***Stegodon trigonocephalus***] [Kriele has '1 incomplete elephant skull with molars and tusks' in Crate 2. M…040-371 below]

(3) banteng horns; lower jaw young stegodon with 4 molars [***Stegodon trigonocephalus***] No. 62 fine sand layer horizontal [even with] the lowest level … downstream; turtles Trinil [*Chitra* or *Batagur*]; complete buffalo skull with 1 horn; [***Axis lydekkeri***] deer antlers Trinil; ribs; hip bone; deer antler hole (pit); shells and small shells Nqalan; ***Bibos*** [***palaesondaicus***] occipital half transition banteng to *Leptobos* (Trinil) Compartment [*Vak*] 5. 157 coarse sand 25 cm below level; 2 nearly complete elephant tusks attached to a skull Trinil (the right tip is missing) [***Stegodon trigonocephalus***] [Kriele has '2 incomplete turtles; a skull of a cow; 18 assorted specimens' in Crate 3]

(4) turtle; tibia Solo valley sandstone at depth of 2.80 M; extremities bones; 2 pieces of upper jaw with molars of *Stegodon* above (?) Tinggang; 3 milk [deciduous] molars of a stegodon Wadogan, East of Pandan; lower jaw near Tinggang village; tibia Wonogari Picket 135 depth 8 meter [Kriele list has '45 assorted specimens' in Crate 4.]

[181] [M…040-369, left]

(5) vertebrae of large animals; stegodon skull with 2 molars (Trinil) [***Stegodon trigonocephalus***] [Kriele has '58 assorted specimens' in Crate 5]

(6) terrestrial turtle [probably **Testudinoidea**] Trinil, fragment of abdominal shield; tibia *Elephas* [***Stegodon trigonocephalus***] Trinil in fine red sandstone 4 to 5 M above highest level; small knuckle bones buffalo (Tawang) along the river Nganjar; pieces of extremities bones of large animals (metacarpal of buffalo with hole caused by crocodile tooth); piece of lower jaw *Elephas* with 1 fragmented molar; fragment of *Elephas* upper jaw with 2 fragmented molars; vertebrae of large animals; molar of young stegodon. [Kriele's list has '36 assorted specimens' in Crate 6. M…040-371, below, records '(6) tibia elephant [***Stegodon trigonocephalus***] (Trinil) fine red sandstone 4 to 5 M above the highest level.']

(7) complete banteng skull with horns in pieces; incomplete banteng skull; lower jaw *Elephas*; piece of turtle; turtle. [Kriele's list has '2 incomplete buffalo skulls' in Crate 7.]

(8) Elephas molar; scapula and bones of ruminants; deer antlers; small [***Axis lydekkeri***] deer skull partially in stone Trinil even with lowest level in coarse sand layer nos. 79-94-81-93. [repeated on M...040-372] [Kriele's list has '59 assorted specimens' in Crate 8.]]

(9) scapula; deer antler hollow (pit); large tusk in pieces; Elephas tusk. [Kriele lists '35 assorted leg bones' in Crate 9.]

(10) Turtle [**Testudinoidea**] (Trinil); scapula; deer antlers; vertebrae of large animals; forward portion of a rhinoceros lower jaw with 4 molars. [Kriele's list has '53 assorted specimens' in Crate 10.]

(11) 2 thigh bones *Elephas*; ribs, scapulas, hip bones and vertebrae of ruminants. [Kriele's list has '45 assorted specimens' in Crate 11.]

(12) pieces of *Elephas* thigh bones; deer antlers; ribs, vertebrae, metacarpals; scapula joint (Trinil); buffalo horn; heel bone; ulna ribs and molars of *Elephas*; small knuckle bone Kedung brubus; Hippopotamus skull Nonko, NW of Trinil

[Page] 182 [MM…040-369, right]

(12) Corner of jaw, piece of thigh bone Buffalo; thigh bones, shoulder blades and molars of *Elephas* (Kedung brubus); *Elephas* molar (Pati Ajam); tibia buffalo Teguan; vertebrae Elephas; piece of vertebra between Dekes and Wadekan. [Kriele's list has '45 complete and incomplete deer antlers and 59 vertebrae' in Crate 12.]

(13) ½ jaw ***Sus*** [***brachygnathus***] (Trinil) in fine sand 0.28 M above level; banteng horns; 2 pieces of [***Stegodon trigonocephalus***] tusk Trinil; skull bones of buffalo and *Elephas*; deer molar; vertebrae and tibia, thigh bones *Elephas*; ribs and extremities bones; skull of elephant in pieces with tusk and molar. [Kriele's list has '1 complete buffalo skull with horns and molars; 2 elephant leg bones; 1 complete lower jaw of an elephant; 1 small chest with 51 molars of buffalos and cows; 1 small chest with 192 deer molars; 1 small chest with 140 deer molars; 1 small chest with 39 assorted small jaws; 1 small chest with 25 assorted small vertebrae; 1 small chest with 24 assorted small leg bones; 1 small chest with 37 alligator (caiman) and crocodile teeth; 1 small chest with 62 assorted specimens' in Crate 13.]

(14) turtle; *Stegodon* molar; jaws and thigh bones of *Elephas*; vertebrae large animals; *Bibos* horns just from occiput and 2 horns weak occipital knob banteng type no frontal excavation (?). [Kriele's list has '1 complete skull of a cow with horns; 2 complete elephant leg bones; 2 complete elephant molars; 14 assorted specimens' in Crate 14.]

(14-15) hip bone Elephas Teguan; vertebrae; ribs, knuckle bones; skull pieces banteng; upper and lower jaws ruminants; heel bones; banteng horns; deer antlers; pieces of Elephas molars; thigh bones Elephas; 1 elephant skull in pieces.

(15) Banteng [***Bibos palaesondaicus***] skull occipital and frontal portions with two horns. 1.50 m below lowest level at Trinil; piece of skull banteng; *Stegodon* molar; buffalo head.

[183] [M…040-370]

(15) 2 small tusks *Elephas*; jaws and molars of Elephas; 2 banteng skulls (back portions); supra-occipital [portion of a] ***Stegodon*** [***trigonocephalus*** cranium], juvenile with molars Trinil, compartment [*vak*] I No. 149 in coarse sand 1 M below [river] level; squamo-temporal of the same; 3 molars 0.50 M below level. [Kriele's list has '1 small chest containing an incomplete antelope skull with 2 horns; [187] [M…040-374] 1 small chest containing a complete pig skull with lower jaw; 1 small chest with 2 deer skulls; 1 small chest with 3 deer skulls; 3 elephant molars; 2 elephant tusks; 1 incomplete buffalo skull; 17 assorted specimens' in Crate 15.]

(16) Crocodile skull [***Crocodylus siamensis*** cranium] (Trinil); *Stegodon* molar; jaw fragment *Sus*; 2 parts of lower jaws *Elephas*; piece of upper jaw and molar buffalo; 3 ribs; crocodile teeth; deer molars and pieces of antlers. [Kriele's list has '1 incomplete lower jaw of an elephant; 1 small chest with assorted teeth and molars and several assorted specimens' in Crate 16.]

A 61 Crocodile

A 282 vertebrae Trinil; deer antlers with skull; shoulder blades *Elephas*

A 263 turtle; several pieces of tusk; molar *Stegodon*; complete deer skull; ribs, vertebrae; knuckle bones; extremities bones; 3 shoulder blades and molar plus tusk of *Elephas*; turtle bones.

A 284 banteng horn; buffalo horns with potion of skull; 3 banteng heads one with incomplete horns

(iii) [Page] 184 [M…040-371]

1900

[Crate] (4) tibia ? Wonogane picket 135 depth 8 M sandstone deep 2.80 M

[Crate] (6) tibia elephant (Trinil) fine red sandstone 4 to 5 M above the highest level; knuckle bones buffalo (Tawang) found along the river; pieces of thigh bones and other long bones; of large animals a piece of lower jaw of an elephant

[Crate] (2) metacarpal (Trinil), fine blue sandstone 1 to 1.5 M below lowest level; lower portion of a tibia (Tawang), fine blue sandstone 1 to 1.5 M below lowest level; piece of rib (Trinil) 1 to 1.5 M below lowest level; back portion of deer skull [***Axis lydekkeri***] (Trinil), fine blue sandstone 1 to 1.5 M below lowest level; piece of vertebra and several bone fragments at the same place; upper arm bone (Tawang) at Picket 135 along the river; elephant molar (Tawang) 1.5 M above the lowest level; pieces of long bones (Tawang) along the river; banteng horn (Tawang) 1.5 M above lowest level, fine yellow sandstone; pieces of turtle [**Testudinoidea**] (Trinil) in fine blue sandstone 1 to 1.5 M below lowest level; molar elephant (Tawang) in coarse blue sandstone; tusk; heel bone; vertebra; back portion of

deer [***Axis lydekkeri***] skull (Trinil) in fine blue sandstone 1 to 1.5 M below lowest level; piece of deer [***Axis lydekkeri***] skull (Trinil) (same place); deer antlers [***Axis lydekkeri***] (same place).
[Crate] (9) tusks elephant; scapula elephant; 1 broken deer hollow [pit]; deer antlers
[Crate] (7) complete skull with horns; head with pieces
[Crate] (15) piece of skull.

Page] 185 [M…040-372] 2

1900

[Crate] (10) pieces of turtle
[Crate] (3) pieces of turtle; ribs; hip bone; deer horn hollows (pits)
[Crate] (13) 2 pieces of tusk; 2 Elephas thigh bones; deer teeth, vertebrae and small tibia
[Crate] (16) elephant jaw
[Crate] (14) jaws and thigh bone of Elephas; Turtle
[Crate] (15) Turtle
[Crate] (8) small deer skull partially in stone (Trinil), even with lowest level, coarse sand layer Nr. 79-94-81-93 [repeat of record in M…0040-369, left]
(A 282) vertebrae (Trinil); deer antlers and small skulls (Trinil); scapula Elephas (Trinil)
(47) Complete banteng skull; incomplete banteng skull; banteng horn
(A288) tusks
(81) elephant molar, scapula and bones of ruminants, antlers and bones of deer
(5) vertebrae of large animals
(10) turtles; scapulas of ruminants; deer antlers.
(A 286) vertebrae; scapulas; banteng horn

(iv) KRIELE'S FINAL LETTER FROM TRINIL

[M…058-480]

Trinil 28-12-1900

Very highly learned Sir!

Hereby I am conveying to you that the chests with fossils have been sent, 16 in total, of which three are heavy and the rest averagely so. Upon your writing I have been trying to obtain cotton, even went to Madiun for it, looked in every store, but no cotton, the Chinese would keep it for me, but when I send someone over the 15th he returned empty handed as what little they got they had used themselves for shipping their own goods. I also include a sketch of the work [of 1900] as it is finished [scan M…059-194, next page; also, -A4m, above] and a listing of where beautiful bones have been found [translated on page after next], I hope you may receive them in good order this time, as I did my best to pack them. The last items found were a complete skull of a cow a complete lower jaw of an elephant and some diverse stuff. I did not receive word of the two Engineers but i did receive a note of the chief of the Solo-works that I am not allowed to assist them.

I also wrote to Head engineer Delprat, but he has not replied as yet.

High very learned Sir!
Your very obedient servant

Kriele

[Kriele's map of the 1900 Trench with his December 28, 1900, letter]

[M…059-194; MM774C-000059-194]

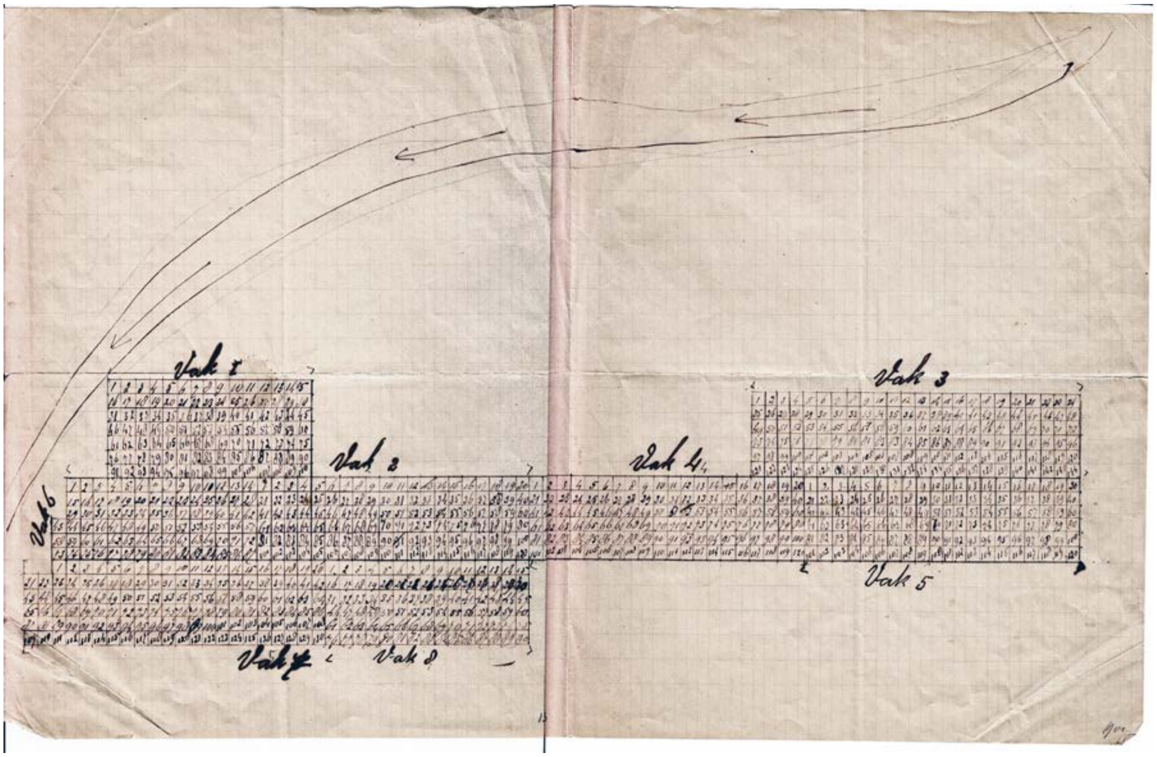


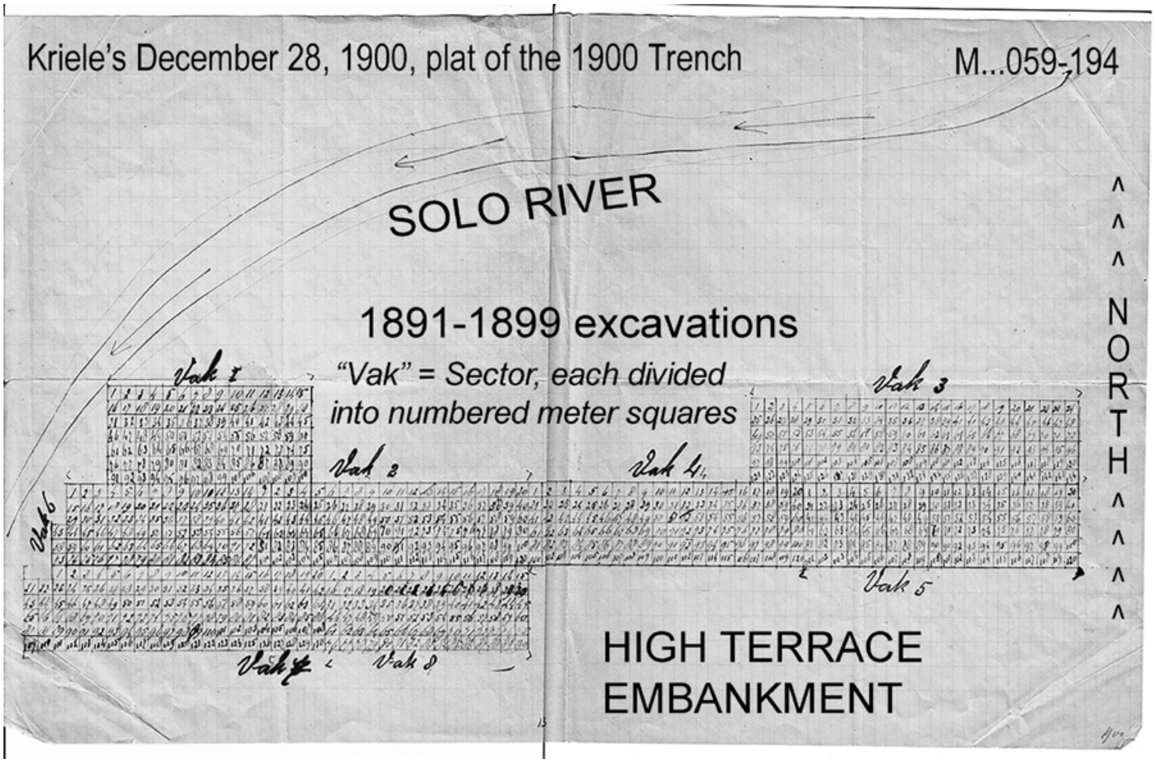

[M…058-482]

Complete contents of 16 filled chests with fossils

| | |
|---|---|
| chest 1 | a complete skull of the elephant with molars and tusks |
| chest 2 | an incomplete skull of the elephant |
| chest 3 | two incomplete turtles, a skull of the cow, 18 pieces diverse material |
| chest 4 | 48 pieces diverse material |
| chest 5 | 58 pieces diverse material |
| chest 6 | 36 pieces diverse material |
| chest 7 | two incomplete skulls of the buffalo |
| chest 8 | 59 pieces diverse material |
| chest 9 | 35 pieces diverse material legs |
| chest 10 | 53 pieces diverse material |
| chest 11 | 45 pieces diverse material |
| chest 12 | 45 complete en incomplete deer antlers and 59 vertebra |
| chest 13 | a complete skull of the buffel with horns and molars |
| | three legs Elephant |
| | a comple lower jaw of Elephant |
| | a small chest 51 molars of buffalo and cow |
| | a small chest 192 molars of deer |
| | a small chest 140 molars of deer |
| | a small chest 39 jaws diverse |
| | a small chest 24 legs diverse |
| | a small chest 37 teeth gharial and crocodile |
| | a small chest 62 teeth diverse |
| chest 14 | a complete skull of a cow with horns |
| | twee complete legs Elephant |
| | twee complete molars Elephant |
| | veertien pieces diverse material |
| chest 15 | a small chest incomplete antilope skull with two horns |
| | a small chest complete skull of a pig with lower jaw |
| | a small chest two deer skulls |
| | a small chest three deer skulls |
| | three molars of Elephant |
| | two tusks of the Elephant |
| | an incomplete skull of the Buffalo |
| | zeventien pieces diverse material |
| chest 16 | a complete lower jaw of Elephant |
| | a small chest with diverse teeth and some diverse material |

Trinil 28 - 12 – 1900

temporary mining overseer

Kriele

[Kriele left Java on about March 20, 1901, a record that was published in ship passenger lists.]

[Eleven entries in Kriele's 16 crates included stratigraphic information on fossils from the 1900 Trench:

(1) an ***Axis lydekkeri*** skull without horns Trinil in fine blue sand at 1 to 1.5 m below low water level;
(2) metacarpal bone from Trinil in fine blue sandstone 1 to 1.5 m below lowest level of the river];
(3) 4 pieces of tibia and a piece of rib, as well as occipital fragment of ***Axis lydekkeri*** and a piece of vertebra and bone fragments, all from Trinil in fine blue sandstone 1 to 1.5 m below lowest river level;
(4) pieces of turtle {**Testudinoidea**} from Trinil in fine blue sandstone 1 to 1.5 m below lowest river level; (5) mandible of a young ***Stegodon trigonocephalus*** with 4 molars, No. 62, from fine sand layer even with the lowest level of the river from a downstream location;
(6) ***Bibos palaesondaicus*** occipital from Trinil compartment (*Vak*) 5. 157 coarse sand 25 cm below level of the river;
(7) ***Axis lydekkeri*** skull partially in stone even with lowest level of the river in coarse sand layer nos. 79-94-81-93;
(8) semi mandible of ***Sus brachygnathus*** in fine sand 0.28 M above the lowest river level;
(9) ***Bibos palaesondaicus*** occipital and frontal with two horns from 1.50 m below lowest level of the river; (10) supra-occipital part of calotte of ***Stegodon trigonocephalus***, a juvenile with molars from Trinil compartment [*vak*] I No. 149 in coarse sand 1 m below [river] level;
(11) tibia *Elephas* {more likely ***Stegodon trigonocephalus***} Trinil in fine red sandstone 4 to 5 m above highest {sic (?)} level of the river.]

[Ten of the LB-HK entries span a 1.75 m depth range (0.28m above to 1.50m below the lowest river level). Six entries (at least nine fossils) refer to sands 1.0-1.5 m below the lowest water level:

A juvenile ***Stegodon trigonocephalus*** supra-occipital with molars, a ***Bibos palaesondaicus*** occipital and frontal with two horns, an ***Axis lydekkeri*** cranium without antlers, an ***Axis lydekkeri*** occipital and turtle remains [**Testudinoidea**], a metacarpal bone, a rib, four tibia fragments, and a vertebral fragment – with four of the six refer to "fine blue sandstone" and one to "coarse sand."

Four entries (four fossils) refer to fine- and coarse-sand at depths ranging from 0.25 below to 0.28 m above the lowest river level**: t**he mandible of a juvenille ***Stegodon trigonocephalus*** with four molars preserved, half of a ***Bibos palaesondaicus*** occipital,a ***Sus brachygnathus*** semi mandible; and an ***Axis lydekkeri*** cranium.

One entry is a "*Elephas*" (***Stegodon trigonocephalus***) tibia from "fine red sandstone 4 to 5 m above highest level," most likely from the *Stegodon* claystone bed that was found in the 1907 Pit II (Figure 5d. Main text). Twelve additional entries (at least 14 fossils) indicate discovery at Trinil but lack stratigraphic data. The fossils are: the mandible of a juvenile ***Stegodon trigonocephalus*** with four molars, an incomplete ***Stegodon trigonocephalus*** cranium with two molars and fragments of tusk, a portion of an ***Axis lydekkeri*** cranium; complete and incomplete ***Axis lydekkeri*** antlers, more ***Axis lydekkeri*** antlers, a ***Stegodon trigonocephalus*** cranium with two nearly complete tusk, only the right tip missing, a ***Stegodon trigonocephalus*** cranium with two molars, the abdominal shield of a terrestrial turtle, additional turtle remains, a scapula joint; and two [***Stegodon trigonocephalus***] tusk fragments.

In September and October Kriele's letters mention several other fossils: "A complete buffalo [***Bubalus palaeokerabau***] skull with complete horns and molars," "a complete elephant [***Stegodon trigonocephalus***] skull with complete tusks and molars," and "a complete lower jaw of an elephant [***Stegodon trigonocephalus***] with molars" (-A4o 'page 23' and 'page 25').]

[De Vos and Sondaar (1982: 47-48) report that one specimen of the small feline *Prionailurus bengalensis* (Kerr) is present in the D.C. (no. 1484), and the fossil "was found in the trench of 1900" (also, de Vos 1989). They also (1982: 48) report, "Coll. Dub., no. 1440, part of the frontal bone with antler {***Axis lydekkeri***}, bearing a label by Kriele: '… Trinil fine blue sandstone 1[.0] a [to] 1.50 m below the lowest water level. The spelling of 'vijn' instead of [the proper Dutch word] fijn, and 'pijl' instead of peil are characteristic of Kriele's labels." The reference to the 'fine blue sandstone…" is a good indication that this find came from the 1900 Trench, because other labels from that excavation bear the same inscription.]

**S II-A4r.**

[**Trial assembly of Kriele's and Selenka's maps of the left bank:**
The 1896 Left-bank Pit (-A4c) and 1897 Up- and Downstream Pits (-A4d) were fitted into Kriele' 1899 map of the 1891-1893 and 1899 excavations (–A4m) and the plat of 1900 vakken (-A4m), guided by other sketches of the excavations (-A4e, -A4f). Conflicts and overlaps arose: The location of the 12x17m 1897 Upstream Pit is clear in Kriele's sketches (-A4e and –A4f; also descriptions in his 1897 letters) but most of the Pit overlaps Kriele's representations of the 1899 and 1900 Trenches. When estimating the size of annual excavations, below, the area of the 1897 Upstream Pit is taken from the 1899 map, placing most of the 12x17m in 1899 and 1900 Trenches. Kriele located the 1897 Downstream Pit west of the 40-m and 1899 Trenches, and southeast of the 1896 Left-bank Pit (-A4f) but 1899 Trench also covers this area. To minimize the conflict, the 1897 Downstream Pit is shifted northward, and placed between the 1896 Left-bank Pit and the 40-m Trench. This created an overlap with the 40-m Trench. The overlapping areas were only counted once in the total area.]

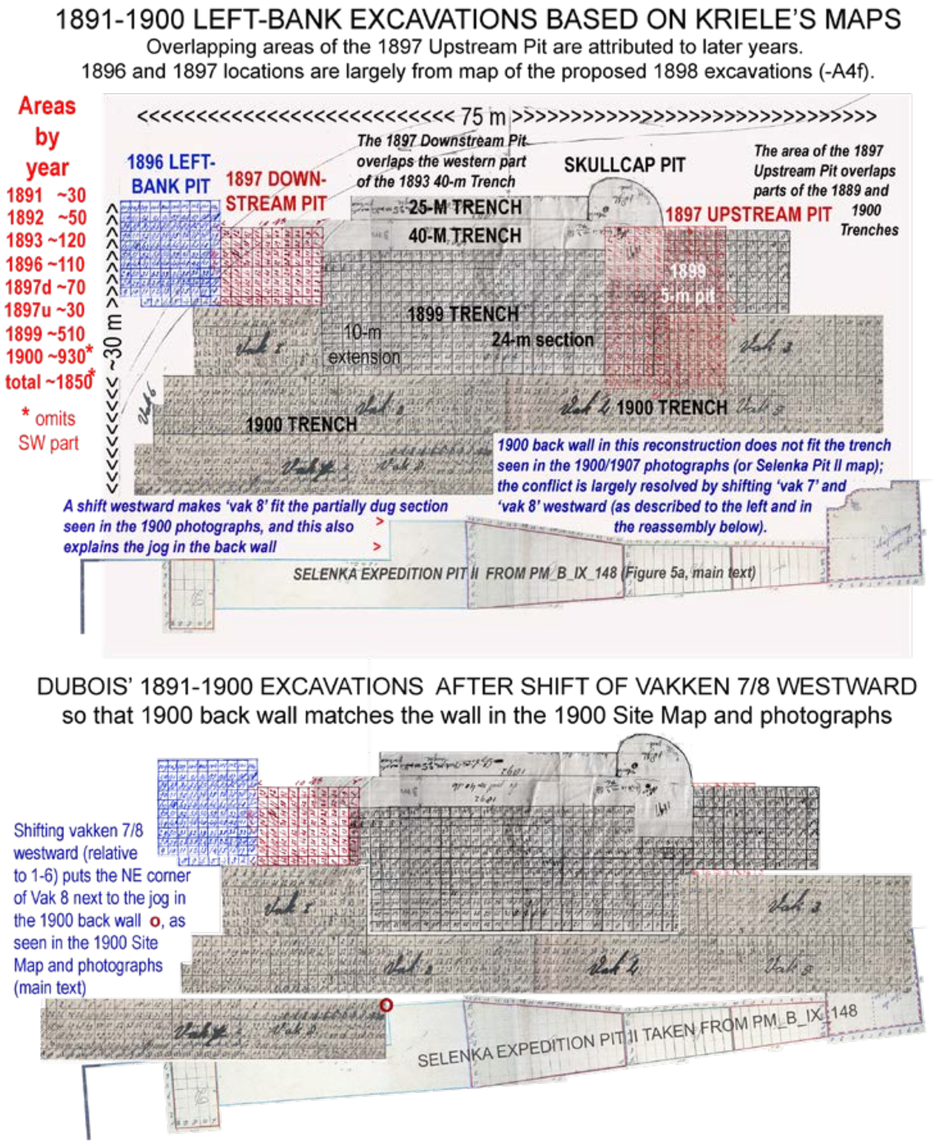


[Left-bank excavations at the main bonebed level: ~1850m$^2$ in the 1891-1900 Dubois excavations and ~350m$^2$ for the 1907-1908 Selenka Expedition Pit II (Table 2, note 4) for totals ~2200m$^2$ for the pits and trenches of 1891-1908.]

**S II-A4s.**

[Letter about Kriele's meeting with L. Selenka in 1902 that lasted 4.5 hours)]

[32] [M…058-485]

Amsterdam
2 October 1902
Most distinguished Sir:

Yesterday afternoon, I sat down with the Lady [Mrs. M. Selenka], and talked with her for as long as 4.5 hours. However, as you know yourself and what the Lady also mentioned, it could take as much as another 7 to 8 months before we can depart for there. That is because a request for a permit has not been submitted. Now, if a permit is granted, the Lady would in that case prefer to leave immediately, but it would actually be best to depart a few months before so that the houses can be built and at the same time also try and finish the dikes etc., because by now everything has been washed away.

I had a very good conversation with her (33) [M…058-486>>]. I did not hear her speak a single harsh word and I believe now that I can travel there without a problem. That is, if she approves it and then possibly [I would go] together with another person.

We have not talked [yet] about monetary matters, but I did receive my travel expenses in the amount of F. 2.10.

In connection with my travels to the East Indies [?; original = 'O Inj'] a request should be submitted to travel at the expense of the Lady, if I will indeed be allowed to go. Would you be willing to submit that request together with my documents?

Regards to all of you,
Faithfully yours, Kriele

**S II-A4t.**
Joede 6 May 1891
Your noble severe Sir

It grieves me to yet again learn from your letter, that you are displeased with me, because of this bamboo, but on my life, I do not write things to please you, all I've written to you is the absolute truth. You know now the amount of money for all the bamboo, please consider how I bought this bamboo. When building the houses every morning free workers came around with bamboo one had a bamboo for eight cents the second had to bamboos for twelve cents, a third had a small bamboo for six cents, et cetera et cetera. You can now understand that the number of suppliers is close to 200.

Please don't think ill of me that I am so free to tell you this. You cannot believe how it grieves me to have to work so hard and so heavy and still not enough to suffice, but the proverb says: once a thief [MM774C-000058-574] always a thief, I've been working now for 10 consecutive days in Djambe, the whole day in the scorching sun, even with the pickaxe in hand, out of fear that the forced labourers will damage the bones. There are many bones to be found in Djambe, and even a lot of jaws and molars of elephants and other large pieces of bone.

If I go home in the evening and I did find a few mandibles, then I think to myself, if the Doctor were to come now, I can show him beautiful bones, but if I get a letter from you like to last one, I only feel weary thinking about the heavy work and being reprimanded on top of it. If bones keep being found like they have I'll be obliged to send bones to you earlier, as within 10 days I've got quite a lot collected. Should you want to come this month I'd rather wait with sending them, as someone them are fragile and I prefer that you saw them here first but if you're not coming I will send bones to you at the 15th or 16th.

My housekeeper has improved a lot so I need not go to Samarang. What the alang alang is concerned, I cannot burn it yet that will take me another half month because it still reigns at night from time to time. I also spoke to a hunter at Pati, and he will deliver me a hogshead with white patches on the cheeks, but I will have to be patient for some days until there is a full moon.

Your serving worker
de Winter

### S II-B. 1890-1893 Trinil discovery narrative in Dubois' governmental submissions.

### S II-B1 (1890)

[Dubois compiled his unpublished monthly memoranda (*Missive* in Dutch) and publishable quarterly reports in a 200-page volume entitled "Report on the Paleontological Investigations of Java 1890-1894 [by] Eugène Dubois" [*Rapporten van de Palaeontologische onderzoekingen op Java 1890-1894 Eug. Dubois* in Dutch] (Dubois 1894d), the title page of which is shown in part below in his handwritting.

Both the memoranda and draft reports were submitted to the Director of the Department of Education, Religious Activities and Industry of The Netherland Indies (Departement van Onderwijs, Eeredienst en Nijverheid). The reports were published anonymously in a government publication, the Reports of the Mining Bureau, Supplement of the Java Daily Report [*Verslag van het Mijnwezen, Extra bijvoegel der Javasche courant* in Dutch] (Dubois 1891, 1892a-d, 1893a,b, and 1894b,c).

The quarterly reports are attributable to Dubois because his handwritten drafts are in "Paleontological Investigations." Our translations of the memoranda are taken from this source. Our translations of the quarterly reports also utilized the "Paleontological Investigations" entries to improve understanding of Dubois' accounts. De Vos and Sondaar (1982), Theunissen (1989: Sources) and our References give formal references to the quarterly reports.]

M…050-052 (MM774C-000050-052)]

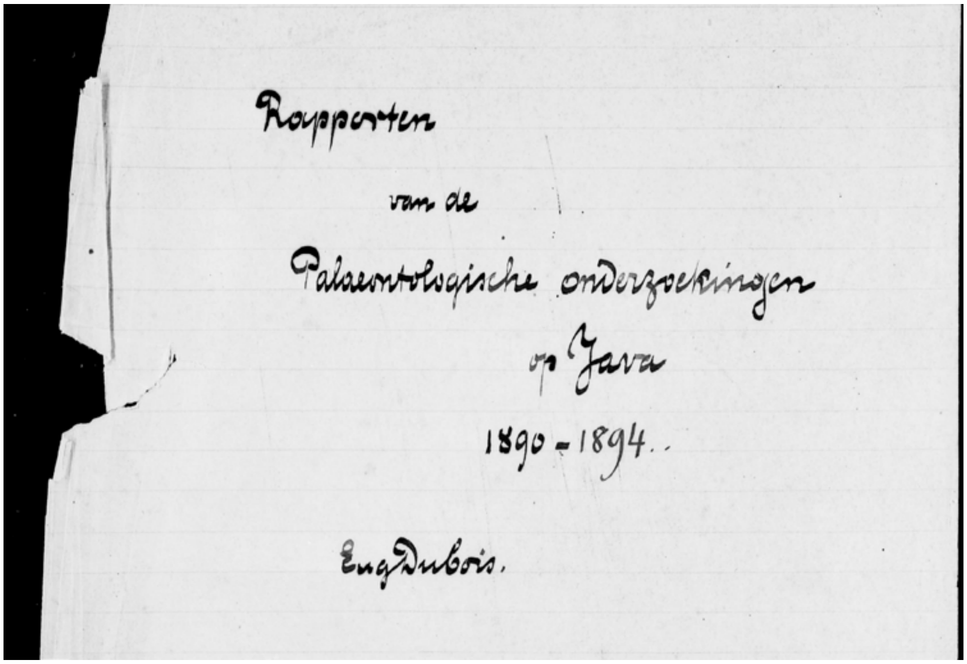


### S II-B1a

[M…050-054. June memorandum contained something of a summary of Dubois activities in the Kendeng Hills. His memoranda were not published.] … fossil bones … seem to be distributed throughout the entire [Kendeng Hills].

[A Java newspaper, The Bataviaasch Handelsblad of 2 July, 1890, carried the announcement – "Extended: Until end of March 1891, a period of one year, the [Army] Health Officer of the 2nd Class M.E.F.T. Dubois has been made available to the Director of Education, Religion and Industry, for doing paleontological investigation in the caves of governmental provinces of Sumatra's West Coast and Java."]

**S II-B1b**

**S II-B1b(i)** [M…050-057/058, left. September memorandum (dated 26 October, 1890).]

[p. 47*] During the beginning of September, we continued searching for Pliocene fauna at Kedungbrubus. Thick cover [of fallen teak leaves] prompted move to Bogo [7 km west of Trinil]. Siwalik fauna bones were found everywhere, in spite of reports of denial by local government. Visited Kendeng mountains in northern Kediri ….. Vertebrate fossils found at … Bangle [east of Kedungbrubus in the Kendeng Hills]. Highlights *Bovina*, *Bubalus platycens*; Bison very different from *Bison sivalensis* Falc.; *Crocodylus* skulls. Bones also at Ngawi [on the Solo River east of Trinil; Figure 1b, main text]. Siwalik deposits along the Solo river near Duku on top of marine marls and limestone. But the fauna points to fresh water deposits. [47>>48*] The sandstone, conglomerates and andesitic breccias and tuffs near Duku at the Solo river, lie directly on top of marine marls and limestones; they are therefore all equivalent to the claystones.

[* The numbers refer to pages in Dubois unpublished drafts in "Paleontological Investigations."]

**S II-B1b(ii)**

Summary Report of the paleontological investigations on Java during the Third Quarter of 1890.

Research on the Siwalik fauna of Java was continued around Kedung Brubus (District Tjaruban, Department Madiun) up to the beginning of September. In the end, the thick cover of Djati [teak] leaves made the search and collection of fossils difficult to the extent that this place had to be temporarily abandoned [the teak plantations of the area form a continuous forest, and once the large leaves drop during the dry season, they cover the ground almost entirely].

Fossilized bones had become known everywhere in the mountains around Kedung Brubus [also Kedungbrubus] as far as across the boundary with Department Ngawi (near Teguan in District Dero) [on the west] and up to the southern slope of the Pandan near Klangon [on the east]. Siwalik vertebrates had also been found north of Kediri, in the District of Lengkong [even farther east]. For the time being, a more westerly point was chosen and a new series of traverses were started from Bogo [7km west of Trinil]. From various sources we had information about the presence of fossil bones at Bogo, as well as the nearby located settlements of Alastoca and Kali Gede and Tawang (at the boundary with Department Pragen [3km west of Trinil]). Several specimens were already known to us from these locales through kindly received shipments from interested parties or through their investigations in place.

Siwalik vertebrate bones proved soon to be present here as well as in the eastern half of the Kendeng mountain range and also everywhere outside of the known locales sometimes in larger numbers and other times more sparsely. Representatives of the named fauna have now been found from the northern to the eastern extremes of this mountain range and thus covering nearly the entire distance from Semarang to Surabaya and in regard to the southern slope also over the entire width of the mountain range. They are also already known to occur at several points on the northern slope but these have not been investigated in any detail. The Siwaliks on Java are thus not less widespread than those of the region south of the Himalayas, even if those occurring outside of the Kendeng mountain range are not included. [see Figure 1b, main text]

In the Department of Ngawi [see Dubois field map of this area, -*E1a*], bones were found, besides in sandstone and finer or coarser conglomerates, also in breccias and tuffs (which all mainly consist of gravels and [veins?] of andesitic material but also include small pieces of limestone (and other rocks) and (near Pengilan and Alastua as well as near Banglee) in white clay stone.

In the incised Solo river one can see these sandstones as well as conglomerates, breccias and tuffs resting on marine marls and limestone (*as well as the white claystone rocks near Pengilan and Alastua* [faintly written in draft, and omitted in the published report])

All these bone bearing rocks have thus been deposited after the marine lime and marls and their full equivalents. They belong to deposits by fresh water flow as becomes clear from, aside of crocodiles and river turtles, the occurrence of *Bulinus* shells and other fresh water shells belonging to the Unionidae [fresh-water mussel] family, which very often accompany the bones.

During a period of somewhat over three months, a large number of attractive fossils of tremendous value were gathered the majority derived from, at least for Java, completely new species of vertebrates. The latter are still increasing in numbers and often different and better-preserved specimens of the already known species are being found. We have definitely only just started to get to know this fauna and no doubt the richness of the Siwalik formation on Java will eventually measure up to that of British India.

Of the animals that have become known to date, the most numerously represented are the elephants and in particular the Stegodonts whereas the sub species *Elephas* is much rarer. Then follow the large ruminants of the families of Bovids and Cervina and then *Hippopotamus*, *Sus*, *Rhinoceros*, *Hyaena*, *Felis*, and even less abundant again are *Crocodylus*, *Garialis* and turtles belonging to the family of Trionychidae.

To date we were able to recognize the following genera and species. Those marked with a "+" are new for Java (or entirely so [not previously recognized anywhere]): + *Hyaena feline* Bose + *Felis* spec. *Stegodon* Combrifous Falc Cantl, ____ Cliftu Falc. Cantl, ____ spec., divers. *Elephas hysondicus* Falc Cantl, _____ *namadicus* + *Rhinoceros* spec. a + *Rhinoceros* spec. b + *Hippopotamus* (*namadicus*?) + *Sus* spec. nov., __ *hysundicus* Falc. Cantl + *Bubalus platyceros* Lyd + *Bison* spec. + (?) *Bovina* spec. c.d. + (?) *Cervus* spec. a [faintly annotated: *Lydekkeri Martin*] + _____ spec. b, + *Capra* spec. nov; + *Crocodylus* spec. a, + (?) ____ spec. b + *Garialis* spec + Trionychidae (fam) spec. a + ________ spec. b.

It is never too difficult to determine the genus but only rarely is it possible to determine the species because of the lack of literature and comparison material. We hope to obtain these in the near future. In any case, those marked with a cross were not yet known from Java.

As far as the collected bones have already been studied, our research initially has not suggested any animals that are extinct or strange for Java. However, towards the end of September an important find was made of a second fossil human skull with the same characteristics as those of the previously discovered one. Any thoughts of an individual evolution (as unreasonable as that is by itself) must therefore be rejected and we should accept the presence on Java in previous times of a race that corresponds to the current Australian population (or Papuas).

**S II-B1c**

[p. 48. M…050-058/059. October memorandum (dated 14 November, 1890).]
Worked out of Bogo [7km west of Trinil] in the Kendeng hills, searching for Siwalik vertebrates [of the kind mentioned in the Third-quarter report, above]. Also at Trinil, Matingan (near the Solo border) new material has come to light about the old Java fauna. The best find spots along the shores of the river will soon be impossible to work [this season] because of the steadily increasing rains.

**S II-B1d**

[p. 50. M…050-058. November 1890 memorandum (submitted on December 12, 1890).]
Solo water [was] too high and on 21 Nov [1890 we] moved to Kedungbrubus. No new species found but much geologic information gained. Core of Kendeng hills [exposes]: white claystone hundreds of meters thick followed upward by sandstone and ending with marine limy marls and limestone. Bone beds are only at the northern and southern rims. Found vertebrates after some searching. …

**S II-B1e**

[p. 52. M…050-060/61. November 1890 memorandum (submitted on December 12, 1890).]
… diagonal bedding (Lyell) [1833 or] oblique lamination (Dana [e.g., 1879) [was] observed [in the sandstone sequence near Kedungbrubus]. The manner of bone burial [there] was in thick slurry (volcanic catastrophe) …. [p. 55.] Vertebrates lived along shores [and] shallow bays for hippos, crocs. [A] last sudden eruption killed many …

**S II-B1f**

[p. 58. M…050-064, right. December memorandum (dated 6 February, 1891).]

On Nov 24, 1890 … a right-side chin portion of a lower jaw with alveoles of the canine tooth and PM-1 and -2, [was] found in the midst of typical representatives of the local fauna. When it got to me, still attached was a piece of sandstone-like andesitic tuff, which is the main sedimentary rock formation around the bones. No other remains were found then or later, which is not surprising since other fossils are usually broken up and distributed. The shape of this bone fragment leaves absolutely no doubt that this is derived from a human.

[Dubois noted in his diary on Monday, November 24 (M…015-195): “l[ower] j[aw] of *Homo fossilis pandanicus* near Kedung Lumbu” (in original Dutch: “O.k. van Homo fossilis pandanicus bij Kedoeng loemboe”). See S I Figure 21.]

**S II-B1g**

p. 61. M…050-066, right. Fourth-quarter report 1890. [Dubois 1891. The submission was delayed until May (?). Dubois announces the discovery of the mandibular fragment from Kedungbrubus; also, S I Figure 21.]

On November 24 [1890] a more important find was made of a human fossil in the midst of representatives of the typical fauna and in the same bed of sandstone-like andesitic tuff: the right chin portion of a mandible with alveoles of the canine and the first and second pre-molars. No further remains of this human were found at the time, nor later, which is not surprising since other bones are usually broken and spread out. There can be no doubt about the human origin of this fragment. The chin must have been even less prominent than that of the humans [Neanderthal] of La Naulette [Belgium] and Schipka. Pronounced flattening and hollowing along the inner surface [of the mandible] due to attachment of the musculus digastricus, which in Pleistocene jaws of Europe is only slightly stronger than in living humans, assigns this fossil to a different and probably lower type than anything we know at this time. In addition, it is now proven that this human lived on Java together with *Stegodon* elephants, hippopotami and hyenas and during a time when Java was connected to mainland Asia (based on the fauna, a Pliocene relationship is assumed by Prof. Martin). This had already been concluded by the genial Wallace when he looked at the present distribution of animals and that sea level would have been lower by not much more than 100 m.

**S II-B1h**

[On January 29, 1891, Dubois (1892a) completed a manuscript (titled in translation) “Preliminary notes on the investigation of the Pleistocene and Tertiary vertebrate fauna of Sumatra and Java during 1890 by Eug. Dubois,” in which he commented on the Kedungbrubus find.]

[94] Of *Homo*, only a small mandible fragment has been found and in one respect the difference with the only extant species in this genus is so great that for the time being it seems prudent to forego a more detailed identification.

[Other notable excerpts from the pages 93-99 are:]

[93] During the first three months of the previous year, searches were continued in the caves along the west coast of Sumatra. Many remains were collected, primarily near the west and east shores of lake Singkarah of a fauna that is completely equivalent to that of the Lida-ajer cave near Pajakombo and some caves in the Sub-Departments Lintou and Bua. This fauna is characterized by the relative frequency of occurrence of orang utans, besides other extant species on Sumatra. Nevertheless, while geologic phenomena of the caves and the distribution of the animals undoubtedly suggest that they are from pre-historic times, this fauna that almost entirely consists of extant species cannot be assigned to the Pleistocene but to the Holocene period.

Neither was a different animal world encountered in the caves of the Ngrowo department, Res. Kediri on Java that we investigated during the last half of the previous year. Its pre-historic character was clearly proved by the occurrence of remains of human skeletons (skulls) [94] which have the characteristics of the Australian race, and therefore most likely represent the autochthonous race of Java.

The fauna of great vertebrates discovered by Junghuhn, Raden-Saleh and others, that later became known through Martin, is undoubtedly an older one. It comprises the area of the Gunung Kendeng at the boundary of the Residencies Kediri, Madiun and Solo with Rembang and Semarang, the Pati-ayam (south of Muria) in Japara, the western part of the Residency Jogjakarta and several other locations in Central Java. During the last part of 1890, we were able to add a large collection to those previously reported on, which at the same time became better known and provided us with information to better judge the geologic nature of the fauna and therefore also enhance knowledge of the geology of Central Java. [A listing and short description of the genera and species that were known by 1891 is presented on page 94 – 97; the following is an updated listing with modern taxonomic names after the = signs; this represents about 50% of the taxa that have been recognized in the *Stegodon-Homo erectus* faunal association]

*Homo, spec. indet.* [ = *Homo erectus* ]
*Felis, spec. indet.* [ = *Panthera tigris* ]
*Hyaena feiina,* Bose. [ = *Hyaena brevirostris* ]
*Elephas indicus,* L. [ = *Elephas hysudrindicus* ]
*Stegodon, spec. indet.* [ = *Stegodon trigonocephalus* ]
*Tapirus, spec. indet*. [ = *Tapirus indicus* ]
*Rhinoceros javanicus* [ = *Rhinoceros javanicus* Cuv. ]
*Bos* (of *Bison*?)*, spec. nov.* [ = *Bos palaeojavanicus* ]
*Bubalus buffelus*, Blum. [ = *Bubalus palaeokerabau*]
*Bibos banteng* [ = *Bibos palaesondaicus*]
*Axis axis*, Erxl. [ = *Axis lydekkeri* (Martin) ]
*Cervus* (incl. *Rusa), spec. indet*. [ = *Rusa* sp. ]
*Anoa sp.* [ = *Duboisia santeng* ]
*Sus celebensis* [ = *Sus brachygnathus* ]
*Sus verrucosis* [ = *Sus macrognathus* ]

From the above it is clear that these fossil vertebrates of Central Java do not correspond to those of the Siwaliks, but that they should definitely be assigned to the Pleistocene. No older formation with land vertebrates has been found anywhere. The fossil-bearing sediments irregularly cover the late Tertiary marine crest [core] of the Kendeng, and only near the Pandan are they present as high as the crest of the mountain.

They [fossil-bearing sediments] are all of a volcanic nature: sandstone-like andesite tuffs as well as coarser tuffs and breccias and sometimes conglomerates. Real bedded deposition can therefore only be observed near the lowlands, but signs of irregular and often cross cutting beds also occur at the higher elevations, evidently related to the slopes of the mountain side along which the mass appears to have flowed as slurry. These bone-bearing rocks completely resemble the same types of sedimentary rocks that cover the lowlands of Madiun, Solo and Kediri. They appear to have been deposited in the same manner [as] and partly they transition into these rocks of the lowlands. Historical eruptions of the Kelut that delivered products to the Kediri lowlands, consisting of sands, sometimes hardened to sandstone, tuffs and breccias, indistinguishable from the Pleistocene on the Kendeng slopes. Sedimentary rock material that encloses the remains of the Pleistocene Java fauna has undoubtedly similarly been carried to its destination by an eruption. This would have been partly in dry condition as volcanic sand, lapilli and bombs etc., but especially during heavy rains that mixed with them in the form of heavy slurry [lahars] flowing down the slope. The animals would have perished in the same manner that the inhabitants of the Kelut slope can now tell us about during the historical eruptions of this volcano. After the last eruption, many cadavers of pigs, kidangs, deer, bantengs, tigers and other forest animals were found on and within the volcanic sand etc.

We cannot tell with certainty which volcano delivered the eruption products that cover the Kendeng. Their distribution makes it most likely [99] that it was the Pandan. It is true that no crater can be observed there, but it is constructed in the same manner that all great volcanoes of Java and Sumatra are, i.e. with a solid core of andesite, surrounded by loose material. However, it is also possible that also both the Wilis and Lawu, or either one of them, were also the likely producers, delivering the volcanic sand, tuffs etc. to the vast lowlands. This possibility becomes more credible with the find of a fossil from the bone bed fauna in the volcanic breccias near

Kedung-Galar and the presence (according to reports that we have not been able to confirm) of such similar fossils at the northern boundary of Department Ponorogo, west of the Wilis. The most northerly place around the Pandan where bones were found, is Gerbogan near Kali-tidu (Distr. Ngumpak) in Bodjonegoro. Only when distribution of the fossils and the sedimentary rocks that contain them become better known, will it be possible to point at the eruption point that delivered them with some certainty.

The black clay that in many places covers the lowlands of Central Java is just an erosion product of the breccias and tuffs and is not an independent formation. This becomes clear at much higher locations in the Kendeng, where a transition can be observed. The erratically formed small lime concretions that are found in the clay are derived from the breccias and tuffs.

## S II-B2 (1891)

### S II-B2a

[M…50-070. July memorandum.]
[In the Ngawi area, the] main finds were at Padasmalang (Sepreh) on the left bank of the Bengawan [Padasmalang is 1.5 west-northwest of Trinil].

### S II-B2b

[M…050-070. August memorandum (dated 18 September, 1891).]

An even more important discovery spot in this region was one in the currently dry portions of the Solo [river] bed. Primarily, found all together [in the LB, evidently all on the left bank] were the remains of *Cervus axis*, *Anoa*, *Bubalus paleoindicus*, *Stegodon,* in andesitic sandstone at Tinil [Trinil] as well as of the fossil land turtle …, besides *Unio* shells and fossilized wood [***Axis lydekkeri***, ***Duboisia santeng***, ***Bubalus palaeokerabau***, ***Stegodon trigonocephalus***, **Mollusca** (mussels) and **wood**, respectively]. This once again proves that these accumulated vertebrate animal species are geologically equivalent [to those elsewhere in the Kendeng beds], while the numerous [Mollusca] shells indicate that the bones settled here in fresh water. Once again they [the vertebrates fossils] are often fractured but no indication of violence [human or carnivore predation] was found. Also, several vertebrae of a small ruminant were preserved in natural articulation, so that because of the preserved connections, proof is given that these fossils lay in their original place of deposition. The rear skull (occipital) half of a fossil *Anoa* with horn was found. ….. The fossil wood appears similar to wood from a European bog (where it is spread around from burning) and also is like this burning peat.

[Dubois' description of the fossil taxa in the LB must have been based upon his September 7-9 visit to Trinil, when just the left bank was being excavated (Skullcap Pit; Figure 3b, main text; the fossilized wood collected by Dubois has not been identified (S I Figure 20e).]

**S II-B2c**

[M…050-070. August memorandum (dated 18 September, 1891).]

[in one search area, the] fossil bones, which are exposed at the surface due to weathering and wasting away of the rock around them, are covered with lime concretions, as is observed often … in the Kendeng. [paraphrased, here]

**S II-B2d**

[M…050-070. September memorandum (dated 27 October, 1891).]

Searching in the Patiayam [near Java' north coast; Figure 1b, main text] was continued till the 12th [of September] but since the fossil fauna proved to be identical to the one in the Kendeng it was decided to discontinue the search there for the time being.

The corporal of the engineering corps, de Winter was also employed at Tinil, where rapid progress of the work was desirable because of the soon expected higher level of the Bengawan [Solo River]. Exposed dry shallow sandstone ledges in the river bed were excavated to below water level on both sides of the river. Advancing water was held back as much as possible with dikes and removed by continuous bailing. The sediments were identified as fresh water deposits by the occurrence of fossil wood and imprints of leaves as well as numerous freshwater shells. A number of often well-preserved remains of vertebrates were found. *Anoa* [***Duboisia santeng***], the small ruminant; many antlers and a skull of the fossil ***Axis*** [***lydekkeri***] deer, buffalo [***Bubalus palaeokerabau***], giant skulls, ***Stegodon*** [***trigonocephalus***] molars and some incomplete skulls connected to extant Java species. ***Batagur*** [sp.], ***Crocodylus*** [***siamensis***] and ***Garialis*** [***Gavialis bengawanicus***, an extinct gharial].

However, the most important find of all was a molar (the upper right third molar) of a chimpanzee (*Anthropopithecus*). This family of human-like apes that currently is found only in West and Central Africa, lived in India during Pliocene times and as is evident from this find, on Java in Pleistocene time.

[*Anthropopithecus*, as used here, approximates *Pan troglodytes*. The 1891 Molar evidently arrived with the fossils that KdW had sent on September 26 without mentioning a primate tooth in the accompanying letter (-A1c). After being notified of this find, KdW began looking for the teeth of humans or apes by October 11 (-A1e). Dubois visited Trinil on October 21-24, shortly after writing this memorandum. He spelled the site name 'Tinil' (see -A1m for fossils in the Dubois Collection bearing inscriptions with this spelling).]

**S II-B2e**

[M…050-071. Dubois, 1892b. Third-quarter report, 1891.]

Also, districts Sepreh and Gendigen in the Ngawi dept. were searched and collected fossils were almost as numerous as last year. The best find spot of all was discovered in the month of August near Tinil [Trinil] in Sepreh distr[ict]. A considerable number of fossils have already been excavated in two shallow sand[stone] ledges that were exposed in the bed of the Bengawan on both sides by low water level.

These fossils are generally better preserved and are distinguished in that they are more complete. Among them some nice pieces of skulls and jaws were found of a curious small ruminant [***Dubosia santeng***], which like the extant *Anoa* of Celebes, forms a link that connects antelopes with cattle, making it possible to sufficiently characterize this species. Also, we found more or less complete skulls of ***Bubalus*** *paleoindicus* [***B. palaeokerabau***], and molars, a tusk and two incomplete skulls of ***Stegodon*** [***trigonocephalus***]. In addition, beautiful leg shields of a terrestrial Turtle (probably ***Batagur***) and the presence was noted of ***Bibos*** [***palaesondaicus***], ***Rhinoceros*** [***sondaicus***], *Sus* [***brachygnathus***], felis and [***Panthera tigris***] a small rodent (*Mus*? [Muridae, presumably the Trinil rat, ***Rattus trinilensis*** Muser 1982])

And one or two lower apes that cannot be further identified with certainty [**Primate**]. Among the reptiles: ***Crocodylus*** [***siamensis***], ***Garialis*** [***bengawanicus***] and *Trinonyx* [**Testudinoidea**].

However, the most important find was a molar (the upper right third molar) of a chimpanzee (***Anthropopithecus***). This family of human-like Apes, currently only present in Central Equatorial Africa, lived in India during Pliocene times and, as is evident from this discovery, in Java during the Pleistocene. Fossil wood and leaves and numerous river fossils clearly identify the formation as a fresh-water deposit. The tree trunks and leaves are always found horizontally [**wood** and **leaves**] and some bones have their natural articulation still preserved. Hence, the animal and plant remains are found at their original place of deposition and the beds have not subsequently been deformed.

**<u>S II-B2f</u>**

[M…050-071. October memorandum (dated 17 November, 1891.)]

After it was discovered in September that a chimpanzee was also part of the Pleistocene fauna of Java through the find of a singular molar, we were now able to judge this species even better. This month, close to the place where the molar was found in volcanic tuff on the left bank of the Bengawan, a magnificent skullcap was excavated that, at first glance, undoubtedly must be ascribed to the species ***Anthropopithecus*** (*Troglodytes*), just like the molar was. It is immediately clear that the two specimens are derived from an anthropoid. The molar conforms entirely in shape to the M.3 of a living chimpanzee and is only distinguished from it by its increased size. The skullcap can be distinguished immediately from that of an orang-utan by its dolicephalic character and from a gorilla by the complete absence of strongly developed ridges, so striking in the great African species. On the other hand, it fully shows that it belongs to a chimpanzee species. The skull of this species is distinguished from that of the living chimpanzee by its larger size as well it's higher vaulting. … [and] flat parietal bones and the little developed occipital. Of all known living and fossil anthropoids, at least certainly of those from which the skull vault is known, the new Java chimpanzee undoubtedly ranks highest.

**<u>S II-B2g</u>**

[M…050 -072. October memorandum (dated 17 November, 1891).]

The enclosed photography shows the skullcap in side view and top view, the forehead oriented to the right.* The upper part of the occipital portion is covered by a stony mass (mainly consisting of the same lime concretion found at many places in the lowlands of central java on bones extracted from the black clay).** This mass is so hard and adheres so strongly to the bone that, for the time being, it cannot be removed without causing damage to this precious fossil. Fossils of other animals were excavated, amongst which a skull of … *Leptobos* [large bovid]. The unexcavated surface area of fossil-bearing strata along the Bengawan near Tinil [Trinil], in which we have now dug [through the LB] to 1.50m below the water level***, offers an opportunity for promising additional searches. However, at this moment, that will have to be postponed, at least for the major part, until the next dry season because of the anticipated rise in the river's water level.

[* We have not found these photographs in the Dubois Collection. ** $CaCO_3$ nodules occur on fossils *in situ* (–B2c, -F6). ** Dubois must have seen the 1.50-m level during his October 21-24 visit. *** Whether he is referring to a depth below the LWL of 1891 or water level at some other time is unclear. He appears to have been less attentive to the position of low-water level than were KdW given that Dubois did not portray it correctly in his 1895-1896 cross sections (B7-, below).]

**S II-B2h**

[M… 052-072. November memorandum (dated 23 December, 1891).]

Water levels of the Bengawan remained low enough into the fourth week of this month for excavations to continue uninterrupted in the shallow sandstone ledges [LB] which at Tinil [Trinil] are part of the river bed. Then, in one night the water rose so much that all work that had progressed quite a bit below this year's particularly low water level [LWL], had to be interrupted in spite of our dikes. After that, it was impossible to do any further work. In the meantime, an excavation was done at a more elevated point on the [incised] river bank [in the upper LB or above it] ... There, many bones were found, amongst them an elephant [***Stegodon trigonocephalus***] skull, which was nowhere nearly as well preserved as those found further down in the harder sandstone [such as the LB] which for most of the year is protected from the air by water …. I consider it quite essential to excavate at Trinil during the next East monsoon and look for the anticipated other parts of remains of ***Pithecanthropus erectus*** [that still should be present in the unexcavated portions the LB along the left bank].

[On December 30, 1891, Dubois wrote to R.C. Kroesen, a high government official in the Indies, that the Skullcap was "truly a new and closer link in the largely buried chain connecting us the 'lower' mammals (in Theunissen, 1989: 55; M…007-486/487). On November 30, December 7 and December 8, KdW reported that de Winter was digging at 0.75m above the LWL "in the side bank" of the right side of the river (-A1h, -Ali and –A1j); we place this elevation at the top of the LB. Dubois continued spelling the site name "Tinil" governments submissions until "Trinil" started being used in early 1892 (-B2i).]

**S II-B2i**

[M…050-073, right. December memorandum (dated 20 January, 1892).]

As excavation of the river embankment near Trinil … has become so important to science …. The best finds were: *Bubalus palaeoindcus* [***B. palaesondaicus***] (skull); *Leptobos* [large bovid] (skull); antlers of ***Axis*** [***lydekkeri***] deer and a *Felis* species (beautiful mandible half) that had become known before at Trinil by loose molars, canines and jaw fragments. With this find, it is now possible to exactly determine this fossil cat [***Panthera tigris***], which must have been about the size of an average royal tiger, but deviates otherwise, when compared to the required materials.

[Dubois visited Trinil on December 25-27; the LB was again accessible. De Vos and Sondaar (1982: 47; also, de Vos, 1989) indicate that Dubois Collection, D.C., no. 1479 is the right mandible of *Panthera tigris*.]

**S II-B2j**

[M… 050-073/074. Dubois, 1892c. Fourth-quarter report (dated 20 January, 1892).]

Up to the middle of November we had continual great success, but when the shallow ledges [*platen* in Dutch] of sandstone-like tuff in the riverbed [LB] became inundated, we had to abandon those [ledges], although the major part of them had already been excavated. At that time, excavations were started at higher points on the [natural] river embankment and these produced at first less well-preserved fossils but later, in the deeper layers, they were in as good a condition as those of the shallow [that is, submerged] ledges [of LB]. But soon, towards the end of December, river level rose, at first intermittently then continuously and to such height that this work also had to be considered done.

The results for this Quarter can again be called outstanding. If in September we increased the list of Pleistocene mammals on Java by the find of a molar of a chimpanzee, no more than a month later we were fortunate to gain more knowledge about this interesting species. Close to the spot on the left shore of the river where the molar was found, a beautiful skullcap was excavated. It undoubtedly must be assigned, like the molar, to the family ***Anthropopithecus*** (Troglodytes). It was immediately clear that both specimens are derived from a large human-like ape. … The skull can immediately be distinguished from that of an Orangutan by its *dolichocephalic* character and from the gorilla by the complete absence of prominently developed ridges.... Instead it shows complete resemblance to the chimpanzee species. There can thus be no doubt about the genus, but in regard to the species, the skull is distinguished from the extant chimpanzee 1) by its larger size 2) by its higher doming. …. Both the skull and the molar indicate the size of an animal in excess of the living chimpanzee. In judging its size, the Java species must have even exceeded the erect posture of a gorilla which can be 2 meters ….

### S II-B3 (mid 1892)

### S II-B3a

[M…050-075, right. March memorandum (dated 4-05-92).]

…. encouraged by the favorable results at Trinil, I decided to try again with a more extensive excavation near Kedungbrubus at the spot [S I Figure 21 at crossed tools] that seemed suitable because of what was found at the surface [previously] …. a very nice specimen … of the lower jaw of a *Gavialis gangeticus* [***G. bengawanicus***] … offered splendid proof of the identical nature of the Java fossil with extant species living in the Ganges.

### S II-B3b

[M…050-076, -077, left. June memorandum (dated 29 July, 1892).]

During the first half of this month [June], [loose] sand had to be removed also on the right shore of the Bengawan at Trinil. This had been carried in by the river during the rainy season and deposited on top of last year's excavation work [Figure 3b, main text]. In addition, the river was initially too high on the left bank to reach the [fossil-] rich, deeper layers, so that excavations were done from above into the [left] river bank, exposing a larger surface area of the fossil-bearing layer [LB]. Due to several factors, only during the last days of the month, bone collecting became just as rewarding as it had been here last year [in the LB].

Besides many horns [antlers] of the fossil ***Axis*** [***lydekkeri***] deer, also three large skull fragments of the same species [***Axis lydekkeri***] with more or less complete horns [antlers] were found, one skull fragment was more important than ever before for the determination of a species {[inserted at bottom] the previously mentioned small ox-like antelope) [***Duboisia santeng***]}. Also, three beautiful skull fragments of ***Stegodon*** [***trigonocephalus***] and the tusks were found of which one was almost complete as well as half a mandible. Also of importance for other reasons were a number of vertebrae and ribs of a large ox [***Bibos palaesondaicus***] in their natural relative position and a piece of the great artery of a relatively small ruminant (probably a deer [***Axis lydekkeri***]), entirely filled with andesite tuff. Both these fossils prove again that they are clearly at a spot where they were originally deposited.

This observation, together with other facts, probably helps explain why the bones are rarely being found in their relative position and are usually separated and fragmented, while they show no characteristics of having been torn apart by the mechanical force of [fast] flowing water …. Separation and fracturing of bones must thus have been caused by animal activity which is positively proven by the piece of artery filled with sandstone-like tuff, because if this strong but flexible organ was not bitten or torn off, it would not have filled in but closed and flattened below the sand. However, savage land animals cannot be the culprits of the osteologic vandalism because then, recognizable tooth marks would be visible. These must have been large predators in the water that separated the bones and fractured them while they were still surrounded by soft tissue and no other species comes to mind than huge crocodiles, whose remains (mainly teeth) are found so abundantly among the other bones. Some spongy bones show well-rounded holes that fully match the voracious water animals.

[Dubois visited Trinil on June 29, while the LB was being dug; he also visited in early June, and went for 9 full days in July on 5-14 and 18-20 (–A2e and –A2f). The left-bank pit from which Femur I came was expanded into the 25-m Trench in late 1892 (Figure 3a, main text). Excavations of some sort were also being done in June-August 1892 on the right bank.]

**S II-B3c**

[M…050-077, left. June memorandum (dated 29 July, 1892).]

Other facts were observed that could serve as evidence for the manner in which the numerous animals, whose remains are now being excavated, found their death. First of all at about 1 meter below the lowest water level of the river near Trinil, a blue-grey clay[stone] variety was found [apparently in an excavation], immediately below the sandstone-like tuff …. This indicates a time of very slow deposition ….. By contrast the superjacent beds of sandstone and conglomerate, which are found horizontally everywhere along the banks of the Bengawan, must have been deposited in faster flowing water.

We were able to determine the direction of the former currents for a certain stretch of these beds, namely between Trinil to beyond Pitu [5.9km east of Trinil]. This was possible by investigating in more detail the thin slanted drift beds (oblique lamination) present everywhere that this formation occurs and which has been mentioned before in these reports.

As is known and clearly could be observed in the sand banks that were formed by the Bengawan during the last West monsoon, parallel beds [sets] dipping 30° or more were formed at the downstream steeply dipping terminations of the growing banks, against which the sand carried along by the stream across the bank came to rest [foreset laminations]. They are on average about 30-cm thick and sandwiched between horizontal thin layers of lighter material [bottom-set and top-set laminations], indicating periods of diminished current when the sand came to rest already before it reached the sloping end of the bank. These slanted drift beds that are truncated in this manner thus provide a reliable way in which we can judge the current direction when they were formed. (+ Compare J.D. Dana, Manual of Geology 3rd Edition 1879, p.663 and A. Geike, Textbook of Geology, 2nd Edition 1855, pp 464-466)

[Dubois noted the presence of claystone below andesitic tuff in his diary entry of Tuesday July 19, 1892 (M…015-364), “claystone beneath andesite of 5 [or 3]; ½ feet higher [shallower] in between at Trinil.” A notebook entry, apparently made in June-July 1892, (M…046-142) states, “claystone below the andesitic tuff at Trinil also thin layers of 10-15cm thickness catastrophe;” also, -B3d. Dubois’ 1895-1896 site cross sections (e.g., Figure 2a, main text) have two units underneath the LB: a conglomeratic facies, and below it, claystone (Figure 2a; -F1). This was based on his observations in 1893. Kriele had reported in October 1893 that the bone-rich layer was underlain by a fossil-poor “coarse layer … almost entirely gravel,” in turn was underlain by “black” claystone (-A3l). Dubois confirmed these relationships (-C6). See -C3 and –C4, below, regarding Dubois’ paleocurrent analysis.]

**S II-B3d**

[M…050-078. Dubois 1892d. Second-quarter report (dated 28 July, 1892).]

Towards the middle of May, the abandoned [Trinil] excavations of last year because of high water … were resumed. First, we had to remove large quantities of sand that were carried in by the river during the West monsoon. Then the river banks [including the left bank, see June memorandum, -B3c, above] were dug into so that after the water sufficiently lowered, a large surface area of the fossil-rich deeper bed [LB] would be obtained [on the left bank].

Because of this preparatory work, the harvest of bones was only similarly plentiful as last year's during the last days of June. Amongst these [new finds] were those of a deer species [***Axis lydekkeri***] that has been buried here in large numbers and the small ox-like antelope [***Duboisia santeng***], as well as ***Stegodon*** [***trigonocephalus***], important specimens to better determine the species. Several vertebrae and ribs of a large ox [***Bibos palaesondaicus***] found in their natural relative position [partially articulated], again provided proof that the bones are still at the spot where they originally came to rest.

However, other facts were observed that seem to give an explanation that the bones are generally isolated and widely distributed and usually broken. Separation by mechanical action of flowing water is not likely because they look well preserved (except for the fractures). It is also hard to imagine that this would lead to breaking of so many bones. Only animal violence could tear those bones apart, break them and distribute them, while they were still surrounded by soft tissue and articulated (and thus be protected against wear at the bottom of the current while it pushed them along). In addition, proof is provided by a fossilized piece of aorta filled with sandstone. However, in no case were the usually recognizable signs of the teeth of land predators undoubtedly observed, instead many breaks of the long bones were seen caused by dentitions large than those of the strongest carnivores. The bones are in fresh water deposits and it is logical to hold large crocodiles responsible, remains of which are found so frequently amongst the other animal remains (primarily teeth). Round holes in some softer bones appear to actually correspond to the cone shaped teeth of these gluttonous water monsters.

But these reptiles cannot be the cause of death of such large numbers of animals of many different species of which we now find the remains accumulated in fresh-water volcanic sand and lapilli now hardened to tuff and spread out over such large distances. At Trinil alone remains of many hundreds of deer were excavated over a few hundreds of square meters of surface area. And many other spots in the Kendeng have a similar such abundance. Only a catastrophe, and a volcanic catastrophe at that, comparable to, but on a larger scale than, those that accompanied eruptions of the Salak (1699), Galungung (1822) and Kelut (1848) [stratovolcanoes in western Java (Salak and Galungung) and East Java (Kelut; Figure 1b)] can explain the reasons for these accumulations and at the same time explain other observations. In the meantime, it appears that some light has been shed on the still obscure question of from which volcano this catastrophe originated.

Between Trinil and Pitu, we were, by means of dip directions of frequently occurring slanted drift beds (oblique diagonal lamination, current lamination, current bedding) of the tuffaceous rocks exposed by the Bengawan, able to determine the principal direction of the paleocurrents according to well-known rules. These [former currents] would have carried both the numerous animal carcasses that served as food for the crocodiles and the volcanic sand and lapilli in which these remains were subsequently buried. This principal direction is clearly SE to NW. The Wilis is located just about SE of the stretch along the Bengawan that is mentioned above and it seems more probable that this once mighty volcano more than any other has been responsible for the destruction of so many lives.

[Inserted at bottom of the page] The bone-bearing tuff beds immediately rest on a bluish-grey clay type at many spots on the banks of the Bengawan (on average at the elevation of the lowest water level) which indicates a time of stagnant or very slowly flowing water. However, the tuff and conglomerate beds themselves, which nowhere reach a thickness of 20 meter, show clear evidence for fairly fast-flowing water and having been deposited in relatively short time.

### S II-B4 (mid 1892)

### S II-B4a

[M…050 -079. July memorandum (dated 28 August, 1892).]

This month, the excavations at Trinil produced in addition to complementary material for the already well-known species, the following: *Verrucosis* (beautiful mandible with all molars and tusk plus canine tooth) [***Sus brachygnathus***; KdW may have mentioned the find in their August 22 letter; -A2b]; *Garialis*

(Pleistocene species of Java): skullcap closely related to *G. gangeticus* [***Gavialis bengawanicus***]; not yet determined *Sus* (lower jaw bone).

The tuff bed at Pitu in which last year bones of a turtle and of *Garialis* were found, proved, after additional excavating, to be fairly thin and did not produce any more fossils of significance. At several other spots along the Bengawan, such as near Porong, fossils were found in the river embankments but there the beds of interest were also only very thin, so that exploiting them would not pay off. Searches are now being continued farther along the river in order to discover richer find spots. In the meantime, for now enough good finds can be expected at Trinil, where we have reached the fossil rich beds, while the river also continues to remain at low level.

**S II-B4b** **Dubois notifies the Indies government of the *Anthropopithecus erectus* discovery**

[M…050-079. August memorandum (dated 23 September, 1892).]

Trinil must certainly be viewed as one of the richest find spots of remains of past mammals. The most abundant species continues to be the small ***Axis*** [***lydekkeri***] deer of which the antlers of as many as 50 individuals were collected this month, as well as several complete skulls ….

However, the most important find was a femur of the chimpanzee previously identified from a molar and skullcap, which evidently are from the same individual. [The antlers of the 50 deer might have come from the Femur I discovery trench, but excavations were also being conducted on the right bank, also; see KdW August letters.] Therewith, such an important fact has seen the light of day as perhaps never before has been gained by a paleontological find ….

This femur possesses in measurements, proportions and shape … such complete human characteristics (deviating only to an insignificant degree) that there can be no doubt that our *Anthropopithecus*, which is so clearly distinguished from the most advanced extant anthropoids by skull, had already adopted an erect posture, which is [generally now] regarded as an exclusive human trait. Thus, ***Anthropopithecus erectus*** … represents a line (or connection; *verbinding* in Dutch) that brings humans in closer relation to their living relatives amongst the mammals.

We don't have to emphasize the huge importance of finding further skeletal parts of this being, especially of the arm, the hand or foot. The chance that these will still be found is very high, because evidently the entire skeleton was buried in the volcanic tuff, although somewhat spread around.

The skull was located about 10 meters … from the femur and the molar was very close to the skull. These parts of the remains were found at the same level of the sediments at such distance from the erosional edge of the river that it is not likely that other parts of the skeleton have already been washed out and removed. At this time (September), a new section of bank wall is being removed so as to expose another portion of the fossil-bearing deeper bed in which these precious remains are to be expected.

[The memorandum was written the day before Dubois went to Trinil for the first time after having received KdW August 31 and September 7 letters explaining the provenience of Femur I (-A2j, -A2k). Dubois was at Trinil on September 24-25.]

**S II-B5 (mid 1892)**

**S II-B5a**

[M…050-079, right. September memorandum (dated 27 October, 1892).]

A new piece of embankment wall was excavated this month on the left shore of the Bengawan near Trinil, between where last year a Skullcap and a molar were found and this year a femur of an anthropoid, 15 meters upstream as defined by the paleo-current direction, with the goal of finding other portions of the skeleton of this extremely important species. The remains of this anthropoid have shown that it should definitely be recognized as one of the sorely lacking extinct transitional forms linking humans to their closest relatives amongst the mammals. This stretch of embankment extends over a length of 25 meters and is 5 – 7 meters wide and will have to be continued to what we now estimate to be a depth of 9 meters, since the level of the remains described above were found and other remains are expected below the water level during the east monsoon. The sedimentary material to be removed is only soft enough for actual digging near the surface, but for the most part we have a fairly hard sandstone-like andesitic tuff which can only be removed with pickaxes and crow bars. Because of this and the low number of forced laborers available this month (which even sank to below 50), the intended [stratigraphic] level of *A. erectus* in which also bones from other species occur, was not reached this month. That is also the reason that very few other finds were made.

[Below are a more-literal and an unedited translation and a scan of the original manuscript.]

September 1892. Missive 24 Oct. No. 17

Between and near the points where, in the left bank of the Bengawan, near Trinil, last year a Skullcap and a molar, and 15 m upstream according to the old stream this year the femur of an anthropoid, was found, which, by these parts with certainty has unveiled itself as one of the so sorely missed (extinct -- transition forms which connect man to its nearest relatives among the mammals, this month a new piece of the bank was dug up aiming to find other parts of the skeleton of this extraordinarily important form. This piece has a length of 25 m, is 5 - 7 m wide and will, as the level in which the intended remains have been found and are to be expected, and below the water level during the east-monsoon and has to be dug until 9 m of depth [that is: This piece of the excavation has a length of 25 M, a width of 5 - 7 M, and since the remains were found at a depth that lies below the water level of the east monsoon, these [parts] would be found at a depth of 9 m, anyhow]. Only the top part of this can be really dug, but mostly it is a relatively hard sandstone-like andesite tuff, which can only be removed slowly with pickaxe and crowbar. Because of this, and because of the low numbers of prisoner laborers available this month (which even went below the number of 50 -- the target level of *A. erectus,* which is also the richest in bones of other species, had not been reached by the end of the month; hence very few other finds were being made.

[Below is a scan of Dubois' handwritten draft of his September 1892 memorandum which is in the Naturalis document "Report on Paleontological Investigations…" (-B1, above)]

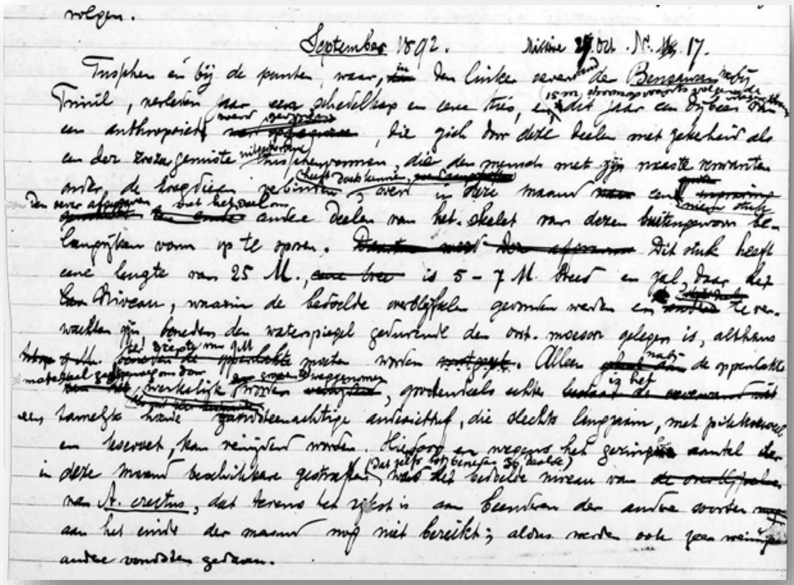

**S II-B5b**

[M…050-080/081. Dubois, 1893a (his Third Quarter 1982 report was published in the First Addendum to the Java Bode on Saturday, 4 February 1893, Nr. 30).]

[p. 88] The most important find, however, was made in the month of August of the left femur of our *Anthropopithecus*, whose existence became known a year ago by a molar and skullcap. This femur was positioned in the same bed in which the other two specimens had been found, but 15 meters upstream with respect to the direction of the former current that had deposited the tuff material. From this observation and from my comparative [anatomical] research it has turned out that the three skeletal parts have belonged to the same individual, probably a female who almost certainly had already reached an advanced age.

[Faintly handwritten at bottom of the page in the "Report on Paleontological … Dubois" is] 15m wide, 5m long; about 1/1,000,000 of the entire terrain when one reduces it 1) to a similarly rich surface [area] over which no human was found, but only a mandible fragment of a higher erect anthropoid. 2) Also many other bones that belong together very often at the same site; jaw and tusks of elephant. 3) At no time complete skeletons, although no violence …. of now soft parts rocks. [The report continued on the next handwritten page.]

[p. 89] Because of this find, an important and also surprising fact has come to light. The Java *Anthropopithecus* which is distinguished by its skull as the highest anthropoid known to date, had already fully adopted an erect posture, which has always been viewed to be the exclusive advantage of humans. This means that, with this early Pleistocene anthropoid of our island, the first transitional species has become known which unquestionably links humans to their closest living relatives amongst the mammals. In all three of the discovered skeletal parts *Anthropopithecus erectus* Eug. Dubois approaches humans more so than any other anthropoid. However, most of this is, through [the nature of] the femur, which fully corresponds to the statement previously made by Lamarck and later expanded on by Darwin and others, that the first step for our antecessors towards becoming human was adoption of an erect posture.

The skullcap, which in its general shape as well as in other aspects relates to the *Anthropopithecus* ….

In the upper 3rd true molar, the rear cusps of the crown are even more reduced than is the case with *A. troglodytes* …. In terms of measurements and shape, the femur shows remarkable correspondence to the human thigh bone and is only distinguished from it by less essential peculiarities. It is just as long and slim, thereby strongly deviating from femurs of the living great anthropoids but less so with that of the chimpanzee. Its length is 455 mm …. [translation of the remainder of p. 89 is omitted]

[p. 90] This creature was thus in no way capable to climb on trees the way chimpanzees, gorillas and orangutans do with their short legs and gripping feet and their long arms and very heavy and long torso. …

From this analysis of the femur, it follows without any doubt whatsoever that *Anthropopithecus* of Java had the same posture when standing and walking as humans do. …

[p. 91] While *A. erectus* thus exclusively used his legs for locomotion, it should be accepted, in view of the division of labor between front and rear limbs, that the hand was already more complete than this organ is in anthropoids. Because once the hands were freed, evolution of this most perfect and universal of all tools, this most reliable sense organ, must have progressed very rapidly and partly as a result of and partly as a reason for that [evolution] of the brain. This is in agreement with an upper torso that must have had human proportions, as well as the higher doming of the brain pan and reduction in the dentition. The reason for this is that this creature had the need and ability to find food and use other (artificial) weapons for defense than his teeth. …

And this then delivers actual proof that the Indies were the cradle of the human race, which some have already long suspected.

[Dubois is not known to have corresponded with professional colleagues about this announcement (however, see an earlier letter to Weber, below, at –E1c).]

**S II-B5c**

[M…050-080. October memorandum (dated 25 November, 1892).]
The desired level with the bonebed [LB] was finally encountered at both ends of the elongated pit [25-m Trench, in addition to the Femur I discovery area, which had been dug into the PFZ in August]. A number of fossils were dug out [from the newly exposed LB], amongst which some very nice ones, but still no other parts of the *Anthropopithecus*. A small but important find was that of a molar of a lower ape, a M.3 [***Macacca*** sp.] …. repeated rising and lowering of the river water, always accompanied by huge masses of sand being deposited on top of our work, forced us to finally abandon the work on November 16th, after only excavating [away] about 1/5th of the level of interest [PFZ], and still not having encountered any of the skeletal parts we were looking for. Further searching for these most important remains in the [80%] unfinished portion can now only be resumed during the next East monsoon [low-water-level season]…

[KdW had reported on November 9, 1892, that they reached "20cm into the target bone layer" [PFZ] at the corners of the trench (-A2q). Among seven Trinil macaque teeth in the Dubois Collection, one (no. 3789) is a *Macaca* $M_3$ dextra (Hooijer, 1962: 50, Figs. 4 and 5) that "was taken in 1892 from the trench of 25 m [25-m Trench], indicated by a note in the box containing the tooth" (de Vos and Sondaar, 1982: 49). Also, -B5e, below. De Vos (1989: 227) also reported that D.C. 3789 came from the "lowest level, ½m below pe," presumably the level at which the *Pithecanthropus erectus* fossils were found. Stremme (1911: 140-141, plate XVII figs. 9-11) described another $M_3$.]

**S II-B5e**

[M…050-081/082. November memorandum (dated 31 December, 1892), December memorandum (M…050-082), and below, Fourth-quarter report (dated 13 January, 1893).]

The excavations at Trinil were continued to as long as the middle of November with the principal goal of finding further remains of the curious anthropoid. This excavation was done in the vicinity of where the skull cap, a molar and the left femur of this transitional form were found [Figure 3a, main text]. However, when after long labor and finally exposure of a new portion of the bed in which these remains may still be expected and we began our search, our hopes were dashed by continued high water levels due to the heavy rains, unusual for this East Monsoon. Among the fossils of other species was also a molar that probably belonged to a probable *Cynocephalus*.*

After the middle of November, systematic searching began in the terrain east of the Pandan [eastern Kendeng Hills, Figure 1b, main text] …. In the area between Bangle and Djeruk …, the fossils were less attractive and similar to what they are at other low spots west of the Pandan … they were covered with difficult-to-remove hard lime concretions. These calcium carbonate deposits are here also intercalated in the tuffs as thin crossing layers which can often been found spread out on the ground as the well-known irregular beet-shaped grayish-white stones as a result of weathering of the tuffs.

{* [Theunissen (1989: 92) noted] "When discussing the molar {1891 Molar in the 1894 monograph} Dubois introduced a new find. This was also a tooth, the second molar from the left side of the upper jaw {1892 Molar}. It was unearthed in October 1892, no more than three metres away from the skullcap [Skullcap]. Dubois claimed that the piece had at first escaped his notice because it lay beneath a large number of other teeth dug up from the same place. This statement notwithstanding, it was probably the molar he had originally classified as belonging to a *Cynocephalus*, a baboon," or other Cercopithecine.}

**S II-B5f** [Reactions in the press of 1893 to Dubois' announcement of *Anthropopithecus erectus*]

[(**i.**) A person using the name Homo erectus (1893) wrote in the "Batavaasch Nieuwsblad" newspaper of February 6, 1893; M…043-305; Shipman, 2001]

IN PURSUANCE OF PALAEONTOLOGICAL INVESTIGATIONS OF JAVA

(Report to the Mining Works, third quarter 1892)

It is well known that etymologists sometimes explain the origin of words through most surprising derivations and on such occasions produce results that are astounding to the uninitiated, but if we compare these attempts with what some paleontologists dare to do with reference to fossils, then their endeavor appears to be child's play.

This flashed across my mind as I read the article mentioned at the head of this piece.

For some time, the military surgeon Eug. Dubois busies himself with paleontological investigations and, because of his predilection for such studies, works with exceptional diligence and assiduity. He is totally absorbed by his work, so to speak. As a firm Darwinist, he dreams of making a discovery which the great master of evolution will greet with joy. Namely, the discovery of the until-recently missing link between the animal world and man.

Should this be taken amiss? I do not believe so. At present Darwinism is the backbone of the education of most high school graduates. The heavy facts that are brought up against Darwin's theory by the most competent authorities -- these leave them cold. Examine their libraries and, ten to one, you will not find a single paper in which Darwin's theory is opposed. It is old-fashioned to think differently, their teachers have told them.

I fear, however, that this time the Darwinian outlook of the esteemed Mr. Dubois has played a trick on him, a danger that an impartial observer would have escaped.

What then is the case?

In the Kediri region, in volcanic beds, he found some time ago a skullcap, not an undamaged skull, but a fragment, along with a loose molar. In August 1892 he found, 15 meters from the place where the skullcap and tooth had been lying, a thighbone with a striking resemblance in measurements and shape to the bone of the leg that supports humans.

Instead of thinking of the remains of a human skeleton (Dubois had already demonstrated the presence of a primitive race here in Java, more like the Papuan race than the present population, in a previous report) he makes comparisons (the details of which are not known yet) with a more surprising result than the most abstruse etymologist has ever put forward!

Given: a molar and one skull fragment found together as well as a left femur, that has been found 15 meters further away.

Until now, Dubois has been skimming the article but he stops when he reaches these words and reads them aloud. "A 'skull fragment'?" he repeats, outraged. "A fragment? I have the better part of a skull, lacking only the face. I have as much of the skull as the original Neanderthal fossil, more than Lydekker's Anthropopithecus. This is no fragment! I have shown it to you, Assistant Resident, haven't I?" "Ja, ja, two times," the official replies a little anxiously. He fears Dubois will insist on showing him all of the thousands of fossils crowded onto the veranda. "This author has set out to make me sound like a fool! What else does he say? Here, ja, here is the place." Dubois reads on aloud. Question: What does this mean?

A non-darwinist would scratch himself through his fur before he would propose a genetic link between the monkey skull and the monkey molar and the femur, which has a close speaking acquaintance with a human femur.

Not so the esteemed Mr. Dubois.

"His sarcasm is really too much," protests Dubois and the Assistant Resident nods in agreement. "'The esteemed Mr. Dubois'! What a backhanded compliment. Now, what else does this scoundrel say?"

With the data mentioned above, he thinks that the theories of Darwin and Lamarck have been confirmed, that the first step on the road of the incarnation of our ancestors has been acquired with erect posture (page 11); That molar, skull fragment, and left thigh bone were once part of an upright female apeman, named Anthropopithecus erectus Eug. Dubois (page 11.) That this creature fed differently from the present-day anthropomorphic apes, despising tree-climbing and even carrying artificial weapons (page 14). Finally, according to the esteemed Mr. Dubois, this is the actual evidence that the Indies were the cradle of the human genus (page 14).

Whew, that's it!  Why is this animal given such a beautiful name? Why wasn't he named Hanoman (*) communis of the family Hanomanaceae....? The historian would have gained something from the find, but Anthropopithecus erectus!

No, I am afraid that the esteemed Mr. Dubois, prejudiced because he has completely swallowed darwinism, has gone too far, and has constructed a connection between the human femur and the monkey skull and molar where none has ever existed. If humans also lived, at the time when the cataclysmic eruption buried so many dead animals in volcanic tuff, no one would be surprised that humans also would be killed and their skeletons or remains of severed body parts would be preserved in the tuff.

In the meantime, this publication of Dr. Dubois will create a furor, especially in the "Land of Intellectuals," and it appears to me that the facts must be reviewed by an impartial committee of experts, before the government should endorse such a report.

"Oh, the man is sly, very sly indeed," Dubois asserts. "And who is he, anyway?"  Dubois glances down at the bottom of the page, to the signature, and reads it, too, aloud.

Signed,
Homo Erectus
Batavia 3 February 1893

[(*) Hanoman is a white monkey, according to the Javanese mythology of Dhewi Handjani, that is also called Dhoyopati or Boyosoeto. Note of the editorial staff.]

[(ii). (Bolsius 1893)

Nr. 20086 Nieuwe Amsterdamsche Courant   66 e jaargang

ALGEMEEN HANDELSBLAD

Friday 28 April 1893.

Phantasm about fossils delivered by Dr. Eug. Dubois on Java.

We all know of phantasms in music, in literature, in the arts such as drawing and painting etc.

Are well executed phantasms welcome – always and everywhere? Even if there is no evidence? Even if there are questions about previously never proven facts?

The phantasm with which Horatius introduces his *Ars poetica* will probably never deliver proof that, during this painter's life, humans or animals existed or had existed the way he had them depicted.

This was one of the thoughts we had upon repeated exposure to the report by Dr. Eug. Dubois of investigations in our Indies: twice extensively (in *Verslag van het Mijnwezen 3 de kwartaal* 1892 and in *Javabode* 4 Feb last, 1st addition), once in the form of a quintessential summary (*Algemeen Handelsblad* 12 March last) and once more as a critical newspaper column (in *Bataviaasch Nieuwsblad*, dated 3 February, signed by *"Homo erectus"*)

In order not to mix sound concepts, scientific data and phantasm pronouncements and so not play into the cards of the nonexperts, we wish to briefly discuss the phantasms of Mr. Eug. Dubois. Since briefness is required we will try to adhere to that.

I. *The positive facts*.

*a*. Dr. Eug. Dubois found among other fossils a left femur in August 1892 at Trinil. A very detailed description is given by "his honor" in which he states up to 15 (read fifteen) times in various wording about all sorts of components and peculiarities that "This femur is just like a human femur."

*b*. A year earlier Dr. Dubois had also found a skull cap and a separate loose lying molar. After his analysis of both specimens the Dr's pronouncement was that "Both belonged to an ape type but still unknown species, but it nevertheless was an *Anthropopithecus* or Anthropoïd.

*c*. The finds of 1891 and of 1892 were made at the same level in the tuff at 15-meters distance from each other.

These are three positive facts.

II *The phantasma explanations*. Dr. Dubois says that the three specimens found at 15-meter distance from each other unquestionable belong to the same individual. Why? Because 1° All three were found at the same level of layers; because 2° the skull cap and molar are still very ape-like and the femur is very human-like and 3° because Lamarck and Darwin indicate that the link between humans and apes is much more human like than we have ever found so far.

I am thoroughly convinced that many who read this reasoning involuntarily must think that I write this nonsense on the account of Dr. Dubois. Well now, my greatest pleasure would be if someone reviews the original [script] and extract other proof from it and make this public so as to shame me. I would still like to add that: after having read the report by Dr. Dubois about what "his honor" calls *Anthropopithecus erectus* at least six times, I recently (12 April) gave a brief announcement of this fact at the meeting of the "Société scientifique de Bruxelles." Among those present were A. de Lapparent, the well known geology professor at Paris and L. Dollo, conservator of the Museum of Natural History in Brussels. The meeting was thus not just attended by dilettantes. Should I even mention that the phantasm declarations by Mr. Dubois caused great hilarity?

III. *Conclusions based on phantasm*.

Now that Dr. Eug. Dubois has screwed together an ape skull, an ape molar and a human femur to construct his *Anthropopithecus erectus*, logical conclusions naturally follow. It could well be possible that *if* there were no doubt about the prior explanations (!) and *if* the femur belonged together with the skull, this animal would have walked upright. If this animal walked upright, it would not have climbed trees. *If* this animal did not climb trees, it would have fed itself differently than tree climbing apes etc. etc. etc.

Dr. Dubois simply reasons those *ifs* away by saying: my explanations are the real ones.

We won't even talk about Eug. Dubois' conclusion that *Anthropopithecus erectus* must have used "artificial weapons" and all the consequences thereof.

IV. *Questions to Dr. Eug. Dubois*. Is it certain that by further excavation no more "surprising" finds could be made in the region where the skull cap, molar and femur were located? If for instance at night or in the morning a new femur were found, a bit less human-like than the previous one? If this were by coincidence a right femur, would your *Anthropopithecus* not become lame? Or if a second left femur were found, almost entirely like the first one, must your *Anthropopithecus* then not be satisfied with having two left legs? Or if a human skull were found within or outside the distance of 15 meter, would *Anthropopithecus* then receive two heads, one ape-like and the other human-like? Or is there a law that says: genetic (original) relationships only exist for specimens that are within 15 meter from each other and to be excavated within a year from each other?

We would lose our breath before coming to the end of our questions, because Dr. Eug. Dubois would have to prove that 1° at this moment no other bone or leg exists within whatever distance from skull and femur to which one of these two belong more so to than they do to each other. 2° that already in the past something of what the volcanic eruptions had buried got lost so that whatever had belonged together still is together.

This space, willingly allotted, prevents us from expounding further and perhaps clearer.

About the conclusion of Dr. Dubois: "and so has become known the first transitional form linking humans more closely with their closest living relatives…… and so has actual proof been delivered……… that the Indies has been the cradle of the human race", about all this we wish to say nothing more than to repeat the words spoken by Mr. L. Dollo at the meeting in Brussels: "C'est ainsi qu'on rend notre science ridicule" [It is in this way that our sciences are made to ridicule].

H. BOLSIUS
Member of the Netherlands Biological Society.

OUDENBOSCH , 21 April 1893.

[The article "Palaeontologische Phantasie" appeared earlier on March 10, 1893, in the 'Sumatra-courant,' and later on August 25, 1893, in the Batavia (now Jakarta, Indonesia) newspaper, 'Bataviaasch Nieuwsblad.']

**S II-B6 (1893)**

**S II-B6a**

[M…050 -086. May memorandum, 1893.]

After searching the entire bone bed region east of the Pandan [and] near Kedungbrubus …. but especially because of the dense growth of plants on the soil, very few fossils were collected here. On May 30, search activities were again transferred to Trinil, where first of all repair and new construction for the forced labor quarters and storage sheds was started.

[The First-quarter report for 1893 (M…050-084) noted "continued with good results east of the Pandan" and concluded that the uplift of Pandan area post-dated the deposition of the fossiliferous tuffaceous strata in which Dubois was collecting.]

**S II-B6b**

[M…050 -086. June memorandum (dated 21 July, 1893).]

Almost the entire first half of this month had to be spent on repair and construction of the forced labor quarters and sheds at Trinil. After that, excavation commenced. Because of the almost permanently high-water level of the Bengawan [Solo River], the rich deeper layer on the left bank which had to be abandoned at the beginning of the previous west monsoon [November 16, 1892], was not yet accessible. Instead, we started a new stretch of the bank, 40 meters in length and 5 meters in width adjacent to it and further landwards. Soon, this work progressed extremely slowly because of the severe hardness of the rocks and the relatively low number (40) of available forced laborers. So as to remove this shallower fossil-poor rock mass as quickly as possible, an additional 10 to 20 paid coolies were temporarily employed.

**S II-B6c**

[(i). M…050 -085. Second-quarter report, 1893.]

….. Towards the end of May, work that had been halted last year was resumed at Trinil (Dept. Ngawi), first of all as construction of the necessary barracks and storage sheds, followed by actual excavations. Because of the continued nearly permanently high-water level of the Bengawan [Solo River], we could not yet access the deeper [previously] exposed rich-bonebed [LB] that we had abandoned at the start of the previous west monsoon. For that reason we instead initiated excavation in a new strip of riverbank more landwards and adjacent to it, with dimensions of 40 m in length and 5 meters wide. This work progressed extremely slowly because of the extraordinarily severe hardness of the rocks, with the result that during this quarter the mentioned rich bonebed did not yet become exposed.

[(ii). M…050 -086. July memorandum (dated 27 September, 1893).]

During this month excavations at Trinil again progressed extremely slowly because of the severe hardness of the rock to be removed [above the LB], although paid coolies were now employed besides the detainees. For this reason and because of the relative paucity of fossils in the beds encountered, only little paleontological material was collected this month. [Dubois visited Trinil on July 18-20.]

**S II-B6d**

[M…050 -087. August memorandum.]

…. At Trinil, the deeper fossil-bearing layer [LB] was partially exposed [in the 40-m Trench]. But the severe hardness of the rocks continued to be a handicap towards speedy progress of the work and also seriously impeded proper retrieval of the fossils by keeping them intact. Another great difficulty was the present abundance of encroaching water. Nevertheless, some attractive specimens were obtained, most of them of a small deer species [***Axis lydekkeri***] of which many hundreds are buried at Trinil and also of *Boselapahus* [***Duboisia santeng***], ***Stegodon*** [***trigonocephalus***] etc. In addition, the first almost complete skull of the previously discovered crocodile was found. It was now possible at this time to assign this species to a position in the system as far as available comparative material permitted; the species cannot be distinguished from *Crocodylus porosus* [later attributed to a new species ***C. siamensis***].

According to a remark made [during Dubois' visit to Buitenzorg (Bogor)] by the Head Engineer and Chief of geological mapping of Java, Dr. R.D.M. Verbeek [on] the 16th of this month, … de Winter was transferred to Tampung Bapang [Sangiran Dome] … to keep oversight of the planned excavations. During this month, they had little success and primarily remains of *Stegodon* were found. In any case, in regard to the distribution of the relevant formation [the bonebed] the presence of it near Surakarta is significant.

**S II-B6e**

[M…050-087. September memorandum with Third-quarter report (dated 8 November, 1893).]

Excavation near Trinil, on the left bank of the Bengawan [40-m Trench] was continued up to a depth of 1 to 1¾ meters below the level at which the skull and femur of *Anthropopithecus* were found previously at 11 or 12 meters below ground level. At that depth a black coaly claystone layer was reached, which turns out to be the underlying formation to the volcanic bone tuffs here and in other places along the Bengawan and sits about 3 meters below river level during the east monsoon. Skeletal parts of previously discovered Kendeng vertebrates were found up to this claystone, but among them were no further parts of the Java *Anthropopithcus*. Also, last year's excavation [25-m Trench] was resumed in which the skull and femur of the above species were found which had been abandoned, only partially worked [through the PFZ], during the start of the west monsoon. First of all, much sand and silt had to be removed since that had been deposited by the river during the West Monsoon period.

Along the left shore of the Kali Tjemoro near Surakarta [now the city of Solo], mainly east of Bapang and also at Tandjung along the right shore of this small river and near Sangiran [geologically at Sangiran Dome], small quantities of not especially attractive [fossil] bones were found of *Stegodon*, *Elephas*, *Bubalus*, *Bibos*, *Boselaphus*, *Cervus*, *Sus*, *Hippopotamus*, *Crocodylus* and a land turtle. These were sufficiently varied to demonstrate the Kendeng fauna character. West as well as east of Bapang very nice "oblique lamination" was observed in the relevant volcanic tuffs. These are so clear here within the horizontally bedded structure, that there is no doubt about their deposition in water. The direction of the [paleo] current must have been from WSW (coming from the direction of the Merapi). The presence of fresh water mollusks of the same kinds as those found at Trinil and Tritik [an area east of Mt. Pandan] indicates that the tuffs were deposited in fresh water and that they are equivalent to those of the Kendeng. These were encountered in large numbers primarily along a small brook above Tjenklik on the left shore of the Tjemoro [Sangiran Dome]. They appear to be mixed with marine shells, but that is because they have been washed into fissures from the very locally outcropping underlying marine layer.

[Dubois had visited Trinil on September 14-15.]

**S II-B6f**

[M…050 -085/086. Dubois 1894b. Third-quarter report (dated 8 November, 1893).]

The new excavation at Trinil on the left river bank of the Bengawan was continued up to a depth of 1 to 1.75 meters below the level [LB] where previously the skull and femur of *Anthropopithecus erectus* had been found and where the black coaly clay bed was reached. This [contact of the LB with the subjacent clay bed] is 11 to 12 meters below ground level and 3 meters below river level during the East Monsoon. This black clay bed proves to be the underlying formation to the bone-bearing volcanic tuffs here, but also elsewhere along the Bengawan. Up to the level of this clay bed, skeletal remains of already known Kendeng vertebrates were found in these tuffs, first of all many antlers of the small *Axis*-like deer species [***A. lydekkeri***] and also remains of ***Stegodon*** [***trigonocephalus***], *Boselaphus* [***Duboisia santeng***], ***Bubalus*** [***palaeosondaicus***], *Hyaena* etc. Also [we found] the first almost complete skull of a crocodile [***Crocodylus siamensis***] …. However, among these remains were none of the Java *Anthropopithecus*. After this [40-m Trench was completed], work was resumed in the old excavation that had to be abandoned last year because of the commencement of the West Monsoon, and in which the skull and femur of the above anthropoid had been found [the 25-m Trench].

About 10 paal northeast of the station of the capital Surakarta, several not particularly nice fossils were collected primarily east of Bapang on the left bank of Kali Tjemoro and at Tandjung on the right bank of this small river as well as at Sangiran. These were of *Stegodon, Elephas, Hippopotamus, Sus, Bibos, Boselaphus, Cervus, Crocodylus* and a land turtle, enough to show that they belong to the *Kendeng* fauna.

Oblique lamination can be observed in the volcanic tuffs west as well as east of Bapang [at Sangiran Dome]. These and the horizontal bedding structures are so clear that there is no doubt of the fact that the sedimentary rocks were deposited sub-aqueously. Judging by these slanted oblique laminations, the flow must have come from the WSW (from the Merapi). The presence of fresh water mollusks of the same kinds as those found at Trinil and Tritik [east of Kedungbrubus and Gunng Pandan] indicate that the tuffs have been deposited in fresh water and are equivalent to those of the Kendeng. Numerous specimens of these were primarily found along a small stream on the north side of Tjenklik [inserted: more than ½km NW of B.T.] on the left bank of the Tjemoro, seemingly mixed with sea shells, because they have been washed into cracks in the underlying marine layer that is locally found in outcrop.

**S II-B6g**

[M…050 -088. October memorandum (dated 23 December, 1893).]

Work on the excavation of last year at Trinil, described in the previous monthly report, was continued. However, the problems caused by repeated flooding and the steady flow of ground water into the pit were so great that progress was extremely slow. No fossils of new species or further remains of *Anthropopithecus* were found.

[Dubois visited Trinil October 19-22; see his notes on examining the 40-m Trench in –C5.]

**S II-B6h** **Dubois informs the Indies government that he is abandoning Trinil excavations**

[M…050 -088. November memorandum (dated 23 December, 1893).]

… (with a small sketch drawing of the Dubois monument) …. Notwithstanding the extremely severe problems caused by the rising river, *we were able to essentially dig away the entire bone-bearing bed in the excavation pit [25-m and 40-m Trenches] before the 26th when the work [site] became hopelessly inundated.* The collected fossils from this period were very carefully analyzed, but it turned out that among them none were of the hoped for additional parts of the curious *Anthropopithecus*. Although they almost certainly must once have been present, it seems that they have thus been washed away during formation of the river bed, together with a large portion of the bone-rich tuff. Further searching for this transitional form appears therefore to be fruitless. However, in order to facilitate recognition of the find spot by later researchers a small pillar was erected before abandoning it [the site] at a point onshore, east of the Klitik settlement that could easily be seen from the river as well as from the land. This pillar has a black marble plate attached with the beginning letters of the further to be named anthropoid *Pithecanthropus erectus*, together with the required orientation and the years in which the investigations have taken place. ….

[Dubois visited Trinil November 11-15. KdW's letter of July 14, 1893, has a sketch of the monument bearing the inscription "*P.e.*" (-A3a).]

**S II-B6i**

[M…050 -088. Fourth-quarter report, 1893. Dubois, 1894d.]

The difficulties with repeated inundations and the steady flow of ground water into the pit that hindered excavations at Trinil were already so severe in October that they frequently had to be interrupted for considerable periods of time and progress was only extremely slow. We were nevertheless able to just about completely remove the bone bed in the excavation pit before November 26th when the work site became hopelessly inundated.

The collected fossils were carefully examined, but it turned out that none of the still anticipated parts of the curious anthropoid (for which the excavations at that location were primarily instigated) were amongst them. Having once almost certainly been present although quite spread out, as is generally the case with other species, these bones must already have been washed away during formation of the current river bed, together with a large portion of the bone rich tuff. It appears therefore to be fruitless to continue searching for this transitional form.

But, to facilitate recognition of the find spot for later investigators, a small pillar was erected at a spot on the right bank of the river, east of the dukuan [<Bahasa] Klitik that could easily be spotted from the river as well as from the land. This small pillar has a black marble plate attached bearing the beginning letters of the further to be named anthropoid *Pithecanthropus erectus* as well as the necessary indications of orientation and the years in which the investigations have taken place.

North of the Kendeng…..

**S II-B6j**

[Dubois'(1894a: 1-2; also, 1895a) monograph on *Pithecanthropus erectus* has a short account of the geological context of the fossils, which follows. The Forward is dated January 1894, three months after Dubois visits Trinil in September 14-19, October 19-22 and November 12-15, 1893. Kriele remained at the site until early December.]

Along the left bank of the Bengawan [Solo River], about 1 meter below the dry period water level of the river and 12 to 15 meters below the surface, into which this river has dug its bed, first a tooth was discovered in September 1891, a right-side M.3 [1891 Molar (Trinil 1)], which was attributed to a species that is related to *Anthropopithecus troglodytes*, but which is larger in size. A month later, only 1 meter away from where the tooth had been buried, a skullcap [Skullcap (Trinil 2)] was found at the same level [PFZ], which apparently belonged to the same individual and because of its [the Skullcap's] significant size and its high doming, clearly approaches resemblance to a human type much more than it does to the living chimpanzee.

Finally the left femur [Femur I (Trinil 3)] was excavated in August 1892, again at the same level [PFZ; Figure 2a, main text] and during the dry period, about 15 meters upstream in the direction of the current that during Pleistocene time had washed out this animal carcass along with the tuff material [ "upstream" refers a paleogeographic direction of transport of the fossils, is based upon paleo-current measurements and is the opposite of the flow direction of the present-day Solo River]. This femur points even more to human like characteristics than the other parts. During the dry period of 1893 further excavations were undertaken in the vicinity of the earlier discovery locations to look for other remains of this species. However, these efforts remained fruitless.

{Footnote: The tooth was mentioned in my report to the Director of the Department of Education, Culture and Industry for the third Quarter of 1891. The skullcap was mentioned in the report for the fourth Quarter of the same year. Both were included in the Report of the Mining Bureau [*Verslag van het Mijnwezen*] for the year 1891. …}. ["the Director…" is *Directeur van Onderwijs, Eredienst en Nijverheld* in Dutch.]

[Dubois' footnote specified the first publications to mention the discoveries, although the Reports were published anonymously, rather than in his name.]

[Theunissen, 1989 (Chapter 3) has an account of Dubois' preparation of the monograph.]

## S II-B7 (1894)

### S II-B7a

[M…050-089/090. First-quarter report, 1894. Dubois 1894d.]

After collection of fossil remains of vertebrates had been stopped [at Trinil and elsewhere], attention was now specifically focused on the geologic formation in which they occur [this is largely the Kabuh Formation of Duyfjes (1936)].

The formation is better developed in the surroundings of Kedungbrubus … than in other locations [see map of Dubois work area in S I Figure 21]. There, inclination of the beds to the SW was previously observed at several places (see report for the 1st quarter 1893, p. 13). Since more specific investigations related to this were initiated by means of excavations it turned out that this [dip] is widespread and is consistent.

However, a better insight into this was obtained from traverses through the ravines of the Kali Brubus and her side branches, the Kali Ngetos and Kali Butak. The strike of the beds is there generally and consistently N 55º [crossed out] W and the dip 8º [crossed out] SW. In the direction [in margin: 5º 15'] perpendicular to the dip of the beds (exactly NE to N) [crossed out to NE] it [the dipping sequence] can be followed from Kedungbrubus to a point at a distance of 3/4km [in margin: 1230m].

That is where the [first of the] underlying andesite breccia is exposed [apparently Duyfjes' upper breccia; S I Figure 21], which in this part of the Kendeng sits between the formation and the marine sedimentary rocks of this range of hills. This breccia, which contains huge blocks of andesite (up to 2 m across) as well as fine gravel, is not covered by other sediment all the way up to the crest of the Kendeng [there is no unconformity within the sequence]. Only just below [down slope from the settlement of] Butak, about 450m [crossed out: in margin 250m] above the point where the bone bearing tuff ends northeast of Kedung Brubus and at a horizontal distance of more than 3km, a small portion has been preserved of only a few hundred meters in length, fifty meters in width and a few meters thick. It [the area of exposure] forms a pedestal on which a small hill emerges, comprised of the same tuff. About 30 years ago, Raden Saléh excavated fossil bones next to it. In the past only remains of land animals were found [Saléh 1867].

During the current investigations in the same location, teeth of crocodiles as well as fresh water mollusks were found, while excavation of the small hill demonstrated that it consists of material that must have been deposited in water [Butak bonebed, Tables 5 and 6-D, main text] . The same strike and dip as those of the tuffs near Kedungbrubus could here be determined with certainty. The placement of the small hill with respect to a point near Kedungbrubus where the deepest tuff beds are exposed aligns perfectly in the direction of dip of the beds. Taking the above horizontal distance between these points and an elevation difference of 450m [crossed out: in margin 250m], it is easily determined through trigonometric calculations that this deepest portion of the bone tuff [from which the crocodile teeth were found] is located in exact alignment with Butak small hill. The material is also the same in both places: claystone like tuff, covered by similar sandstone-like beds. There can thus not be any doubt about their perfect correlation [Note: Field observations on this are on pages 270-271 of Dubois' notebook].

The tuff formation [referring largely to the Kabuh Formation of Duyfjes, S I Figure 21] is distinctly and beautifully bedded near Kedungbrubus and consists primarily of fine or a bit coarser sandstone-like andesite tuff, alternating with clay-stone beds of greater or lesser thickness and only a few thin beds of conglomerate. This formation contains fossil bones in its deepest as well as its shallowest portions. A thickness can be calculated from its base to Kedungbrubus of about 140 m [crossed out: 100m after reducing by 14½ m of elevation difference]. The bone bearing tuffs can be further followed for at least 3 km in a southwesterly direction (perpendicular to the strike of the beds). This means that, if no faults are present, of which none have been found so far, and with a dip of 8º, the total thickness of the formation in this portion of the Kendeng must be more than 500 m.

Almost 40km further east, near Bangle (District Lenkong of the Department Ngandjuk in the Residency Kediri) the same strike and similar dip of about the same magnitude were observed.

[Pages 265 to 273 in Dubois' "Notes related to the paleontological investigations…." (M…046-153 to -157) have numerous observations and analyses concerning the stratigraphy of the greater Kedungbrubus area. Also, Albers and de Vos 2010 (especially pages 64-72) have photographs that Dubois took in the vicinity of Kedungbrubus and at the small hill near Butak.]

[The Dubois Collection (D.C.) from the Kedung Brubus area (referred to in Tables 5 and 6-C, main text) consists of the following taxa (listed in approximate order of the number of identified specimens and minimum number of individuals from Storm 2012): [a] *Stegodon trigonocephalus* (NISP of 129, MNI of 9); [b] *Bibos palaesondaicus* (28, 7); [c] *Dubois santeng* (26, 5); [d] *Axis lydekkeri* (39, 4); [e] *Bubalis palaeokerabau* (25, 4); [f] *Rhinoceros sondaicus* (26, 3); [g] *Rhinoceros unicornis kendengindicus* (13, 3); [h] *Elephas hysudrindicus* (23, 2); [i] *Tapirus indicus* (3, 2); [j] *Hexaprotodon sivajavanicus* (35, 1); [k] *Manis palaeojavanica* (29, 1); [l] *Panthera tigris* (15, 1); [m] *Epileptobos groeneveldtii* (9, 1); [n] *Sus brachygnathus* (4, 1); [o] *Muntiacus muntjac* (2, 1); [p] *Rusa* sp. (5,1); [q] *Lutrogale palaeojavanica* (1, 1); [r] *Crocuta brevirostris* (1, 1); [s] and *Homo erectus* (1, 1).]

[The D.C. contains the following specimens from "Boetak, Java" (referred to in Tables 5 and 6-D, main text, and S I Figure 21). This assemblage is separate from the larger- and better-known one from Kedungbrubus. Also, Tables 5, note 3, and 6-D. The most-relevant specimens in the D.C. are the following: No. 1575A *Crocodilus* "*ossifragus* Dubois, 1908" Teeth; 1575B *Cervus* (*Axis*) "*lydekkeri* Martin, 1892" M1/M2 superior; 1575C *Cervus* (*Axis*) "*lydekkeri* Martin, 1892" M1 or M2 inferior; 1575 *Cervus* (*Axis*) "lydekkeri Martin, 1892" M1/M2 superior; 1575E *Panthera* sp. indet. Phalange I; 1575F *Cervus* (*Axis*) "*lydekkeri* Martin, 1892" Antler fragment; 1575G *Cervus* (*Axis*) "*lydekkeri* Martin, 1892" Phalange II; 1575H *Stegodon* "*trigonocephalus* Martin, 1887" Molar fragment; 1575I *Cervus* sp. indet. Antler fragment; 1575J *Cervus* sp. indet. Phalange fragment; 1575K *Sus* sp. indet. Canine fragment.]

**S II-B7b**

[M…050-090/091, left. Second-quarter report, 1894. Dubois 1894d.]

Results obtained during the previous quarter on the geologic formation that contains remains of vertebrate animals in Central Java, made it incumbent upon us to check the facts on which they are based. For this purpose, at several important locations in the Kendeng, distances were measured and elevation differences were surveyed in and where possible strike and dip of the beds were determined with a leveling instrument, more accurately than had been possible with an inclinometer.

The most important of these more precisely measured values are the following: >>1200 meter northeast of Kedeng Brubus, just north of the point where the Kali Brubus crosses the boundary between Rembang and Madiun, is the point where the bone bearing tuffs begin resting concordantly on a massive layer of coarse andesitic breccias, locally called Kedeng Panas. Here, the dip of the beds is 5¼º SW up to Kedengbrubus (113m. above sea level). They consist almost entirely of fine sandy andesite tuff, about 100 meters thick. Intercalations occur of only two very thin claystone beds and a conglomeratic bed of 1½ meter but near the settlement [of Kedungbrubus] the tuffs are partially replaced by a claystone interval of 36 meters [in thickness]. These are again followed by beds of sandstone like andesite tuff, also generally consisting of very fine material up to the alluvium of the lowlands (97 meter above sea level) where the formation generally disappears below the surface. A few thin claystone beds and just as thin a conglomerate bed occur south of Kedungbrubus. The dip decreases slowly in this direction and at Notopuro, 5km southwest from the point where the bone beds begin above Kedungbrubus, it is only 1½º. With these facts in hand, the thickness of the formation can be calculated up to near Notopuro to at least 250 meter. >>About 6km east of Kedengbrubus and 1km north of Ando, also along the boundary of the named residencies, the bone beds begin at a few hundred meters higher elevation with a dip of 13º to 15º. However, this dip soon decreases and as nearby as Kedung Galeh (located just about along the strike of the foot of the formation above Kedengbrubus) it is no more than 6º. Here the dip angle gradually diminishes down the inclination, so that near Kali Gedeh it measures only 2º. Even further south, the dip becomes steadily smaller and between Saradan and Wilangan, the tuff beds are no longer visibly tilted and have virtually retained their original horizontal attitude [Notopuro Formation, Duyfjes 1936]. This southern portion of the formation, that is also 5km wide generally contains increasingly smaller and larger pebbles, but otherwise has the same nature as that of the northern half and also contains bones of the same animal species as the northern deeper beds, albeit not as numerous. For this southern half of the formation that reaches as high as 125 meter above sea level near Saradan, a thickness can be calculated of 100 meter. In this region, the entire thickness of the formation must therefore be more than 350 meters. The entire formation contains bones in its deepest parts as well as the most elevated beds. Throughout its thickness it has a fluvial character, indicated by the widespread presence of freshwater mollusks and current bedding observed in all tuff layers.

The latter suggests, also in other regions of the Kendeng, a general current direction from west to east. Only near Ngawi could a broadly northwesterly direction be determined from the current bedding. It appears therefore that in the past a branch similar to the Madiun river already controlled the water shed through the valley between the Wilis and Lawu. This very fine material that generally makes up the tuffs and even more so the claystone prove that these must have settled downstream in a quietly flowing river, that is in agreement with other facts. Claystone rarely contains bones, but the richest layers are generally those of sandstone-like andesite tuffs, resting in sudden transition immediately on top of the claystone and starting with lapilli. On the other hand, transition from sandstone tuffs into claystone is always very gradual. This supports the theory that the animals perished during volcanic eruptions. Claystone reflects periods of slow and quiet sedimentation that was suddenly interrupted and that then gradually returned.

Very much the same layers of sand and clay make up the alluvium in the Ganges valley, but these are much younger in age and are not disturbed by uplift. Drill holes near Calcutta to 150 meter below the surface, or 140 meter below sea level and near Lucknow to more than 400 meter below the surface or 300 meter below sea level, did not reach the base of the river deposits.

The river deposits in Java with preserved animal bones from a late Pliocene or early Pleistocene fauna are present to at least 150 meter below sea level and since they were deposited in the downstream portion of a river, it must then be assumed that, similar to the situation in the Ganges, considerable subsidence of the land has taken place with respect to sea level during the formation of the deposits. When the lowest bone bearing tuffs were deposited, the land must have been about 150 meter higher with respect to sea level than at present. Thus, from the geology of this formation as well as the nature of its fauna whose remains it preserved, confirmation is given for the hypothesis by Wallace about the mainland character of Java.

**S II-B7c, d**

[M…050-091/092,right/093. Summary report, Third-quarter 1894 (Dubois 1894d).

Summary Report on Paleontologic Investigations on Java
during the Third Quarter, 1894.

An attempt was made to gather further geologic information in the western regions of the Kendeng in regard to the bone bearing formation. This was primarily with the intent to check its distribution and thickness, similar to what had been done in the region near the Pandan.

Use was mainly made of the natural incision of the Bengawan once again, starting at the point where it crosses the western boundary of the Madiun residency up to Ngluwak in the Padangan District of the Bodjonegoro Department in the Rembang Residency, and further from the western boundary of Surabaya to her delta. Even if gathered results for the distribution and thickness of the formation might not even closely approach the accuracy of that in the Pandan area, better control was nevertheless obtained for the relationship with the underlying marine beds. Over this long distance, the beds are not everywhere disturbed in the same manner, nor with the same intensity [of structural tilting] and often the original horizontal attitude has been retained.

Near the western boundary of Madiun they are inclined to the SSW with dip angles of 6º to 8º, but they are soon replaced by inclined marine marls, limestones, sandstones and breccias, dipping 20º to 30º to the SSW up to Gendingan. Below Gendingan, dip is only 10º to 20º to the S or SE. Here and there they are concordantly overlain by not very thick horizontal bone carrying conglomerates and sandstone-like andesite tuffs. However near Golan the latter show dip of 10º S, while the underlying marine beds dip only 15º in the same direction.

From Trinil to below Ngawi, the marine substratum is everywhere completely hidden below the bone-bearing beds which, only at Trinil [are] horizontal, [and] over a length of 11km are dipping 5º to the SE, S and SW. Since this inclination can be traced very nicely in its proper direction [perpendicular to the strike] for at least 3km, the thickness of the formation may here be estimated at a few hundred meters or more. Here, the width of this strip over which it is distributed from W to E must surely amount to 5km.

Just north of Ngawi marine rocks are exposed again, while only at Ngluwak overlying bone tuffs of measurable thickness are present with virtually horizontal attitude. These are replaced further north by the alluvium of the Bengawan, and reappear in the hills of Tinggang as the SW border of this alluvium. Evidently, the bone bearing formation is also along the Bengawan of fluvial origin.

Current bedding [sets of cross-laminated beds], observed in many places indicates a general current direction from west to east, corresponding to the derived current direction closer to the Pandan. Severely upturned beds occur below the alluvium in the lower course of the Bengawan, at least from a point where it enters the Residency Surabaya to her most northern bend past Wringinanom. They are similar to those of the Kendeng along the northern boundary of the Madiun and Kediri Residencies and composed of andesite tuffs but they are softer (probably because of their lower altitude and less limy ground water). It is also quite probable that they are equivalent to the bone beds, although neither bones nor other fossils were found in them. They have an inclination of 15º NE and extend in that direction over 15 km, so that [we conclude that] these layers of sand and clay-like material reach a considerable thickness of 4,000 meter.

According to comments by the Chief of Geologic Mapping on Java, bones in verifiable Tertiary beds were found near Bodjongbitung in the south of Department Tjiandur of the Preanger Regencies. Since this find spot can be of great importance for the age determination of remains found at other find spots, further investigations were initiated specifically looking for those bones. From these we found that they occur as rare fragments, accompanied by marine shells, corals and other lower marine animals and all from further to be determined turtles, recognized as marine turtles. It thus turned out that comparison with other find spots of vertebrate bones was not possible. Tulung Agung the 19th of November 1894 Eug Dubois

[A copy of this report, basically repeating -B7c above, was rewritten by Dubois (1894d) in elegant formal handwriting, and thereby was far more carefully prepared than were normal draft in the "Report on Paleontological …".]

**S II-B8 (1896 letter about Dubois cross section of Trinil)**

Haag, 28.2.96

Dear Fürbringer,

This evening I received the galley proofs together with your letter and post card and I'm yet again very grateful for all the effort you have put into my article. It makes me feel rather silly looking at it to see from the corrections you have made how many errors I have overlooked.

I completely agree with the new arrangement of the figures. As you propose, the teeth, the second real tooth in two views, are best reproduced at ⅔ of my original figures (which is also ⅔ of their natural size). The other photographs, I think, will be best reproduced at the size in which I sent them, so the skull at ½ and the Femur ¼ of their natural size.

You will also watch over the figures being well and nicely spaced within the text.

With regard to Fig. 1 [below] I wish the lines H and I to be set horizontal, as you would expect a water level to be {* so parallel with the lines of text.} So slightly turn the whole figure like the hand of a clockwork; the stratigraphic layers will subsequently be slanting a bit more than they already are, as they should be. Nothing needs to be changed about the figure itself, only its direction.

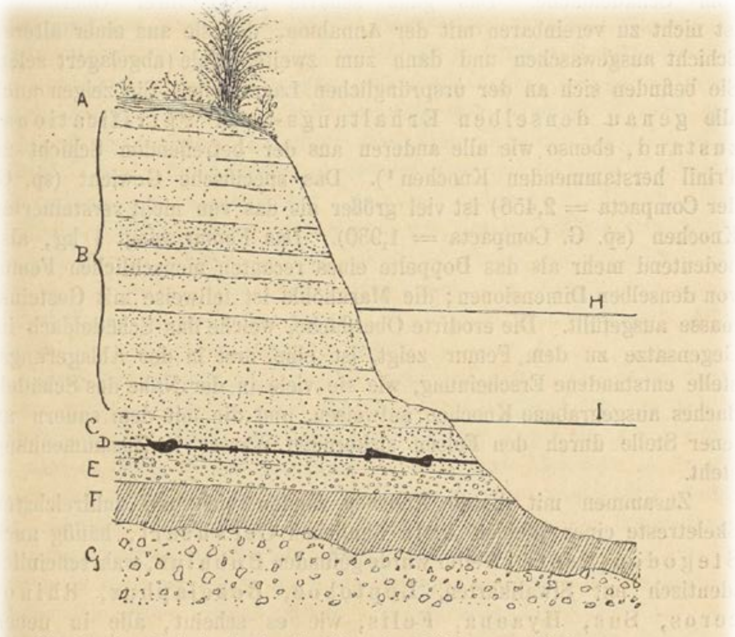


Fig. 1. Durchschnitt der knochenführenden Schichten zu Trinil.
*A* Culturboden. *B* weicher Sandstein. *C* Lapilli-Schicht. *D* Niveau, in welchem
ie Skeletreste gefunden sind. *E* Conglomerat. *F* Thonstein. *G* Marine Breccie. *H*

[Fig 1 as published in Dubois 1896f… see S II-F4; also, -E4]

I have made some corrections to figure 10: the words "Pliocaen" on the line underneath need to move 1½ mm up, the word "Miocän" needs to go 4 mm down.

I've also made a few small text changes, that I thought important; the one about the "Ueberzeugung", because the level of solidity is several times as [??gersi??g?], that interpretation could truly be ambiguous — the other because arguments were already put to me that particularly in Cebus a larger inclination was to be proven. Accurate measurements however have shown me that it is smaller than in Ateles.

I have erased a (*). Hopefully it is possible to move both the transverse line as well as the * in both of the Calvaria-Figurs 1 mm backward. Leaving my little article with you until its publication (unless you urgently see a necessity to do otherwise) I gratefully thank you yet again in advance for your tremendous effort and send warm greetings to you and yours.

Your

Eug Dubois

PS I am finally sending you a cast of *P.e.* Calvaria, which will have reached you by now.

$^{Y}$ED

**S II-C. 1890 and 1893 notes on Dubois' fieldwork along the Solo River.**

**S II-C1**

[Notes on 1890 paleocurrent observations at numerous outcrops along the Solo River; image of the original, below, and translation, next page.]

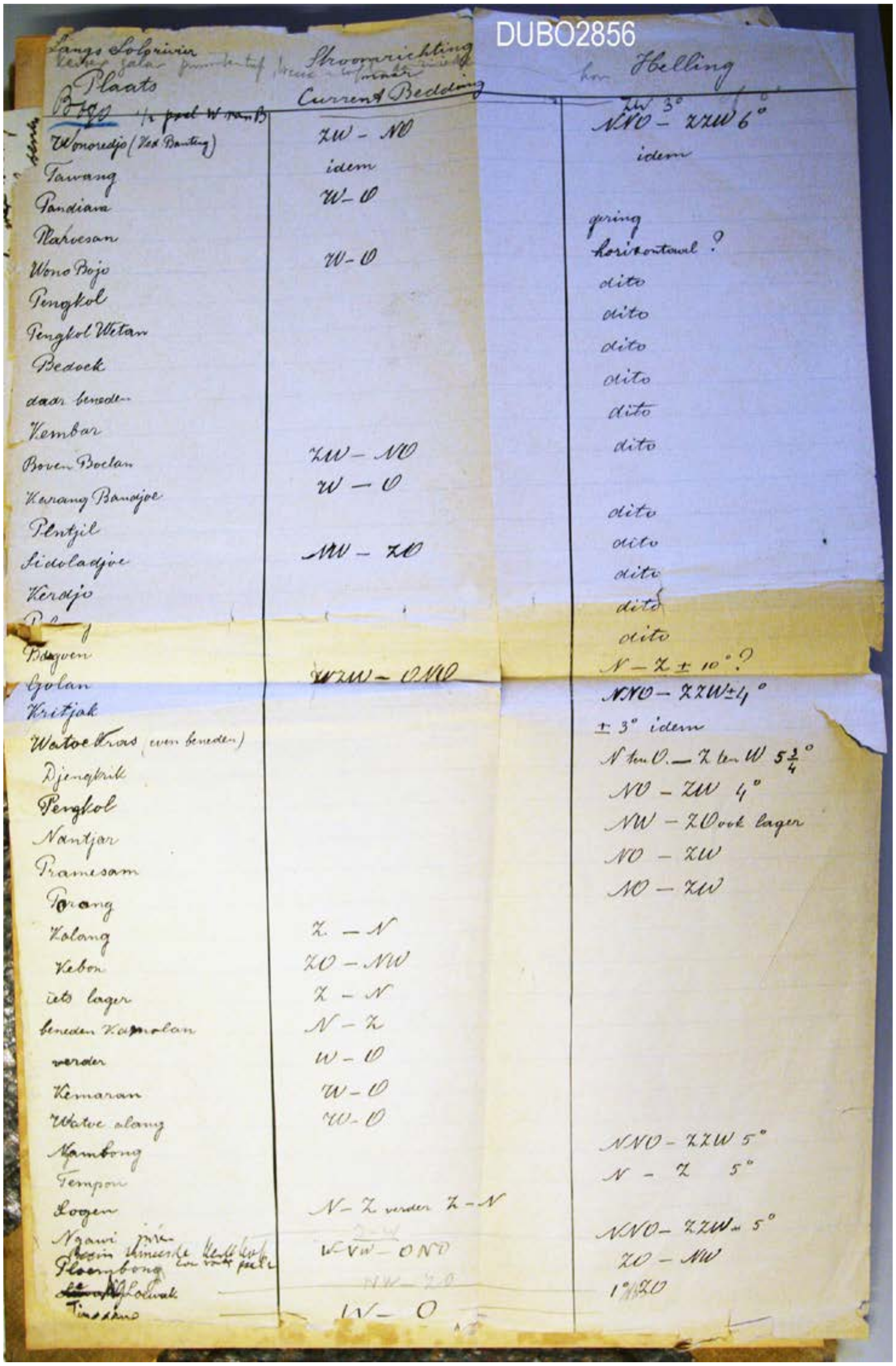


Langs Solorivier

| Plaats | Stroomrichting / Current Bedding | Helling |
|---|---|---|
| Bogo ½ paal W van B | | ZW 3° NNO – ZZW 6° |
| Wonoredjo (Ned Banting) | ZW – NO | idem |
| Tawang | idem | |
| Pandiara | W – O | |
| Nahoesan | | gering |
| Wono Bojo | W – O | horizontaal ? |
| Pengkol | | dito |
| Pengkol Wetan | | dito |
| Bedoek | | dito |
| daar beneden | | dito |
| Kembar | | dito |
| Boven Boelan | ZW – NO | dito |
| Karang Banadjoe | W – O | |
| Plentjil | | dito |
| Sidoladjoe | NW – ZO | dito |
| Kerdjo | | dito |
| [illegible] | | dito |
| Bagoen | | dito |
| Golan | WZW – ONO | N – Z ± 10° ? |
| Kritjak | | NNO – ZZW ± 4° |
| Watoe Kras (even beneden) | | ± 3° idem |
| Djengkrik | | N ten O. – Z ten W 5¾° |
| Pengkol | | NO – ZW 4° |
| Nantjar | | NW – ZO ook lager |
| Pramesam | | NO – ZW |
| Porong | | NO – ZW |
| Kolang | Z – N | |
| Kebon | ZO – NW | |
| iets lager | Z – N | |
| beneden Kamolan | N – Z | |
| verder | W – O | |
| Kemaran | W – O | |
| Watoe alang | W – O | |
| Ngambong | | NNO – ZZW 5° |
| Tempon | | N – Z 5° |
| Sogen | N – Z verder Z – N | |
| Ngawi [illegible] | WZW – ONO | NNO – ZZW 5° |
| Ploembong | | ZO – NW |
| [illegible] Kolwak | NW – ZO | 1° NZO |
| Tinadhuo | W – O | |

[Some of the handwriting in the list is Dubois' (red and blue – pencil, below), but the majority of the notes are the handwriting of someone else's (black). The column reading "Flow direction but Current Bedding" refers to paleocurrent directions determined from the inclination of cross-laminated beds. See –E1a, below, for map of the section of the Solo River along which most of these localities occur.]

Along Solo river

*(Kedung zalar(* [?] *pumice tuff, breccia along the river – horizontal)*

| Place: | Flow direction but Current Bedding | Dip |
|---|---|---|
| (below) Bogo | | |
| *½ paal west of B.* | | SW 3º or 0º |
| Wonoredjo (Ved Banteng) | SW – NE | NNE – SSW 6º |
| Tawang | same | same |
| Pandiana | W - E | |
| Plahusan | | little |
| Wono Bojo | W - E | horizontal ? |
| Pengkol | | same |
| Pengkol Wetan | | same |
| Beduk | | same |
| below (Beduk) | | same |
| Kembar | | same |
| above (upstream) Bulan | SW – NE | same |
| Karang Bandju | W - E | |
| Pentjil | | same |
| Sidoladju | NW – SE | same |
| Kerdjo | | same |
| Pelang [? *Hard to decipher*] | | same |
| Bangun | | same |
| Golan | WSW – ENE | N – S ± 10 º ? |
| Kritjak | | NNE – SSW ± 4 º |
| Watukras (just below) | | same ± 3 º |
| Djengkrik | | N to E - S to W 5 ¾ º |
| Pengkol | | NE – SW 4 º |
| Nantjar [*near Trinil*] | | NW – SE also lower |
| Pramesam | | NE - SW |
| Porang | | NE - SW |
| Kalang | S - N | |
| Kebon | SE - NW | |
| somewhat below [*Kebon*] | S - N | |
| below Kamolan | N - S | |
| further [*downstream*] | W - E | |
| Kemaran | W - E | |
| Watu alang | W - E | |
| Ngambong | | NNE - SSW 5 º |
| Tempon | | N - S 5 º |
| Logen | N – S (beyond S – N) | |
| Ngawi | E - W | NNE - SSW 5 º |
| beginning of Chinese cemetery just before paal 2 | WNW [*and*] ENE | SE - NW |
| Plumbong | | SE - NW |
| Gluwak | NW - SE | 1 º SE (NNE) |

**S II-C2**

[M…046-141; also, M…049-240. 1890 notes about the geology at Trinil.]

1890 At Trinil, on the left shore of the [Solo] river, [a] findspot of lapilli tuff; on the right shore, a very large 100-meter long ledge [*‘plaat’* in the original] of sandstone-like andesite tuff, which towards the south rests on andesite lava containing smaller and larger andesite blocks. Another 100 meter further southwards … and within the steep shoreline embankment, several meters above the water level [of the river], smaller bone-bearing sandstone intervals also occur. At the more northerly find spot along the left shore, the lava [lavaresten?] also crops out. Based on these facts, it becomes clear that the bonebed formation does not dip or have a concordant [base]. [Instead the bonebed formation] has horizontal bedding and rests in an irregular fashion on top of the lava.

**S II-C3**

[M….046-141. [Dubois’ notebook is entitled “*Notes related to the paleontological investigations of the West coast of Sumatra*” [Aantekeningen betreffende de paleontologische onderzoekingen der Sumatra's Westkust in Dutch] contains more information about Java than Sumatra. Page 243 includes an artful sketch of a modern migrating gravel bar of the Solo River between Trinil and Ngawi. An annotated scanned version of the lower portion of page shows Dubois observations and inferences.

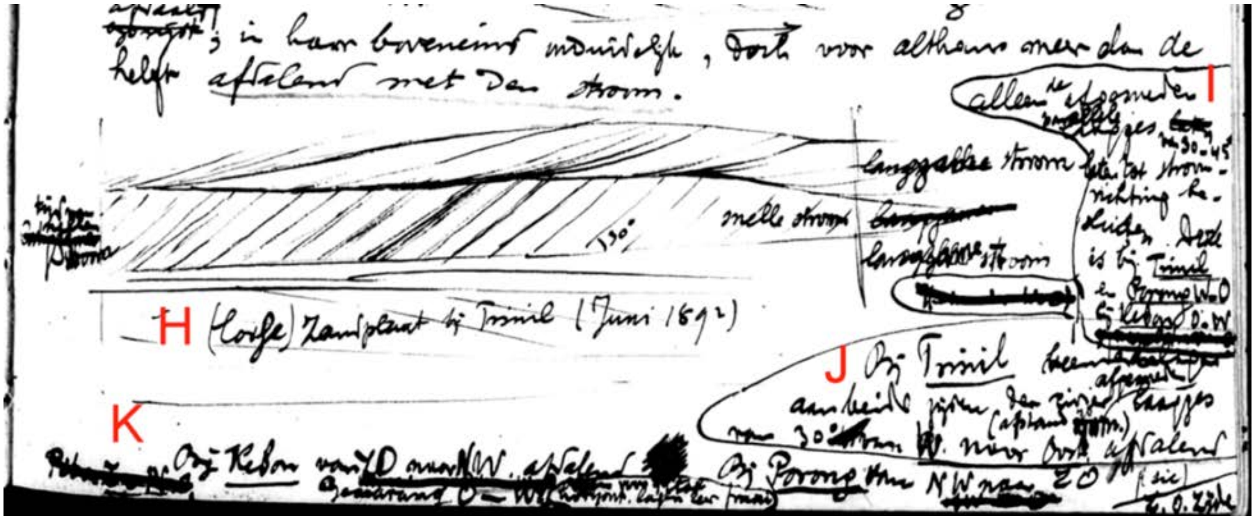


[the H portion, above, reads] (High) sandy shoal at Trinil (June 1892) [and show a cross-sectional view of bedding planes inclined 30 degrees, which Dubois interpreted to represent] “quick current,” [whereas the flat-lying laminations that indicate] “slow current.” [In a nearby note, I, Dubois asserts that only the thin parallel sets of laminations that have truncated tops and dip at 30-45 degrees allow for determination of current direction. He added in J that] at Trinil[,] bonebeds on both sides of the river (separated by a distance of 70m) have truncated thin beds dipping 30 degrees from west. [in part paraphrased]

[Bedding relations in modern point bars along the Solo River apparently inspired Dubois to observe cross beds in the bone-bearing formations that were exposed in the river embankments. In his monograph on *Pithecanthropus erectus*, Dubois concludes that the LB was deposited by fluvial currents that flowed in a direction opposite of that in those of the modern river (-B6j).]

**S II-C4**

[M…046-142 (part). Annotated image of Dubois' July 1892 notes on fluvial bedding along the Solo River, Page 244 in "*Notes related to the paleontological investigations….,*" includes an expressive sketch of the Solo River in plan view at Pramesan, east of Trinil.]

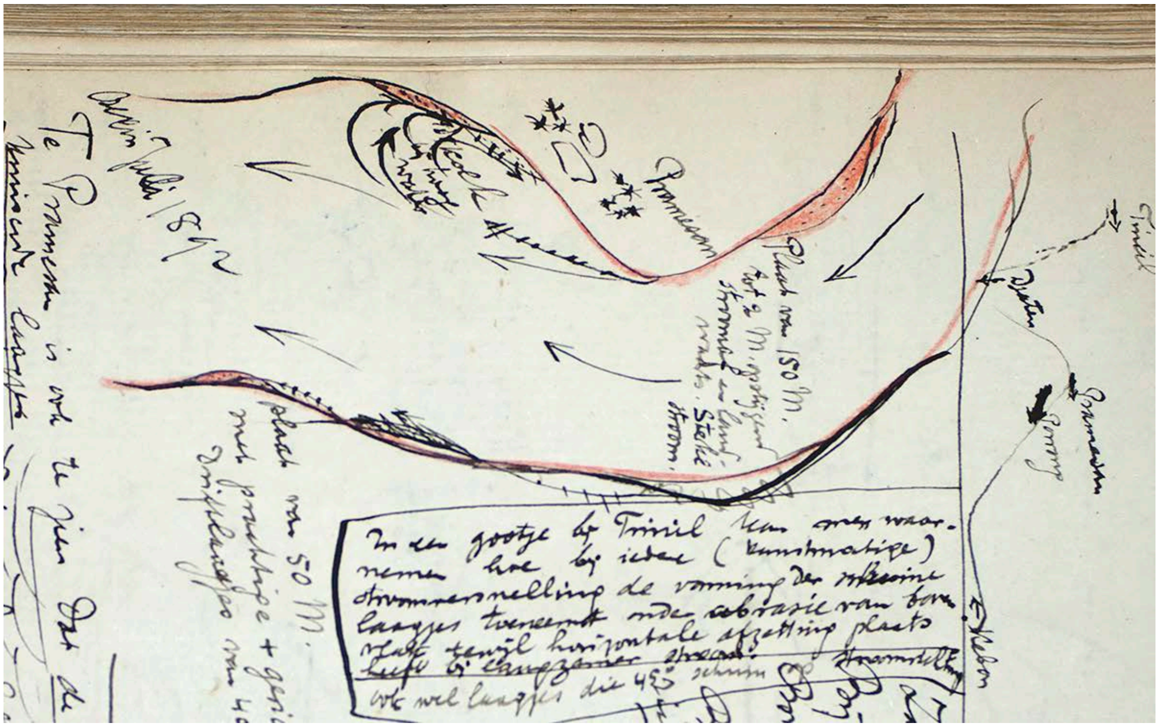


[Dubois' observations were made] "at the beginning of July 1892" [along the east side of the Solo at a] "bank 50m long with beautiful oriented small drift beds [cross lamination] of 40 degrees [of inclination]." [Elsewhere on the page, he lists paleo-current measurements at various places along the river] "Slanted beds within the sediments at Trinil [are] from W to E[ast in a] down dip [direction.]

[The prominent note that Dubois outlined on the page, relates] In a small channel at Trinil, one can observe how an increase in the formation of thin-slanted layers [cross laminations] takes place with each (artificial) flow acceleration accompanied by abrasion from the top surface, while horizontal deposition takes place at slower flow. [An additional note states] Also at times thin slanted layers that are 45 degr[es] cross-wise to the flow acceleration. [A summation on page 84, he concludes that] oblique lamination in the bone tuff primarily dips downwards from SE to NW between Trinil and Pitu [east of Trinil], here and there vortex currents (always in fine sand) [form cross bed sets] in the reverse direction.

[At the bottom of the preceding page, Dubois noted that the cross-bed orientation is from west to east in a down-dip direction, while near Porong the down-dip inclination is from northwest to southeast, Djaten from north to south, Kebon from east to west, Kalang from east to northwest, Watualang from east to northwest, and at Pitu is from east to northwest.]

**S II-C5**

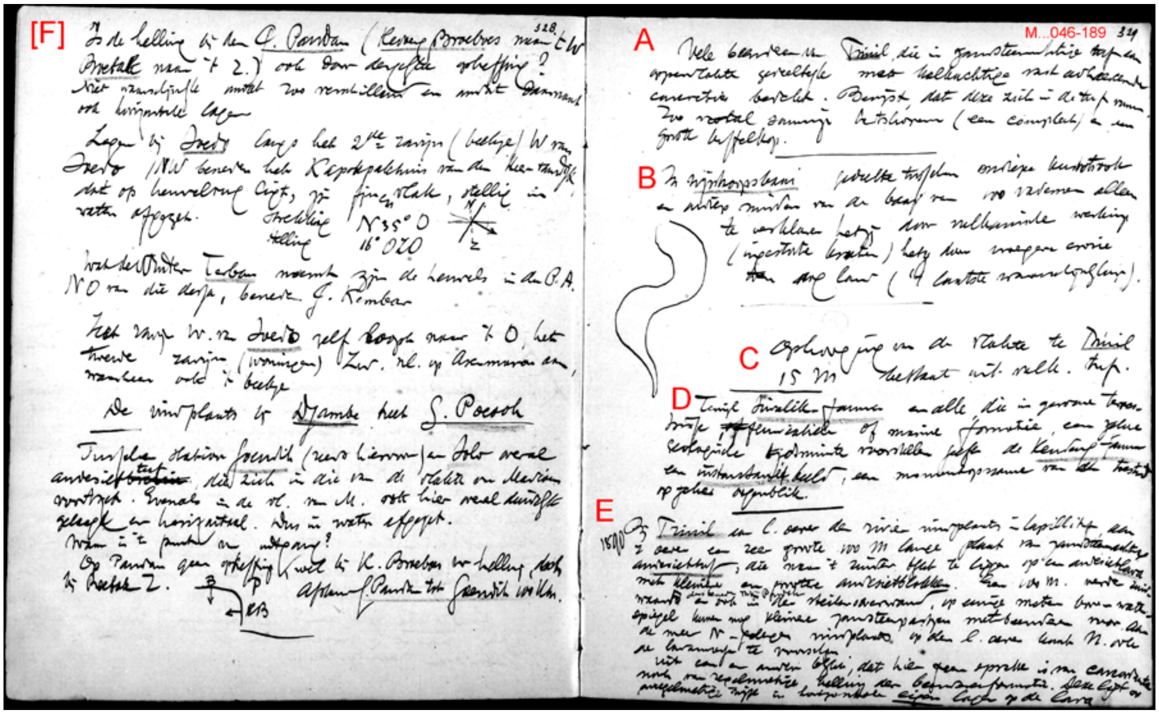


M…046-189, Page 329, on right, in "Notes related to the paleontological investigations…."

A. Many bones from Trinil, which in sandstone-like tuffs at the surface [of outcrops are] partially covered with limy [$CaCO_3$] concretions, that adhere tightly to its surface. This proves that they are formed wiithin the tuff. This is especially so with some [*Axis lydekkeri*] deer antlers (one is complete) and a large buffalo skull [*Bubalus palaekeratau*]. [see also, -C6, below]

B. In [the] "Wijnkoopsbaai" [a bay] [the] portion between [the] shallow coastal strip and shallow(?) center of the bay [at] 100 fatoms [can] only be explained by either volcanic action (collapsed crater) or by previous erosion [when] the land was dry (the latter most probable).

C. Raised part of the flats at Trinil [which are] 15 meters [above the river] consists of volcanic tuff [underlain by 'tuffs'].

D. While [the] Siwalik fauna and all those in normal terrestrial, fluvial or marine formation(s) reflect a certain geologic span of time, the Kendeng fauna yields an instantaneous image, a snapshot of the situation at a certain moment in [geological] time.

E. [Apparently copied from notes made at Trinil in 1890] *1890* At Trinil, on the left shore of the river [a] find spot of lapilli tuff [a fossil locality in lapilli-bearing volcaniclastic sandstone, LB, of the 1891 Skullcap Pit]; on the right shore a very large 100 meters long ledge [*plaat* in Dutch] of sandstone-like andesite tuff [the other LB outcrop], which towards the south turns out to rest on an andesite lava [of the later-defined Pucangan Formation] that contains smaller and larger andesite blocks [as clasts in a 'tuff breccia']. Another 100 meters further southwards (hence below Mig? Pandoh? [perhaps referring to 'my cabin,' built in 1892]) and within the steep-shore embankment, several meters above the water level [of the river] also smaller bone-bearing sandstone intervals occur [the situation is portrayed in the I.J.R.C.P. (1979) mapping shown in S I Figure 18]. At the more northerly find spot along the left shore [probably the locality known as Batu Gadja], the lava [deposit; *lavaresten* in Dutch] also crop out in the N[orth]. Based on these facts it becomes clear that this is absolutely neither a case of concordant nor regular dip of the bone bed formation. [Instead] it rests in irregular fashion [but] with horizontal internal beds on top of the lava [that is, the LB is horizontal but rests discordantly along an irregular surface that had formed on the breccias of the Pucangan Formation].

**S II-C6**

[M…012-463-466. Letters to Verbeek, diary- and notebook-entries regarding limy, **$CaCO_3$**, concretions]

(i) [Dubois' diary under 'Remarks' for May 31, 1890 (M…015-350)] Limestone concretions [occur] near Trinil at various levels (even up to 10 meters below the surface [of the top of the terraced embankment] between thin beds of 1-, 2- to 3-cm thickness. Appears as if limy solutions penetrate the formation and lime has precipitated. Same [phenomena occurs] at Bogo (lesser [?]) along the [Solo] river bank near the small bridge, just as [it does] near Patiayam (G[unung] Strompret). Also [lime precipitate is found] in diagonal cracks in the rock formation, confirming the explanation give [here] above.

(ii) [Excerpt from letter to R.M. Verbeek, December 23, 1890 (M…012-463)] At many locations in the [Kendeng] hills, they [the strata] contain truly weathered small andesite rock fragments, and these irregularly formed white rock fragments also occur in weathered breccias (of which the largest part, admittedly, in fact consist of [secondary] lime concretions).

(iii) (M…046-189, page 328-329 of "Notes related to be the paleontological investigations …" [undated]) Many bones from Trinil that are [found] in sandstone-like tuff at the surface [are] partially covered with a limy concretion that adheres tightly to them. This is proof that it is formed within the tuff. Especially noticeable on some deer skulls (one complete) and a large buffalo skull.

**S II-C7**

[Notes on stratigraphy of left-bank excavation (combined 25-m and 40-m Trenches) in the October 1893] [M…046-175] [page] 304 [of *"Notes related to the paleontological investigations…;"* the scan of which is shown on the next page] [an unrelated note in the upper left margin has been omitted from the translation] Notations on the beds at Trinil

21 Oct. 1893

Large trench (of 40 X 6 meters), at upper end (NE) [portion of the trench, nearest to the Skullcap discovery point, the excavation reached the] top of black claystone [which is] about 1 meter deeper than the Chimp skull [Skullcap]; [the] lapilli bed (in which he [the Skullcap] occurred) [is] about 2 meters thick. On top of that [is] sandstone. On top of that [is] sandstone.

Lower (SW) end [ of the trench, west of the Femur I discovery point was] excavated 1.30 meters deeper [than at the NE end, and] into the conglomerate after [below] alternating lapilli and sand. Bones [are] still [present] in the conglomerate. This is only ½ meter thick, then [follows below the] black clay layer, which is thus 1.80 meters deeper than at upper end. Deepest spots are about 3 meter below the current river level (East Monsoon level).

If 5° 48' [dip] Sin = 0.101 corresponds to a length of 1.80/0.101 = 17.9 meters or about 18 meters However, that has probably also been the true difference in the dip direction

[inserted along the left margin] Trinil: Kriele (16 Nov 1895) in [a] pit of 25 meter more than 2 meter below lowest river level

[Page 304 of the "Notes …" (M…046-175; scan on the next page) are nearly verbatum from "Remarks" of October 31, 1893, diary entry below (M…016- 079). The "Remarks" reads "…..zwarte kleisteen ongeveer 1 m dieper dan chimp- schedel. Lapilli laag (waarin kop) ongeveer 2 m dik," while the diary is "….zwarte kleisteen ongeveer 1 m dieper dan chimp. schedel. Lapilli laag (waarin hij lag) ongeveer 2 m dik."

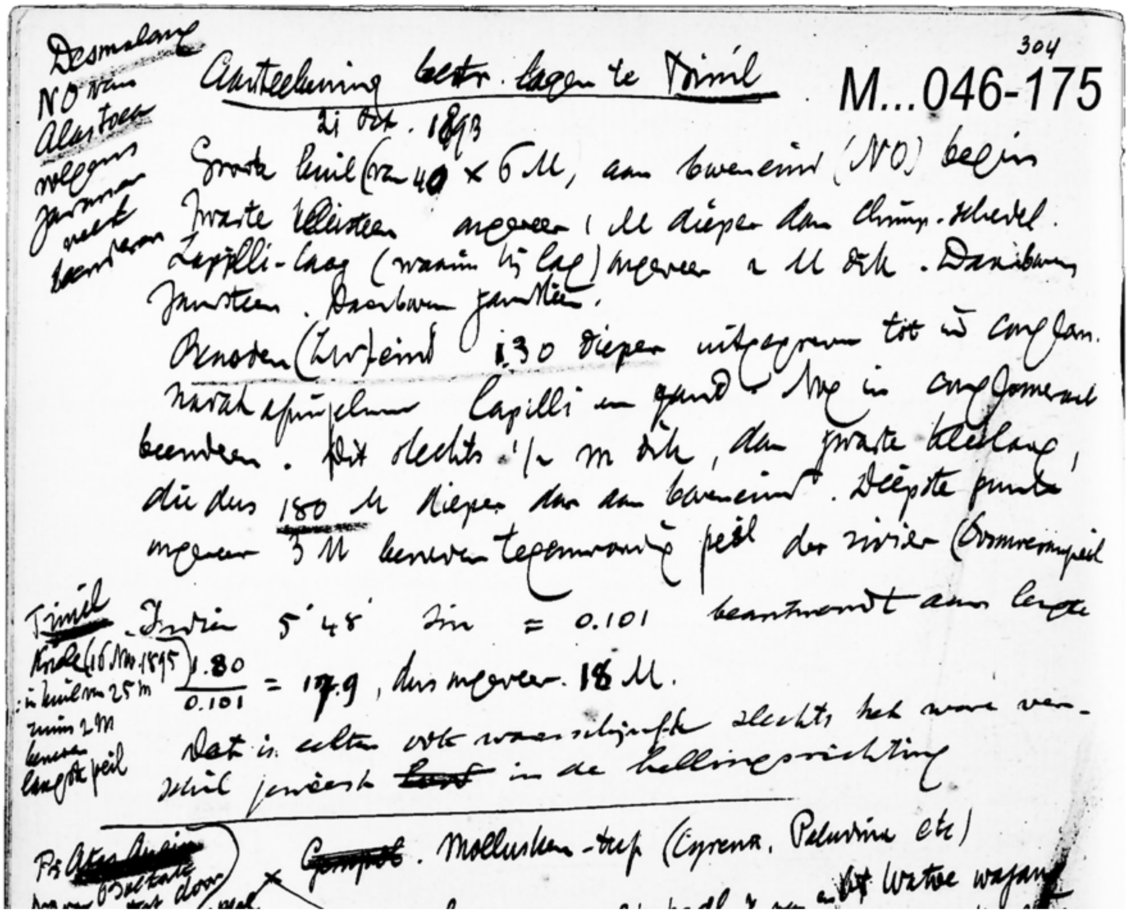
304

M...046-175

Aanteekening betr. lagen te Trinil

21 Oct. 1893

[Dubois referred to two observation points in the excavation open on October 21, 1893: the “upper end (NE),” and “Lower (SW) end,” both which were inside of the “Large trench (of 40 x 6 meters).” According to his 1895 sketch of the trench (Figure 2b, main text) and Kriele’s 1899 map of it (Figure 3a, main text) the long dimension of the excavation was approximately east-west, and it was much narrower from north-to-south. Dubois reference to two observation points appear to refer to (i) a point in the northeastern corner of the trench, adjacent to the Ledge and near the Skullcap Pit, and (ii) a spot along the 1893 backwall of the 40-m Trench that was close to the Femur I discovery point, 18 m away.

The stratigraphic circumstances he describes are as follows (see diagram on the next page):

(i) At the eastern end of the 40-m Trench, near the Skullcap Pit, the discovery level of the *Homo erectus* specimen (that is, the PFZ) was ~1m above the top of the claystone, and the LB was ~2m thick, so that the top of the LB was ~1m above the PFZ; the LB was overlain by sandstone, which he did not further characterize. (see S II–B2g for note about reaching 1.5 m below LWL in October 1891).

(ii) Towards the west in the 40-m Trench, the excavation went 1.3m deeper than it did near the Skullcap pit, and to a total depth of ~3m below the LWL; the claystone at the western end underlay 0.5m of conglomerate, which occurred below small-pebble conglomerate interlayered with sandstone; the conglomerate-claystone contact was 1.8m lower on the west compared to the east, and had a dip of nearly 6$^{o}$ or more. Ambiguity in the Dubois description make it possible to infer that the top of the LB occurred at a substantially higher elevation at one end of the trench that the other. However, if the 6$^{o}$ dip that he described at the top of claystone is applied to the top of the LB, the upper portion of the LB would have risen 5m over the 50 m length of the 1891-1893 excavations, which did not occur.]

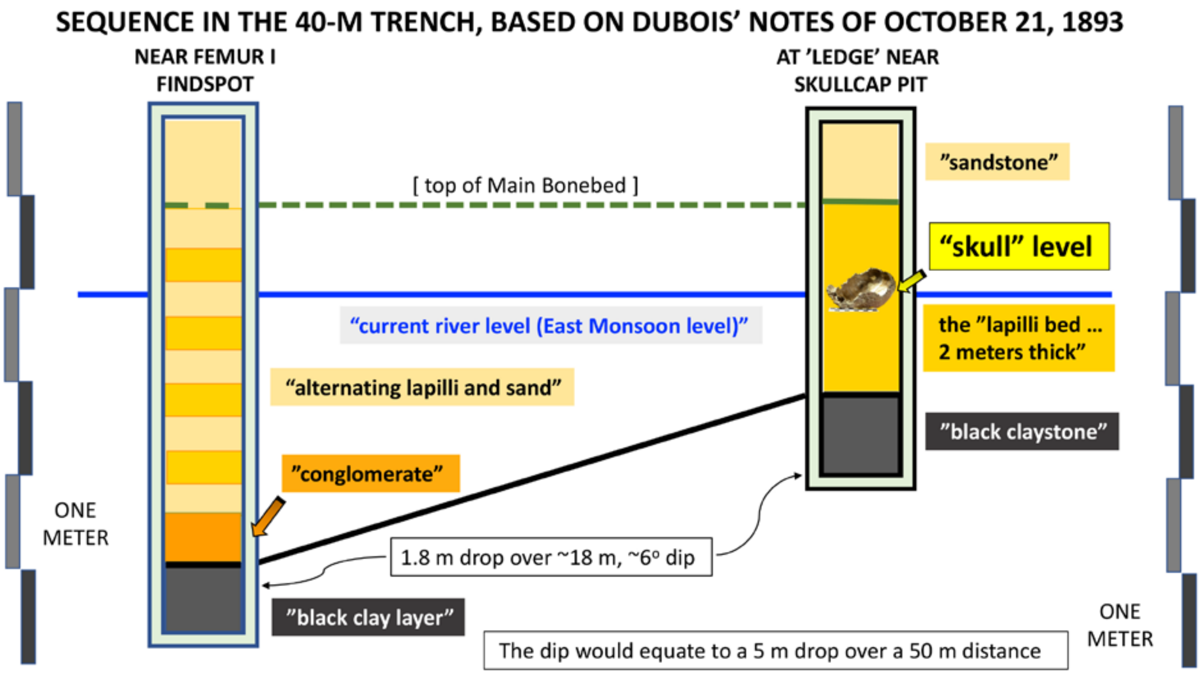


[The claystone was ~1m deeper than the level at which the Skullcap was found (our PFZ). However, the top of the claystone dipped 1.8m westward across 18m within the 40-m Trench, and the deepest spots of excavation reached 3 meters below dry-season river level (LWL). The dip at the top of the black claystone appears to indicate a local thickening of the overlying sandstone, pebble conglomerate (with the "lapilli") and coarser-grained conglomerate.

In his 1895–1896 published accounts, Dubois separated the lower portion of the lapilli bed into a "conglomerate" bed (e.g., Figure 2a, main text, and –F1, below).]

### <u>S II-C (Examples of geological exchanges between E. Dubois (ED) and R.D.M. Verbeek (RV)</u>

[In 1890, Rogier D.M. Verbeek (1845-1926) was the head of the Mining Bureau in the Netherland Indies. He was a renowned geologist of the Dutch East Indies and famed globally for reporting on the 1883 Krakatau volcanic eruption; he was the principal author of "Geology of Java and Madura" (Verbeek and Fennema 1896). Dubois' consultations and interchanges with Verbeek about the geology of Java are partially preserved in letters between the men.].

### <u>S II-D1</u>

[M…012-437. ED to RV; 12 November, 1890.]

[Following a 3-day boat trip down the Solo River past Trinil (S II-E1a, below), Dubois had determined that the core of the Kendeng Hills consisted of folded claystone, sandstone, marl and limestone, and] "andesitic breccias, conglomerates, and andesitic sandstones or tuffs," [striking N75° E and dipping 15° southward on average, made up of younger volcaniclastic formations on the flanks of the range;] "after a bit of searching, I could find fossil bones everywhere in these 'Pliocene' rocks" [partially paraphrased].

**S II-D2**

[M…012-442/443. ED to RV; 30 November, 1890.] [S I Figure 21]

[Over the course of November,] I have paid attention to the elevations at Kedungbrubus …. [With respect to] the distribution of bones … sandstone-like andesite tuff was most prolific …. The tuffs, as well as the breccia, only have traces of bedding …. [while] sand forms thin cross beds that separate sandstone lenses … [the beds are] several cm thick …. [and having been] derived from the rocks due to weathering … [were] deposited under the influence of currents". [With reference to the sketch cross section included in the letter, below], Dubois commented that the south-dipping sequence extends downward stratigraphically from the] "bone tuff" [in the flat-lands around Kedungbrubus up the mountain slope to the] "andesite breccia" [of Gunung Butak at 450 m elevation, where Raden Saléh collected vertebrate fossils.] "No remains of sea animals," [occur, and rather there are] "many remains of fresh water mollusks.

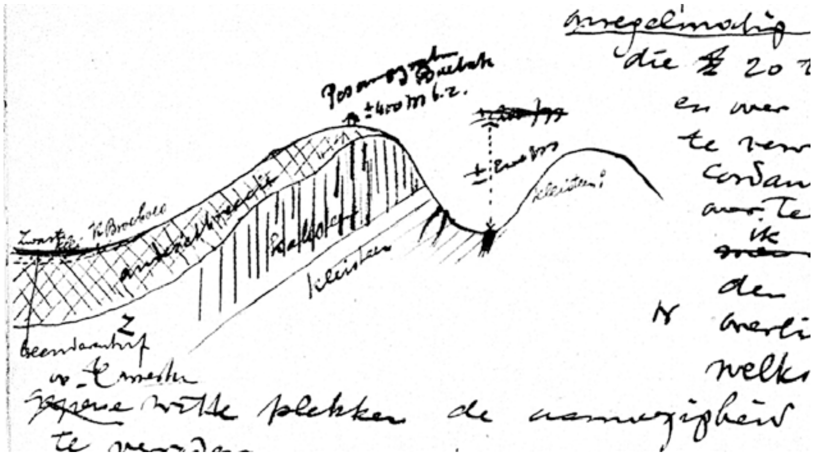

[Other notes: Dubois observed dark clay in lowland (on left of the cross section, near K. Brubus). He characterized the principal formation with vertebrate fossils as "andesite breccia." It held up the hills north of K. Brubus, and underlay a high point of +/-450 m at Gunung Butak. The "breccia" unit was later divided into the Pucangan, Kabuh and Notopuro Formations (Duyfjes 1936 in S I Figure 21).]

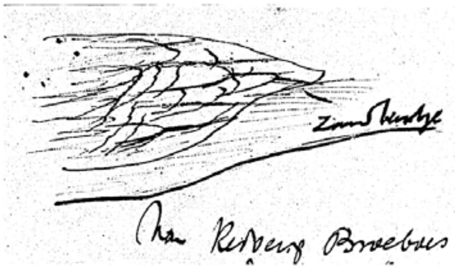

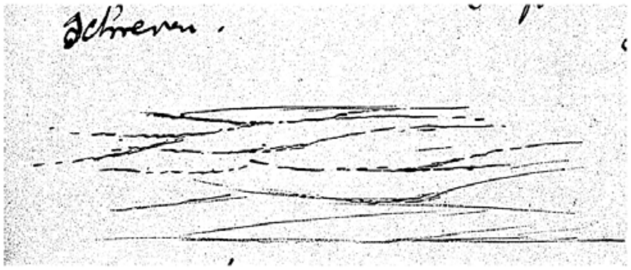

[Small sketches in this letter reveal Dubois' interest in and close observation of the sedimentological origin of water-laid cross laminations, in relation to the flow direction of fluvial currents. The sketches also reveal Dubois' ability to illustrate small-scale geological features he saw in the field, just as his geological cross section, above, shows his capability to characterize large-scale relationships. ]

[Dubois Collection at Naturalis has many other unpublished documents of Dubois' concerning Kedungbrubus. See also –B7, above, S I Figure 21, and Albers and de Vos, 2010.]

**S II-D3**

[M…012-443/444. ED to RV; 30 November, 1890.]

[Regarding the origin of the fossil-bearing volcaniclastic formations,] …. considerable eruptions of andesite breccia [accumulated around a lake, where] large Kendeng vertebrates, herds of buffalo and bison, deer and elephants passed [and …] in which hippopotami, crocodiles and river turtles lived … many of these animals perished, [and] heavy rains … washed the tuff downward as a slurry … and swelled the rivers, where animal remains became buried in the gravels and other materials.

**S II-D4**

[M…012-445. ED to RV; 30 November, 1890.]

[On December 1, 1890] before sending this to you, I wanted first to climb the [Gunung] Pandan again [a volcanic peak east of Kedung Brubus; S Figure 21]. Although the eruption products of andesite breccia and tuff [there] may have originated from large volcanoes of central Java, including Wilis and Lawu, [and] the resulting flows spread into the flats …. Large water animals could have lived near Kedungbrubus and in the Ngawi area …. In marshy, abandoned meanders of the large rivers …. That carried water from the alpine lands of Java [on the south] to the Indian Ocean … and [also] partly across the bottom of the Java Sea (then to the mainland).

**S II-D5**

[M…012-446, M…012-450-459, M…012-463/466; also, 008-337/338. ED to RV, 30 November and 6 December, 1890.]

I am very delighted about all the facts that are so rapidly becoming clear, now that my eyes have been opened following that wonderful trip along the Solo River [-C1, above].

[Concerning Verbeek' corrections of Dubois' account of the geology around Mount Butak (-D2, above), and discussion about confirmatory information on other topics, ED opened with] my cross section of Butak is correct …. I did see exactly what I sketched; [and then, regarding] big volcanoes. It may seem presumptuous that the novice [geologist] Dubois dares to differ in this with the very experienced geologist Verbeek [but] I cannot agree with your identification of the bone-bearing tuff beds, etc., as lapilli [among other differences with Verbeek].

### S II-C. Other E. Dubois' documents relating to Trinil

**S II-E1a**

[M…016-101 colorized and annotated with some place name spellings modernized. The segment of the river shown was studied and collected intensely by Dubois and KdW in the 1890s. The east-west road with the kilometer-markers (e.g., 28, 27, 26 …) was the major access route to Trinil from Ngawi, the nearest city and military post. See –A1c for portion around Trinil.]

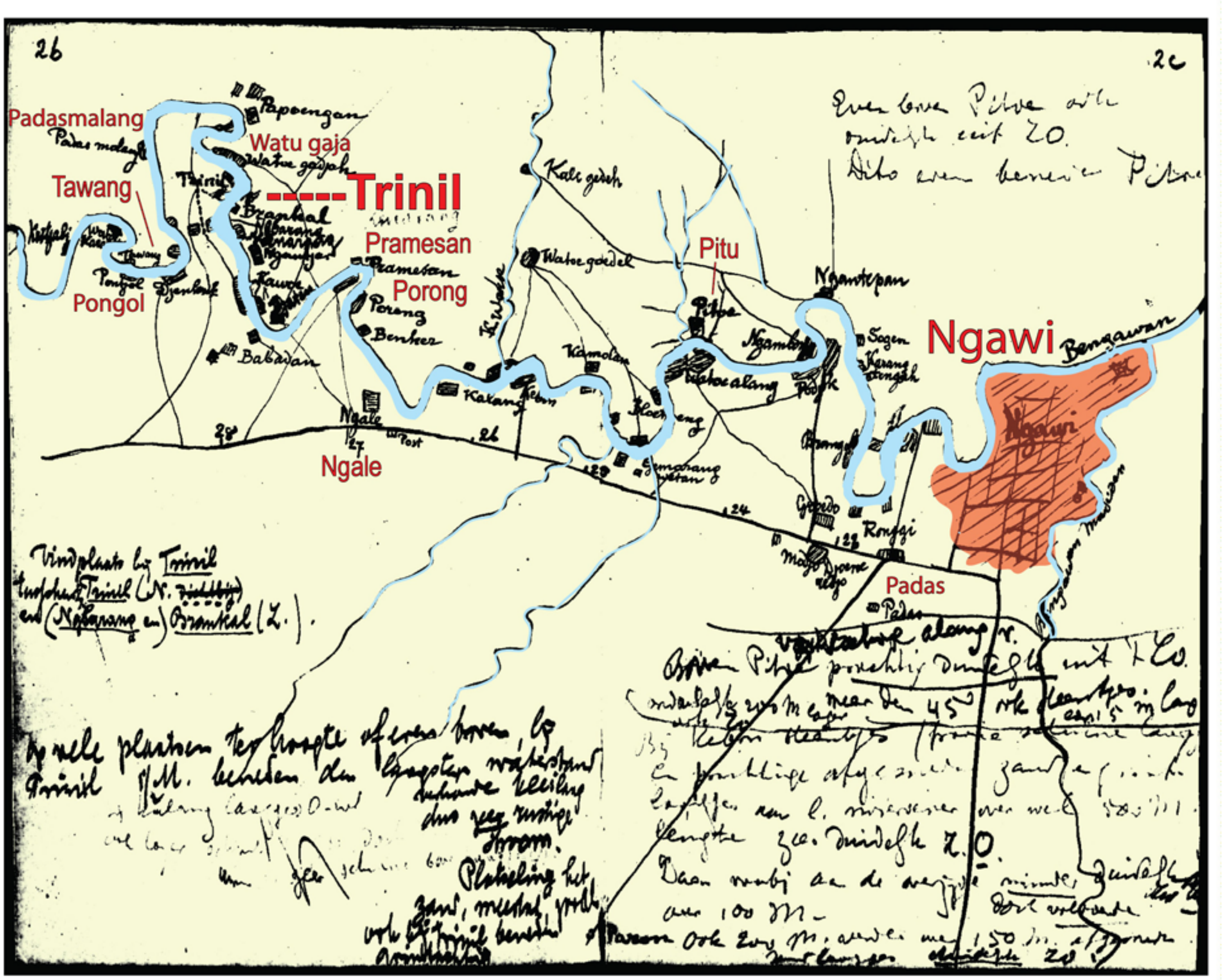

**S II-E1b**

[M…033-663/664. Letter to an unidentified friend in Java concerning the Femur I discovery 1892.]

Tulung Agung, 23 September 1892

Dear Friend,

Although I was happy to receive the Transactions of the Zoological Series containing the article by Owen now already more than a week ago, I have not yet thanked you for it. I should therefore offer you my apologies at this time.

Notwithstanding my [current] location in cooler weather, I have not been able to get rid of the nervous tension and general malaise of which I have been suffering more and more over the last two years. Unfortunately it was even worse than ever before during the last few days, perhaps just prompted by that find of the chimpanzee femur.

By forwarding me the relative article of Owen [in the Transactions of Zoology series], you have once again been of great service to me. This has allowed me to progress with the studies of the discovered specimen as far as is possible without comparison with real skeletons. The conclusion that I was thus able to draw is the following: the bone shows the greatest correspondence amongst the anthropoids with the chimpanzee, just like the skull and the molar.

However, for the femur this [correspondence] is even much greater with humans. In dimensions and proportions, in the curvatures of the joint distal end(s) [condyles] and their placement with respect to the diaphysis [spinal process of the tibia] and with respect each other, the bone is completely human. Measured lengths, thicknesses, radii and angles are so strikingly human that I was amazed myself when I compared them.

This key to the skeletal mechanics surely proves that the animal possessed human proportions of trunk and legs and that it walked upright. However, only a few insignificant peculiarities prove that it is not a human leg, but that of an anthropoid, which by the way is also indicated by the [fact that the] discovery spot [is] near the spot where the skull was found.

If the weather were to stay dry for awhile, there is a real chance that [other] parts will be excavated in the same river bank of the arm, [or] the hand and foot, thereby providing evidence that would confirm the above conclusion (which for me is already a certainty)

For myself, it is also a pleasant experience that my prediction, made 4 years ago in the N.T. [Natuurkundig Tijdschrift?] that in our archipelago forms would be found that are transitional between humans and their highest living relatives, has now been proven correct. Again my sincere thanks and in the hope that you are enjoying a better health situation than me, I remain with friendly regards,

Yours faithfully

Eug. Dubois

**S II-E1c**

[M…013-184/185. Draft of letter to Max Weber, December 19, 1892.]

[Max W.C. Weber (1852-1937) was the Professor in Zoology in Amsterdam, and a Dubois friend (M…013-171 to M…013-241). See Albers and de Vos, 2010.]

19-12-1892
Highly esteemed Professor

Although I received your letter of the 21st of October already the 28th of last month, the post package mentioned therein that was simultaneously sent only arrived yesterday. I had in particular been desiring and looking out so much for the chimpanzee skull that I postponed my answer – on your earlier received generous writing – up until now. I am very grateful to you, more than I can say, for that you keep having good intentions with me, and kept keeping trust in me. The help you have given me is even more appreciated because of what will come from it and I am so grateful for your benevolence, your trust and your help to the success that is my part of which I have occasionally dreamed, but never expected, and has now come to pass.

Failed to find more of Erectus because of high water [see –B5d and –B5e, above]

Only slowly have started to realize of what importance *A.e.* is and yesterday the comparison with the skull was an extremely exciting moment.

And now you would much oblige me to [?? unreadable ??] *Hylobatus* and Chimpanzee skulls sequentially listed

| Siamang | Chimpanzee | Erectus | |
|---|---|---|---|
| 10 | 13 | 18 | length skull |
| 7 | 9 | 13 | width |

Femur width walking upright and appearance of upper [ ??? ] human
But now most importantly
Calculation of skull capacity

I have written the above firstly to prove to you that your benevolence has led to my success, and secondly, because I hope to make this known via you, to be in some way able to whip out the impression that my leaving Amsterdam has made on you. I hope I will succeed to do that with all my teachers and particularly with the still, by me, highly regarded prof. Place! [Thomas Place, physiologist]

For myself I am of the opinion I have done my best in the Indies and not have been less loving or loyal to science than I would have been had I stayed in Amsterdam. But I would like to feel aware again, as used to be, that those who bestowed that love [for science] upon me, or made it grow, would give me their sympathy.

Together with post cheque [equivalent thereof] of 27 Guilders (45 Mark ....

**S II-E1d**
[M…008-356] [J.K.L. Martin (1851-1942) was Professor in Geology, director of the National Geology Museum, Leiden. See Albers and de Vos, 2010.]

Prof. K. Martin in Leiden

Tulung Agung, 20 Nov 1894

Highly learned and esteemed Sir,

Already several weeks ago now, I sent to your address some photographs for which circumstances have prevented me of writing an explanation. The photographs, seven in total were taken of those specimens within my collection which I consider to be the most precious. Although to me the prints of these do not suffice (mainly because of the poor quality of the light-sensitive paper), they may yet serve as an addition to the images in my treatise of which I hope you have received a copy. Given that the whole of my collection is intended for the Leyden Museum, I assumed to do well by sending these images in advance of the shipment of the specimens, since even by the most secure road there is always some risk. The hue on the photographs nearly matches their natural hue, as well as that of all other bones originating from below the lowest water level in Trinil and elsewhere. The photograph was taken with a 64 mm Steinheil's Antiplanet [*camera*], the outline [M…008-357] is thus most accurate, more so than that of the femurs on page 17 of my treatise which were taken with a smaller lens. Also, the femur frontal view is not sufficiently accurate in the middle which in print has been made too thin. Please read on page 8 fifth line from above "*viel*" [*much*] instead of "*nicht viel*" [*not much*].

Through the intervention of the Ministry of Colonies, my collection will be sent to the Leyden Museum in three shipments of about 100 crates each. The first shipment can be sent right away as soon as I have received authorization from Batavia. The other two can then follow quickly. I hope to finish packing them in no more than a few weeks.

I hope there is no objection to leave the crates unpacked until I have arrived, which, if the plans of the government are coming true, will be somewhere in the middle of next year. Before that, I will visit the Indies Museum in Calcutta and find spots in the Siwalik hills and the Narbada valley.

Summarizing the findings and results of my research I cannot say that these have fulfilled my expectations as far as the completeness of the specimens is concerned. Most are only very fragmentary ones leaving for instance species that are not very rare like the small *Boselaphus* (*kroeseni*) [*Duboisia santeng*] without even a complete skull and of *Stegodon*, of which hundreds of molars and fragments of molars were collected, but just two skulls, which only in liberal terms can be called almost complete. However, the number of them is large enough that most can be mentally reconstructed since they complement each other. But a few finds, although equally disappointing in regard to the just mentioned incompleteness, have exceeded my expectations as to their nature. The very first by far, are those of *Pithecanthropus erectus*, although I had suspected the existence of such a form (my essay in the '*Natuurkundig Tijdschrift van Nederlandsch Indië*' of 1888), and further the giant pangolin which I propose to name *Manis palaejavanica*, of which some 20 bones or bone fragments all correspond to their similarly named counterparts of *M. javanica* [>>M…008-358] albeit that those of the first mentioned [?] are slightly reduced and that they exceed that of the extant species 2½ to 3 times in size.

Uncertainty that has constantly accompanied my work has had a detrimental influence, since I never knew if my research would be continued. If I am allowed to express a personal feeling I would say that if I could have foreseen the misery resulting from this, it would have been sufficient to have prevented me from even thinking of such an enterprise, to which I have sacrificed my whole career, my health and my good humor. I will not even mention my wife and children.

I believe it to be justified in sharing these personal experiences, because I flatter myself that you might find thus a reason to allow me, after my arrival in Leyden, to pluck as many fruits from this preliminary work as possible. In any event, I am gratefully convinced that you are prepared to cooperate with me.

Whilst offering my polite greetings I remain with the highest esteem,

your willing servant, Eug Dubois

[The photographs of Dubois' precious specimens have not been located. The prints must have been sepia tone to match the "natural hue" of the lB fossils.]

**S II-E2**

[February 15, 1940, letter to a colleague concerning the Femur I discovery (DUBO9907)]

Haelen near Roermond

February 15 1940

Dear friend Woerdeman:

I am not sending you a bundle of letters from Kriele and de Winter, but additional letters from these Army corps of engineers (Genie) workers, extracted from a bundle of all sorts of letters and papers that are only indirectly related to my paleontological investigations on Java such as monetary accounts etc., similar to a number that I have previously sent that were intended only to recognize the nature of the exchanges. These are the only two important letters in which something is written about the exact spot and level from which the femur of *Pithecanthropus erectus* was excavated in 1892.

At that time, Kriele and de Winter were both placed at Trinil and sometime later I received from the pit under supervision of de Winter (very close to the one of Kriele) to my great surprise a very human like femur, not recognized by them, which is logical since no one had ever shown them the nature of this kind of bone.

From these letters we can mainly determine that the femur was found in exactly the same bed as the Skullcap and at a distance of only about 12 meters.

After I had found the femur in the shipment of 6 crates of August 15 (1892), I immediately inquired about the pieces of bone that were missing in the popliteal region. The answer to that is contained in the first two paragraphs of the letter of 31 August 1892.

With my best greetings and wishes for you

Eug. Dubois

**S II-E3**

[Femur II, III, IV and V attributed to the 1900 excavations.]

[In June, 1932, during the final stages of registering fossils from his Java collection, nearly 40 years after the discovery of Femur I, Dubois and his assistants recognized three new partial *Homo erectus* femora, found as eight femoral fragments in “a lot of inconsiderable fragments of ribs from different Trinil mammals” (Dubois, 1932: 718). This particular lot was collected in 1900 with Dubois (1934: 140) noting by way of further explanation,

> “During the time of my residence in Java, I had direct occasion of examining all the fossils obtained in excavation under my direction, and at the end of 1895, I did not dispatch to Holland such voluminous objects of acknowledged lesser value as fragments of real large ribs. Through unfavorable circumstances I did not recognize the femur fragments in the year 1900.”

Two of the femoral fragments “bears the inscription *Trinil*, in the handwriting of my former technical assistant at the explorations and excavations, the late sergeant-sapper Kriele” (Dubois, 1934: 139-140).

However, according to Dubois (1932: 719), “the exact site of these fossil remains in the [1900] excavation is not known, because they were not remarked as uncommonly important and [therefore became] mixed up with fragments of different ribs and some other fossils of small value. It is, however, possible to ascertain that the distance of the three was between 16 and 48 m. Now considering that the skullcap and femur I were found at only about 12 m the one from the other (perhaps only 10 m, certainly not 15 m, as appears from my notes of that time), in part of the excavations where corrosion was generally less, whereas the surface of the three new thigh bones is intensely corroded... [footnoted, as follows] Femur I is quite unaffected by corrosion, the skullcap yet less than the new femora.”

The nearest point of discovery for these femora, by Dubois' reckoning, was 16 m from the Skullcap and Femur I find spots. Skullcap and Femur I were found 12 m apart, perhaps on 10 m but not the 15 m Dubois earlier stated.

By "corrosion" Dubois (1934: 140) means that the original "periosteal bone layer … was partially or entirely worn away." In the case of Femur II-IV "the corrosion was restricted to the … periosteal bone layer." Dubois (1932: 718-719) further reported that:

- Femur II was "a foot long, still partially covered with rock …. a pyritous impregnation of andesite," the pyrite of which was unweathered, and is the most complete of the three new specimens with partial preservation of the superior diaphysis; the surface of Femur II, a right-side element, is the least corroded of the new specimens; as a complete element, it would have been 4 cm longer than Femur I;
- Femur III, 32.5 cm long and a left-side long bone, was found as two pieces of shaft that refit; like Femur IV, Femur III was found in the collection "nearly bare of rock;"
- Femur IV, 31.5 cm long and a right-side element, was originally about the same length as Femur I

An additional fragment was recognized in late 1932, Femur V; "corrosion most strongly acted on femur V, and laid bare the internal structure over the whole surface of the fragment;" presumably the s Femur V also was inscribed by Kriele with the site name Trinil. See Shipman (2001: 407) for a photograph of the five Trinil femora.

**<u>S II-E4</u>** [M…059-220. Dubois' '1900 Site Map" used in Figures 6a, main text, and S I Figure 7.]

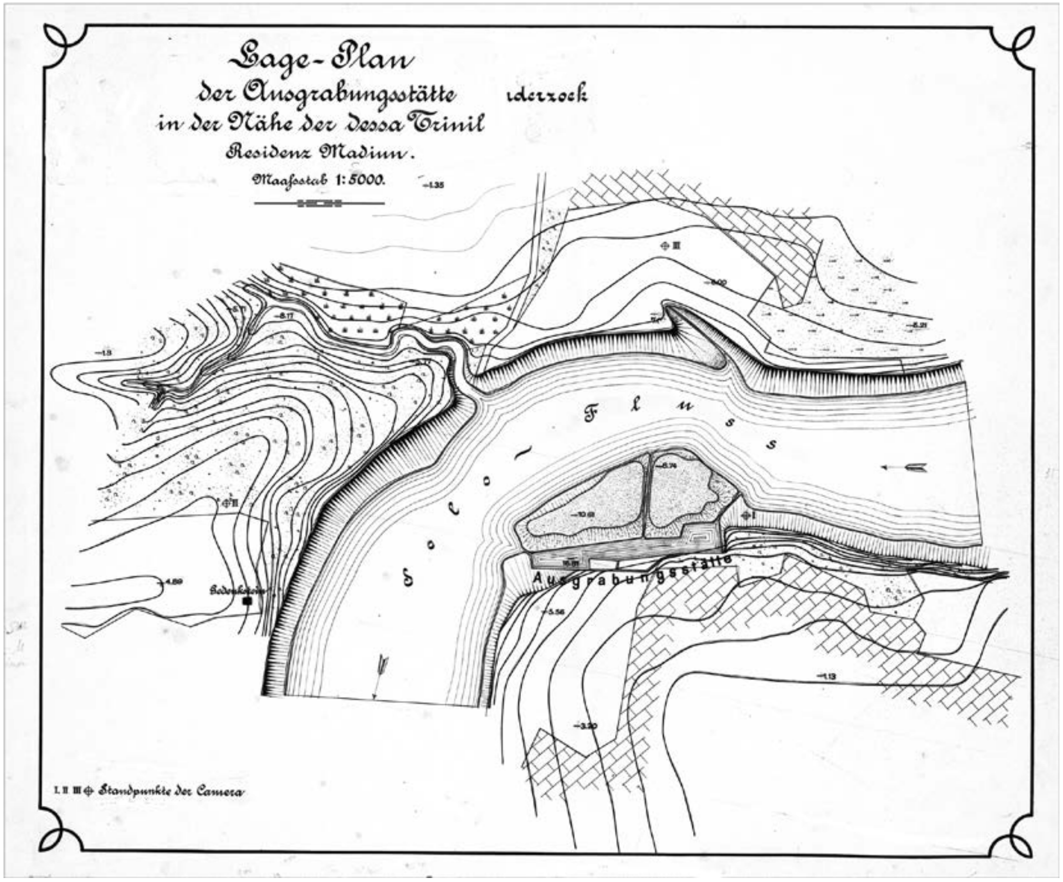

**S II-E5.**

[Dubois' lantern slides (of unknown dates) showing unpublished versions of his cross section of the *Pithecanthropus erectus* discovery site (upper image is scan DUBO9785; lower one is DUBO3236).]

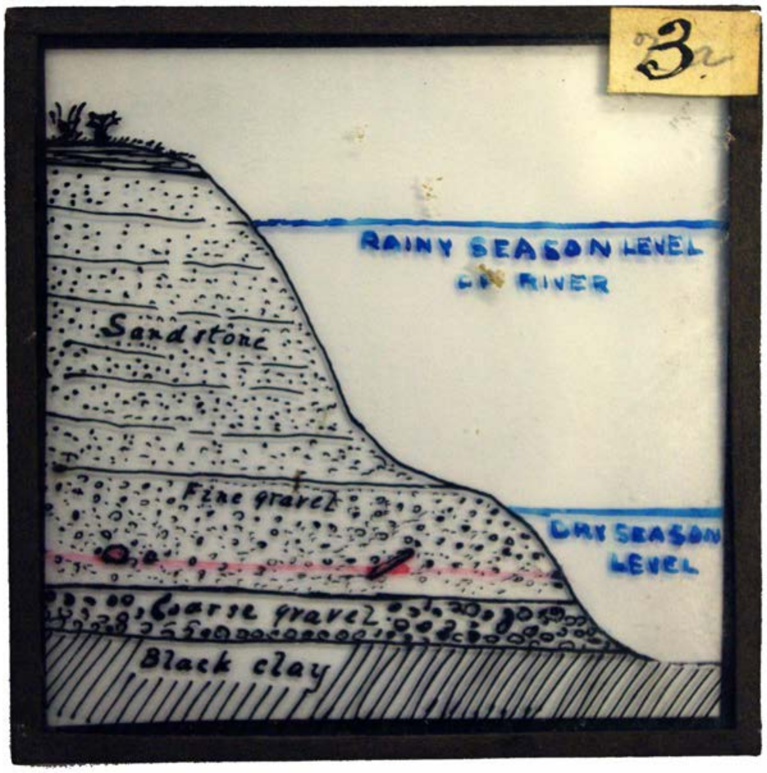


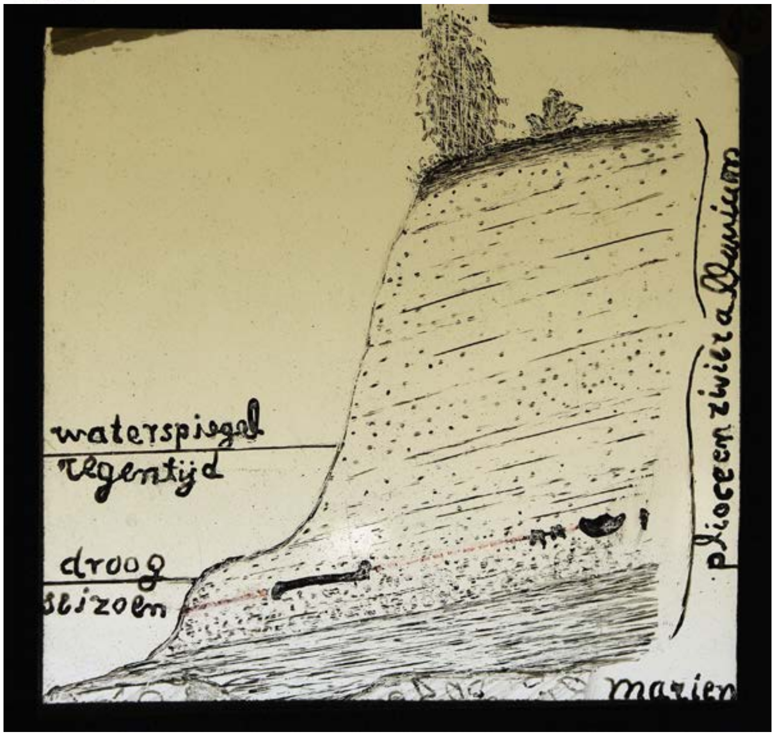

**S II-E6.**

[Dubois' unpublished map of the Trinil area, ca 1900 (de Vos and Sondaar, 1982)]

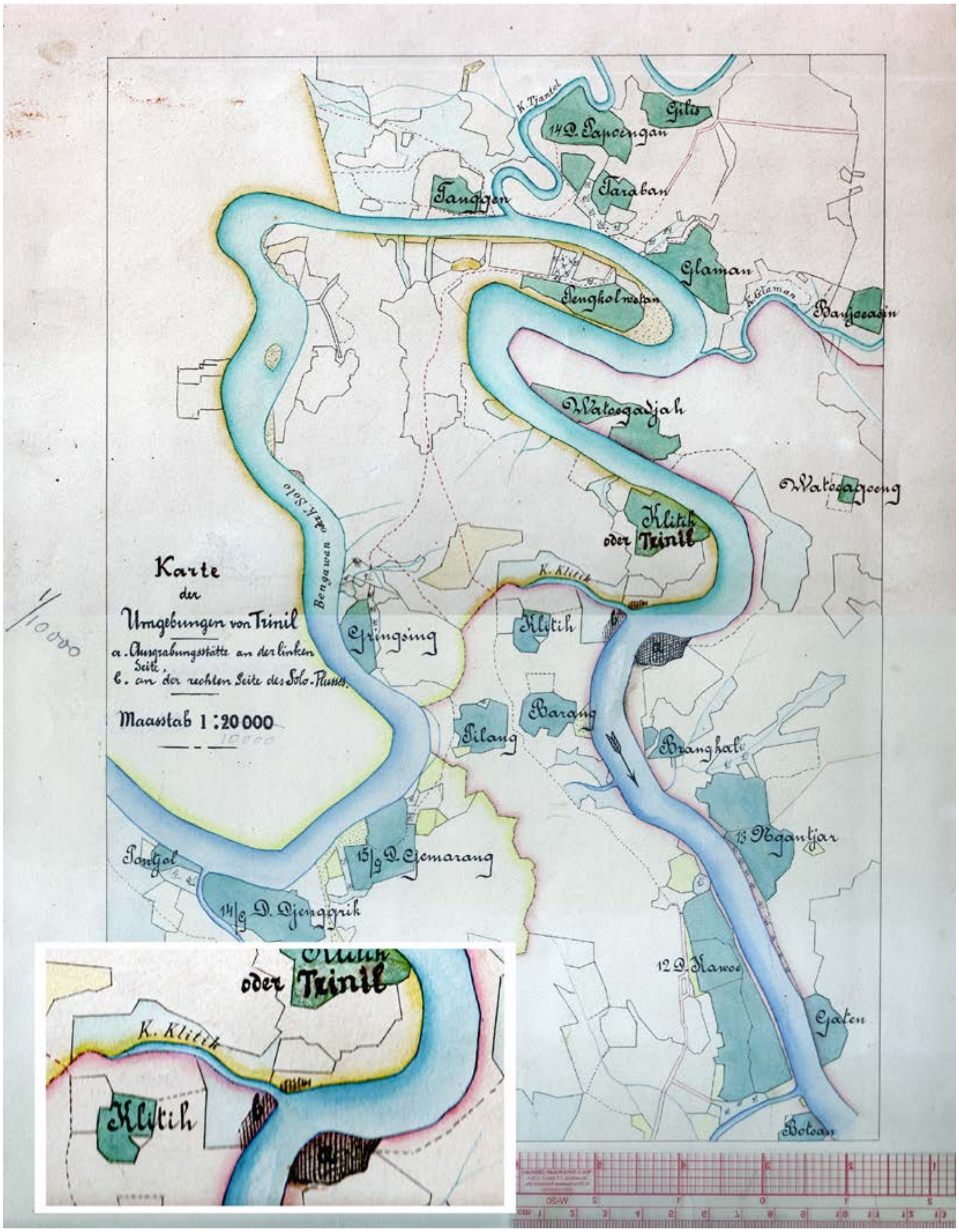


[Translation of the title and legend: "Map of the Trinil area *a*. Excavation locations on the left bank. *b*. On the right bank of the Solo River. Scale 1:2,000 [crossed out and '1:10,000' is written]. Our inset shows the left-bank ("*a*") and right-bank ("*b*") excavations, with the latter lying astride the outlet of Klitik Creek, east and northeast of the present-day Trinil Museum. See –A1b and –A4c, above, for Kriele's only representations of the right-bank excavation.]

**S II-E7.**

[Modern maps reveal many changes in place names compared to 1890-1908, as illustrated here by a portion of the modern topographic maps covering the Trinil area]

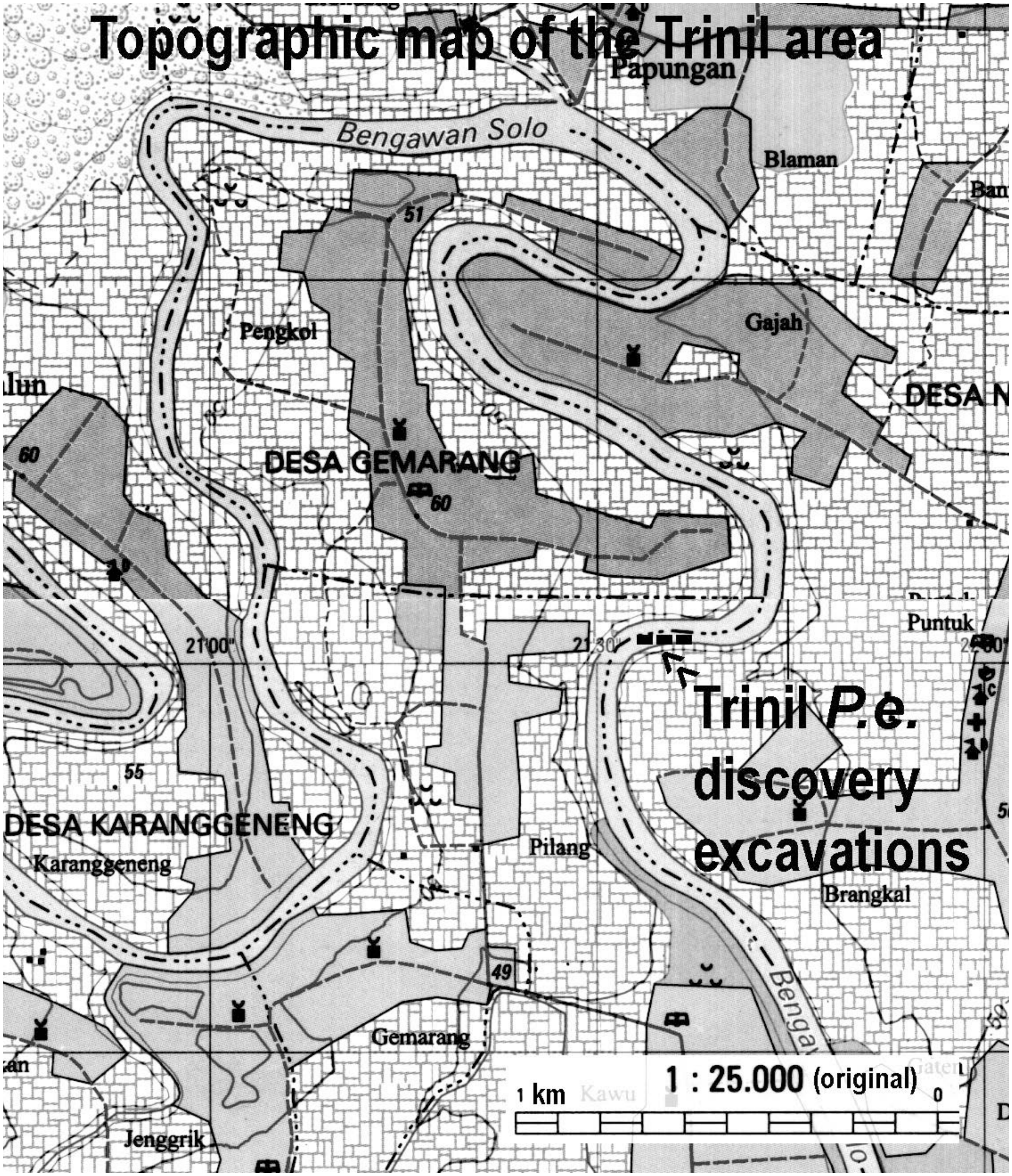

### S II-F. Dubois' published accounts of the main bonebed, 1895-1896, 1907-1908 and 1924

[The fossils that came from the left bank are in red while those on the left bank remain in black.]

**Foreword regarding Dubois' various versions of the site cross section (Figure 2a, main text)**

Dubois produced multiple versions of his site cross section in 1895-1896. They were accompanied by legends on the stratigraphy written in several languages with unit characterization which differ slightly when translated into English. The invariant features of the cross sections are: The topographic profile of the bank; the wet- and dry-season river levels relative to this profile; the presence of a conformable stratigraphic sequence under the bank; the occurrence of Skullcap and Femur I at one stratigraphic level; a wide spatial separation between the two finds within this level; and the presence of claystone below the discovery unit; and the Skullcap having been embedded deep beneath the incised embankment of the Solo River instead of the actual discovery location where the incision had exposed the discovery bed (Figures 2a, 3b, main text).

Despite diagrammatic elements and details which differ from one version to next, the cross sections portray accurately the stratigraphic fundamentals of the left bank. For instance, the version presented in the main text shows the *"Bed of lapilli-rock"* (our LB) and an overlying sequence of *"Sand rock"* (the rest of our units 2 to 5) capped by soil along the terrace upland south of the river (Figure 2a, main text). No younger formation is portrayed in between.

However, Dubois' 1895-1896 cross sections suggest that he left Java unsure of key aspects of the site geology, or was error-prone in making this type of geological illustration. Readily noticed, the strata have a north dip toward the river in some editions of cross sections (-F1, -F3 and –F4; also, -F10). Other versions of the cross section do show flat-lying beds (Figures 2a, main text). In 1894, Dubois had noted that *"the bone bearing beds of sandstone-like andesite tuffs and conglomerates … are horizontal only at Trinil"* (-B7c and –B7d). His 1895-1896 papers, and several later geological studies, refer to a south-dipping monocline in the Trinil area. Evidence present here shows the strata in the excavations on the left bank were essentially horizontal. No apparent north dip is possible in either situation.

The dip of the strata is not the only disconcerting inconsistency in the 1895-1896 cross sections. The top of the LB is lower in its relative position to the invariant features in one edition compared to the others, and the thicknesses of the LB and conglomerate change slightly one version to the next, despite Dubois' consistent reporting that together they were 1.5m thick (-F1 to -F5 and –F10).

While these puzzling miscues are reasonably attributable to illustrative imprecision on his part, there is one inconsistency between Dubois' graphic portrayals of the stratigraphy and contemporaneous accounts that is still inexplicable. The LB unquestionably cropped out above the LWL in 1891 (Figure 3b, main text). According to Kriele's letters from the field, the LB was exposed where the Skullcap Pit was dug; some fossils in the outcrop originated as high as 0.75m above the water line. Dubois himself even initially reported that the *"discovery spot … was one in the currently dry portion of the Solo River."* KdW also prescribed that the Skullcap and Femur I were from the Principal Fossil Zone (PFZ) laying at the seasonal low-water level or 0.20 m below it, as mention above (-A1e, -A2j, -A2k, and -A2q).

This well-documented relationship between the LB and LWL is not how Dubois later showed and reported the situation. By 1895-1896, his *"Level in which the four* [hominin] *remains were found"* (the PFZ) sat *"about 1 meter below the dry period water level"* (Figure 2a, main text). Indeed, all of *"Bed of lapilli-rock"* (the LB) in his cross sections lies below during the dry-season water level (Figure 2a, main text), despite Dubois' concurrent contention that *"the first bones"* came from exposures in a *"flat rocky ledge"* above the river level. This portrayal evidently was intentional, since Dubois showed the same relationship in each version of the cross section (-F1 to –F5). We have not determined the reasons behind the change from original reporting. We give priority to that reporting, however, because of its firsthand contemporaneous character, and therefore accept that the LB was initially exposed and the PFZ lay at and just below the 1891 low-water level, the LWL.

**S II-F1**

[Dubois (1896b: 251-255). Translated from German. Dubois' first formal European presentation was made on Saturday morning, September 21, 1895, in Leiden, The Netherlands, to Third International Zoological Congress (TIGC). At this point in time Dubois' knowledge of Trinil was limited to the information he had from 1891-1894 operations there.]

During the period 1890 to 1895, I conducted investigations on Java … vertebrate animals that inhabited this area in earlier geologic times. … [An] older fauna is primarily composed of extinct species and points to an unmistakable association with the late Tertiary and Pleistocene fauna of British India. The main area for discovery of this second and older fauna is in the hilly Kendeng range, which stretches out over a distance of about 100km… . The terrain in which the vertebrate remains were found can be as wide as about 3 to 5 km on average.

Bones are present within beds of tight and hardened volcanic tuffs, consisting of clay, sand and lapilli rocks. These tuffs suggest a fluvial origin, especially indicated so by a strong general presence of fresh water animals and by certain fluvial structures that English geologists call current bedding.

In the entire theater of their presence, the beds have undergone considerable deformation from folding, with the result that from east to west they have an overall dip angle ranging from 3° to 15°, generally in a southerly direction. The formation reaches a maximum thickness of more than 350 meter. It is underlain by discordant bedded Pliocene marine marls, sandstone and limestone beds. The vertebrate fauna within [the tuffs] is uniform everywhere in the Kendeng and other discovery regions on Java … . This fauna is most likely of late Pliocene age, but in any event not younger than very early Pleistocene… . [and] consists almost exclusively of extant genera (of these, *Stegodon* and *Hexaprotodon* are only sub-genera). …

In August 1891, I came upon a particularly rich find spot near Trinil … at the foot of the Kendeng range. In that year and the following ones, I discovered amongst a large number of skeletal remains of other vertebrates there, the bones and teeth of a large human-like mammal. … The steep banks of the river Bengawan or Solo from upstream of Trinil [eastward] to Ngawi, 12km wide *linea recta*, are exclusively built from the described rocks of hardened volcanic sands and lapilli. The beds in this strip of land have a general north to south dip of about 5º [see also, -B7c]. They are also exposed in the nearly flat surroundings of Trinil, at many places just below the thin agricultural topsoil [Figures 2a and 8, main text]. [Here] the river has incised into these sediments as deep as 12 to 15 meters, relative to the dry period water level. … The first bones were found in the rocky shores … . were an antler [***Axis lydekkeri***] …, a molar of a ***Stegodon*** [***trigonocephalus***] and several other skeletal remains…

[The fossils] were chiseled out of the flat rocky ledge that reaches out into the river from the foot of the steep river bank [referring to the left-bank outcrop and the Skullcap Pit; -A1b, -B2d, etc., above]. Excavations were organized such that in the beginning sediments were carefully removed in thin layers. The beds in this outcrop, which is on the left shore of the river, consist from top to bottom of variously colored and loose sandstones that become slightly coarser below dry-season ground-water level with increasingly more lapilli becoming part of the composition ([Figure 2a]). These predominate in the deepest bed, about 1-meter thick, which in turn transitions downward into a ½ meter-thick conglomerate bed that primarily consists of about walnut-sized rock fragments. Distinctly separated and below this bed follows an almost black claystone, rich in coal but in which bones no longer occur.

The latter increase in numbers from top to bottom [of the sequence beneath the terrace upland], so that the lapilli bed becomes the richest [in fossils]. The conglomerate bed, on the other hand, contains only a few bones. At the rocky ledges [in which the excavations of 1891 began], the top sandstone beds [those above the LB] had already been removed by river erosion, making the richest beds readily accessible.

The described molar [**1891 Molar**] and skullcap [**Skullcap**] were found in the lapilli bed on the left shore among hundreds of other skeletal remains and were extracted from exactly the same level [LB; see also, -B6e and –B6f]. The [***Pithecanthropus erectus***] molar was found first, in September. Then after expansion of the trench in October, the Skullcap was found at a distance of only 1 meter [away]. The mammalian species occurring with them in the same bed are for the most part extinct, most probably no longer living on Java. The most numerous finds were of a *Cervus* [***Axis lydekkeri***] species… . There are also many ***Stegodon*** [***trigonocephalus***] remains. One or two *Bubalus* [***B. palaeakerabau*** and presumably *Bibos palaesondaicus*] species appear … . A *Boselaphus* [***Dubosia santeng***] in fossil form is certainly distinct from what we know of living species. A ***Garialis*** [***Gavialis bengawanicus***] and a ***Crocodylus*** [***siamensis***] are very similar to extant species in India… . Further represented were the genera *Leptobos* [there are no *Leptobos* in the Trinil Dubois Collection], ***Rhinoceros*** [***sondaicus***], ***Sus*** [***brachygnathus***], *Felis* [***Panthera tigris***], *Hyaena* [there are no *Hyaena* in the Trinil Dubois Collection today] and others.

Compte-Rendu 3me Congrès Intern. de Zoologie. Planche II.

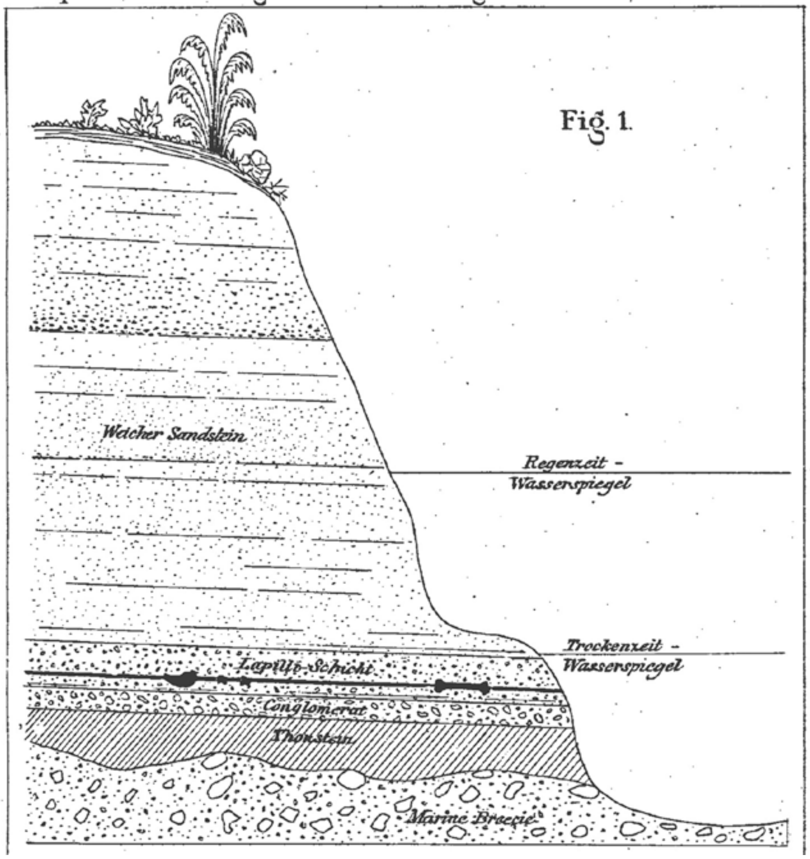


[As drawn, the top of the Lapilli Schicht (Lapilli Bed) is just below the Trockenzeit Wasser-spiegel (dry-season water level); the Lapilli Schicht is about twice as thick as the “Conglomerat” unit below it; the Skullcap-Femur horizon is in the lower part of the Lapilli-Schicht. However, given a ~1m thickness for the the Skullcap/Femur I stratigraphic level would have been ~¾ meter below the dry season river, rather than the 0.00-0.20 below the LWL, which KdW reported while excavation was underway.]

After our work of that year had been halted due to a swollen river resulting from the start of the rainy season, it was resumed again in May 1892 at the beginning of the dry season. This time, a new trench was cut on the left shore about 25 meters long and 12 meters wide which included the not yet excavated remaining portion of the older, almost circular pit whose diameter was about 12 meter. This resulted in the discovery of large numbers of bones, particularly in the deeper beds. In August we found amongst these [fossils] the left femur [**Femur I**] at the same level of the lapilli bed in which the skullcap and molar [**Skullcap** and **1891 Molar**] had been found and at a distance of about 15 meters from them. And finally, in October, at about 3 meter from the spot where the skull cap had been found (probably even closer) a second molar [**1892 Molar**] was found in the direction where the femur had been excavated. I have not yet described this specimen, because I found it only later among a number of teeth that were excavated at this spot.

This explanation of the geologic relationships under which the four skeletal parts have been found is hopefully sufficient to convince you that the finds surely are not only derived from one and the same completely undisturbed bed, most likely of late Pliocene age, but that they also were deposited at exactly the same level and thus must all be of the same age. The rather flat relief of the surface [horizontal bedding], among other things, argues against the possible theory that these specimens originally belonged to an older bed and were subsequently reworked. ….

The femur shows, as you gentlemen can easily observe, exactly the same degree of preservation and fossilization as the skullcap and molars, as well as all other bones that were excavated at Trinil. [The femoral fossil] is more than twice as heavy as a recent human femur of the same size. Its color is chocolate brown, and it is hard as marble and … . weighs 1kg, which is more than double the dry weight of a recent human thigh bone. ….

**S II-F2**

[Dubois (1895b: 151 -160). Translated from French. Resume d'une Communication de le Dr. Eug. Dubois. Summary of an October 1895 oral presentation about the *Pithecanthropus erectus* of the Java Pliocene. Bulletin de la Société Bèlge de Géologie, de Paleontologie et d' Hydrologie, Bruxelles.]

[151] Session of October 29, 1895 Proceedings

The bones were discovered at Trinil, located very close to the town of Ngawi in the central part of the island of Java, during searches related to the excavation of a fossil fauna, which Mr. Dubois was in charge of during a period of six years and which was subsidized by the Netherlands Indies Government. The explored sedimentary rocks consist mostly of sandstone and hard clay and localized conglomerates. The entire sequence represents a sandy formation that is more than 100 km in length and is found in outcrop over 1 to 5 kilometers.

[151>>152] Mr. Dubois has discovered a large quantity of bones. He has reported that he found almost 400 crates worth of bones from these excavations, containing remnants of vertebrate animals, in particular those of all kinds of mammals. The exhibited bones were not found all together, but in relatively close proximity and at the same level within the deposits. The skullcap was found at a distance of 1 to 3 meters from the molars, the femur about 15 meters away from the skull cap, but in the same bed and at the same stratigraphic level. The nature and appearance of the bones [of *Pithecanthropus erectus* presented] are similar to those of a variety of animals that were recovered from the sandstone formation. Mr. Dubois considers these remains to have belonged to one and the same skeleton. He has observed that in cases where a number of animal bones that undoubtedly belong to the same skeleton, these were found at similar distances from each other.

The femur [Femur I] has a strong resemblance to a human femur. The shape is very human in character and in this respect there is perfect agreement by at least the majority of anatomists and zoologists. Diverging opinions have been formed in particular about the skull cap [Skullcap]: some see it undoubtedly as a human skull cap, others with the same strong conviction as one of an ape.

It is highly unlikely that the geologic formation from which these bones are derived is Pleistocene, because the encountered species of mammals and reptiles did not have any of the characteristics of the modern fauna. In regard to the beds themselves, these are dipping over a long [outcrop] range at 3º to 15º and this dip cannot be attributed to any local or regional volcanic activity. The geology and the fauna indicate that the geologic bed in question is of Tertiary age and belongs to the Pliocene.

The fauna appears to be older than that of the early Pleistocene [Diluvium] of the Narbudah valley, which is considered to be part of the oldest Pleistocene, but it is certainly more recent than the early Pliocene fauna of the Sivalik mountains. It can therefore only belong to the late Pliocene, if the faunas to which it is compared have been placed at their correct geologic level.

If the femur [**Femur I**] is human and the skullcap [**Skullcap**] that of an ape, we would then have at a distance of 15 meters in a horizontal direction and in exactly the same level of deposition, meaning being deposited at the same time, the first Tertiary human and the most paradoxical anthropoid of all known extant and fossil apes that we have here, which would hardly be likely at all. There have been suggestions that attribute the skullcap to that of a gibbon, but it is much too large for justifying this hypothesis and would at least concede that the [152>>153] fossil gibbon would have had a much larger stature and a weight three times heavier than that of a large gorilla.

The capacity of this skull may be evaluated at 1,000 cubic centimeters. Mr. Manouvrier has arrived at the same number via a different method. The cranial volume is thus clearly well established. There are human craniums that have a similar small capacity, but only in humans of small stature. Now, if the femur belongs to the same creature, it would have had a height of 165 to 170 centimeters. The skullcap would thus be much too small for a normal human cranium.

Mr. Cunningham has suggested that the skulls of micro-cephalids would be comparable especially in profile, but the parietal region is much more flattened in this actual case than it is in the profile of the micro-cephalid presented by Mr. Cunningham as well as those of all other micro-cephalids. The bony frontal brow ridge is also more prominent and in particular the zygomatic apophysis of this bone is much thicker than in all human skulls even the pathologic ones. The absolute size of the skull is also much larger than those of micro-cephalic humans.

Next, Mr. Dubois moves on to a comparison with the Spy skulls, as described by Mr. Fraipont. He reaches the same conclusions as with the previous comparison with micro-cephalic humans. There are even larger differences in the lateral orbital parts.

The frontal zygomatic apophysis of the Trinil skull cap is completely ape-like. It is also two times thicker than in the most ape-like one from Spy and the temporal narrowing is also located towards the back like it is in apes. It is only possible to convince oneself of this important special character by examining the original [specimen]. Based on these facts, the Spy skulls are clearly human. The Spy skulls are also much larger.

SÉANCE DU 29 OCTOBRE 1895 157

FIG. 1.

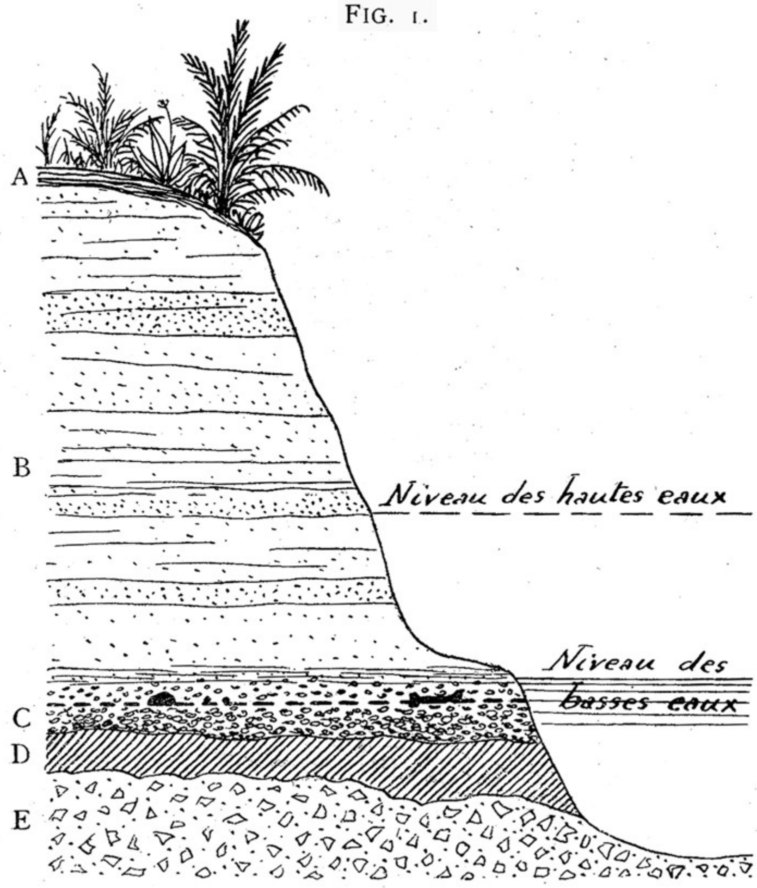


*Coupe des couches ossifères à Trinil.*

A. Terre végétale. — B. Tuf gréseux volcanique. — C. Couche de lapilli et de cailloux. — D. Limon durci. — E. Brèche marine.

[The discovery horizon is in the "C. Couche de lapilli et caillaux," in which the upper lapilli facies is combined with a lower, conglomeratic and fossil-poor one. In his previous version of the cross section (-F1), the beds dip slightly toward the river (north), while in the cross section used in Figure 2a, main tekst, and in this paper (right), the beds are portrayed as flat lying.]

Related to the femur, the majority of anatomists and zoologists consider it to be human. There are some differences. Besides, human femurs exhibit also differences amongst themselves, but the deviation is much too great in this case.

Mr. Dubois has compared 150 specimens and Mr. Manouvrier has likewise examined human femurs for the same purpose but nearly twice as many and the result of these comparisons and examinations is that the fossil femur does not range within the bounds of the human variations. Great differences are mostly in the popliteal region: the presented femur is nearly round in the section of this region. The internal facet of the diaphysis is very round, which ordinarily is not the case with human femurs, which almost always display an internal angle. The crest between the greater and lesser trochanters is less raised [153>>154] than it is on average in the human femur in which, however, one occasionally can find this characteristic, but which never can take on the form that this one has.

There is an objection that such differences can be found in extinct human races. This is a possibility that we therefore propose, but a possibility is not proof. Besides, as per Mr. Dubois, a femur of an anthropoid biped which in my opinion this Trinil creature has been, could not be very different from a human femur.

The femur is affected by pathological exostoses: all experts are in agreement that this has no influence on the general shape of the femur.

FIG. 2.

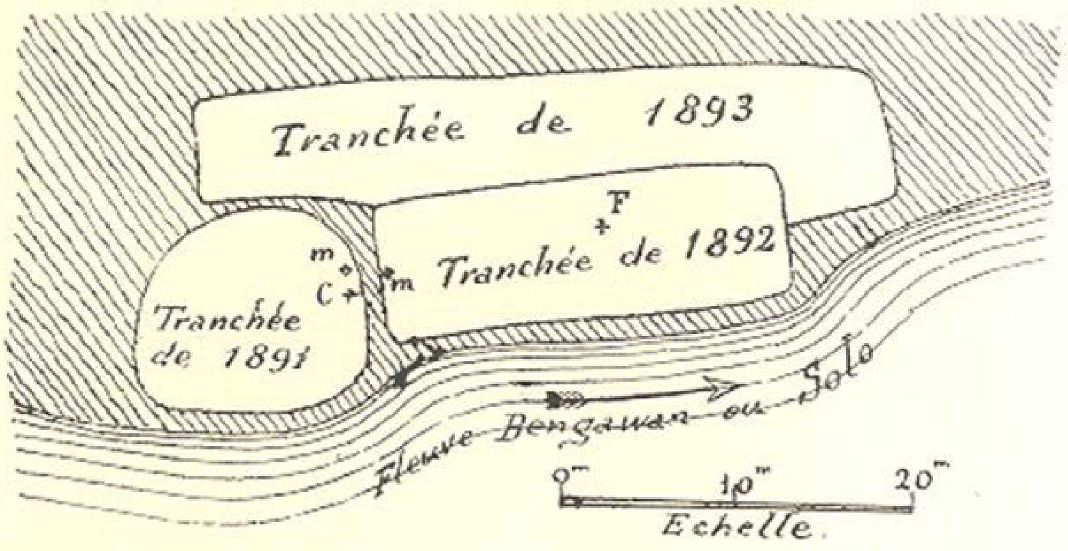


Virchow says that according to him, the healed decay which would thus be indicated appears to contradict the supposition that the creature who suffered from this illness could have been an ape. This creature which had been victim of it was given proper care. Nevertheless Virchow did not believe that the skull cap is derived from a human.

Continuing then with the teeth, Mr. Dubois shows the gathering that they consist of a third right upper molar and of a second left upper molar. They are not of the same size; the second one is less wide and is longer. The third one is wider and smaller in its antero- posterior diameter. Virchow claimed that these teeth could not possibly be derived from the same individual, but he had to admit that the dentition of two anthropoid apes that Mr. Dubois could show him at the Congress of Leiden, confirmed this.

The degree of fossilization of the teeth as well as their color, are the same. All of them show similarities in characteristics of roots as well as crowns. These teeth clearly belong to one and the same skull and this can only be the one near which they have been found. Judging by the shape of their crowns and roots they are rather ape-like.

Mr. Dubois ends his presentation with the following remarks which he has forwarded to the editors:

Finally, allow me to say a word or so about the communication by Mr. Pohlig about the anthropoid femur of Eppelsheim, a communication that Mr. President has just read to us. I am glad to be able to show you here a good cast of this ape bone, which initially was attributed to that of a child and subsequently much later to that of *Dryopithecus*. Whereas the fossil ape of Saint Gaudens belongs to a geologic level below the fossil bearing beds of Eppelsheim, [154>>155] Mr. Pohlig himself {Footnote: Sitzungsberichte der niederrhine. Gesellschaft für Natur unde Heilfunde zu Bonn, 1892, pp. 41-43}reported again in 1892, that he thought that by certain anatomical characteristics it has been proven that locomotion in this biped was already much more advanced than in the extant anthropoids that are closest to humans.

Careful examination of the fossil in the Darmstadt museum and numerous comparisons that I was able to make there and in addition also between the original and bones of living anthropoid species as well as between the latter and the reference cast, have convinced me that the femur by no means approaches that of a human femur nor of any gibbon femurs. It is above all very different from that of a 12 year old child, in which the epiphyses are very thick and also not welded together. On the contrary, in its entire shape and down to the smallest details, it is a femur that resembles that of gibbons. The only difference is that in this one is bit larger.

What Mr. Pohlig has taken for the 'rough line' is nothing more than its exterior lip which, just like it is in gibbons, has developed in the shape of a crest. And about the two other lines that Mr. Pohlig discusses, indicated in his sketch by *aa* and which he regards as proof of the use of the bone for bipedal locomotion, I could only find a single one on the original. The other one, by the way, does not occur any longer on the human femur. The third one (*b*) is the gluteal tuberosity which is equally well expressed in the gibbon femurs.

By comparing the cast alongside gibbon femurs, it will then not be necessary to insist, based on these facts, that the condyles do not resemble in any way those of human femurs or of the one of *Pithecanthropus* and that their shape is of the same class as those of *Hylobates*, and any other ape not indicating bipedal motion.

There are thus good reasons for seeing in the ape which has left us its femur at Eppelsheim a close ancestor of the gibbons. But since a single bone is not always sufficient to determine a species or a genera, I rather prefer to separate the fossil ape from the genus *Hylobates* of which Mr. Pohlig is speaking and names it *Pliohylobates Eppelsheimensis*, to indicate at the same time its relationship with *Hylobates* and its provenience.

In ending his presentation, Mr. Dubois thanked Mr. President and the Assembly for the opportunity that he was given to show, in front of an attentive and knowledgeable gathering, the paradoxical bones [155>>156] of *Pithecanthropus* and he offered them at the disposal of the members of the Society who would like to obtain additional information.

Mr. President thanked Dr. Dubois for his presentation and expressed the hope that the author would be willing to offer a written summary for our publications and he then declared the discussion period as opened.

Mr. Van den Broeck expressed his wish for Mr. Dubois to pay particular attention to further details and additional refinement of the nature of the deposits. That is the recognized rational basis on which one could attribute full value to the interesting discovery and no detail in this matter would be superfluous.

Mr. Dubois then provided the detailed illustration of the deposits, shown here in Fig. 1, which allows one to make a synthetic evaluation. The deposits are located in the central region of the island of Java, in a fossil bearing terrain composed of fluvial beds with a total thickness of 350 meters. In this place, they were deposited as a sandstone formation which is usually yellowish – grey in color, but blueish in places where it has not been aerially exposed owing to protection under water such as at the river edges. This sandstone deposit is in fact best described as bedded volcanic tuff (See Fig. 1 B).

At Trinil, the Solo River (Bengawan), the largest on Java, has incised this formation to a depth of 12 to 15 meters below the overall surface of the terrain. In this natural outcrop, the beds show a slight dip of about 5º to the south. One can follow this formation in outcrop along a zone that extends from east to west over a length of 100 kilometers. Fresh water (fluvial) mollusk shells are found of the species *Unio, Corbicula, Melania*, *Paludina*. The structures within the sedimentary deposits are equally fluvial.

At Trinil, the base of Formation B (see Fig.1), which is also the richest fossiliferous zone, consists of a kind of coarse sand of 1 meter in thickness, composed of lapilli, which rests on a zone of fine gravel (C) that has a thickness of 0.50 meters.

The base of this sedimentary sequence is a kind of indurated limestone (D), very dark in color, almost black, with an appearance of limy clay without stratification. On breaking, it does not split into thin sheets but into conchoidal pieces. In this respect, Figure 1 should actually be corrected since the contact between C and D should be drawn as a straight and horizontal line.

Near Trinil, these beds rest concordantly on a marine formation (Fig. 1 E) which, according to Martin, belongs to the lower Pliocene and is composed of marl, limestone and marine breccias. [156>>157]

It was at Trinil, a small hamlet near the village of Ngawi, that Mr. Dubois ordered excavations into these sedimentary rocks along the Solo river that produced the bones that have been shown at this meeting. He directed digging into the rocks up to 13 meters in depth of the original wall of the escarpment. The bones were chiseled out of the hard rock after they were exposed and these excavations at Trinil were carried out over three periods of 6 months duration, with the aid of 50 workers and under the auspices of the Netherlands Indies Government. The bones do not appear to be rounded; nevertheless, the formation [157>>158] is a fluvial one, which is confirmed by the type of mollusk shells mentioned above.

Among the vertebrate remains from this same formation, one can list the Tertiary species of *Stegodon* as well as *Bubalus, Bosephalus, Cervus, Hyaena, Felis* etc., whose fossil bones have the same color and show the same degree of mineralization [***Stegodon trigonocephalus***, ***Bubalus palaeosondaicus***, ***Dubosia santeng***, ***Axis lydekkeri***, ***Hyaena brevirostris*** (there is no fossil of Hyaena in the Dubois Collection), and ***Panthera tigris***, respectively (Table 1, main text)]. The bones are very heavy and the shown femur [**Femur I**] weighs at least 1 kilogram.

After showing this evidence, Mr. Dubois drew a map of the excavations and the stratigraphic sequence on the blackboard, material that has not yet been published and that is represented in Figure 2 just below, sent to us by Mr. Dubois, that clearly shows the relationship of the locations where the bones were extracted.

Mr. Lemonnier asked if the river had developed any alluvial deposits. Mr. Dubois answered he had done so very sparsely. Instead, the fluvial regime had been more of an eroding one than of alluvial deposits. There were even stretches, one as much as 12 kilometers in length, where the basement rocks were denuded.

**S II-F3(i)**

[Dubois, 1896c. Scientific Transactions of the Royal Dublin Society. Volume VI. I. “On Pithecanthropus Erectus: A transitional form between, Man and The Apes. By Dr. Eugene Dubois.” {Read November 20, 1895. {Published in English}. {Selections; see original for full text.}]

[1] In speaking of the remains, which are the subject of this Paper, I think the best way for me to take will be, first, to give an account of the circumstances attending the discovery, and further, to treat of the principal interpretations which have been given of them, including my own considerations.*

By order of the Netherlands Indies Government I conducted in Java, from· 1890 to 1895, explorations for a fossil vertebrate fauna, of which already some remains had been discovered, many years ago, by Junghuhn and others, and later extensively described by Professor K. Martin, of Leiden. I found a very large quantity of remains of mammals and reptiles, for the most part derived from extinct species, which show, as might be expected, an unmistakable relation to the later Tertiary and Pleistocene fauna of India.

The chief localities of these finds are in the southern slope of a range of low hills, the Kendengs, which extends between the residences Kediri, Madiun, and Surakarta on one side, and Rembang and Samarang on the other, in a length of about 60 miles. The area in which these vertebrate remains are abundantly found, {* The chief substance of this Paper was contained already in a discourse which I delivered in Leiden in a. session of the Second Section of the Third International Zoological Congress, on the 2lst September, 1895.} [1 > 2] in: many places, may have on an average a breadth of from one to three miles. They are contained in beds of cemented volcanic tuff, consisting of clay, sand, lapilli stone, which especially, through the very general occurrence of the remains of freshwater animals, and of that fluviatile structure which English geologists call current-bedding, or false bedding, prove to be of fluviatile origin. The strata have undergone, in the whole area, considerable disturbances by folding, on account of which they have, from east to west, dips of 3° to 15° in a general southerly direction. The whole formation reaches a maximum thickness of more than 350 metres. The strata rest, unconformably, upon beds of marine marl, sand, and limestone, recently determined by Professor K. Martin to be of Pliocene age. The fossil vertebrate fauna, which they contain, is everywhere in the Kendeng, and also in other places in Java, the same, and a homogeneous one. Its age can only be judged when the description of my collection, which I intend to give in the course of a few years, shall be published. But I have studied it already a little, and it can be said, in accordance with geological circumstances, and the relations which this fauna has with the Post-Tertiary and Pleistocene vertebrate fauna of India, that, most probably, it is young Pliocene; in no case, however, can it be younger than the oldest Pleistocene. For, whilst on the one hand the species surely belong almost exclusively to living genera-only the genus Leptobos and the sub-genera *Stegodon* and *Hexaprotodon* are extinct--and it must therefore be younger than the principal part of the Upper Miocene or Lower Pliocene Siwalik-fauna, including not a few extinct genera; on the other hand, the number of the extinct species seems to be in proportion somewhat greater than that of the Narbada-fauna, which is put in the early Pleistocene, Further, the inclination which the strata show does not well agree with a Pleistocene age.

In August, 1891, in the neighbourhood of Trinil (in the regency of Ngawi, of the Residency Madiun), at the foot of the Kendeng, I came upon a place particu-larly rich in fossil bones, and found there, in that and the following year, among a great number of remains of other vertebrates, bones and teeth of a great man-like mammal, which I have named *Pithecanthropus erectus*, considering it as a link con-necting together Apes and Man. These remains I held to be so important that I thought it necessary, notwithstanding the great incompleteness of my resources of comparison, to publish a provisional description in Java, especially because, through my very short reports given to the Netherland Indian Government, these :finds had · already raised more or less scientific discussions in Europe. . It is now a year ago since my description came into the hands of the most renowned anatomists and zoologists of Europe and America; it gave occasion to so much criticism that the great importance of the remains is now unquestionable. But in the interpretation of them a very large divergence is also apparent. This divergence may be [2 > 3] ascribed, chiefly to the very divergence of the characters of the remains, partly to the shortness of my statements. It appeared especially necessary to enter more in detail into the circumstances under which the remains were found. Concerning these circumstances, many conceptions have been formed which in no way answer to the reality. In the first place, then, let me explain the circumstances under which the remains were found.

From Trinil to Ngawi the steep banks of the Bengawan or Solo river, for an extent of 7½ miles, consist exclusively of the above-mentioned volcanic sands and lapilli, cemented into soft rocks, very much like the rocks which I saw in the Siwalik hills. The strata have in this area a general dip S. of about 5°, and are only concealed by a thin covering of vegetable soil. In these strata the Solo river has cut its channel, 12 to 15 metres deep, near Trinil. North and west of Trinil the Pliocene marl and limestone appear under them. When I first, species of deer, which is among the commonest of the fauna, a molar tooth of *Stegodon*, and a few other remains belonging to the same fauna. They were dug out of the rock by means of chisel and hammer, and the excavations which becomes coarser, whilst more and more lapilli occur in it, and the latter prevail in the deepest bed, about 1 metre thick, passing downward over into a conglomerate bed. Under this follows a bed of hardened blackish clay, sharply separated, which does not contain any bones. The latter in the sand-rock increase in number from higher to lower , so that the lapilli bed is the richest; the conglomerate bed, however, contains but few bones.

Among hundreds of other skeleton remains, in the lapilli bed on the left bank [3 > 4] of the river, the third molar tooth was first found in September; then, the hole having been enlarged, the cranium a month later, at about 1 metre distant from the former, but in the very same level of that bed. The species of mammals; of which remains were found in the same bed, are, for the greater part at least, extinct ones, and almost certainly none of them are at present living in Java. Among these remains we find a great number of the above-mentioned small species of Cervus, which certainly is not extant in the Malayan isles. Also many bones of Stegodon were found. One or two Bubalus species seem to be identical with Siwalik species; a Boselaphus undoubtedly differs from the known species, living and fossil. Further on there were found the extinct genus Leptobos, the genera Rhinoceros, Sus, Felis, Hyaena, and others; a G[h]arial and a Crocodile, differing little from the existing species in India, but which cannot be classed among them.

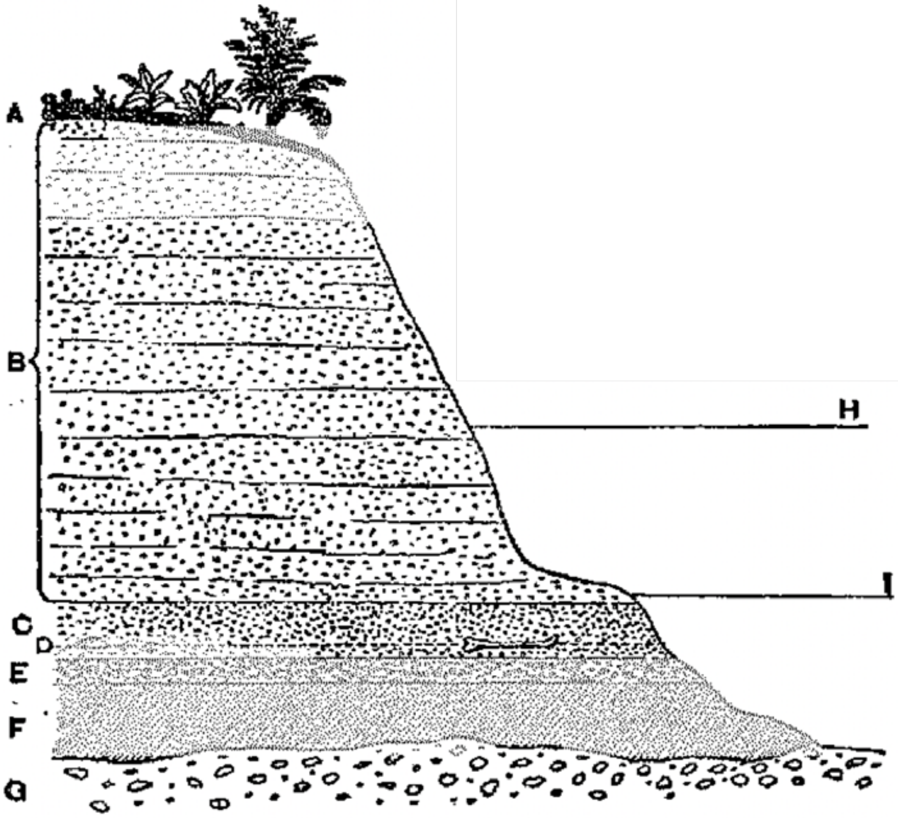


FIG. 1.—SECTION OF THE BONE STRATA AT TRINIL.

A—Vegetable soil.
B—Sand-rock.
C—Bed of lapilli-rock.
D—Level in which the four remains were found.
E—Conglomerate.
F—Clay-rock
G—Marine breccia.
H—Rainy season level of river.
I—Dry season level of river.

A doubt whether the four remains were once organically connected is certainly comprehensible, and was pronounced from different sides. Nevertheless, it seems to me that this doubt is hardly allowable, on account of the short distance of the places of discovery from ono another, for a distance of 15 metres is so small that as an argument against the supposition that the bones belonged to the same 'Skeleton, it cannot be considered as of more importance than if the bones had been found in contact with one another. I often found bones from the self-same skeleton, and even fragments of one bone at corresponding distances. I dare say that every paleontologist who has made any excavations for fossil vertebrate remains has had the same experience. I never found in one place anything like a complete skeleton, and, as certainly the bones once belonged all to complete skeletons, the bones must have been all dispersed. I have good reason to think that the animals perished in volcanic catastrophies, and that their corpses were brought down in the current of a large Pliocene river. Before, then, the bones were definitely deposited and buried in the old alluvia, they must generally have been separated through the rotting of the flesh, and torn the one from the other, and dragged away with the adhering flesh by crocodiles. Many remains of these preying water-reptiles, and also the traces of their teeth in spongy parts of bones, were found. So this argument against the assumption that the femur ascribed by me to the *Pithecanthropus* belonged to the same skeleton as the skull-cap, fails.

Seemingly as a better contra-argument it has repeatedly been said that the result of their anatomical analysis is opposed to that assumption [5 >6] others to an Anthropoid Ape, which, through the size of its skull, and some other characteristics, is the nearest to Man of _all known, living and fossil, Anthropoids. The molar teeth are judged of in the same manner. The femur, however, is so human-like that nearly all, after reading my description and seeing the specimen itself, did not hesitate to declare it to be human.

But up to the p1·esent moment no human remains have been found in the Lower Pleistocene; the oldest only reach down to about the middle of that period. From the Tertiary, however, every certain trace of the existence of Man is still m1ssmg. Moreover it is a fact that, with the exception of eome isolated teeth, only three small specimens of remains of :fossil Anthropoid Apes have been found.·

Considering, on the one side, this scarcity of remains of Anthropoids which all the researches in the whole world have brought to light up to the present moment, and on the other hand the complete absence of human remains from older strata than the Middle Pleistocene, it is certainly in the highest degree improbable that now, at once, should have been found the oldest human bone, which almost certainly goes back to the Tertiary period, and the largest fossil specimen yet known of an Anthropoid Ape, so paradoxically anthropoid that very experienced anatomists and zoologists have even taken it to be human. During five years' researches in an area hundreds of square miles in extent, and particularly rich in remains of one fauna, I did not find (with perhaps a single exception) anything which could in the slightest degree suggest the idea of a great Anthropoid or of a Man. And yet, at Trinil, those skeleton remains, as already stated, were lying close together in exactly the same level of the self-same strata, which are more than 350 metres thick, and include a homogeneous fauna. Let us imagine the proportion on a smaller scale: we have a layer of 1 millimetre (representing the thickness of the bones) in a strata-complex 3½ metres this and having an extent of some thousand square metres, and therein the four anthropoid remains ( the like of which were never found elsewhere) at a distance of almost 15 centimetres, the one from the other. Certainly the probability that these remains, being exactly contemporary, are from the same skeleton is --unless the result of the anatomical examination should be decidedly contradictory--many thousand times greater than the reverse, viz. that the bones were never parts of the same animal.

Now, however, the total result of anatomical considerations which have been taken by zoologists and anatomists respecting these remains is in no way in decisive contradiction to their belonging together. Some, as Professor Cunningham.* ….

**S II-F3(ii)**

[Dubois, 1896d: 241-243. Published in English.]

In the neighbourhood of Trinil the Bengawan or Solo river runs as far as Ngawi, a distance of 7 miles, between steep banks, composed almost exclusively of the above-mentioned volcanic sand and lapilli, cemented together and hardened into soft rock, similar to those in the Siwalik hills. In this area the strata have a general dip S. of about 5°, and are only concealed by a thin covering of vegetable mould; through these, in the immediate vicinity of Trinil, the river has cut its channel to a depth of 12 to 15 meters.

At points above and below Trinil, the Pliocene marl and limestone appear under these strata.

The first fossil bones discovered here were cut out of the rock by means of hammer and chisel, and consisted of the antler of a species of small deer, which is among the commonest of the fauna, a molar tooth of ***Stegodon*** [***trigonocephalus***], and a few other bones, all belonging to the same homogeneous fauna met with in other parts of the Kendeng Hills.

In making the excavations care was taken that the layers were removed one by one. They consisted from above downwards of alternately coloured sand rock which became coarser owing to a greater admixture of lapilli especially in the deepest layer, which measured about one meter in thickness, and which merged below with a conglomerate layer. Under this there is a sharply defined layer of hardened blackish clay. In the sand-rock the bones were found in increasing numbers from above downwards, so that the lapilli layer contained the most; in the conglomerate layer but few were met with ; whilst in the clay none were discovered.

It was in the lapilli layer on the left bank of the river, amidst hundreds of other skeletal remains, that the third-molar tooth [**1891 Molar**] was found. This was in September 1891; here also, a month later, on exactly the same level in that layer the cranium [**Skullcap**] was discovered at a point one meter distant from the spot where the tooth was picked up. The mammalian remains found in the same layer are for the most part those of extinct species, but it is almost certain that some of them are now living in Java.

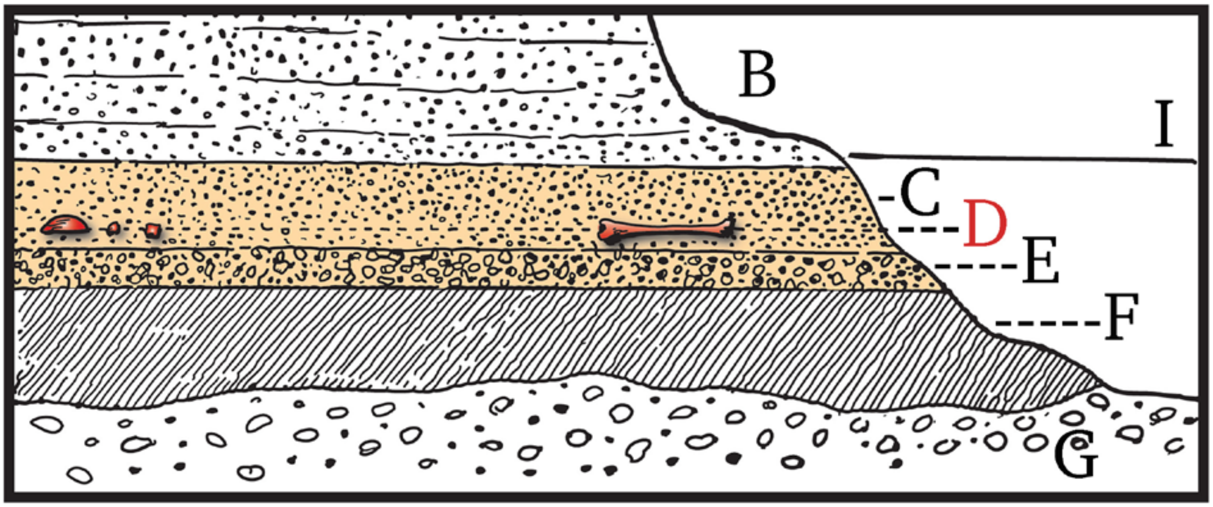


[Detail of the lower portion of Figure 2a, main text. The Top of the "C -- Bed of lapilli rock" (LB) is shown just below the "I - Dry seasonal level of river" (LWL), and the Skullcap/Femur I level (D) is placed in the lower portion of the LB, representations that conflict with the discovery of the LB in outcrop above the river level.]

These comprised the bones of a species of *Cervus* [***Axis lydekkeri***] which still survives in the Malayan Isles. Many remains of ***Stegodon*** [***trigoncephalus***], besides some of a species of ***Bubalus*** [***palaesondaicus***] apparently identical with the Siwalik species. At the same time a *Boselephas* [***Duboisia santeng***] differing from the known species, living or fossil, was discovered. In addition the genera rhinoceros, Sus, felis, hyena, and others were represented, whilst a gavial and crocodile were found which differ somewhat from the existing species in India, were unearthed. [See Table 1, main text.]

The setting in of the rainy season prevented the continuance of the excavations until next May, when work was resumed, with the result that in August 1892, the left femur [**Femur I**] was discovered in precisely the same layer at a distance of 15 metres from the spot where the Skullcap and molar tooth were found in the previous year.

In the October following a second molar tooth was discovered [**1892 Molar**]. This lay 3 meters from the place where the skullcap [**Skullcap**] was picked up, in a direction towards the spot where the femur was found. This tooth was not described in Dr. Dubois' first monograph, as it was only subsequently met with among a number of teeth taken from the same place. ….

These details as to the geological conditions under which the four bones were discovered were advanced to prove that they not only came from the same bed, most probably late Pliocene, but that they were all found at precisely the same level in it, hence most probably they must be of equal age. The fact that their contours are sharp is opposed to the view that they have been washed out of an older (later ?) bed. Dr. Dubois therefore thought that there was little reason to doubt their origin from the rocky strata.

All the bones were in the same state of fossilization, harder than marble, very heavy, and of a chocolate-brown color. The femur was twice the weight of a recent human femur of the same dimensions. The erosion on the surface of the skull-cap was not the result of disease as some critics had suggested, but was due to the corroding influence of the water, which had similarly affected all the bones found in this locality.

In discussing the question as to whether the bones belonged to the same individual or not, Dr. Dubois stated that in his opinion the distances apart at which they were found was no argument against their common origin. In his experience he had frequently found portions of the same skeleton as widely separated and, in some instances, fragments of the same bone were met with as far apart. Nor had he ever found anything like a complete skeleton, which seemed to prove that the bones must have been widely scattered. Dr. Dubois advanced the theory that in all probability the remains were those of animals which had perished in volcanic catastrophies, their corpses being washed down in the course of a large Pliocene river.

Most probably the bones, before they were deposited and buried in the old alluvia, were separated from one another by the rotting of the flesh or it may be together with the flesh were torn apart by crocodiles. In support of the latter view Dr. Dubois noted the fact that many remains of crocodiles were found as well as the marks of their teeth in the spongy parts of many fossil bones.

Considering the few examples there are of the fossil remains of anthropoid apes, and the absence of human remains from strata older than the middle Pleistocene, the lecturer urged that it was almost impossible 'to conceive that the thigh bone now discovered, which most certainly belonged to the Tertiary Period, could be human, whilst the skull found alongside of it should be regarded as that of a great anthropoid ape, a supposition which would necessarily involve their separate origin; much more reasonable to him appeared the view that they belonged to one and the same intermediate form.

**<u>S II-F4</u>**

[Dubois (1896e), translated from German. December 14, 1895, oral presentation to the Berlin Society for Anthropology. Primarily translated from reprint of Anatomischer Anzeiger. Centralblatt für die gesamte wischenschaftliche Anatomie. Amtliches Organ der Anatomischen Gesellshaft. Herausgegeben von Prof. K. von Bardeleben in Jena–Verslag von Gustav Fisher in Jena XII. Band, Nr. 1, 1896 [Anatomical Journal. Central publication for all scientific anatomy. Professional Journal of the Anatomical Society. Edited by prof. K von Bardeleben in Jena-Published by Gustav Fisher ….; also, Dubois 1895c, 1896a, g]

[3] At the discovery site of Trinil, the beds, which throughout consist of volcanic tuff, are exposed in the rocky slopes on the banks of a relatively large river, the Bengawan or Solo. They consist here primarily of (Fig. 1) not very consolidated sandstone, which becomes coarser towards the bottom near the level of the dry season watermark of the river, with increasing amounts of lapilli occurring in the composition. The bones are found throughout the entire sandstone section [3>>4], but are more numerous in the lower half and most numerous in a lapilli bed of about 1-meter thickness. I found very few within the conglomerate bed below it and none in the indurated clay bed.

The four skeletal remains of *Pithecanthropus* were found in different years, since each rainy season marked the end of the excavations, due to swelling of the river. These could then only be resumed in the following dry season. Also one was uncovered later than the other within the same working period, when the rocks had to be carefully removed, layer by layer and divided into surface blocks.

The four remains, however, were found at exactly the same level within the fully developed lapilli bed (Fig. 1). They have thus been deposited at the same time, i.e. they are of the same age. The molars were located within 1 to at most 3 meters distance from the skullcap; the femur at a distance of 15 meters. The very sharp contours of their outer surfaces does not fit the theory that they have been reworked from an older bed and then were re-deposited a second time. They are within the original deposits. Also, all of them show the exact same state of preservation and petrification, as is the case with all other bones that are derived from this specific bed at Trinil {FN: Also the color of the femur [**Femur I**] is exactly the same chocolate brown as that of the calvaria; the latter only appears somewhat different, since the calvaria was treated with varnish etc. before a mold was made} The specific gravity (s.g of the compacted (?) = 2.456) is much greater than that of non-fossilized bones (s.g. compacted (?) = 1.930). The femur [**Femur I**] weighs 1 kilogram, thus significantly more than a recent human femur of the same dimensions.

The marrow canal has been partly filled with a stony mass. The eroded surface that is shown on the skull cap, but by contrast is not present on the femur, is a result of a process that took place at the site of deposition. This is demonstrated by [the ? similar degree of corrosion of] many bones recovered from the vicinity of the Skullcap, and [is a *in situ* process] which is related to the acidic water that percolates throughout the rocks.

[And then quoting a translation of "*Verhandlungen der Berliner Gesellschaft für Anthropologie, Ethnologie und Urgeschichte"* {Proceedings of the Berlin Society for Anthropology, Ethnology and Early History} Redigirt von Rud. Virchow {Edited by Rudi Virchow} Aussertordentliche Sitzung vom 14. Dezember 1895 {Special Session of December 14, 1895} Zeitschrift fuer Ethnologie 27: 723-738,749. Verslag von A. Asher & Co. 1895. [Dubois 1895c]

[725] Together with these finds, the most numerous skeletal remains found were those of a small *Axis*-like deer [***Axis lydekkeri***], namely hundreds of complete antler beams and fragments, as well as numerous ***Stegodon*** [***trigonocephalus***] remains. Furthermore *Bubalus* [***palaeokerabau***] was found, probably identical to the Siwalik species. Also, the genus *Leptobos* [there are no Epileptobos fossils in the Dubois Collection] indicates [725>>726] the same relationships to the Neo Tertiary and Pleistocene faunas of the near Indies. Also, there is a *Bosephalus* (*Portax*) [***Duboisia santeng***] which differs from the extant one as well as from the fossil Narbada species. Additional genera were represented of ***Rhinoceros*** [***sondaicus***], ***Sus*** [***brachygnathus***], *Hyaena* [there are no Hyaena fossils in the Trinil material of the Dubois Collection], and *Felis* [***Panthera tigris***], apparently all new species.

Among the species from the same beds, found at different locations, a giant scaly animal (*Pholidota?*)

(*Manis*) occurs, which exceeds the size of the extant species threefold, as well as a *Hippopotamus*, belonging to the same sub-genus, *Hexaprotodon* [there are no hippopotamus in the Trinil material of the Dubois Collection today], as the species from the Siwalik beds in the near Indies.

[Fig. 1 Cross section of the bone bearing beds at Trinil
A. cultivated land. B. soft sandstone. C. lapilli bed. D. level at which the skeletal remains were found. E. conglomerate. F. indurated clay. G. marine breccias. H. Rainy season water mark of the river. I. Dry season water mark of the river.]

Figur 1.

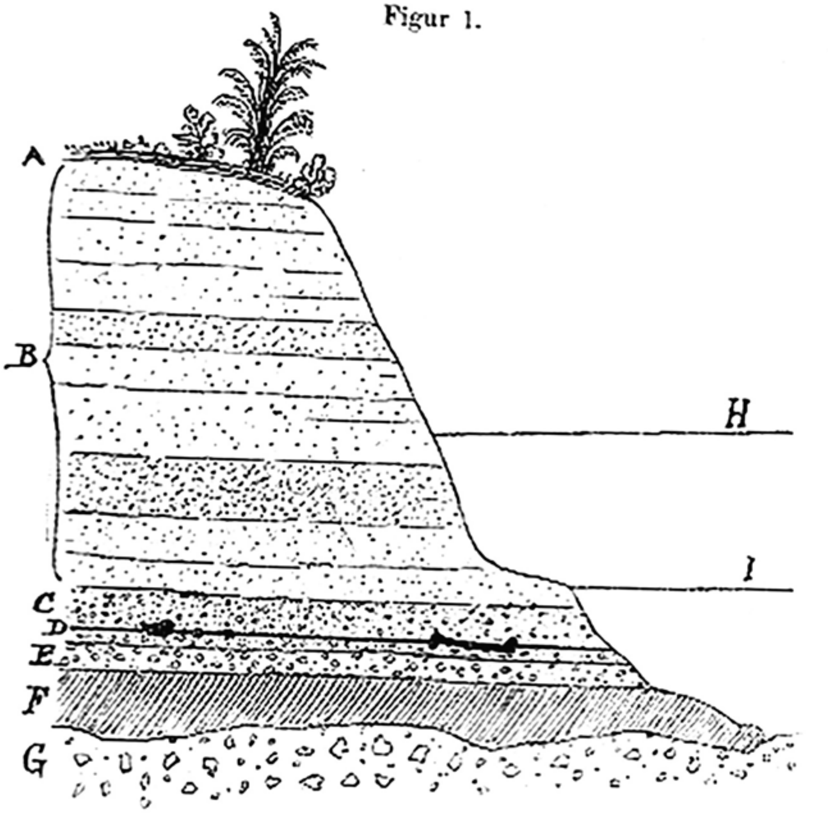


Durchschnitt der knochenführenden Schichten zu Trinil.
*A.* Culturboden. *B.* Weicher Sandstein. *C.* Lapilli-Schicht.
*D.* Niveau, in welchem die vier Skeletreste gefunden sind.
*E.* Conglomerat. *F.* Thonstein. *G.* Marine Breccie.
*H.* Regenzeitpegel des Flusses. *I.* Trockenzeitpegel des Flusses.

**S II-F5**

[Dubois (1896f), Translated from French. Bulletin of the Society of Anthropology of Paris. 645th Session, 24 June 1896, Chaired by Mr. Hervé, Fourteenth Annual Transformist Conference *Pithecanthropus erectus* and the origin of Man, By Mesrs Dr. Eug, Dubois and Dr. L. Manouvrier.]

[page 463] See below the geologic cross section of the [sedimentary] rocks at Trinil (previously published in: *Royal Dublin Society* Fig. 1).

The *P.e.* femur was found at a distance of 15 meters from the cranial skullcap. Our mind would more easily understand the dimensional relationships if they are smaller rather than larger and it seems useful to me to consider them at a reduced scale.

Let us assume for the extent of the terrain of 100,000 meters: 10 meters, [then] we get for the distance between the two main bones of our *Pithecanthropus* 1½ millimeter. If you keep in mind this [463>>464] account of the distance and if one does not forget the fact that among the immense numbers of bones that I have collected in the field, I have never found bones, anthropoid as well as human ones, that at Trinil are such a small distance apart and in addition exactly in the same horizon of the [sedimentary] layers; that is to say deposited at the same time !

I even believe that the probability that exists here – that the bones are derived from the same individual – is large enough that we may view it practically as a certainty.

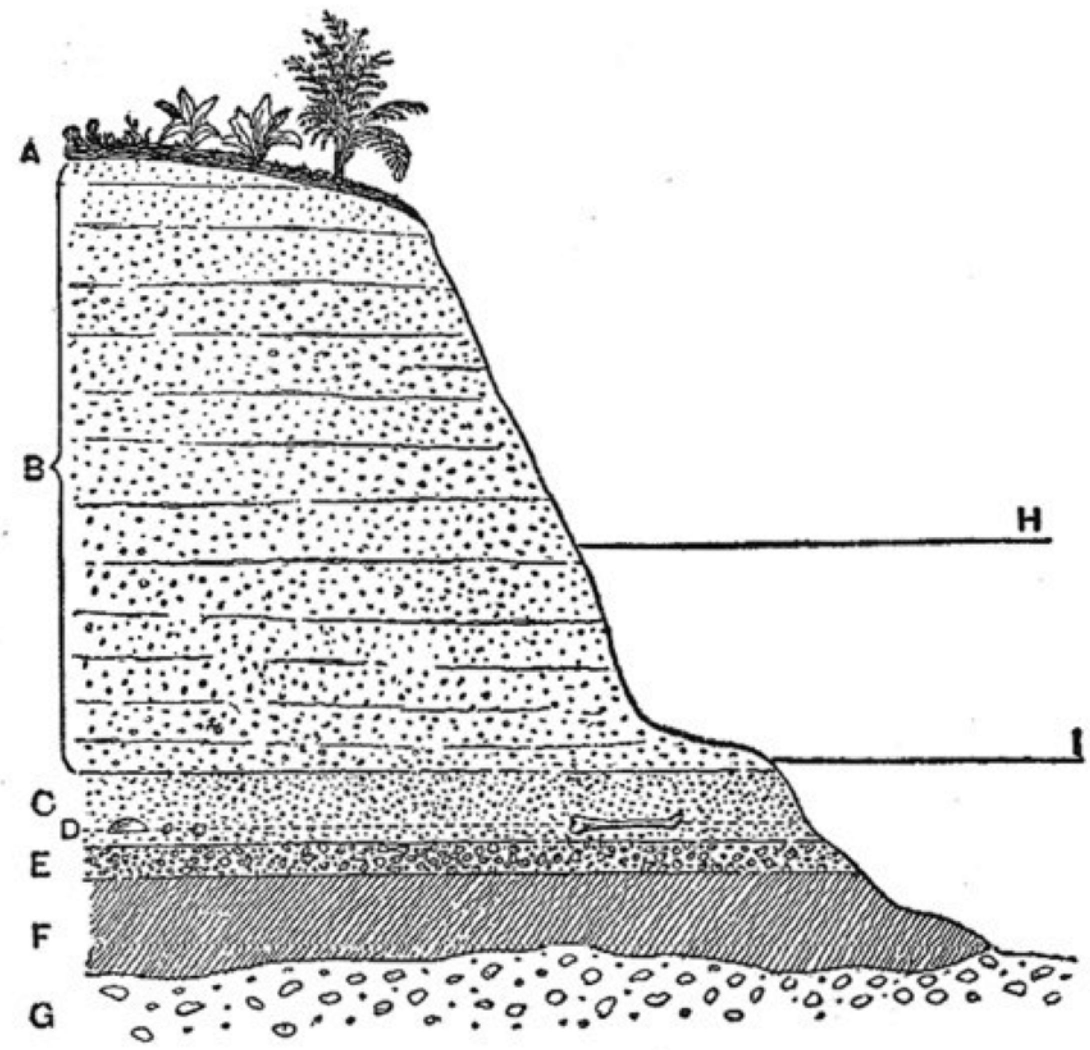


Fig. 1. — Coupe de Trinil.

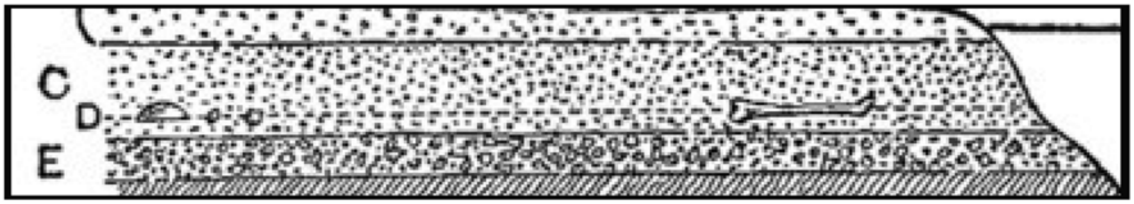


A. Vegetable soil [not shown]. B. Sands [not shown]. C. Lapilli. D. Bed of bones. E. Conglomerate. F. Clay. G. Marine breccia. II. Level of the river (rainy season) [not shown]. I. Level of the river (dry season).

**S II-F6**

[Dubois (1907: 452) translated from Dutch. Communications of the Journal of the Royal Netherlands Geographic Society. From "Some results from the Netherlands regarding the knowledge of the Kendeng (Trinil) fauna."]

Due to weathering and washing out of the loose sandstone, the bones appear at the surface and mix with the black soil, likewise a product of erosion. Very many bones can thus readily be picked up off the ground. Whereas the tuffs contain many limestone concretions, these occur not only as grayish-white irregular nodules (*kankar* of the geologists in India) within the black soil, but also usually coat the bones, as can be observed frequently and especially near Bangle, Kali Gede (between Trinil and Ngawi) and numerous other places. ….

The huge majority of animals that perished [including those recovered as fossils in the LB] apparently died during volcanic eruptions. It is a well-known fact that during the initial violent stages mostly coarse material is produced that becomes finer and finer as the eruption proceeds. This corresponds indeed with the fact that now most fossil bones are found in the coarser (lapilli) tuffs. In places such as at Trinil, where considerable accumulation of bones has taken place, it may be assumed that these animal cadavers were washed together in a bend of the old river. Within a surface area of only a few numbers of Ares [Note: 1 Are = .024710 acres], horns [antlers] of hundreds of deer have been excavated and train wagon loads full of other bones. The [this] most productive of all discovery sites was found in 1891, in the form of two rocky sandstone ledges, which were exposed during the dry season along both shores of the Bengawan and covered by a few meters of water during the wet season. However, small dams were dug around them and subsequently also the 10-12 m high, steep shore banks were excavated from the top downward to a few meters below the lowest level of the river.

**S II-F7**

[Dubois (1908: 1235 -1270), translated from German. Journal of the Royal Netherlands Geographic Society. From "The geologic age of the Kendeng- or Trinil-fauna."]

In the last few years an opinion has increasingly emerged to consider a relatively young age for *Pithecanthropus erectus* and its corresponding fauna, as well as the beds which have yielded the fossil remains of the Trinil fauna on Java, that is, at the oldest, an early Pleistocene age.

Some have been led to this opinion through non-geologic considerations, namely from certain ideas about the origin of man, others drew support from geological – paleontological facts.

I will subject to a discussion here only those facts that fall within the scope of this journal, mainly the actual paleontological ones, since with respect to determining the question of geologic age, guidance from paleontology has always proved to be most reliable.

W. Volz made an attempt in previous years to investigate more closely the question of age of the Trinil layers from a purely geologic point of view, occasioned by a short visit to the area of interest [1*) The geologic age of the Pithecanthropus beds near Trinil, eastern Java. Neues Jahrbuch fuer Mineralogie, Geologie and Paleontologie. Hardback 1907, pp. 256-271. Also: "Globus", Vol. 92, Nr. 22]. According to his view, the, as he believes, irregularly deposited, generally non-bedded tuffs, which as muddy streams would have embedded the bones, could only be derived from the volcano Lawu-Kukusan, during the main activity of this volcano which had started in the early Pleistocene and which has presently come to an almost complete halt. This mountain, in fact, belongs to a class of younger pyroxene- andesite volcanoes of Java and shows its young age in that the typical volcanic shape has not been lost through denudation. Based on these concepts, Volz concludes that the depositional beds in question are at any rate not older than early Pleistocene and that they probably can be placed in the middle Pleistocene.

I consider the basis for this view to be quite inadequate, but I can refrain from an exhaustive argument against it since the paleontological facts clearly lead to a contrary result. I will only describe here the following.

Only the Wilis and the Lawu-Kukusan [volcanic centers] are candidates for the production of the bone tuff. This can be easily concluded from the distribution of the tuff, as is shown on the attached sketch (Plate XXXIX, Fig. A [below]), and from the fact that they (the bone tuffs) have the same petrologic composition as the rocks in these volcanoes. Because of the identical composition of these two, as is the case with most of the volcanic mountains in Java (the younger ones all belong to the pyroxene-andesite types), it is impossible to determine petrologically which one of them has exclusively delivered the bone tuff material or has been the principal source.

The distribution of the tuff[aceous formation of the Kendeng Hills] and its massive thickness in the west, suggests that the Wilis is the principal source [1*) My own research led to this conclusion already in 1892 [Dubois 1892d] (Verslag van het Mijnwezen, tweede kwartaal, Batavia 1892, p. 17). I had then well abandoned my initial opinion of 1890 to view the Pandan more so than the Lawu and the Wilis as producer of the ash (Volz, e.g. p.265)], although the Lawu could also have contributed a considerable quantity [of volcaniclastic sediment]. Considering also that both volcanoes came to rest only at a late stage, nothing can be argued against the possibility that they had started their activity already in the Pliocene, because the petrographic analyses of similarly looking tuffs which are paleontologically distinct, does not indicate the opposite.

According to Verbeek, eruptions of pyroxene-andesite lava did take place early on in the Miocene and continued later in that period and probably lasted into the Pliocene [1*) Verbeek and Fennema, Geologic description of Java and Madura, Amsterdam, 1896, Part I, pp. 39 and 40]. The considerable Quaternary deposits, which according to Volz must be present in the vicinity of this huge, partly still active volcano, are in fact present. These are the beds (q on the sketch map) of the great plains on which the volcanoes emerged and which Verbeek also designated as Quaternary.

However, where the flat Quaternary beds thin and disappear to the north, and where a bit further north the chained hills of the Kendeng emerge, the older (bone bearing) beds are exposed. These have been primarily uplifted from their original horizontal attitude and where they are developed thickest, such as in the vicinity of Pandan, the dip angle increases gradually to the north, from 1.5 degrees near Notopuro to 8 degrees near Kedungbrubus (See Fig. A [provided below]) and even 15 degrees near Terbalan. On the plains, the bone bearing beds are probably continued below the Quaternary cover at considerable depth.

One of the geologic facts which most strongly supports an age older than Quaternary for the bone bearing tuff, is precisely this uplift of the formations which has led to their considerable height above the plains. West of Pandan and 4 km in a northeasterly direction from Kedungbrubus, they reach a height of almost 400 m. above sea-level. The volcanic tuffs from which a small hill is formed near Butak (see the sketch map [provided below]), are also clearly bedded, and show, at the same time, cross-bedding of river deposits, and contain not only bones of land vertebrates of the Kendeng fauna, but also the same fresh water crocodiles and shells of freshwater mollusks as found in all other bone tuffs. These same bone-bearing tuffs also occur at some 300 m. above sea level east of Pandan (between Djeruk and Tritik).

Related to the fact that the tuffs are indeed bedded everywhere and exhibit the full character of fluvial deposits, contrary to the opinion expressed by Volz, is also the great distance which they could overcome, away from the volcanic craters. This explains why bone tuffs with considerable thickness and extent occur near Tinggang (see sketch map [provided below]) and even further north of the Kendeng hills (up to a distance of 60 km from the Lawu crater).

When in 1890, I recognized that volcanic tuffs formed the layers which enclose the bones [1*) Initially not far from the abandoned village Kedinglumbu, from where most of the bones collected by Raden Saléh originated [e.g., Saléh 1867]. My first find in May 1890, was the upper jaw of a hippopotamus (Hexaprotodon), which I extracted from the tuffs near Kedunglumbu], that the fauna by its character was younger than the Siwalik fauna, and that the bones nevertheless occur at considerable height above the plains, I originally imagined that the tuffs had formed in Pleistocene times from mud flows. Further studies and investigations of the layers, however, soon taught me that fluvial bedding of these tuffs is a more likely explanation. They are found along almost the entire length of the Kendeng hills, over a distance of more than 100 km in an east-west direction. Only east of Ngawi, a larger interruption is found and bone discovery sites form an island in the middle of the foraminiferous marl, which usually underlies the tuffs. As an aside, the build-up of the beds west of Ngawi is much less than it is east of there, on both sides of (the) Pandan. Only the coincidence that the western extension of the beds has been incised by the Bengawan led to the discovery of the Trinil location. Furthermore, the thick bone-bearing tuffs are also found north of the Kendeng hills, on the east side of the Bengawan, near Ngluwak and in the vicinity of Tinggang, along the west side of the river, and further north in the neighborhood of Tjepu.

Throughout the entire extent of the tuffs, remains of fresh water mollusks, fresh water inhabiting crocodiles and turtles (*Garialis* [***Gavialis bengawanensis***] and a *Crocodylus palustris* [*siamensis*], closely resembling the established species *Hardella* and *Trinonyx* [**Testudinoidea**]) are found, and near [at] Trinil at least 7 fresh water fish species (***Anabis microcephalus***, ***Clarias magur*** and also 4 other Silurids as well as several **Ophiocephalids** species), which all belong to one and the same fauna. In general, the tuffs show clear bedding and in very many places also cross bedding typical of fluvial deposits. It must be emphasized here again strongly: there is no doubt about the fluvial nature of the entire formation. From the nature of the cross bedding, I concluded already in 1894, that the direction of flow of the ancient river was west to east in the entire Kendeng area as well as north of the chain of hills (in the vicinity of Tinggang). To a large extent, only near Ngawi a general direction to the northwest could be derived. I concluded then that already in these distant times, the drainage of the plains between the Wilis and the Lawu was accomplished by a river much like the present day Madiun [2*) Verslag van het Mijnwezen, 1894, 2nd Quarter, p. 14 {Dubois 1894d}].

The west to east flowing river had also several branches, one of which ran along the north side of the Kendeng hills, or it temporarily changed its course to there; otherwise one cannot explain the thick tuff deposits near Ngluwak and Tinggang . At any rate, the Kendeng hills were not present in their current form as proved by the elevated tuff layers. The break-through valley of the Bengawan near Ngawi can be explained by the theory of Powell-Tietze of gradual emergence of the Kendeng Hill chain while the river maintained its deepest valley point at the same level.

Further, it must be emphasized that the bones are found in their original layers, and in no way could have washed out of older beds, as Martin [1*) K. Martin: The age of the beds of Sondee and Trinil on Java. Transactions Royal Academy for the Sciences, Amsterdam, 1908, pp. 7-16], among others, has held for possible. It is true that in the Kendeng Hills the bones are often completely or partially exposed due to the rain, which can wash away the surrounding weaker sandstone [2*) In some places the tuffs are completely washed away and the bones lie directly on the Foraminifera marls; adjacent to that then, the tuff bed commences again and gradually increases in thickness]. Those bones show then, however, clearly the effects of erosion, whereas those that are still encased in the formations, as is the case near Trinil, the original, most delicate, surface pattern has been retained.

It is true that many of the bones that have been dug up near Trinil (among them the skull cap of the *Pithecanthropus* at its outer surface) have suffered from corrosion caused by groundwater that contains sulfuric acid. The tuffs in this location happen to contain many sulfur compounds; pyrite has been deposited on the bones and lignites, and the water that springs free during the digging had a noticeable acidic reaction, was milky in appearance caused by sulfur precipitation and smelled like hydrogen sulfide. Fortunately most of the Trinil bones have not suffered from corrosion, the effect of which on the bones, it should be noted, is completely different than that of transportation in a river bed. One cannot imagine that delicate bone platelets and skull fragments, the long and thin deer antlers still attached to the skull, the entire jaw of ***Garialis*** with its numerous thin teeth that were in extremely fragile fossil condition, had undergone a small amount of transportation or that they were washed out locally from older beds, and then still had been remained undamaged.

The facts force us to accept that the bones are still at the very same place where they were deposited in fresh condition, before their fossilization and that they are located in their primary resting place.

These bone beds have in common with all other fluvial deposits that very rarely all bones of an animal are found together and also the fact that most of the bones have been fragmented, as well as the great enrichment in some and the relative paucity in other locations in addition to the fact that in some places species are missing which are abundant elsewhere, all this can be explained very naturally. I imagine the events to have taken place about in the following manner {1*) See my Note in the "Jaarboek van het Mijnwezen", 1892, 2nd Quarter, pp.16-17 [Dubois 1892d]}:

From the nature of the bedding, it is logical that the animals perished during volcanic eruptions in a manner similar to, but even more violently than, they took place often during ancient times in Java. The eruptions, to which we indirectly can attribute the fossil bones, must at times have repeated themselves again and again, even if they all occurred in the same geologic period. Most of the bones in the richest discovery places such as near Trinil, Kedungbrubus and Bangle, occur in a lapilli bed, which rests upon a gray-black calcium-rich and, when dry, very crumbly claystone, containing freshwater mollusks (mostly ***Melania*** and ***Unio*** [gastropods and mussels]). One such clay layer, which must have been deposited in very quietly flowing water, reaches a thickness of about 35 meters near Kedungbrubus, while near Trinil it is on average only 1 meter thick. The lapilli bed signifies then, the beginning of the volcanic break-through and animal carcasses must have been washed together exactly at those quiet places (where under normal circumstances clay was deposited). The main bone bed near Trinil was deposited in this manner.

During times of diminishing volcanic activity, or after most animals had perished, the animal remains in the tuffs at those times must have become less abundant. This explains the relative paucity of remains in the other beds of Trinil. The cadavers in those quiet places fell apart to some extent through decay, but much more likely they must have been ripped to pieces by crocodiles, which often live in such places, preying on dead and living animals. The bones were then separated from each other and were broken many times. By strongly slapping the prey back and forth and thereby causing it to hit the ground, those fearsome animals of prey try to partly crush them and rip them apart with the help of their front paws. Animals as large as humans, together with bones, could then be swallowed. It is well known that crocodiles have an extremely powerful bite. This allows them, as has been determined experimentally, to swallow almost three times their body weight, which could amount to thousands of kilograms. Such power is sufficient to cause the breaks in the bones that have been found near Trinil and in other discovery places.

In many cases I noticed tooth marks of these reptiles on the less fossilized bone fragments. Colossal **crocodile** teeth are, by the way, the most common fossils of Trinil.

The fact that hundreds of antlers of the same deer species (*Cervus liriocerus* [*Axis lydekkeri*]) have been found near Trinil, is simply explained by the simultaneous extermination of the entire herd of these *Axis*-like deer [1* It is well known that the currently living *Axis* species also builds large herds], and were swept together at these tranquil places. Precisely these antlers, which are not covered by flesh, are left mostly untouched by the crocodiles, while the other bones were almost always broken, as they are in fact encountered.

It must be pointed out here in particular, that many shed antlers were also found. This can be explained by the fact that these quiet locations in the river were also used as watering places. Such shed deer antlers are also common in European clay deposits.

An important fact of a different nature must not remain unmentioned. While remains of *Elephas* and *Hippopotamus* in general neither belong to the rarest nor exactly to the most frequent fossils in the Kendeng beds, both these species are curiously enough missing in my extensive collections of Trinil. From this, one should not conclude, however, that there is a difference in age of these beds, because in the closest vicinity of Trinil (near Nongko and at other places) remains of both these species are not rare. I believe that the paucity of certain species is related to the manner in which the animals lived together. Hippos were relatively common in their waterholes of the plains before the eruptions of the distant volcanoes. The elephants may have lived nearby and from there, their carcasses would have easily been washed to Trinil during a volcanic break-out, whereas Stegodonts, whose remains are abundant, were already present in the collection area.

Many other circumstances can explain the abundance or sparseness of the various mammal species, depending on the discovery location of their remains. Similar to the inability to decide on a difference in climate from a lack of plant fossils in one spot and an abundance in another, one cannot conclude from the above facts that there is a difference in fauna.

The preceding discussion from which one can imagine a probable scenario for the build up of the bone tuffs of the Kendeng or Trinil beds, based on the available facts, appeared to be necessary in a discussion of the problem of the age of these beds, since erroneous ideas thereof from various sources could be used and lead to a wrong conclusion regarding this age. Also, questions often asked, such as: Why has only one *Pithecanthropus* been found? Why could his bones be separated so much? These are naturally answered when one realizes to some degree what happened in this distant time.

Let us now direct our attention to paleontology to judge the question of age. It should be stated first of all, that all discovery locations of the entire Kendeng are entirely of the same age, and that the vertebrate fauna of the bone-bearing tuffs is a very uniform unit. It is true that, as we have seen, in some places, certain species are missing which are relatively abundant elsewhere, or conversely there may be a species present which otherwise is nowhere to be found; these differences can however, as already discussed for Trinil, be viewed as facies variations caused by the locally differing circumstances during deposition. Most of the species are common in all finding places, a considerable number of strongly characteristic species can readily be considered as “diagnostic fossils”.  These include the very typical species *Cervus liriocerus*, *Bubalus palaeokerabau*, *Leptobus Groeneveldtii*, *Tetracerus Kroesenii*, *Rhinocerus sivasondaicus*, *Stegodon javanoganesa*, *Garialis bengawanicus*, *Crocodilus ossifragus*, *Hardella isoclina* and several others [see below, and Table 3, main text]. Also, those species that are missing in some places, which are common elsewhere, do occur again in the most immediate vicinity.

What is it now that the paleontology teaches us about the age of these bone-bearing tuffs?

The underlying beds of course determine their maximum age. These are mostly composed of marine foraminifera marls, locally also of carbonates, marine breccias and very locally, probably representing the very youngest bed, the sandy clay- marl of Sonde and Padasmalang [1-3 km west of Trinil]. All these marine beds on which the tuffs are discordantly deposited belong to Stage M2 of Verbeek, which he describes, based on the most common rocks, as the marl stage, and to the Java stage of Martin. Their age can go back as far as the Oligocene.

Of these underlying beds, the only one that requires further discussion is the age of the clayey marls of Sonde (and Palismalang), based on Martin’s careful research on the well-preserved mollusk fauna. According to Martin’s most recent determinations, this fauna contains 54 % of species that are still alive today; he concludes therefrom that the marine sediments of Sonde cannot be older than Pliocene {1*) Loco citato, p.11 Prof. Martin has expressed himself previously repeatedly in similar terms, most recently in the Journal of the German Geological Society, 1900, in which he indicated an almost identical percentage, namely 53}.

I have expressed my views about this conclusion previously in this magazine {2*) Second series, part 24 (1907), p.456, Note}, in the following manner: “The underlying beds of Sonde, deposited in marine waters cannot tell us anything definitive about the age of the bone-bearing beds. If the Sonde deposits indeed contain about 53% of still living species, as is assumed by Prof. Martin, then these deposits of the Indies must be regarded as Miocene, if one considers that a careful investigation of such older marine Pliocene deposits in England showed they contain exactly such a percentage of still living species.” Martin remarks on this that this standard is inadmissible, because in the determination of the stratigraphic level, one must first of all consider the relationship of the beds of Sonde with the other sediments in Java and neighboring areas {1*) Loco citato, p. 11}.

This last statement is very true, but unfortunately the relationship of the Sondee deposits with the other sediments of Java as well as with the deposits in Burma, researched by Noetling {2*) Fauna of the Miocene beds of Burma. Palaeontologia Indica, New Series, Vol. I. 3. Calcutta 1901}, gives us only a relative age of these beds with respect to each other, but not their absolute geologic age. It has so far remained true that the identifications “Eocene, Miocene and Pliocene” which were used in this context, refer solely to the deposits in the tropics, without thereby intending to indicate any equivalence with European Tertiary strata {‘3*) K. Martin. The Tertiary deposits on Java. Leiden 1880. General Section, p. 25}. This fact has been forgotten over time; that’s why it seems to me that the cited remark is fitting. Till now, Verbeek’s research on the Orbitoids of Java has taught us the first absolute age boundary in the Java Tertiary beds, namely that between the Eocene and Oligocene. I believe that the bone tuffs of the Kendeng represent a second section of the Tertiary that can be correlated to beds outside of the tropics.

As is well known, Deshayes has used the jumps in percentage of still living species contained in the European Tertiary deposits as the basis for their age determination. By comparison of 3,000 Tertiary and 5,000 recent species, he concluded that the still living species become more numerous as the bed becomes younger and takes up a higher position and vice versa.

He had, however, also clearly recognized that this steady change in fauna can be connected with a steadily lowering temperature during the Tertiary period in Europe, which has decreased from an equatorial–like condition to one of about that of the current European climate {4*) G.P. Deshayes, Description of fossil shells from the area around Paris. Mollusks. Second Volume. Paris 1824, p. 775 and on, and: Description of characteristic field fossils. Paris 1831}. Lyell [1830–1833] based his tripartite division of the European Tertiary in Eocene, Miocene and Pliocene on the numbers of Deshayes, and designated 35 – 50% of recent species for the Pliocene, 17 – 25% for the Miocene and 3.5 – 5% for the Eocene {5*) Principles of Geology, First edition, Vol. III, Appendix 1833}.

These percentages can no longer be accepted. According to our current knowledge, up to 90 % of recent species could be attributed to the Pliocene. In the series of beds of the English Pliocene the percentage from the oldest to the youngest division gradually rises from about 60 to 90. The faunas of these layered complexes can be compared very precisely with the well known faunas from the relevant seas studied by various researchers (Searles Wood, S.P. Woodward, C. Reid, Harmer and others); one may therefore assume that the computed numbers deviate very little from the actual ones.

In consideration of the unmistakable fact that in Europe the changing of the faunas was not caused by the gradual transition of species as the principal cause, but the lowering of temperature during the entire Tertiary period, and especially strongly so near the end and thereafter, that in the tropical seas, on the other hand, this prevailing factor was not or hardly in force, it must be judged that a percentage of 53 for recent species for the Pliocene of Java is much too low. Martin himself says "that with relatively contemporaneous beds of the Indies on the one hand and Europe's on the other hand, we encounter a much higher percentage of recent species in the Tertiary of the Indies as compared to that of Europe and the difference should even increase as the contemporaneously deposited beds in both places get younger" 1*). {FN: The Tertiary beds of Java. 1880. General Section p. 24}.

The actual percentage of recent species in the Tertiary beds of Java must in fact be significantly higher than computed, because a large number of species that have been described as new, were later in fact discovered to be still living. For this reason Martin thought it justified to assume at least 50% instead of the computed number of 35 % for the Miocene of Java (later corrected to 30%).

If the number computed for the Pliocene must now be proportionally increased even more so in order to attain the proper percentage figure, what value remains then in this computed amount! However, we don't know that the correction to be applied is large; in the past, it may well have been significantly overestimated by Martin. In a very young fauna from Celebes, in which one might expect nearly 100%, Schepman determined that 80% is present as recent species {2*) Martin, The age of the Sondee beds, p. 10 [Martin]}.

This provides also, according to Martin, a handle on the judgement of the size of the correction. Consequently, the computed percentage number for Sondee may not be much below the actual number so that this fauna can be attributed the Pliocene. In addition, we may accept as fact that the changes in the marine faunas in the tropics must have taken place imperceptibly slow towards the end of the Tertiary in comparison with those caused by the strong climatic changes in Europe.

For these reasons the underlying beds cannot yield any certainty about the age of the bone bearing tuffs, although perhaps the most reasonable assumption is that the Sondee bed belongs to the Miocene, which in Europe can contain up to 40% of living species.

From the bone bed in Trinil itself 7 freshwater gastropods, of which 5 *Melania* species, one *Paludina* and one *Ampullaria,* were named by Martin, as well as a land snail, a *Bulimus* {1*) In W. Branca, Preliminary announcement of the results of the Trinil Expedition of the Academic Jubilee Foundation of the City of Berlin. Royal Bavarian Academy of Sciences and: K. Martin, The age of the Sondee and Trinil beds on Java: Report of the Meeting of the Mathematics and Natural Sciences Department of the Royal Academy of Sciences, Amsterdam, dated 12 June 1908 [Martin 1908]]}. "All eight species are still living, only one variety differs a little bit from the recent representative". This common *Melania* variety in Trinil is likewise common in the Sonde bed. Two *Unio* species remain undetermined and if I understand it correctly also three gastropod species.

Two species of river mussels, one large the other small, however, belong to the most common mollusks of Trinil. While Trinil-type *Melania* [*Elongaria orientalis*] are very frequently found in the rivers and other fresh water bodies of Java, the large *Unio* [*Pseudodon vondembuschiamus trinilensis*] is remarkably missing. I have made a great effort therefore to look for it in various fresh-water, as well as brackish-water, bodies, but without success. It is thus not improbable that this represents an extinct species; I am therefore including an image of it below and tentatively name it *Unio trinilensis* [*Pseudodon vondembuschiamus trinilensis*] (Plate XXXIX, Fig. B [provided below]).

As a third species of lamellibranchiates, which I encountered at Trinil, I am introducing a *Corbicula* [mussel] which still must be identified, and which in other bone bed sites belongs to the most frequently occurring species; also two gastropods a *Limnaeus* and an *Isodoria* [*S I-U*].

In addition to the eight species named by Martin, there are therefore at least five others, increasing the total number to 13. Apart from the fact that the first mentioned number is small in an absolute sense, one is then not justified to conclude, with Martin, from the 8 determined species "that based on this, a post Tertiary age of the relevant layers can be concluded with absolute certainty", and " Therefore the main bone bed can only be a post Tertiary deposit" {1*) W. Branca, Preliminary announcement, p. 270 and: Martin, The age of the layers of Trinil and Sondee, p. 15; see Berkhout and Huffman 2021}. If among the 8 determined gastropods, upon further inspection, a single species proves yet to be different from recent species, which may still be somewhat possible (the opinions on species, variety and individual deviations are differing), then a percentage number below 88 can be computed for the still living species which have been identified. If the single *Unio* is indeed an extinct species, then this percentage would become even considerably smaller. And if among the minimum of 13 species, of which 5 are still undetermined, only one species were indeed extinct, then the percentage of about 92 for living species would be entirely sufficient to place the beds from which they have been derived in the late Pliocene.

In the youngest Pliocene deposits of England, the percentage number of still living species reaches 90%, and in the tropics we must find " a significantly higher percentage of recent species" for beds of the same age " and the difference becomes even larger as the deposits become younger".

It seems to me that the mollusks don't have the power of persuasion that Martin ascribes to them.

I can be brief about the disclosures of Messrs. Elbert and Carthaus {1*) FN: Dr. Joh. Elbert, The latest investigations on the question of Pithecanthropus. Natuurkundig Tijdschrift for Netherlands Indies. Issue 67, Weltevreden 1907, pp. 125-142; also: About the age of the Kendeng beds containing Pithecanthropus erectus Dubois. Neues Jahrbuch fuer Mineralogie, Geologie und Palaeontologie. Appendix Volume 25, pp. 648-662 (1908). Dr. E. Carthaus, The Selenka Expedition to Trinil. "Globus". Vol 93, Nr. 4 1908. From an announcement in a newspaper I understand that the latest discovery of Elbert in the Kendeng hills consisted of a fossilized banteng (species of wild ox) skull that had a bronze bullet imbedded. This bullet must have made an extreme chronological error in placing itself there.} The one by Elbert in which he divides the deposits in alignment with the European Pleistocene (Diluvium), I consider, foregoing a stronger expression, completely unfounded. The plant species, which by the way are very questionable, and which would point to a cooler climate, are not justifiable reasons for the assumption of an ice age on Java, because the remains may well have simply been washed down from higher mountainous zones, and thus do not prove a Pleistocene age. The plant remains which have been determined with certainty by Dr. Valeton of Buitenzorg, the outstanding expert of the flora of Java, do not lead to the conclusion of a cooler climate {2*) Branca, loco citato p. 271}. Besides laterites also real clayey deposits occur in the tropics, and the presence of the latter does then also not give proof of a colder climate. And finally concerning the human artifacts and the cooking hearth " consisting of three cooking pots, a Javanese kitchen (pavon) in its simplest form", which Elbert and Carthaus discovered in the river deposits of the Kendeng, I can assure the reader that during the entire length of my seven years of research on Sumatra and Java and especially during the excavations of Trinil, I always keenly looked for fossil human artifacts, without any success. Our brown and yellow brothers are, however, very innovative and also very skilled, if they know what we are looking for, to satisfy this wish and simultaneously hope to make a little money.

When, during my excavations near Trinil, and in order to promote a very thorough search of the excavated sand, I offered very modest prices for small teeth which looked like monkey or human teeth, within a few days, I collected from my prisoner-laborers several child or monkey teeth, which, as I could readily discern, were only "sub fossil", although they had gone to the trouble of making them look like real fossils, by rubbing them with ash and in one case by gluing on some Trinil sand [1*) Of the two teeth that Mrs. Selenka showed me as her most important finds, one looked exactly like one of those recent and whitish appearing human lower molar, with Trinil sand glued to its rootless bottom, although it "was not found at Trinil". The other one that indeed was found at Trinil was an upper pre-molar of a pig.].

At this point I must once again remind the reader that the bones from the Kendeng beds are all fossilized similar to Tertiary bones, for instance those of the Siwalik mountain chain.

Marsh and other paleontologists who inspected my collections immediately noticed this aspect of them. The bones have a very high specific gravity, organic material is only present in traces, and not only the amount of calcium carbonate but even more so of calcium phosphate has increased significantly. Also, the fluorine content conforms with that which is found by Carnot in Pliocene bones [2*) J.M. van Bemmelen in: Journal for Organic Chemistry. Hamburg and Leipzig, Vol. 15 (1897), pp 84 and following. ]. Lime spar and pyrite have been deposited in the marrow pipes and up into the smallest bone canals. The bones of the Narbada deposits appear completely different; they are much less fossilized and they stick to the tongue.

Dr. Carthaus mentions in his message to Branca {3*) Loco citato p. 271} [-*G4a*] "large pieces of wood, which are so well preserved, that based solely on that he could conclude that the deposit was really young." According to Branca, the few wood pieces that have been shipped to Berlin, show "that these are dark colored but little carbonized wood pieces, which remind him of the wood from the peat bogs." In contrast with that, I must note that the frequently found fossilized wood at Trinil [**wood**] has been altered into lignite, although not all to the same degree; this appears to depend on the type of wood. When freshly excavated and preserved wet, the wood often shows some flexibility, once it is dried it becomes brittle and looks in fact almost like black lignite. Taxodium and other Pliocene woods, which are found in Europe, are preserved in the same manner.

The remains of mammals (and in part the other vertebrates) are of much larger value for the age determination of the Kendeng beds than plant remains or even mollusk shells. First of all, the living mammal fauna is pretty well known, which can not at all be said of the mollusk fauna, and secondly our knowledge of each single mammal species is far more extensive which allows for a much more detailed comparison considering the complex construction of the mammal skeletons. In this context, the skull of a mammal species has the diagnostic value of many shells of mollusk species.

The mammal and other vertebrate animal faunas indicate that the Kendeng tuffs most certainly belong to the Upper Pliocene, although the Siwalik beds are considered to be not older than Pliocene.

This conclusion is justified in the following brief statements:

The main characteristics of the Pleistocene are the strong and periodic decreases in temperature, which in those times were felt over the entire earth. Since during later Tertiary time the gradual zonal differentiation of the climate had also particularly intensified, the climatic changes occurring during the end were more sudden events, which in middle and higher latitudes had the greatest influence on changes in the mammal world, because of the resulting expansion of the ice sheets in these already cold zones. Yet, the Pleistocene, Diluvial or Quarternary time represents, as the first name indeed signifies, as is well known, that slice of history of the world during which most of the species now living in Europe, were already developed. This was particularly true for the mammal and other vertebrate faunas.

In the equatorial regions, on the other hand, such climatic changes must have taken place almost without notice; the character of the mammal fauna cannot have changed noticeably as a result of it. Because, a lowering of temperature of 4 deg. C, as is assumed for the Pleistocene ice ages, in the mountainous areas corresponds to an increase in elevation of about 800 m.; and the spreading of the most common mammals in such warm areas is demonstrably fully independent from such an increase in elevation as far as it influences the temperature. This is true even for apes who especially love the warmth.

While in Europe and other areas at higher latitudes the Pleistocene fauna is composed then of a relatively large number of extinct mammals besides the still living species or temporarily assimilated species from colder climates, the Pleistocene climatic changes in the tropical regions of the earth therefore could have hardly changed the mammal world. In agreement herewith, the discovery by Lydekker {1*) Indian Tertiary and Post Tertiary Vertebrata. Vol. IV, p. 26. Calcutta 1886} of 32 with certainty determined mammal species in the south India karst holes (?) at latitude 15 deg. 25 min., included no fewer than 26 living species, only 6 (according to Lydekker 5) extinct species and 2 (according to Lydekker 3) species which should still be alive in Africa.

The fossil mammal fauna of the many caves that I investigated in Sumatra and Java, is composed exclusively of species which are still alive on those islands. There is no reason to assume that the bones which those animals left behind for us are all of a more recent time than those of the European fossil cave fauna; evidence for an older age is not only found in the condition of preservation and the geological conditions, but also the current different dissemination of several species e.g. the occurrence of the orangutan 400 to 500 km on the other side of his current range boundary in Sumatra, the Australian human races etc.

With regard to the much discussed, but relatively poorly known Narbada fauna, which entails a considerable number of extinct mammal species: it was assigned to the older Pleistocene purely for this reason, and that in the Near Indies it represents, as far as the specimens found to date, the next younger fauna to the Pliocene Sawalik. There is, however, no good reason, besides perhaps the degree of fossilization of the bones, to prevent us from assigning it to the late Pliocene.

Besides, it has too often been forgotten that the determination of several of these faunas represented by relatively little-known species, such as for instance *Stegodon ganesa*, is certainly not solid. The Narbada fauna can thus not be simply used as a basis for a judgement about the age of the Kendeng fauna, since the Narbada beds also have universally been declared as Quaternary by the geologists of India. The generic correspondence between the two faunas {1*) F. Frech in: Lethea geognostica. Part III, Vol. 2. Portion I: Flora and Fauna of the Quaternary, p. 31}, is not helpful as this also exists, and even to a stronger degree, between the Kendeng fauna and the younger portion of the Siwalik fauna; it yields a closer comparison of these two with even more intimate relationships of the species.

More than anything, the description of the mammal species, whose skeletal parts have been preserved in sufficient completeness to establish a true recognition of their relationships to known living and fossil species, may well lead to a correct judgment of the geologic age of the Kendeng fauna.

Among these faunas the *Stegodons* are of major interest to us, not only because the remains of these intermediate forms between Mastodon and Elephas belong to the most common ones, but primarily because the determination of species of these east Asian Proboscidians by itself, provides, up to a certain elevation, a conclusion about the geologic age.

Research into the very abundant *Stegodon* material from the entire Kendeng area, among which also several skulls of young and old animals, has led me to the conclusion that all these remains belong to a single species. It is distinguished from *Stegodon ganesa* by not much more than its significantly smaller size.

The skull possesses the same broad temporal groove, is just as characteristically strongly brachycephaloid and for the remainder both species look very similar. Only older skulls deviate a bit in some aspects from the typical Ganesa form, in that the frontal and occipital portions flatten towards each other and are more or less sharply separated. This does not need to prevent us, however, from ascribing these forms to the same species, because the skull form of the same species of elephant may, within certain limits, show considerable variation. In our species, the Ganesa type remains always recognizable.

Even the molars do not allow a specific distinction from *S. ganesa*, only the number of lamellas in the lower jaw reaches at least 14 instead of 13; the number of lamella has, however, as we know from the work of Pohlig, not the great significance for the distinction of elephant species as was previously thought.

The two *Stegodon trigonocephalus* skulls described by Martin {1*) Collections of the National Museum in Leiden. Vol. IV. Fossil mammals from Java and Japan, p.36 and Plates II, II and IV}, also belong to this species. If they had been well preserved, they would have shown that the triangular shape was not an original one and not characteristic of the species. The name which refers thereto can thus also be scratched. I propose at this point to distinguish this smaller Kendeng species only as a variant of the more robust mainland species, whom it clearly closely resembles, and introduce her then as *Stegodon ganesa* var. *javanicus*.

Even if one can not really admit that *Stegodon ganesa* and *Stegodon insignis* belong to one single species, the two are in any case closely related (the molars are hardly or not all distinguishable), *Stegodon insignis* can however, according to M. Schlosser {2*) M. Schlosser, The fossil mammals of China. Transactions Mathematics and Physics Class K, Bavarian Academy of Sciences. Vol. 12 (1903), p. 218} be regarded as a diagnostic fossil for the late Pliocene fauna of east Asia. From this alone it is probable that our Javanese *Stegodon ganesa* belongs to the same late Pliocene beds. Indisputable remains of *Stegodon ganesa* have not been found in any beds younger than Pliocene, because the tusk found in the Narbada beds can no longer be ascribed to this species with any certainty {3*) Lydekker, Indian Tertiary and post Tertiary vertebrata. Vol. I, p. 274}.

Also, among my Kendeng fossils there is only a single species of *Elephas*, which have allowed me precise insight into the many collected molars and in particular great familiarity with the shape of the skull as well. This species very closely resembles Elephas hysudricus and even more so the still living *Elephas indicus*, so that I therefore will refer to it as *Elephas hysurindicus* n.sp. Had this species already been recognized from the Siwalik beds as the probable antecessor of the living Asian elephants, this familial relationship becomes even clearer with the new Java species.

The great similarity of the molars had originally even caused me to assume that E. indicus was part of the Kendeng fauna, although the lamella count of the fossil Java molars does not exceed 19, thus considerably less than for *E. indicus*. The skull (which became known later) is however significantly more like the Siwalik skull than that of the living species, particularly in its profile, because of the larger lateral development of the parieto-frontal protuberances and the straight alveoles for the tusks. Yet the distance between the temple grooves has already increased; in this, the extinct Java species is extremely close to the living Indian species, whose immediate antecessor we undoubtedly have found in the first mentioned one.

The *Rhinoceros* class is, as I have reported before in this journal, represented by two single horned species. The most common of these is a close relative of the *R. sondaicus*, which is still living on Java as well as the mainland. This living species is certainly little different from *R. sivalensis*, and Lydekker considers the latter then as its forefather. It is then very peculiar that the Kendeng species bridges the small gap that still separates these two. Exactly the few differences which Lydekker describes, have become even less significant in the Kendeng species. In particular, the ratio of the length and width of the upper molars is adjusted as a transition between the two. I therefore name it *Rhinoceros sivasondaicus* n. sp. [*R. sondaicus*]. Yet, in any case this species was not identical to the modern one. Another difference with *R. sivalensis* is that it has a very small medial incisor of the lower jaw, with *R. karnuliensis* through the absence of a cingulum on the inner surface of the frontal cheek bone and small bumps in the cross valley of the upper molars.

The second species of *Rhinoceros* is associated with the exclusively on the continent living *R. indicus*, but is only different from it in minor points. The back cheek bone of the upper molars is relatively wide, the nose bones are narrower and have sharper edges. I distinguish this second species as *Rhinoceros kendengindicus* n. sp.

Among the most significant species of the Kendeng fauna belongs the deer species which I have introduced already in the past in this magazine as *Cervus liriocerus* n. sp. [*Axis lydekkeri*]. It concerns here a species that judging from its teeth and antlers belongs to the *Axis* group [Tables 3 and 7]. Just like the living *Axis* species, which it is very close to at any rate, it must have formed large herds, which we must be thankful for in that, near Trinil, it is represented by many hundreds of individuals. This graceful deer was somewhat smaller than the living *Axis* and distinguishes itself from all living and known fossil deer species by the antler beams which in their lower parts get very close together (in certain cases to within a few centimeters), as a result of which a perfect lyre shape is obtained. Also, the inside branches of the antlers are relatively short and not conjoined as is the case with the living *Axis*, as well as slightly turned backwards, the eye tine is on the other hand relatively long and strong.

*Cervus Lydekkeri* Martin {1*) Collections etc. Vol. IV, p.63 and Plate VII, Figs. 1 and 1a} is based only on the immature antlers specimen of the deer species that I just described and can therefore not really be maintained as a species. It is remarkable that one of the deer species, closely resembling living *Cervus hippelaphus* of Java [*Rusa timorensis*], *Cervus kendengensis* n. sp. [*Rusa* sp.], which is distinguished from the former by thicker and shorter antler beams, was much less common. The same is true for the second large fossil deer species, *Cervus palaeomendjangan* n. sp. [*Rusa* sp.], characterized by the small size of its outward and forward bending pronged branch, which thus had this in a more exaggerated proportion than it is in the modern species of Java.

The (*Axis*-) species that is rare in the Malay archipelago, is exactly the one that was most common in the past.

Several horns of *Cervulus* [*Muntiacus muntjak*]may well belong to a Pliocene species, although they are very similar to those of the still living species; these horns, as is well known, can vary greatly. Of greater significance for the determination of age of the Kendeng beds are some different ruminants, in the first place an antelope species, which already has been named *Tetracerus Kroesenii* n. sp. {2*) It is the same species which I originally suspected to be related to *Anoa* (*Anoa santeng*), when I only knew it by the horn core and a few broken skull pieces.} [*Duboisia santeng*; Table 3, main text]. This represents a highly peculiar transitional form. Because, while the shape of the skull, in particular the brow bone (on which the locations of the second horn pair of the living *Tetracerus quadricornus* [Four-horned antelope or Chousingha], similar to the two horned variety, is indicated by the deviation angles of the very roughened pre-cornutus ridges) as well as forward shifting of the infra- orbital holes correspond to that of an antelope species of the Indies, it is different again from this and from *Tetracerus Daviesi* from the Siwalik beds by well-developed prisms (“basalt pilings”) on the upper molars. Because of this, and the mostly straight lined parieto-frontal profile, it comes close to the living *Boselaphus tragocamelus* [the Nilgai of South Asia] and thus ties together these two living antelope species of the Indies, which, when following Ruetimeyer, were already known to be closely related. The strong correspondence with the latter species in the built up of the teeth, has led me in the past to include it in the same family. Considering, however, that on the one hand the second pair of horns is not always developed in *Tetracerus quadricornus*, and that on the other hand traces of accessory prisms are sometimes encountered on the upper molars, I still believe that it should be classified under *Tetracerus*. It is clearly distinguished from *Boselaphus namadicus* by the small size, little more than the living *Tetracerus* species, as well as by the different shape of the horn cores and the forward portion of the brow bone, which contains the rough growths.

Besides this antelope species an oxen species is even more important to the evolutional history relations. I name it *Leptobos groeneveldtii* [*Epiletobos groeneveldtii*], to honor the man who besides Kroese (his predecessor as vice president of the Indies Council) I owe most for the promotion of my paleontological research in Sumatra and Java.

*Leptobos groeneveldtii* n. sp. was as beautiful a species as it is significant to us. The skull is in size and general shape similar to the banteng skull, although the horn cores are rounder, rejuvenate themselves much less towards the tips and while, starting out as shoulders, they bend at first strongly upward they then turn downward and inward to the back until the tips are in front of the eye sockets. Between the horn bases the bony distention of the bovids is completely missing, the forward skull surface on the other hand is deepened between the horns and is continued upward up to a parieto-occipitalis protuberance. Also the measured height between the roof of the mouth and the nose bone is significantly smaller and the palate curvature is much flatter.

In a somewhat deviating variety of this same species, which I consider to be the female form, but for the time being still designate as *L. dependicornus*, the horns are weaker and are bent more backwards and the mentioned bony growth is much less developed.

The *Leptobos* of Java is distinguished from *Leptobos falconeri* of the Siwalik beds, as well as from *Leptobos elatus* (*L. etruscus*) of the southern European Pliocene, by its elongated shape of the horned skull and the orientation of the horn cones. Of *Leptobos frazeri* from the Narbada beds, only a horn-less variety is known, whose skull is also developed differently in other aspects from that of the Kendeng species. This one has most probably horns in both sexes, at least only horned skulls are being found. Also, the significant size of the *Leptobos* from Java is closer to that of the banteng then the remaining species are.

The teeth of this *Leptobos* are distinguished from the banteng species (Bibos) in the manner described by Ruetimeyer {1*) L. Ruetimeyer, Investigation of the natural history of the oxen. New commemorative journal of the General Swiss Society for collective Natural Sciences, Zuerich 1867, p. 91, 97, 100. Refer to the skull shape, ibid p. 71 and on. Also: L. Ruetimeyer, The Oxen of the Tertiary epoch, *Transactions of the Swiss Paleontological Society*, Vol. V (1878), pp. 153-168}. That is to say that the cross diameter of the upper molars (more pronounced with the m3) is smaller (pm4 is shorter as well). The lateral folds are about as strongly developed as with *L. elatus*. Also, besides this *Leptobos*, several *Bibos* species occur, of which the majority are more or less closely associated with the living banteng.

I collectively call these *Bibos palaeosondicus* n. sp. Others are transition forms halfway between our *Leptobos* species and this primitive banteng.

Very peculiar is, however, that besides this species there is a form which indistinguishably resembles the living gaur by its very high frontal swelling between the horns and the entire brow surface: it is *Bibos* *protocavifrons* n. sp. In all probability we are dealing here with the evolution of Bibos species from a *Leptobos* species that still somewhat resembled the antelope, an evolution which the genial Ruetimeyer had already suspected.

What must strike us as special, and something similar to this must capture our attention with other species of the Kendeng fauna, is the fact that the various stages of evolution occur simultaneously and side by side. This points more towards a sudden transitional evolution rather than a gradual and very slow one.

In the past, I assumed that the Kendeng buffalo was *Bubalus palaeindicus*. The fossil species of Java, however, resembles little in the elongated shape of the skull of the Narbada species, nor in the almost straight orientation and the nearly square cross section of its hollow horn cones. The shape of the skull is short as is the case with the living Java species, and the frontal surface of the horn cores is flat, with sharp upper and lower edges, while the orbital and temporal surfaces are rounded; and while these are also separated by a very blunt edge, the cross-sectional area of the horn cones can change from triangular to almost semi- circular. The Kendeng buffalo resembles in the shape of the hollow horn cones somewhat the Siwalik's *Bubalos platyceros* and differs in this from the living kerabau species. Apart from that, the skulls of both Java species are very similar; the living one is probably descended from the fossil species, as the name *Bubalus palaeokerabau* would indicate.

The two wild pig species of the Kendeng fauna belong to the group of *Celebensis verrucosis*, as is particularly shown in the shape of the cross section of the eye tooth, which according to Nathusius and Nehring {1*) A. Nehring, About Sus celebensis and related species. Nr. 2 of the Transactions and Announcements of the Royal Zoological and Anthropological Ethnographic Museum of Dresden. 1889}, forms the most important characteristic for the group classification of pigs. It is then again remarkable that based on the shape of the lower back molars, the one species, *Sus brachygnathus* n. sp., is very similar to the present-day Celebes pig [Sulawesi Warty Pig, *Sus celebensis*], while the other, *Sus macrognathus* n. sp., corresponds to *Sus verrucosus* [Java Warty Pig] now living on Java. Yet, the latter has peculiarities, such as the relative strength and shape of the lower canine as well as the bumpy rise in the lower jaw, which reminds one of *Sus celebensis*. The fossil species are therefore a bit closer than the mentioned living species and also are considered to be between the first transitions. Nehring, however, already suspected an earlier existence of a common ancestral form to the living species. Important is also the fact that the most common living wild pig of Java and Sumatra, closely related to the Papua pig, *Sus vittatus*, is not represented in the Kendeng fauna. It appears that this species entered at a later time from the Australian region.

The lower jaw from the Kendeng beds [1*) Collections of the National Geologic Museum of Leiden, Vol. IV. Fossil mammals from Java and Japan, p. 59 and Plate VII, Fig. 3 and 3a] ascribed by Martin to the Siwalik *Sus hysudricus*, belongs without doubt to *Sus brachygnathus*. As is well known, several species are grouped together into *Sus hysudricus* from the Siwalik beds; one of these may have been close to our *Sus brachygnathus*, but was most certainly not identical.

We are now getting to the extremely important hippopotamus species of the Kendeng. This is a *Hexaprotodon*, similar to the fossil hippopotami of the mainland of India. She is, however, far removed from the two Narbada species: *Hexaprotodon namadicus* and *H. palaeindicus*. Because of the length of the lower jaw symphysis and the relatively large size of the incisors as well as the shape of the skull, she corresponds on these points relatively closely to *Hexaprotodon sivalensis*, from which she is distinguished, apart from her somewhat smaller size, by the greater distance in proportion to the length between the palate and the nose bone, by the relatively wide upper jaw and especially also by the non rectilinear spots of wear on the lower incisors. She is clearly very close to the Siwalik species and while the overwhelming development of the lateral incisors makes the Narbada species known as an end branch of the *Hexaprotodon* family tree, *Hexaprotodon sivajavanicus* n. sp., is certainly even closer to the family of African hiipopotami, even more emphatically so than *Hexaprotodon sivalensis*, when comparing the relatively strong medial incisors.

Martin has attributed a piece of broken tooth from a drill hole at Ngembak in the residency Samarang, to a *Hippopotamus* "of tremendous size" {2*) Collections of the National Geological Museum of Leiden. Vol. III (Leiden 1883). Paleontological results of deep wells on Java, p. 12 and Plate I, Figs. 2, 2a, and 2b}. This turns out to be, however, the forward half of the right last lower jaw molar of a *Bubalus*. It must therefore not be assumed that a second species of Java *Hippopotamus* exists.

*Tapirus pandanicus* n. sp. [*Tapirus indicus*] from the Kendeng is very close to *Tapirus indicus*, but is distinguished from it by its smaller size and the broader development of the cross valleys on the upper molars.

Further, of great importance is a *Hyaena* belonging to the Crocuta group, which in the Eurasian Pliocene is represented by many species. This large species, the size of a lion, is very akin to the south European Pliocene *Hyaena brevirostris*, is similar in size and among the Siwalik hyena is closest to *Hyaena felina*. I name her *Hyaena bathygnatha* n. sp. [*Hyaena brevirostris*], because she is distinguished from this Siwalik species by the large height of the lower jaw, although this characteristic is not exceeding that of the European species. She is distinguished from the latter species by the somewhat greater length of the forward jaw portion (the shortness of which lends its name to *Hyaena brevirostris*) and thus by having not such a short face and consequently by the not so steep symphysis of the lower jaw, and furthermore by the more forward located foramen mentale.

There are three species of *Felis*. For two of them the body size was about the same as that of the tiger and lion, the third one strongly resembled *Felis minuta*. None of these can be identified with living species and neither can I find any close relationship with described fossil species. Strangely enough, the so common tiger that currently lives on Sumatra and Java as well as the mainland, is not even once represented by a related species. Their very peculiar current distribution indicates, by the way, that they have entered their habitat late.

a. *Felis oxygnatha* n. sp. is instantly distinguished from *Felis leo*, *Felis tigris*, *Felis pardus* and *Felis nebulosa*, with which he can be compared in size, by the narrow chin and related to this characteristic, the very obliquely outward oriented canines of the lower jaw; at the same time the diastema is short, the lower pm3 is proportionally large and on the pm4 the back cingulum has a knobby shape, and the lower m 1 is proportionally short. Whereas *Felis pardus* corresponds in short diastema and the general shortness of the forward jaw portion, as well as the snout, with the living *Felis jubata* of India, in the latter characteristics, however it does so even more with the American *Felis uncia*. Both of these, however, at the same time have broad snouts. He resembles *Felis nebulosa* by its narrow chin; but the diastema is very large in this living species and the forward jaw is very long. There are no close relationships to the *Felis* species from the Siwalik beds.

b. *Felis trinilensis* n. sp. [*Panthera tigris trinilensis*] This name replaces *Felis Groeneveldtii*, which I introduced in the past in this journal for the second large Felis species. The chin is of normal width, the gap between the teeth of the lower jaw is large, the pm 3 proportionally small, the m 1 large. There is a smaller pm2 in the upper jaw, pm3 and pm4 were shorter and more simply built than with the tiger. The skull portion in front of the eye openings, the snout, is proportionally short. Our species is different from lions among other things in the shape of the cavitas glenoidalis.

c. *Felis microgale* n. sp. [*Prionailurus bengalensis*], differs from the living *Felis minuta*, with which he corresponds in size, by the absence of a pm 2 in the lower jaw, by the presence of a small accessory hump on the lower pm3 and by a stronger cingulum on the pm4. With respect to these last two characteristics, he corresponds to *Felis catus*, but is distinguished then from him and corresponds in that regard with *Felis minuta*, by a deep fold on the outside of the lower canine. He is different from *Felis rubiginosa* in the humps on the front part of the pm4, and from *Felis chaus* by his much smaller size.

As sole representative of the otters, I discovered *Lutra palaeoleptonyx* n. sp. {see Willemsen, G.F., 1986. *Lutrogale palaeoleptonyx* (Dubois 1908), a fossil otter from Java In the Dubois Collection. *Proc. Kon. Akad. Wetenschappen* B, 89: 195-200.} This species is very similar to the present day *Lutra leptonyx*, living on Java and also on the mainland, but was significantly larger, even somewhat larger than the African *Lutra Aonyx*. Besides this, it is distinct from *Lutra leptonyx* through the straight outside circumference outline of the upper pm4, which only has a small protuberance at the front edge, as well as through the weakly expressed cingulum of these teeth, while, by contrast, this outer edge for the living species is concave for the front half and convex for the back half and a well pronounced cingulum is indicated for the latter half. In addition the forward peak of this tooth is proportionally higher.

A very peculiar species is the large *Manis palaejavanica* n. sp. This scaly ant eater, of which we have a number of bones of extremities, in particular also of the foot and a broken piece of skull bone, all belonging to one individual, reached one and a half time the size of the African *Manis gigantea*. The Kendeng species can be distinguished from this African species in many aspects. The shape of the second phalanx of the middle finger is also very different from the same sized bone of *Chalicotherium sivalense* from the Siwalik beds.

Lydekker has described the end phalanx of the middle finger of a *Manis* species from the karst holes (?), which equaled in size that of the African *Manis gigantea* and which he "by the absence of any proof to the contrary," could view as specifically identical to this African species {1*) Indian Tertiary and Post tertiary Vertebrata. Vol. IV, p. 50 and Plate VIII, Figs. 8 and 8a.}. In this, I find it difficult to follow this prominent worker of the Siwalik fauna, because when comparing them more carefully, there are indeed important differences. The principal ones are the continuing split and the blunt base of the bone from the karst hole, as well as the absence of spanning of the large blood vessel imprint on the palm of the hand (volar). In this respect the species from India is similar to our *Manis palaejavanica*, but is also different from it, besides in size, in the sharp palm side boundary of the joint surface and several other features. The karst hole *Manis*, which has been used for far reaching conclusions about the family relations between the animal world from the Indies and Africa, must therefore be considered a new species, for which I propose the name *Manis lydekkeri* n. sp.

*Manis palaejavanica* is, judging by the available bones, certainly closer related to the living *Manis javanica* than the African species as well as the other species from the Indies. The extremities were, apart from the enormous difference in size, proportionally stocker as well as relatively stronger than in the living Java species.

Tafel XXXIX.

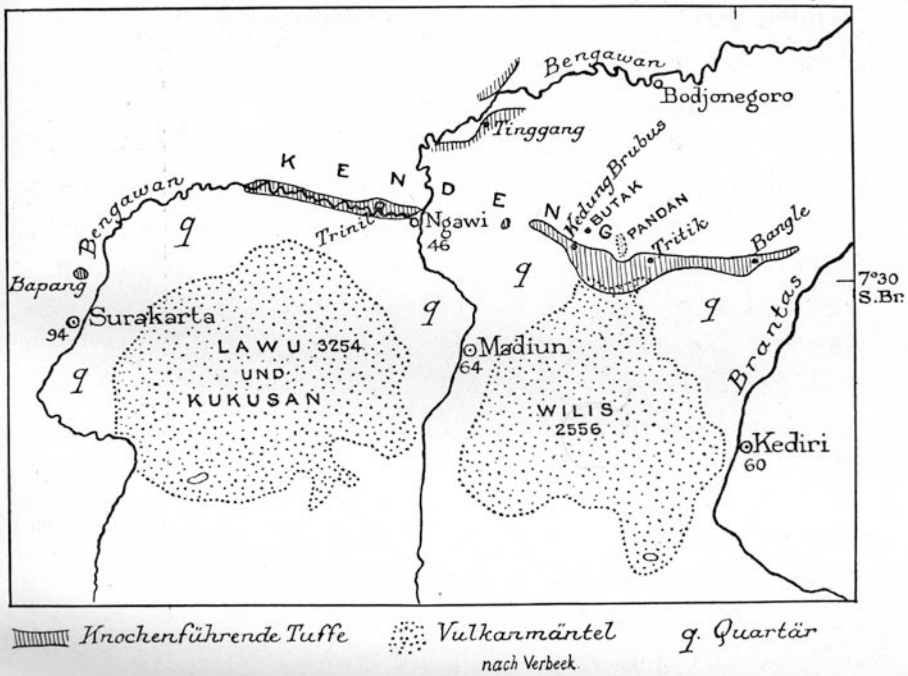


*a.* Kartenskizze der Umgebung von Trinil. Masstab 1 : 1 500 000.

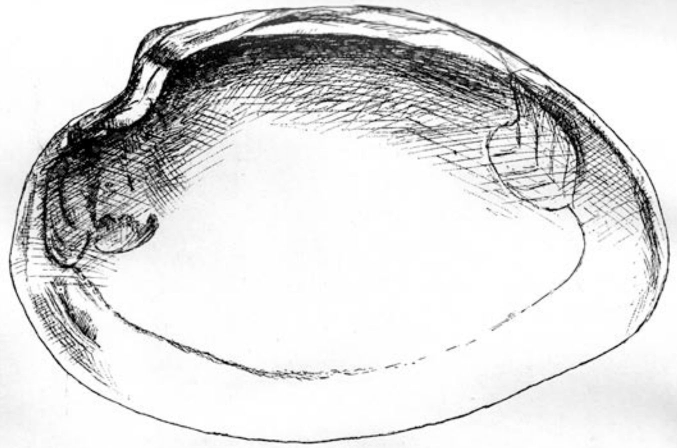

*b.* Unio trinilensis. $\frac{1}{1}$
Trinil. „Hauptknochenschicht".

**S II-F8**

[Dubois (1924: 265-278), written in English. “On the Principal Characters of the Cranium and the Brain, the Mandible and the Teeth of *Pithecanthropus erectus*.”]

[265>>266] Before the morphological characters of the fossil remains of this Primate are discussed, a few remarks may be made about the state of perfect mineralization in which they are. This state is entirely different from that of the oldest of the human remains known. Their specific gravity, like that of the bones of the other mammals which were dug up with them at Trinil (which bones have the same outward appearance), has risen to about 2.7 and increase of about 35% compared with dry fresh bony substance. …. According to the analysis of the late Prof. J.M. van Bemmelen both phosphate and carbonate of calcium have taken the place of the ossein, and they contain fluorium in the quantity which, according to Ad. Carnot’s investigations, is characteristic of fossil bones of the Pliocene. The particular pseudomorphs known as mineralization, petrification or fossilization, have very strongly affected the bones of Trinil. Their great antiquity appears also from this that calcite and pyrites have crystallized in many cavities and canals in the bones …. The morphological investigation of the cranium is restricted or hampered by three circumstances: first that only the upper part of the calvaria [**Skullcap**] as calotte or calvarium has been preserved; secondly that the outer surface has been greatly corroded by sulfuric acid, formed from pyrites in the volcanic tuff; thirdly that the cranium has been deformed in a natural way (through trigoncephalism, though in a small degree). … Particularly, however, about the original state of the external surface of the calvaria the fossil alone can give an accurate idea. It must have been smooth on the whole, about as at the skull of a small gibbon.. .

[268] … of the occipital bone …. the loss of substance greatly increases towards the edge of the fragment, so that this edge only still consists in the knife-like lamina interna.

[274>>275] … at Trinil … a nearly a year before the discovery there of the first fossil remain of *Pithecanthropus* (the hindmost right upper molaris), a mandibular fragment, a small piece on the right of the symphysis, was found in the same Kendeng-layers, but 40 km. distance on the E.S.E. of Trinil, namely at Kendung Brubus, among other fossil remains of the Kendeng-fauna. Its specific gravity is the same as that of the teeth and the other remains of *Pithecanthropus*. A brief description of the mandibular fragement appeared at Batavia in 1891, in the ‘Verslag van het Mijnwezen,’ and I mentioned it, in a few words, in the ‘Natuurkundig Tijdschrift voor Nederlansch-Indie” of the same year. I then considered it a remain of a not exactly determinable human species, “of another and probably lower type” than those existing and the extinct European diluvial species.

**S II-F9(i)**

[Dubois (1926a. Palaeontology. – "On the Principal Characters of the Femur of *Pithecanthropus erectus*." By Prof. Eug. Dubois ….meeting of March 27, 1926, pages 730-743).]

The left femur was dug up at Trinil in August 1892, during low-water level of the Bengawan, when the most fossiliferous layer of the andesite tuff was again accessible. Ten months earlier the calvarium had been found in that deposit of an ancient river, at 15m distance *in the same stratigraphic plane*. This circumstance, added to the anatomical relations between the two objects, renders it almost infinitely probably that they represent parts of the same individual.

The state of petrification is also the same. The femur has the same deep brown colour as the calvarium. It weighs 1018gr., which is more than twice the weight of a human femur of the same size, to which the fossil, on the whole, bears a striking resemblance. The volume is 485cm$^3$. Without the exostosis and without the comparatively small defects this would have been 467cm$^3$. …. I estimate that more than 100 cm$^3$, about 2/5 of all the cavities, are filled up with calcite and some pyrite. The existence of this filing can be observed through the hole in the popliteal surface and by rontgenograms …..

That hole in the popliteal surface has been caused during the excavation, a piece of compacta of a length of almost 4 cm and of a breath of from 1 to 1½ cm broke off and got lost. In the same way a piece in the foss intercondyloidea, about 2cm long, has got lost, and also a small fragment at the anterior extremity of the condylus medialis. Immediately above its lower edge the condylus lateralis further shows a round impression of the point of a crocodile tooth. Some marks of crocodile teeth are to be seen at the upper portions of the femur. Thus on the front side: in the collum, beside the caput, a shallow impression accompanied with small fractures, and between the linea intertrochanterica anterior and the lower part of the troncanter major, a deeper, round impression, directed obliquely from the inner side and above toward the outer side and below. At the back side: in the middle of the collum a semi-circular impression obliquely across fragments going from above and the inner side towards below outward; another inthe upper corner of the trochanter major, which corner is further broken off; on the outer side of the crista intertrochanterica a large shallow impression 1 cm lower; a double shallow impression on the trochanter major, in the middle of the surface for the tendon of the musculus glutaeus medius. The caput femoris, preserved for the most part, presents however extensive defects on the margin of the globular articular surface, which were probably caused by crocodiles, so that of that margin only a small part has remained preserved on the upper side, and a still smaller part on the lower side.

The large exostosis below the trochanter minor takes the place of the intermuscular connective tissue between the vastus medialis and the adductores, accompanying the arteria and the vena profunda femoris and their rami perforantes. The course of these blood-vessels can clearly be recognized by the grooves and perforations of the exostosis. ….

**S II-F9(ii)**

[Dubois (1926b. Palaeontology. – "Figures of the Femur of *Pithecanthropus Erectus*" (Plates I to IV). By Prof. Eug. Dubois … meeting of June 26, 1926, pages 1275-1277; also, Dubois 1932, 1934, 1935]

The photographic figures 1 tot [sic] 6 and the rontgenograms, figures 8 and 9….

Of points not mentioned in "The Principal characters of the Femur," the figure shows, in the upper two-third part of the shaft, a certain roughness caused by adherent small pyrtoid concretions, moreover, as consequences of the excavating work, some superficial loss of flakes and a somewhat larger one at the inner side of the lower end (about 9cm from below). Of the anterior extremity of the condylus medialis a small fragment of bone got lost, exposing the spongiosa: this part appears smooth in the figure, as a consequence of the preparation for making the plaster cast. Conspicuous is on the epicondylus lateralis a flat eminence for the insertion of the lower part of the intermuscular septum behind the vastus lateralis crureus. ….

DUBO1407

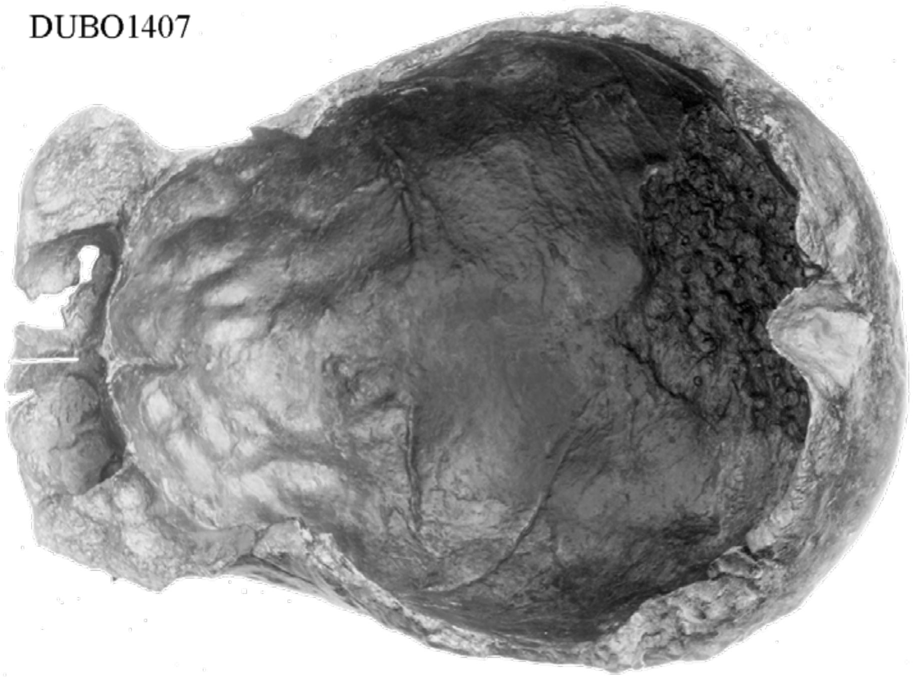

[The basal view, above, is similar to the image that Dubois used in his Plate IV. The superior view, below, is similar to the one used in Plate I. The DUBO####s refer to the Naturalis image used.]

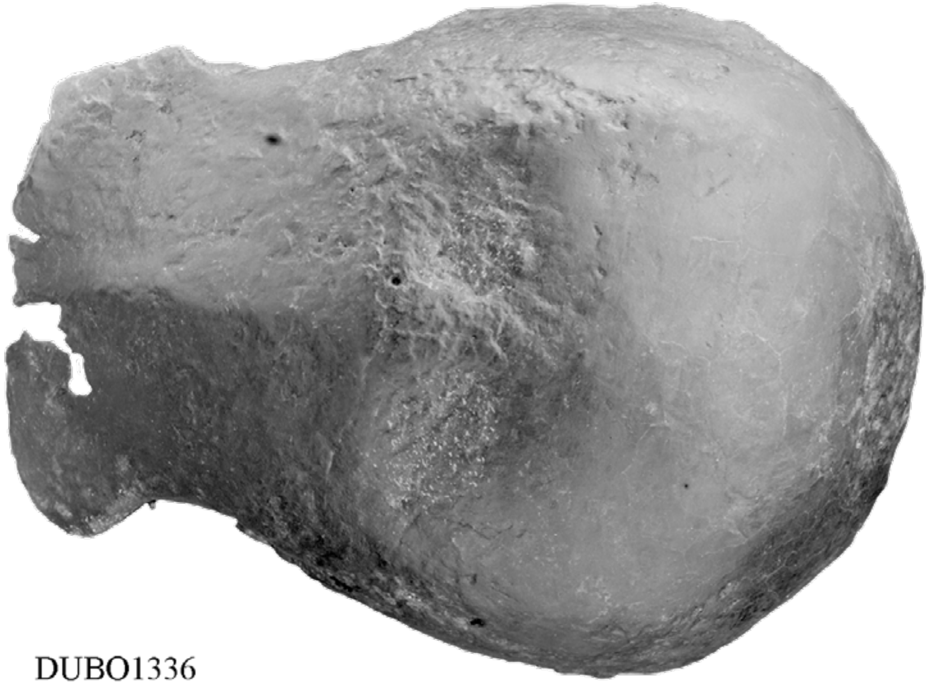

DUBO1336

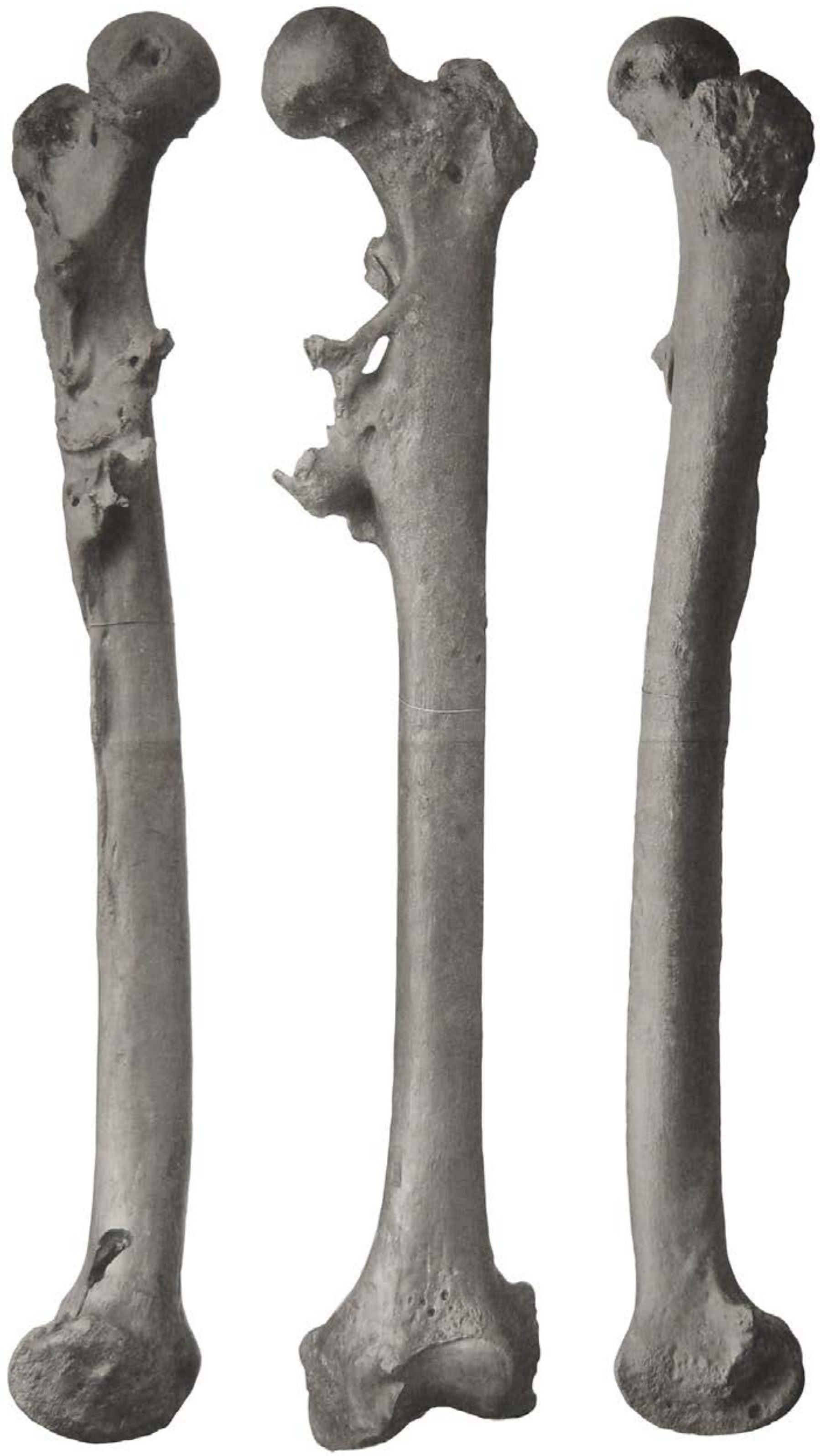

[Fig. 1. – (Left) Femur of *Pithecanthropus erectus*, front view. … of the most prominent back parts of the bone. [colorized version of a black and white photograph].

Dana, J.D. 1879. Manual of Geology, Third Edition. Trübner & Co., New York/Chicago. 716 p.

de Vos, J. 1989. The environment of *Homo erectus* from Trinil H.K. In *Hominidae, Proceedings of the 2nd International Congress of Human Paleontology, Turin, (September 28-October 3, 1987),* Editorial Jaca Book. pp. 225-229.

de Vos, J., and Aziz, F., 1989. The excavations by Dubois (1891-1900), Selenka (1906-1908), and the geological survey by the Indonesian-Japanese team (1976-1977) at Trinil (Java, Indonesia). *Anthropological Society of Nippon* 97, 407-421. https://www.jstage.jst.go.jp/article/ase1911/97/3/97_3_407/_pdf

de Vos, J., and Sondaar, P.Y. 1982. The importance of the "Dubois Collection" reconsidered. *Modern Quaternary Research in Southeast Asia* 7, 35-63.

Dubois, E. 1891. *Palaeontologische onderzoekingen op Java: 3rd quarter 1890.* Verslag van het Mijnwezen. Extra bijvoegel der Javasche courant 9, 12-15.

Dubois, E. 1892a. Voorloopig bericht omtrent het onderzoek naar de Pleistocene en Tertiaire vertebraten-fauna van Sumatra en Java, gedurende het jaar 1890 [Preliminary report on the investigation into the Tertiary and Pleistocene vertebrate fauna of Sumatra and Java, during the year 1890]. *Natuurkundig Tijdschrift voor Nederlandsch-Indië* 51, 93–100 [Dubois' paper is indicated to have been completed on Jan. 29, 1891, at Tulung-Agung, Java].

Dubois, E. 1892b. *Palaeontologische onderzoekingen op Java: 3rd quarter 1891.* Verslag van het Mijnwezen. Extra bijvoegsel der Javasche Courant 1891, 12-14. [also, Dubois, E. 1894e, below, and Naturalis scan M…050-071].

Dubois, E. 1892c. *Palaeontologische onderzoekingen op Java: 4th quarter 1891.* Verslag van het Mijnwezen. Extra bijvoegsel der Javasche Courant 1891, 12-15. [also, Dubois, E. 1894e, below, and Naturalis scan M…050-075].

Dubois, E. 1892d. *Palaeontologische onderzoekingen op Java: 2nd quarter 1892.* Verslag van het Mijnwezen. Extra bijvoegel der Javasche courant 87, 14-18. [also, Dubois, E. 1894e, below, and Naturalis scan M…050-077 and -078].

Dubois, E. 1893a. *Palaeontologische onderzoekingen op Java: 3rd quarter 1892.* Verslag van het Mijnwezen. Extra bijvoegel der Javasche courant 10, 10-14. [also, Dubois, E. 1894e, below, and Naturalis scan M…050-077 and -078].

Dubois, E. 1893b. *Palaeontologische onderzoekingen op Java: 4th quarter 1892.* Verslag van het Mijnwezen. Extra bijvoegel der Javasche courant 36, 11-12. [also, Dubois, E. 1894e, below].

Dubois, E. 1894a. *Pithecanthropus erectus. Eine Menschenaehliche Uebergangsform aus Java.* Batavia: Landsdrukkerij [Foreword dated January 1894] 39 p. [with 2 plates]. https://www.biodiversitylibrary.org/item/132258#page/7/mode/1up

Dubois, E. 1894b. *Palaeontologische onderzoekingen op Java: 3rd quarter of 1893.* Verslag van het Mijnwezen. Extra bijvoegel der Javasche courant 2, 15-17.

Dubois, E. 1894c. *Palaeontologische onderzoekingen op Java: 4th quarter 1893.* Verslag van het Mijnwezen. Extra bijvoegel der Javasche courant 81, 12-15.

Dubois, E. 1894e. *Rapporten van de Palaeontologische onderzoekingen op Java 1890-1894 Eug. Dubois.* [Reports of the Paleontologic investigations on Java 1890-94 Eug. Dubois]. First Quarter 1894 [p.

104, Naturalis scan M…050-090, Right]; Second Quarter 1894; Third Quarter 1894 [p.108 –112, Naturalis scan M…0050 -091].

Dubois, E., 1895a. *Pithecanthropus erectus. Eine Menschenaehliche Uebergangsform aus Java.* J*aarboek van het Mijnwezen Nederlandsch-Oost-Indie* 24. pp. 1-77. Stemler (Amsterdam).

Dubois, E. 1895b. Resumé d'une Communications de M. Le Dr. Eug. Dubois sur le Pithecanthropus erectus du Pliocène de Java [Summary of an oral presentation by Dr. Eugene Dubois on the *Pithecanthropus erectus* from the Pliocene of Java]. In *Bulletin de la Société Bèlge de Géologie, de Paleontologie et d' Hydrologie, Bruxelles.* [Bulletin of the Belgian Society for Geology, Paleontology and Hydrology, Brussels] Proces-vertaux Tome IX [1895]. Bruxelles (Polleunis et Ceuterick, Imprimeurs, Décembre 1896). [p. 157 has "Fig. 1, Coupe des couches ossiferes a Trinil;" p. 158 has Fig. 2, an uncaptioned map which is the source of Figure 2b, main text].

Dubois, E., 1895c. *Pithecanthropus erectus*, betrachtet als eine wirkliche Ueberigansform und als Stammform des Menchen. In *Verhandlungen der Berliner Gesellschaft für Anthropologie, Ethnologie und Urgeschichte* [Proceedings of the Berlin Society for Anthropology, Ethnology and Early History]. Virchow, R. (ed.). Sitzung vom 14 December 1895 [Special Session of December 14, 1895], pp. 723-738 [p. 725, cross section]. Berlin A. Asher & Co.

Dubois, E., 1896a. Näheres über den *Pithecanthropus erectu*s als menschenähnliche Übergangsform. *Internationale Monatschrift fur Anatomie und Physiologie* 13, 1-26.

Dubois, E., 1896b. *Pithecanthropus erectus*, einen menschenaehnliche Uebergangsform. In *Compte-rendu des seances du Troisieme Congres International de Zoologie, Leyde, 16-21 Septembre, 1895.* Hoek, P.P.C. (ed.). Leiden: E.J. Brill. pp. 251–273 [the first paper on *Pithecanthropus erectus* that Dubois delivered in Europe, the presentation was given in Quatrieme Séance, Saturday morning, September 21 1895, and included notes from discussion, a geological cross section (Fig. 1), and a comparison of cranial profiles (Fig. 2)].

Dubois, E., 1896c. On *Pithecanthropus erectus*: a transitional form between man the apes. *Scientific Transactions of the Royal Dublin Society* S.2. 6 [read November 20, 1895; published February 1896], 1-18. https://www.biodiversitylibrary.org/item/51748#page/7/mode/1up

Dubois, E., 1896d. On *Pithecanthropus erectus*: a transitional form between man and the apes. Abstract. *Journal of the Anthropological Institute of Great Britain and Ireland.* 25, 240-248 [An "abstract" based upon the Royal Dublin Society paper read on November 20, 1895, with material already published in Dubois's first European paper (Dubois, 1896b, above), and published more fully in the February 1896 Transactions of the Royal Dublin Society vi (I), 1-18].

Dubois, E. 1896e. *Pithecanthropus erectus*, eine Stammform des Menchen. *Anatomischer Anzeiger* 12, 1-22.

Dubois, E. 1896f. Le *Pithecanthropus erectus* et l'origine de l'homme. *Bulletins de la Société d'Anthropologie de Paris, Tome Septième IVe Série* 7, 460 – 467.

Dubois, E. 1899. Abstract of remarks on the brain-cast of *Pithecanthropus erectus* [Read at a General Meeting of the Fourth International Congress of Zoology, Cambridge, 26th August 1898. Communicated by W.L.H. Duckworth]. *Journal of Anatomical Physiology* 33, 273–276.

Dubois, E. 1907. Eenige van Nederlandschen kant verkregen uitkomsten met betrekking tot de kennis der Këndengfauna (fauna van Trinil) [Some results from the Netherlands regarding the knowledge of the Kendeng (Trinil) fauna]. *Tijdschrift van het Koninklijk Nederlandsch Aardrijkskundig Genootschap* 24, 449-458.

Dubois, E. 1908. Das geologische Alter der Kendeng oder Trinil Fauna [The geologic age of the

Kendeng- or Trinil-fauna]. *Tijdschrift van het Koninklijk Nederlandsch Aardrijkskundig Genootschap* 25, 1235–1270.

Dubois, E. 1924. On the principal characters of the cranium and the brain, the mandible and the teeth of *Pithecanthropus erectus*. Proceedings of the Koninklijke Nederlandse Akademie van Wetenschappen Amsterdam. 27: 265-278

Dubois, E. 1926a. On the principal characters of the femur of *Pithecanthropus erectus*. *Proceedings of the Koninklijke Nederlandse Akademie van Wetenschappen Amsterdam* 29, 730-743.

Dubois, E. 1926b. Figures of the femur of *Pithecanthropus erectus*. *Proceedings of the Koninklijke Nederlandse Akademie van Wetenschappen Amsterdam* 29, 1275-1277.

Dubois, E. 1932. The distinct organization of *Pithecanthropus* of which the femur bears evidence, now confirmed from other individuals of the described species. *Proceedings of the Koninklijke Nederlandse Akademie van Wetenschappen Amsterdam* 35, 716-22.

Dubois, E. 1934. New evidence of the distinct organization of *Pithecanthropus*. *Proceedings of the Koninklijke Nederlandse Akademie van Wetenschappen Amsterdam* 37, 139-145.

Dubois, E. 1935. The sixth (fifth new) femur of *Pithecanthropus erectus*. *Proceedings of the Koninklijke Nederlandse Akademie van Wetenschappen Amsterdam* 38, 850-53.

Duyfjes, J. 1933. *Javakaarteering. Maandverslag over Mei/Juni 1933 een Tournee in Blad 93B Ngawi* [Java Mapping Program Monthly report for May/June 1933 a survey in Quadrangle 93B Ngawi (which include Trinil)]. [Internal unpublished type-written report of the Geologische Dienst, Bandung, Java; Reference no. E33-79 with 1:25,000 geological map]. Library of the Geololgical Research and Development Center (Bandung). 18 p.

Duyfjes, J. 1936. Zur Geologie und Stratigraphie des Kendenggebietes zwischen Trinil und Soerabaja (Java) [On the Geology and Stratigraphy of the Kendeng Hills between Trinil and Surabaya (Java)]. *De Ingenieur in Nederlandsch-Indië, IV. Mijnbouw & Geologie, De Mijningenieur*. 3, 136-149 [translation in Huffman 2020].

Geike, A. 1885. Textbook of Geology, Second Edition. Macmillan, London. 992 p. ISBN: B0024TKU1E

Hooijer, D.A. 1946a. Prehistoric and fossil rhinoceroses from the Malay Archipelago and India. *Zoologische Mededelingen Museum Leiden* 26, 1-138.

Hooijer, D.A. 1958a. Fossil Bovidae from the Malay Archipelago and the Punjab. *Zoologische Mededelingen Museum Leiden* 38, 1-112.

Hooijer, D.A. 1962. Quaternary langurs and macaques from the Malay Archipelago. *Zoologische Mededelingen Museum Leiden* 55, 1-64.

Homo erectus. 1893. Naar Aanleiding van Paleontologische Onderzoeken op Java [In reference to paleontological research on Java]. Verslag van het Mijnwezen in Ned. Indie, 3e kwartaal 1892. *Derde Blad Bataviaansch Nieuwsblad*, Maandag 6 Februari, 1893, No. 57. [Report of Mining Bureau in Netherlands Indies, Third Quarter 1892. Third page of Batavia Newspaper, Monday 6 February 1893, No. 57].

I.J.R.C.P. (Indonesia – Japan Research Cooperation Programme CTA-41). 1979. Progress report on the Indonesia – Japan joint research project on geology of human fossil bearing formations in Java. *Bulletin of the Geological Research and Development Centre* 1, 47-60. [also, Soeradi et al. 1985].

Indriati, E. 2004. Indonesian hominid fossil discovery of 1889-2003: catalogue and problems. In *Proceedings of the Fifth and Sixth Symposia on Collection Building and Natural History Studies in Asia and Pacific Rim*, Akiyami, S., Miyawaki, R., Kubodora, T., Higuchi, M. (eds.). National Science Museums Monographs 24, pp. 163-177.

Jacob, T., 1975. Indonesia. In Oakley, K.P., Campbell, B.G., and Molleson, T.I. (eds.). *Catalogue of Fossil Hominids. Part III: Americas, Asia, Australasia.* The British Museum London, London, 104-112 (217 p.).

Leakey, R. E., and Slikkerveer L. J. 1993. *Man-ape ape-man. The quest for human's place in nature and Dubois' 'missing link.'* Leiden (Netherlands Foundation for Kenya Wildlife Service). 184 p. ISBN-10 : 0710304803 ISBN-13 : 978-0710304803

Lyell, C. 1830–1833. Principles of Geology, Being an Attempt to Explain the Former Changes of the Earth's Surface, by Reference to Causes Now in Operation. John Murray, London. 3 vols.

Oppenoorth, W.F.F. 1911. Arbeitsbericht über die Ausgrabungen. I. Teil. Die Arbeiten des Jahres 1907 bis August [Report on the Excavation Work. Part I. Work up to August 1907]. In *Die Pithecanthropus-Schichten auf Java: Geologische und Palaeontologische Ergebnisse der Trinil-Expedition (1907 und 1908)*, Selenka, M.L., and Blanckenhorn, M. (eds.). Wilhelm Engelmann, Leipzig. pp. XXVI-XXXVIII (288 p.). [also, Berkhout and Huffman 2021]

Saléh, R. 1867. Over fossiele beenderen van den Pandan. *Natuurkundig Tijdschrift voor Nederlandsch Indië* 29, 422-423, 426-429, 433-437, 448-451, 455-459.

Schwartz, J., Tattersall, I. 2005. The Human Fossil Record. Vol. 4, Craniodental Morphology of Early Hominids (Genera *Australopithecus, Parathropus, Orrorin*): An Overview. John Wiley & Sons, Hoboken.

Selenka, M.L., and Blanckenhorn, M., 1911. Die *Pithecanthropus*-Schichten auf Java: Geologische und Palaeontologische Ergebnisse der Trinil-Expedition (1907 und 1908) [The *Pithecanthropus* beds in Java: Geological and Paleontological Results of the 1907 and 1908 Trinil Expedition], Leipzig: Wilhelm Engelmann. 288 p. http://www.biodiversitylibrary.org [also, Berkhout and Huffman 2021]

Shipman, P. 2001. *The man who found the missing link. Eugène Dubois and his lifelong quest to prove Darwin right.* Simon and Schuster, New York. 514p.

Smith, T.M., Olejniczak, A.J., Kupcizik, K., Lazzari, V., de Vos, J., Kullmer, O., Schrenk, F., Hublin, J.-J., Jacob, T., and Tafforeau, P. 2009. Taxonomic assessment of the Trinil molars using non-destructive 3D structural and development analysis. *PaleoAnthropology* 2009, 117-129.

Storm, P. 2012. A carnivorous niche for Java Man? A preliminary consideration of the abundance of fossils in Middle Pleistocene Java. *Comptes Rendus Palevol* 11, 191–202.

Stremme, H. 1911. Die Saugetiere mit Ausnahme der Proboscidier. In *Die Pithecanthropus-Schichten auf Java: Geologische und Palaeontologische Ergebnisse der Trinil-Expedition (1907 und 1908),* Selenka, M.L., and Blanckenhorn, M. (eds.). Engelmann, Leipzig. pp. 82-160 [also, Berkhout and Huffman 2021].

Theunissen, B., 1989. *Eugene Dubois and the Ape-Man from Java. The history of the "Missing Link" and its discoverer.* Dordrecht (Kluwer). 216 p. ISBN: 978-94-009-2209-9

van Veen, F.R., 2004. Gustaaf Molengraaff 1860-1942 Een avontuurlijk geleerde. Delft University Press Satellite, Delft

Verbeek, R.D.M. Fennema, R., 1896. *Geologische beschrijving van Java en Madoera.* J.G. Stemler Cz., Amsterdam.

Willemsen, G.F.,1986. Lutrogale palaeoleptonyx (Dubois 1908), a fossil otter from Java In the Dubois Collection. Proceedings of the Koninklijke Nederlandse Akademie van Wetenschappen Amsterdam. 89, 195-200.
